# Tensor-decomposition-based unsupervised feature extraction applied to prostate cancer multiomics data

**DOI:** 10.1101/2020.07.17.208538

**Authors:** Y-h. Taguchi, Turki Turki

**Affiliations:** Department of Physics, Chuo University, Tokyo 112-8551, Japan; Department of Computer Science, King Abdulaziz University, Jeddah, Saudi Arabia

**Keywords:** prostate cancer, gene expression, genomic regions, protien-coding genes, tensor decomposition, unsupervised learning

## Abstract

The large *p* small *n* problem is a challenge without a de facto standard method available to it. In this study, we propose a tensor-decomposition (TD)-based unsupervised feature extraction (FE) formalism applied to multiomics datasets, in which the number of features is more than 100,000 whereas the number of samples is as small as about 100, hence constituting a typical large *p* small *n* problem. The proposed TD-based unsupervised FE outperformed other conventional supervised feature selection methods, random forest, categorical regression (also known as analysis of variance, or ANOVA), penalized linear discriminant analysis, and two unsupervised methods, multiple non-negative matrix factorization and principal component analysis (PCA) based unsupervised FE when applied to synthetic datasets and four methods other than PCA based unsupervised FE when applied to multiomics datasets. The genes selected by TD-based unsupervised FE were enriched in genes known to be related to tissues and transcription factors measured. TD-based unsupervised FE was demonstrated to be not only the superior feature selection method but also the method that can select biologically reliable genes. To our knowledge, this is the first study in which TD-based unsupervised FE has been successfully applied to the integration of this variety of multiomics measurements.

## 1. Introduction

The term “big data” often indicates a high number of instances as well as features [1,2]. Typical big data comprise a few million images (photos) each composed of several million pixels. The number of features is as high as the number of instances, or often higher. Most recently developed machine-learning methods including deep learning (DL) aim to address this manner of problem [1,3–8]. Nonetheless, although less popular than these typical big data problems, there is another branch of big data problem known as the “large *p* small *n*” problem, or “high-dimensional data analysis”. For these cases, the number of instances is typically smaller than the number of features [9–11]. In this case, there can be many unique problems that do not exist for the typical big data problems mentioned above. For example, in discrimination tasks, because there are more features than instances, if we do not reduce the number of features, the accuracy will always be 100% for no other reason than overfitting. Alternatively, the so-called “curse of dimensionality” problem describes a shortage of instances covering sufficiently large spaces. Because of these difficulties, compared with the popular big data problems, the “large *p* small *n*” problem has relatively rarely been investigated comprehensively, although there are many studies to tackle “large *p* small *n*” problem [12–15].

This might not seem significant, as the “large *p* small *n*” problem is relatively rare and not as important. Nevertheless, dependent on the domains, the “large *p* small *n*” problem will inevitably become typical. For example, in genomic science, the typical number of features is as many as or more than that of genes (i.e., about 10^4^), whereas the number of samples is the number of patients, typically at most a few hundred. The length of the DNA sequence of the human genome is approximately 3 × 10^9^ [16], whereas the human population is about 10^10^. Nevertheless, only a very small fraction of these populations can be considered in an individual study. Thus, in typical genomic studies, the ratio of the number of features to the number of samples can be in the region of 10^2^ or more.

Nonetheless, Taguchi has proposed a very different method to the typical machine-learning methods that are applicable to large *p* small *n* problems: tensor-decomposition (TD)-based unsupervised feature extraction (FE) [17]. *m*-mode tensor is associated with more than two suffix whereas matrix is associated with two suffix, row and column. In this method, a smaller number of representative features, referred to as singular value vectors, are generated with linear combinations of the original large number of features, without considering labeling. These singular value vectors are attributed to both samples and original features. We have investigated the singular value vectors attributed to samples. Finally, the selected singular value vectors attributed to the original features are used to evaluate the importance of the original features; because the singular value vectors are the linear combinations of original features, their coefficients of linear combinations can be used to evaluate the importance of individual original features. The original features with larger absolute values of coefficients are selected. As a result, we can avoid many difficulties in the “large *p* small *n*” problem; for example, we can perform standard discriminant analysis when there is a higher number of features than instances.

In this paper, we propose a TD-based unsupervised FE formalism, applied to synthetic data that imitate typical genomic datasets. The proposed method is first compared with four conventional methods applicable to the “large *p* small *n*” problem: random forest (RF), penalized linear discriminant analysis (LDA) [18], categorical regression, and multiple non-negative matrix factorization (MNMF); the latter has been used especially often for analysis of multiomics datasets [19]. These five methods are then also applied to a real dataset consisting of multiomics measurements of prostate cancers, a typical “large *p* small *n*” dataset. The results for the synthetic and real datasets demonstrate the superiority of our proposed method when compared against the four conventional methods in feature selections.

In this study, we need to deal with categorical data set. Since the labeling is neither binary nor numeric, no standard regression-based approaches are applicable. Thus, the methods applicable are restricted to either supervised methods that can deal with categorical labeling, e.g., tissues or unsupervised methods that do not require labeling and do not consider sample annotations at all. The above six methods were selected as representative methods that can satisfy the above requirements.

To our knowledge, this is the first study in which TD-based unsupervised FE has been successfully applied to the integration of this variety of multiomics measurements. The performance of TD-based unsupervised FE is highly data-type-dependent, since we cannot intentionally generate singular value vectors of interest. When singular value vectors generated by TD is not of interest, we cannot do anything since TD does not have tunable parameters. In this sense, it is challenging to estimate how many types of multiomics datasets (here, three transcription factor bindings as well as four histone modifications and one chromosomal state measurement, making eight types in total) can be successfully integrated with this method.

The types of multiomics data [20] that are possibly dealt with the proposed technology are as follows: gene (or mRNA) expression profiles, DNA methylation profiles (e.g., promoter methylation), various histone modification, chromatin structure, protein binding to genome (e.g., transcription factor biding to DNA), and RNA modification and so on.

A recent review [20] listed handful multi-omics data sets analysis tools (Table 1). As can be seen, the number of samples are from 10^2^ to 10^3^ and numbers of omics data used are at most four. TD based unsupervised FE (see below) can deal with mode extreme case; the number of samples per omics and tissue is from three to thirty while number of omics is as many as eight. We do not know any methods other than ours that can deal with this many kinds of omics data set as well as this small number of samples.

**Table 1.**
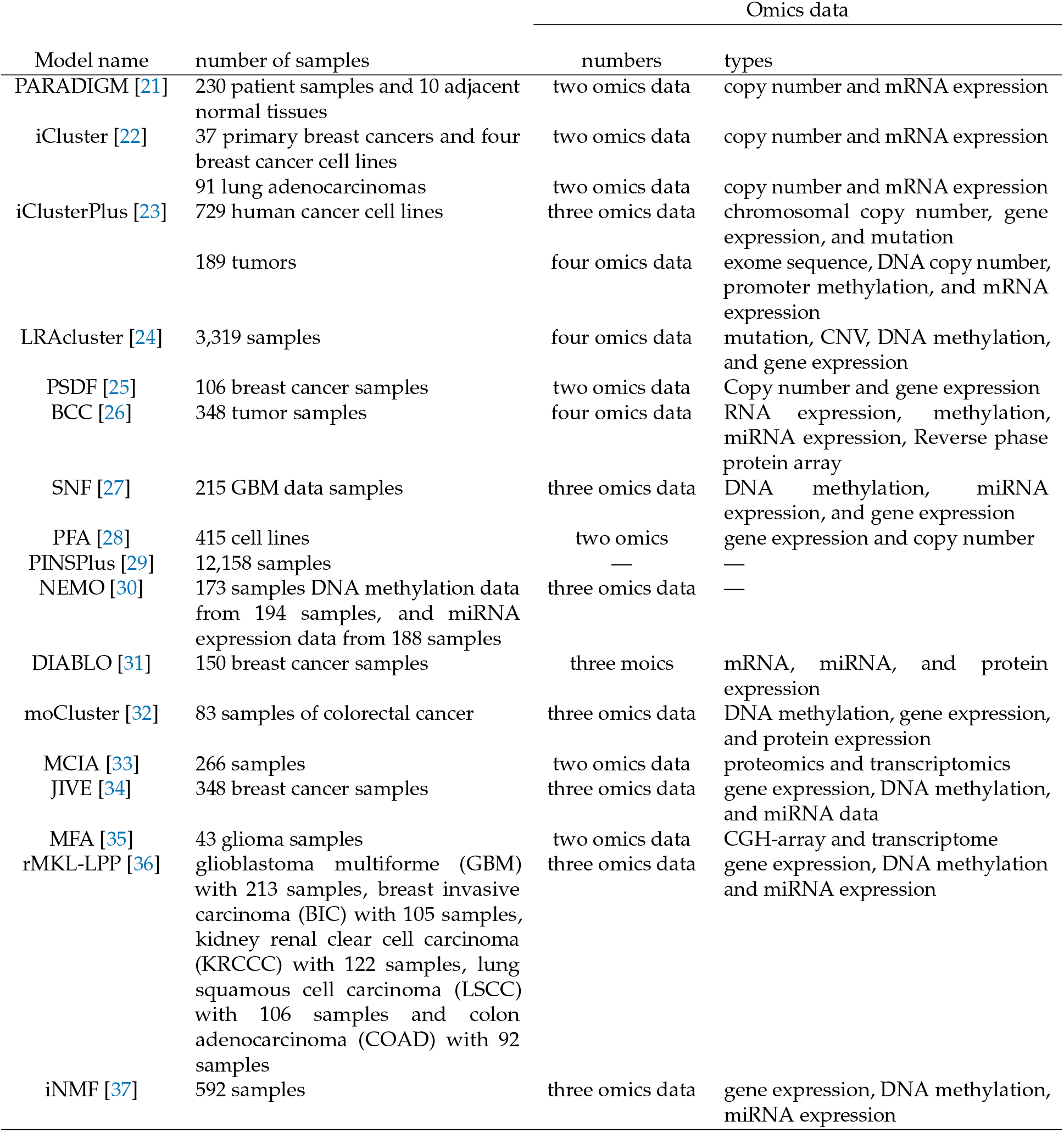
List of number of sample as well as types of omics data analyzed in the review [20]

The way by which multiomics data is formatted as a tensor as follows. Suppose that 𝒳 ∈ ℝ^*N*×*M*×*K*×*S*^ composed of *x*_*ijks*_ that represents *j*th kind of omics data (e.g., gene expression, promoter methylation, histone modification etc) attributed to *i*th gene of *s*th biological replicates of *k*th organism. In this case, for *k*th patients, *M* kind of omics data was measured using microarray or next generation sequencing technology measuring variables attributed to *N* genes at once. Thus, the multiomics measurements are naturally formatted as a tensor without any special reformat.

The primary difference between PCA based unsupervised FE and TD based unsupervised FE is as follows. When condition of experiments is restricted to one, e.g., “patients vs healthy control”, PCA is very useful. However, if it is a combination of multiple ones, e.g., “patients vs healthy controls” and “multiple tissues”, since tensor is more reasonable format than matrix, TD based unsupervised FE is more suitable method than PCA based unsupervised FE.

## 2. Materials and Methods

### 2.1. Synthetic data

In the synthetic dataset, we assumed that there are 81 samples composed of three tissue subclasses (each class has three replicates) and three disease subclasses (each class also has three replicates). That is, the 81 samples are composed of all possible pairs of one of the nine tissue samples (= three tissues × three replicates) and one of the nine disease samples (= three diseases × three replicates). Typically, the three tissues are supposed to be, for example, heart, brain, and skin, while the three diseases are supposed to be a few exclusive diseases, and healthy controls; this means that we do not have any biological knowledge of which combinations are more likely to be associated with one another. The number of genes, which are features, is as many as 10^5^; although so-called protein-coding genes are at most a few tens of thousands, a higher number of non-coding genes have come to be considered. Unlike the synthetic data, in the real dataset to which the methods are applied in the later part of this paper, the number of features corresponds to the number of genomic regions with fixed DNA sequence length. Thus, it can be even higher than that of the synthetic data. Among these 10^5^ features, only the first 100 features are supposed to be associated with distinction between diseases and tissues. The purpose of the analysis is to identify these 100 features correctly based upon the available datasets in a fully data-driven approach. To emulate this situation, we introduced tensor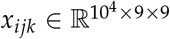, where *i, j, k* stand for features, tissues, and diseases, as

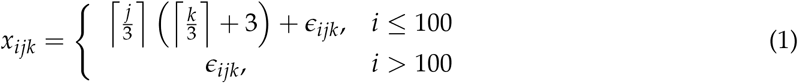

where *∈*_*ijk*_ ∼ *N* (0, 3), *N* (*µ, σ*) is a Gaussian distribution with mean *µ* and standard deviation *σ*, and *x* is a ceiling function that gives the smallest integer that is not less than *x*. The *x*_ijk_ values are classified into nine subclasses dependent upon pairs 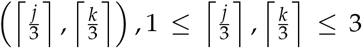 because 1 ≤ *j, k* ≤ 9.

The synthetic data set intentionally model the simplest cases of integrated analysis of multiple omics data set. *j* and *k* are assumed to be two independent omics measurements while replicates are supposed to be biological replicates in real experiments.

All performances reported in this study is averaged value over one hundred independent trials.

### 2.2. Real dataset

The real dataset consists of multiomics measurements of prostate cancer retrieved from the Gene Expression Omnibus (GEO) [38], using GEO ID GSE130408. The processed file named GSE130408_RAW.tar was downloaded and individual files with names starting with “GSM” were extracted from it. The individual files are composed of eight omics measurements: three transcription factor bindings, AR, FOXA1, and HOXB13; four histone modifications, H3K27AC, H3K27me3, H3K4me3, and K4me2; and ATAC-seq. Two additional subclasses, tumor and normal tissue, were attributed to each. Because the number of biological replicates varies from subclass to subclass, we randomly selected the following number of ChIP-seq subclasses: two AR, four FOXA1, four HOXB13, ten H3K27AC, one H3K27me3, one H3K4me3, one K4me2, and one ATAC-seq. In total, 24 multiomics measurement groups were constructed. Each multiomics measurement group is composed of six samples, comprising two tissue subclasses (tumor and normal prostate), each of which has three biological replicates (Table 2).

**Table 2.** Sumary of real dataset

| Omics | number of |  |  |
| --- | --- | --- | --- |
|  | multiomics | tissues | biological replicates |
| AR | 2 | 2 | 3 |
| FOXA1 | 4 | 2 | 3 |
| HOXB13 | 4 | 2 | 3 |
| H3K27AC | 10 | 2 | 3 |
| H3K27me3 | 1 | 2 | 3 |
| H3K4me3 | 1 | 2 | 3 |
| K4me2 | 1 | 2 | 3 |
| ATAC | 1 | 2 | 3 |
| total | 24 | — | — |

The values in each file were averaged over 25,000 DNA sequence intervals, giving a total number of 123,817, as 3 × 10^9^, the total DNA length of the human genome, divided by 25,000 is equal to 120,000. The averaged values were treated as a dataset to be analyzed further.

### 2.3. Categorical regression

Categorical regression, also known as analysis of variance or ANOVA, is expressed as

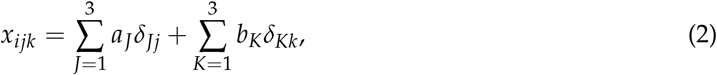

where *δ*_*Jj*_ and *δ*_*Kk*_ take 1 only when *j* or *k* belong to the *J*th tissue or *K*th disease subclass, respectively. *a*_*J*_ and *b*_*K*_ are regression coefficients (for synthetic data).

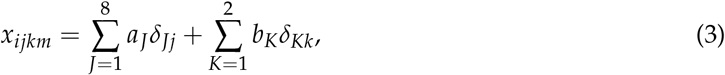

where *δ*_*Jj*_ and *δ*_*Kk*_ take 1 only when *j* or *k* belong to the *J*th multiomics measurement group or *K*th tissue (*K* = 1: tumor, *K* = 2: normal prostate) subclass, respectively. *a*_*J*_ and *b*_*K*_ are regression coefficients (for real data).

The computation was performed by the lm function in the base package in R [39]. The computed *P*-values were adjusted by the BH criterion [17] and the *i*th features associated with adjusted *P*-values less than 0.01 were selected.

### 2.4. RF

RF was performed using the randomForest function implemented in the randomForest package [40] in R. Synthetic and real data were regarded as 3 × 3 = 9 and 8 × 2 = 16 subclasses, respectively, each of which corresponds to one of the (*J, K*) pairs defined in Eqs. (2) or (3). Features included in the out-of-bag error were selected that had non-zero importance given by the importance function implemented in the randomForest package.

### 2.5. PenalizeLDA

PenalizedLDA was performed using the PenalizedLDA.cv function implemented in the PenalizedLDA package [41] in R. Synthetic and real data were regarded as 3 × 3 = 9 and 8 × 2 = 16 subclasses, respectively, each of which corresponds to one of the (*J, K*) pairs defined in Eqs. (2) or (3). *λ* was taken to be 0.01, 0.02, 0.03, 0.04 for synthetic data. PenalizedLDA could not be performed for the real dataset because of the zero in-subclass standard deviations of some features (for more details, see [18]).

### 2.6 TD-based unsupervised FE

Although the process was fully described in the recently published book [17], here we propose a variant, which is outlined briefly. First, a dataset must be formatted as a tensor. For synthetic data,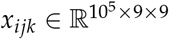. For real data, 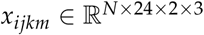 represents the averaged value of the *i*th interval of the *j*th omics measurement group of the *k*th tissue (*k* = 1: tumor, *k* = 2: normal prostate) of the *m*th biological replicates (1 ≤ *m* ≤ 3). Here, *N* = 123,817 is the total number of genomic regions of 25,000 DNA sequence length.

Higher-order singular value decomposition (HOSVD) [17] was then applied to the tensor, obtaining

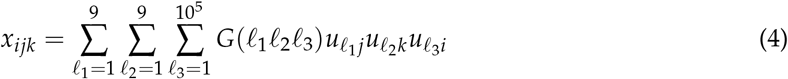

for synthetic data. 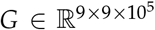 is a core tensor and 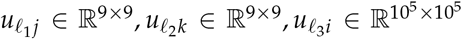 are singular value matrices that are orthogonal. Conversely, for the real dataset,

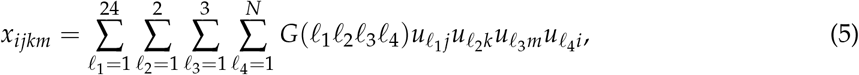

where *G* ∈ ℝ ^24×2×3×*N*^ is a core tensor and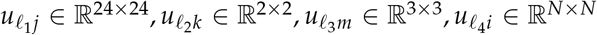 are singular value matrices that are orthogonal.

The first step is to check which singular value vectors attributed to sample subclasses (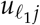attributed to tissue subclasses and *u*_*𝓁k*_ to disease subclasses for the synthetic dataset, and 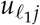 attributed to multiomics measurement groups and 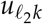 to tissues for the real dataset) represent distinction between samples of interest. After identifying biologically interesting singular value vectors attributed to samples (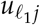 and 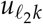 for the synthetic data, and 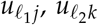 and 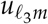 for the real data), we next attempt to find singular value vectors attributed to features (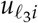 for the synthetic data and 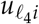 for the real data) that share core tensor *G* with larger absolute values with those identified singular value vectors attributed to samples. For example, suppose that 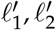 are selected for synthetic data and 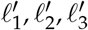 are selected for real data. In this case we seek *𝓁*_3_ that has 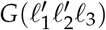 with larger absolute values for the synthetic data and *𝓁*_4_ that has 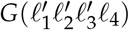 with larger absolute values for the synthetic data. Next, using the selected 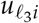 and 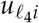, *P*-values are attributed to the *i*th feature as [17]

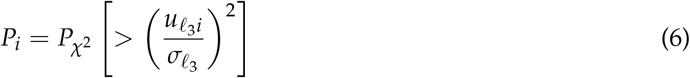

for synthetic data and

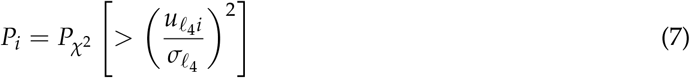

for real data.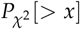 is the cumulative probability of *𝒳* ^2^ distribution that the argument is larger than *x*. 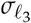 and 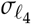are the standard deviations. The computed *P*-values are corrected by the BH criterion and features associated with adjusted *P*-values less than 0.01 are selected.

### 2.7 MNMF

Suppose that we have multiple matrices sharing the same number of rows, i.e.,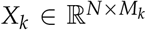. When there are no non-positive components in these matrices, we can apply MNMF as

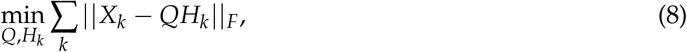

where *Q* ∈ R^*N*×*n*^ and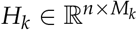, the column and row vectors of which are *n* latent variable vectors attributed to *i*s and samples, respectively, and there are no non-positive components in *Q* and *H*_*k*_s. || · ||_*F*_ is the Frobenius norm. *n* is an integer smaller than any *N* or *M*_*k*_. MNMF can be easily performed by applying non-negative matrix factorization (NMF) to the contraction matrix of 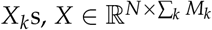, which is an unfolded matrix of tensors in this study (see Fig. 1 and below), because

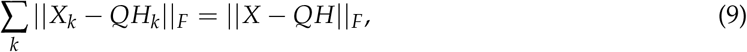

where 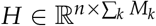 is a contraction matrix of *H*_*k*_s. NMF was performed by the nmf function in the NMF package of R.

When MNMF is applied to the synthetic dataset, it is applied to the matrix 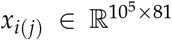 generated by unfolding tensor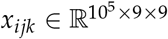. When MNMF is applied to the real dataset, it is applied to the matrix *x*_*i*(*jkm*)_ ∈ R^123817×144^ generated by unfolding tensor *x*_*ijkm*_ ∈ R^123817×24×2×3^. Because NMF cannot be applied to matrices with non-positive values, min(*x*_*ijk*_) are extracted from *x*_*ijk*_ so that NMF can be applied to matrices without non-positive values when applied to synthetic data. When NMF is applied to the real dataset, 1 × 10^−10^ is added to *x*_*ijkm*_ when *x*_*ijkm*_ = 0 (no negative values are included in *x*_*ijkm*_).

### 2.8. PCA based unsupervised FE

Although the process was fully described in the recently published book [17], we briefly explain the method. PCA was applied to matrices to whch MNMF was applied in the previous subsection so that PC scores are attributed to *i* whereas PC loading applied to either (*j*) (for synthetic data) or (*jkm*) (for real data). After identifying PC loading of interest, using corresponding PC score, *P*-values are attributed to *i*s as [17]

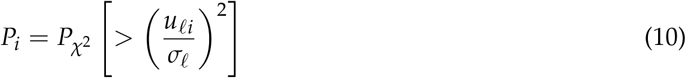

where *u*_*𝓁i*_ is the *𝓁*th PC score that corresponds to the PC loading of interest. P-values are corrected by BH criterion and *i*s associated with adjusted P-values less than 0.01 are selected.

## 3. Results

Figure 1 shows the flowchart of analyses performed in this study.

**Figure 1.**
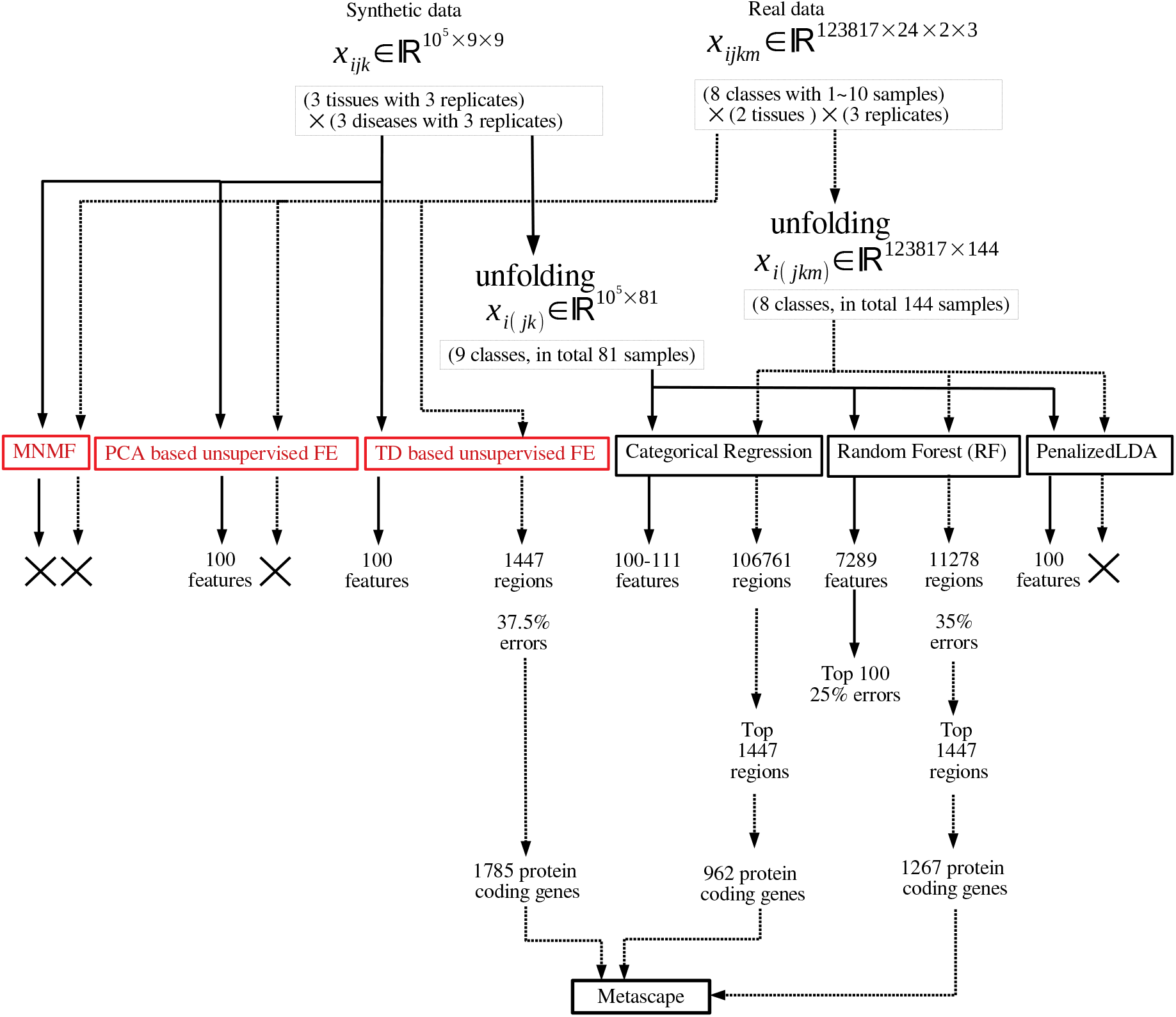
Flowchart of analyses performed in this study. Those in red and black are unsupervised and supervised feature selection, respectively.

### 3.1. Synthetic data

We applied RF, PenalizedLDA, categorical regression, TD-based unsupervised FE, and PCA based unsupervised FE to the synthetic dataset. Please note that all performances reported in the below are averaged over one hundred independent trials as denoted in Materials and Methods. When RF was applied to the synthetic dataset, there were up to 8456 features in average with non-zero importance, although the 100 features with subclass dependence were correctly selected. Thus, RF has the ability to select all the correct features. Nevertheless, because it selects too many false positives, it is ineffective. Reflecting this poor ability, as small as 16 samples among 81 samples were correctly classified in average. Although one might wonder if we select top 100 features with larger importance values provided by random forest, only as small as 73 features are correctly selected among top ranked 100 features by random forest. In contrast, when PenalizedLDA was applied to the synthetic dataset, the 100 features with subclass dependence were selected without any false positives being selected. Thus, PenalizedLDA can outperform RF. The only disadvantage is that we have to find the optimal *λ*, which is the weight coefficient of the L1 norm. Through massive trials and errors, we found that when *λ* is taken to be 0.01, it achieves the ideal performance mentioned above for all of one hundred independent trials. Reflecting this, PenalizedLDA can classify 81 samples with the accuracy of as high as 84 % in average. In contrast, if we select *λ* as 0.02, it selects as small as only nine features in average, although no features with subclass dependence are correctly selected for *λ* = 0.03 and 0.04. If we do not have prior knowledge that the correct number is 100, it is impossible to tune the optimal parameter. Thus, in this sense, PenalizedLDA cannot be regarded as a perfect method, although it can achieve perfect performance if we can successfully tune the optimal parameter. Categorical regression is far superior to PenalizedLDA, as it achieves good performances regardless of the threshold *P*-value (Table 3). It exhibits a maximum of just 11 false positives even if the threshold *P*-values vary from 0.01 to 0.1. This is in significant contrast to PenalizedLDA that requires sophisticated parameter tuning.

**Table 3.** Confusion matrices when categorical regression is applied to synthetic data. *P* is the threshold *P*-value. Features are selected when adjusted *P*-values less than the threshold P-values are associated with them. Averaged over 100 independent trials.

| $P$ | 0.1 | | 0.05 | | 0.01 | | 0.001 | |
| --- | --- | --- | --- | --- | --- | --- | --- | --- |
| | $i > 100$ | $i \leq 100$ | $i > 100$ | $i \leq 100$ | $i > 100$ | $i \leq 100$ | $i > 100$ | $i \leq 100$ |
| not selected | 99889 | 0 | 99895 | 0 | 99899 | 0 | 99900 | 0 |
| selected | 11 | 100 | 5 | 100 | 1 | 100 | 0 | 100 |

Nevertheless, TD-based unsupervised FE generated good performance results when compared against categorical regression. Initially, after applying HOSVD to the synthetic dataset, we noticed that *u*_1*j*_ and *u*_2*k*_ were associated with distinction between the three subclasses (Fig. 2). Thus, we attempted to find which *𝓁*_3_ is associated with *G*(1, 1, *𝓁*_3_) of the largest absolute value (Fig. 3). From this, it was obvious that *G*(1, 1, 1) had the largest absolute values. Equation (6) with *𝓁*_3_ = 1 was used to attribute *P*_*i*_ to the *i*th feature. *P*_*i*_ was corrected by the BH criterion and features associated with adjusted *P*-values less than the threshold values were selected (Table 4). As a result, TD-based unsupervised FE achieved perfect performances regardless of the threshold *P*-values.

**Figure 2.**
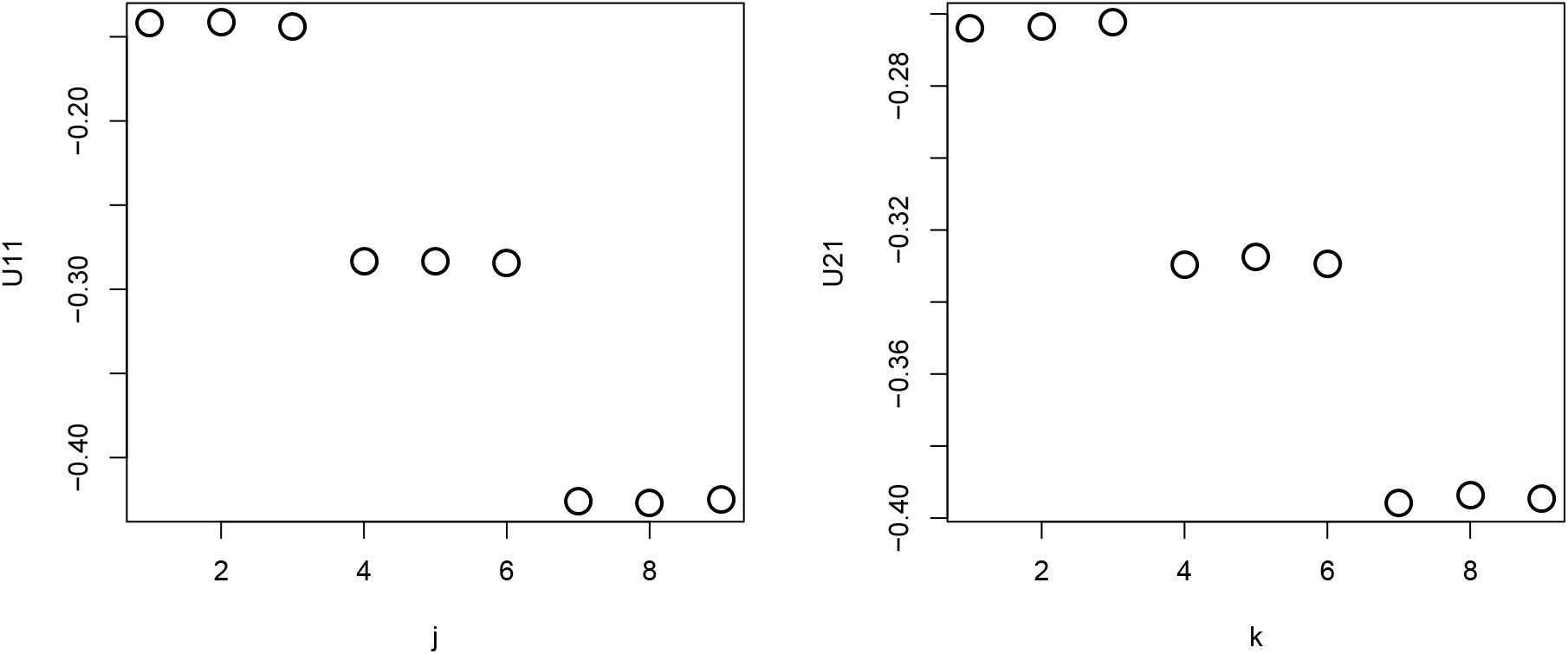
*u*_1*j*_ and *u*_1*k*_ obtained when HOSVD is applied to the synthetic dataset. Averaged over 100 independent trials.

**Figure 3.**
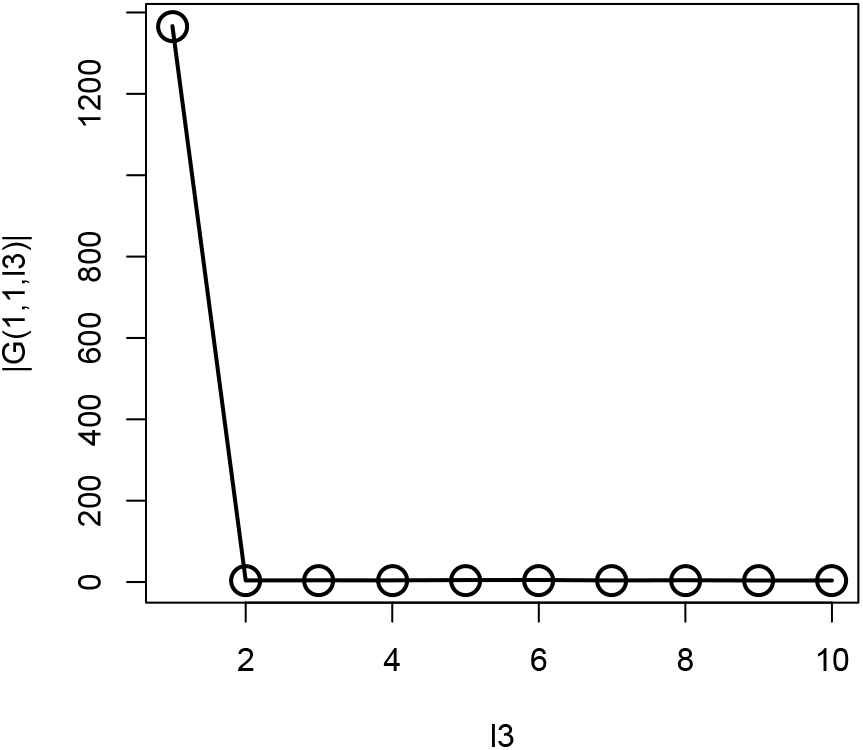
*G*(1, 1, *𝓁*3) obtained when HOSVD is applied to the synthetic dataset. Averaged over 100 independent trials.

**Table 4.**
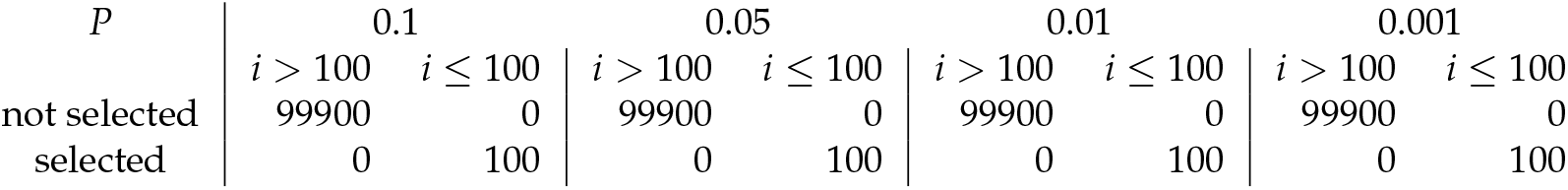
Confusion matrices when TD-based unsupervised FE is applied to the synthetic data. *P* is the threshold *P*-value. Features are selected when adjusted *P*-values less than the threshold P-values are associated with them. 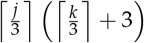. Averaged over 100 independent trials.

### 3.2. Real data

Next, we apply TD-based unsupervised FE to a real dataset, as it was shown to be the best of the four methods tested on synthetic data in the previous subsection. After applying HOSVD to *x*_*ijkm*_, we realized that *u*_1*j*_ and *u*_2*j*_ have significant biological dependence upon multiomics measurement groups (Fig. 4). To explain why *u*_1*j*_ and *u*_2*j*_ are biologically reliable, we need to introduce genomic science briefly as follows:

**Figure 4.**
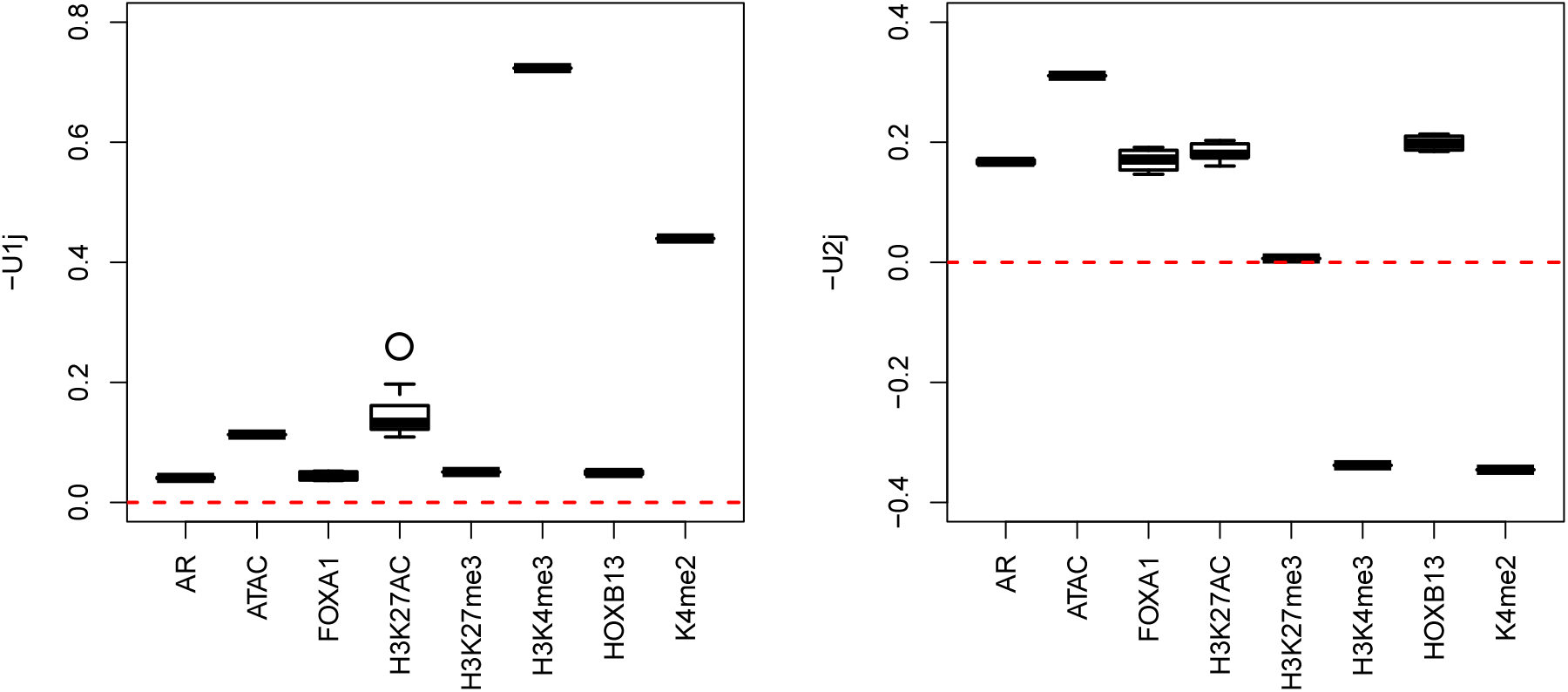
Boxplot of *u*_1*j*_ and *u*_2*j*_ when HOSVD is applied to the real dataset.

A genome, which is a sequence of DNA, stores the information to be translated into a protein. Because all cells store the same genome, the distinct functions and structures of individual cells are controlled by whichever parts of the genome are translated into proteins through transcription to RNA. This process is controlled by the various proteins that bind to genomes. In these measurements, the amounts of three such proteins, AR, FOXA1, and HOXB13, are measured. Another factor that controls this process is the small molecules that bind to histone. Histone is a core protein around which DNA wraps itself, and the tightness of this wrapping is the factor that controls the process from DNA to RNA. H3K27AC, H3K27me3, H3K4me3, and K4me2 are the measures of which small molecules bind to which parts of histone. Finally, ATAC-seq measures how “open” the DNA is. As DNA becomes more open, it can produce more RNA. Thus, the number of proteins that bind to DNA, the number of small molecules that bind to histone, and ATAC-seq are the measures of how many genomes are transcribed to RNA. The difficulty in multiomics analysis is to identify the combinations of proteins binding to DNA and small molecules binding to histone that can enhance the openness of DNA, which is assumed to be observed by ATAC-seq. As we do not know what combination is likely to be observed, the supervised approach is not easy to employ.

We now explain why *u*_1*j*_ and *u*_2*j*_ are biologically reliable. H3K4me3 and H3K27ac are known to be active/open chromatin marks, whereas H3K27me3 is known to be an inactive mark [42]. *u*_1*j*_ has higher values for H3K4me3 and H3K27ac but smaller values for H3K27me3, with larger values for ATAC-seq, which measures the amount of openness of genome. Thus, the dependences of *u*_1*j*_ upon H3K4me3, H3K27ac, H3K27me3, and ATAC-seq are reliable. H3K4me2 [43] is also known to activate gene transcription in tissue-specific ways. The overall dependence of *u*_1*j*_ upon multiomics measurement groups is reasonable. Nonetheless, three proteins, AR [44], FOXA1 [45], and HOXB13 [46], are also known to mediate prostate cancer progression, and their enrichment corresponded with *u*_2*j*_. We decided that *u*_1*j*_ and *u*_2*j*_ successfully captured the biological aspects of multiomics measurements.

In contrast, *u*_2*k*_ represents the distinction between a tumor (*k* = 1) and normal prostate (*k* = 2), while *u*_1*m*_ represents the independence of biological replicates (Fig. 5). As we are seeking multiomics measurements that are distinct between a tumor and normal prostate as well as common between biological replicates, *u*_2*j*_ and *u*_1*m*_ represent the properties that we seek. Although we can select either *𝓁*_1_ = 1 or *𝓁*_1_ = 2, as both are biologically reliable, we select *𝓁*_1_ = 1 because it is the first (i.e., mathematically primary) factor. To find the 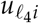 used for attributing *P*-values to genes *i*, we seek *G*(1, 2, 1, *𝓁*_4_) with the largest absolute value (Fig. 6).

**Figure 5.**
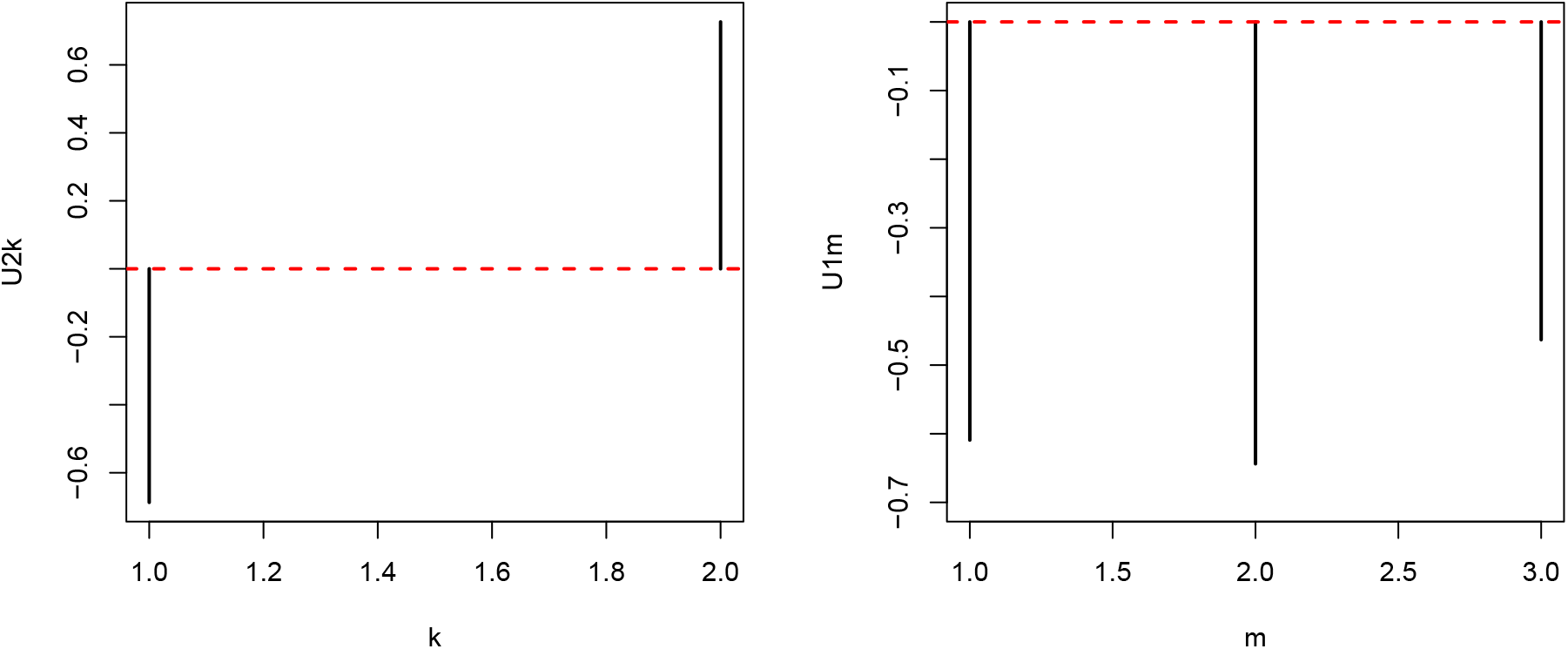
*u*_2*k*_ and *u*_1*m*_ when HOSVD is applied to the real dataset.

**Figure 6.**
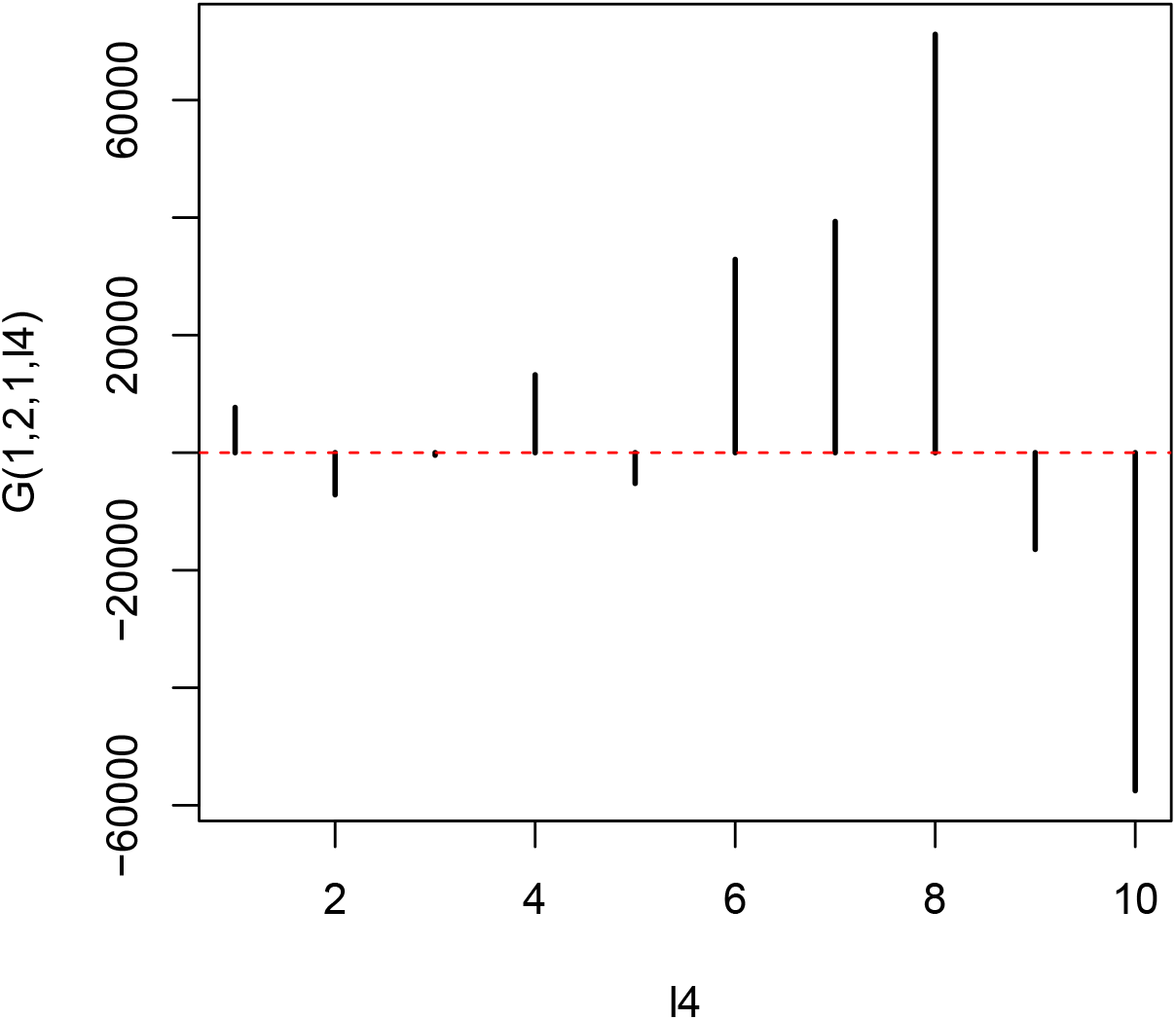
*G*(1, 2, 1, *𝓁*_4_) when HOSVD is applied to the real dataset.

It is obvious that *G*(1, 2, 1, 8) has the largest absolute value. Hence, *u*_8*i*_ is employed to assign *P*-values to the *i*th genomic regions using (7) with *𝓁*_4_ = 8. The *P*_*i*_ values are corrected using the BH criterion, and 1447 genomic regions associated with adjusted *P*_*i*_ less than 0.01 are selected from the total 123,817 genomic regions. This indicates that TD-based unsupervised FE selects approximately 2% of genomics regions of the whole human genome.

Various biological evaluations of selected regions are possible, one of which is the number of protein-coding regions included in these 1447 selected regions. As the genomic regions that include protein-coding genes comprise less than a few percent of the human genome, and the total number of human protein-coding genes is up to 2 × 10^4^, the expected number of protein-coding genes included in these 1447 selected genomic regions is at most a few hundred. Nevertheless, as many as 1785 protein-coding genes can be counted in these regions, which is much higher than expected. This indicates that TD-based unsupervised FE can select genomic regions that include protein-coding genes, correctly considering the altered multiomics variables between normal and tumor tissues, although non-coding RNAs (ncRNAs) have also a key role in regulating the behavior of cells and their over- and underexpression strongly correlated with cancer.

Another evaluation of these selected 1447 genomic regions is the biological properties of the 1785 protein-coding genes included in these 1447 genomic regions. If biologically reliable protein-coding genes are selected, they should be enriched with the genes related to prostate cancer. To confirm this point, we uploaded these genes to Metascape [47], which evaluates the enrichment of various biological terms with reducing redundancy. Figure 7 shows the results of the DisGeNet [48] category of Metascape. It is obvious that prostatic neoplasms are not only top ranked but also outstandingly enriched. This suggests that TD-based unsupervised FE can specifically select genes related to prostate cancer. Figure 8 shows the results of the PaGenBase [49] category of Metascape. It is obvious that LNCaP cells are not only top ranked but also outstandingly enriched. LNCAP [50] is a model cell line of a human prostatic carcinoma. This also suggests that TD-based unsupervised FE can specifically select genes related to prostate cancer. Figure 9 shows the results of the TRRUST [51] category of Metascape. It is obvious that the androgen receptor (AR) is not only top ranked but also outstandingly enriched. As mentioned above, AR is a protein that binds to the genome and mediates prostate cancer progression. This also suggests that TD-based unsupervised FE can specifically select genes related to prostate cancer. Although there are more convincing results, the supplementary materials include the full report provided by Metascape.

**Figure 7.**
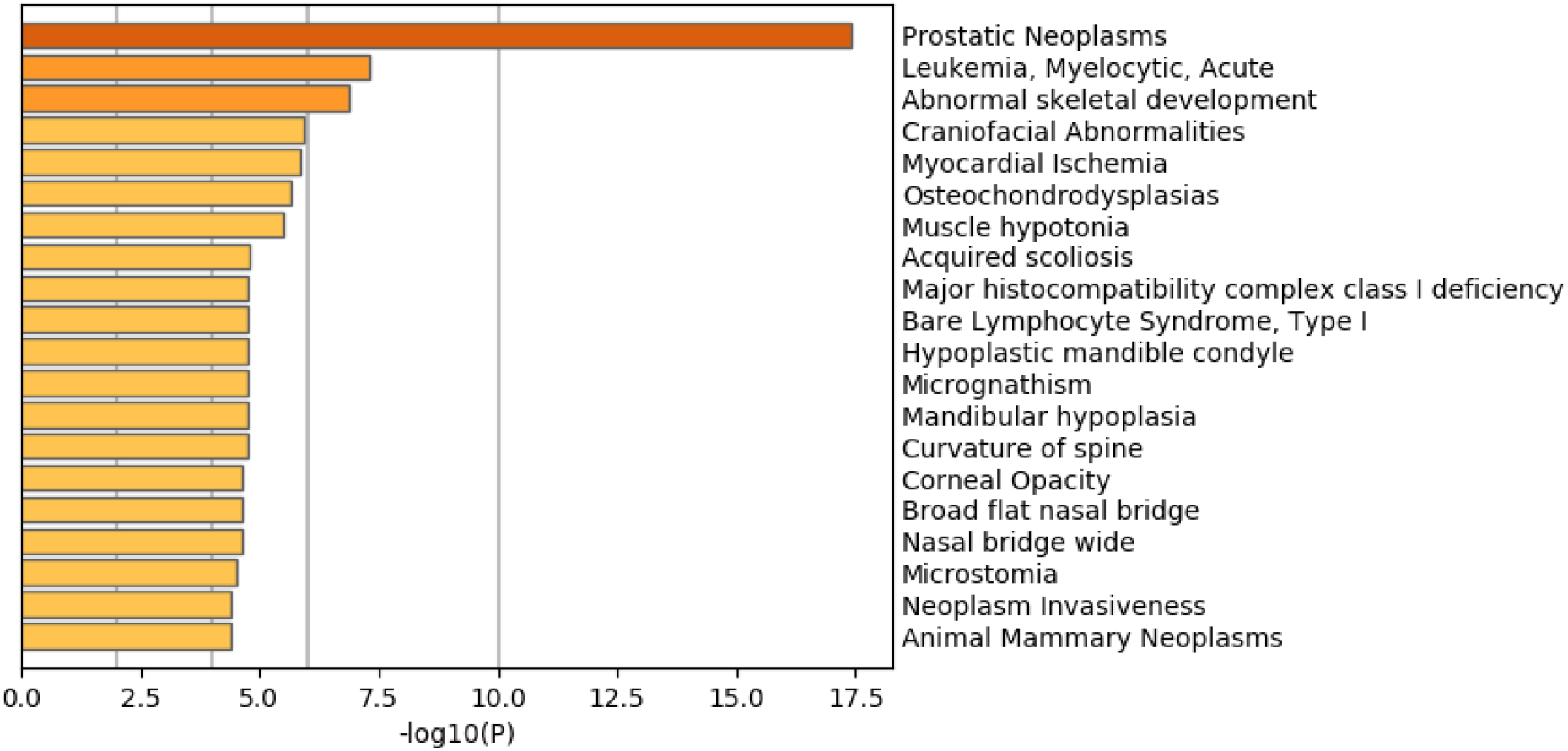
Summary of enrichment of the DisGeNet category of Metascape when 1785 protein-coding genes included in 1447 genomic regions selected by TD-based unsupervised FE are considered.

**Figure 8.**
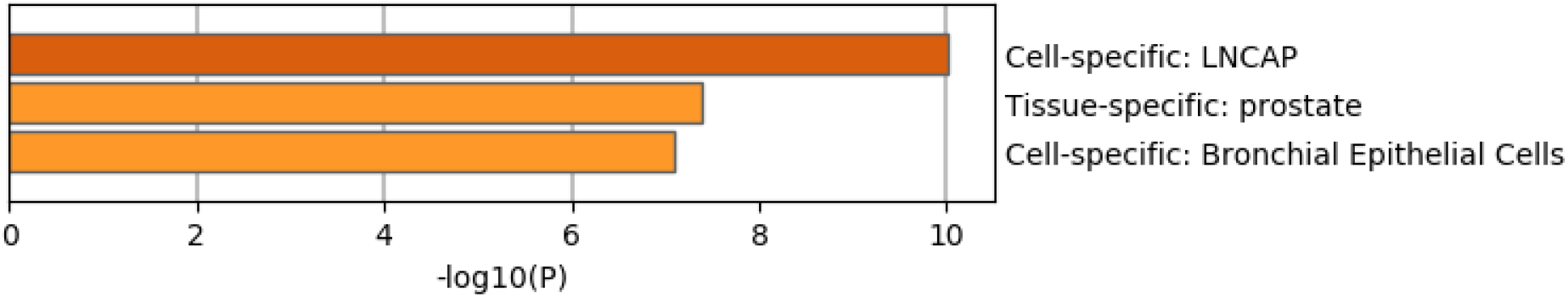
Summary of enrichment of the PaGenBase category of Metascape when 1785 protein-coding genes included in 1447 genomic regions selected by TD-based unsupervised FE are considered.

**Figure 9.**
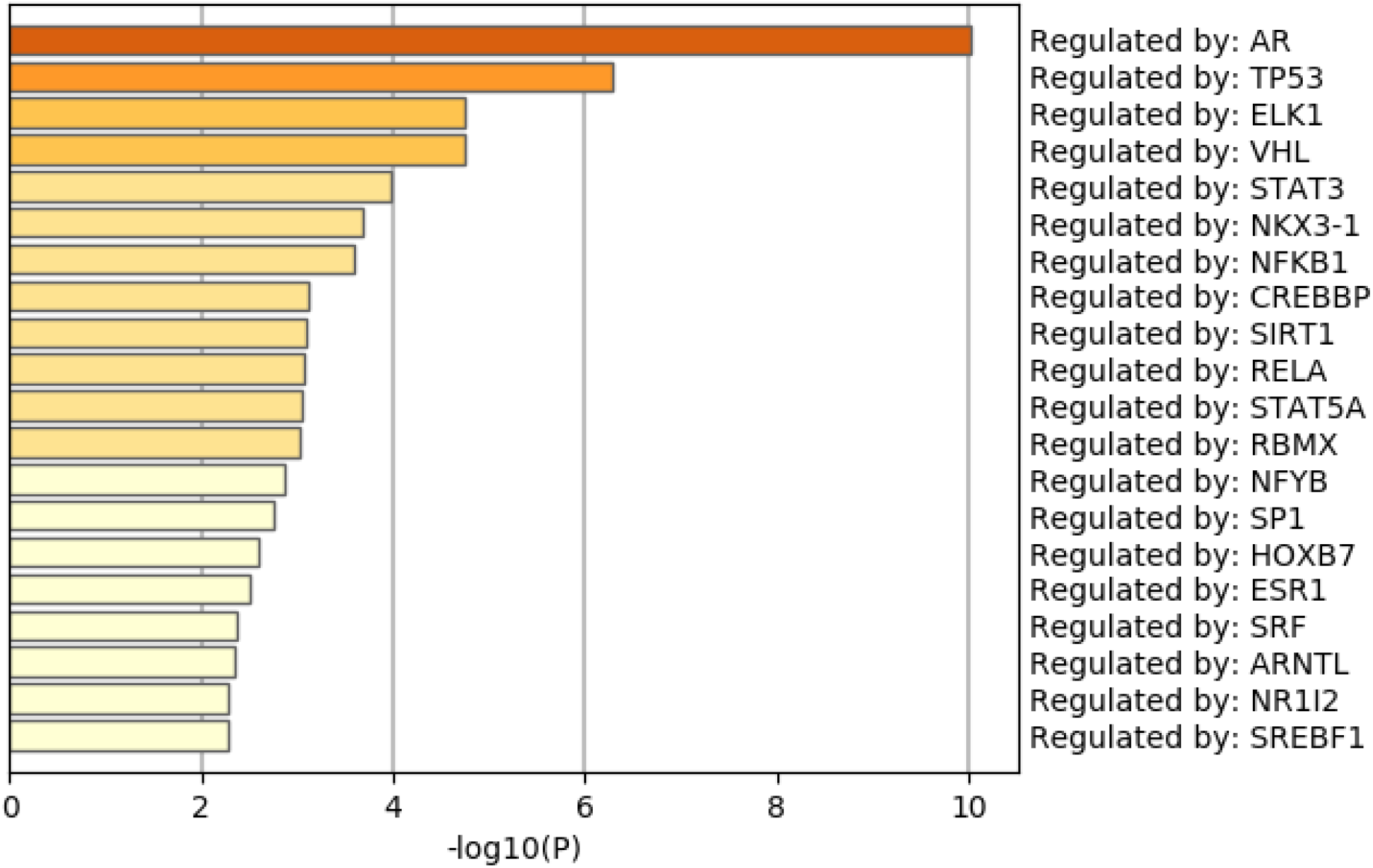
Summary of enrichment of the TRRUST category of Metascape when 1785 protein-coding genes included in 1447 genomic regions selected by TD-based unsupervised FE are considered.

## Discussion

### Synthetic data

To determine whether TD-based unsupervised FE outperformed the three supervised feature selections because of its unsupervised nature, we also applied an alternative unsupervised feature selection, MNMF, to the synthetic data, specifying *n* = 3 as the number of vectors each of which is composed of latent variables. Figures 10 shows the latent variable vectors attributed to the samples and *i*s, respectively, computed by MNMF. Up to the third latent vector, none is coincident with 9 classes defined in eq. (1). Thus, it is obvious that MNMF is inferior to TD based unsupervised FE. Thus the reason why TD based unsupervised FE could outperform other supervised methods is not simply because it is a unsupervised method.

**Figure 10.**
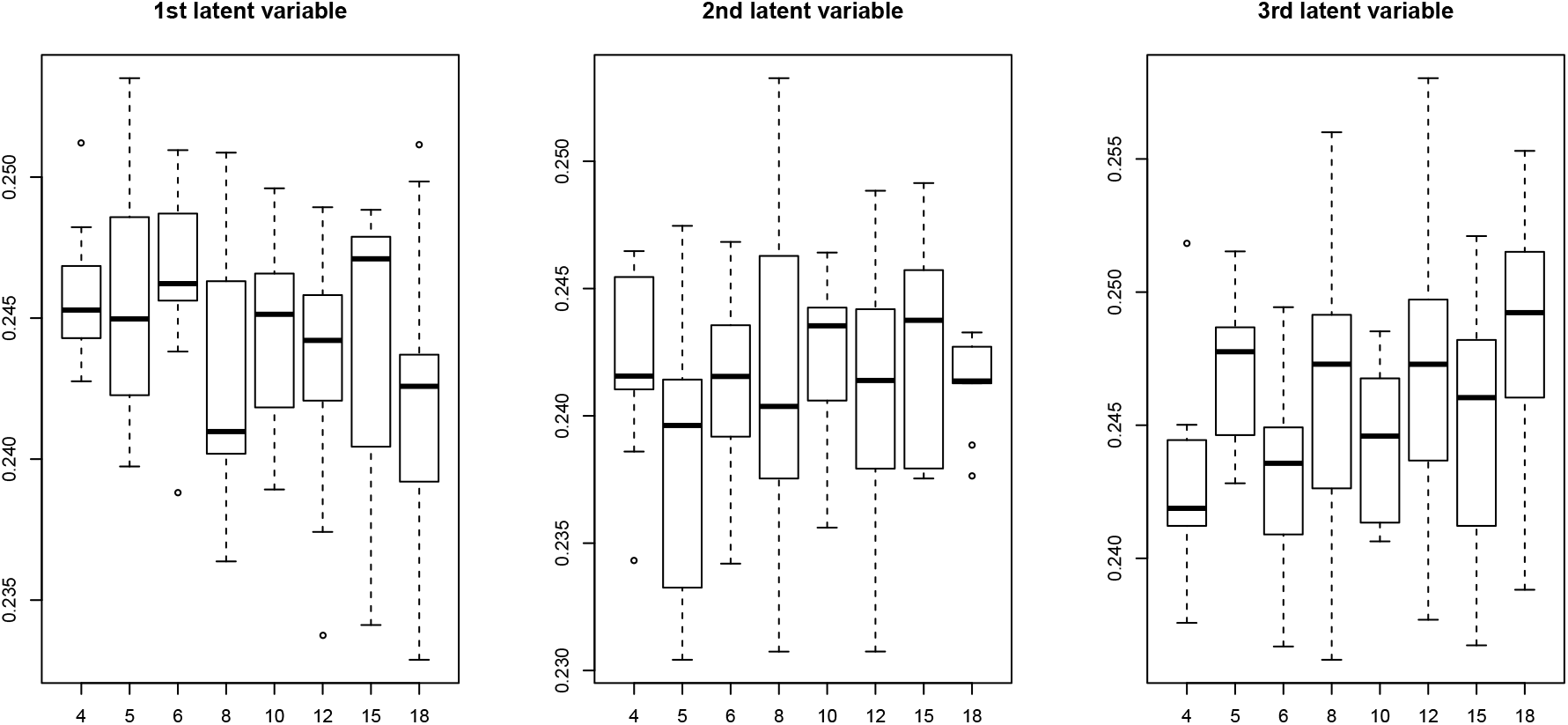
Boxplot of the first to third latent variable vectors attributed to the 81 samples. Horizontal axis: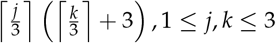. Averaged over 100 independent trials.

**Figure 11.**
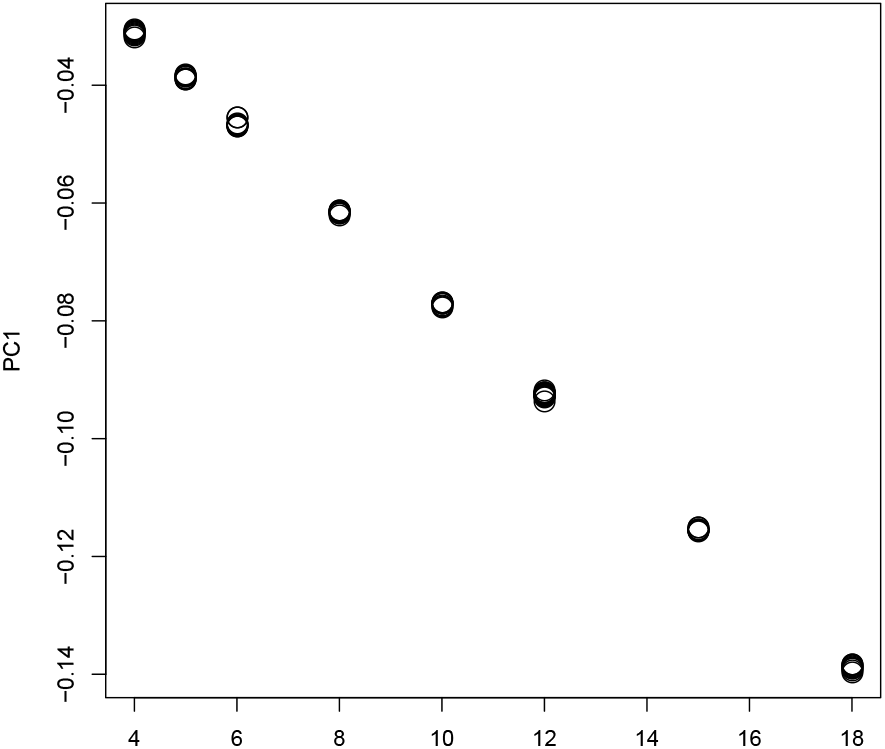
The 1st PC loading computed by PCA applied to synthetic data. Horizontal axis: 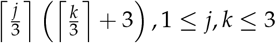. Averaged over 100 independent trials.

In order to see if TD based unsupervised FE is a only unsupervised method that can outperform other supervised method, we also tried PCA based unsupervised FE [17] from which TD based unsupervised FE developed. It is obvious that it is well coincident with :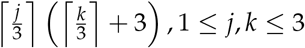. Then we attributed P-values to *i* using eq. (10) with *𝓁* = 1. Table 5 shows the confusion matrix when PCA based unsupervised FE was applied to synthetic data. It is identical to Table 4. Thus Thus, reported results show that TD and PCA-based unsupervised FE outperforms the other six methods tested on the synthetic dataset.

**Table 5.**
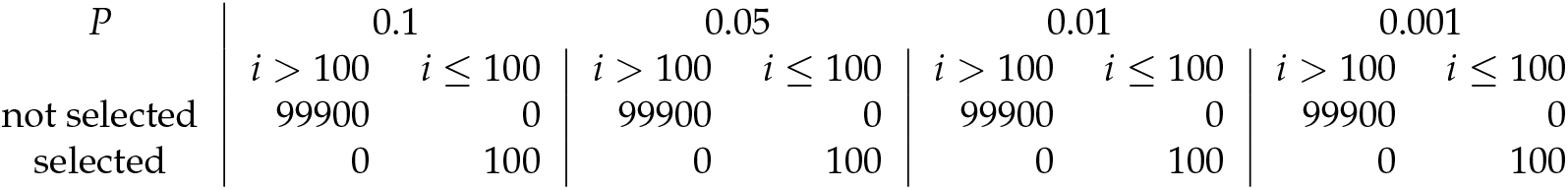
Confusion matrices when PCA-based unsupervised FE is applied to the synthetic data. *P* is the threshold *P*-value. Features are selected when adjusted *P*-values less than the threshold P-values are associated with them. 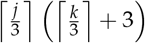.Averaged over 100 independent trials.

TD-based unsupervised FE also has the advantage of its CPU time (Table 6). PCA and TD-based unsupervised FE are the fastest of the six methods tested on the synthetic data. It can finish within a few seconds, whereas the others take longer, ranging from a few tens of seconds for categorical regression to a few minutes for RF. In this sense, PCA and TD-based unsupervised FE are the fastest, most accurate, and most robust of the six methods tested on the synthetic dataset.

**Table 6.**
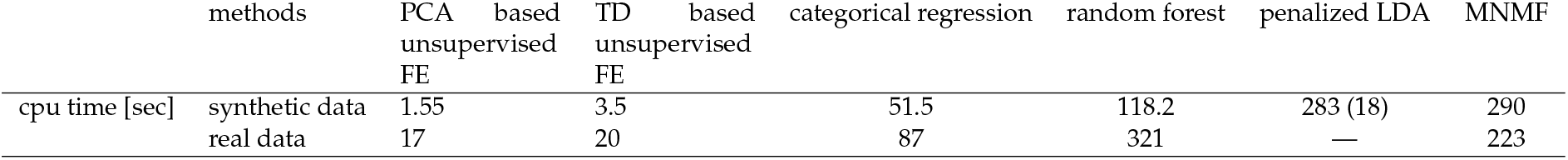
CPU times of the five methods required for the synthetic and real datasets. The CPU time for PenalizedLDA is for testing four *λ* values with between one and eight discriminant functions. The CPU time for an individual run is 18 s. Cpu times for synthetic data were averaged over 100 independent trials.

### Real data

We now compare the performances of TD-based unsupervised FE with those of the other five methods: RF, PenalizedLDA, categorical regression, MNMF and PCA based unsupervised FE. First, RF was applied to the real dataset, which is classified into 16 categorical classes composed of any pairs of eight mulitiomics measurement groups and two tissues (tumor and normal prostate). RF results in as many as 11,278 genomic regions having non-zero importance. This number is almost eight times larger than the number of regions selected by TD-based unsupervised FE (1447). One of the evaluations of RF performance is its accuracy of discrimination, as RF is a supervised method. RF should have selected the minimum features that can successfully discriminate between the 16 classes. The error rate given by RF was as small as 0.35 (Table 7). This indicates that 65% of the 144 samples were correctly classified into the 16 classes (see Table 2 for details of how the 144 samples were classified into the 16 subclasses). To validate the discrimination performances of the 1447 genomic regions selected by TD-based unsupervised FE, we carried out the following. *u*_1*j*_, *u*_2*k*_, *u*_3*k*_ were recomputed by HOSVD using only the 1447 genomic regions selected by TD-based unsupervised FE. The 144 samples were then discriminated using linear discriminant analysis, employing the product *u*_1*j*_*u*_2*k*_*u*_3*k*_ as input vectors (Table 8 with leave-one-out cross-validation). The error rate was as small as 37.5%, similar to that of RF (0.35). Considering the fact that TD-based unsupervised FE employed only one-eighth of the number of genomic regions selected by RF, we concluded that TD-based unsupervised FE can outperform RF.

**Table 7.** Confusion matrix for RF applied to the real dataset. N: normal prostate, T: tumor. The error rate is 35%.

|  | AR |  | ATAC |  | FOXA1 |  | H3K27AC |  | H3K27me3 |  | H3K4me3 |  | HOXB13 |  | K4me2 |  | class error |
| --- | --- | --- | --- | --- | --- | --- | --- | --- | --- | --- | --- | --- | --- | --- | --- | --- | --- |
|  | N | T | N | T | N | T | N | T | N | T | N | T | N | T | N | T |  |
| AR_N | 5 | 0 | 0 | 0 | 1 | 0 | 0 | 0 | 0 | 0 | 0 | 0 | 0 | 0 | 0 | 0 | 1/6 |
| AR_T | 0 | 0 | 0 | 0 | 1 | 3 | 1 | 0 | 0 | 0 | 0 | 0 | 0 | 1 | 0 | 0 | 1 |
| ATAC_N | 0 | 0 | 0 | 2 | 0 | 0 | 0 | 1 | 0 | 0 | 0 | 0 | 0 | 0 | 0 | 0 | 1 |
| ATAC_T | 0 | 0 | 2 | 1 | 0 | 0 | 0 | 0 | 0 | 0 | 0 | 0 | 0 | 0 | 0 | 0 | 2/3 |
| FOXA1_N | 1 | 0 | 0 | 0 | 9 | 0 | 0 | 0 | 0 | 0 | 0 | 0 | 2 | 0 | 0 | 0 | 1/4 |
| FOXA1_T | 0 | 0 | 0 | 0 | 4 | 5 | 0 | 0 | 0 | 0 | 0 | 0 | 0 | 3 | 0 | 0 | 7/12 |
| H3K27AC_N | 0 | 0 | 0 | 0 | 0 | 0 | 26 | 4 | 0 | 0 | 0 | 0 | 0 | 0 | 0 | 0 | 2/15 |
| H3K27AC_T | 0 | 0 | 0 | 0 | 0 | 0 | 6 | 24 | 0 | 0 | 0 | 0 | 0 | 0 | 0 | 0 | 1/5 |
| H3K27me3_N | 0 | 0 | 0 | 0 | 0 | 0 | 0 | 0 | 1 | 2 | 0 | 0 | 0 | 0 | 0 | 0 | 2/3 |
| H3K27me3_T | 0 | 0 | 0 | 0 | 0 | 0 | 0 | 0 | 1 | 2 | 0 | 0 | 0 | 0 | 0 | 0 | 1/3 |
| H3K4me3_N | 0 | 0 | 0 | 0 | 0 | 0 | 0 | 0 | 0 | 0 | 0 | 3 | 0 | 0 | 0 | 0 | 1 |
| H3K4me3_T | 0 | 0 | 0 | 0 | 0 | 0 | 0 | 0 | 0 | 0 | 3 | 0 | 0 | 0 | 0 | 0 | 1 |
| HOXB13_N | 0 | 0 | 0 | 0 | 2 | 0 | 0 | 0 | 0 | 0 | 0 | 0 | 9 | 1 | 0 | 0 | 1/4 |
| HOXB13_T | 0 | 0 | 0 | 0 | 0 | 0 | 0 | 0 | 0 | 0 | 0 | 0 | 1 | 11 | 0 | 0 | 1/12 |
| K4me2_N | 0 | 0 | 0 | 0 | 0 | 0 | 0 | 0 | 0 | 0 | 0 | 0 | 0 | 0 | 0 | 3 | 1 |
| K4me2_T | 0 | 0 | 0 | 0 | 0 | 0 | 0 | 1 | 0 | 0 | 0 | 0 | 0 | 0 | 2 | 0 | 1 |

**Table 8.**
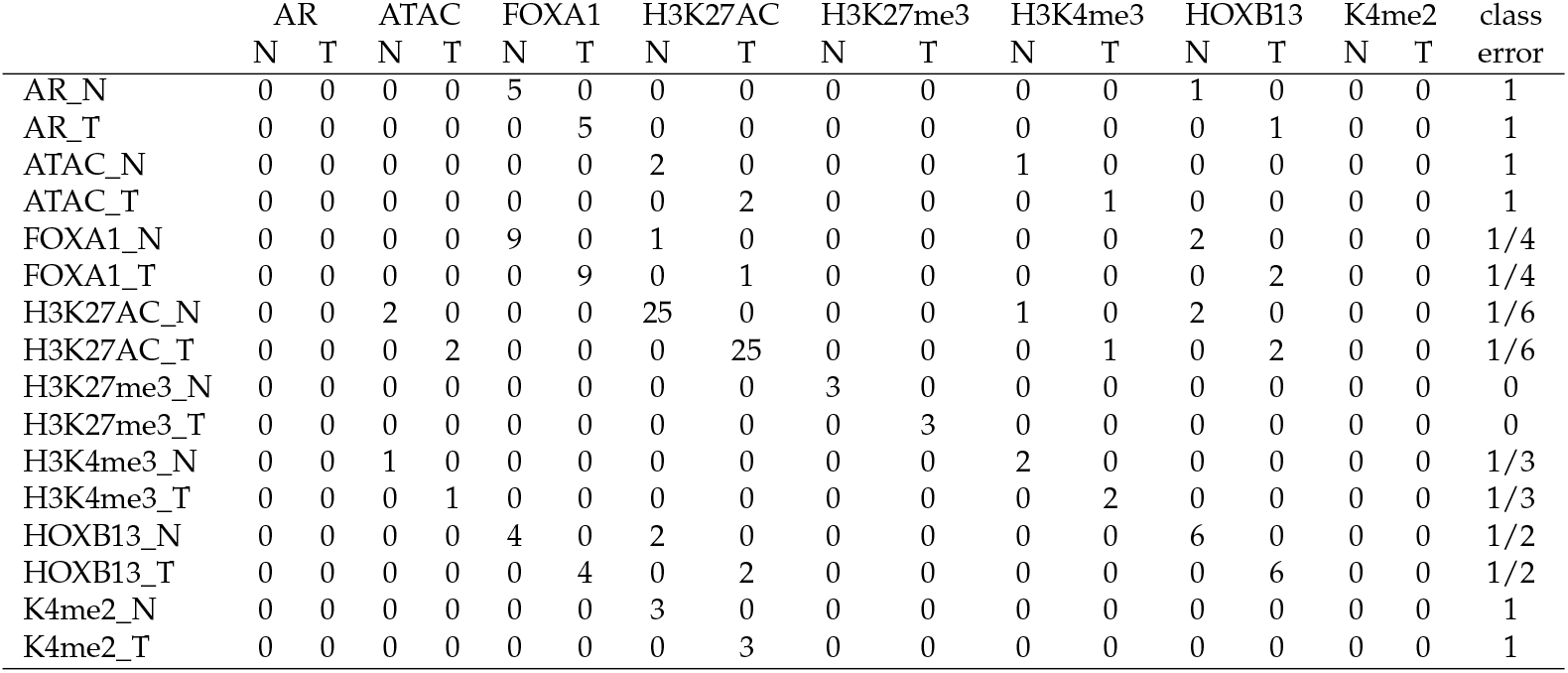
Confusion matrix for LDA using *u*_1*j*_*u*_2*k*_*u*3*m* given by TD applied to 1447 genomic regions selected by TD-based unsupervised FE applied to the real dataset. N: normal prostate, T: tumor. The error rate is 37.5%.

In spite of the comparisons presented in the above, one might wonder that RF can achieve comparative performances if top ranked restricted number of genomic regions are intentionally selected. In order to this, we selected 1447 top ranked regions (they are as many as those selected by TD based unsupervised FE) and identified as many as 1267 gene symbols included in these genomic regions. These 1267 genes are uploaded to Metascape to evaluate them biologically. Figure 12 shows the results of the DisGeNet category of Metascape. In contrast to Fig. 7 where prostatic neoplasms was top ranked, it was even not ranked at all in Figure 12. Thus, it is obvious that genes selected by RF is inferior to those by TD based unsupervised FE. Figure 13 shows the results of the PaGenBase category of Metascape. In cotrast to Fig. 8 where prostate related cell lines are top ranked, no prostate related cell lines are top ranked in Fig. 13. Thus, again, it is obvious that genes selected by RF is inferior to those by TD based unsupervised FE. Figure 14 shows the results of the TRRUST category of Metascape. In contrast to Fig. 9 where AR is top ranked, AR is not ranked at all. Thus, it is obvious that genes selected by RF is inferior to those by TD based unsupervised FE as well.

**Figure 12.**
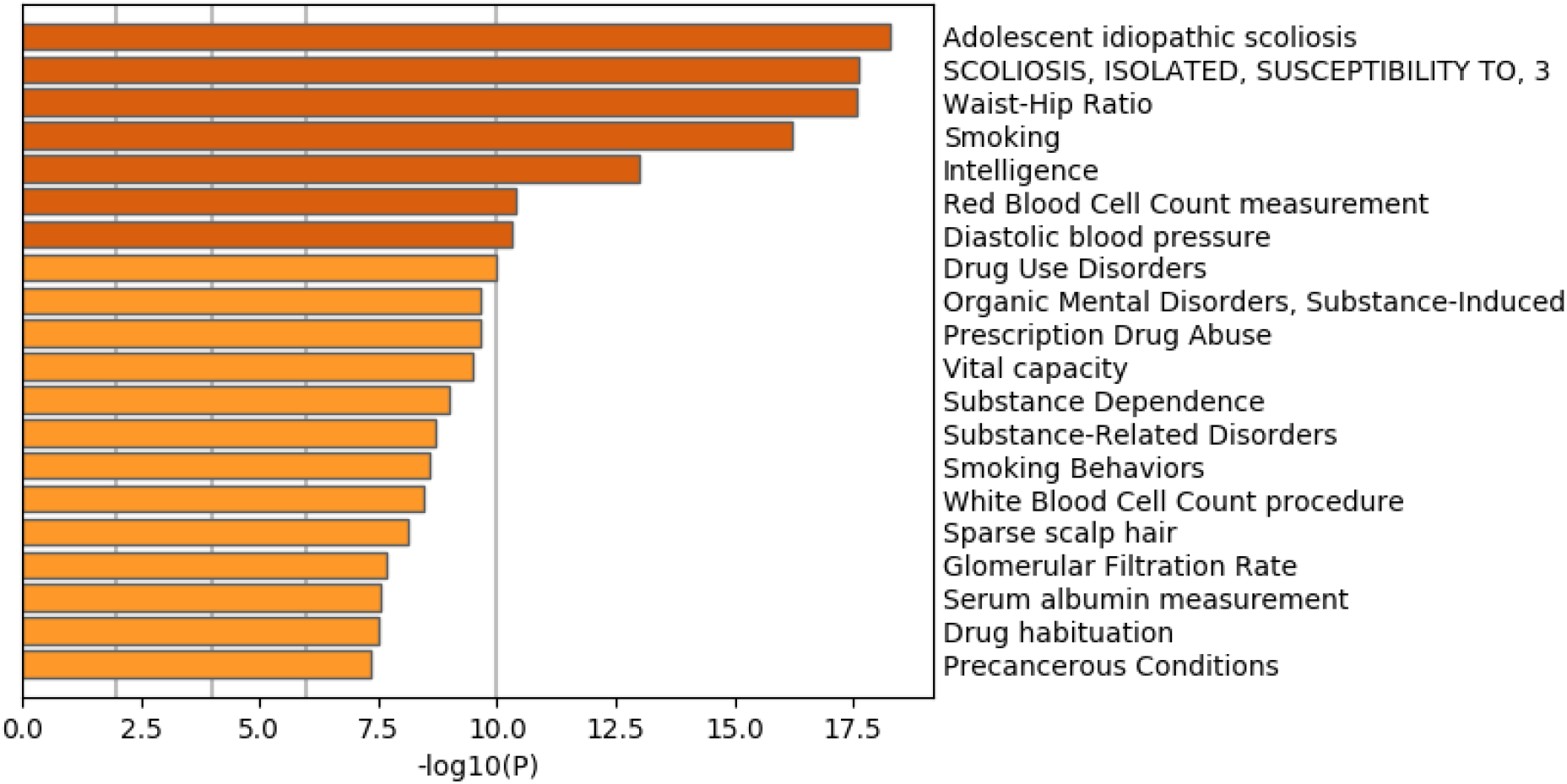
Summary of enrichment of the DisGeNet category of Metascape when 11267 protein-coding genes included in 1447 genomic regions selected by RF are considered.

**Figure 13.**
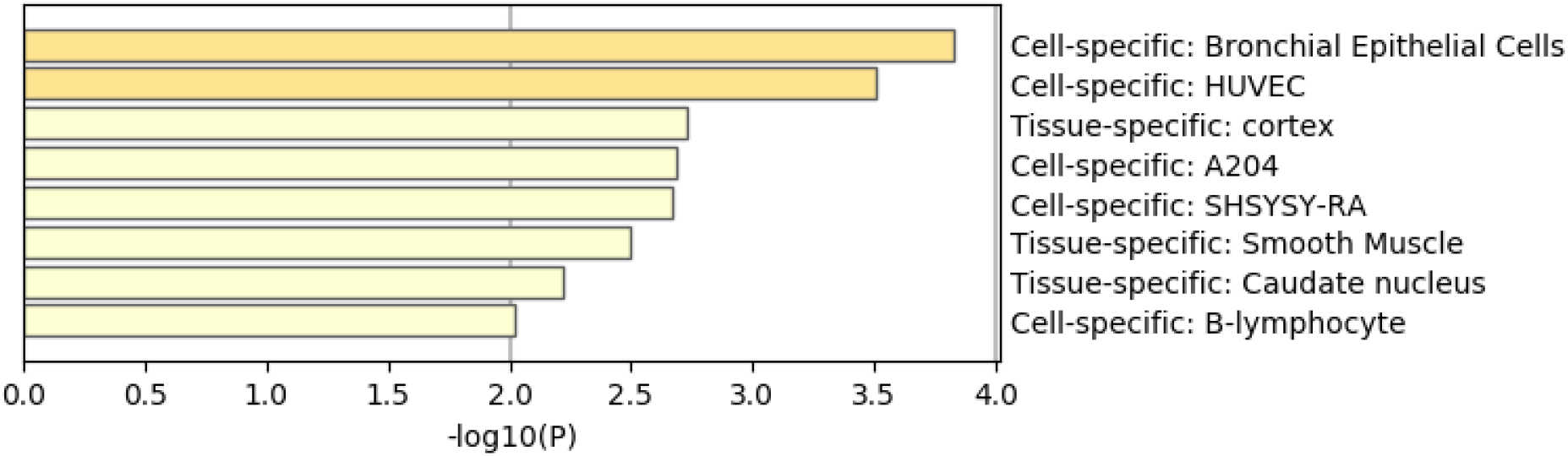
Summary of enrichment of the PaGenBase category of Metascape when 1267 protein-coding genes included in 1447 genomic regions selected by RF are considered.

**Figure 14.**
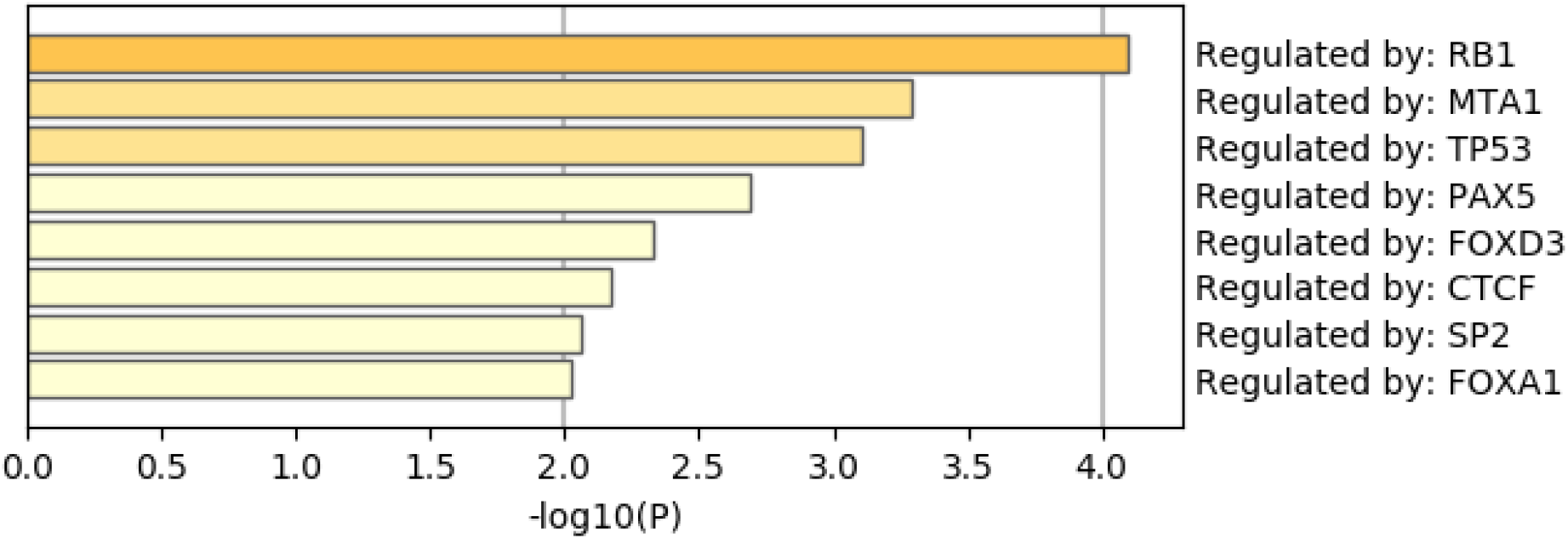
Summary of enrichment of the TRRUST category of Metascape when 1267 protein-coding genes included in 1447 genomic regions selected by RF are considered.

Next, we attempted to apply PenalizedLDA to the real dataset, but found that we could not. To perform LDA, all in-subclass variance must be non-zero. This condition was not fulfilled for the real data. Hence, PenalizedLDA cannot be employed to process real datasets.

Third, we applied categorical regression to the real dataset by regarding it as 16 categorical subclasses. The number of features associated with adjusted *P*-values less than 0.01 was as high as 106,761 among all 123,817 features. This indicates that categorical regression selected almost all features and is thus clearly ineffective. The reason categorical regression failed, despite being relatively successful (Table 3) when applied to the synthetic dataset, is as follows. In the synthetic dataset, there was only one dependence upon subclasses. Selecting features with subclass dependence automatically gives us targeted features. In the real dataset, this cannot stand as it is. There are many variations of subclass dependence. In TD-based unsupervised FE, by assessing the dependence of singular value vectors upon subclasses (see Fig. 4), we can specify the type of dependence upon subclasses that should be considered. This allows us to identify a restricted number of biologically reliable genomic regions. Categorical regression cannot distinguish between various subclass dependences and can only identify genomic regions associated with any type of subclass dependence. This results in the identification of almost all genomic regions, as most are likely to be associated with a type of subclass dependence.

In spite of the comparisons presented in the above, one might wonder that categorical regression can achieve comparative performances if top ranked restricted number of genomic regions are intentionally selected. In order to this, we selected 1447 top ranked regions (they are as many as those selected by TD based unsupervised FE) and identified as many as 962 gene symbols included in these genomic regions. These 1267 genes are uploaded to Metascape to evaluate them biologically. Figure 15 shows the results of the DisGeNet category of Metascape. In contrast to Fig. 7 where prostatic neoplasms was top ranked, it was even not ranked at all in Figure 15. Thus, it is obvious that genes selected by RF is inferior to those by TD based unsupervised FE. Figure 16 shows the results of the PaGenBase category of Metascape. In cotrast to Fig. 8 where prostate related cell lines are top ranked, no prostate related cell lines are top ranked in Fig. 16. Thus, again, it is obvious that genes selected by RF is inferior to those by TD based unsupervised FE. In contrast to Fig. 9 where AR is top ranked, we could find no significant enrichment in TRRUST category of Metascape. Thus, it is obvious that genes selected by RF is inferior to those by TD based unsupervised FE as well.

**Figure 15.**
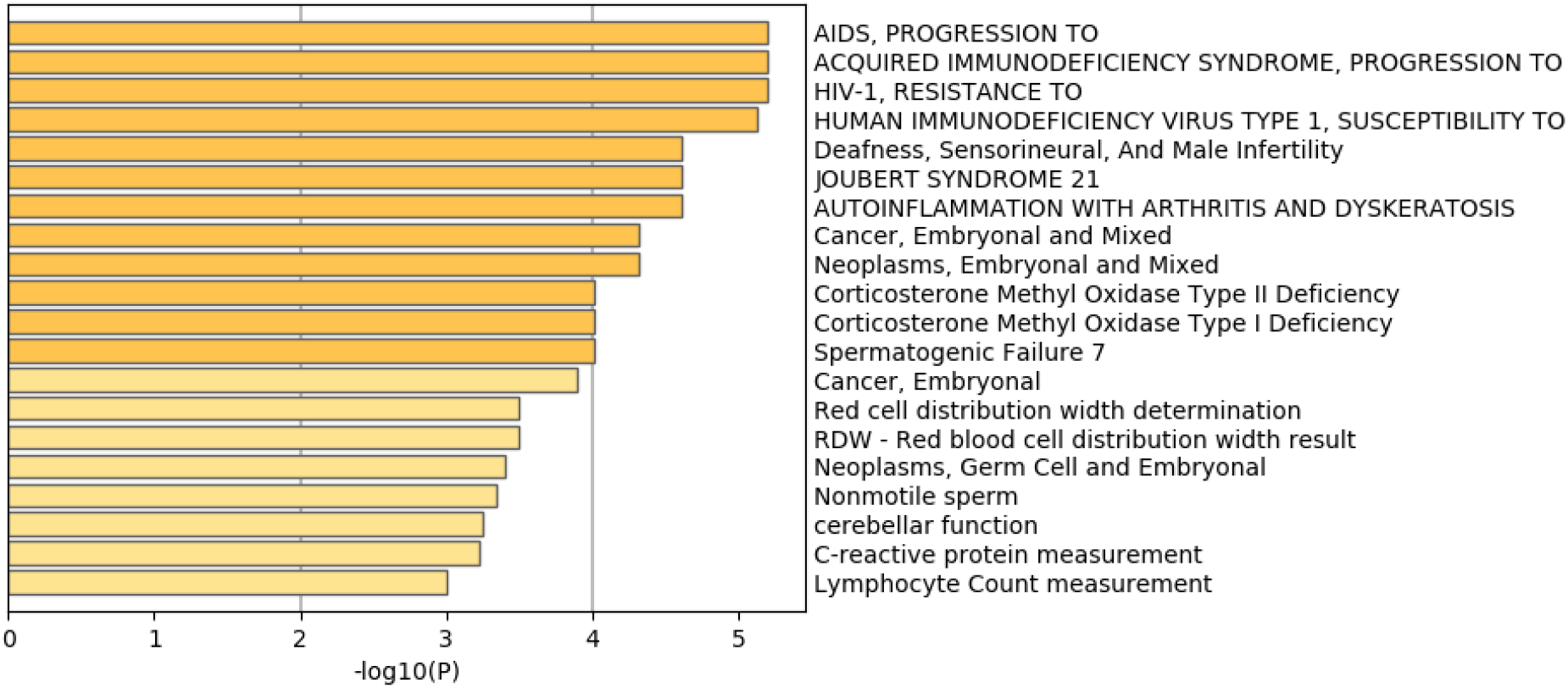
Summary of enrichment of the DisGeNet category of Metascape when 962 protein-coding genes included in 1447 genomic regions selected by categorical regression are considered.

**Figure 16.**
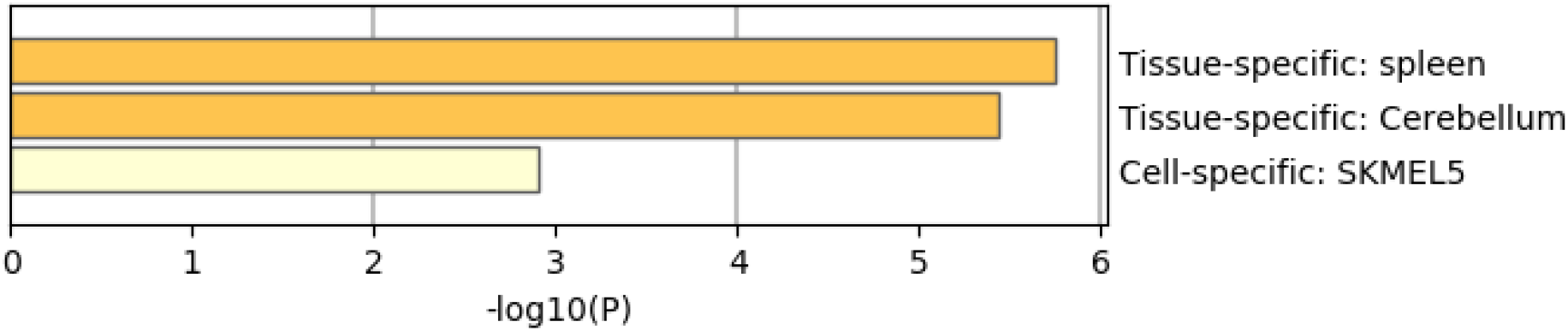
Summary of enrichment of the PaGenBase category of Metascape when 962 protein-coding genes included in 1447 genomic regions selected by categorical regression are considered.

Finally, MNMF (*n* = 3), one of the state-of-the-art methods that are often applied to multiomics datasets [19], and PCA were applied to *x*_*ijkm*_ as described in Materials and Methods. Figs. 17 and 18 show the latent variable vectors and PC loading attributed to 144 samples, respectively. Although the dependence upon eight multiomics measurements are similar to those of TD-based unsupervised FE, because the correspondence with eight types of measurement is clearly inferior to that of TD-based unsupervised FE (within class variances much larger than those of the singular value vectors obtained by TD-based unsupervised FE, Fig. 4), we employ TD-based unsupervised FE. An additional reason not to employ MNMF instead of TD-based unsupervised FE is that, although singular value vectors derived by HOSVD can be assumed to obey a Gaussian distribution, latent variables computed by MNMF are unlikely to because they do not take negative values. Thus, there is no way to attribute *P*-values to genes, as there is no suitable null hypothesis by which we can compute the *P*-values.

**Figure 17.**
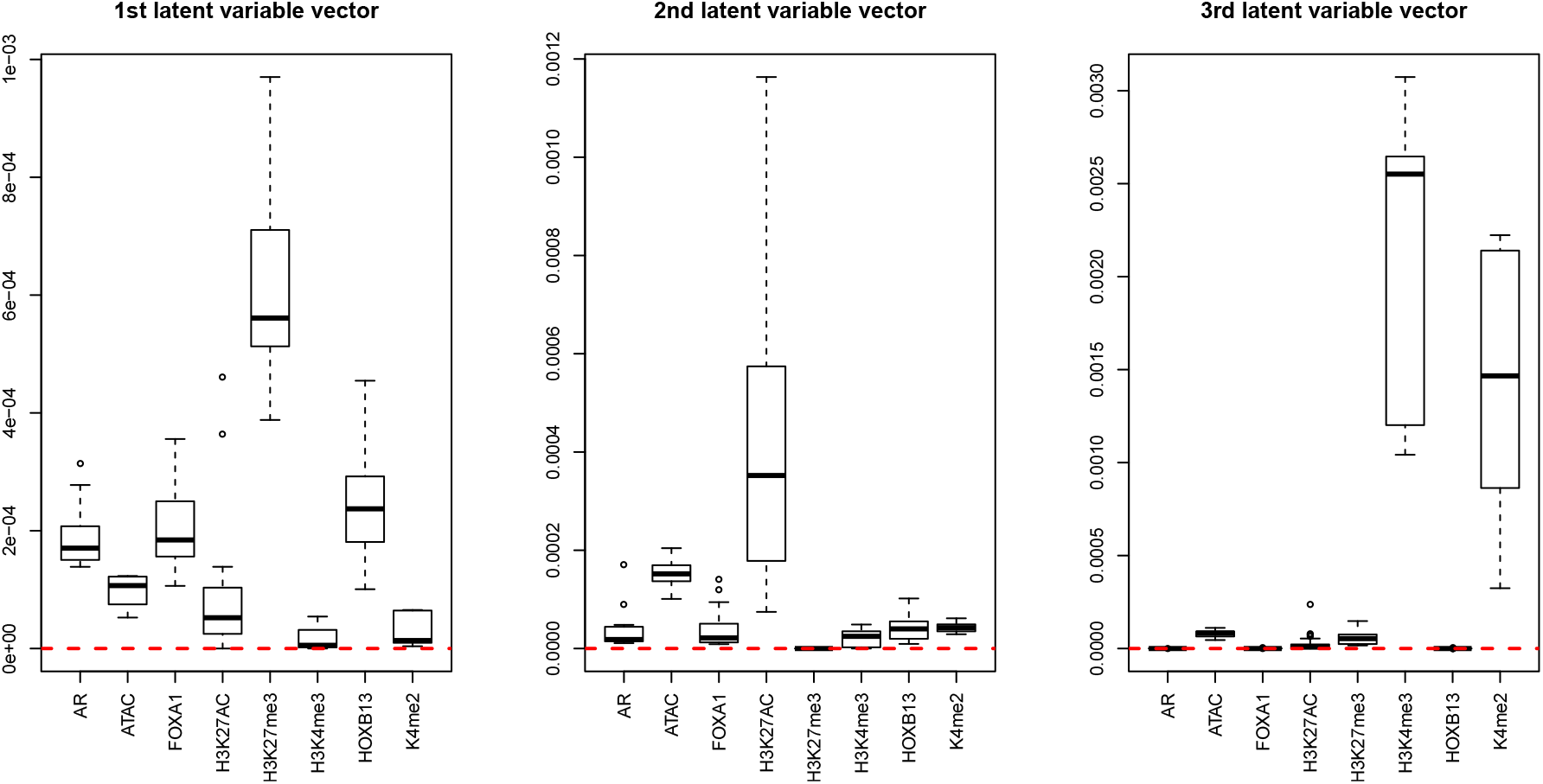
Boxplots of the first to third latent variable vectors attributed to 144 samples classified into eight subclasses, each of which corresponds to a multiomics measurement. The horizontal red broken lines are base lines (zero).

**Figure 18.**
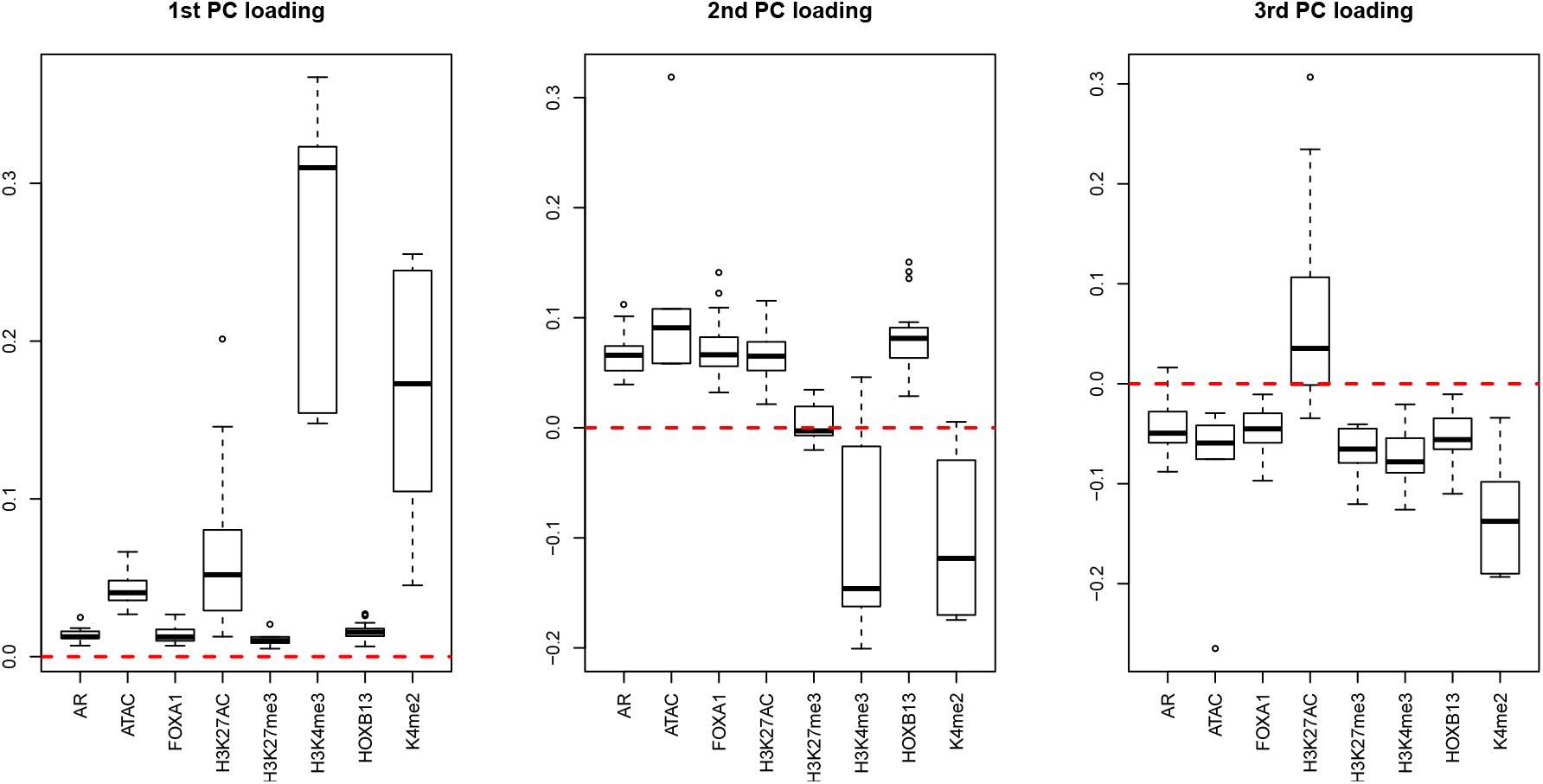
Boxplots of the first to third PC loading attributed to 144 samples classified into eight subclasses, each of which corresponds to a multiomics measurement. The horizontal red broken lines are base lines (zero).

As a result, none of the five methods (RF, PenalizedLDA, categorical regression, MNMF, or PCA) can compete with TD-based unsupervised FE when applied to a real dataset.

In addition, the CPU time required of PCA and TD-based unsupervised FE are also the shortest compared to the other four methods when applied to the real dataset (Table 6). From this perspective, TD-based unsupervised FE outperforms the other four methods.

### Discussions not to specific to either synthetic or real data

Unlike “genomic regions + samples” structure used in traditional unsupervised learning that requires a matrix representation, Our TD employs “genomic regions + samples + tissues + biological replicates” structure that requires a four-axes tensor representation. Reported results via enrichment analysis show the superiority of our TD, attributed to taking into account the relationships among genomic regions, samples, tissues, and biological replicates at once. One might wonder why TD based unsupervised FE is more coincident with subclasses than other supervised or unsupervised methods. It is simple because of the nature of tensor. As can be seen in eqs. (4) and (5), using TD, dependence of *x*_*ijk*_ or *x*_*ijkm*_ upon *i, j, k* or *i, j, k, m* is decomposed. Thus distinction between *i, j, k, m* or *i, j, k* is specifically identified. This means that we can identify distinction of *x*_*ijkm*_ or *x*_*ijk*_ between distinct *i* independent of other index, *j, k* or *j, k, m*. Such a detection is impossible since the distinction between *x*_*ijk*_1 and *x*_*ij*_1 _*k*_ or that between *x*_*ijk*_1 _*m*_ and *x*_*ij*_1 _*km*_ where more than one index are altered simultaneously. This weaken the ability for methods other than TD based unsupervised FE to identify subclass dependence specifically. This is possibly because TD based unsupervised FE can outperform other methods on the identification of subclasses.

## Conclusions

In this study, a TD-based unsupervised FE formalism is successfully applied for the first time to a variety of multiomics measurements for both synthetic and real datasets that represented a typical large *p* small *n* problem. The proposed method outperformed the conventional supervised feature selection methods of RF, categorical regression, PenalizedLDA, and MNMF, and was demonstrated to be superior not only in feature selection but also in selection of biologically reliable genes.

## Supporting information

output from metascape

## Acknowledgment

This study was supported by KAKENHI, 20K12067, 20H04848, and 19H05270. This project was also funded by the Deanship of Scientific Research (DSR) at King Abdulaziz University, Jeddah, under grant no. KEP-8-611-38. The authors, therefore, acknowledge DSR with thanks for providing technical and financial support.

## Supplementary data

Output from Metascape.

