## Supplementary material for "Tensor-decomposition-based unsupervised feature extraction applied to prostate cancer multiomics data": output from metascape: AnalysisReport.html

### Metascape Gene List Analysis Report

metascape.org^1^

#### Bar Graph Summary

Figure 1. Bar graph of enriched terms across input gene lists, colored by p-values.

Metascape only visualizes the top 20 clusters. Up to 100 enriched clusters can be viewed here.

The top-level Gene Ontology biological processes can be viewed here.

#### Gene Lists

User-provided gene identifiers are first converted into their corresponding H. sapiens Entrez gene IDs using the latest version of the database (last updated on 2020-03-19). If multiple identifiers correspond to the same Entrez gene ID, they will be considered as a single Entrez gene ID in downstream analyses. The gene lists are summarized in Table 1.

Table 1. Statistics of input gene lists.

| Name | Total | Unique |
| --- | --- | --- |
| Input ID | 1785 | 1775 |

#### Gene Annotation

The following are the list of annotations retrieved from the latest version of the database (last updated on 2020-03-19) (Table 2).

Table 2. Gene annotations extracted

| Name | Type | Description |
| --- | --- | --- |
| Gene Symbol | Description | Primary HUGO gene symbol. |
| Description | Description | Short description. |
| Biological Process (GO) | Function/Location | Descriptions summarized based on gene ontology database, where up to three most informative GO terms are kept. |
| Kinase Class (UniProt) | Function/Location | Detailed kinase classes. |
| Protein Function (Protein Atlas) | Function/Location | Protein Function (Protein Atlas) |
| Subcellular Location (Protein Atlas) | Function/Location | Sucellular Location (Protein Atlas) |
| Drug (DrugBank) | Genotype/Phenotype/Disease | Drug information for the given gene as target. |
| Canonical Pathways | Ontology | Canonical Pathways |
| Hallmark Gene Sets | Ontology | Hallmark Gene Sets |

#### Pathway and Process Enrichment Analysis

For each given gene list, pathway and process enrichment analysis has been carried out with the following ontology sources: KEGG Pathway, GO Biological Processes, Reactome Gene Sets, Canonical Pathways, CORUM, TRRUST, DisGeNET and PaGenBase. All genes in the genome have been used as the enrichment background. Terms with a p-value < 0.01, a minimum count of 3, and an enrichment factor > 1.5 (the enrichment factor is the ratio between the observed counts and the counts expected by chance) are collected and grouped into clusters based on their membership similarities. More specifically, p-values are calculated based on the accumulative hypergeometric distribution^2^, and q-values are calculated using the Banjamini-Hochberg procedure to account for multiple testings^3^. Kappa scores^4^ are used as the similarity metric when performing hierachical clustering on the enriched terms, and sub-trees with a similarity of > 0.3 are considered a cluster. The most statistically significant term within a cluster is chosen to represent the cluster.

Table 3. Top 20 clusters with their representative enriched terms (one per cluster). "Count" is the number of genes in the user-provided lists with membership in the given ontology term. "%" is the percentage of all of the user-provided genes that are found in the given ontology term (only input genes with at least one ontology term annotation are included in the calculation). "Log10(P)" is the p-value in log base 10. "Log10(q)" is the multi-test adjusted p-value in log base 10.

| GO | Category | Description | Count | % | Log10(P) | Log10(q) |
| --- | --- | --- | --- | --- | --- | --- |
| GO:0043009 | GO Biological Processes | chordate embryonic development | 91 | 5.83 | -11.98 | -7.68 |
| GO:0001568 | GO Biological Processes | blood vessel development | 102 | 6.53 | -11.46 | -7.62 |
| GO:0048732 | GO Biological Processes | gland development | 68 | 4.35 | -10.80 | -7.08 |
| GO:0030855 | GO Biological Processes | epithelial cell differentiation | 100 | 6.40 | -10.45 | -6.84 |
| GO:0002521 | GO Biological Processes | leukocyte differentiation | 75 | 4.80 | -10.38 | -6.84 |
| GO:0045444 | GO Biological Processes | fat cell differentiation | 44 | 2.82 | -10.25 | -6.78 |
| GO:0009611 | GO Biological Processes | response to wounding | 89 | 5.70 | -9.63 | -6.31 |
| GO:0045596 | GO Biological Processes | negative regulation of cell differentiation | 95 | 6.08 | -9.57 | -6.29 |
| GO:0034329 | GO Biological Processes | cell junction assembly | 44 | 2.82 | -9.21 | -6.01 |
| hsa05202 | KEGG Pathway | Transcriptional misregulation in cancer | 36 | 2.30 | -9.06 | -5.92 |
| GO:0061061 | GO Biological Processes | muscle structure development | 86 | 5.51 | -9.06 | -5.92 |
| GO:0045664 | GO Biological Processes | regulation of neuron differentiation | 84 | 5.38 | -8.65 | -5.59 |
| GO:0034248 | GO Biological Processes | regulation of cellular amide metabolic process | 69 | 4.42 | -8.62 | -5.59 |
| GO:0030099 | GO Biological Processes | myeloid cell differentiation | 60 | 3.84 | -8.22 | -5.30 |
| GO:0071383 | GO Biological Processes | cellular response to steroid hormone stimulus | 42 | 2.69 | -8.10 | -5.21 |
| GO:0008285 | GO Biological Processes | negative regulation of cell proliferation | 91 | 5.83 | -7.86 | -5.02 |
| GO:0001503 | GO Biological Processes | ossification | 57 | 3.65 | -7.60 | -4.80 |
| GO:0061448 | GO Biological Processes | connective tissue development | 44 | 2.82 | -7.59 | -4.80 |
| GO:0030856 | GO Biological Processes | regulation of epithelial cell differentiation | 31 | 1.98 | -7.56 | -4.79 |
| R-HSA-9006931 | Reactome Gene Sets | Signaling by Nuclear Receptors | 46 | 2.94 | -7.52 | -4.76 |

To further capture the relationships between the terms, a subset of enriched terms have been selected and rendered as a network plot, where terms with a similarity > 0.3 are connected by edges. We select the terms with the best p-values from each of the 20 clusters, with the constraint that there are no more than 15 terms per cluster and no more than 250 terms in total. The network is visualized using Cytoscape^5^, where each node represents an enriched term and is colored first by its cluster ID (Figure 2.a) and then by its p-value (Figure 2.b). These networks can be interactively viewed in Cytoscape through the .cys files (contained in the Zip package, which also contains a publication-quality version as a PDF) or within a browser by clicking on the web icon. For clarity, term labels are only shown for one term per cluster, so it is recommended to use Cytoscape or a browser to visualize the network in order to inspect all node labels. We can also export the network into a PDF file within Cytoscape, and then edit the labels using Adobe Illustrator for publication purposes. To switch off all labels, delete the "Label" mapping under the "Style" tab within Cytoscape, and then export the network view.

Figure 2. Network of enriched terms: (a) colored by cluster ID, where nodes that share the same cluster ID are typically close to each other; (b) colored by p-value, where terms containing more genes tend to have a more significant p-value.

#### Protein-protein Interaction Enrichment Analysis

For each given gene list, protein-protein interaction enrichment analysis has been carried out with the following databases: BioGrid^6^, InWeb\_IM^7^, OmniPath^8^. The resultant network contains the subset of proteins that form physical interactions with at least one other member in the list. If the network contains between 3 and 500 proteins, the Molecular Complex Detection (MCODE) algorithm^9^ has been applied to identify densely connected network components.

#### Quality Control and Association Analysis

Gene list enrichments are identified in the following ontology categories: TRRUST, DisGeNET, PaGenBase. All genes in the genome have been used as the enrichment background. Terms with a p-value < 0.01, a minimum count of 3, and an enrichment factor > 1.5 (the enrichment factor is the ratio between the observed counts and the counts expected by chance) are collected and grouped into clusters based on their membership similarities. The top few enriched clusters (one term per cluster) are shown in the Figure 3-5. The algorithm used here is the same as that is used for pathway and process enrichment analysis.

Figure 3. Summary of enrichment analysis in TRRUST.

| GO | Description | Count | % | Log10(P) | Log10(q) |
| --- | --- | --- | --- | --- | --- |
| TRR00011 | Regulated by: AR | 26 | 1.70 | -10.00 | -6.30 |
| TRR00714 | Regulated by: TP53 | 29 | 1.90 | -6.30 | -3.00 |
| TRR00140 | Regulated by: ELK1 | 8 | 0.51 | -4.80 | -1.90 |
| TRR00745 | Regulated by: VHL | 8 | 0.51 | -4.80 | -1.90 |
| TRR00662 | Regulated by: STAT3 | 22 | 1.40 | -4.00 | -1.50 |
| TRR00466 | Regulated by: NKX3-1 | 5 | 0.32 | -3.70 | -1.30 |
| TRR00452 | Regulated by: NFKB1 | 36 | 2.30 | -3.60 | -1.30 |
| TRR00082 | Regulated by: CREBBP | 7 | 0.45 | -3.10 | -0.99 |
| TRR00610 | Regulated by: SIRT1 | 10 | 0.64 | -3.10 | -0.98 |
| TRR00575 | Regulated by: RELA | 34 | 2.20 | -3.10 | -0.95 |
| TRR00665 | Regulated by: STAT5A | 5 | 0.32 | -3.00 | -0.95 |
| TRR00572 | Regulated by: RBMX | 4 | 0.26 | -3.00 | -0.95 |
| TRR00459 | Regulated by: NFYB | 5 | 0.32 | -2.90 | -0.86 |
| TRR00641 | Regulated by: SP1 | 47 | 3.00 | -2.80 | -0.79 |
| TRR00283 | Regulated by: HOXB7 | 4 | 0.26 | -2.60 | -0.69 |
| TRR00152 | Regulated by: ESR1 | 12 | 0.77 | -2.50 | -0.62 |
| TRR00655 | Regulated by: SRF | 6 | 0.38 | -2.40 | -0.52 |
| TRR00016 | Regulated by: ARNTL | 3 | 0.19 | -2.30 | -0.52 |
| TRR00478 | Regulated by: NR1I2 | 6 | 0.38 | -2.30 | -0.48 |
| TRR00653 | Regulated by: SREBF1 | 6 | 0.38 | -2.30 | -0.48 |

Figure 4. Summary of enrichment analysis in DisGeNET^10^.

| GO | Description | Count | % | Log10(P) | Log10(q) |
| --- | --- | --- | --- | --- | --- |
| C0033578 | Prostatic Neoplasms | 87 | 5.60 | -17.00 | -13.00 |
| C0023467 | Leukemia, Myelocytic, Acute | 26 | 1.70 | -7.30 | -3.90 |
| C4280567 | Abnormal skeletal development | 17 | 1.10 | -6.90 | -3.60 |
| C0376634 | Craniofacial Abnormalities | 27 | 1.70 | -6.00 | -2.80 |
| C0151744 | Myocardial Ischemia | 29 | 1.90 | -5.90 | -2.70 |
| C0029422 | Osteochondrodysplasias | 17 | 1.10 | -5.70 | -2.50 |
| C0026827 | Muscle hypotonia | 66 | 4.20 | -5.50 | -2.40 |
| C0700208 | Acquired scoliosis | 38 | 2.40 | -4.80 | -1.90 |
| C0451695 | Major histocompatibility complex class I deficiency | 4 | 0.26 | -4.80 | -1.90 |
| C1858266 | Bare Lymphocyte Syndrome, Type I | 4 | 0.26 | -4.80 | -1.90 |
| C0025990 | Micrognathism | 37 | 2.40 | -4.80 | -1.90 |
| C0240295 | Mandibular hypoplasia | 37 | 2.40 | -4.80 | -1.90 |
| C1857130 | Hypoplastic mandible condyle | 37 | 2.40 | -4.80 | -1.90 |
| C0037932 | Curvature of spine | 38 | 2.40 | -4.70 | -1.90 |
| C0010038 | Corneal Opacity | 13 | 0.83 | -4.70 | -1.90 |
| C1839764 | Broad flat nasal bridge | 33 | 2.10 | -4.70 | -1.90 |
| C1849367 | Nasal bridge wide | 33 | 2.10 | -4.70 | -1.90 |
| C0026034 | Microstomia | 16 | 1.00 | -4.50 | -1.80 |
| C0024667 | Animal Mammary Neoplasms | 23 | 1.50 | -4.40 | -1.70 |
| C0027626 | Neoplasm Invasiveness | 23 | 1.50 | -4.40 | -1.70 |

Figure 5. Summary of enrichment analysis in PaGenBase^11^.

| GO | Description | Count | % | Log10(P) | Log10(q) |
| --- | --- | --- | --- | --- | --- |
| PGB:00091 | Cell-specific: LNCAP | 16 | 1.00 | -10.00 | -6.30 |
| PGB:00010 | Tissue-specific: prostate | 36 | 2.30 | -7.40 | -3.90 |
| PGB:00076 | Cell-specific: Bronchial Epithelial Cells | 26 | 1.70 | -7.10 | -3.70 |
