## Supplementary material for "Tensor-decomposition-based unsupervised feature extraction applied to prostate cancer multiomics data": output from metascape: AnalysisReport.pptx

### Slide 1
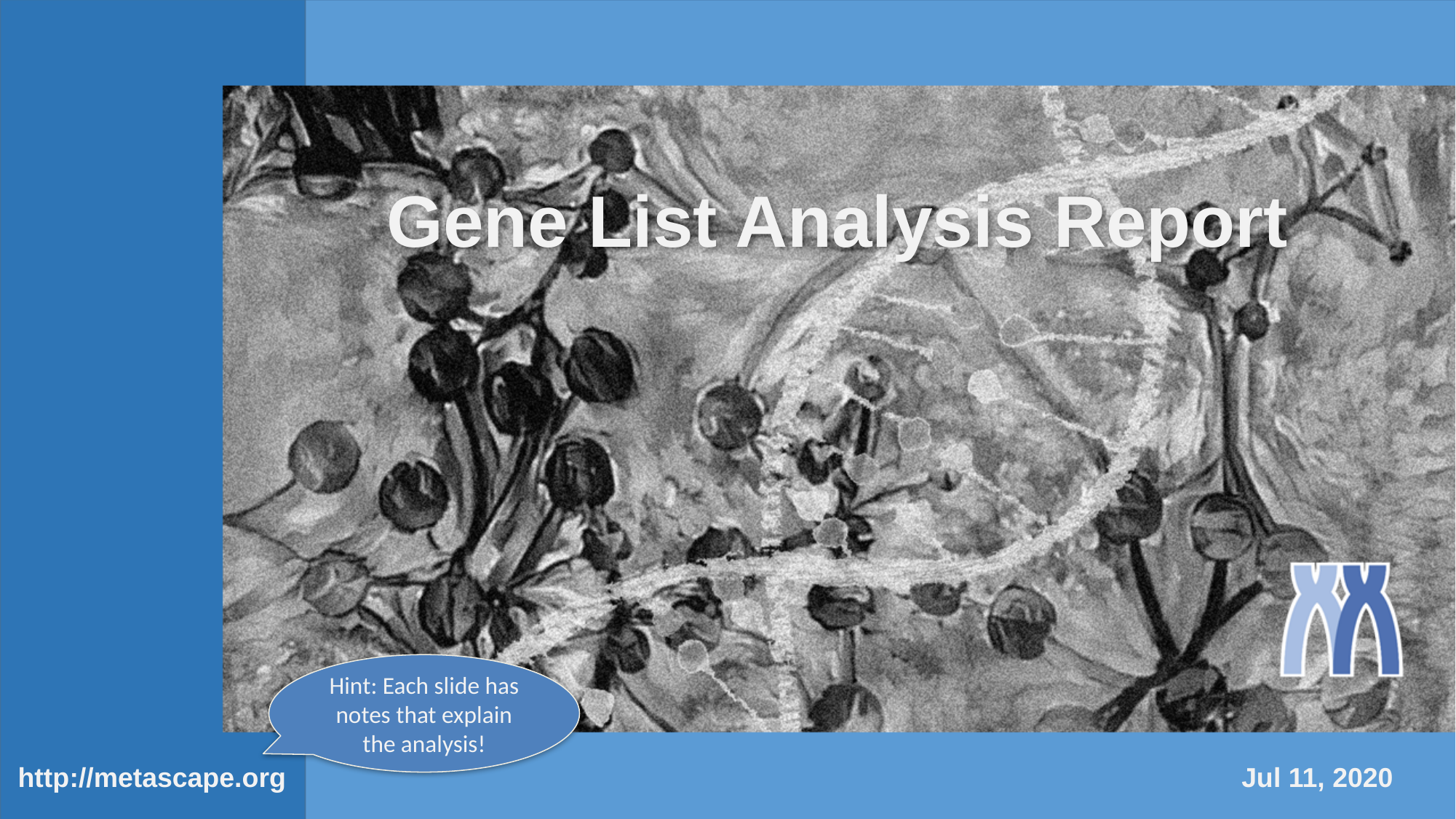

#### Gene List Analysis Report
Hint: Each slide has notes that explain the analysis!
http://metascape.org
Jul 11, 2020

### Slide 2
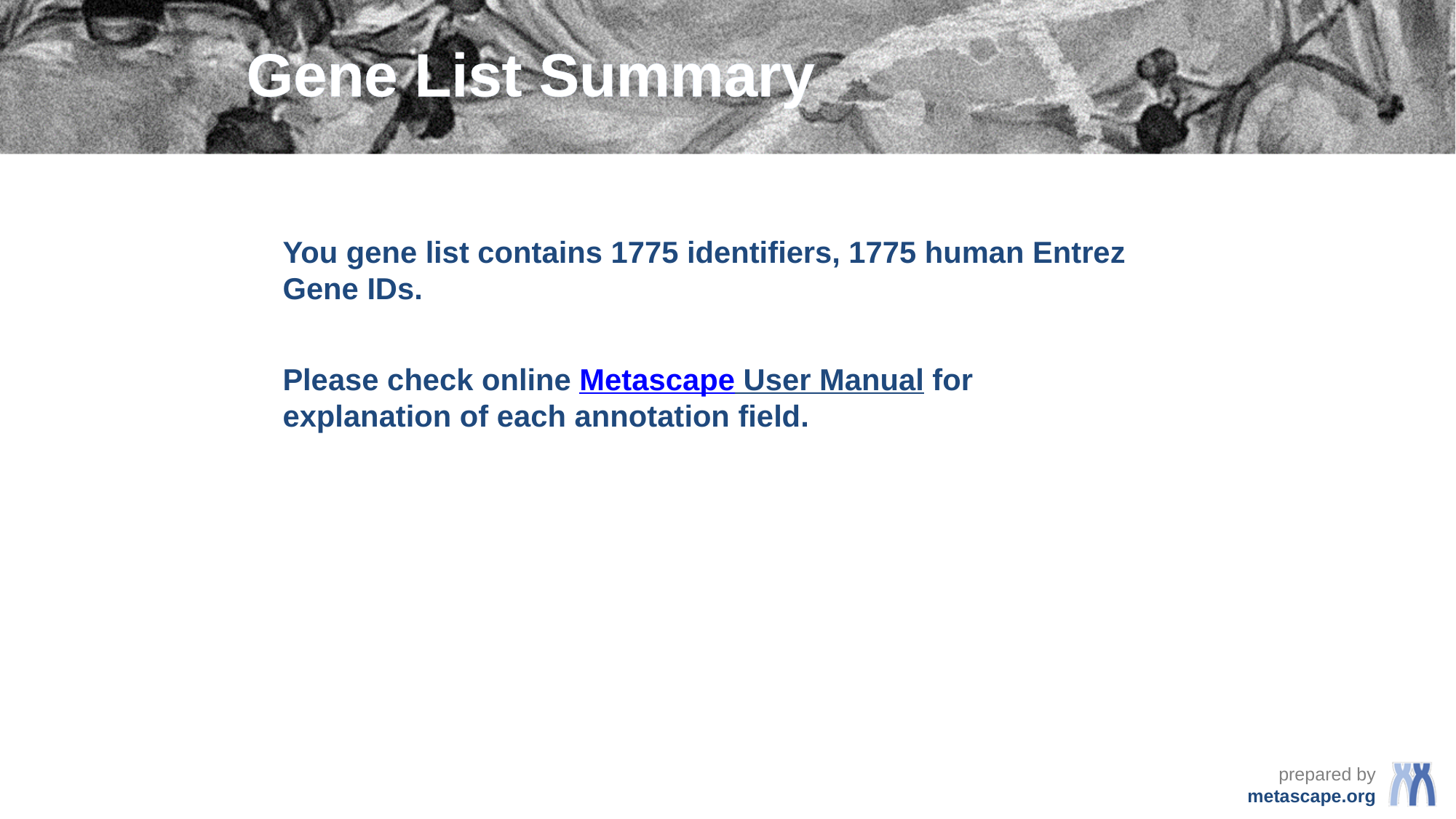

#### Gene List Summary
You gene list contains 1775 identifiers, 1775 human Entrez Gene IDs.
Please check online Metascape User Manual for explanation of each annotation field.

### Slide 3
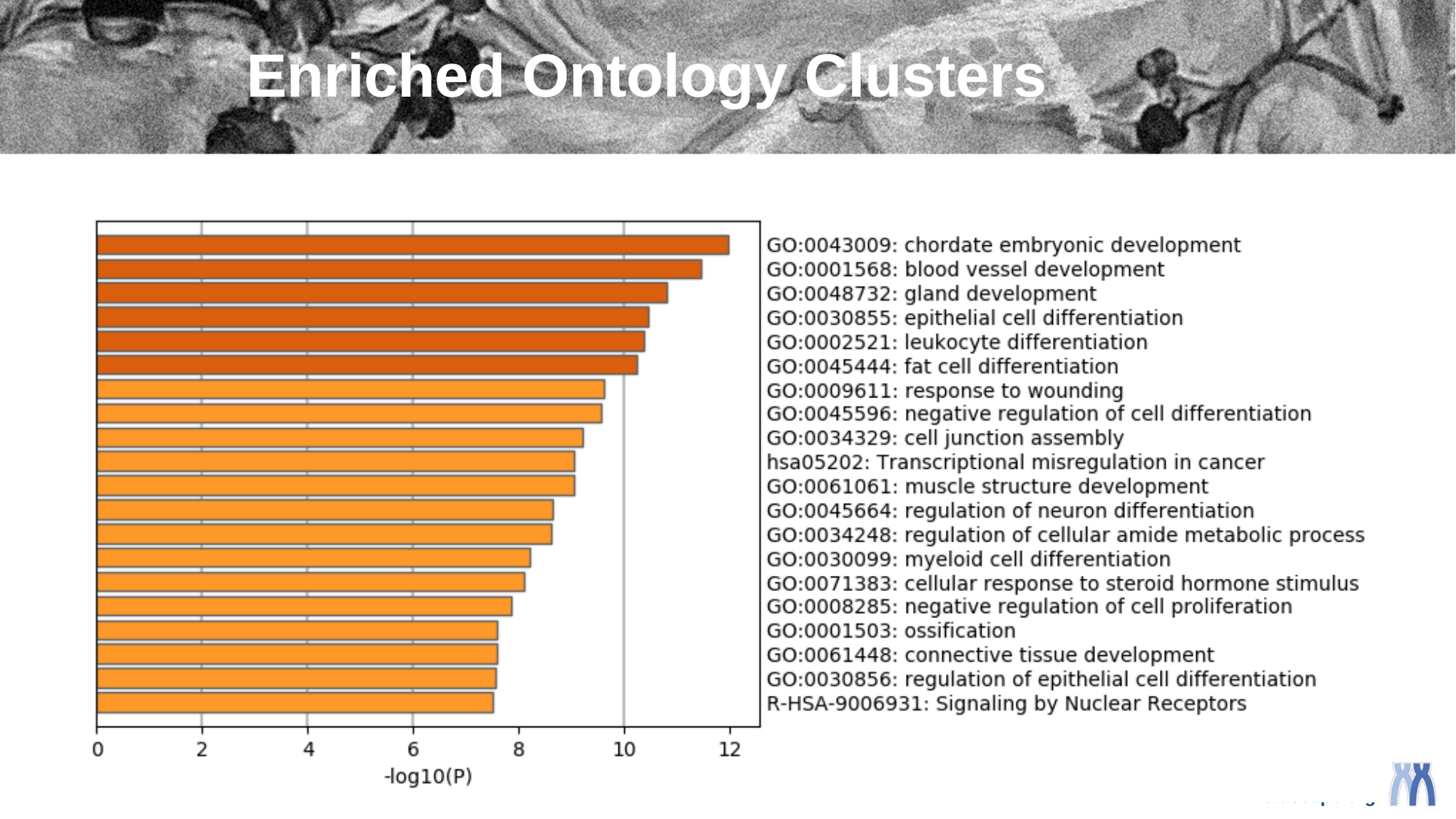

#### Enriched Ontology Clusters

### Slide 4
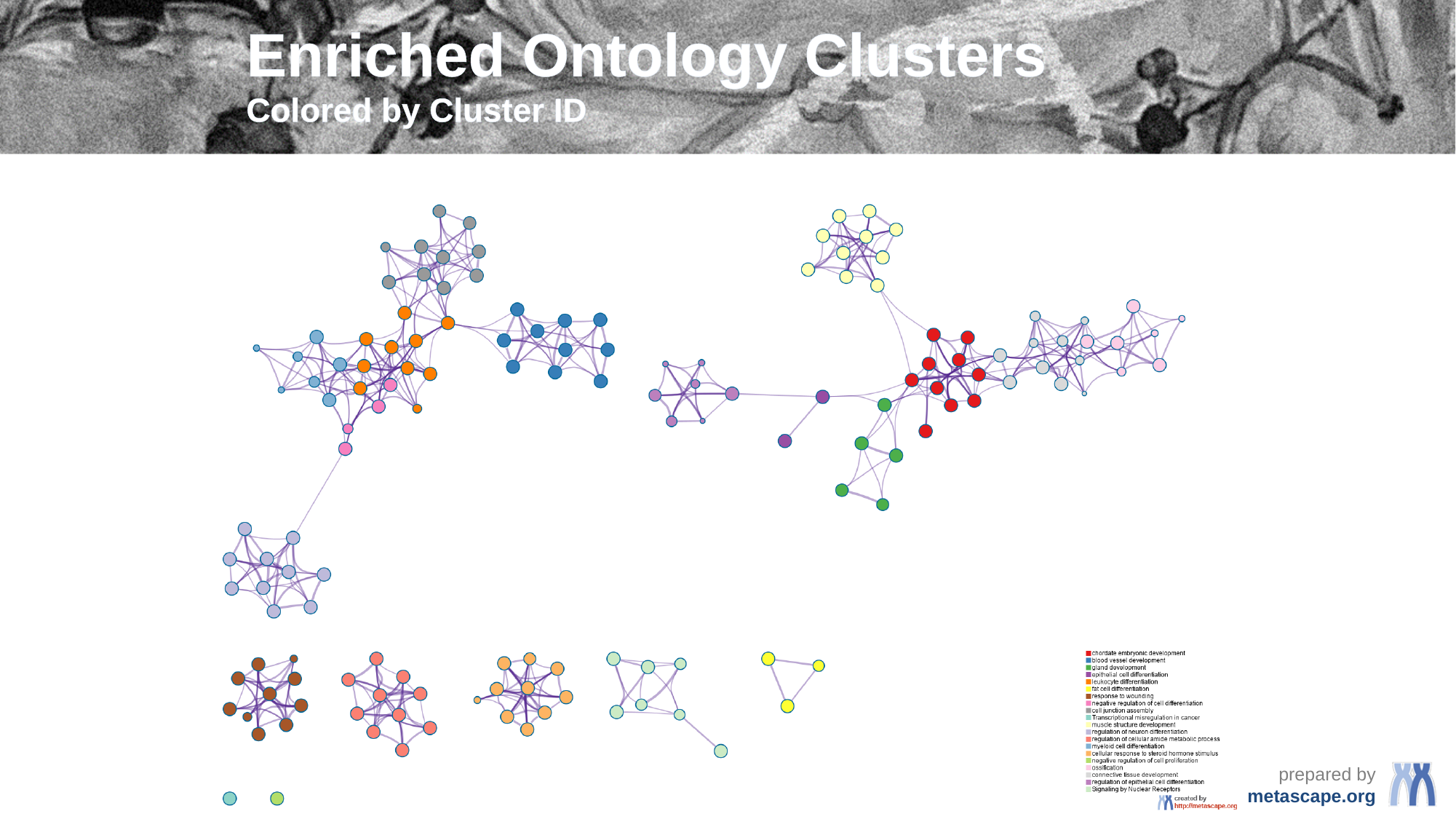

#### Enriched Ontology Clusters Colored by Cluster ID

### Slide 5
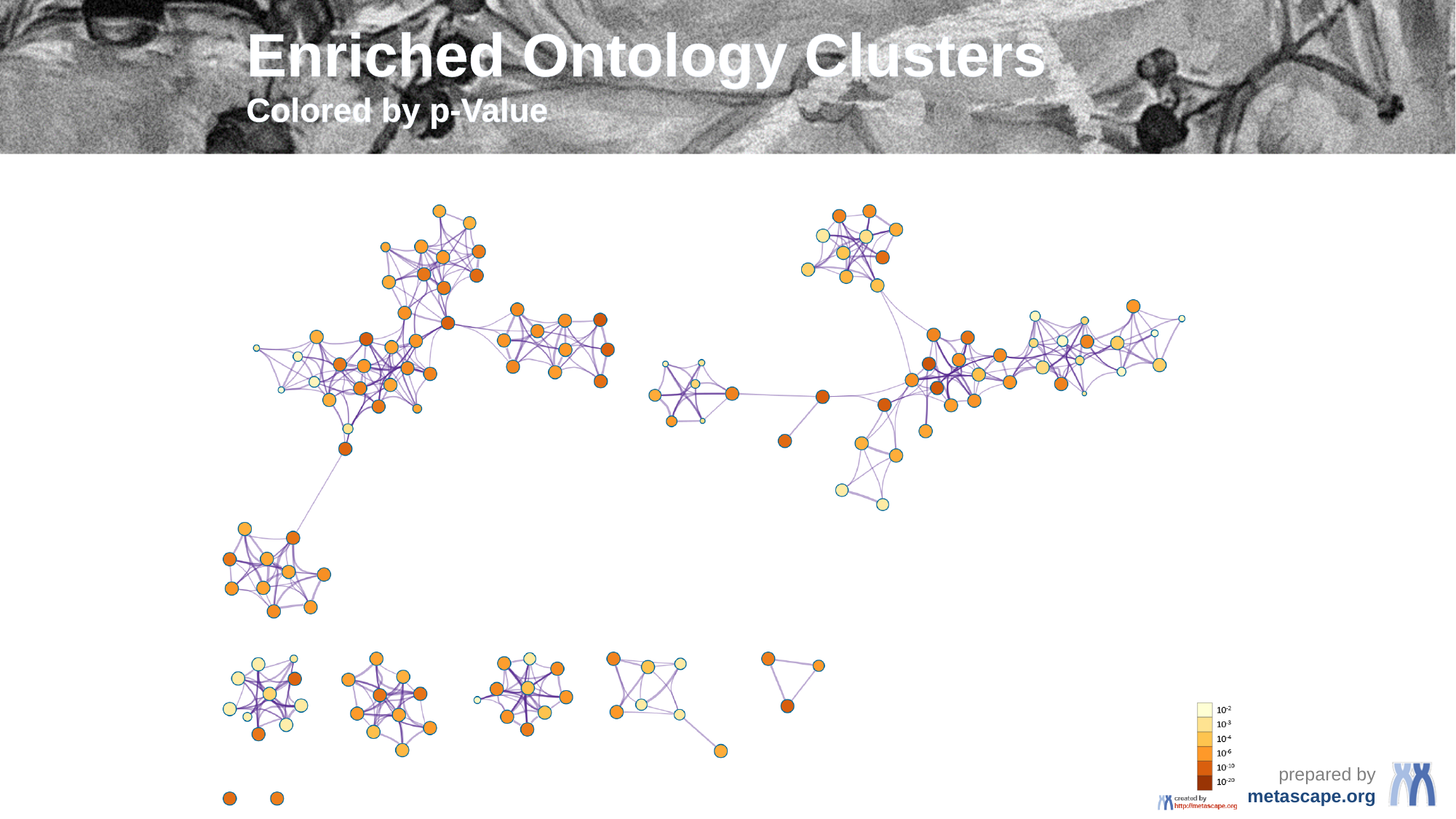

#### Enriched Ontology Clusters Colored by p-Value

### Slide 6
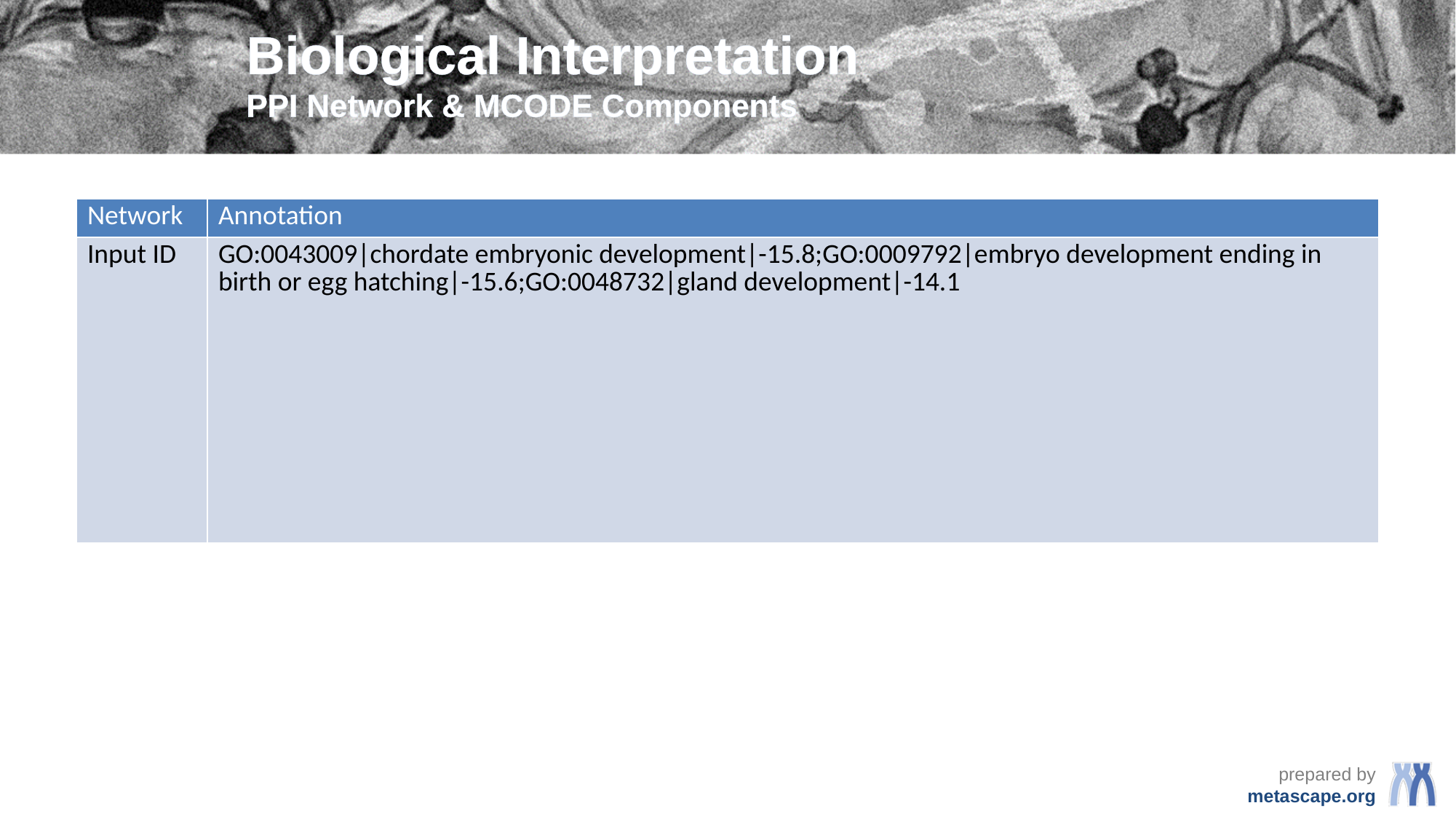

#### Biological Interpretation PPI Network & MCODE Components
| Network | Annotation |
| --- | --- |
| Input ID | GO:0043009|chordate embryonic development|-15.8;GO:0009792|embryo development ending in birth or egg hatching|-15.6;GO:0048732|gland development|-14.1 |

### Slide 7
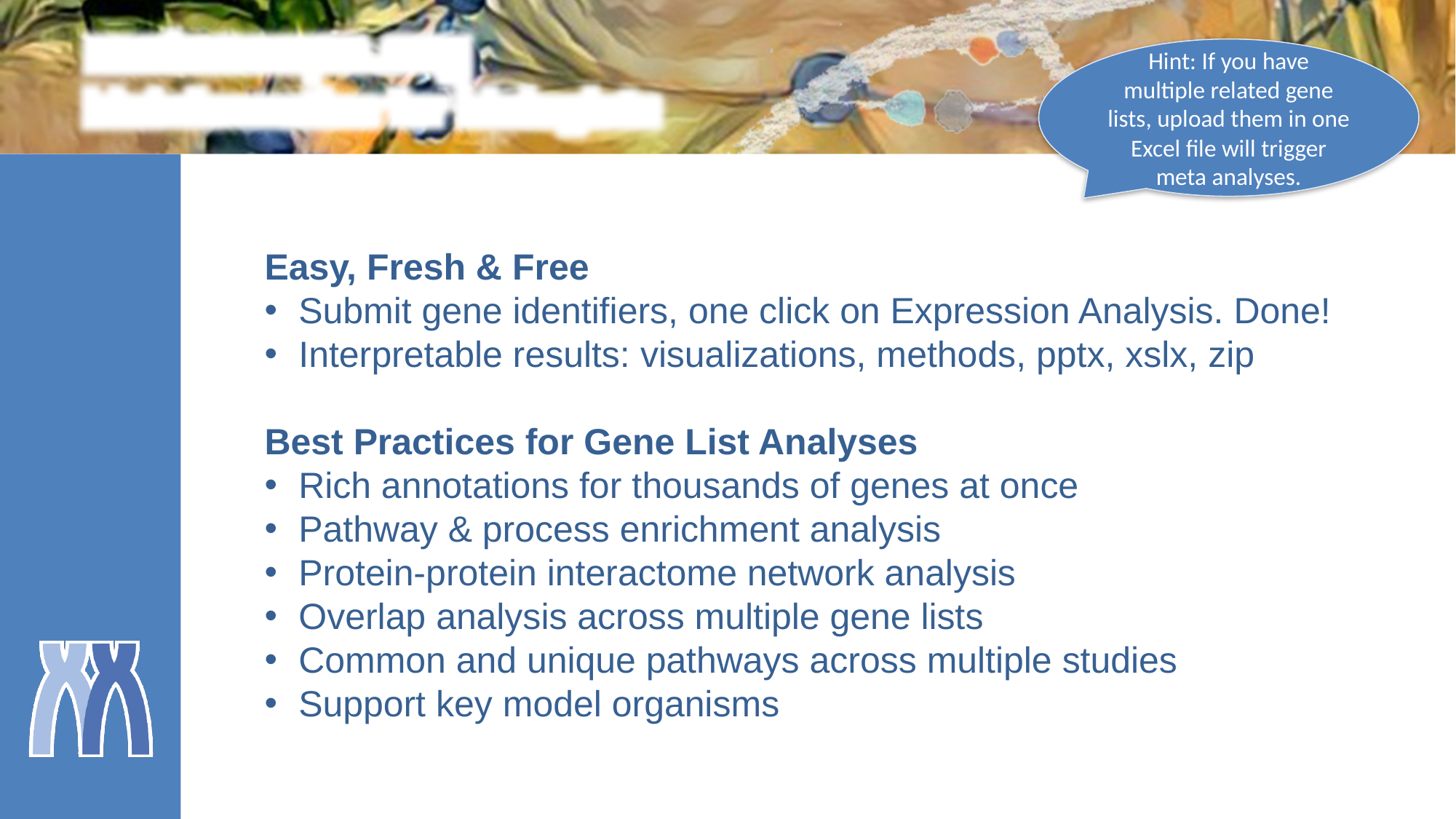

metascape.org
bioinformatics for biologists
Hint: If you have multiple related gene lists, upload them in one Excel file will trigger meta analyses.
Easy, Fresh & Free
Submit gene identifiers, one click on Expression Analysis. Done!
Interpretable results: visualizations, methods, pptx, xslx, zip
Best Practices for Gene List Analyses
Rich annotations for thousands of genes at once
Pathway & process enrichment analysis
Protein-protein interactome network analysis
Overlap analysis across multiple gene lists
Common and unique pathways across multiple studies
Support key model organisms
