## Supplementary material for "Tensor-decomposition-based unsupervised feature extraction applied to prostate cancer multiomics data": output from metascape: 65d77273.vendor.js

```
if(function(t,e){"object"==typeof module&&"object"==typeof module.exports?module.exports=t.document?e(t,!0):function(t){if(!t.document)throw new Error("jQuery requires a window with a document");return e(t)}:e(t)}("undefined"!=typeof window?window:this,function(f,t){function s(t){var e="length"in t&&t.length,n=U.type(t);return"function"!==n&&!U.isWindow(t)&&(!(1!==t.nodeType||!e)||("array"===n||0===e||"number"==typeof e&&0<e&&e-1 in t))}function e(t,n,i){if(U.isFunction(n))return U.grep(t,function(t,e){return!!n.call(t,e,t)!==i});if(n.nodeType)return U.grep(t,function(t){return t===n!==i});if("string"==typeof n){if(tt.test(n))return U.filter(n,t,i);n=U.filter(n,t)}return U.grep(t,function(t){return 0<=V.call(n,t)!==i})}function n(t,e){for(;(t=t[e])&&1!==t.nodeType;);return t}function i(){Y.removeEventListener("DOMContentLoaded",i,!1),f.removeEventListener("load",i,!1),U.ready()}function r(){Object.defineProperty(this.cache={},0,{get:function(){return{}}}),this.expando=U.expando+r.uid++}function l(t,e,n){var i;if(void 0===n&&1===t.nodeType)if(i="data-"+e.replace(dt,"-$1").toLowerCase(),"string"==typeof(n=t.getAttribute(i))){try{n="true"===n||"false"!==n&&("null"===n?null:+n+""===n?+n:ht.test(n)?U.parseJSON(n):n)}catch(t){}ct.set(t,e,n)}else n=void 0;return n}function o(){return!0}function u(){return!1}function a(){try{return Y.activeElement}catch(t){}}function c(t,e){return U.nodeName(t,"table")&&U.nodeName(11!==e.nodeType?e:e.firstChild,"tr")?t.getElementsByTagName("tbody")[0]||t.appendChild(t.ownerDocument.createElement("tbody")):t}function g(t){return t.type=(null!==t.getAttribute("type"))+"/"+t.type,t}function v(t){var e=Pt.exec(t.type);return e?t.type=e[1]:t.removeAttribute("type"),t}function m(t,e){for(var n=0,i=t.length;n<i;n++)ut.set(t[n],"globalEval",!e||ut.get(e[n],"globalEval"))}function d(t,e){var n,i,r,o,a,s,l,u;if(1===e.nodeType){if(ut.hasData(t)&&(o=ut.access(t),a=ut.set(e,o),u=o.events))for(r in delete a.handle,a.events={},u)for(n=0,i=u[r].length;n<i;n++)U.event.add(e,r,u[r][n]);ct.hasData(t)&&(s=ct.access(t),l=U.extend({},s),ct.set(e,l))}}function y(t,e){var n=t.getElementsByTagName?t.getElementsByTagName(e||"*"):t.querySelectorAll?t.querySelectorAll(e||"*"):[];return void 0===e||e&&U.nodeName(t,e)?U.merge([t],n):n}function h(t,e){var n,i=U(e.createElement(t)).appendTo(e.body),r=f.getDefaultComputedStyle&&(n=f.getDefaultComputedStyle(i[0]))?n.display:U.css(i[0],"display");return i.detach(),r}function b(t){var e=Y,n=Rt[t];return n||("none"!==(n=h(t,e))&&n||((e=(Nt=(Nt||U("<iframe frameborder='0' width='0' height='0'/>")).appendTo(e.documentElement))[0].contentDocument).write(),e.close(),n=h(t,e),Nt.detach()),Rt[t]=n),n}function p(t,e,n){var i,r,o,a,s=t.style;return(n=n||qt(t))&&(a=n.getPropertyValue(e)||n[e]),n&&(""!==a||U.contains(t.ownerDocument,t)||(a=U.style(t,e)),Vt.test(a)&&Ft.test(e)&&(i=s.width,r=s.minWidth,o=s.maxWidth,s.minWidth=s.maxWidth=s.width=a,a=n.width,s.width=i,s.minWidth=r,s.maxWidth=o)),void 0!==a?a+"":a}function x(t,e){return{get:function(){return t()?void delete this.get:(this.get=e).apply(this,arguments)}}}function w(t,e){if(e in t)return e;for(var n=e[0].toUpperCase()+e.slice(1),i=e,r=Gt.length;r--;)if((e=Gt[r]+n)in t)return e;return i}function $(t,e,n){var i=Xt.exec(e);return i?Math.max(0,i[1]-(n||0))+(i[2]||"px"):e}function k(t,e,n,i,r){for(var o=n===(i?"border":"content")?4:"width"===e?1:0,a=0;o<4;o+=2)"margin"===n&&(a+=U.css(t,n+mt[o],!0,r)),i?("content"===n&&(a-=U.css(t,"padding"+mt[o],!0,r)),"margin"!==n&&(a-=U.css(t,"border"+mt[o]+"Width",!0,r))):(a+=U.css(t,"padding"+mt[o],!0,r),"padding"!==n&&(a+=U.css(t,"border"+mt[o]+"Width",!0,r)));return a}function C(t,e,n){var i=!0,r="width"===e?t.offsetWidth:t.offsetHeight,o=qt(t),a="border-box"===U.css(t,"boxSizing",!1,o);if(r<=0||null==r){if(((r=p(t,e,o))<0||null==r)&&(r=t.style[e]),Vt.test(r))return r;i=a&&(X.boxSizingReliable()||r===t.style[e]),r=parseFloat(r)||0}return r+k(t,e,n||(a?"border":"content"),i,o)+"px"}function E(t,e){for(var n,i,r,o=[],a=0,s=t.length;a<s;a++)(i=t[a]).style&&(o[a]=ut.get(i,"olddisplay"),n=i.style.display,e?(o[a]||"none"!==n||(i.style.display=""),""===i.style.display&&pt(i)&&(o[a]=ut.access(i,"olddisplay",b(i.nodeName)))):(r=pt(i),"none"===n&&r||ut.set(i,"olddisplay",r?n:U.css(i,"display"))));for(a=0;a<s;a++)(i=t[a]).style&&(e&&"none"!==i.style.display&&""!==i.style.display||(i.style.display=e?o[a]||"":"none"));return t}function S(t,e,n,i,r){return new S.prototype.init(t,e,n,i,r)}function T(){return setTimeout(function(){Zt=void 0}),Zt=U.now()}function D(t,e){var n,i=0,r={height:t};for(e=e?1:0;i<4;i+=2-e)r["margin"+(n=mt[i])]=r["padding"+n]=t;return e&&(r.opacity=r.width=t),r}function _(t,e,n){for(var i,r=(oe[e]||[]).concat(oe["*"]),o=0,a=r.length;o<a;o++)if(i=r[o].call(n,e,t))return i}function P(o,t,e){var n,a,i=0,r=re.length,s=U.Deferred().always(function(){delete l.elem}),l=function(){if(a)return!1;for(var t=Zt||T(),e=Math.max(0,u.startTime+u.duration-t),n=1-(e/u.duration||0),i=0,r=u.tweens.length;i<r;i++)u.tweens[i].run(n);return s.notifyWith(o,[u,n,e]),n<1&&r?e:(s.resolveWith(o,[u]),!1)},u=s.promise({elem:o,props:U.extend({},t),opts:U.extend(!0,{specialEasing:{}},e),originalProperties:t,originalOptions:e,startTime:Zt||T(),duration:e.duration,tweens:[],createTween:function(t,e){var n=U.Tween(o,u.opts,t,e,u.opts.specialEasing[t]||u.opts.easing);return u.tweens.push(n),n},stop:function(t){var e=0,n=t?u.tweens.length:0;if(a)return this;for(a=!0;e<n;e++)u.tweens[e].run(1);return t?s.resolveWith(o,[u,t]):s.rejectWith(o,[u,t]),this}}),c=u.props;for(function(t,e){var n,i,r,o,a;for(n in t)if(r=e[i=U.camelCase(n)],o=t[n],U.isArray(o)&&(r=o[1],o=t[n]=o[0]),n!==i&&(t[i]=o,delete t[n]),(a=U.cssHooks[i])&&"expand"in a)for(n in o=a.expand(o),delete t[i],o)n in t||(t[n]=o[n],e[n]=r);else e[i]=r}(c,u.opts.specialEasing);i<r;i++)if(n=re[i].call(u,o,c,u.opts))return n;return U.map(c,_,u),U.isFunction(u.opts.start)&&u.opts.start.call(o,u),U.fx.timer(U.extend(l,{elem:o,anim:u,queue:u.opts.queue})),u.progress(u.opts.progress).done(u.opts.done,u.opts.complete).fail(u.opts.fail).always(u.opts.always)}function M(o){return function(t,e){"string"!=typeof t&&(e=t,t="*");var n,i=0,r=t.toLowerCase().match(at)||[];if(U.isFunction(e))for(;n=r[i++];)"+"===n[0]?(n=n.slice(1)||"*",(o[n]=o[n]||[]).unshift(e)):(o[n]=o[n]||[]).push(e)}}function A(e,r,o,a){function s(t){var i;return l[t]=!0,U.each(e[t]||[],function(t,e){var n=e(r,o,a);return"string"!=typeof n||u||l[n]?u?!(i=n):void 0:(r.dataTypes.unshift(n),s(n),!1)}),i}var l={},u=e===xe;return s(r.dataTypes[0])||!l["*"]&&s("*")}function N(t,e){var n,i,r=U.ajaxSettings.flatOptions||{};for(n in e)void 0!==e[n]&&((r[n]?t:i||(i={}))[n]=e[n]);return i&&U.extend(!0,t,i),t}function O(n,t,i,r){var e;if(U.isArray(t))U.each(t,function(t,e){i||Ee.test(n)?r(n,e):O(n+"["+("object"==typeof e?t:"")+"]",e,i,r)});else if(i||"object"!==U.type(t))r(n,t);else for(e in t)O(n+"["+e+"]",t[e],i,r)}function I(t){return U.isWindow(t)?t:9===t.nodeType&&t.defaultView}function B(t,e){return e.toUpperCase()}var L=[],j=L.slice,R=L.concat,F=L.push,V=L.indexOf,q={},z=q.toString,H=q.hasOwnProperty,X={},Y=f.document,U=function(t,e){return new U.fn.init(t,e)},W=/^[\s\uFEFF\xA0]+|[\s\uFEFF\xA0]+$/g,G=/^-ms-/,Z=/-([\da-z])/gi;U.fn=U.prototype={jquery:"2.1.4",constructor:U,selector:"",length:0,toArray:function(){return j.call(this)},get:function(t){return null!=t?t<0?this[t+this.length]:this[t]:j.call(this)},pushStack:function(t){var e=U.merge(this.constructor(),t);return e.prevObject=this,e.context=this.context,e},each:function(t,e){return U.each(this,t,e)},map:function(n){return this.pushStack(U.map(this,function(t,e){return n.call(t,e,t)}))},slice:function(){return this.pushStack(j.apply(this,arguments))},first:function(){return this.eq(0)},last:function(){return this.eq(-1)},eq:function(t){var e=this.length,n=+t+(t<0?e:0);return this.pushStack(0<=n&&n<e?[this[n]]:[])},end:function(){return this.prevObject||this.constructor(null)},push:F,sort:L.sort,splice:L.splice},U.extend=U.fn.extend=function(){var t,e,n,i,r,o,a=arguments[0]||{},s=1,l=arguments.length,u=!1;for("boolean"==typeof a&&(u=a,a=arguments[s]||{},s++),"object"==typeof a||U.isFunction(a)||(a={}),s===l&&(a=this,s--);s<l;s++)if(null!=(t=arguments[s]))for(e in t)n=a[e],a!==(i=t[e])&&(u&&i&&(U.isPlainObject(i)||(r=U.isArray(i)))?(o=r?(r=!1,n&&U.isArray(n)?n:[]):n&&U.isPlainObject(n)?n:{},a[e]=U.extend(u,o,i)):void 0!==i&&(a[e]=i));return a},U.extend({expando:"jQuery"+("2.1.4"+Math.random()).replace(/\D/g,""),isReady:!0,error:function(t){throw new Error(t)},noop:function(){},isFunction:function(t){return"function"===U.type(t)},isArray:Array.isArray,isWindow:function(t){return null!=t&&t===t.window},isNumeric:function(t){return!U.isArray(t)&&0<=t-parseFloat(t)+1},isPlainObject:function(t){return"object"===U.type(t)&&!t.nodeType&&!U.isWindow(t)&&!(t.constructor&&!H.call(t.constructor.prototype,"isPrototypeOf"))},isEmptyObject:function(t){var e;for(e in t)return!1;return!0},type:function(t){return null==t?t+"":"object"==typeof t||"function"==typeof t?q[z.call(t)]||"object":typeof t},globalEval:function(t){var e,n=eval;(t=U.trim(t))&&(1===t.indexOf("use strict")?((e=Y.createElement("script")).text=t,Y.head.appendChild(e).parentNode.removeChild(e)):n(t))},camelCase:function(t){return t.replace(G,"ms-").replace(Z,B)},nodeName:function(t,e){return t.nodeName&&t.nodeName.toLowerCase()===e.toLowerCase()},each:function(t,e,n){var i=0,r=t.length,o=s(t);if(n){if(o)for(;i<r&&!1!==e.apply(t[i],n);i++);else for(i in t)if(!1===e.apply(t[i],n))break}else if(o)for(;i<r&&!1!==e.call(t[i],i,t[i]);i++);else for(i in t)if(!1===e.call(t[i],i,t[i]))break;return t},trim:function(t){return null==t?"":(t+"").replace(W,"")},makeArray:function(t,e){var n=e||[];return null!=t&&(s(Object(t))?U.merge(n,"string"==typeof t?[t]:t):F.call(n,t)),n},inArray:function(t,e,n){return null==e?-1:V.call(e,t,n)},merge:function(t,e){for(var n=+e.length,i=0,r=t.length;i<n;i++)t[r++]=e[i];return t.length=r,t},grep:function(t,e,n){for(var i=[],r=0,o=t.length,a=!n;r<o;r++)!e(t[r],r)!=a&&i.push(t[r]);return i},map:function(t,e,n){var i,r=0,o=t.length,a=[];if(s(t))for(;r<o;r++)null!=(i=e(t[r],r,n))&&a.push(i);else for(r in t)null!=(i=e(t[r],r,n))&&a.push(i);return R.apply([],a)},guid:1,proxy:function(t,e){var n,i,r;return"string"==typeof e&&(n=t[e],e=t,t=n),U.isFunction(t)?(i=j.call(arguments,2),(r=function(){return t.apply(e||this,i.concat(j.call(arguments)))}).guid=t.guid=t.guid||U.guid++,r):void 0},now:Date.now,support:X}),U.each("Boolean Number String Function Array Date RegExp Object Error".split(" "),function(t,e){q["[object "+e+"]"]=e.toLowerCase()});var K=function(n){function x(t,e,n,i){var r,o,a,s,l,u,c,h,d,p;if((e?e.ownerDocument||e:F)!==A&&M(e),n=n||[],s=(e=e||A).nodeType,"string"!=typeof t||!t||1!==s&&9!==s&&11!==s)return n;if(!i&&O){if(11!==s&&(r=mt.exec(t)))if(a=r[1]){if(9===s){if(!(o=e.getElementById(a))||!o.parentNode)return n;if(o.id===a)return n.push(o),n}else if(e.ownerDocument&&(o=e.ownerDocument.getElementById(a))&&j(e,o)&&o.id===a)return n.push(o),n}else{if(r[2])return K.apply(n,e.getElementsByTagName(t)),n;if((a=r[3])&&b.getElementsByClassName)return K.apply(n,e.getElementsByClassName(a)),n}if(b.qsa&&(!I||!I.test(t))){if(h=c=R,d=e,p=1!==s&&t,1===s&&"object"!==e.nodeName.toLowerCase()){for(u=E(t),(c=e.getAttribute("id"))?h=c.replace(bt,"\\$&"):e.setAttribute("id",h),h="[id='"+h+"'] ",l=u.length;l--;)u[l]=h+g(u[l]);d=yt.test(t)&&f(e.parentNode)||e,p=u.join(",")}if(p)try{return K.apply(n,d.querySelectorAll(p)),n}catch(t){}finally{c||e.removeAttribute("id")}}}return T(t.replace(st,"$1"),e,n,i)}function t(){var i=[];return function t(e,n){return i.push(e+" ")>$.cacheLength&&delete t[i.shift()],t[e+" "]=n}}function l(t){return t[R]=!0,t}function i(t){var e=A.createElement("div");try{return!!t(e)}catch(t){return!1}finally{e.parentNode&&e.parentNode.removeChild(e),e=null}}function e(t,e){for(var n=t.split("|"),i=t.length;i--;)$.attrHandle[n[i]]=e}function u(t,e){var n=e&&t,i=n&&1===t.nodeType&&1===e.nodeType&&(~e.sourceIndex||1<<31)-(~t.sourceIndex||1<<31);if(i)return i;if(n)for(;n=n.nextSibling;)if(n===e)return-1;return t?1:-1}function r(e){return function(t){return"input"===t.nodeName.toLowerCase()&&t.type===e}}function o(n){return function(t){var e=t.nodeName.toLowerCase();return("input"===e||"button"===e)&&t.type===n}}function a(a){return l(function(o){return o=+o,l(function(t,e){for(var n,i=a([],t.length,o),r=i.length;r--;)t[n=i[r]]&&(t[n]=!(e[n]=t[n]))})})}function f(t){return t&&void 0!==t.getElementsByTagName&&t}function s(){}function g(t){for(var e=0,n=t.length,i="";e<n;e++)i+=t[e].value;return i}function h(a,t,e){var s=t.dir,l=e&&"parentNode"===s,u=q++;return t.first?function(t,e,n){for(;t=t[s];)if(1===t.nodeType||l)return a(t,e,n)}:function(t,e,n){var i,r,o=[V,u];if(n){for(;t=t[s];)if((1===t.nodeType||l)&&a(t,e,n))return!0}else for(;t=t[s];)if(1===t.nodeType||l){if((i=(r=t[R]||(t[R]={}))[s])&&i[0]===V&&i[1]===u)return o[2]=i[2];if((r[s]=o)[2]=a(t,e,n))return!0}}}function d(r){return 1<r.length?function(t,e,n){for(var i=r.length;i--;)if(!r[i](t,e,n))return!1;return!0}:r[0]}function w(t,e,n,i,r){for(var o,a=[],s=0,l=t.length,u=null!=e;s<l;s++)!(o=t[s])||n&&!n(o,i,r)||(a.push(o),u&&e.push(s));return a}function y(p,f,g,v,m,t){return v&&!v[R]&&(v=y(v)),m&&!m[R]&&(m=y(m,t)),l(function(t,e,n,i){var r,o,a,s=[],l=[],u=e.length,c=t||function(t,e,n){for(var i=0,r=e.length;i<r;i++)x(t,e[i],n);return n}(f||"*",n.nodeType?[n]:n,[]),h=!p||!t&&f?c:w(c,s,p,n,i),d=g?m||(t?p:u||v)?[]:e:h;if(g&&g(h,d,n,i),v)for(r=w(d,l),v(r,[],n,i),o=r.length;o--;)(a=r[o])&&(d[l[o]]=!(h[l[o]]=a));if(t){if(m||p){if(m){for(r=[],o=d.length;o--;)(a=d[o])&&r.push(h[o]=a);m(null,d=[],r,i)}for(o=d.length;o--;)(a=d[o])&&-1<(r=m?J(t,a):s[o])&&(t[r]=!(e[r]=a))}}else d=w(d===e?d.splice(u,d.length):d),m?m(null,e,d,i):K.apply(e,d)})}function p(t){for(var r,e,n,i=t.length,o=$.relative[t[0].type],a=o||$.relative[" "],s=o?1:0,l=h(function(t){return t===r},a,!0),u=h(function(t){return-1<J(r,t)},a,!0),c=[function(t,e,n){var i=!o&&(n||e!==D)||((r=e).nodeType?l(t,e,n):u(t,e,n));return r=null,i}];s<i;s++)if(e=$.relative[t[s].type])c=[h(d(c),e)];else{if((e=$.filter[t[s].type].apply(null,t[s].matches))[R]){for(n=++s;n<i&&!$.relative[t[n].type];n++);return y(1<s&&d(c),1<s&&g(t.slice(0,s-1).concat({value:" "===t[s-2].type?"*":""})).replace(st,"$1"),e,s<n&&p(t.slice(s,n)),n<i&&p(t=t.slice(n)),n<i&&g(t))}c.push(e)}return d(c)}function v(t,e,n){var i="0x"+e-65536;return i!=i||n?e:i<0?String.fromCharCode(65536+i):String.fromCharCode(i>>10|55296,1023&i|56320)}function c(){M()}var m,b,$,k,C,E,S,T,D,_,P,M,A,N,O,I,B,L,j,R="sizzle"+1*new Date,F=n.document,V=0,q=0,z=t(),H=t(),X=t(),Y=function(t,e){return t===e&&(P=!0),0},U={}.hasOwnProperty,W=[],G=W.pop,Z=W.push,K=W.push,Q=W.slice,J=function(t,e){for(var n=0,i=t.length;n<i;n++)if(t[n]===e)return n;return-1},tt="checked|selected|async|autofocus|autoplay|controls|defer|disabled|hidden|ismap|loop|multiple|open|readonly|required|scoped",et="[\\x20\\t\\r\\n\\f]",nt="(?:\\\\.|[\\w-]|[^\\x00-\\xa0])+",it=nt.replace("w","w#"),rt="\\["+et+"*("+nt+")(?:"+et+"*([*^$|!~]?=)"+et+"*(?:'((?:\\\\.|[^\\\\'])*)'|\"((?:\\\\.|[^\\\\\"])*)\"|("+it+"))|)"+et+"*\\]",ot=":("+nt+")(?:\\((('((?:\\\\.|[^\\\\'])*)'|\"((?:\\\\.|[^\\\\\"])*)\")|((?:\\\\.|[^\\\\()[\\]]|"+rt+")*)|.*)\\)|)",at=new RegExp(et+"+","g"),st=new RegExp("^"+et+"+|((?:^|[^\\\\])(?:\\\\.)*)"+et+"+$","g"),lt=new RegExp("^"+et+"*,"+et+"*"),ut=new RegExp("^"+et+"*([>+~]|"+et+")"+et+"*"),ct=new RegExp("="+et+"*([^\\]'\"]*?)"+et+"*\\]","g"),ht=new RegExp(ot),dt=new RegExp("^"+it+"$"),pt={ID:new RegExp("^#("+nt+")"),CLASS:new RegExp("^\\.("+nt+")"),TAG:new RegExp("^("+nt.replace("w","w*")+")"),ATTR:new RegExp("^"+rt),PSEUDO:new RegExp("^"+ot),CHILD:new RegExp("^:(only|first|last|nth|nth-last)-(child|of-type)(?:\\("+et+"*(even|odd|(([+-]|)(\\d*)n|)"+et+"*(?:([+-]|)"+et+"*(\\d+)|))"+et+"*\\)|)","i"),bool:new RegExp("^(?:"+tt+")$","i"),needsContext:new RegExp("^"+et+"*[>+~]|:(even|odd|eq|gt|lt|nth|first|last)(?:\\("+et+"*((?:-\\d)?\\d*)"+et+"*\\)|)(?=[^-]|$)","i")},ft=/^(?:input|select|textarea|button)$/i,gt=/^h\d$/i,vt=/^[^{]+\{\s*\[native \w/,mt=/^(?:#([\w-]+)|(\w+)|\.([\w-]+))$/,yt=/[+~]/,bt=/'|\\/g,xt=new RegExp("\\\\([\\da-f]{1,6}"+et+"?|("+et+")|.)","ig");try{K.apply(W=Q.call(F.childNodes),F.childNodes),W[F.childNodes.length].nodeType}catch(t){K={apply:W.length?function(t,e){Z.apply(t,Q.call(e))}:function(t,e){for(var n=t.length,i=0;t[n++]=e[i++];);t.length=n-1}}}for(m in b=x.support={},C=x.isXML=function(t){var e=t&&(t.ownerDocument||t).documentElement;return!!e&&"HTML"!==e.nodeName},M=x.setDocument=function(t){var e,n,l=t?t.ownerDocument||t:F;return l!==A&&9===l.nodeType&&l.documentElement?(N=(A=l).documentElement,(n=l.defaultView)&&n!==n.top&&(n.addEventListener?n.addEventListener("unload",c,!1):n.attachEvent&&n.attachEvent("onunload",c)),O=!C(l),b.attributes=i(function(t){return t.className="i",!t.getAttribute("className")}),b.getElementsByTagName=i(function(t){return t.appendChild(l.createComment("")),!t.getElementsByTagName("*").length}),b.getElementsByClassName=vt.test(l.getElementsByClassName),b.getById=i(function(t){return N.appendChild(t).id=R,!l.getElementsByName||!l.getElementsByName(R).length}),b.getById?($.find.ID=function(t,e){if(void 0!==e.getElementById&&O){var n=e.getElementById(t);return n&&n.parentNode?[n]:[]}},$.filter.ID=function(t){var e=t.replace(xt,v);return function(t){return t.getAttribute("id")===e}}):(delete $.find.ID,$.filter.ID=function(t){var n=t.replace(xt,v);return function(t){var e=void 0!==t.getAttributeNode&&t.getAttributeNode("id");return e&&e.value===n}}),$.find.TAG=b.getElementsByTagName?function(t,e){return void 0!==e.getElementsByTagName?e.getElementsByTagName(t):b.qsa?e.querySelectorAll(t):void 0}:function(t,e){var n,i=[],r=0,o=e.getElementsByTagName(t);if("*"!==t)return o;for(;n=o[r++];)1===n.nodeType&&i.push(n);return i},$.find.CLASS=b.getElementsByClassName&&function(t,e){return O?e.getElementsByClassName(t):void 0},B=[],I=[],(b.qsa=vt.test(l.querySelectorAll))&&(i(function(t){N.appendChild(t).innerHTML="<a id='"+R+"'></a><select id='"+R+"-\f]' msallowcapture=''><option selected=''></option></select>",t.querySelectorAll("[msallowcapture^='']").length&&I.push("[*^$]="+et+"*(?:''|\"\")"),t.querySelectorAll("[selected]").length||I.push("\\["+et+"*(?:value|"+tt+")"),t.querySelectorAll("[id~="+R+"-]").length||I.push("~="),t.querySelectorAll(":checked").length||I.push(":checked"),t.querySelectorAll("a#"+R+"+*").length||I.push(".#.+[+~]")}),i(function(t){var e=l.createElement("input");e.setAttribute("type","hidden"),t.appendChild(e).setAttribute("name","D"),t.querySelectorAll("[name=d]").length&&I.push("name"+et+"*[*^$|!~]?="),t.querySelectorAll(":enabled").length||I.push(":enabled",":disabled"),t.querySelectorAll("*,:x"),I.push(",.*:")})),(b.matchesSelector=vt.test(L=N.matches||N.webkitMatchesSelector||N.mozMatchesSelector||N.oMatchesSelector||N.msMatchesSelector))&&i(function(t){b.disconnectedMatch=L.call(t,"div"),L.call(t,"[s!='']:x"),B.push("!=",ot)}),I=I.length&&new RegExp(I.join("|")),B=B.length&&new RegExp(B.join("|")),e=vt.test(N.compareDocumentPosition),j=e||vt.test(N.contains)?function(t,e){var n=9===t.nodeType?t.documentElement:t,i=e&&e.parentNode;return t===i||!(!i||1!==i.nodeType||!(n.contains?n.contains(i):t.compareDocumentPosition&&16&t.compareDocumentPosition(i)))}:function(t,e){if(e)for(;e=e.parentNode;)if(e===t)return!0;return!1},Y=e?function(t,e){if(t===e)return P=!0,0;var n=!t.compareDocumentPosition-!e.compareDocumentPosition;return n||(1&(n=(t.ownerDocument||t)===(e.ownerDocument||e)?t.compareDocumentPosition(e):1)||!b.sortDetached&&e.compareDocumentPosition(t)===n?t===l||t.ownerDocument===F&&j(F,t)?-1:e===l||e.ownerDocument===F&&j(F,e)?1:_?J(_,t)-J(_,e):0:4&n?-1:1)}:function(t,e){if(t===e)return P=!0,0;var n,i=0,r=t.parentNode,o=e.parentNode,a=[t],s=[e];if(!r||!o)return t===l?-1:e===l?1:r?-1:o?1:_?J(_,t)-J(_,e):0;if(r===o)return u(t,e);for(n=t;n=n.parentNode;)a.unshift(n);for(n=e;n=n.parentNode;)s.unshift(n);for(;a[i]===s[i];)i++;return i?u(a[i],s[i]):a[i]===F?-1:s[i]===F?1:0},l):A},x.matches=function(t,e){return x(t,null,null,e)},x.matchesSelector=function(t,e){if((t.ownerDocument||t)!==A&&M(t),e=e.replace(ct,"='$1']"),b.matchesSelector&&O&&(!B||!B.test(e))&&(!I||!I.test(e)))try{var n=L.call(t,e);if(n||b.disconnectedMatch||t.document&&11!==t.document.nodeType)return n}catch(t){}return 0<x(e,A,null,[t]).length},x.contains=function(t,e){return(t.ownerDocument||t)!==A&&M(t),j(t,e)},x.attr=function(t,e){(t.ownerDocument||t)!==A&&M(t);var n=$.attrHandle[e.toLowerCase()],i=n&&U.call($.attrHandle,e.toLowerCase())?n(t,e,!O):void 0;return void 0!==i?i:b.attributes||!O?t.getAttribute(e):(i=t.getAttributeNode(e))&&i.specified?i.value:null},x.error=function(t){throw new Error("Syntax error, unrecognized expression: "+t)},x.uniqueSort=function(t){var e,n=[],i=0,r=0;if(P=!b.detectDuplicates,_=!b.sortStable&&t.slice(0),t.sort(Y),P){for(;e=t[r++];)e===t[r]&&(i=n.push(r));for(;i--;)t.splice(n[i],1)}return _=null,t},k=x.getText=function(t){var e,n="",i=0,r=t.nodeType;if(r){if(1===r||9===r||11===r){if("string"==typeof t.textContent)return t.textContent;for(t=t.firstChild;t;t=t.nextSibling)n+=k(t)}else if(3===r||4===r)return t.nodeValue}else for(;e=t[i++];)n+=k(e);return n},($=x.selectors={cacheLength:50,createPseudo:l,match:pt,attrHandle:{},find:{},relative:{">":{dir:"parentNode",first:!0}," ":{dir:"parentNode"},"+":{dir:"previousSibling",first:!0},"~":{dir:"previousSibling"}},preFilter:{ATTR:function(t){return t[1]=t[1].replace(xt,v),t[3]=(t[3]||t[4]||t[5]||"").replace(xt,v),"~="===t[2]&&(t[3]=" "+t[3]+" "),t.slice(0,4)},CHILD:function(t){return t[1]=t[1].toLowerCase(),"nth"===t[1].slice(0,3)?(t[3]||x.error(t[0]),t[4]=+(t[4]?t[5]+(t[6]||1):2*("even"===t[3]||"odd"===t[3])),t[5]=+(t[7]+t[8]||"odd"===t[3])):t[3]&&x.error(t[0]),t},PSEUDO:function(t){var e,n=!t[6]&&t[2];return pt.CHILD.test(t[0])?null:(t[3]?t[2]=t[4]||t[5]||"":n&&ht.test(n)&&(e=E(n,!0))&&(e=n.indexOf(")",n.length-e)-n.length)&&(t[0]=t[0].slice(0,e),t[2]=n.slice(0,e)),t.slice(0,3))}},filter:{TAG:function(t){var e=t.replace(xt,v).toLowerCase();return"*"===t?function(){return!0}:function(t){return t.nodeName&&t.nodeName.toLowerCase()===e}},CLASS:function(t){var e=z[t+" "];return e||(e=new RegExp("(^|"+et+")"+t+"("+et+"|$)"))&&z(t,function(t){return e.test("string"==typeof t.className&&t.className||void 0!==t.getAttribute&&t.getAttribute("class")||"")})},ATTR:function(n,i,r){return function(t){var e=x.attr(t,n);return null==e?"!="===i:!i||(e+="","="===i?e===r:"!="===i?e!==r:"^="===i?r&&0===e.indexOf(r):"*="===i?r&&-1<e.indexOf(r):"$="===i?r&&e.slice(-r.length)===r:"~="===i?-1<(" "+e.replace(at," ")+" ").indexOf(r):"|="===i&&(e===r||e.slice(0,r.length+1)===r+"-"))}},CHILD:function(p,t,e,f,g){var v="nth"!==p.slice(0,3),m="last"!==p.slice(-4),y="of-type"===t;return 1===f&&0===g?function(t){return!!t.parentNode}:function(t,e,n){var i,r,o,a,s,l,u=v!=m?"nextSibling":"previousSibling",c=t.parentNode,h=y&&t.nodeName.toLowerCase(),d=!n&&!y;if(c){if(v){for(;u;){for(o=t;o=o[u];)if(y?o.nodeName.toLowerCase()===h:1===o.nodeType)return!1;l=u="only"===p&&!l&&"nextSibling"}return!0}if(l=[m?c.firstChild:c.lastChild],m&&d){for(s=(i=(r=c[R]||(c[R]={}))[p]||[])[0]===V&&i[1],a=i[0]===V&&i[2],o=s&&c.childNodes[s];o=++s&&o&&o[u]||(a=s=0)||l.pop();)if(1===o.nodeType&&++a&&o===t){r[p]=[V,s,a];break}}else if(d&&(i=(t[R]||(t[R]={}))[p])&&i[0]===V)a=i[1];else for(;(o=++s&&o&&o[u]||(a=s=0)||l.pop())&&((y?o.nodeName.toLowerCase()!==h:1!==o.nodeType)||!++a||(d&&((o[R]||(o[R]={}))[p]=[V,a]),o!==t)););return(a-=g)===f||a%f==0&&0<=a/f}}},PSEUDO:function(t,o){var e,a=$.pseudos[t]||$.setFilters[t.toLowerCase()]||x.error("unsupported pseudo: "+t);return a[R]?a(o):1<a.length?(e=[t,t,"",o],$.setFilters.hasOwnProperty(t.toLowerCase())?l(function(t,e){for(var n,i=a(t,o),r=i.length;r--;)t[n=J(t,i[r])]=!(e[n]=i[r])}):function(t){return a(t,0,e)}):a}},pseudos:{not:l(function(t){var i=[],r=[],s=S(t.replace(st,"$1"));return s[R]?l(function(t,e,n,i){for(var r,o=s(t,null,i,[]),a=t.length;a--;)(r=o[a])&&(t[a]=!(e[a]=r))}):function(t,e,n){return i[0]=t,s(i,null,n,r),i[0]=null,!r.pop()}}),has:l(function(e){return function(t){return 0<x(e,t).length}}),contains:l(function(e){return e=e.replace(xt,v),function(t){return-1<(t.textContent||t.innerText||k(t)).indexOf(e)}}),lang:l(function(n){return dt.test(n||"")||x.error("unsupported lang: "+n),n=n.replace(xt,v).toLowerCase(),function(t){var e;do{if(e=O?t.lang:t.getAttribute("xml:lang")||t.getAttribute("lang"))return(e=e.toLowerCase())===n||0===e.indexOf(n+"-")}while((t=t.parentNode)&&1===t.nodeType);return!1}}),target:function(t){var e=n.location&&n.location.hash;return e&&e.slice(1)===t.id},root:function(t){return t===N},focus:function(t){return t===A.activeElement&&(!A.hasFocus||A.hasFocus())&&!!(t.type||t.href||~t.tabIndex)},enabled:function(t){return!1===t.disabled},disabled:function(t){return!0===t.disabled},checked:function(t){var e=t.nodeName.toLowerCase();return"input"===e&&!!t.checked||"option"===e&&!!t.selected},selected:function(t){return t.parentNode&&t.parentNode.selectedIndex,!0===t.selected},empty:function(t){for(t=t.firstChild;t;t=t.nextSibling)if(t.nodeType<6)return!1;return!0},parent:function(t){return!$.pseudos.empty(t)},header:function(t){return gt.test(t.nodeName)},input:function(t){return ft.test(t.nodeName)},button:function(t){var e=t.nodeName.toLowerCase();return"input"===e&&"button"===t.type||"button"===e},text:function(t){var e;return"input"===t.nodeName.toLowerCase()&&"text"===t.type&&(null==(e=t.getAttribute("type"))||"text"===e.toLowerCase())},first:a(function(){return[0]}),last:a(function(t,e){return[e-1]}),eq:a(function(t,e,n){return[n<0?n+e:n]}),even:a(function(t,e){for(var n=0;n<e;n+=2)t.push(n);return t}),odd:a(function(t,e){for(var n=1;n<e;n+=2)t.push(n);return t}),lt:a(function(t,e,n){for(var i=n<0?n+e:n;0<=--i;)t.push(i);return t}),gt:a(function(t,e,n){for(var i=n<0?n+e:n;++i<e;)t.push(i);return t})}}).pseudos.nth=$.pseudos.eq,{radio:!0,checkbox:!0,file:!0,password:!0,image:!0})$.pseudos[m]=r(m);for(m in{submit:!0,reset:!0})$.pseudos[m]=o(m);return s.prototype=$.filters=$.pseudos,$.setFilters=new s,E=x.tokenize=function(t,e){var n,i,r,o,a,s,l,u=H[t+" "];if(u)return e?0:u.slice(0);for(a=t,s=[],l=$.preFilter;a;){for(o in n&&!(i=lt.exec(a))||(i&&(a=a.slice(i[0].length)||a),s.push(r=[])),n=!1,(i=ut.exec(a))&&(n=i.shift(),r.push({value:n,type:i[0].replace(st," ")}),a=a.slice(n.length)),$.filter)!(i=pt[o].exec(a))||l[o]&&!(i=l[o](i))||(n=i.shift(),r.push({value:n,type:o,matches:i}),a=a.slice(n.length));if(!n)break}return e?a.length:a?x.error(t):H(t,s).slice(0)},S=x.compile=function(t,e){var n,i=[],r=[],o=X[t+" "];if(!o){for(e||(e=E(t)),n=e.length;n--;)(o=p(e[n]))[R]?i.push(o):r.push(o);(o=X(t,function(v,m){function t(t,e,n,i,r){var o,a,s,l=0,u="0",c=t&&[],h=[],d=D,p=t||b&&$.find.TAG("*",r),f=V+=null==d?1:Math.random()||.1,g=p.length;for(r&&(D=e!==A&&e);u!==g&&null!=(o=p[u]);u++){if(b&&o){for(a=0;s=v[a++];)if(s(o,e,n)){i.push(o);break}r&&(V=f)}y&&((o=!s&&o)&&l--,t&&c.push(o))}if(l+=u,y&&u!==l){for(a=0;s=m[a++];)s(c,h,e,n);if(t){if(0<l)for(;u--;)c[u]||h[u]||(h[u]=G.call(i));h=w(h)}K.apply(i,h),r&&!t&&0<h.length&&1<l+m.length&&x.uniqueSort(i)}return r&&(V=f,D=d),c}var y=0<m.length,b=0<v.length;return y?l(t):t}(r,i))).selector=t}return o},T=x.select=function(t,e,n,i){var r,o,a,s,l,u="function"==typeof t&&t,c=!i&&E(t=u.selector||t);if(n=n||[],1===c.length){if(2<(o=c[0]=c[0].slice(0)).length&&"ID"===(a=o[0]).type&&b.getById&&9===e.nodeType&&O&&$.relative[o[1].type]){if(!(e=($.find.ID(a.matches[0].replace(xt,v),e)||[])[0]))return n;u&&(e=e.parentNode),t=t.slice(o.shift().value.length)}for(r=pt.needsContext.test(t)?0:o.length;r--&&(a=o[r],!$.relative[s=a.type]);)if((l=$.find[s])&&(i=l(a.matches[0].replace(xt,v),yt.test(o[0].type)&&f(e.parentNode)||e))){if(o.splice(r,1),!(t=i.length&&g(o)))return K.apply(n,i),n;break}}return(u||S(t,c))(i,e,!O,n,yt.test(t)&&f(e.parentNode)||e),n},b.sortStable=R.split("").sort(Y).join("")===R,b.detectDuplicates=!!P,M(),b.sortDetached=i(function(t){return 1&t.compareDocumentPosition(A.createElement("div"))}),i(function(t){return t.innerHTML="<a href='#'></a>","#"===t.firstChild.getAttribute("href")})||e("type|href|height|width",function(t,e,n){return n?void 0:t.getAttribute(e,"type"===e.toLowerCase()?1:2)}),b.attributes&&i(function(t){return t.innerHTML="<input/>",t.firstChild.setAttribute("value",""),""===t.firstChild.getAttribute("value")})||e("value",function(t,e,n){return n||"input"!==t.nodeName.toLowerCase()?void 0:t.defaultValue}),i(function(t){return null==t.getAttribute("disabled")})||e(tt,function(t,e,n){var i;return n?void 0:!0===t[e]?e.toLowerCase():(i=t.getAttributeNode(e))&&i.specified?i.value:null}),x}(f);U.find=K,U.expr=K.selectors,U.expr[":"]=U.expr.pseudos,U.unique=K.uniqueSort,U.text=K.getText,U.isXMLDoc=K.isXML,U.contains=K.contains;var Q=U.expr.match.needsContext,J=/^<(\w+)\s*\/?>(?:<\/\1>|)$/,tt=/^.[^:#\[\.,]*$/;U.filter=function(t,e,n){var i=e[0];return n&&(t=":not("+t+")"),1===e.length&&1===i.nodeType?U.find.matchesSelector(i,t)?[i]:[]:U.find.matches(t,U.grep(e,function(t){return 1===t.nodeType}))},U.fn.extend({find:function(t){var e,n=this.length,i=[],r=this;if("string"!=typeof t)return this.pushStack(U(t).filter(function(){for(e=0;e<n;e++)if(U.contains(r[e],this))return!0}));for(e=0;e<n;e++)U.find(t,r[e],i);return(i=this.pushStack(1<n?U.unique(i):i)).selector=this.selector?this.selector+" "+t:t,i},filter:function(t){return this.pushStack(e(this,t||[],!1))},not:function(t){return this.pushStack(e(this,t||[],!0))},is:function(t){return!!e(this,"string"==typeof t&&Q.test(t)?U(t):t||[],!1).length}});var et,nt=/^(?:\s*(<[\w\W]+>)[^>]*|#([\w-]*))$/;(U.fn.init=function(t,e){var n,i;if(!t)return this;if("string"!=typeof t)return t.nodeType?(this.context=this[0]=t,this.length=1,this):U.isFunction(t)?void 0!==et.ready?et.ready(t):t(U):(void 0!==t.selector&&(this.selector=t.selector,this.context=t.context),U.makeArray(t,this));if(!(n="<"===t[0]&&">"===t[t.length-1]&&3<=t.length?[null,t,null]:nt.exec(t))||!n[1]&&e)return!e||e.jquery?(e||et).find(t):this.constructor(e).find(t);if(n[1]){if(e=e instanceof U?e[0]:e,U.merge(this,U.parseHTML(n[1],e&&e.nodeType?e.ownerDocument||e:Y,!0)),J.test(n[1])&&U.isPlainObject(e))for(n in e)U.isFunction(this[n])?this[n](e[n]):this.attr(n,e[n]);return this}return(i=Y.getElementById(n[2]))&&i.parentNode&&(this.length=1,this[0]=i),this.context=Y,this.selector=t,this}).prototype=U.fn,et=U(Y);var it=/^(?:parents|prev(?:Until|All))/,rt={children:!0,contents:!0,next:!0,prev:!0};U.extend({dir:function(t,e,n){for(var i=[],r=void 0!==n;(t=t[e])&&9!==t.nodeType;)if(1===t.nodeType){if(r&&U(t).is(n))break;i.push(t)}return i},sibling:function(t,e){for(var n=[];t;t=t.nextSibling)1===t.nodeType&&t!==e&&n.push(t);return n}}),U.fn.extend({has:function(t){var e=U(t,this),n=e.length;return this.filter(function(){for(var t=0;t<n;t++)if(U.contains(this,e[t]))return!0})},closest:function(t,e){for(var n,i=0,r=this.length,o=[],a=Q.test(t)||"string"!=typeof t?U(t,e||this.context):0;i<r;i++)for(n=this[i];n&&n!==e;n=n.parentNode)if(n.nodeType<11&&(a?-1<a.index(n):1===n.nodeType&&U.find.matchesSelector(n,t))){o.push(n);break}return this.pushStack(1<o.length?U.unique(o):o)},index:function(t){return t?"string"==typeof t?V.call(U(t),this[0]):V.call(this,t.jquery?t[0]:t):this[0]&&this[0].parentNode?this.first().prevAll().length:-1},add:function(t,e){return this.pushStack(U.unique(U.merge(this.get(),U(t,e))))},addBack:function(t){return this.add(null==t?this.prevObject:this.prevObject.filter(t))}}),U.each({parent:function(t){var e=t.parentNode;return e&&11!==e.nodeType?e:null},parents:function(t){return U.dir(t,"parentNode")},parentsUntil:function(t,e,n){return U.dir(t,"parentNode",n)},next:function(t){return n(t,"nextSibling")},prev:function(t){return n(t,"previousSibling")},nextAll:function(t){return U.dir(t,"nextSibling")},prevAll:function(t){return U.dir(t,"previousSibling")},nextUntil:function(t,e,n){return U.dir(t,"nextSibling",n)},prevUntil:function(t,e,n){return U.dir(t,"previousSibling",n)},siblings:function(t){return U.sibling((t.parentNode||{}).firstChild,t)},children:function(t){return U.sibling(t.firstChild)},contents:function(t){return t.contentDocument||U.merge([],t.childNodes)}},function(i,r){U.fn[i]=function(t,e){var n=U.map(this,r,t);return"Until"!==i.slice(-5)&&(e=t),e&&"string"==typeof e&&(n=U.filter(e,n)),1<this.length&&(rt[i]||U.unique(n),it.test(i)&&n.reverse()),this.pushStack(n)}});var ot,at=/\S+/g,st={};U.Callbacks=function(r){r="string"==typeof r?st[r]||function(t){var n=st[t]={};return U.each(t.match(at)||[],function(t,e){n[e]=!0}),n}(r):U.extend({},r);var e,n,i,o,a,s,l=[],u=!r.once&&[],c=function(t){for(e=r.memory&&t,n=!0,s=o||0,o=0,a=l.length,i=!0;l&&s<a;s++)if(!1===l[s].apply(t[0],t[1])&&r.stopOnFalse){e=!1;break}i=!1,l&&(u?u.length&&c(u.shift()):e?l=[]:h.disable())},h={add:function(){if(l){var t=l.length;!function i(t){U.each(t,function(t,e){var n=U.type(e);"function"===n?r.unique&&h.has(e)||l.push(e):e&&e.length&&"string"!==n&&i(e)})}(arguments),i?a=l.length:e&&(o=t,c(e))}return this},remove:function(){return l&&U.each(arguments,function(t,e){for(var n;-1<(n=U.inArray(e,l,n));)l.splice(n,1),i&&(n<=a&&a--,n<=s&&s--)}),this},has:function(t){return t?-1<U.inArray(t,l):!(!l||!l.length)},empty:function(){return l=[],a=0,this},disable:function(){return l=u=e=void 0,this},disabled:function(){return!l},lock:function(){return u=void 0,e||h.disable(),this},locked:function(){return!u},fireWith:function(t,e){return!l||n&&!u||(e=[t,(e=e||[]).slice?e.slice():e],i?u.push(e):c(e)),this},fire:function(){return h.fireWith(this,arguments),this},fired:function(){return!!n}};return h},U.extend({Deferred:function(t){var o=[["resolve","done",U.Callbacks("once memory"),"resolved"],["reject","fail",U.Callbacks("once memory"),"rejected"],["notify","progress",U.Callbacks("memory")]],r="pending",a={state:function(){return r},always:function(){return s.done(arguments).fail(arguments),this},then:function(){var r=arguments;return U.Deferred(function(i){U.each(o,function(t,e){var n=U.isFunction(r[t])&&r[t];s[e[1]](function(){var t=n&&n.apply(this,arguments);t&&U.isFunction(t.promise)?t.promise().done(i.resolve).fail(i.reject).progress(i.notify):i[e[0]+"With"](this===a?i.promise():this,n?[t]:arguments)})}),r=null}).promise()},promise:function(t){return null!=t?U.extend(t,a):a}},s={};return a.pipe=a.then,U.each(o,function(t,e){var n=e[2],i=e[3];a[e[1]]=n.add,i&&n.add(function(){r=i},o[1^t][2].disable,o[2][2].lock),s[e[0]]=function(){return s[e[0]+"With"](this===s?a:this,arguments),this},s[e[0]+"With"]=n.fireWith}),a.promise(s),t&&t.call(s,s),s},when:function(t){function e(e,n,i){return function(t){n[e]=this,i[e]=1<arguments.length?j.call(arguments):t,i===r?u.notifyWith(n,i):--l||u.resolveWith(n,i)}}var r,n,i,o=0,a=j.call(arguments),s=a.length,l=1!==s||t&&U.isFunction(t.promise)?s:0,u=1===l?t:U.Deferred();if(1<s)for(r=new Array(s),n=new Array(s),i=new Array(s);o<s;o++)a[o]&&U.isFunction(a[o].promise)?a[o].promise().done(e(o,i,a)).fail(u.reject).progress(e(o,n,r)):--l;return l||u.resolveWith(i,a),u.promise()}}),U.fn.ready=function(t){return U.ready.promise().done(t),this},U.extend({isReady:!1,readyWait:1,holdReady:function(t){t?U.readyWait++:U.ready(!0)},ready:function(t){(!0===t?--U.readyWait:U.isReady)||((U.isReady=!0)!==t&&0<--U.readyWait||(ot.resolveWith(Y,[U]),U.fn.triggerHandler&&(U(Y).triggerHandler("ready"),U(Y).off("ready"))))}}),U.ready.promise=function(t){return ot||(ot=U.Deferred(),"complete"===Y.readyState?setTimeout(U.ready):(Y.addEventListener("DOMContentLoaded",i,!1),f.addEventListener("load",i,!1))),ot.promise(t)},U.ready.promise();var lt=U.access=function(t,e,n,i,r,o,a){var s=0,l=t.length,u=null==n;if("object"===U.type(n))for(s in r=!0,n)U.access(t,e,s,n[s],!0,o,a);else if(void 0!==i&&(r=!0,U.isFunction(i)||(a=!0),u&&(e=a?(e.call(t,i),null):(u=e,function(t,e,n){return u.call(U(t),n)})),e))for(;s<l;s++)e(t[s],n,a?i:i.call(t[s],s,e(t[s],n)));return r?t:u?e.call(t):l?e(t[0],n):o};U.acceptData=function(t){return 1===t.nodeType||9===t.nodeType||!+t.nodeType},r.uid=1,r.accepts=U.acceptData,r.prototype={key:function(e){if(!r.accepts(e))return 0;var n={},i=e[this.expando];if(!i){i=r.uid++;try{n[this.expando]={value:i},Object.defineProperties(e,n)}catch(t){n[this.expando]=i,U.extend(e,n)}}return this.cache[i]||(this.cache[i]={}),i},set:function(t,e,n){var i,r=this.key(t),o=this.cache[r];if("string"==typeof e)o[e]=n;else if(U.isEmptyObject(o))U.extend(this.cache[r],e);else for(i in e)o[i]=e[i];return o},get:function(t,e){var n=this.cache[this.key(t)];return void 0===e?n:n[e]},access:function(t,e,n){var i;return void 0===e||e&&"string"==typeof e&&void 0===n?void 0!==(i=this.get(t,e))?i:this.get(t,U.camelCase(e)):(this.set(t,e,n),void 0!==n?n:e)},remove:function(t,e){var n,i,r,o=this.key(t),a=this.cache[o];if(void 0===e)this.cache[o]={};else{n=(i=U.isArray(e)?e.concat(e.map(U.camelCase)):(r=U.camelCase(e),e in a?[e,r]:(i=r)in a?[i]:i.match(at)||[])).length;for(;n--;)delete a[i[n]]}},hasData:function(t){return!U.isEmptyObject(this.cache[t[this.expando]]||{})},discard:function(t){t[this.expando]&&delete this.cache[t[this.expando]]}};var ut=new r,ct=new r,ht=/^(?:\{[\w\W]*\}|\[[\w\W]*\])$/,dt=/([A-Z])/g;U.extend({hasData:function(t){return ct.hasData(t)||ut.hasData(t)},data:function(t,e,n){return ct.access(t,e,n)},removeData:function(t,e){ct.remove(t,e)},_data:function(t,e,n){return ut.access(t,e,n)},_removeData:function(t,e){ut.remove(t,e)}}),U.fn.extend({data:function(i,t){var e,n,r,o=this[0],a=o&&o.attributes;if(void 0!==i)return"object"==typeof i?this.each(function(){ct.set(this,i)}):lt(this,function(e){var t,n=U.camelCase(i);if(o&&void 0===e){if(void 0!==(t=ct.get(o,i)))return t;if(void 0!==(t=ct.get(o,n)))return t;if(void 0!==(t=l(o,n,void 0)))return t}else this.each(function(){var t=ct.get(this,n);ct.set(this,n,e),-1!==i.indexOf("-")&&void 0!==t&&ct.set(this,i,e)})},null,t,1<arguments.length,null,!0);if(this.length&&(r=ct.get(o),1===o.nodeType&&!ut.get(o,"hasDataAttrs"))){for(e=a.length;e--;)a[e]&&(0===(n=a[e].name).indexOf("data-")&&(n=U.camelCase(n.slice(5)),l(o,n,r[n])));ut.set(o,"hasDataAttrs",!0)}return r},removeData:function(t){return this.each(function(){ct.remove(this,t)})}}),U.extend({queue:function(t,e,n){var i;return t?(e=(e||"fx")+"queue",i=ut.get(t,e),n&&(!i||U.isArray(n)?i=ut.access(t,e,U.makeArray(n)):i.push(n)),i||[]):void 0},dequeue:function(t,e){e=e||"fx";var n=U.queue(t,e),i=n.length,r=n.shift(),o=U._queueHooks(t,e);"inprogress"===r&&(r=n.shift(),i--),r&&("fx"===e&&n.unshift("inprogress"),delete o.stop,r.call(t,function(){U.dequeue(t,e)},o)),!i&&o&&o.empty.fire()},_queueHooks:function(t,e){var n=e+"queueHooks";return ut.get(t,n)||ut.access(t,n,{empty:U.Callbacks("once memory").add(function(){ut.remove(t,[e+"queue",n])})})}}),U.fn.extend({queue:function(e,n){var t=2;return"string"!=typeof e&&(n=e,e="fx",t--),arguments.length<t?U.queue(this[0],e):void 0===n?this:this.each(function(){var t=U.queue(this,e,n);U._queueHooks(this,e),"fx"===e&&"inprogress"!==t[0]&&U.dequeue(this,e)})},dequeue:function(t){return this.each(function(){U.dequeue(this,t)})},clearQueue:function(t){return this.queue(t||"fx",[])},promise:function(t,e){function n(){--r||o.resolveWith(a,[a])}var i,r=1,o=U.Deferred(),a=this,s=this.length;for("string"!=typeof t&&(e=t,t=void 0),t=t||"fx";s--;)(i=ut.get(a[s],t+"queueHooks"))&&i.empty&&(r++,i.empty.add(n));return n(),o.promise(e)}});function pt(t,e){return t=e||t,"none"===U.css(t,"display")||!U.contains(t.ownerDocument,t)}var ft,gt,vt=/[+-]?(?:\d*\.|)\d+(?:[eE][+-]?\d+|)/.source,mt=["Top","Right","Bottom","Left"],yt=/^(?:checkbox|radio)$/i;ft=Y.createDocumentFragment().appendChild(Y.createElement("div")),(gt=Y.createElement("input")).setAttribute("type","radio"),gt.setAttribute("checked","checked"),gt.setAttribute("name","t"),ft.appendChild(gt),X.checkClone=ft.cloneNode(!0).cloneNode(!0).lastChild.checked,ft.innerHTML="<textarea>x</textarea>",X.noCloneChecked=!!ft.cloneNode(!0).lastChild.defaultValue;var bt="undefined";X.focusinBubbles="onfocusin"in f;var xt=/^key/,wt=/^(?:mouse|pointer|contextmenu)|click/,$t=/^(?:focusinfocus|focusoutblur)$/,kt=/^([^.]*)(?:\.(.+)|)$/;U.event={global:{},add:function(e,t,n,i,r){var o,a,s,l,u,c,h,d,p,f,g,v=ut.get(e);if(v)for(n.handler&&(n=(o=n).handler,r=o.selector),n.guid||(n.guid=U.guid++),(l=v.events)||(l=v.events={}),(a=v.handle)||(a=v.handle=function(t){return typeof U!=bt&&U.event.triggered!==t.type?U.event.dispatch.apply(e,arguments):void 0}),u=(t=(t||"").match(at)||[""]).length;u--;)p=g=(s=kt.exec(t[u])||[])[1],f=(s[2]||"").split(".").sort(),p&&(h=U.event.special[p]||{},p=(r?h.delegateType:h.bindType)||p,h=U.event.special[p]||{},c=U.extend({type:p,origType:g,data:i,handler:n,guid:n.guid,selector:r,needsContext:r&&U.expr.match.needsContext.test(r),namespace:f.join(".")},o),(d=l[p])||((d=l[p]=[]).delegateCount=0,h.setup&&!1!==h.setup.call(e,i,f,a)||e.addEventListener&&e.addEventListener(p,a,!1)),h.add&&(h.add.call(e,c),c.handler.guid||(c.handler.guid=n.guid)),r?d.splice(d.delegateCount++,0,c):d.push(c),U.event.global[p]=!0)},remove:function(t,e,n,i,r){var o,a,s,l,u,c,h,d,p,f,g,v=ut.hasData(t)&&ut.get(t);if(v&&(l=v.events)){for(u=(e=(e||"").match(at)||[""]).length;u--;)if(p=g=(s=kt.exec(e[u])||[])[1],f=(s[2]||"").split(".").sort(),p){for(h=U.event.special[p]||{},d=l[p=(i?h.delegateType:h.bindType)||p]||[],s=s[2]&&new RegExp("(^|\\.)"+f.join("\\.(?:.*\\.|)")+"(\\.|$)"),a=o=d.length;o--;)c=d[o],!r&&g!==c.origType||n&&n.guid!==c.guid||s&&!s.test(c.namespace)||i&&i!==c.selector&&("**"!==i||!c.selector)||(d.splice(o,1),c.selector&&d.delegateCount--,h.remove&&h.remove.call(t,c));a&&!d.length&&(h.teardown&&!1!==h.teardown.call(t,f,v.handle)||U.removeEvent(t,p,v.handle),delete l[p])}else for(p in l)U.event.remove(t,p+e[u],n,i,!0);U.isEmptyObject(l)&&(delete v.handle,ut.remove(t,"events"))}},trigger:function(t,e,n,i){var r,o,a,s,l,u,c,h=[n||Y],d=H.call(t,"type")?t.type:t,p=H.call(t,"namespace")?t.namespace.split("."):[];if(o=a=n=n||Y,3!==n.nodeType&&8!==n.nodeType&&!$t.test(d+U.event.triggered)&&(0<=d.indexOf(".")&&(d=(p=d.split(".")).shift(),p.sort()),l=d.indexOf(":")<0&&"on"+d,(t=t[U.expando]?t:new U.Event(d,"object"==typeof t&&t)).isTrigger=i?2:3,t.namespace=p.join("."),t.namespace_re=t.namespace?new RegExp("(^|\\.)"+p.join("\\.(?:.*\\.|)")+"(\\.|$)"):null,t.result=void 0,t.target||(t.target=n),e=null==e?[t]:U.makeArray(e,[t]),c=U.event.special[d]||{},i||!c.trigger||!1!==c.trigger.apply(n,e))){if(!i&&!c.noBubble&&!U.isWindow(n)){for(s=c.delegateType||d,$t.test(s+d)||(o=o.parentNode);o;o=o.parentNode)h.push(o),a=o;a===(n.ownerDocument||Y)&&h.push(a.defaultView||a.parentWindow||f)}for(r=0;(o=h[r++])&&!t.isPropagationStopped();)t.type=1<r?s:c.bindType||d,(u=(ut.get(o,"events")||{})[t.type]&&ut.get(o,"handle"))&&u.apply(o,e),(u=l&&o[l])&&u.apply&&U.acceptData(o)&&(t.result=u.apply(o,e),!1===t.result&&t.preventDefault());return t.type=d,i||t.isDefaultPrevented()||c._default&&!1!==c._default.apply(h.pop(),e)||!U.acceptData(n)||l&&U.isFunction(n[d])&&!U.isWindow(n)&&((a=n[l])&&(n[l]=null),n[U.event.triggered=d](),U.event.triggered=void 0,a&&(n[l]=a)),t.result}},dispatch:function(t){t=U.event.fix(t);var e,n,i,r,o,a=[],s=j.call(arguments),l=(ut.get(this,"events")||{})[t.type]||[],u=U.event.special[t.type]||{};if((s[0]=t).delegateTarget=this,!u.preDispatch||!1!==u.preDispatch.call(this,t)){for(a=U.event.handlers.call(this,t,l),e=0;(r=a[e++])&&!t.isPropagationStopped();)for(t.currentTarget=r.elem,n=0;(o=r.handlers[n++])&&!t.isImmediatePropagationStopped();)t.namespace_re&&!t.namespace_re.test(o.namespace)||(t.handleObj=o,t.data=o.data,void 0!==(i=((U.event.special[o.origType]||{}).handle||o.handler).apply(r.elem,s))&&!1===(t.result=i)&&(t.preventDefault(),t.stopPropagation()));return u.postDispatch&&u.postDispatch.call(this,t),t.result}},handlers:function(t,e){var n,i,r,o,a=[],s=e.delegateCount,l=t.target;if(s&&l.nodeType&&(!t.button||"click"!==t.type))for(;l!==this;l=l.parentNode||this)if(!0!==l.disabled||"click"!==t.type){for(i=[],n=0;n<s;n++)void 0===i[r=(o=e[n]).selector+" "]&&(i[r]=o.needsContext?0<=U(r,this).index(l):U.find(r,this,null,[l]).length),i[r]&&i.push(o);i.length&&a.push({elem:l,handlers:i})}return s<e.length&&a.push({elem:this,handlers:e.slice(s)}),a},props:"altKey bubbles cancelable ctrlKey currentTarget eventPhase metaKey relatedTarget shiftKey target timeStamp view which".split(" "),fixHooks:{},keyHooks:{props:"char charCode key keyCode".split(" "),filter:function(t,e){return null==t.which&&(t.which=null!=e.charCode?e.charCode:e.keyCode),t}},mouseHooks:{props:"button buttons clientX clientY offsetX offsetY pageX pageY screenX screenY toElement".split(" "),filter:function(t,e){var n,i,r,o=e.button;return null==t.pageX&&null!=e.clientX&&(i=(n=t.target.ownerDocument||Y).documentElement,r=n.body,t.pageX=e.clientX+(i&&i.scrollLeft||r&&r.scrollLeft||0)-(i&&i.clientLeft||r&&r.clientLeft||0),t.pageY=e.clientY+(i&&i.scrollTop||r&&r.scrollTop||0)-(i&&i.clientTop||r&&r.clientTop||0)),t.which||void 0===o||(t.which=1&o?1:2&o?3:4&o?2:0),t}},fix:function(t){if(t[U.expando])return t;var e,n,i,r=t.type,o=t,a=this.fixHooks[r];for(a||(this.fixHooks[r]=a=wt.test(r)?this.mouseHooks:xt.test(r)?this.keyHooks:{}),i=a.props?this.props.concat(a.props):this.props,t=new U.Event(o),e=i.length;e--;)t[n=i[e]]=o[n];return t.target||(t.target=Y),3===t.target.nodeType&&(t.target=t.target.parentNode),a.filter?a.filter(t,o):t},special:{load:{noBubble:!0},focus:{trigger:function(){return this!==a()&&this.focus?(this.focus(),!1):void 0},delegateType:"focusin"},blur:{trigger:function(){return this===a()&&this.blur?(this.blur(),!1):void 0},delegateType:"focusout"},click:{trigger:function(){return"checkbox"===this.type&&this.click&&U.nodeName(this,"input")?(this.click(),!1):void 0},_default:function(t){return U.nodeName(t.target,"a")}},beforeunload:{postDispatch:function(t){void 0!==t.result&&t.originalEvent&&(t.originalEvent.returnValue=t.result)}}},simulate:function(t,e,n,i){var r=U.extend(new U.Event,n,{type:t,isSimulated:!0,originalEvent:{}});i?U.event.trigger(r,null,e):U.event.dispatch.call(e,r),r.isDefaultPrevented()&&n.preventDefault()}},U.removeEvent=function(t,e,n){t.removeEventListener&&t.removeEventListener(e,n,!1)},U.Event=function(t,e){return this instanceof U.Event?(t&&t.type?(this.originalEvent=t,this.type=t.type,this.isDefaultPrevented=t.defaultPrevented||void 0===t.defaultPrevented&&!1===t.returnValue?o:u):this.type=t,e&&U.extend(this,e),this.timeStamp=t&&t.timeStamp||U.now(),void(this[U.expando]=!0)):new U.Event(t,e)},U.Event.prototype={isDefaultPrevented:u,isPropagationStopped:u,isImmediatePropagationStopped:u,preventDefault:function(){var t=this.originalEvent;this.isDefaultPrevented=o,t&&t.preventDefault&&t.preventDefault()},stopPropagation:function(){var t=this.originalEvent;this.isPropagationStopped=o,t&&t.stopPropagation&&t.stopPropagation()},stopImmediatePropagation:function(){var t=this.originalEvent;this.isImmediatePropagationStopped=o,t&&t.stopImmediatePropagation&&t.stopImmediatePropagation(),this.stopPropagation()}},U.each({mouseenter:"mouseover",mouseleave:"mouseout",pointerenter:"pointerover",pointerleave:"pointerout"},function(t,r){U.event.special[t]={delegateType:r,bindType:r,handle:function(t){var e,n=t.relatedTarget,i=t.handleObj;return n&&(n===this||U.contains(this,n))||(t.type=i.origType,e=i.handler.apply(this,arguments),t.type=r),e}}}),X.focusinBubbles||U.each({focus:"focusin",blur:"focusout"},function(n,i){function r(t){U.event.simulate(i,t.target,U.event.fix(t),!0)}U.event.special[i]={setup:function(){var t=this.ownerDocument||this,e=ut.access(t,i);e||t.addEventListener(n,r,!0),ut.access(t,i,(e||0)+1)},teardown:function(){var t=this.ownerDocument||this,e=ut.access(t,i)-1;e?ut.access(t,i,e):(t.removeEventListener(n,r,!0),ut.remove(t,i))}}}),U.fn.extend({on:function(t,e,n,i,r){var o,a;if("object"==typeof t){for(a in"string"!=typeof e&&(n=n||e,e=void 0),t)this.on(a,e,n,t[a],r);return this}if(null==n&&null==i?(i=e,n=e=void 0):null==i&&("string"==typeof e?(i=n,n=void 0):(i=n,n=e,e=void 0)),!1===i)i=u;else if(!i)return this;return 1===r&&(o=i,(i=function(t){return U().off(t),o.apply(this,arguments)}).guid=o.guid||(o.guid=U.guid++)),this.each(function(){U.event.add(this,t,i,n,e)})},one:function(t,e,n,i){return this.on(t,e,n,i,1)},off:function(t,e,n){var i,r;if(t&&t.preventDefault&&t.handleObj)return i=t.handleObj,U(t.delegateTarget).off(i.namespace?i.origType+"."+i.namespace:i.origType,i.selector,i.handler),this;if("object"!=typeof t)return!1!==e&&"function"!=typeof e||(n=e,e=void 0),!1===n&&(n=u),this.each(function(){U.event.remove(this,t,n,e)});for(r in t)this.off(r,e,t[r]);return this},trigger:function(t,e){return this.each(function(){U.event.trigger(t,e,this)})},triggerHandler:function(t,e){var n=this[0];return n?U.event.trigger(t,e,n,!0):void 0}});var Ct=/<(?!area|br|col|embed|hr|img|input|link|meta|param)(([\w:]+)[^>]*)\/>/gi,Et=/<([\w:]+)/,St=/<|&#?\w+;/,Tt=/<(?:script|style|link)/i,Dt=/checked\s*(?:[^=]|=\s*.checked.)/i,_t=/^$|\/(?:java|ecma)script/i,Pt=/^true\/(.*)/,Mt=/^\s*<!(?:\[CDATA\[|--)|(?:\]\]|--)>\s*$/g,At={option:[1,"<select multiple='multiple'>","</select>"],thead:[1,"<table>","</table>"],col:[2,"<table><colgroup>","</colgroup></table>"],tr:[2,"<table><tbody>","</tbody></table>"],td:[3,"<table><tbody><tr>","</tr></tbody></table>"],_default:[0,"",""]};At.optgroup=At.option,At.tbody=At.tfoot=At.colgroup=At.caption=At.thead,At.th=At.td,U.extend({clone:function(t,e,n){var i,r,o,a,s,l,u,c=t.cloneNode(!0),h=U.contains(t.ownerDocument,t);if(!(X.noCloneChecked||1!==t.nodeType&&11!==t.nodeType||U.isXMLDoc(t)))for(a=y(c),i=0,r=(o=y(t)).length;i<r;i++)s=o[i],l=a[i],void 0,"input"===(u=l.nodeName.toLowerCase())&&yt.test(s.type)?l.checked=s.checked:"input"!==u&&"textarea"!==u||(l.defaultValue=s.defaultValue);if(e)if(n)for(o=o||y(t),a=a||y(c),i=0,r=o.length;i<r;i++)d(o[i],a[i]);else d(t,c);return 0<(a=y(c,"script")).length&&m(a,!h&&y(t,"script")),c},buildFragment:function(t,e,n,i){for(var r,o,a,s,l,u,c=e.createDocumentFragment(),h=[],d=0,p=t.length;d<p;d++)if((r=t[d])||0===r)if("object"===U.type(r))U.merge(h,r.nodeType?[r]:r);else if(St.test(r)){for(o=o||c.appendChild(e.createElement("div")),a=(Et.exec(r)||["",""])[1].toLowerCase(),s=At[a]||At._default,o.innerHTML=s[1]+r.replace(Ct,"<$1></$2>")+s[2],u=s[0];u--;)o=o.lastChild;U.merge(h,o.childNodes),(o=c.firstChild).textContent=""}else h.push(e.createTextNode(r));for(c.textContent="",d=0;r=h[d++];)if((!i||-1===U.inArray(r,i))&&(l=U.contains(r.ownerDocument,r),o=y(c.appendChild(r),"script"),l&&m(o),n))for(u=0;r=o[u++];)_t.test(r.type||"")&&n.push(r);return c},cleanData:function(t){for(var e,n,i,r,o=U.event.special,a=0;void 0!==(n=t[a]);a++){if(U.acceptData(n)&&((r=n[ut.expando])&&(e=ut.cache[r]))){if(e.events)for(i in e.events)o[i]?U.event.remove(n,i):U.removeEvent(n,i,e.handle);ut.cache[r]&&delete ut.cache[r]}delete ct.cache[n[ct.expando]]}}}),U.fn.extend({text:function(t){return lt(this,function(t){return void 0===t?U.text(this):this.empty().each(function(){1!==this.nodeType&&11!==this.nodeType&&9!==this.nodeType||(this.textContent=t)})},null,t,arguments.length)},append:function(){return this.domManip(arguments,function(t){1!==this.nodeType&&11!==this.nodeType&&9!==this.nodeType||c(this,t).appendChild(t)})},prepend:function(){return this.domManip(arguments,function(t){if(1===this.nodeType||11===this.nodeType||9===this.nodeType){var e=c(this,t);e.insertBefore(t,e.firstChild)}})},before:function(){return this.domManip(arguments,function(t){this.parentNode&&this.parentNode.insertBefore(t,this)})},after:function(){return this.domManip(arguments,function(t){this.parentNode&&this.parentNode.insertBefore(t,this.nextSibling)})},remove:function(t,e){for(var n,i=t?U.filter(t,this):this,r=0;null!=(n=i[r]);r++)e||1!==n.nodeType||U.cleanData(y(n)),n.parentNode&&(e&&U.contains(n.ownerDocument,n)&&m(y(n,"script")),n.parentNode.removeChild(n));return this},empty:function(){for(var t,e=0;null!=(t=this[e]);e++)1===t.nodeType&&(U.cleanData(y(t,!1)),t.textContent="");return this},clone:function(t,e){return t=null!=t&&t,e=null==e?t:e,this.map(function(){return U.clone(this,t,e)})},html:function(t){return lt(this,function(t){var e=this[0]||{},n=0,i=this.length;if(void 0===t&&1===e.nodeType)return e.innerHTML;if("string"==typeof t&&!Tt.test(t)&&!At[(Et.exec(t)||["",""])[1].toLowerCase()]){t=t.replace(Ct,"<$1></$2>");try{for(;n<i;n++)1===(e=this[n]||{}).nodeType&&(U.cleanData(y(e,!1)),e.innerHTML=t);e=0}catch(t){}}e&&this.empty().append(t)},null,t,arguments.length)},replaceWith:function(){var e=arguments[0];return this.domManip(arguments,function(t){e=this.parentNode,U.cleanData(y(this)),e&&e.replaceChild(t,this)}),e&&(e.length||e.nodeType)?this:this.remove()},detach:function(t){return this.remove(t,!0)},domManip:function(n,i){n=R.apply([],n);var t,e,r,o,a,s,l=0,u=this.length,c=this,h=u-1,d=n[0],p=U.isFunction(d);if(p||1<u&&"string"==typeof d&&!X.checkClone&&Dt.test(d))return this.each(function(t){var e=c.eq(t);p&&(n[0]=d.call(this,t,e.html())),e.domManip(n,i)});if(u&&(e=(t=U.buildFragment(n,this[0].ownerDocument,!1,this)).firstChild,1===t.childNodes.length&&(t=e),e)){for(o=(r=U.map(y(t,"script"),g)).length;l<u;l++)a=t,l!==h&&(a=U.clone(a,!0,!0),o&&U.merge(r,y(a,"script"))),i.call(this[l],a,l);if(o)for(s=r[r.length-1].ownerDocument,U.map(r,v),l=0;l<o;l++)a=r[l],_t.test(a.type||"")&&!ut.access(a,"globalEval")&&U.contains(s,a)&&(a.src?U._evalUrl&&U._evalUrl(a.src):U.globalEval(a.textContent.replace(Mt,"")))}return this}}),U.each({appendTo:"append",prependTo:"prepend",insertBefore:"before",insertAfter:"after",replaceAll:"replaceWith"},function(t,a){U.fn[t]=function(t){for(var e,n=[],i=U(t),r=i.length-1,o=0;o<=r;o++)e=o===r?this:this.clone(!0),U(i[o])[a](e),F.apply(n,e.get());return this.pushStack(n)}});var Nt,Ot,It,Bt,Lt,jt,Rt={},Ft=/^margin/,Vt=new RegExp("^("+vt+")(?!px)[a-z%]+$","i"),qt=function(t){return t.ownerDocument.defaultView.opener?t.ownerDocument.defaultView.getComputedStyle(t,null):f.getComputedStyle(t,null)};function zt(){jt.style.cssText="-webkit-box-sizing:border-box;-moz-box-sizing:border-box;box-sizing:border-box;display:block;margin-top:1%;top:1%;border:1px;padding:1px;width:4px;position:absolute",jt.innerHTML="",Bt.appendChild(Lt);var t=f.getComputedStyle(jt,null);Ot="1%"!==t.top,It="4px"===t.width,Bt.removeChild(Lt)}Bt=Y.documentElement,Lt=Y.createElement("div"),(jt=Y.createElement("div")).style&&(jt.style.backgroundClip="content-box",jt.cloneNode(!0).style.backgroundClip="",X.clearCloneStyle="content-box"===jt.style.backgroundClip,Lt.style.cssText="border:0;width:0;height:0;top:0;left:-9999px;margin-top:1px;position:absolute",Lt.appendChild(jt),f.getComputedStyle&&U.extend(X,{pixelPosition:function(){return zt(),Ot},boxSizingReliable:function(){return null==It&&zt(),It},reliableMarginRight:function(){var t,e=jt.appendChild(Y.createElement("div"));return e.style.cssText=jt.style.cssText="-webkit-box-sizing:content-box;-moz-box-sizing:content-box;box-sizing:content-box;display:block;margin:0;border:0;padding:0",e.style.marginRight=e.style.width="0",jt.style.width="1px",Bt.appendChild(Lt),t=!parseFloat(f.getComputedStyle(e,null).marginRight),Bt.removeChild(Lt),jt.removeChild(e),t}})),U.swap=function(t,e,n,i){var r,o,a={};for(o in e)a[o]=t.style[o],t.style[o]=e[o];for(o in r=n.apply(t,i||[]),e)t.style[o]=a[o];return r};var Ht=/^(none|table(?!-c[ea]).+)/,Xt=new RegExp("^("+vt+")(.*)$","i"),Yt=new RegExp("^([+-])=("+vt+")","i"),Ut={position:"absolute",visibility:"hidden",display:"block"},Wt={letterSpacing:"0",fontWeight:"400"},Gt=["Webkit","O","Moz","ms"];U.extend({cssHooks:{opacity:{get:function(t,e){if(e){var n=p(t,"opacity");return""===n?"1":n}}}},cssNumber:{columnCount:!0,fillOpacity:!0,flexGrow:!0,flexShrink:!0,fontWeight:!0,lineHeight:!0,opacity:!0,order:!0,orphans:!0,widows:!0,zIndex:!0,zoom:!0},cssProps:{float:"cssFloat"},style:function(t,e,n,i){if(t&&3!==t.nodeType&&8!==t.nodeType&&t.style){var r,o,a,s=U.camelCase(e),l=t.style;return e=U.cssProps[s]||(U.cssProps[s]=w(l,s)),a=U.cssHooks[e]||U.cssHooks[s],void 0===n?a&&"get"in a&&void 0!==(r=a.get(t,!1,i))?r:l[e]:("string"===(o=typeof n)&&(r=Yt.exec(n))&&(n=(r[1]+1)*r[2]+parseFloat(U.css(t,e)),o="number"),void(null!=n&&n==n&&("number"!==o||U.cssNumber[s]||(n+="px"),X.clearCloneStyle||""!==n||0!==e.indexOf("background")||(l[e]="inherit"),a&&"set"in a&&void 0===(n=a.set(t,n,i))||(l[e]=n))))}},css:function(t,e,n,i){var r,o,a,s=U.camelCase(e);return e=U.cssProps[s]||(U.cssProps[s]=w(t.style,s)),(a=U.cssHooks[e]||U.cssHooks[s])&&"get"in a&&(r=a.get(t,!0,n)),void 0===r&&(r=p(t,e,i)),"normal"===r&&e in Wt&&(r=Wt[e]),""===n||n?(o=parseFloat(r),!0===n||U.isNumeric(o)?o||0:r):r}}),U.each(["height","width"],function(t,r){U.cssHooks[r]={get:function(t,e,n){return e?Ht.test(U.css(t,"display"))&&0===t.offsetWidth?U.swap(t,Ut,function(){return C(t,r,n)}):C(t,r,n):void 0},set:function(t,e,n){var i=n&&qt(t);return $(0,e,n?k(t,r,n,"border-box"===U.css(t,"boxSizing",!1,i),i):0)}}}),U.cssHooks.marginRight=x(X.reliableMarginRight,function(t,e){return e?U.swap(t,{display:"inline-block"},p,[t,"marginRight"]):void 0}),U.each({margin:"",padding:"",border:"Width"},function(r,o){U.cssHooks[r+o]={expand:function(t){for(var e=0,n={},i="string"==typeof t?t.split(" "):[t];e<4;e++)n[r+mt[e]+o]=i[e]||i[e-2]||i[0];return n}},Ft.test(r)||(U.cssHooks[r+o].set=$)}),U.fn.extend({css:function(t,e){return lt(this,function(t,e,n){var i,r,o={},a=0;if(U.isArray(e)){for(i=qt(t),r=e.length;a<r;a++)o[e[a]]=U.css(t,e[a],!1,i);return o}return void 0!==n?U.style(t,e,n):U.css(t,e)},t,e,1<arguments.length)},show:function(){return E(this,!0)},hide:function(){return E(this)},toggle:function(t){return"boolean"==typeof t?t?this.show():this.hide():this.each(function(){pt(this)?U(this).show():U(this).hide()})}}),((U.Tween=S).prototype={constructor:S,init:function(t,e,n,i,r,o){this.elem=t,this.prop=n,this.easing=r||"swing",this.options=e,this.start=this.now=this.cur(),this.end=i,this.unit=o||(U.cssNumber[n]?"":"px")},cur:function(){var t=S.propHooks[this.prop];return t&&t.get?t.get(this):S.propHooks._default.get(this)},run:function(t){var e,n=S.propHooks[this.prop];return this.options.duration?this.pos=e=U.easing[this.easing](t,this.options.duration*t,0,1,this.options.duration):this.pos=e=t,this.now=(this.end-this.start)*e+this.start,this.options.step&&this.options.step.call(this.elem,this.now,this),n&&n.set?n.set(this):S.propHooks._default.set(this),this}}).init.prototype=S.prototype,(S.propHooks={_default:{get:function(t){var e;return null==t.elem[t.prop]||t.elem.style&&null!=t.elem.style[t.prop]?(e=U.css(t.elem,t.prop,""))&&"auto"!==e?e:0:t.elem[t.prop]},set:function(t){U.fx.step[t.prop]?U.fx.step[t.prop](t):t.elem.style&&(null!=t.elem.style[U.cssProps[t.prop]]||U.cssHooks[t.prop])?U.style(t.elem,t.prop,t.now+t.unit):t.elem[t.prop]=t.now}}}).scrollTop=S.propHooks.scrollLeft={set:function(t){t.elem.nodeType&&t.elem.parentNode&&(t.elem[t.prop]=t.now)}},U.easing={linear:function(t){return t},swing:function(t){return.5-Math.cos(t*Math.PI)/2}},U.fx=S.prototype.init,U.fx.step={};var Zt,Kt,Qt,Jt,te,ee=/^(?:toggle|show|hide)$/,ne=new RegExp("^(?:([+-])=|)("+vt+")([a-z%]*)$","i"),ie=/queueHooks$/,re=[function(e,t,n){var i,r,o,a,s,l,u,c=this,h={},d=e.style,p=e.nodeType&&pt(e),f=ut.get(e,"fxshow");for(i in n.queue||(null==(s=U._queueHooks(e,"fx")).unqueued&&(s.unqueued=0,l=s.empty.fire,s.empty.fire=function(){s.unqueued||l()}),s.unqueued++,c.always(function(){c.always(function(){s.unqueued--,U.queue(e,"fx").length||s.empty.fire()})})),1===e.nodeType&&("height"in t||"width"in t)&&(n.overflow=[d.overflow,d.overflowX,d.overflowY],"inline"===("none"===(u=U.css(e,"display"))?ut.get(e,"olddisplay")||b(e.nodeName):u)&&"none"===U.css(e,"float")&&(d.display="inline-block")),n.overflow&&(d.overflow="hidden",c.always(function(){d.overflow=n.overflow[0],d.overflowX=n.overflow[1],d.overflowY=n.overflow[2]})),t)if(r=t[i],ee.exec(r)){if(delete t[i],o=o||"toggle"===r,r===(p?"hide":"show")){if("show"!==r||!f||void 0===f[i])continue;p=!0}h[i]=f&&f[i]||U.style(e,i)}else u=void 0;if(U.isEmptyObject(h))"inline"===("none"===u?b(e.nodeName):u)&&(d.display=u);else for(i in f?"hidden"in f&&(p=f.hidden):f=ut.access(e,"fxshow",{}),o&&(f.hidden=!p),p?U(e).show():c.done(function(){U(e).hide()}),c.done(function(){var t;for(t in ut.remove(e,"fxshow"),h)U.style(e,t,h[t])}),h)a=_(p?f[i]:0,i,c),i in f||(f[i]=a.start,p&&(a.end=a.start,a.start="width"===i||"height"===i?1:0))}],oe={"*":[function(t,e){var n=this.createTween(t,e),i=n.cur(),r=ne.exec(e),o=r&&r[3]||(U.cssNumber[t]?"":"px"),a=(U.cssNumber[t]||"px"!==o&&+i)&&ne.exec(U.css(n.elem,t)),s=1,l=20;if(a&&a[3]!==o)for(o=o||a[3],r=r||[],a=+i||1;a/=s=s||".5",U.style(n.elem,t,a+o),s!==(s=n.cur()/i)&&1!==s&&--l;);return r&&(a=n.start=+a||+i||0,n.unit=o,n.end=r[1]?a+(r[1]+1)*r[2]:+r[2]),n}]};U.Animation=U.extend(P,{tweener:function(t,e){for(var n,i=0,r=(t=U.isFunction(t)?(e=t,["*"]):t.split(" ")).length;i<r;i++)n=t[i],oe[n]=oe[n]||[],oe[n].unshift(e)},prefilter:function(t,e){e?re.unshift(t):re.push(t)}}),U.speed=function(t,e,n){var i=t&&"object"==typeof t?U.extend({},t):{complete:n||!n&&e||U.isFunction(t)&&t,duration:t,easing:n&&e||e&&!U.isFunction(e)&&e};return i.duration=U.fx.off?0:"number"==typeof i.duration?i.duration:i.duration in U.fx.speeds?U.fx.speeds[i.duration]:U.fx.speeds._default,null!=i.queue&&!0!==i.queue||(i.queue="fx"),i.old=i.complete,i.complete=function(){U.isFunction(i.old)&&i.old.call(this),i.queue&&U.dequeue(this,i.queue)},i},U.fn.extend({fadeTo:function(t,e,n,i){return this.filter(pt).css("opacity",0).show().end().animate({opacity:e},t,n,i)},animate:function(e,t,n,i){function r(){var t=P(this,U.extend({},e),a);(o||ut.get(this,"finish"))&&t.stop(!0)}var o=U.isEmptyObject(e),a=U.speed(t,n,i);return r.finish=r,o||!1===a.queue?this.each(r):this.queue(a.queue,r)},stop:function(r,t,o){function a(t){var e=t.stop;delete t.stop,e(o)}return"string"!=typeof r&&(o=t,t=r,r=void 0),t&&!1!==r&&this.queue(r||"fx",[]),this.each(function(){var t=!0,e=null!=r&&r+"queueHooks",n=U.timers,i=ut.get(this);if(e)i[e]&&i[e].stop&&a(i[e]);else for(e in i)i[e]&&i[e].stop&&ie.test(e)&&a(i[e]);for(e=n.length;e--;)n[e].elem!==this||null!=r&&n[e].queue!==r||(n[e].anim.stop(o),t=!1,n.splice(e,1));!t&&o||U.dequeue(this,r)})},finish:function(a){return!1!==a&&(a=a||"fx"),this.each(function(){var t,e=ut.get(this),n=e[a+"queue"],i=e[a+"queueHooks"],r=U.timers,o=n?n.length:0;for(e.finish=!0,U.queue(this,a,[]),i&&i.stop&&i.stop.call(this,!0),t=r.length;t--;)r[t].elem===this&&r[t].queue===a&&(r[t].anim.stop(!0),r.splice(t,1));for(t=0;t<o;t++)n[t]&&n[t].finish&&n[t].finish.call(this);delete e.finish})}}),U.each(["toggle","show","hide"],function(t,i){var r=U.fn[i];U.fn[i]=function(t,e,n){return null==t||"boolean"==typeof t?r.apply(this,arguments):this.animate(D(i,!0),t,e,n)}}),U.each({slideDown:D("show"),slideUp:D("hide"),slideToggle:D("toggle"),fadeIn:{opacity:"show"},fadeOut:{opacity:"hide"},fadeToggle:{opacity:"toggle"}},function(t,i){U.fn[t]=function(t,e,n){return this.animate(i,t,e,n)}}),U.timers=[],U.fx.tick=function(){var t,e=0,n=U.timers;for(Zt=U.now();e<n.length;e++)(t=n[e])()||n[e]!==t||n.splice(e--,1);n.length||U.fx.stop(),Zt=void 0},U.fx.timer=function(t){U.timers.push(t),t()?U.fx.start():U.timers.pop()},U.fx.interval=13,U.fx.start=function(){Kt||(Kt=setInterval(U.fx.tick,U.fx.interval))},U.fx.stop=function(){clearInterval(Kt),Kt=null},U.fx.speeds={slow:600,fast:200,_default:400},U.fn.delay=function(i,t){return i=U.fx&&U.fx.speeds[i]||i,t=t||"fx",this.queue(t,function(t,e){var n=setTimeout(t,i);e.stop=function(){clearTimeout(n)}})},Qt=Y.createElement("input"),Jt=Y.createElement("select"),te=Jt.appendChild(Y.createElement("option")),Qt.type="checkbox",X.checkOn=""!==Qt.value,X.optSelected=te.selected,Jt.disabled=!0,X.optDisabled=!te.disabled,(Qt=Y.createElement("input")).value="t",Qt.type="radio",X.radioValue="t"===Qt.value;var ae,se=U.expr.attrHandle;U.fn.extend({attr:function(t,e){return lt(this,U.attr,t,e,1<arguments.length)},removeAttr:function(t){return this.each(function(){U.removeAttr(this,t)})}}),U.extend({attr:function(t,e,n){var i,r,o=t.nodeType;if(t&&3!==o&&8!==o&&2!==o)return typeof t.getAttribute==bt?U.prop(t,e,n):(1===o&&U.isXMLDoc(t)||(e=e.toLowerCase(),i=U.attrHooks[e]||(U.expr.match.bool.test(e)?ae:void 0)),void 0===n?i&&"get"in i&&null!==(r=i.get(t,e))?r:null==(r=U.find.attr(t,e))?void 0:r:null!==n?i&&"set"in i&&void 0!==(r=i.set(t,n,e))?r:(t.setAttribute(e,n+""),n):void U.removeAttr(t,e))},removeAttr:function(t,e){var n,i,r=0,o=e&&e.match(at);if(o&&1===t.nodeType)for(;n=o[r++];)i=U.propFix[n]||n,U.expr.match.bool.test(n)&&(t[i]=!1),t.removeAttribute(n)},attrHooks:{type:{set:function(t,e){if(!X.radioValue&&"radio"===e&&U.nodeName(t,"input")){var n=t.value;return t.setAttribute("type",e),n&&(t.value=n),e}}}}}),ae={set:function(t,e,n){return!1===e?U.removeAttr(t,n):t.setAttribute(n,n),n}},U.each(U.expr.match.bool.source.match(/\w+/g),function(t,e){var o=se[e]||U.find.attr;se[e]=function(t,e,n){var i,r;return n||(r=se[e],se[e]=i,i=null!=o(t,e,n)?e.toLowerCase():null,se[e]=r),i}});var le=/^(?:input|select|textarea|button)$/i;U.fn.extend({prop:function(t,e){return lt(this,U.prop,t,e,1<arguments.length)},removeProp:function(t){return this.each(function(){delete this[U.propFix[t]||t]})}}),U.extend({propFix:{for:"htmlFor",class:"className"},prop:function(t,e,n){var i,r,o=t.nodeType;if(t&&3!==o&&8!==o&&2!==o)return(1!==o||!U.isXMLDoc(t))&&(e=U.propFix[e]||e,r=U.propHooks[e]),void 0!==n?r&&"set"in r&&void 0!==(i=r.set(t,n,e))?i:t[e]=n:r&&"get"in r&&null!==(i=r.get(t,e))?i:t[e]},propHooks:{tabIndex:{get:function(t){return t.hasAttribute("tabindex")||le.test(t.nodeName)||t.href?t.tabIndex:-1}}}}),X.optSelected||(U.propHooks.selected={get:function(t){var e=t.parentNode;return e&&e.parentNode&&e.parentNode.selectedIndex,null}}),U.each(["tabIndex","readOnly","maxLength","cellSpacing","cellPadding","rowSpan","colSpan","useMap","frameBorder","contentEditable"],function(){U.propFix[this.toLowerCase()]=this});var ue=/[\t\r\n\f]/g;U.fn.extend({addClass:function(e){var t,n,i,r,o,a,s="string"==typeof e&&e,l=0,u=this.length;if(U.isFunction(e))return this.each(function(t){U(this).addClass(e.call(this,t,this.className))});if(s)for(t=(e||"").match(at)||[];l<u;l++)if(i=1===(n=this[l]).nodeType&&(n.className?(" "+n.className+" ").replace(ue," "):" ")){for(o=0;r=t[o++];)i.indexOf(" "+r+" ")<0&&(i+=r+" ");a=U.trim(i),n.className!==a&&(n.className=a)}return this},removeClass:function(e){var t,n,i,r,o,a,s=0===arguments.length||"string"==typeof e&&e,l=0,u=this.length;if(U.isFunction(e))return this.each(function(t){U(this).removeClass(e.call(this,t,this.className))});if(s)for(t=(e||"").match(at)||[];l<u;l++)if(i=1===(n=this[l]).nodeType&&(n.className?(" "+n.className+" ").replace(ue," "):"")){for(o=0;r=t[o++];)for(;0<=i.indexOf(" "+r+" ");)i=i.replace(" "+r+" "," ");a=e?U.trim(i):"",n.className!==a&&(n.className=a)}return this},toggleClass:function(r,e){var o=typeof r;return"boolean"==typeof e&&"string"==o?e?this.addClass(r):this.removeClass(r):U.isFunction(r)?this.each(function(t){U(this).toggleClass(r.call(this,t,this.className,e),e)}):this.each(function(){if("string"==o)for(var t,e=0,n=U(this),i=r.match(at)||[];t=i[e++];)n.hasClass(t)?n.removeClass(t):n.addClass(t);else o!=bt&&"boolean"!=o||(this.className&&ut.set(this,"__className__",this.className),this.className=this.className||!1===r?"":ut.get(this,"__className__")||"")})},hasClass:function(t){for(var e=" "+t+" ",n=0,i=this.length;n<i;n++)if(1===this[n].nodeType&&0<=(" "+this[n].className+" ").replace(ue," ").indexOf(e))return!0;return!1}});var ce=/\r/g;U.fn.extend({val:function(n){var i,t,r,e=this[0];return arguments.length?(r=U.isFunction(n),this.each(function(t){var e;1===this.nodeType&&(null==(e=r?n.call(this,t,U(this).val()):n)?e="":"number"==typeof e?e+="":U.isArray(e)&&(e=U.map(e,function(t){return null==t?"":t+""})),(i=U.valHooks[this.type]||U.valHooks[this.nodeName.toLowerCase()])&&"set"in i&&void 0!==i.set(this,e,"value")||(this.value=e))})):e?(i=U.valHooks[e.type]||U.valHooks[e.nodeName.toLowerCase()])&&"get"in i&&void 0!==(t=i.get(e,"value"))?t:"string"==typeof(t=e.value)?t.replace(ce,""):null==t?"":t:void 0}}),U.extend({valHooks:{option:{get:function(t){var e=U.find.attr(t,"value");return null!=e?e:U.trim(U.text(t))}},select:{get:function(t){for(var e,n,i=t.options,r=t.selectedIndex,o="select-one"===t.type||r<0,a=o?null:[],s=o?r+1:i.length,l=r<0?s:o?r:0;l<s;l++)if(((n=i[l]).selected||l===r)&&(X.optDisabled?!n.disabled:null===n.getAttribute("disabled"))&&(!n.parentNode.disabled||!U.nodeName(n.parentNode,"optgroup"))){if(e=U(n).val(),o)return e;a.push(e)}return a},set:function(t,e){for(var n,i,r=t.options,o=U.makeArray(e),a=r.length;a--;)((i=r[a]).selected=0<=U.inArray(i.value,o))&&(n=!0);return n||(t.selectedIndex=-1),o}}}}),U.each(["radio","checkbox"],function(){U.valHooks[this]={set:function(t,e){return U.isArray(e)?t.checked=0<=U.inArray(U(t).val(),e):void 0}},X.checkOn||(U.valHooks[this].get=function(t){return null===t.getAttribute("value")?"on":t.value})}),U.each("blur focus focusin focusout load resize scroll unload click dblclick mousedown mouseup mousemove mouseover mouseout mouseenter mouseleave change select submit keydown keypress keyup error contextmenu".split(" "),function(t,n){U.fn[n]=function(t,e){return 0<arguments.length?this.on(n,null,t,e):this.trigger(n)}}),U.fn.extend({hover:function(t,e){return this.mouseenter(t).mouseleave(e||t)},bind:function(t,e,n){return this.on(t,null,e,n)},unbind:function(t,e){return this.off(t,null,e)},delegate:function(t,e,n,i){return this.on(e,t,n,i)},undelegate:function(t,e,n){return 1===arguments.length?this.off(t,"**"):this.off(e,t||"**",n)}});var he=U.now(),de=/\?/;U.parseJSON=function(t){return JSON.parse(t+"")},U.parseXML=function(t){var e;if(!t||"string"!=typeof t)return null;try{e=(new DOMParser).parseFromString(t,"text/xml")}catch(t){e=void 0}return e&&!e.getElementsByTagName("parsererror").length||U.error("Invalid XML: "+t),e};var pe=/#.*$/,fe=/([?&])_=[^&]*/,ge=/^(.*?):[ \t]*([^\r\n]*)$/gm,ve=/^(?:GET|HEAD)$/,me=/^\/\//,ye=/^([\w.+-]+:)(?:\/\/(?:[^\/?#]*@|)([^\/?#:]*)(?::(\d+)|)|)/,be={},xe={},we="*/".concat("*"),$e=f.location.href,ke=ye.exec($e.toLowerCase())||[];U.extend({active:0,lastModified:{},etag:{},ajaxSettings:{url:$e,type:"GET",isLocal:/^(?:about|app|app-storage|.+-extension|file|res|widget):$/.test(ke[1]),global:!0,processData:!0,async:!0,contentType:"application/x-www-form-urlencoded; charset=UTF-8",accepts:{"*":we,text:"text/plain",html:"text/html",xml:"application/xml, text/xml",json:"application/json, text/javascript"},contents:{xml:/xml/,html:/html/,json:/json/},responseFields:{xml:"responseXML",text:"responseText",json:"responseJSON"},converters:{"* text":String,"text html":!0,"text json":U.parseJSON,"text xml":U.parseXML},flatOptions:{url:!0,context:!0}},ajaxSetup:function(t,e){return e?N(N(t,U.ajaxSettings),e):N(U.ajaxSettings,t)},ajaxPrefilter:M(be),ajaxTransport:M(xe),ajax:function(t,e){function n(t,e,n,i){var r,o,a,s,l,u=e;2!==w&&(w=2,p&&clearTimeout(p),c=void 0,d=i||"",$.readyState=0<t?4:0,r=200<=t&&t<300||304===t,n&&(s=function(t,e,n){for(var i,r,o,a,s=t.contents,l=t.dataTypes;"*"===l[0];)l.shift(),void 0===i&&(i=t.mimeType||e.getResponseHeader("Content-Type"));if(i)for(r in s)if(s[r]&&s[r].test(i)){l.unshift(r);break}if(l[0]in n)o=l[0];else{for(r in n){if(!l[0]||t.converters[r+" "+l[0]]){o=r;break}a||(a=r)}o=o||a}return o?(o!==l[0]&&l.unshift(o),n[o]):void 0}(g,$,n)),s=function(t,e,n,i){var r,o,a,s,l,u={},c=t.dataTypes.slice();if(c[1])for(a in t.converters)u[a.toLowerCase()]=t.converters[a];for(o=c.shift();o;)if(t.responseFields[o]&&(n[t.responseFields[o]]=e),!l&&i&&t.dataFilter&&(e=t.dataFilter(e,t.dataType)),l=o,o=c.shift())if("*"===o)o=l;else if("*"!==l&&l!==o){if(!(a=u[l+" "+o]||u["* "+o]))for(r in u)if((s=r.split(" "))[1]===o&&(a=u[l+" "+s[0]]||u["* "+s[0]])){!0===a?a=u[r]:!0!==u[r]&&(o=s[0],c.unshift(s[1]));break}if(!0!==a)if(a&&t.throws)e=a(e);else try{e=a(e)}catch(t){return{state:"parsererror",error:a?t:"No conversion from "+l+" to "+o}}}return{state:"success",data:e}}(g,s,$,r),r?(g.ifModified&&((l=$.getResponseHeader("Last-Modified"))&&(U.lastModified[h]=l),(l=$.getResponseHeader("etag"))&&(U.etag[h]=l)),204===t||"HEAD"===g.type?u="nocontent":304===t?u="notmodified":(u=s.state,o=s.data,r=!(a=s.error))):(a=u,!t&&u||(u="error",t<0&&(t=0))),$.status=t,$.statusText=(e||u)+"",r?y.resolveWith(v,[o,u,$]):y.rejectWith(v,[$,u,a]),$.statusCode(x),x=void 0,f&&m.trigger(r?"ajaxSuccess":"ajaxError",[$,g,r?o:a]),b.fireWith(v,[$,u]),f&&(m.trigger("ajaxComplete",[$,g]),--U.active||U.event.trigger("ajaxStop")))}"object"==typeof t&&(e=t,t=void 0),e=e||{};var c,h,d,i,p,r,f,o,g=U.ajaxSetup({},e),v=g.context||g,m=g.context&&(v.nodeType||v.jquery)?U(v):U.event,y=U.Deferred(),b=U.Callbacks("once memory"),x=g.statusCode||{},a={},s={},w=0,l="canceled",$={readyState:0,getResponseHeader:function(t){var e;if(2===w){if(!i)for(i={};e=ge.exec(d);)i[e[1].toLowerCase()]=e[2];e=i[t.toLowerCase()]}return null==e?null:e},getAllResponseHeaders:function(){return 2===w?d:null},setRequestHeader:function(t,e){var n=t.toLowerCase();return w||(t=s[n]=s[n]||t,a[t]=e),this},overrideMimeType:function(t){return w||(g.mimeType=t),this},statusCode:function(t){var e;if(t)if(w<2)for(e in t)x[e]=[x[e],t[e]];else $.always(t[$.status]);return this},abort:function(t){var e=t||l;return c&&c.abort(e),n(0,e),this}};if(y.promise($).complete=b.add,$.success=$.done,$.error=$.fail,g.url=((t||g.url||$e)+"").replace(pe,"").replace(me,ke[1]+"//"),g.type=e.method||e.type||g.method||g.type,g.dataTypes=U.trim(g.dataType||"*").toLowerCase().match(at)||[""],null==g.crossDomain&&(r=ye.exec(g.url.toLowerCase()),g.crossDomain=!(!r||r[1]===ke[1]&&r[2]===ke[2]&&(r[3]||("http:"===r[1]?"80":"443"))===(ke[3]||("http:"===ke[1]?"80":"443")))),g.data&&g.processData&&"string"!=typeof g.data&&(g.data=U.param(g.data,g.traditional)),A(be,g,e,$),2===w)return $;for(o in(f=U.event&&g.global)&&0==U.active++&&U.event.trigger("ajaxStart"),g.type=g.type.toUpperCase(),g.hasContent=!ve.test(g.type),h=g.url,g.hasContent||(g.data&&(h=g.url+=(de.test(h)?"&":"?")+g.data,delete g.data),!1===g.cache&&(g.url=fe.test(h)?h.replace(fe,"$1_="+he++):h+(de.test(h)?"&":"?")+"_="+he++)),g.ifModified&&(U.lastModified[h]&&$.setRequestHeader("If-Modified-Since",U.lastModified[h]),U.etag[h]&&$.setRequestHeader("If-None-Match",U.etag[h])),(g.data&&g.hasContent&&!1!==g.contentType||e.contentType)&&$.setRequestHeader("Content-Type",g.contentType),$.setRequestHeader("Accept",g.dataTypes[0]&&g.accepts[g.dataTypes[0]]?g.accepts[g.dataTypes[0]]+("*"!==g.dataTypes[0]?", "+we+"; q=0.01":""):g.accepts["*"]),g.headers)$.setRequestHeader(o,g.headers[o]);if(g.beforeSend&&(!1===g.beforeSend.call(v,$,g)||2===w))return $.abort();for(o in l="abort",{success:1,error:1,complete:1})$[o](g[o]);if(c=A(xe,g,e,$)){$.readyState=1,f&&m.trigger("ajaxSend",[$,g]),g.async&&0<g.timeout&&(p=setTimeout(function(){$.abort("timeout")},g.timeout));try{w=1,c.send(a,n)}catch(t){if(!(w<2))throw t;n(-1,t)}}else n(-1,"No Transport");return $},getJSON:function(t,e,n){return U.get(t,e,n,"json")},getScript:function(t,e){return U.get(t,void 0,e,"script")}}),U.each(["get","post"],function(t,r){U[r]=function(t,e,n,i){return U.isFunction(e)&&(i=i||n,n=e,e=void 0),U.ajax({url:t,type:r,dataType:i,data:e,success:n})}}),U._evalUrl=function(t){return U.ajax({url:t,type:"GET",dataType:"script",async:!1,global:!1,throws:!0})},U.fn.extend({wrapAll:function(e){var t;return U.isFunction(e)?this.each(function(t){U(this).wrapAll(e.call(this,t))}):(this[0]&&(t=U(e,this[0].ownerDocument).eq(0).clone(!0),this[0].parentNode&&t.insertBefore(this[0]),t.map(function(){for(var t=this;t.firstElementChild;)t=t.firstElementChild;return t}).append(this)),this)},wrapInner:function(n){return U.isFunction(n)?this.each(function(t){U(this).wrapInner(n.call(this,t))}):this.each(function(){var t=U(this),e=t.contents();e.length?e.wrapAll(n):t.append(n)})},wrap:function(e){var n=U.isFunction(e);return this.each(function(t){U(this).wrapAll(n?e.call(this,t):e)})},unwrap:function(){return this.parent().each(function(){U.nodeName(this,"body")||U(this).replaceWith(this.childNodes)}).end()}}),U.expr.filters.hidden=function(t){return t.offsetWidth<=0&&t.offsetHeight<=0},U.expr.filters.visible=function(t){return!U.expr.filters.hidden(t)};var Ce=/%20/g,Ee=/\[\]$/,Se=/\r?\n/g,Te=/^(?:submit|button|image|reset|file)$/i,De=/^(?:input|select|textarea|keygen)/i;U.param=function(t,e){function n(t,e){e=U.isFunction(e)?e():null==e?"":e,r[r.length]=encodeURIComponent(t)+"="+encodeURIComponent(e)}var i,r=[];if(void 0===e&&(e=U.ajaxSettings&&U.ajaxSettings.traditional),U.isArray(t)||t.jquery&&!U.isPlainObject(t))U.each(t,function(){n(this.name,this.value)});else for(i in t)O(i,t[i],e,n);return r.join("&").replace(Ce,"+")},U.fn.extend({serialize:function(){return U.param(this.serializeArray())},serializeArray:function(){return this.map(function(){var t=U.prop(this,"elements");return t?U.makeArray(t):this}).filter(function(){var t=this.type;return this.name&&!U(this).is(":disabled")&&De.test(this.nodeName)&&!Te.test(t)&&(this.checked||!yt.test(t))}).map(function(t,e){var n=U(this).val();return null==n?null:U.isArray(n)?U.map(n,function(t){return{name:e.name,value:t.replace(Se,"\r\n")}}):{name:e.name,value:n.replace(Se,"\r\n")}}).get()}}),U.ajaxSettings.xhr=function(){try{return new XMLHttpRequest}catch(t){}};var _e=0,Pe={},Me={0:200,1223:204},Ae=U.ajaxSettings.xhr();f.attachEvent&&f.attachEvent("onunload",function(){for(var t in Pe)Pe[t]()}),X.cors=!!Ae&&"withCredentials"in Ae,X.ajax=Ae=!!Ae,U.ajaxTransport(function(o){var a;return X.cors||Ae&&!o.crossDomain?{send:function(t,e){var n,i=o.xhr(),r=++_e;if(i.open(o.type,o.url,o.async,o.username,o.password),o.xhrFields)for(n in o.xhrFields)i[n]=o.xhrFields[n];for(n in o.mimeType&&i.overrideMimeType&&i.overrideMimeType(o.mimeType),o.crossDomain||t["X-Requested-With"]||(t["X-Requested-With"]="XMLHttpRequest"),t)i.setRequestHeader(n,t[n]);a=function(t){return function(){a&&(delete Pe[r],a=i.onload=i.onerror=null,"abort"===t?i.abort():"error"===t?e(i.status,i.statusText):e(Me[i.status]||i.status,i.statusText,"string"==typeof i.responseText?{text:i.responseText}:void 0,i.getAllResponseHeaders()))}},i.onload=a(),i.onerror=a("error"),a=Pe[r]=a("abort");try{i.send(o.hasContent&&o.data||null)}catch(t){if(a)throw t}},abort:function(){a&&a()}}:void 0}),U.ajaxSetup({accepts:{script:"text/javascript, application/javascript, application/ecmascript, application/x-ecmascript"},contents:{script:/(?:java|ecma)script/},converters:{"text script":function(t){return U.globalEval(t),t}}}),U.ajaxPrefilter("script",function(t){void 0===t.cache&&(t.cache=!1),t.crossDomain&&(t.type="GET")}),U.ajaxTransport("script",function(n){var i,r;if(n.crossDomain)return{send:function(t,e){i=U("<script>").prop({async:!0,charset:n.scriptCharset,src:n.url}).on("load error",r=function(t){i.remove(),r=null,t&&e("error"===t.type?404:200,t.type)}),Y.head.appendChild(i[0])},abort:function(){r&&r()}}});var Ne=[],Oe=/(=)\?(?=&|$)|\?\?/;U.ajaxSetup({jsonp:"callback",jsonpCallback:function(){var t=Ne.pop()||U.expando+"_"+he++;return this[t]=!0,t}}),U.ajaxPrefilter("json jsonp",function(t,e,n){var i,r,o,a=!1!==t.jsonp&&(Oe.test(t.url)?"url":"string"==typeof t.data&&!(t.contentType||"").indexOf("application/x-www-form-urlencoded")&&Oe.test(t.data)&&"data");return a||"jsonp"===t.dataTypes[0]?(i=t.jsonpCallback=U.isFunction(t.jsonpCallback)?t.jsonpCallback():t.jsonpCallback,a?t[a]=t[a].replace(Oe,"$1"+i):!1!==t.jsonp&&(t.url+=(de.test(t.url)?"&":"?")+t.jsonp+"="+i),t.converters["script json"]=function(){return o||U.error(i+" was not called"),o[0]},t.dataTypes[0]="json",r=f[i],f[i]=function(){o=arguments},n.always(function(){f[i]=r,t[i]&&(t.jsonpCallback=e.jsonpCallback,Ne.push(i)),o&&U.isFunction(r)&&r(o[0]),o=r=void 0}),"script"):void 0}),U.parseHTML=function(t,e,n){if(!t||"string"!=typeof t)return null;"boolean"==typeof e&&(n=e,e=!1),e=e||Y;var i=J.exec(t),r=!n&&[];return i?[e.createElement(i[1])]:(i=U.buildFragment([t],e,r),r&&r.length&&U(r).remove(),U.merge([],i.childNodes))};var Ie=U.fn.load;U.fn.load=function(t,e,n){if("string"!=typeof t&&Ie)return Ie.apply(this,arguments);var i,r,o,a=this,s=t.indexOf(" ");return 0<=s&&(i=U.trim(t.slice(s)),t=t.slice(0,s)),U.isFunction(e)?(n=e,e=void 0):e&&"object"==typeof e&&(r="POST"),0<a.length&&U.ajax({url:t,type:r,dataType:"html",data:e}).done(function(t){o=arguments,a.html(i?U("<div>").append(U.parseHTML(t)).find(i):t)}).complete(n&&function(t,e){a.each(n,o||[t.responseText,e,t])}),this},U.each(["ajaxStart","ajaxStop","ajaxComplete","ajaxError","ajaxSuccess","ajaxSend"],function(t,e){U.fn[e]=function(t){return this.on(e,t)}}),U.expr.filters.animated=function(e){return U.grep(U.timers,function(t){return e===t.elem}).length};var Be=f.document.documentElement;U.offset={setOffset:function(t,e,n){var i,r,o,a,s,l,u=U.css(t,"position"),c=U(t),h={};"static"===u&&(t.style.position="relative"),s=c.offset(),o=U.css(t,"top"),l=U.css(t,"left"),r=("absolute"===u||"fixed"===u)&&-1<(o+l).indexOf("auto")?(a=(i=c.position()).top,i.left):(a=parseFloat(o)||0,parseFloat(l)||0),U.isFunction(e)&&(e=e.call(t,n,s)),null!=e.top&&(h.top=e.top-s.top+a),null!=e.left&&(h.left=e.left-s.left+r),"using"in e?e.using.call(t,h):c.css(h)}},U.fn.extend({offset:function(e){if(arguments.length)return void 0===e?this:this.each(function(t){U.offset.setOffset(this,e,t)});var t,n,i=this[0],r={top:0,left:0},o=i&&i.ownerDocument;return o?(t=o.documentElement,U.contains(t,i)?(typeof i.getBoundingClientRect!=bt&&(r=i.getBoundingClientRect()),n=I(o),{top:r.top+n.pageYOffset-t.clientTop,left:r.left+n.pageXOffset-t.clientLeft}):r):void 0},position:function(){if(this[0]){var t,e,n=this[0],i={top:0,left:0};return"fixed"===U.css(n,"position")?e=n.getBoundingClientRect():(t=this.offsetParent(),e=this.offset(),U.nodeName(t[0],"html")||(i=t.offset()),i.top+=U.css(t[0],"borderTopWidth",!0),i.left+=U.css(t[0],"borderLeftWidth",!0)),{top:e.top-i.top-U.css(n,"marginTop",!0),left:e.left-i.left-U.css(n,"marginLeft",!0)}}},offsetParent:function(){return this.map(function(){for(var t=this.offsetParent||Be;t&&!U.nodeName(t,"html")&&"static"===U.css(t,"position");)t=t.offsetParent;return t||Be})}}),U.each({scrollLeft:"pageXOffset",scrollTop:"pageYOffset"},function(e,r){var o="pageYOffset"===r;U.fn[e]=function(t){return lt(this,function(t,e,n){var i=I(t);return void 0===n?i?i[r]:t[e]:void(i?i.scrollTo(o?f.pageXOffset:n,o?n:f.pageYOffset):t[e]=n)},e,t,arguments.length,null)}}),U.each(["top","left"],function(t,n){U.cssHooks[n]=x(X.pixelPosition,function(t,e){return e?(e=p(t,n),Vt.test(e)?U(t).position()[n]+"px":e):void 0})}),U.each({Height:"height",Width:"width"},function(o,a){U.each({padding:"inner"+o,content:a,"":"outer"+o},function(i,t){U.fn[t]=function(t,e){var n=arguments.length&&(i||"boolean"!=typeof t),r=i||(!0===t||!0===e?"margin":"border");return lt(this,function(t,e,n){var i;return U.isWindow(t)?t.document.documentElement["client"+o]:9===t.nodeType?(i=t.documentElement,Math.max(t.body["scroll"+o],i["scroll"+o],t.body["offset"+o],i["offset"+o],i["client"+o])):void 0===n?U.css(t,e,r):U.style(t,e,n,r)},a,n?t:void 0,n,null)}})}),U.fn.size=function(){return this.length},U.fn.andSelf=U.fn.addBack,"function"==typeof define&&define.amd&&define("jquery",[],function(){return U});var Le=f.jQuery,je=f.$;return U.noConflict=function(t){return f.$===U&&(f.$=je),t&&f.jQuery===U&&(f.jQuery=Le),U},typeof t==bt&&(f.jQuery=f.$=U),U}),function(){function t(s){return function(t,e,n,i){e=b(e,i,4);var r=!S(t)&&y.keys(t),o=(r||t).length,a=0<s?0:o-1;return arguments.length<3&&(n=t[r?r[a]:a],a+=s),function(t,e,n,i,r,o){for(;0<=r&&r<o;r+=s){var a=i?i[r]:r;n=e(n,t[a],a,t)}return n}(t,e,n,r,a,o)}}function e(o){return function(t,e,n){e=x(e,n);for(var i=E(t),r=0<o?0:i-1;0<=r&&r<i;r+=o)if(e(t[r],r,t))return r;return-1}}function n(o,a,s){return function(t,e,n){var i=0,r=E(t);if("number"==typeof n)0<o?i=0<=n?n:Math.max(n+r,i):r=0<=n?Math.min(n+1,r):n+r+1;else if(s&&n&&r)return t[n=s(t,e)]===e?n:-1;if(e!=e)return 0<=(n=a(h.call(t,i,r),y.isNaN))?n+i:-1;for(n=0<o?i:r-1;0<=n&&n<r;n+=o)if(t[n]===e)return n;return-1}}function i(t,e){var n=M.length,i=t.constructor,r=y.isFunction(i)&&i.prototype||l,o="constructor";for(y.has(t,o)&&!y.contains(e,o)&&e.push(o);n--;)(o=M[n])in t&&t[o]!==r[o]&&!y.contains(e,o)&&e.push(o)}function r(){}var o=this,a=o._,s=Array.prototype,l=Object.prototype,u=Function.prototype,c=s.push,h=s.slice,d=l.toString,p=l.hasOwnProperty,f=Array.isArray,g=Object.keys,v=u.bind,m=Object.create,y=function(t){return t instanceof y?t:this instanceof y?void(this._wrapped=t):new y(t)};"undefined"!=typeof exports?("undefined"!=typeof module&&module.exports&&(exports=module.exports=y),exports._=y):o._=y,y.VERSION="1.8.3";var b=function(r,o,t){if(void 0===o)return r;switch(null==t?3:t){case 1:return function(t){return r.call(o,t)};case 2:return function(t,e){return r.call(o,t,e)};case 3:return function(t,e,n){return r.call(o,t,e,n)};case 4:return function(t,e,n,i){return r.call(o,t,e,n,i)}}return function(){return r.apply(o,arguments)}},x=function(t,e,n){return null==t?y.identity:y.isFunction(t)?b(t,e,n):y.isObject(t)?y.matcher(t):y.property(t)};y.iteratee=function(t,e){return x(t,e,1/0)};function w(l,u){return function(t){var e=arguments.length;if(e<2||null==t)return t;for(var n=1;n<e;n++)for(var i=arguments[n],r=l(i),o=r.length,a=0;a<o;a++){var s=r[a];u&&void 0!==t[s]||(t[s]=i[s])}return t}}function $(t){if(!y.isObject(t))return{};if(m)return m(t);r.prototype=t;var e=new r;return r.prototype=null,e}function k(e){return function(t){return null==t?void 0:t[e]}}var C=Math.pow(2,53)-1,E=k("length"),S=function(t){var e=E(t);return"number"==typeof e&&0<=e&&e<=C};y.each=y.forEach=function(t,e,n){var i,r;if(e=b(e,n),S(t))for(i=0,r=t.length;i<r;i++)e(t[i],i,t);else{var o=y.keys(t);for(i=0,r=o.length;i<r;i++)e(t[o[i]],o[i],t)}return t},y.map=y.collect=function(t,e,n){e=x(e,n);for(var i=!S(t)&&y.keys(t),r=(i||t).length,o=Array(r),a=0;a<r;a++){var s=i?i[a]:a;o[a]=e(t[s],s,t)}return o},y.reduce=y.foldl=y.inject=t(1),y.reduceRight=y.foldr=t(-1),y.find=y.detect=function(t,e,n){var i;return void 0!==(i=S(t)?y.findIndex(t,e,n):y.findKey(t,e,n))&&-1!==i?t[i]:void 0},y.filter=y.select=function(t,i,e){var r=[];return i=x(i,e),y.each(t,function(t,e,n){i(t,e,n)&&r.push(t)}),r},y.reject=function(t,e,n){return y.filter(t,y.negate(x(e)),n)},y.every=y.all=function(t,e,n){e=x(e,n);for(var i=!S(t)&&y.keys(t),r=(i||t).length,o=0;o<r;o++){var a=i?i[o]:o;if(!e(t[a],a,t))return!1}return!0},y.some=y.any=function(t,e,n){e=x(e,n);for(var i=!S(t)&&y.keys(t),r=(i||t).length,o=0;o<r;o++){var a=i?i[o]:o;if(e(t[a],a,t))return!0}return!1},y.contains=y.includes=y.include=function(t,e,n,i){return S(t)||(t=y.values(t)),"number"==typeof n&&!i||(n=0),0<=y.indexOf(t,e,n)},y.invoke=function(t,n){var i=h.call(arguments,2),r=y.isFunction(n);return y.map(t,function(t){var e=r?n:t[n];return null==e?e:e.apply(t,i)})},y.pluck=function(t,e){return y.map(t,y.property(e))},y.where=function(t,e){return y.filter(t,y.matcher(e))},y.findWhere=function(t,e){return y.find(t,y.matcher(e))},y.max=function(t,i,e){var n,r,o=-1/0,a=-1/0;if(null==i&&null!=t)for(var s=0,l=(t=S(t)?t:y.values(t)).length;s<l;s++)n=t[s],o<n&&(o=n);else i=x(i,e),y.each(t,function(t,e,n){r=i(t,e,n),(a<r||r===-1/0&&o===-1/0)&&(o=t,a=r)});return o},y.min=function(t,i,e){var n,r,o=1/0,a=1/0;if(null==i&&null!=t)for(var s=0,l=(t=S(t)?t:y.values(t)).length;s<l;s++)(n=t[s])<o&&(o=n);else i=x(i,e),y.each(t,function(t,e,n){((r=i(t,e,n))<a||r===1/0&&o===1/0)&&(o=t,a=r)});return o},y.shuffle=function(t){for(var e,n=S(t)?t:y.values(t),i=n.length,r=Array(i),o=0;o<i;o++)(e=y.random(0,o))!==o&&(r[o]=r[e]),r[e]=n[o];return r},y.sample=function(t,e,n){return null==e||n?(S(t)||(t=y.values(t)),t[y.random(t.length-1)]):y.shuffle(t).slice(0,Math.max(0,e))},y.sortBy=function(t,i,e){return i=x(i,e),y.pluck(y.map(t,function(t,e,n){return{value:t,index:e,criteria:i(t,e,n)}}).sort(function(t,e){var n=t.criteria,i=e.criteria;if(n!==i){if(i<n||void 0===n)return 1;if(n<i||void 0===i)return-1}return t.index-e.index}),"value")};function T(a){return function(i,r,t){var o={};return r=x(r,t),y.each(i,function(t,e){var n=r(t,e,i);a(o,t,n)}),o}}y.groupBy=T(function(t,e,n){y.has(t,n)?t[n].push(e):t[n]=[e]}),y.indexBy=T(function(t,e,n){t[n]=e}),y.countBy=T(function(t,e,n){y.has(t,n)?t[n]++:t[n]=1}),y.toArray=function(t){return t?y.isArray(t)?h.call(t):S(t)?y.map(t,y.identity):y.values(t):[]},y.size=function(t){return null==t?0:S(t)?t.length:y.keys(t).length},y.partition=function(t,i,e){i=x(i,e);var r=[],o=[];return y.each(t,function(t,e,n){(i(t,e,n)?r:o).push(t)}),[r,o]},y.first=y.head=y.take=function(t,e,n){return null==t?void 0:null==e||n?t[0]:y.initial(t,t.length-e)},y.initial=function(t,e,n){return h.call(t,0,Math.max(0,t.length-(null==e||n?1:e)))},y.last=function(t,e,n){return null==t?void 0:null==e||n?t[t.length-1]:y.rest(t,Math.max(0,t.length-e))},y.rest=y.tail=y.drop=function(t,e,n){return h.call(t,null==e||n?1:e)},y.compact=function(t){return y.filter(t,y.identity)};var D=function(t,e,n,i){for(var r=[],o=0,a=i||0,s=E(t);a<s;a++){var l=t[a];if(S(l)&&(y.isArray(l)||y.isArguments(l))){e||(l=D(l,e,n));var u=0,c=l.length;for(r.length+=c;u<c;)r[o++]=l[u++]}else n||(r[o++]=l)}return r};y.flatten=function(t,e){return D(t,e,!1)},y.without=function(t){return y.difference(t,h.call(arguments,1))},y.uniq=y.unique=function(t,e,n,i){y.isBoolean(e)||(i=n,n=e,e=!1),null!=n&&(n=x(n,i));for(var r=[],o=[],a=0,s=E(t);a<s;a++){var l=t[a],u=n?n(l,a,t):l;e?(a&&o===u||r.push(l),o=u):n?y.contains(o,u)||(o.push(u),r.push(l)):y.contains(r,l)||r.push(l)}return r},y.union=function(){return y.uniq(D(arguments,!0,!0))},y.intersection=function(t){for(var e=[],n=arguments.length,i=0,r=E(t);i<r;i++){var o=t[i];if(!y.contains(e,o)){for(var a=1;a<n&&y.contains(arguments[a],o);a++);a===n&&e.push(o)}}return e},y.difference=function(t){var e=D(arguments,!0,!0,1);return y.filter(t,function(t){return!y.contains(e,t)})},y.zip=function(){return y.unzip(arguments)},y.unzip=function(t){for(var e=t&&y.max(t,E).length||0,n=Array(e),i=0;i<e;i++)n[i]=y.pluck(t,i);return n},y.object=function(t,e){for(var n={},i=0,r=E(t);i<r;i++)e?n[t[i]]=e[i]:n[t[i][0]]=t[i][1];return n},y.findIndex=e(1),y.findLastIndex=e(-1),y.sortedIndex=function(t,e,n,i){for(var r=(n=x(n,i,1))(e),o=0,a=E(t);o<a;){var s=Math.floor((o+a)/2);n(t[s])<r?o=s+1:a=s}return o},y.indexOf=n(1,y.findIndex,y.sortedIndex),y.lastIndexOf=n(-1,y.findLastIndex),y.range=function(t,e,n){null==e&&(e=t||0,t=0),n=n||1;for(var i=Math.max(Math.ceil((e-t)/n),0),r=Array(i),o=0;o<i;o++,t+=n)r[o]=t;return r};function _(t,e,n,i,r){if(!(i instanceof e))return t.apply(n,r);var o=$(t.prototype),a=t.apply(o,r);return y.isObject(a)?a:o}y.bind=function(t,e){if(v&&t.bind===v)return v.apply(t,h.call(arguments,1));if(!y.isFunction(t))throw new TypeError("Bind must be called on a function");var n=h.call(arguments,2),i=function(){return _(t,i,e,this,n.concat(h.call(arguments)))};return i},y.partial=function(r){var o=h.call(arguments,1),a=function(){for(var t=0,e=o.length,n=Array(e),i=0;i<e;i++)n[i]=o[i]===y?arguments[t++]:o[i];for(;t<arguments.length;)n.push(arguments[t++]);return _(r,a,this,this,n)};return a},y.bindAll=function(t){var e,n,i=arguments.length;if(i<=1)throw new Error("bindAll must be passed function names");for(e=1;e<i;e++)t[n=arguments[e]]=y.bind(t[n],t);return t},y.memoize=function(i,r){var o=function(t){var e=o.cache,n=""+(r?r.apply(this,arguments):t);return y.has(e,n)||(e[n]=i.apply(this,arguments)),e[n]};return o.cache={},o},y.delay=function(t,e){var n=h.call(arguments,2);return setTimeout(function(){return t.apply(null,n)},e)},y.defer=y.partial(y.delay,y,1),y.throttle=function(n,i,r){var o,a,s,l=null,u=0;r||(r={});function c(){u=!1===r.leading?0:y.now(),l=null,s=n.apply(o,a),l||(o=a=null)}return function(){var t=y.now();u||!1!==r.leading||(u=t);var e=i-(t-u);return o=this,a=arguments,e<=0||i<e?(l&&(clearTimeout(l),l=null),u=t,s=n.apply(o,a),l||(o=a=null)):l||!1===r.trailing||(l=setTimeout(c,e)),s}},y.debounce=function(e,n,i){var r,o,a,s,l,u=function(){var t=y.now()-s;t<n&&0<=t?r=setTimeout(u,n-t):(r=null,i||(l=e.apply(a,o),r||(a=o=null)))};return function(){a=this,o=arguments,s=y.now();var t=i&&!r;return r||(r=setTimeout(u,n)),t&&(l=e.apply(a,o),a=o=null),l}},y.wrap=function(t,e){return y.partial(e,t)},y.negate=function(t){return function(){return!t.apply(this,arguments)}},y.compose=function(){var n=arguments,i=n.length-1;return function(){for(var t=i,e=n[i].apply(this,arguments);t--;)e=n[t].call(this,e);return e}},y.after=function(t,e){return function(){return--t<1?e.apply(this,arguments):void 0}},y.before=function(t,e){var n;return function(){return 0<--t&&(n=e.apply(this,arguments)),t<=1&&(e=null),n}},y.once=y.partial(y.before,2);var P=!{toString:null}.propertyIsEnumerable("toString"),M=["valueOf","isPrototypeOf","toString","propertyIsEnumerable","hasOwnProperty","toLocaleString"];y.keys=function(t){if(!y.isObject(t))return[];if(g)return g(t);var e=[];for(var n in t)y.has(t,n)&&e.push(n);return P&&i(t,e),e},y.allKeys=function(t){if(!y.isObject(t))return[];var e=[];for(var n in t)e.push(n);return P&&i(t,e),e},y.values=function(t){for(var e=y.keys(t),n=e.length,i=Array(n),r=0;r<n;r++)i[r]=t[e[r]];return i},y.mapObject=function(t,e,n){e=x(e,n);for(var i,r=y.keys(t),o=r.length,a={},s=0;s<o;s++)a[i=r[s]]=e(t[i],i,t);return a},y.pairs=function(t){for(var e=y.keys(t),n=e.length,i=Array(n),r=0;r<n;r++)i[r]=[e[r],t[e[r]]];return i},y.invert=function(t){for(var e={},n=y.keys(t),i=0,r=n.length;i<r;i++)e[t[n[i]]]=n[i];return e},y.functions=y.methods=function(t){var e=[];for(var n in t)y.isFunction(t[n])&&e.push(n);return e.sort()},y.extend=w(y.allKeys),y.extendOwn=y.assign=w(y.keys),y.findKey=function(t,e,n){e=x(e,n);for(var i,r=y.keys(t),o=0,a=r.length;o<a;o++)if(e(t[i=r[o]],i,t))return i},y.pick=function(t,e,n){var i,r,o={},a=t;if(null==a)return o;y.isFunction(e)?(r=y.allKeys(a),i=b(e,n)):(r=D(arguments,!1,!1,1),i=function(t,e,n){return e in n},a=Object(a));for(var s=0,l=r.length;s<l;s++){var u=r[s],c=a[u];i(c,u,a)&&(o[u]=c)}return o},y.omit=function(t,e,n){if(y.isFunction(e))e=y.negate(e);else{var i=y.map(D(arguments,!1,!1,1),String);e=function(t,e){return!y.contains(i,e)}}return y.pick(t,e,n)},y.defaults=w(y.allKeys,!0),y.create=function(t,e){var n=$(t);return e&&y.extendOwn(n,e),n},y.clone=function(t){return y.isObject(t)?y.isArray(t)?t.slice():y.extend({},t):t},y.tap=function(t,e){return e(t),t},y.isMatch=function(t,e){var n=y.keys(e),i=n.length;if(null==t)return!i;for(var r=Object(t),o=0;o<i;o++){var a=n[o];if(e[a]!==r[a]||!(a in r))return!1}return!0};var A=function(t,e,n,i){if(t===e)return 0!==t||1/t==1/e;if(null==t||null==e)return t===e;t instanceof y&&(t=t._wrapped),e instanceof y&&(e=e._wrapped);var r=d.call(t);if(r!==d.call(e))return!1;switch(r){case"[object RegExp]":case"[object String]":return""+t==""+e;case"[object Number]":return+t!=+t?+e!=+e:0==+t?1/+t==1/e:+t==+e;case"[object Date]":case"[object Boolean]":return+t==+e}var o="[object Array]"===r;if(!o){if("object"!=typeof t||"object"!=typeof e)return!1;var a=t.constructor,s=e.constructor;if(a!==s&&!(y.isFunction(a)&&a instanceof a&&y.isFunction(s)&&s instanceof s)&&"constructor"in t&&"constructor"in e)return!1}i=i||[];for(var l=(n=n||[]).length;l--;)if(n[l]===t)return i[l]===e;if(n.push(t),i.push(e),o){if((l=t.length)!==e.length)return!1;for(;l--;)if(!A(t[l],e[l],n,i))return!1}else{var u,c=y.keys(t);if(l=c.length,y.keys(e).length!==l)return!1;for(;l--;)if(u=c[l],!y.has(e,u)||!A(t[u],e[u],n,i))return!1}return n.pop(),i.pop(),!0};y.isEqual=function(t,e){return A(t,e)},y.isEmpty=function(t){return null==t||(S(t)&&(y.isArray(t)||y.isString(t)||y.isArguments(t))?0===t.length:0===y.keys(t).length)},y.isElement=function(t){return!(!t||1!==t.nodeType)},y.isArray=f||function(t){return"[object Array]"===d.call(t)},y.isObject=function(t){var e=typeof t;return"function"==e||"object"==e&&!!t},y.each(["Arguments","Function","String","Number","Date","RegExp","Error"],function(e){y["is"+e]=function(t){return d.call(t)==="[object "+e+"]"}}),y.isArguments(arguments)||(y.isArguments=function(t){return y.has(t,"callee")}),"function"!=typeof/./&&"object"!=typeof Int8Array&&(y.isFunction=function(t){return"function"==typeof t||!1}),y.isFinite=function(t){return isFinite(t)&&!isNaN(parseFloat(t))},y.isNaN=function(t){return y.isNumber(t)&&t!==+t},y.isBoolean=function(t){return!0===t||!1===t||"[object Boolean]"===d.call(t)},y.isNull=function(t){return null===t},y.isUndefined=function(t){return void 0===t},y.has=function(t,e){return null!=t&&p.call(t,e)},y.noConflict=function(){return o._=a,this},y.identity=function(t){return t},y.constant=function(t){return function(){return t}},y.noop=function(){},y.property=k,y.propertyOf=function(e){return null==e?function(){}:function(t){return e[t]}},y.matcher=y.matches=function(e){return e=y.extendOwn({},e),function(t){return y.isMatch(t,e)}},y.times=function(t,e,n){var i=Array(Math.max(0,t));e=b(e,n,1);for(var r=0;r<t;r++)i[r]=e(r);return i},y.random=function(t,e){return null==e&&(e=t,t=0),t+Math.floor(Math.random()*(e-t+1))},y.now=Date.now||function(){return(new Date).getTime()};function N(e){function n(t){return e[t]}var t="(?:"+y.keys(e).join("|")+")",i=RegExp(t),r=RegExp(t,"g");return function(t){return t=null==t?"":""+t,i.test(t)?t.replace(r,n):t}}var O={"&":"&amp;","<":"&lt;",">":"&gt;",'"':"&quot;","'":"&#x27;","`":"&#x60;"},I=y.invert(O);y.escape=N(O),y.unescape=N(I),y.result=function(t,e,n){var i=null==t?void 0:t[e];return void 0===i&&(i=n),y.isFunction(i)?i.call(t):i};var B=0;y.uniqueId=function(t){var e=++B+"";return t?t+e:e},y.templateSettings={evaluate:/<%([\s\S]+?)%>/g,interpolate:/<%=([\s\S]+?)%>/g,escape:/<%-([\s\S]+?)%>/g};function L(t){return"\\"+R[t]}var j=/(.)^/,R={"'":"'","\\":"\\","\r":"r","\n":"n","\u2028":"u2028","\u2029":"u2029"},F=/\\|'|\r|\n|\u2028|\u2029/g;y.template=function(o,t,e){!t&&e&&(t=e),t=y.defaults({},t,y.templateSettings);var n=RegExp([(t.escape||j).source,(t.interpolate||j).source,(t.evaluate||j).source].join("|")+"|$","g"),a=0,s="__p+='";o.replace(n,function(t,e,n,i,r){return s+=o.slice(a,r).replace(F,L),a=r+t.length,e?s+="'+\n((__t=("+e+"))==null?'':_.escape(__t))+\n'":n?s+="'+\n((__t=("+n+"))==null?'':__t)+\n'":i&&(s+="';\n"+i+"\n__p+='"),t}),s+="';\n",t.variable||(s="with(obj||{}){\n"+s+"}\n"),s="var __t,__p='',__j=Array.prototype.join,print=function(){__p+=__j.call(arguments,'');};\n"+s+"return __p;\n";try{var i=new Function(t.variable||"obj","_",s)}catch(t){throw t.source=s,t}function r(t){return i.call(this,t,y)}var l=t.variable||"obj";return r.source="function("+l+"){\n"+s+"}",r},y.chain=function(t){var e=y(t);return e._chain=!0,e};function V(t,e){return t._chain?y(e).chain():e}y.mixin=function(n){y.each(y.functions(n),function(t){var e=y[t]=n[t];y.prototype[t]=function(){var t=[this._wrapped];return c.apply(t,arguments),V(this,e.apply(y,t))}})},y.mixin(y),y.each(["pop","push","reverse","shift","sort","splice","unshift"],function(e){var n=s[e];y.prototype[e]=function(){var t=this._wrapped;return n.apply(t,arguments),"shift"!==e&&"splice"!==e||0!==t.length||delete t[0],V(this,t)}}),y.each(["concat","join","slice"],function(t){var e=s[t];y.prototype[t]=function(){return V(this,e.apply(this._wrapped,arguments))}}),y.prototype.value=function(){return this._wrapped},y.prototype.valueOf=y.prototype.toJSON=y.prototype.value,y.prototype.toString=function(){return""+this._wrapped},"function"==typeof define&&define.amd&&define("underscore",[],function(){return y})}.call(this),function(f,J,tt){"use strict";function k(o){return function(){var t,e,n,i=arguments[0],r=arguments;for(t=(t="["+(o?o+":":"")+i+"] "+arguments[1].replace(/\{\d+\}/g,function(t){var e,n=+t.slice(1,-1);return 2+n<r.length?"function"==typeof(e=r[2+n])?e.toString().replace(/ ?\{[\s\S]*$/,""):void 0===e?"undefined":"string"!=typeof e?A(e):e:t}))+"\nhttp://errors.angularjs.org/1.2.28/"+(o?o+"/":"")+i,e=2;e<arguments.length;e++)t=t+(2==e?"?":"&")+"p"+(e-2)+"="+encodeURIComponent("function"==typeof(n=arguments[e])?n.toString().replace(/ \{[\s\S]*$/,""):void 0===n?"undefined":"string"!=typeof n?JSON.stringify(n):n);return new Error(t)}}function P(t){if(null==t||h(t))return!1;var e=t.length;return!(1!==t.nodeType||!e)||(it(t)||en(t)||0===e||"number"==typeof e&&0<e&&e-1 in t)}function et(t,e,n){var i;if(t)if(rt(t))for(i in t)"prototype"==i||"length"==i||"name"==i||t.hasOwnProperty&&!t.hasOwnProperty(i)||e.call(n,t[i],i);else if(en(t)||P(t))for(i=0;i<t.length;i++)e.call(n,t[i],i);else if(t.forEach&&t.forEach!==et)t.forEach(e,n);else for(i in t)t.hasOwnProperty(i)&&e.call(n,t[i],i);return t}function R(t){var e=[];for(var n in t)t.hasOwnProperty(n)&&e.push(n);return e.sort()}function g(n){return function(t,e){n(e,t)}}function r(){for(var t,e=tn.length;e;){if(57==(t=tn[--e].charCodeAt(0)))return tn[e]="A",tn.join("");if(90!=t)return tn[e]=String.fromCharCode(t+1),tn.join("");tn[e]="0"}return tn.unshift("0"),tn.join("")}function u(t,e){e?t.$$hashKey=e:delete t.$$hashKey}function M(n){var t=n.$$hashKey;return et(arguments,function(t){t!==n&&et(t,function(t,e){n[e]=t})}),u(n,t),n}function $(t){return parseInt(t,10)}function v(t,e){return M(new(M(function(){},{prototype:t})),e)}function E(){}function m(t){return t}function y(t){return function(){return t}}function S(t){return void 0===t}function F(t){return void 0!==t}function nt(t){return null!=t&&"object"==typeof t}function it(t){return"string"==typeof t}function l(t){return"number"==typeof t}function b(t){return"[object Date]"===Ke.call(t)}function rt(t){return"function"==typeof t}function c(t){return"[object RegExp]"===Ke.call(t)}function h(t){return t&&t.document&&t.location&&t.alert&&t.setInterval}function d(t){return t&&t.$evalAsync&&t.$watch}function x(t){return t&&rt(t.then)}function e(t){return!(!t||!(t.nodeName||t.prop&&t.attr&&t.find))}function C(t,e){if(t.indexOf)return t.indexOf(e);for(var n=0;n<t.length;n++)if(e===t[n])return n;return-1}function p(t,e){var n=C(t,e);return 0<=n&&t.splice(n,1),e}function T(t,n,e,i){if(h(t)||d(t))throw Qe("cpws","Can't copy! Making copies of Window or Scope instances is not supported.");if(n){if(t===n)throw Qe("cpi","Can't copy! Source and destination are identical.");if(e=e||[],i=i||[],nt(t)){var r=C(e,t);if(-1!==r)return i[r];e.push(t),i.push(n)}var o;if(en(t))for(var a=n.length=0;a<t.length;a++)o=T(t[a],null,e,i),nt(t[a])&&(e.push(t[a]),i.push(o)),n.push(o);else{var s=n.$$hashKey;for(var l in en(n)?n.length=0:et(n,function(t,e){delete n[e]}),t)o=T(t[l],null,e,i),nt(t[l])&&(e.push(t[l]),i.push(o)),n[l]=o;u(n,s)}}else(n=t)&&(en(t)?n=T(t,[],e,i):b(t)?n=new Date(t.getTime()):c(t)?(n=new RegExp(t.source,t.toString().match(/[^\/]*$/)[0])).lastIndex=t.lastIndex:nt(t)&&(n=T(t,{},e,i)));return n}function ot(t,e){if(en(t)){e=e||[];for(var n=0;n<t.length;n++)e[n]=t[n]}else if(nt(t))for(var i in e=e||{},t)!qe.call(t,i)||"$"===i.charAt(0)&&"$"===i.charAt(1)||(e[i]=t[i]);return e||t}function at(t,e){if(t===e)return!0;if(null===t||null===e)return!1;if(t!=t&&e!=e)return!0;var n,i,r,o=typeof t;if(o==typeof e&&"object"==o){if(!en(t)){if(b(t))return!!b(e)&&(isNaN(t.getTime())&&isNaN(e.getTime())||t.getTime()===e.getTime());if(c(t)&&c(e))return t.toString()==e.toString();if(d(t)||d(e)||h(t)||h(e)||en(e))return!1;for(i in r={},t)if("$"!==i.charAt(0)&&!rt(t[i])){if(!at(t[i],e[i]))return!1;r[i]=!0}for(i in e)if(!r.hasOwnProperty(i)&&"$"!==i.charAt(0)&&e[i]!==tt&&!rt(e[i]))return!1;return!0}if(!en(e))return!1;if((n=t.length)==e.length){for(i=0;i<n;i++)if(!at(t[i],e[i]))return!1;return!0}}return!1}function D(t,e,n){return t.concat(Ge.call(e,n))}function st(t,e){return Ge.call(t,e||0)}function _(t,e){var n=2<arguments.length?st(arguments,2):[];return!rt(e)||e instanceof RegExp?e:n.length?function(){return arguments.length?e.apply(t,n.concat(Ge.call(arguments,0))):e.apply(t,n)}:function(){return arguments.length?e.apply(t,arguments):e.call(t)}}function n(t,e){var n=e;return"string"==typeof t&&"$"===t.charAt(0)?n=tt:h(e)?n="$WINDOW":e&&J===e?n="$DOCUMENT":d(e)&&(n="$SCOPE"),n}function A(t,e){return void 0===t?tt:JSON.stringify(t,n,e?"  ":null)}function a(t){return it(t)?JSON.parse(t):t}function N(t){if("function"==typeof t)t=!0;else if(t&&0!==t.length){var e=Ve(""+t);t=!("f"==e||"0"==e||"false"==e||"no"==e||"n"==e||"[]"==e)}else t=!1;return t}function lt(t){t=Xe(t).clone();try{t.empty()}catch(t){}var e=Xe("<div>").append(t).html();try{return 3===t[0].nodeType?Ve(e):e.match(/^(<[^>]+>)/)[1].replace(/^<([\w\-]+)/,function(t,e){return"<"+Ve(e)})}catch(t){return Ve(e)}}function o(t){try{return decodeURIComponent(t)}catch(t){}}function s(t){var n,i,r={};return et((t||"").split("&"),function(t){if(t&&(n=t.replace(/\+/g,"%20").split("="),F(i=o(n[0])))){var e=!F(n[1])||o(n[1]);qe.call(r,i)?en(r[i])?r[i].push(e):r[i]=[r[i],e]:r[i]=e}}),r}function i(t){var n=[];return et(t,function(t,e){en(t)?et(t,function(t){n.push(O(e,!0)+(!0===t?"":"="+O(t,!0)))}):n.push(O(e,!0)+(!0===t?"":"="+O(t,!0)))}),n.length?n.join("&"):""}function w(t){return O(t,!0).replace(/%26/gi,"&").replace(/%3D/gi,"=").replace(/%2B/gi,"+")}function O(t,e){return encodeURIComponent(t).replace(/%40/gi,"@").replace(/%3A/gi,":").replace(/%24/g,"$").replace(/%2C/gi,",").replace(/%20/g,e?"%20":"+")}function I(n,i){function e(){if((n=Xe(n)).injector()){var t=n[0]===J?"document":lt(n);throw Qe("btstrpd","App Already Bootstrapped with this Element '{0}'",t.replace(/</,"&lt;").replace(/>/,"&gt;"))}(i=i||[]).unshift(["$provide",function(t){t.value("$rootElement",n)}]),i.unshift("ng");var e=wt(i);return e.invoke(["$rootScope","$rootElement","$compile","$injector","$animate",function(t,e,n,i,r){t.$apply(function(){e.data("$injector",i),n(e)(t)})}]),e}var t=/^NG_DEFER_BOOTSTRAP!/;return f&&!t.test(f.name)?e():(f.name=f.name.replace(t,""),void(Je.resumeBootstrap=function(t){et(t,function(t){i.push(t)}),e()}))}function ut(t,n){return n=n||"_",t.replace(on,function(t,e){return(e?n:"")+t.toLowerCase()})}function ct(t,e,n){if(!t)throw Qe("areq","Argument '{0}' is {1}",e||"?",n||"required");return t}function B(t,e,n){return n&&en(t)&&(t=t[t.length-1]),ct(rt(t),e,"not a function, got "+(t&&"object"==typeof t?t.constructor.name||"Object":typeof t)),t}function L(t,e){if("hasOwnProperty"===t)throw Qe("badname","hasOwnProperty is not a valid {0} name",e)}function j(t,e,n){if(!e)return t;for(var i,r=e.split("."),o=t,a=r.length,s=0;s<a;s++)i=r[s],t&&(t=(o=t)[i]);return!n&&rt(t)?_(o,t):t}function V(t){var e=t[0],n=t[t.length-1];if(e===n)return Xe(e);var i=e,r=[i];do{if(!(i=i.nextSibling))break;r.push(i)}while(i!==n);return Xe(r)}function q(t){return t.replace(hn,function(t,e,n,i){return i?n.toUpperCase():n}).replace(dn,"Moz$1")}function t(t,c,h,d){function e(t){var e,n,i,r,o,a,s,l=h&&t?[this.filter(t)]:[this],u=c;if(!d||null!=t)for(;l.length;)for(n=0,i=(e=l.shift()).length;n<i;n++)for(r=Xe(e[n]),u?r.triggerHandler("$destroy"):u=!u,o=0,a=(s=r.children()).length;o<a;o++)l.push(Ye(s[o]));return p.apply(this,arguments)}var p=Ye.fn[t];p=p.$original||p,e.$original=p,Ye.fn[t]=e}function ht(t){return!gn.test(t)}function z(t,e){var n;return e=e||J,(n=fn.exec(t))?[e.createElement(n[1])]:function(t,e){var n,i,r,o,a,s,l=e.createDocumentFragment(),u=[];if(ht(t))u.push(e.createTextNode(t));else{for(n=l.appendChild(e.createElement("div")),i=(vn.exec(t)||["",""])[1].toLowerCase(),r=yn[i]||yn._default,n.innerHTML="<div>&#160;</div>"+r[1]+t.replace(mn,"<$1></$2>")+r[2],n.removeChild(n.firstChild),o=r[0];o--;)n=n.lastChild;for(a=0,s=n.childNodes.length;a<s;++a)u.push(n.childNodes[a]);(n=l.firstChild).textContent=""}return l.textContent="",l.innerHTML="",u}(t,e)}function H(t){if(t instanceof H)return t;if(it(t)&&(t=nn(t)),!(this instanceof H)){if(it(t)&&"<"!=t.charAt(0))throw pn("nosel","Looking up elements via selectors is not supported by jqLite! See: http://docs.angularjs.org/api/angular.element");return new H(t)}it(t)?(pt(this,z(t)),Xe(J.createDocumentFragment()).append(this)):pt(this,t)}function dt(t){return t.cloneNode(!0)}function X(t){U(t);for(var e=0,n=t.childNodes||[];e<n.length;e++)X(n[e])}function Y(n,t,e,i){if(F(i))throw pn("offargs","jqLite#off() does not support the `selector` argument");var r=W(n,"events");W(n,"handle")&&(S(t)?et(r,function(t,e){cn(n,e,t),delete r[e]}):et(t.split(" "),function(t){S(e)?(cn(n,t,r[t]),delete r[t]):p(r[t]||[],e)}))}function U(t,e){var n=t.ng339,i=sn[n];if(i){if(e)return void delete sn[n].data[e];i.handle&&(i.events.$destroy&&i.handle({},"$destroy"),Y(t)),delete sn[n],t.ng339=tt}}function W(t,e,n){var i=t.ng339,r=sn[i||-1];return F(n)?(r||(t.ng339=i=++ln,r=sn[i]={}),void(r[e]=n)):r&&r[e]}function G(t,e,n){var i=W(t,"data"),r=F(n),o=!r&&F(e),a=o&&!nt(e);if(i||a||W(t,"data",i={}),r)i[e]=n;else{if(!o)return i;if(a)return i&&i[e];M(i,e)}}function Z(t,e){return!!t.getAttribute&&-1<(" "+(t.getAttribute("class")||"")+" ").replace(/[\n\t]/g," ").indexOf(" "+e+" ")}function K(e,t){t&&e.setAttribute&&et(t.split(" "),function(t){e.setAttribute("class",nn((" "+(e.getAttribute("class")||"")+" ").replace(/[\n\t]/g," ").replace(" "+nn(t)+" "," ")))})}function Q(t,e){if(e&&t.setAttribute){var n=(" "+(t.getAttribute("class")||"")+" ").replace(/[\n\t]/g," ");et(e.split(" "),function(t){t=nn(t),-1===n.indexOf(" "+t+" ")&&(n+=t+" ")}),t.setAttribute("class",nn(n))}}function pt(t,e){if(e){e=e.nodeName||!F(e.length)||h(e)?[e]:e;for(var n=0;n<e.length;n++)t.push(e[n])}}function ft(t,e){return gt(t,"$"+(e||"ngController")+"Controller")}function gt(t,e,n){9==t.nodeType&&(t=t.documentElement);for(var i=en(e)?e:[e];t;){for(var r=0,o=i.length;r<o;r++)if((n=Xe.data(t,i[r]))!==tt)return n;t=t.parentNode||11===t.nodeType&&t.host}}function vt(t){for(var e=0,n=t.childNodes;e<n.length;e++)X(n[e]);for(;t.firstChild;)t.removeChild(t.firstChild)}function mt(t,e){var n=xn[e.toLowerCase()];return n&&$n[t.nodeName]&&n}function yt(t,e){var n,i=typeof t;return"function"==i||"object"==i&&null!==t?"function"==typeof(n=t.$$hashKey)?n=t.$$hashKey():n===tt&&(n=t.$$hashKey=(e||r)()):n=t,i+":"+n}function bt(t,e){if(e){var n=0;this.nextUid=function(){return++n}}et(t,this.put,this)}function xt(t){var i,e;return"function"==typeof t?(i=t.$inject)||(i=[],t.length&&et(t.toString().replace(Tn,"").match(Cn)[1].split(En),function(t){t.replace(Sn,function(t,e,n){i.push(n)})}),t.$inject=i):en(t)?(B(t[e=t.length-1],"fn"),i=t.slice(0,e)):B(t,"fn",!0),i}function wt(t){function e(n){return function(t,e){return nt(t)?void et(t,g(n)):n(t,e)}}function n(t,e){if(L(t,"service"),(rt(e)||en(e))&&(e=d.instantiate(e)),!e.$get)throw Dn("pget","Provider '{0}' must define $get factory method.",t);return h[t+s]=e}function i(t,e){return n(t,{$get:e})}function r(n,t){function l(e){if(n.hasOwnProperty(e)){if(n[e]===a)throw Dn("cdep","Circular dependency found: {0}",e+" <- "+u.join(" <- "));return n[e]}try{return u.unshift(e),n[e]=a,n[e]=t(e)}catch(t){throw n[e]===a&&delete n[e],t}finally{u.shift()}}function o(t,e,n){var i,r,o,a=[],s=xt(t);for(r=0,i=s.length;r<i;r++){if("string"!=typeof(o=s[r]))throw Dn("itkn","Incorrect injection token! Expected service name as string, got {0}",o);a.push(n&&n.hasOwnProperty(o)?n[o]:l(o))}return en(t)&&(t=t[i]),t.apply(e,a)}return{invoke:o,instantiate:function(t,e){function n(){}var i,r;return n.prototype=(en(t)?t[t.length-1]:t).prototype,nt(r=o(t,i=new n,e))||rt(r)?r:i},get:l,annotate:xt,has:function(t){return h.hasOwnProperty(t+s)||n.hasOwnProperty(t)}}}var a={},s="Provider",u=[],c=new bt([],!0),h={$provide:{provider:e(n),factory:e(i),service:e(function(t,e){return i(t,["$injector",function(t){return t.instantiate(e)}])}),value:e(function(t,e){return i(t,y(e))}),constant:e(function(t,e){L(t,"constant"),h[t]=e,o[t]=e}),decorator:function(t,e){var n=d.get(t+s),i=n.$get;n.$get=function(){var t=l.invoke(i,n);return l.invoke(e,null,{$delegate:t})}}}},d=h.$injector=r(h,function(){throw Dn("unpr","Unknown provider: {0}",u.join(" <- "))}),o={},l=o.$injector=r(o,function(t){var e=d.get(t+s);return l.invoke(e.$get,e)});return et(function i(t){var r,o,a,s,l=[];return et(t,function(e){if(!c.get(e)){c.put(e,!0);try{if(it(e))for(r=Ue(e),l=l.concat(i(r.requires)).concat(r._runBlocks),o=r._invokeQueue,a=0,s=o.length;a<s;a++){var t=o[a],n=d.get(t[0]);n[t[1]].apply(n,t[2])}else rt(e)?l.push(d.invoke(e)):en(e)?l.push(d.invoke(e)):B(e,"module")}catch(t){throw en(e)&&(e=e[e.length-1]),t.message&&t.stack&&-1==t.stack.indexOf(t.message)&&(t=t.message+"\n"+t.stack),Dn("modulerr","Failed to instantiate module {0} due to:\n{1}",e,t.stack||t.message||t)}}}),l}(t),function(t){l.invoke(t||E)}),l}function $t(){var o=!0;this.disableAutoScrolling=function(){o=!1},this.$get=["$window","$location","$rootScope",function(n,i,t){function e(){var t,e=i.hash();e?(t=r.getElementById(e))?t.scrollIntoView():(t=function(t){var e=null;return et(t,function(t){e||"a"!==Ve(t.nodeName)||(e=t)}),e}(r.getElementsByName(e)))?t.scrollIntoView():"top"===e&&n.scrollTo(0,0):n.scrollTo(0,0)}var r=n.document;return o&&t.$watch(function(){return i.hash()},function(){t.$evalAsync(e)}),e}]}function kt(){this.$get=["$$rAF","$timeout",function(e,n){return e.supported?function(t){return e(t)}:function(t){return n(t,0,!1)}}]}function Ct(i,t,s,r){function o(t){try{t.apply(null,st(arguments,1))}finally{if(0===--p)for(;f.length;)try{f.pop()()}catch(t){s.error(t)}}}function e(){m!=a.url()&&(m=a.url(),et(x,function(t){t(a.url())}))}var a=this,l=t[0],u=i.location,c=i.history,h=i.setTimeout,n=i.clearTimeout,d={};a.isMock=!1;var p=0,f=[];a.$$completeOutstandingRequest=o,a.$$incOutstandingRequestCount=function(){p++},a.notifyWhenNoOutstandingRequests=function(t){et(v,function(t){t()}),0===p?t():f.push(t)};var g,v=[];a.addPollFn=function(t){return S(g)&&function(e,n){!function t(){et(v,function(t){t()}),g=n(t,e)}()}(100,h),v.push(t),t};var m=u.href,y=t.find("base"),b=null;a.url=function(t,e){if(u!==i.location&&(u=i.location),c!==i.history&&(c=i.history),t){if(m==t)return;var n=m&&Wt(m)===Wt(t);return m=t,!n&&r.history?e?c.replaceState(null,"",t):(c.pushState(null,"",t),y.attr("href",y.attr("href"))):(n||(b=t),e?u.replace(t):u.href=t),a}return b||u.href.replace(/%27/g,"'")};var x=[],w=!1;a.onUrlChange=function(t){return w||(r.history&&Xe(i).on("popstate",e),r.hashchange?Xe(i).on("hashchange",e):a.addPollFn(e),w=!0),x.push(t),t},a.$$checkUrlChange=e,a.baseHref=function(){var t=y.attr("href");return t?t.replace(/^(https?\:)?\/\/[^\/]*/,""):""};var $={},k="",C=a.baseHref();a.cookies=function(t,e){var n,i,r,o,a;if(!t){if(l.cookie!==k)for(i=(k=l.cookie).split("; "),$={},o=0;o<i.length;o++)0<(a=(r=i[o]).indexOf("="))&&(t=unescape(r.substring(0,a)),$[t]===tt&&($[t]=unescape(r.substring(a+1))));return $}e===tt?l.cookie=escape(t)+"=;path="+C+";expires=Thu, 01 Jan 1970 00:00:00 GMT":!it(e)||4096<(n=(l.cookie=escape(t)+"="+escape(e)+";path="+C).length+1)&&s.warn("Cookie '"+t+"' possibly not set or overflowed because it was too large ("+n+" > 4096 bytes)!")},a.defer=function(t,e){var n;return p++,n=h(function(){delete d[n],o(t)},e||0),d[n]=!0,n},a.defer.cancel=function(t){return!!d[t]&&(delete d[t],n(t),o(E),!0)}}function Et(){this.$get=["$window","$log","$sniffer","$document",function(t,e,n,i){return new Ct(t,i,e,n)}]}function St(){this.$get=function(){function t(t,e){function n(t){t!=u&&(c?c==t&&(c=t.n):c=t,i(t.n,t.p),i(t,u),(u=t).n=null)}function i(t,e){t!=e&&(t&&(t.p=e),e&&(e.n=t))}if(t in h)throw k("$cacheFactory")("iid","CacheId '{0}' is already taken!",t);var r=0,o=M({},e,{id:t}),a={},s=e&&e.capacity||Number.MAX_VALUE,l={},u=null,c=null;return h[t]={put:function(t,e){s<Number.MAX_VALUE&&n(l[t]||(l[t]={key:t}));if(!S(e))return t in a||r++,a[t]=e,s<r&&this.remove(c.key),e},get:function(t){if(s<Number.MAX_VALUE){var e=l[t];if(!e)return;n(e)}return a[t]},remove:function(t){if(s<Number.MAX_VALUE){var e=l[t];if(!e)return;e==u&&(u=e.p),e==c&&(c=e.n),i(e.n,e.p),delete l[t]}delete a[t],r--},removeAll:function(){a={},r=0,l={},u=c=null},destroy:function(){l=o=a=null,delete h[t]},info:function(){return M({},o,{size:r})}}}var h={};return t.info=function(){var n={};return et(h,function(t,e){n[e]=t.info()}),n},t.get=function(t){return h[t]},t}}function Tt(){this.$get=["$cacheFactory",function(t){return t("templates")}]}function Dt(n,e){var p={},f="Directive",E=/^\s*directive\:\s*([\d\w_\-]+)\s+(.*)$/,S=/(([\d\w_\-]+)(?:\:([^;]+))?;?)/,u=/^(on[a-z]+|formaction)$/;this.directive=function t(a,e){return L(a,"directive"),it(a)?(ct(e,"directiveFactory"),p.hasOwnProperty(a)||(p[a]=[],n.factory(a+f,["$injector","$exceptionHandler",function(i,r){var o=[];return et(p[a],function(t,e){try{var n=i.invoke(t);rt(n)?n={compile:y(n)}:!n.compile&&n.link&&(n.compile=y(n.link)),n.priority=n.priority||0,n.index=e,n.name=n.name||a,n.require=n.require||n.controller&&n.name,n.restrict=n.restrict||"A",o.push(n)}catch(t){r(t)}}),o}])),p[a].push(e)):et(a,g(t)),this},this.aHrefSanitizationWhitelist=function(t){return F(t)?(e.aHrefSanitizationWhitelist(t),this):e.aHrefSanitizationWhitelist()},this.imgSrcSanitizationWhitelist=function(t){return F(t)?(e.imgSrcSanitizationWhitelist(t),this):e.imgSrcSanitizationWhitelist()},this.$get=["$injector","$interpolate","$exceptionHandler","$http","$templateCache","$parse","$controller","$rootScope","$document","$sce","$animate","$$sanitizeUri",function(d,B,L,t,e,j,R,o,n,s,r,l){function F(l,t,e,n,i){l instanceof Xe||(l=Xe(l)),et(l,function(t,e){3==t.nodeType&&t.nodeValue.match(/\S+/)&&(l[e]=t=Xe(t).wrap("<span></span>").parent()[0])});var u=T(l,t,l,e,n,i);return V(l,"ng-scope"),function(t,e,n,i){ct(t,"scope");var r=e?bn.clone.call(l):l;et(n,function(t,e){r.data("$"+e+"Controller",t)});for(var o=0,a=r.length;o<a;o++){var s=r[o].nodeType;1!==s&&9!==s||r.eq(o).data("$scope",t)}return e&&e(r,t),u&&u(t,r,r,i),r}}function V(t,e){try{t.addClass(e)}catch(t){}}function T(t,p,e,n,i,r){for(var o,a,s,l,u,c,f=[],h=0;h<t.length;h++)o=new K,(s=(a=q(t[h],[],o,0===h?n:tt,i)).length?_(a,t[h],o,p,e,null,[],[],r):null)&&s.scope&&V(o.$$element,"ng-scope"),u=s&&s.terminal||!(l=t[h].childNodes)||!l.length?null:T(l,s?(s.transcludeOnThisElement||!s.templateOnThisElement)&&s.transclude:p),f.push(s,u),c=c||s||u,r=null;return c?function(t,e,n,i){var r,o,a,s,l,u,c,h=e.length,d=new Array(h);for(l=0;l<h;l++)d[l]=e[l];for(c=l=0,u=f.length;l<u;c++)a=d[c],r=f[l++],o=f[l++],r?(r.scope?(s=t.$new(),Xe.data(a,"$scope",s)):s=t,r(o,s,a,n,r.transcludeOnThisElement?D(t,r.transclude,i):!r.templateOnThisElement&&i?i:!i&&p?D(t,p):null)):o&&o(t,a.childNodes,tt,i)}:null}function D(o,a,s){return function(t,e,n){var i=!1;t||(i=(t=o.$new()).$$transcluded=!0);var r=a(t,e,n,s);return i&&r.on("$destroy",function(){t.$destroy()}),r}}function q(t,e,n,i,r){var o,a,s=t.nodeType,l=n.$attr;switch(s){case 1:w(e,_t(We(t).toLowerCase()),"E",i,r);for(var u,c,h,d,p,f,g=t.attributes,v=0,m=g&&g.length;v<m;v++){var y=!1,b=!1;if(u=g[v],!He||8<=He||u.specified){c=u.name,p=nn(u.value),d=_t(c),(f=C.test(d))&&(c=ut(d.substr(6),"-"));var x=d.replace(/(Start|End)$/,"");d===x+"Start"&&(b=(y=c).substr(0,c.length-5)+"end",c=c.substr(0,c.length-6)),l[h=_t(c.toLowerCase())]=c,!f&&n.hasOwnProperty(h)||(n[h]=p,mt(t,h)&&(n[h]=!0)),k(t,e,p,h),w(e,h,"A",i,r,y,b)}}if(it(a=t.className)&&""!==a)for(;o=S.exec(a);)w(e,h=_t(o[2]),"C",i,r)&&(n[h]=nn(o[3])),a=a.substr(o.index+o[0].length);break;case 3:!function(t,e){var o=B(e,!0);o&&t.push({priority:0,compile:function(t){var r=t.parent().length;return r&&V(t.parent(),"ng-binding"),function(t,e){var n=e.parent(),i=n.data("$binding")||[];i.push(o),n.data("$binding",i),r||V(n,"ng-binding"),t.$watch(o,function(t){e[0].nodeValue=t})}}})}(e,t.nodeValue);break;case 8:try{(o=E.exec(t.nodeValue))&&(w(e,h=_t(o[1]),"M",i,r)&&(n[h]=nn(o[2])))}catch(t){}}return e.sort($),e}function z(t,e,n){var i=[],r=0;if(e&&t.hasAttribute&&t.hasAttribute(e))do{if(!t)throw Mn("uterdir","Unterminated attribute, found '{0}' but no matching '{1}' found.",e,n);1==t.nodeType&&(t.hasAttribute(e)&&r++,t.hasAttribute(n)&&r--),i.push(t),t=t.nextSibling}while(0<r);else i.push(t);return Xe(i)}function H(o,a,s){return function(t,e,n,i,r){return e=z(e[0],a,s),o(t,e,n,i,r)}}function _(t,v,m,e,n,i,y,b,r){function o(t,e,n,i){t&&(n&&(t=H(t,n,i)),t.require=l.require,t.directiveName=u,$!==l&&!l.$$isolateScope||(t=Z(t,{isolateScope:!0})),y.push(t)),e&&(n&&(e=H(e,n,i)),e.require=l.require,e.directiveName=u,$!==l&&!l.$$isolateScope||(e=Z(e,{isolateScope:!0})),b.push(e))}function x(e,t,n,i){var r,o="data",a=!1;if(it(t)){for(;"^"==(r=t.charAt(0))||"?"==r;)t=t.substr(1),"^"==r&&(o="inheritedData"),a=a||"?"==r;if(r=null,i&&"data"===o&&(r=i[t]),!(r=r||n[o]("$"+t+"Controller"))&&!a)throw Mn("ctreq","Controller '{0}', required by directive '{1}', can't be found!",t,e);return r}return en(t)&&(r=[],et(t,function(t){r.push(x(e,t,n,i))})),r}function a(t,c,e,n,i){var h,r,o,a,s,l,d,u,p={};if(h=v===e?m:ot(m,new K(Xe(e),m.$attr)),r=h.$$element,$){var f=/^\s*([@=&])(\??)\s*(\w*)\s*$/;d=c.$new(!0),!k||k!==$&&k!==$.$$originalDirective?r.data("$isolateScopeNoTemplate",d):r.data("$isolateScope",d),V(r,"ng-isolate-scope"),et($.scope,function(t,e){var n,i,r,o,a=t.match(f)||[],s=a[3]||e,l="?"==a[2],u=a[1];switch(d.$$isolateBindings[e]=u+s,u){case"@":h.$observe(s,function(t){d[e]=t}),h.$$observers[s].$$scope=c,h[s]&&(d[e]=B(h[s])(c));break;case"=":if(l&&!h[s])return;i=j(h[s]),o=i.literal?at:function(t,e){return t===e||t!=t&&e!=e},r=i.assign||function(){throw n=d[e]=i(c),Mn("nonassign","Expression '{0}' used with directive '{1}' is non-assignable!",h[s],$.name)},n=d[e]=i(c),d.$watch(function(){var t=i(c);return o(t,d[e])||(o(t,n)?r(c,t=d[e]):d[e]=t),n=t},null,i.literal);break;case"&":i=j(h[s]),d[e]=function(t){return i(c,t)};break;default:throw Mn("iscp","Invalid isolate scope definition for directive '{0}'. Definition: {... {1}: '{2}' ...}",$.name,e,t)}})}for(u=i&&function(t,e){var n;return arguments.length<2&&(e=t,t=tt),E&&(n=p),i(t,e,n)},w&&et(w,function(t){var e,n={$scope:t===$||t.$$isolateScope?d:c,$element:r,$attrs:h,$transclude:u};"@"==(l=t.controller)&&(l=h[t.name]),e=R(l,n),p[t.name]=e,E||r.data("$"+t.name+"Controller",e),t.controllerAs&&(n.$scope[t.controllerAs]=e)}),o=0,a=y.length;o<a;o++)try{(s=y[o])(s.isolateScope?d:c,r,h,s.require&&x(s.directiveName,s.require,r,p),u)}catch(t){L(t,lt(r))}var g=c;for($&&($.template||null===$.templateUrl)&&(g=d),t&&t(g,e.childNodes,tt,i),o=b.length-1;0<=o;o--)try{(s=b[o])(s.isolateScope?d:c,r,h,s.require&&x(s.directiveName,s.require,r,p),u)}catch(t){L(t,lt(r))}}r=r||{};for(var s,l,u,c,h,d,p=-Number.MAX_VALUE,w=r.controllerDirectives,$=r.newIsolateScopeDirective,k=r.templateDirective,f=r.nonTlbTranscludeDirective,g=!1,C=!1,E=r.hasElementTranscludeDirective,S=m.$$element=Xe(v),T=i,D=e,_=0,P=t.length;_<P;_++){var M=(l=t[_]).$$start,A=l.$$end;if(M&&(S=z(v,M,A)),c=tt,p>l.priority)break;if((d=l.scope)&&(s=s||l,l.templateUrl||(W("new/isolated scope",$,l,S),nt(d)&&($=l))),u=l.name,!l.templateUrl&&l.controller&&(d=l.controller,W("'"+u+"' controller",(w=w||{})[u],l,S),w[u]=l),(d=l.transclude)&&(g=!0,l.$$tlb||(W("transclusion",f,l,S),f=l),D="element"==d?(E=!0,p=l.priority,c=S,S=m.$$element=Xe(J.createComment(" "+u+": "+m[u]+" ")),v=S[0],G(n,st(c),v),F(c,e,p,T&&T.name,{nonTlbTranscludeDirective:f})):(c=Xe(dt(v)).contents(),S.empty(),F(c,e))),l.template)if(C=!0,W("template",k,l,S),d=rt((k=l).template)?l.template(S,m):l.template,d=Q(d),l.replace){if(T=l,c=ht(d)?[]:Xe(nn(d)),v=c[0],1!=c.length||1!==v.nodeType)throw Mn("tplrt","Template for directive '{0}' must have exactly one root element. {1}",u,"");G(n,S,v);var N={$attr:{}},O=q(v,[],N),I=t.splice(_+1,t.length-(_+1));$&&X(O),t=t.concat(O).concat(I),Y(m,N),P=t.length}else S.html(d);if(l.templateUrl)C=!0,W("template",k,l,S),(k=l).replace&&(T=l),a=U(t.splice(_,t.length-_),S,m,n,g&&D,y,b,{controllerDirectives:w,newIsolateScopeDirective:$,templateDirective:k,nonTlbTranscludeDirective:f}),P=t.length;else if(l.compile)try{rt(h=l.compile(S,m,D))?o(null,h,M,A):h&&o(h.pre,h.post,M,A)}catch(t){L(t,lt(S))}l.terminal&&(a.terminal=!0,p=Math.max(p,l.priority))}return a.scope=s&&!0===s.scope,a.transcludeOnThisElement=g,a.templateOnThisElement=C,a.transclude=D,r.hasElementTranscludeDirective=E,a}function X(t){for(var e=0,n=t.length;e<n;e++)t[e]=v(t[e],{$$isolateScope:!0})}function w(t,e,n,i,r,o,a){if(e===r)return null;var s=null;if(p.hasOwnProperty(e))for(var l,u=d.get(e+f),c=0,h=u.length;c<h;c++)try{l=u[c],(i===tt||i>l.priority)&&-1!=l.restrict.indexOf(n)&&(o&&(l=v(l,{$$start:o,$$end:a})),t.push(l),s=l)}catch(t){L(t)}return s}function Y(n,i){var r=i.$attr,o=n.$attr,a=n.$$element;et(n,function(t,e){"$"!=e.charAt(0)&&(i[e]&&i[e]!==t&&(t+=("style"===e?";":" ")+i[e]),n.$set(e,t,!0,r[e]))}),et(i,function(t,e){"class"==e?(V(a,t),n.class=(n.class?n.class+" ":"")+t):"style"==e?(a.attr("style",a.attr("style")+";"+t),n.style=(n.style?n.style+";":"")+t):"$"==e.charAt(0)||n.hasOwnProperty(e)||(n[e]=t,o[e]=r[e])})}function U(d,p,f,g,v,m,y,b){var x,w,$=[],k=p[0],C=d.shift(),E=M({},C,{templateUrl:null,transclude:null,replace:null,$$originalDirective:C}),S=rt(C.templateUrl)?C.templateUrl(p,f):C.templateUrl;return p.empty(),t.get(s.getTrustedResourceUrl(S),{cache:e}).success(function(t){var n,e,i,r;if(t=Q(t),C.replace){if(i=ht(t)?[]:Xe(nn(t)),n=i[0],1!=i.length||1!==n.nodeType)throw Mn("tplrt","Template for directive '{0}' must have exactly one root element. {1}",C.name,S);e={$attr:{}},G(g,p,n);var o=q(n,[],e);nt(C.scope)&&X(o),d=o.concat(d),Y(f,e)}else n=k,p.html(t);for(d.unshift(E),x=_(d,n,f,v,p,C,m,y,b),et(g,function(t,e){t==n&&(g[e]=p[0])}),w=T(p[0].childNodes,v);$.length;){var a=$.shift(),s=$.shift(),l=$.shift(),u=$.shift(),c=p[0];if(s!==k){var h=s.className;b.hasElementTranscludeDirective&&C.replace||(c=dt(n)),G(l,Xe(s),c),V(Xe(c),h)}r=x.transcludeOnThisElement?D(a,x.transclude,u):u,x(w,a,c,g,r)}$=null}).error(function(t,e,n,i){throw Mn("tpload","Failed to load template: {0}",i.url)}),function(t,e,n,i,r){var o=r;$?($.push(e),$.push(n),$.push(i),$.push(o)):(x.transcludeOnThisElement&&(o=D(e,x.transclude,r)),x(w,e,n,i,o))}}function $(t,e){var n=e.priority-t.priority;return 0!=n?n:t.name!==e.name?t.name<e.name?-1:1:t.index-e.index}function W(t,e,n,i){if(e)throw Mn("multidir","Multiple directives [{0}, {1}] asking for {2} on: {3}",e.name,n.name,t,lt(i))}function k(r,t,e,o){var a=B(e,!0);if(a){if("multiple"===o&&"SELECT"===We(r))throw Mn("selmulti","Binding to the 'multiple' attribute is not supported. Element: {0}",lt(r));t.push({priority:100,compile:function(){return{pre:function(t,e,n){var i=n.$$observers||(n.$$observers={});if(u.test(o))throw Mn("nodomevents","Interpolations for HTML DOM event attributes are disallowed.  Please use the ng- versions (such as ng-click instead of onclick) instead.");(a=B(n[o],!0,function(t,e){if("srcdoc"==e)return s.HTML;var n=We(t);return"xlinkHref"==e||"FORM"==n&&"action"==e||"IMG"!=n&&("src"==e||"ngSrc"==e)?s.RESOURCE_URL:void 0}(r,o)))&&(n[o]=a(t),(i[o]||(i[o]=[])).$$inter=!0,(n.$$observers&&n.$$observers[o].$$scope||t).$watch(a,function(t,e){"class"===o&&t!=e?n.$updateClass(t,e):n.$set(o,t)}))}}}})}}function G(t,e,n){var i,r,o=e[0],a=e.length,s=o.parentNode;if(t)for(i=0,r=t.length;i<r;i++)if(t[i]==o){t[i++]=n;for(var l=i,u=l+a-1,c=t.length;l<c;l++,u++)u<c?t[l]=t[u]:delete t[l];t.length-=a-1;break}s&&s.replaceChild(n,o);var h=J.createDocumentFragment();h.appendChild(o),n[Xe.expando]=o[Xe.expando];for(var d=1,p=e.length;d<p;d++){var f=e[d];Xe(f).remove(),h.appendChild(f),delete e[d]}e[0]=n,e.length=1}function Z(t,e){return M(function(){return t.apply(null,arguments)},t,e)}var K=function(t,e){this.$$element=t,this.$attr=e||{}};K.prototype={$normalize:_t,$addClass:function(t){t&&0<t.length&&r.addClass(this.$$element,t)},$removeClass:function(t){t&&0<t.length&&r.removeClass(this.$$element,t)},$updateClass:function(t,e){var n=Pt(t,e),i=Pt(e,t);0===n.length?r.removeClass(this.$$element,i):0===i.length?r.addClass(this.$$element,n):r.setClass(this.$$element,n,i)},$set:function(t,e,n,i){var r,o=mt(this.$$element[0],t);o&&(this.$$element.prop(t,e),i=o),this[t]=e,i?this.$attr[t]=i:(i=this.$attr[t])||(this.$attr[t]=i=ut(t,"-")),("A"===(r=We(this.$$element))&&"href"===t||"IMG"===r&&"src"===t)&&(this[t]=e=l(e,"src"===t)),!1!==n&&(null===e||e===tt?this.$$element.removeAttr(i):this.$$element.attr(i,e));var a=this.$$observers;a&&et(a[t],function(t){try{t(e)}catch(t){L(t)}})},$observe:function(t,e){var n=this,i=n.$$observers||(n.$$observers={}),r=i[t]||(i[t]=[]);return r.push(e),o.$evalAsync(function(){r.$$inter||e(n[t])}),e}};var i=B.startSymbol(),a=B.endSymbol(),Q="{{"==i||"}}"==a?m:function(t){return t.replace(/\{\{/g,i).replace(/}}/g,a)},C=/^ngAttr[A-Z]/;return F}]}function _t(t){return q(t.replace(An,""))}function Pt(t,e){var n="",i=t.split(/\s+/),r=e.split(/\s+/);t:for(var o=0;o<i.length;o++){for(var a=i[o],s=0;s<r.length;s++)if(a==r[s])continue t;n+=(0<n.length?" ":"")+a}return n}function Mt(){var l={},u=/^(\S+)(\s+as\s+(\w+))?$/;this.register=function(t,e){L(t,"controller"),nt(t)?M(l,t):l[t]=e},this.$get=["$injector","$window",function(a,s){return function(t,e){var n,i,r,o;if(it(t)&&(r=(i=t.match(u))[1],o=i[3],B(t=l.hasOwnProperty(r)?l[r]:j(e.$scope,r,!0)||j(s,r,!0),r,!0)),n=a.instantiate(t,e),o){if(!e||"object"!=typeof e.$scope)throw k("$controller")("noscp","Cannot export controller '{0}' as '{1}'! No $scope object provided via `locals`.",r||t.name,o);e.$scope[o]=n}return n}}]}function At(){this.$get=["$window",function(t){return Xe(t.document)}]}function Nt(){this.$get=["$log",function(n){return function(t,e){n.error.apply(n,arguments)}}]}function Ot(t){var e,n,i,r={};return t&&et(t.split("\n"),function(t){i=t.indexOf(":"),e=Ve(nn(t.substr(0,i))),n=nn(t.substr(i+1)),e&&(r[e]=r[e]?r[e]+", "+n:n)}),r}function It(e){var n=nt(e)?e:tt;return function(t){return n||(n=Ot(e)),t?n[Ve(t)]||null:n}}function Bt(e,n,t){return rt(t)?t(e,n):(et(t,function(t){e=t(e,n)}),e)}function Lt(t){return 200<=t&&t<300}function jt(){var e=/^\s*(\[|\{[^\{])/,n=/[\}\]]\s*$/,i=/^\)\]\}',?\n/,t={"Content-Type":"application/json;charset=utf-8"},m=this.defaults={transformResponse:[function(t){return it(t)&&(t=t.replace(i,""),e.test(t)&&n.test(t)&&(t=a(t))),t}],transformRequest:[function(t){return!nt(t)||function(t){return"[object File]"===Ke.call(t)}(t)||function(t){return"[object Blob]"===Ke.call(t)}(t)?t:A(t)}],headers:{common:{Accept:"application/json, text/plain, */*"},post:ot(t),put:ot(t),patch:ot(t)},xsrfCookieName:"XSRF-TOKEN",xsrfHeaderName:"X-XSRF-TOKEN"},r=this.interceptors=[],o=this.responseInterceptors=[];this.$get=["$httpBackend","$browser","$cacheFactory","$rootScope","$q","$injector",function(h,d,t,p,f,i){function g(t){function n(t){var e=M({},t,{data:Bt(t.data,t.headers,i.transformResponse)});return Lt(t.status)?e:f.reject(e)}var i={method:"get",transformRequest:m.transformRequest,transformResponse:m.transformResponse},r=function(t){var e,n,i,r,o,a=m.headers,s=M({},t.headers);a=M({},a.common,a[Ve(t.method)]);t:for(e in a){for(i in n=Ve(e),s)if(Ve(i)===n)continue t;s[e]=a[e]}return et(r=s,function(t,e){rt(t)&&(null!=(o=t())?r[e]=o:delete r[e])}),s}(t);M(i,t),i.headers=r,i.method=ze(i.method);var e=[function(t){r=t.headers;var e=Bt(t.data,It(r),t.transformRequest);return S(e)&&et(r,function(t,e){"content-type"===Ve(e)&&delete r[e]}),S(t.withCredentials)&&!S(m.withCredentials)&&(t.withCredentials=m.withCredentials),function(r,t,e){function o(t,e,n,i){(Lt(e=Math.max(e,0))?s.resolve:s.reject)({data:t,status:e,headers:It(n),config:r,statusText:i})}function n(){var t=C(g.pendingRequests,r);-1!==t&&g.pendingRequests.splice(t,1)}var a,i,s=f.defer(),l=s.promise,u=function(t,e){if(!e)return t;var n=[];return function(t,e,n){for(var i=R(t),r=0;r<i.length;r++)e.call(n,t[i[r]],i[r])}(e,function(t,e){null===t||S(t)||(en(t)||(t=[t]),et(t,function(t){nt(t)&&(t=b(t)?t.toISOString():A(t)),n.push(O(e)+"="+O(t))}))}),0<n.length&&(t+=(-1==t.indexOf("?")?"?":"&")+n.join("&")),t}(r.url,r.params);if(g.pendingRequests.push(r),l.then(n,n),!r.cache&&!m.cache||!1===r.cache||"GET"!==r.method&&"JSONP"!==r.method||(a=nt(r.cache)?r.cache:nt(m.cache)?m.cache:v),a)if(F(i=a.get(u))){if(x(i))return i.then(n,n),i;en(i)?o(i[1],i[0],ot(i[2]),i[3]):o(i,200,{},"OK")}else a.put(u,l);if(S(i)){var c=xe(r.url)?d.cookies()[r.xsrfCookieName||m.xsrfCookieName]:tt;c&&(e[r.xsrfHeaderName||m.xsrfHeaderName]=c),h(r.method,u,t,function(t,e,n,i){a&&(Lt(t)?a.put(u,[t,e,Ot(n),i]):a.remove(u)),o(e,t,n,i),p.$$phase||p.$apply()},e,r.timeout,r.withCredentials,r.responseType)}return l}(t,e,r).then(n,n)},tt],o=f.when(i);for(et(l,function(t){(t.request||t.requestError)&&e.unshift(t.request,t.requestError),(t.response||t.responseError)&&e.push(t.response,t.responseError)});e.length;){var a=e.shift(),s=e.shift();o=o.then(a,s)}return o.success=function(e){return o.then(function(t){e(t.data,t.status,t.headers,i)}),o},o.error=function(e){return o.then(null,function(t){e(t.data,t.status,t.headers,i)}),o},o}var v=t("$http"),l=[];return et(r,function(t){l.unshift(it(t)?i.get(t):i.invoke(t))}),et(o,function(t,e){var n=it(t)?i.get(t):i.invoke(t);l.splice(e,0,{response:function(t){return n(f.when(t))},responseError:function(t){return n(f.reject(t))}})}),g.pendingRequests=[],function(t){et(arguments,function(n){g[n]=function(t,e){return g(M(e||{},{method:n,url:t}))}})}("get","delete","head","jsonp"),function(t){et(arguments,function(i){g[i]=function(t,e,n){return g(M(n||{},{method:i,url:t,data:e}))}})}("post","put","patch"),g.defaults=m,g}]}function Rt(t){if(He<=8&&(!t.match(/^(get|post|head|put|delete|options)$/i)||!f.XMLHttpRequest))return new f.ActiveXObject("Microsoft.XMLHTTP");if(f.XMLHttpRequest)return new f.XMLHttpRequest;throw k("$httpBackend")("noxhr","This browser does not support XMLHttpRequest.")}function Ft(){this.$get=["$browser","$window","$document",function(t,e,n){return function(g,v,m,y,b){return function(t,o,e,i,n,r,a,s){function l(){c=-1,d&&d(),p&&p.abort()}function u(t,e,n,i,r){f&&m.cancel(f),d=p=null,0===e&&(e=n?200:"file"==be(o).protocol?404:0),t(e=1223===e?204:e,n,i,r=r||""),g.$$completeOutstandingRequest(E)}var c;if(g.$$incOutstandingRequestCount(),o=o||g.url(),"jsonp"==Ve(t)){var h="_"+(y.counter++).toString(36);y[h]=function(t){y[h].data=t,y[h].called=!0};var d=function(t,i,r){var o=b.createElement("script"),a=null;return o.type="text/javascript",o.src=t,o.async=!0,a=function(t){cn(o,"load",a),cn(o,"error",a),b.body.removeChild(o),o=null;var e=-1,n="unknown";t&&("load"!==t.type||y[i].called||(t={type:"error"}),n=t.type,e="error"===t.type?404:200),r&&r(e,n)},un(o,"load",a),un(o,"error",a),He<=8&&(o.onreadystatechange=function(){it(o.readyState)&&/loaded|complete/.test(o.readyState)&&(o.onreadystatechange=null,a({type:"load"}))}),b.body.appendChild(o),a}(o.replace("JSON_CALLBACK","angular.callbacks."+h),h,function(t,e){u(i,t,y[h].data,"",e),y[h]=E})}else{var p=v(t);if(p.open(t,o,!0),et(n,function(t,e){F(t)&&p.setRequestHeader(e,t)}),p.onreadystatechange=function(){if(p&&4==p.readyState){var t=null,e=null,n="";-1!==c&&(t=p.getAllResponseHeaders(),e="response"in p?p.response:p.responseText),-1===c&&He<10||(n=p.statusText),u(i,c||p.status,e,t,n)}},a&&(p.withCredentials=!0),s)try{p.responseType=s}catch(t){if("json"!==s)throw t}p.send(e||null)}if(0<r)var f=m(l,r);else x(r)&&r.then(l)}}(t,Rt,t.defer,e.angular.callbacks,n[0])}]}function Vt(){var m="{{",y="}}";this.startSymbol=function(t){return t?(m=t,this):m},this.endSymbol=function(t){return t?(y=t,this):y},this.$get=["$parse","$exceptionHandler","$sce",function(d,p,f){function t(o,t,a){for(var e,n,i,r,s=0,l=[],u=o.length,c=!1,h=[];s<u;)-1!=(e=o.indexOf(m,s))&&-1!=(n=o.indexOf(y,e+g))?(s!=e&&l.push(o.substring(s,e)),l.push(i=d(r=o.substring(e+g,n))),i.exp=r,s=n+v,c=!0):(s!=u&&l.push(o.substring(s)),s=u);if((u=l.length)||(l.push(""),u=1),a&&1<l.length)throw Nn("noconcat","Error while interpolating: {0}\nStrict Contextual Escaping disallows interpolations that concatenate multiple expressions when a trusted value is required.  See http://docs.angularjs.org/api/ng.$sce",o);return!t||c?(h.length=u,(i=function(t){try{for(var e,n=0,i=u;n<i;n++){if("function"==typeof(e=l[n]))if(e=e(t),null==(e=a?f.getTrusted(a,e):f.valueOf(e)))e="";else switch(typeof e){case"string":break;case"number":e=""+e;break;default:e=A(e)}h[n]=e}return h.join("")}catch(t){var r=Nn("interr","Can't interpolate: {0}\n{1}",o,t.toString());p(r)}}).exp=o,i.parts=l,i):void 0}var g=m.length,v=y.length;return t.startSymbol=function(){return m},t.endSymbol=function(){return y},t}]}function qt(){this.$get=["$rootScope","$window","$q",function(c,h,d){function t(t,e,n,i){var r=h.setInterval,o=h.clearInterval,a=d.defer(),s=a.promise,l=0,u=F(i)&&!i;return n=F(n)?n:0,s.then(null,null,t),s.$$intervalId=r(function(){a.notify(l++),0<n&&n<=l&&(a.resolve(l),o(s.$$intervalId),delete p[s.$$intervalId]),u||c.$apply()},e),p[s.$$intervalId]=a,s}var p={};return t.cancel=function(t){return!!(t&&t.$$intervalId in p)&&(p[t.$$intervalId].reject("canceled"),h.clearInterval(t.$$intervalId),delete p[t.$$intervalId],!0)},t}]}function zt(){this.$get=function(){return{id:"en-us",NUMBER_FORMATS:{DECIMAL_SEP:".",GROUP_SEP:",",PATTERNS:[{minInt:1,minFrac:0,maxFrac:3,posPre:"",posSuf:"",negPre:"-",negSuf:"",gSize:3,lgSize:3},{minInt:1,minFrac:2,maxFrac:2,posPre:"¤",posSuf:"",negPre:"(¤",negSuf:")",gSize:3,lgSize:3}],CURRENCY_SYM:"$"},DATETIME_FORMATS:{MONTH:"January,February,March,April,May,June,July,August,September,October,November,December".split(","),SHORTMONTH:"Jan,Feb,Mar,Apr,May,Jun,Jul,Aug,Sep,Oct,Nov,Dec".split(","),DAY:"Sunday,Monday,Tuesday,Wednesday,Thursday,Friday,Saturday".split(","),SHORTDAY:"Sun,Mon,Tue,Wed,Thu,Fri,Sat".split(","),AMPMS:["AM","PM"],medium:"MMM d, y h:mm:ss a",short:"M/d/yy h:mm a",fullDate:"EEEE, MMMM d, y",longDate:"MMMM d, y",mediumDate:"MMM d, y",shortDate:"M/d/yy",mediumTime:"h:mm:ss a",shortTime:"h:mm a"},pluralCat:function(t){return 1===t?"one":"other"}}}}function Ht(t){for(var e=t.split("/"),n=e.length;n--;)e[n]=w(e[n]);return e.join("/")}function Xt(t,e,n){var i=be(t,n);e.$$protocol=i.protocol,e.$$host=i.hostname,e.$$port=$(i.port)||In[i.protocol]||null}function Yt(t,e,n){var i="/"!==t.charAt(0);i&&(t="/"+t);var r=be(t,n);e.$$path=decodeURIComponent(i&&"/"===r.pathname.charAt(0)?r.pathname.substring(1):r.pathname),e.$$search=s(r.search),e.$$hash=decodeURIComponent(r.hash),e.$$path&&"/"!=e.$$path.charAt(0)&&(e.$$path="/"+e.$$path)}function Ut(t,e){return 0===e.indexOf(t)?e.substr(t.length):void 0}function Wt(t){var e=t.indexOf("#");return-1==e?t:t.substr(0,e)}function Gt(t){return t.substr(0,Wt(t).lastIndexOf("/")+1)}function Zt(o,a){this.$$html5=!0,a=a||"";var s=Gt(o);Xt(o,this,o),this.$$parse=function(t){var e=Ut(s,t);if(!it(e))throw Bn("ipthprfx",'Invalid url "{0}", missing path prefix "{1}".',t,s);Yt(e,this,o),this.$$path||(this.$$path="/"),this.$$compose()},this.$$compose=function(){var t=i(this.$$search),e=this.$$hash?"#"+w(this.$$hash):"";this.$$url=Ht(this.$$path)+(t?"?"+t:"")+e,this.$$absUrl=s+this.$$url.substr(1)},this.$$parseLinkUrl=function(t,e){var n,i,r;return(n=Ut(o,t))!==tt?r=(n=Ut(a,i=n))!==tt?s+(Ut("/",n)||n):o+i:(n=Ut(s,t))!==tt?r=s+n:s==t+"/"&&(r=s),r&&this.$$parse(r),!!r}}function Kt(l,u){var c=Gt(l);Xt(l,this,l),this.$$parse=function(t){var e,n,i,r,o,a=Ut(l,t)||Ut(c,t),s="#"==a.charAt(0)?Ut(u,a):this.$$html5?a:"";if(!it(s))throw Bn("ihshprfx",'Invalid url "{0}", missing hash prefix "{1}".',t,u);Yt(s,this,l),this.$$path=(e=this.$$path,i=l,o=/^\/[A-Z]:(\/.*)/,0===(n=s).indexOf(i)&&(n=n.replace(i,"")),o.exec(n)?e:(r=o.exec(e))?r[1]:e),this.$$compose()},this.$$compose=function(){var t=i(this.$$search),e=this.$$hash?"#"+w(this.$$hash):"";this.$$url=Ht(this.$$path)+(t?"?"+t:"")+e,this.$$absUrl=l+(this.$$url?u+this.$$url:"")},this.$$parseLinkUrl=function(t,e){return Wt(l)==Wt(t)&&(this.$$parse(t),!0)}}function Qt(r,o){this.$$html5=!0,Kt.apply(this,arguments);var a=Gt(r);this.$$parseLinkUrl=function(t,e){var n,i;return r==Wt(t)?n=t:(i=Ut(a,t))?n=r+o+i:a===t+"/"&&(n=a),n&&this.$$parse(n),!!n},this.$$compose=function(){var t=i(this.$$search),e=this.$$hash?"#"+w(this.$$hash):"";this.$$url=Ht(this.$$path)+(t?"?"+t:"")+e,this.$$absUrl=r+o+this.$$url}}function Jt(t){return function(){return this[t]}}function te(e,n){return function(t){return S(t)?this[e]:(this[e]=n(t),this.$$compose(),this)}}function ee(){var d="",p=!1;this.hashPrefix=function(t){return F(t)?(d=t,this):d},this.html5Mode=function(t){return F(t)?(p=t,this):p},this.$get=["$rootScope","$browser","$sniffer","$rootElement",function(r,o,t,a){function n(t){r.$broadcast("$locationChangeSuccess",s.absUrl(),t)}var s,e,i,l=o.baseHref(),u=o.url();e=p?(i=function(t){return t.substring(0,t.indexOf("/",t.indexOf("//")+2))}(u)+(l||"/"),t.history?Zt:Qt):(i=Wt(u),Kt),(s=new e(i,"#"+d)).$$parseLinkUrl(u,u);var c=/^\s*(javascript|mailto):/i;a.on("click",function(t){if(!t.ctrlKey&&!t.metaKey&&2!=t.which){for(var e=Xe(t.target);"a"!==Ve(e[0].nodeName);)if(e[0]===a[0]||!(e=e.parent())[0])return;var n=e.prop("href"),i=e.attr("href")||e.attr("xlink:href");nt(n)&&"[object SVGAnimatedString]"===n.toString()&&(n=be(n.animVal).href),c.test(n)||!n||e.attr("target")||t.isDefaultPrevented()||s.$$parseLinkUrl(n,i)&&(t.preventDefault(),s.absUrl()!=o.url()&&(r.$apply(),f.angular["ff-684208-preventDefault"]=!0))}}),s.absUrl()!=u&&o.url(s.absUrl(),!0),o.onUrlChange(function(e){s.absUrl()!=e&&(r.$evalAsync(function(){var t=s.absUrl();s.$$parse(e),r.$broadcast("$locationChangeStart",e,t).defaultPrevented?(s.$$parse(t),o.url(t)):n(t)}),r.$$phase||r.$digest())});var h=0;return r.$watch(function(){var t=o.url(),e=s.$$replace;return h&&t==s.absUrl()||(h++,r.$evalAsync(function(){r.$broadcast("$locationChangeStart",s.absUrl(),t).defaultPrevented?s.$$parse(t):(o.url(s.absUrl(),e),n(t))})),s.$$replace=!1,h}),s}]}function ne(){var n=!0,i=this;this.debugEnabled=function(t){return F(t)?(n=t,this):n},this.$get=["$window",function(r){function t(t){var n=r.console||{},i=n[t]||n.log||E,e=!1;try{e=!!i.apply}catch(t){}return e?function(){var e=[];return et(arguments,function(t){e.push(function(t){return t instanceof Error&&(t.stack?t=t.message&&-1===t.stack.indexOf(t.message)?"Error: "+t.message+"\n"+t.stack:t.stack:t.sourceURL&&(t=t.message+"\n"+t.sourceURL+":"+t.line)),t}(t))}),i.apply(n,e)}:function(t,e){i(t,null==e?"":e)}}return{log:t("log"),info:t("info"),warn:t("warn"),error:t("error"),debug:(e=t("debug"),function(){n&&e.apply(i,arguments)})};var e}]}function ie(t,e){if("__defineGetter__"===t||"__defineSetter__"===t||"__lookupGetter__"===t||"__lookupSetter__"===t||"__proto__"===t)throw jn("isecfld","Attempting to access a disallowed field in Angular expressions! Expression: {0}",e);return t}function re(t,e){if(t){if(t.constructor===t)throw jn("isecfn","Referencing Function in Angular expressions is disallowed! Expression: {0}",e);if(t.document&&t.location&&t.alert&&t.setInterval)throw jn("isecwindow","Referencing the Window in Angular expressions is disallowed! Expression: {0}",e);if(t.children&&(t.nodeName||t.prop&&t.attr&&t.find))throw jn("isecdom","Referencing DOM nodes in Angular expressions is disallowed! Expression: {0}",e);if(t===Object)throw jn("isecobj","Referencing Object in Angular expressions is disallowed! Expression: {0}",e)}return t}function oe(t,e,n,i,r){re(t,i),r=r||{};for(var o,a=e.split(".");1<a.length;0){var s=re(t[o=ie(a.shift(),i)],i);s||(s={},t[o]=s),(t=s).then&&r.unwrapPromises&&(Ln(i),"$$v"in t||function(e){e.then(function(t){e.$$v=t})}(t),t.$$v===tt&&(t.$$v={}),t=t.$$v)}return re(t[o=ie(a.shift(),i)],i),t[o]=n}function ae(t){return"constructor"==t}function se(r,o,a,s,l,u,t){ie(r,u),ie(o,u),ie(a,u),ie(s,u),ie(l,u);function e(t){return re(t,u)}var n=t.expensiveChecks,c=n||ae(r)?e:m,h=n||ae(o)?e:m,d=n||ae(a)?e:m,p=n||ae(s)?e:m,f=n||ae(l)?e:m;return t.unwrapPromises?function(t,e){var n,i=e&&e.hasOwnProperty(r)?e:t;return null==i?i:((i=c(i[r]))&&i.then&&(Ln(u),"$$v"in i||((n=i).$$v=tt,n.then(function(t){n.$$v=c(t)})),i=c(i.$$v)),o?null==i?tt:((i=h(i[o]))&&i.then&&(Ln(u),"$$v"in i||((n=i).$$v=tt,n.then(function(t){n.$$v=h(t)})),i=h(i.$$v)),a?null==i?tt:((i=d(i[a]))&&i.then&&(Ln(u),"$$v"in i||((n=i).$$v=tt,n.then(function(t){n.$$v=d(t)})),i=d(i.$$v)),s?null==i?tt:((i=p(i[s]))&&i.then&&(Ln(u),"$$v"in i||((n=i).$$v=tt,n.then(function(t){n.$$v=p(t)})),i=p(i.$$v)),l?null==i?tt:((i=f(i[l]))&&i.then&&(Ln(u),"$$v"in i||((n=i).$$v=tt,n.then(function(t){n.$$v=f(t)})),i=f(i.$$v)),i):i):i):i):i)}:function(t,e){var n=e&&e.hasOwnProperty(r)?e:t;return null==n?n:(n=c(n[r]),o?null==n?tt:(n=h(n[o]),a?null==n?tt:(n=d(n[a]),s?null==n?tt:(n=p(n[s]),l?null==n?tt:n=f(n[l]):n):n):n):n)}}function le(t,r,o){var a=r.expensiveChecks,e=a?Wn:Un;if(e.hasOwnProperty(t))return e[t];var n,s=t.split("."),l=s.length;if(r.csp)n=l<6?se(s[0],s[1],s[2],s[3],s[4],o,r):function(t,e){for(var n,i=0;n=se(s[i++],s[i++],s[i++],s[i++],s[i++],o,r)(t,e),e=tt,t=n,i<l;);return n};else{var u="var p;\n";a&&(u+="s = eso(s, fe);\nl = eso(l, fe);\n");var c=a;et(s,function(t,e){ie(t,o);var n=(e?"s":'((l&&l.hasOwnProperty("'+t+'"))?l:s)')+'["'+t+'"]',i=a||ae(t);i&&(n="eso("+n+", fe)",c=!0),u+="if(s == null) return undefined;\ns="+n+";\n",r.unwrapPromises&&(u+='if (s && s.then) {\n pw("'+o.replace(/(["\r\n])/g,"\\$1")+'");\n if (!("$$v" in s)) {\n p=s;\n p.$$v = undefined;\n p.then(function(v) {p.$$v='+(i?"eso(v)":"v")+";});\n}\n s="+(i?"eso(s.$$v)":"s.$$v")+"\n}\n")}),u+="return s;";var i=new Function("s","l","pw","eso","fe",u);i.toString=y(u),(c||r.unwrapPromises)&&(i=function(n,i){return function(t,e){return n(t,e,Ln,re,i)}}(i,o)),n=i}return"hasOwnProperty"!==t&&(e[t]=n),n}function ue(){var l={},u={},c={csp:!1,unwrapPromises:!1,logPromiseWarnings:!0,expensiveChecks:!1};this.unwrapPromises=function(t){return F(t)?(c.unwrapPromises=!!t,this):c.unwrapPromises},this.logPromiseWarnings=function(t){return F(t)?(c.logPromiseWarnings=t,this):c.logPromiseWarnings},this.$get=["$filter","$sniffer","$log",function(a,t,e){c.csp=t.csp;var s={csp:c.csp,unwrapPromises:c.unwrapPromises,logPromiseWarnings:c.logPromiseWarnings,expensiveChecks:!0};return Ln=function(t){c.logPromiseWarnings&&!Rn.hasOwnProperty(t)&&(Rn[t]=!0,e.warn("[$parse] Promise found in the expression `"+t+"`. Automatic unwrapping of promises in Angular expressions is deprecated."))},function(t,e){var n;switch(typeof t){case"string":var i=e?u:l;if(i.hasOwnProperty(t))return i[t];var r=e?s:c,o=new Xn(r);return n=new Yn(o,a,r).parse(t),"hasOwnProperty"!==t&&(i[t]=n),n;case"function":return t;default:return E}}}]}function ce(){this.$get=["$rootScope","$exceptionHandler",function(e,t){return function(u,c){function h(t){return t}function d(t){return g(t)}var p=function(){var s,e,l=[];return e={resolve:function(t){if(l){var i=l;l=tt,s=f(t),i.length&&u(function(){for(var t,e=0,n=i.length;e<n;e++)t=i[e],s.then(t[0],t[1],t[2])})}},reject:function(t){e.resolve(n(t))},notify:function(n){if(l){var i=l;l.length&&u(function(){for(var t=0,e=i.length;t<e;t++)i[t][2](n)})}},promise:{then:function(e,n,i){function t(t){try{a.resolve((rt(e)?e:h)(t))}catch(t){a.reject(t),c(t)}}function r(t){try{a.resolve((rt(n)?n:d)(t))}catch(t){a.reject(t),c(t)}}function o(t){try{a.notify((rt(i)?i:h)(t))}catch(t){c(t)}}var a=p();return l?l.push([t,r,o]):s.then(t,r,o),a.promise},catch:function(t){return this.then(null,t)},finally:function(i){function r(t,e){var n=p();return e?n.resolve(t):n.reject(t),n.promise}function e(t,e){var n=null;try{n=(i||h)()}catch(t){return r(t,!1)}return x(n)?n.then(function(){return r(t,e)},function(t){return r(t,!1)}):r(t,e)}return this.then(function(t){return e(t,!0)},function(t){return e(t,!1)})}}}},f=function(n){return x(n)?n:{then:function(t){var e=p();return u(function(){e.resolve(t(n))}),e.promise}}},g=function(t){var e=p();return e.reject(t),e.promise},n=function(i){return{then:function(t,e){var n=p();return u(function(){try{n.resolve((rt(e)?e:d)(i))}catch(t){n.reject(t),c(t)}}),n.promise}}};return{defer:p,reject:g,when:function(t,e,n,i){function r(t){try{return(rt(e)?e:h)(t)}catch(t){return c(t),g(t)}}function o(t){try{return(rt(n)?n:d)(t)}catch(t){return c(t),g(t)}}function a(t){try{return(rt(i)?i:h)(t)}catch(t){c(t)}}var s,l=p();return u(function(){f(t).then(function(t){s||(s=!0,l.resolve(f(t).then(r,o,a)))},function(t){s||(s=!0,l.resolve(o(t)))},function(t){s||l.notify(a(t))})}),l.promise},all:function(t){var n=p(),i=0,r=en(t)?[]:{};return et(t,function(t,e){i++,f(t).then(function(t){r.hasOwnProperty(e)||(r[e]=t,--i||n.resolve(r))},function(t){r.hasOwnProperty(e)||n.reject(t)})}),0===i&&n.resolve(r),n.promise}}}(function(t){e.$evalAsync(t)},t)}]}function he(){this.$get=["$window","$timeout",function(t,n){var i=t.requestAnimationFrame||t.webkitRequestAnimationFrame||t.mozRequestAnimationFrame,r=t.cancelAnimationFrame||t.webkitCancelAnimationFrame||t.mozCancelAnimationFrame||t.webkitCancelRequestAnimationFrame,e=!!i,o=e?function(t){var e=i(t);return function(){r(e)}}:function(t){var e=n(t,16.66,!1);return function(){n.cancel(e)}};return o.supported=e,o}]}function de(){var x=10,w=k("$rootScope"),$=null;this.digestTtl=function(t){return arguments.length&&(x=t),x},this.$get=["$injector","$exceptionHandler","$parse","$browser",function(t,g,f,v){function n(){this.$id=r(),this.$$phase=this.$parent=this.$$watchers=this.$$nextSibling=this.$$prevSibling=this.$$childHead=this.$$childTail=null,this.this=this.$root=this,this.$$destroyed=!1,this.$$asyncQueue=[],this.$$postDigestQueue=[],this.$$listeners={},this.$$listenerCount={},this.$$isolateBindings={}}function m(t){if(e.$$phase)throw w("inprog","{0} already in progress",e.$$phase);e.$$phase=t}function y(){e.$$phase=null}function l(t,e){var n=f(t);return B(n,e),n}function o(t,e,n){for(;t.$$listenerCount[n]-=e,0===t.$$listenerCount[n]&&delete t.$$listenerCount[n],t=t.$parent;);}function b(){}n.prototype={constructor:n,$new:function(t){var e;return t?((e=new n).$root=this.$root,e.$$asyncQueue=this.$$asyncQueue,e.$$postDigestQueue=this.$$postDigestQueue):(this.$$childScopeClass||(this.$$childScopeClass=function(){this.$$watchers=this.$$nextSibling=this.$$childHead=this.$$childTail=null,this.$$listeners={},this.$$listenerCount={},this.$id=r(),this.$$childScopeClass=null},this.$$childScopeClass.prototype=this),e=new this.$$childScopeClass),(e.this=e).$parent=this,e.$$prevSibling=this.$$childTail,this.$$childHead?(this.$$childTail.$$nextSibling=e,this.$$childTail=e):this.$$childHead=this.$$childTail=e,e},$watch:function(t,e,n){var i=l(t,"watch"),r=this.$$watchers,o={fn:e,last:b,get:i,exp:t,eq:!!n};if($=null,!rt(e)){var a=l(e||E,"listener");o.fn=function(t,e,n){a(n)}}if("string"==typeof t&&i.constant){var s=o.fn;o.fn=function(t,e,n){s.call(this,t,e,n),p(r,o)}}return r||(r=this.$$watchers=[]),r.unshift(o),function(){p(r,o),$=null}},$watchCollection:function(t,n){var i,r,o,a=this,s=1<n.length,l=0,u=f(t),c=[],h={},d=!0,p=0;return this.$watch(function(){var t,e;if(nt(i=u(a)))if(P(i)){r!==c&&(p=(r=c).length=0,l++),t=i.length,p!==t&&(l++,r.length=p=t);for(var n=0;n<t;n++)r[n]!=r[n]&&i[n]!=i[n]||r[n]===i[n]||(l++,r[n]=i[n])}else{for(e in r!==h&&(r=h={},p=0,l++),t=0,i)i.hasOwnProperty(e)&&(t++,r.hasOwnProperty(e)?r[e]!=r[e]&&i[e]!=i[e]||r[e]===i[e]||(l++,r[e]=i[e]):(p++,r[e]=i[e],l++));if(t<p)for(e in l++,r)r.hasOwnProperty(e)&&!i.hasOwnProperty(e)&&(p--,delete r[e])}else r!==i&&(r=i,l++);return l},function(){if(d?(d=!1,n(i,i,a)):n(i,o,a),s)if(nt(i))if(P(i)){o=new Array(i.length);for(var t=0;t<i.length;t++)o[t]=i[t]}else for(var e in o={},i)qe.call(i,e)&&(o[e]=i[e]);else o=i})},$digest:function(){var t,e,n,i,r,o,a,s,l,u,c,h=this.$$asyncQueue,d=this.$$postDigestQueue,p=x,f=[];m("$digest"),v.$$checkUrlChange(),$=null;do{for(o=!1,s=this;h.length;){try{(c=h.shift()).scope.$eval(c.expression)}catch(t){y(),g(t)}$=null}t:do{if(i=s.$$watchers)for(r=i.length;r--;)try{if(t=i[r])if((e=t.get(s))===(n=t.last)||(t.eq?at(e,n):"number"==typeof e&&"number"==typeof n&&isNaN(e)&&isNaN(n))){if(t===$){o=!1;break t}}else o=!0,($=t).last=t.eq?T(e,null):e,t.fn(e,n===b?e:n,s),p<5&&(f[l=4-p]||(f[l]=[]),u=rt(t.exp)?"fn: "+(t.exp.name||t.exp.toString()):t.exp,u+="; newVal: "+A(e)+"; oldVal: "+A(n),f[l].push(u))}catch(t){y(),g(t)}if(!(a=s.$$childHead||s!==this&&s.$$nextSibling))for(;s!==this&&!(a=s.$$nextSibling);)s=s.$parent}while(s=a);if((o||h.length)&&!p--)throw y(),w("infdig","{0} $digest() iterations reached. Aborting!\nWatchers fired in the last 5 iterations: {1}",x,A(f))}while(o||h.length);for(y();d.length;)try{d.shift()()}catch(t){g(t)}},$destroy:function(){if(!this.$$destroyed){var t=this.$parent;this.$broadcast("$destroy"),this.$$destroyed=!0,this!==e&&(et(this.$$listenerCount,_(null,o,this)),t.$$childHead==this&&(t.$$childHead=this.$$nextSibling),t.$$childTail==this&&(t.$$childTail=this.$$prevSibling),this.$$prevSibling&&(this.$$prevSibling.$$nextSibling=this.$$nextSibling),this.$$nextSibling&&(this.$$nextSibling.$$prevSibling=this.$$prevSibling),this.$parent=this.$$nextSibling=this.$$prevSibling=this.$$childHead=this.$$childTail=this.$root=null,this.$$listeners={},this.$$watchers=this.$$asyncQueue=this.$$postDigestQueue=[],this.$destroy=this.$digest=this.$apply=E,this.$on=this.$watch=function(){return E})}},$eval:function(t,e){return f(t)(this,e)},$evalAsync:function(t){e.$$phase||e.$$asyncQueue.length||v.defer(function(){e.$$asyncQueue.length&&e.$digest()}),this.$$asyncQueue.push({scope:this,expression:t})},$$postDigest:function(t){this.$$postDigestQueue.push(t)},$apply:function(t){try{return m("$apply"),this.$eval(t)}catch(t){g(t)}finally{y();try{e.$digest()}catch(t){throw g(t),t}}},$on:function(e,n){var i=this.$$listeners[e];i||(this.$$listeners[e]=i=[]),i.push(n);for(var t=this;t.$$listenerCount[e]||(t.$$listenerCount[e]=0),t.$$listenerCount[e]++,t=t.$parent;);var r=this;return function(){var t=C(i,n);-1!==t&&(i[t]=null,o(r,1,e))}},$emit:function(t,e){var n,i,r,o=[],a=this,s=!1,l={name:t,targetScope:a,stopPropagation:function(){s=!0},preventDefault:function(){l.defaultPrevented=!0},defaultPrevented:!1},u=D([l],arguments,1);do{for(n=a.$$listeners[t]||o,l.currentScope=a,i=0,r=n.length;i<r;i++)if(n[i])try{n[i].apply(null,u)}catch(t){g(t)}else n.splice(i,1),i--,r--;if(s)return l;a=a.$parent}while(a);return l},$broadcast:function(t,e){for(var n,i,r,o=this,a=this,s={name:t,targetScope:this,preventDefault:function(){s.defaultPrevented=!0},defaultPrevented:!1},l=D([s],arguments,1);o=a;){for(i=0,r=(n=(s.currentScope=o).$$listeners[t]||[]).length;i<r;i++)if(n[i])try{n[i].apply(null,l)}catch(t){g(t)}else n.splice(i,1),i--,r--;if(!(a=o.$$listenerCount[t]&&o.$$childHead||o!==this&&o.$$nextSibling))for(;o!==this&&!(a=o.$$nextSibling);)o=o.$parent}return s}};var e=new n;return e}]}function pe(){var r=/^\s*(https?|ftp|mailto|tel|file):/,o=/^\s*((https?|ftp|file):|data:image\/)/;this.aHrefSanitizationWhitelist=function(t){return F(t)?(r=t,this):r},this.imgSrcSanitizationWhitelist=function(t){return F(t)?(o=t,this):o},this.$get=function(){return function(t,e){var n,i=e?o:r;return He&&!(8<=He)||(""===(n=be(t).href)||n.match(i))?t:"unsafe:"+n}}}function fe(t){var e=[];return F(t)&&et(t,function(t){e.push(function(t){if("self"===t)return t;if(it(t)){if(-1<t.indexOf("***"))throw Gn("iwcard","Illegal sequence *** in string matcher.  String: {0}",t);return t=function(t){return t.replace(/([-()\[\]{}+?*.$\^|,:#<!\\])/g,"\\$1").replace(/\x08/g,"\\x08")}(t).replace("\\*\\*",".*").replace("\\*","[^:/.?&;]*"),new RegExp("^"+t+"$")}if(c(t))return new RegExp("^"+t.source+"$");throw Gn("imatcher",'Matchers may only be "self", string patterns or RegExp objects')}(t))}),e}function ge(){this.SCE_CONTEXTS=Zn;var a=["self"],s=[];this.resourceUrlWhitelist=function(t){return arguments.length&&(a=fe(t)),a},this.resourceUrlBlacklist=function(t){return arguments.length&&(s=fe(t)),s},this.$get=["$injector",function(t){function o(t,e){return"self"===t?xe(e):!!t.exec(e.href)}function e(t){function e(t){this.$$unwrapTrustedValue=function(){return t}}return t&&(e.prototype=new t),e.prototype.valueOf=function(){return this.$$unwrapTrustedValue()},e.prototype.toString=function(){return this.$$unwrapTrustedValue().toString()},e}var i=function(t){throw Gn("unsafe","Attempting to use an unsafe value in a safe context.")};t.has("$sanitize")&&(i=t.get("$sanitize"));var n=e(),r={};return r[Zn.HTML]=e(n),r[Zn.CSS]=e(n),r[Zn.URL]=e(n),r[Zn.JS]=e(n),r[Zn.RESOURCE_URL]=e(r[Zn.URL]),{trustAs:function(t,e){var n=r.hasOwnProperty(t)?r[t]:null;if(!n)throw Gn("icontext","Attempted to trust a value in invalid context. Context: {0}; Value: {1}",t,e);if(null===e||e===tt||""===e)return e;if("string"!=typeof e)throw Gn("itype","Attempted to trust a non-string value in a content requiring a string: Context: {0}",t);return new n(e)},getTrusted:function(t,e){if(null===e||e===tt||""===e)return e;var n=r.hasOwnProperty(t)?r[t]:null;if(n&&e instanceof n)return e.$$unwrapTrustedValue();if(t===Zn.RESOURCE_URL){if(function(t){var e,n,i=be(t.toString()),r=!1;for(e=0,n=a.length;e<n;e++)if(o(a[e],i)){r=!0;break}if(r)for(e=0,n=s.length;e<n;e++)if(o(s[e],i)){r=!1;break}return r}(e))return e;throw Gn("insecurl","Blocked loading resource from url not allowed by $sceDelegate policy.  URL: {0}",e.toString())}if(t===Zn.HTML)return i(e);throw Gn("unsafe","Attempting to use an unsafe value in a safe context.")},valueOf:function(t){return t instanceof n?t.$$unwrapTrustedValue():t}}}]}function ve(){var s=!0;this.enabled=function(t){return arguments.length&&(s=!!t),s},this.$get=["$parse","$sniffer","$sceDelegate",function(e,t,n){if(s&&t.msie&&t.msieDocumentMode<8)throw Gn("iequirks","Strict Contextual Escaping does not support Internet Explorer version < 9 in quirks mode.  You can fix this by adding the text <!doctype html> to the top of your HTML document.  See http://docs.angularjs.org/api/ng.$sce for more information.");var r=ot(Zn);r.isEnabled=function(){return s},r.trustAs=n.trustAs,r.getTrusted=n.getTrusted,r.valueOf=n.valueOf,s||(r.trustAs=r.getTrusted=function(t,e){return e},r.valueOf=m),r.parseAs=function(n,t){var i=e(t);return i.literal&&i.constant?i:function(t,e){return r.getTrusted(n,i(t,e))}};var i=r.parseAs,o=r.getTrusted,a=r.trustAs;return et(Zn,function(e,t){var n=Ve(t);r[q("parse_as_"+n)]=function(t){return i(e,t)},r[q("get_trusted_"+n)]=function(t){return o(e,t)},r[q("trust_as_"+n)]=function(t){return a(e,t)}}),r}]}function me(){this.$get=["$window","$document",function(t,e){var n,i,r={},o=$((/android (\d+)/.exec(Ve((t.navigator||{}).userAgent))||[])[1]),a=/Boxee/i.test((t.navigator||{}).userAgent),s=e[0]||{},l=s.documentMode,u=/^(Moz|webkit|O|ms)(?=[A-Z])/,c=s.body&&s.body.style,h=!1,d=!1;if(c){for(var p in c)if(i=u.exec(p)){n=(n=i[0]).substr(0,1).toUpperCase()+n.substr(1);break}n||(n="WebkitOpacity"in c&&"webkit"),h=!!("transition"in c||n+"Transition"in c),d=!!("animation"in c||n+"Animation"in c),!o||h&&d||(h=it(s.body.style.webkitTransition),d=it(s.body.style.webkitAnimation))}return{history:!(!t.history||!t.history.pushState||o<4||a),hashchange:"onhashchange"in t&&(!l||7<l),hasEvent:function(t){if("input"==t&&9==He)return!1;if(S(r[t])){var e=s.createElement("div");r[t]="on"+t in e}return r[t]},csp:rn(),vendorPrefix:n,transitions:h,animations:d,android:o,msie:He,msieDocumentMode:l}}]}function ye(){this.$get=["$rootScope","$browser","$q","$exceptionHandler",function(s,l,u,c){function t(t,e,n){var i,r=u.defer(),o=r.promise,a=F(n)&&!n;return i=l.defer(function(){try{r.resolve(t())}catch(t){r.reject(t),c(t)}finally{delete h[o.$$timeoutId]}a||s.$apply()},e),o.$$timeoutId=i,h[i]=r,o}var h={};return t.cancel=function(t){return!!(t&&t.$$timeoutId in h)&&(h[t.$$timeoutId].reject("canceled"),delete h[t.$$timeoutId],l.defer.cancel(t.$$timeoutId))},t}]}function be(t){var e=t;return He&&(Kn.setAttribute("href",e),e=Kn.href),Kn.setAttribute("href",e),{href:Kn.href,protocol:Kn.protocol?Kn.protocol.replace(/:$/,""):"",host:Kn.host,search:Kn.search?Kn.search.replace(/^\?/,""):"",hash:Kn.hash?Kn.hash.replace(/^#/,""):"",hostname:Kn.hostname,port:Kn.port,pathname:"/"===Kn.pathname.charAt(0)?Kn.pathname:"/"+Kn.pathname}}function xe(t){var e=it(t)?be(t):t;return e.protocol===Qn.protocol&&e.host===Qn.host}function we(){this.$get=y(f)}function $e(i){function r(t,e){if(nt(t)){var n={};return et(t,function(t,e){n[e]=r(e,t)}),n}return i.factory(t+o,e)}var o="Filter";this.register=r,this.$get=["$injector",function(e){return function(t){return e.get(t+o)}}],r("currency",Ce),r("date",Pe),r("filter",ke),r("json",Me),r("limitTo",Ae),r("lowercase",ii),r("number",Ee),r("orderBy",Ne),r("uppercase",ri)}function ke(){return function(t,n,r){if(!en(t))return t;var e=typeof r,i=[];i.check=function(t){for(var e=0;e<i.length;e++)if(!i[e](t))return!1;return!0},"function"!=e&&(r="boolean"==e&&r?function(t,e){return Je.equals(t,e)}:function(t,e){if(t&&e&&"object"==typeof t&&"object"==typeof e){for(var n in t)if("$"!==n.charAt(0)&&qe.call(t,n)&&r(t[n],e[n]))return!0;return!1}return e=(""+e).toLowerCase(),-1<(""+t).toLowerCase().indexOf(e)});var o=function(t,e){if("string"==typeof e&&"!"===e.charAt(0))return!o(t,e.substr(1));switch(typeof t){case"boolean":case"number":case"string":return r(t,e);case"object":switch(typeof e){case"object":return r(t,e);default:for(var n in t)if("$"!==n.charAt(0)&&o(t[n],e))return!0}return!1;case"array":for(var i=0;i<t.length;i++)if(o(t[i],e))return!0;return!1;default:return!1}};switch(typeof n){case"boolean":case"number":case"string":n={$:n};case"object":for(var a in n)!function(e){void 0!==n[e]&&i.push(function(t){return o("$"==e?t:t&&t[e],n[e])})}(a);break;case"function":i.push(n);break;default:return t}for(var s=[],l=0;l<t.length;l++){var u=t[l];i.check(u)&&s.push(u)}return s}}function Ce(t){var n=t.NUMBER_FORMATS;return function(t,e){return S(e)&&(e=n.CURRENCY_SYM),Se(t,n.PATTERNS[1],n.GROUP_SEP,n.DECIMAL_SEP,2).replace(/\u00A4/g,e)}}function Ee(t){var n=t.NUMBER_FORMATS;return function(t,e){return Se(t,n.PATTERNS[0],n.GROUP_SEP,n.DECIMAL_SEP,e)}}function Se(t,e,n,i,r){if(null==t||!isFinite(t)||nt(t))return"";var o=t<0,a=(t=Math.abs(t))+"",s="",l=[],u=!1;if(-1!==a.indexOf("e")){var c=a.match(/([\d\.]+)e(-?)(\d+)/);c&&"-"==c[2]&&c[3]>r+1?(a="0",t=0):(s=a,u=!0)}if(u)0<r&&-1<t&&t<1&&(s=t.toFixed(r));else{var h=(a.split(Jn)[1]||"").length;S(r)&&(r=Math.min(Math.max(e.minFrac,h),e.maxFrac)),0===(t=+(Math.round(+(t.toString()+"e"+r)).toString()+"e"+-r))&&(o=!1);var d=(""+t).split(Jn),p=d[0];d=d[1]||"";var f,g=0,v=e.lgSize,m=e.gSize;if(p.length>=v+m)for(g=p.length-v,f=0;f<g;f++)(g-f)%m==0&&0!==f&&(s+=n),s+=p.charAt(f);for(f=g;f<p.length;f++)(p.length-f)%v==0&&0!==f&&(s+=n),s+=p.charAt(f);for(;d.length<r;)d+="0";r&&"0"!==r&&(s+=i+d.substr(0,r))}return l.push(o?e.negPre:e.posPre),l.push(s),l.push(o?e.negSuf:e.posSuf),l.join("")}function Te(t,e,n){var i="";for(t<0&&(i="-",t=-t),t=""+t;t.length<e;)t="0"+t;return n&&(t=t.substr(t.length-e)),i+t}function De(n,i,r,o){return r=r||0,function(t){var e=t["get"+n]();return(0<r||-r<e)&&(e+=r),0===e&&-12==r&&(e=12),Te(e,i,o)}}function _e(i,r){return function(t,e){var n=t["get"+i]();return e[ze(r?"SHORT"+i:i)][n]}}function Pe(a){var h=/^(\d{4})-?(\d\d)-?(\d\d)(?:T(\d\d)(?::?(\d\d)(?::?(\d\d)(?:\.(\d+))?)?)?(Z|([+-])(\d\d):?(\d\d))?)?$/;return function(e,t){var n,i,r="",o=[];if(t=t||"mediumDate",t=a.DATETIME_FORMATS[t]||t,it(e)&&(e=ni.test(e)?$(e):function(t){var e;if(e=t.match(h)){var n=new Date(0),i=0,r=0,o=e[8]?n.setUTCFullYear:n.setFullYear,a=e[8]?n.setUTCHours:n.setHours;e[9]&&(i=$(e[9]+e[10]),r=$(e[9]+e[11])),o.call(n,$(e[1]),$(e[2])-1,$(e[3]));var s=$(e[4]||0)-i,l=$(e[5]||0)-r,u=$(e[6]||0),c=Math.round(1e3*parseFloat("0."+(e[7]||0)));return a.call(n,s,l,u,c),n}return t}(e)),l(e)&&(e=new Date(e)),!b(e))return e;for(;t;)t=(i=ei.exec(t))?(o=D(o,i,1)).pop():(o.push(t),null);return et(o,function(t){n=ti[t],r+=n?n(e,a.DATETIME_FORMATS):t.replace(/(^'|'$)/g,"").replace(/''/g,"'")}),r}}function Me(){return function(t){return A(t,!0)}}function Ae(){return function(t,e){if(!en(t)&&!it(t))return t;if(e=Math.abs(Number(e))===1/0?Number(e):$(e),it(t))return e?0<=e?t.slice(0,e):t.slice(e,t.length):"";var n,i,r=[];for(e>t.length?e=t.length:e<-t.length&&(e=-t.length),i=0<e?(n=0,e):(n=t.length+e,t.length);n<i;n++)r.push(t[n]);return r}}function Ne(s){return function(t,r,e){function o(n,t){return N(t)?function(t,e){return n(e,t)}:n}function a(t,e){var n=typeof t,i=typeof e;return n==i?(b(t)&&b(e)&&(t=t.valueOf(),e=e.valueOf()),"string"==n&&(t=t.toLowerCase(),e=e.toLowerCase()),t===e?0:t<e?-1:1):n<i?-1:1}return P(t)?(0===(r=en(r)?r:[r]).length&&(r=["+"]),r=function(t,i,r){var o=[];return et(t,function(t,e,n){o.push(i.call(r,t,e,n))}),o}(r,function(t){var e=!1,n=t||m;if(it(t)){if("+"!=t.charAt(0)&&"-"!=t.charAt(0)||(e="-"==t.charAt(0),t=t.substring(1)),""===t)return o(function(t,e){return a(t,e)},e);if((n=s(t)).constant){var i=n();return o(function(t,e){return a(t[i],e[i])},e)}}return o(function(t,e){return a(n(t),n(e))},e)}),Ge.call(t).sort(o(function(t,e){for(var n=0;n<r.length;n++){var i=r[n](t,e);if(0!==i)return i}return 0},e))):t}}function Oe(t){return rt(t)&&(t={link:t}),t.restrict=t.restrict||"AC",y(t)}function Ie(n,t,e,i){function r(t,e){e=e?"-"+ut(e,"-"):"",i.setClass(n,(t?wi:$i)+e,(t?$i:wi)+e)}var o=this,a=n.parent().controller("form")||si,s=0,l=o.$error={},u=[];o.$name=t.name||t.ngForm,o.$dirty=!1,o.$pristine=!0,o.$valid=!0,o.$invalid=!1,a.$addControl(o),n.addClass(ki),r(!0),o.$addControl=function(t){L(t.$name,"input"),u.push(t),t.$name&&(o[t.$name]=t)},o.$removeControl=function(n){n.$name&&o[n.$name]===n&&delete o[n.$name],et(l,function(t,e){o.$setValidity(e,!0,n)}),p(u,n)},o.$setValidity=function(t,e,n){var i=l[t];if(e)i&&(p(i,n),i.length||(--s||(r(e),o.$valid=!0,o.$invalid=!1),r(!(l[t]=!1),t),a.$setValidity(t,!0,o)));else{if(s||r(e),i){if(function(t,e){return-1!=C(t,e)}(i,n))return}else l[t]=i=[],s++,r(!1,t),a.$setValidity(t,!1,o);i.push(n),o.$valid=!1,o.$invalid=!0}},o.$setDirty=function(){i.removeClass(n,ki),i.addClass(n,Ci),o.$dirty=!0,o.$pristine=!1,a.$setDirty()},o.$setPristine=function(){i.removeClass(n,Ci),i.addClass(n,ki),o.$dirty=!1,o.$pristine=!0,et(u,function(t){t.$setPristine()})}}function Be(t,e,n,i){return t.$setValidity(e,n),n?i:tt}function Le(t,e){var n;if(e)for(n=0;n<e.length;++n)if(t[e[n]])return!0;return!1}function je(i,r,o,a,t,e){var s=r.prop(Fe),l=r[0].placeholder,u={},c=Ve(r[0].type);if(a.$$validityState=s,!t.android){var h=!1;r.on("compositionstart",function(t){h=!0}),r.on("compositionend",function(){h=!1,n()})}var n=function(t){if(!h){var e=r.val();if(He&&"input"===(t||u).type&&r[0].placeholder!==l)return void(l=r[0].placeholder);"password"!==c&&N(o.ngTrim||"T")&&(e=nn(e));var n=s&&a.$$hasNativeValidators;(a.$viewValue!==e||""===e&&n)&&(i.$root.$$phase?a.$setViewValue(e):i.$apply(function(){a.$setViewValue(e)}))}};if(t.hasEvent("input"))r.on("input",n);else{var d,p=function(){d||(d=e.defer(function(){n(),d=null}))};r.on("keydown",function(t){var e=t.keyCode;91===e||15<e&&e<19||37<=e&&e<=40||p()}),t.hasEvent("paste")&&r.on("paste cut",p)}r.on("change",n),a.$render=function(){r.val(a.$isEmpty(a.$viewValue)?"":a.$viewValue)};var f,g,v=o.ngPattern;if(v){var m=function(t,e){return Be(a,"pattern",a.$isEmpty(e)||t.test(e),e)};f=(g=v.match(/^\/(.*)\/([gim]*)$/))?(v=new RegExp(g[1],g[2]),function(t){return m(v,t)}):function(t){var e=i.$eval(v);if(!e||!e.test)throw k("ngPattern")("noregexp","Expected {0} to be a RegExp but was {1}. Element: {2}",v,e,lt(r));return m(e,t)},a.$formatters.push(f),a.$parsers.push(f)}if(o.ngMinlength){var y=$(o.ngMinlength),b=function(t){return Be(a,"minlength",a.$isEmpty(t)||t.length>=y,t)};a.$parsers.push(b),a.$formatters.push(b)}if(o.ngMaxlength){var x=$(o.ngMaxlength),w=function(t){return Be(a,"maxlength",a.$isEmpty(t)||t.length<=x,t)};a.$parsers.push(w),a.$formatters.push(w)}}function Re(d,p){return d="ngClass"+d,["$animate",function(u){function c(t,e){var n=[];t:for(var i=0;i<t.length;i++){for(var r=t[i],o=0;o<e.length;o++)if(r==e[o])continue t;n.push(r)}return n}function h(t){if(en(t))return t;if(it(t))return t.split(" ");if(nt(t)){var n=[];return et(t,function(t,e){t&&(n=n.concat(e.split(" ")))}),n}return t}return{restrict:"AC",link:function(r,o,a){function s(t){var e=l(t,1);a.$addClass(e)}function l(t,e){var n=o.data("$classCounts")||{},i=[];return et(t,function(t){(0<e||n[t])&&(n[t]=(n[t]||0)+e,n[t]===+(0<e)&&i.push(t))}),o.data("$classCounts",n),i.join(" ")}function e(t){if(!0===p||r.$index%2===p){var e=h(t||[]);if(n){if(!at(t,n))!function(t,e){var n=c(e,t),i=c(t,e);i=l(i,-1),0===(n=l(n,1)).length?u.removeClass(o,i):0===i.length?u.addClass(o,n):u.setClass(o,n,i)}(h(n),e)}else s(e)}n=ot(t)}var n;r.$watch(a[d],e,!0),a.$observe("class",function(t){e(r.$eval(a[d]))}),"ngClass"!==d&&r.$watch("$index",function(t,e){var n=1&t;if(n!=(1&e)){var i=h(r.$eval(a[d]));n===p?s(i):function(t){var e=l(t,-1);a.$removeClass(e)}(i)}})}}}]}var Fe="validity",Ve=function(t){return it(t)?t.toLowerCase():t},qe=Object.prototype.hasOwnProperty,ze=function(t){return it(t)?t.toUpperCase():t};"i"!=="I".toLowerCase()&&(Ve=function(t){return it(t)?t.replace(/[A-Z]/g,function(t){return String.fromCharCode(32|t.charCodeAt(0))}):t},ze=function(t){return it(t)?t.replace(/[a-z]/g,function(t){return String.fromCharCode(-33&t.charCodeAt(0))}):t});var He,Xe,Ye,Ue,We,Ge=[].slice,Ze=[].push,Ke=Object.prototype.toString,Qe=k("ng"),Je=f.angular||(f.angular={}),tn=["0","0","0"];He=$((/msie (\d+)/.exec(Ve(navigator.userAgent))||[])[1]),isNaN(He)&&(He=$((/trident\/.*; rv:(\d+)/.exec(Ve(navigator.userAgent))||[])[1])),E.$inject=[],m.$inject=[];var en=rt(Array.isArray)?Array.isArray:function(t){return"[object Array]"===Ke.call(t)},nn=String.prototype.trim?function(t){return it(t)?t.trim():t}:function(t){return it(t)?t.replace(/^\s\s*/,"").replace(/\s\s*$/,""):t};We=He<9?function(t){return(t=t.nodeName?t:t[0]).scopeName&&"HTML"!=t.scopeName?ze(t.scopeName+":"+t.nodeName):t.nodeName}:function(t){return t.nodeName?t.nodeName:t[0].nodeName};var rn=function(){if(F(rn.isActive_))return rn.isActive_;var e=!(!J.querySelector("[ng-csp]")&&!J.querySelector("[data-ng-csp]"));if(!e)try{new Function("")}catch(t){e=!0}return rn.isActive_=e},on=/[A-Z]/g,an={full:"1.2.28",major:1,minor:2,dot:28,codeName:"finnish-disembarkation"};H.expando="ng339";var sn=H.cache={},ln=1,un=f.document.addEventListener?function(t,e,n){t.addEventListener(e,n,!1)}:function(t,e,n){t.attachEvent("on"+e,n)},cn=f.document.removeEventListener?function(t,e,n){t.removeEventListener(e,n,!1)}:function(t,e,n){t.detachEvent("on"+e,n)},hn=(H._data=function(t){return this.cache[t[this.expando]]||{}},/([\:\-\_]+(.))/g),dn=/^moz([A-Z])/,pn=k("jqLite"),fn=/^<(\w+)\s*\/?>(?:<\/\1>|)$/,gn=/<|&#?\w+;/,vn=/<([\w:]+)/,mn=/<(?!area|br|col|embed|hr|img|input|link|meta|param)(([\w:]+)[^>]*)\/>/gi,yn={option:[1,'<select multiple="multiple">',"</select>"],thead:[1,"<table>","</table>"],col:[2,"<table><colgroup>","</colgroup></table>"],tr:[2,"<table><tbody>","</tbody></table>"],td:[3,"<table><tbody><tr>","</tr></tbody></table>"],_default:[0,"",""]};yn.optgroup=yn.option,yn.tbody=yn.tfoot=yn.colgroup=yn.caption=yn.thead,yn.th=yn.td;var bn=H.prototype={ready:function(t){function e(){n||(n=!0,t())}var n=!1;"complete"===J.readyState?setTimeout(e):(this.on("DOMContentLoaded",e),H(f).on("load",e))},toString:function(){var e=[];return et(this,function(t){e.push(""+t)}),"["+e.join(", ")+"]"},eq:function(t){return Xe(0<=t?this[t]:this[this.length+t])},length:0,push:Ze,sort:[].sort,splice:[].splice},xn={};et("multiple,selected,checked,disabled,readOnly,required,open".split(","),function(t){xn[Ve(t)]=t});var wn,$n={};function kn(t,e){var n=wn[t.nodeType];return S(e)?n?t[n]:"":void(t[n]=e)}et("input,select,option,textarea,button,form,details".split(","),function(t){$n[ze(t)]=!0}),et({data:G,removeData:U},function(t,e){H[e]=t}),et({data:G,inheritedData:gt,scope:function(t){return Xe.data(t,"$scope")||gt(t.parentNode||t,["$isolateScope","$scope"])},isolateScope:function(t){return Xe.data(t,"$isolateScope")||Xe.data(t,"$isolateScopeNoTemplate")},controller:ft,injector:function(t){return gt(t,"$injector")},removeAttr:function(t,e){t.removeAttribute(e)},hasClass:Z,css:function(t,e,n){var i;if(e=q(e),!F(n))return He<=8&&(""===(i=t.currentStyle&&t.currentStyle[e])&&(i="auto")),i=i||t.style[e],He<=8&&(i=""===i?tt:i),i;t.style[e]=n},attr:function(t,e,n){var i=Ve(e);if(xn[i]){if(!F(n))return t[e]||(t.attributes.getNamedItem(e)||E).specified?i:tt;n?(t[e]=!0,t.setAttribute(e,i)):(t[e]=!1,t.removeAttribute(i))}else if(F(n))t.setAttribute(e,n);else if(t.getAttribute){var r=t.getAttribute(e,2);return null===r?tt:r}},prop:function(t,e,n){return F(n)?void(t[e]=n):t[e]},text:(wn=[],He<9?(wn[1]="innerText",wn[3]="nodeValue"):wn[1]=wn[3]="textContent",kn.$dv="",kn),val:function(t,e){if(S(e)){if("SELECT"===We(t)&&t.multiple){var n=[];return et(t.options,function(t){t.selected&&n.push(t.value||t.text)}),0===n.length?null:n}return t.value}t.value=e},html:function(t,e){if(S(e))return t.innerHTML;for(var n=0,i=t.childNodes;n<i.length;n++)X(i[n]);t.innerHTML=e},empty:vt},function(u,t){H.prototype[t]=function(t,e){var n,i,r=this.length;if(u!==vt&&(2==u.length&&u!==Z&&u!==ft?t:e)===tt){if(nt(t)){for(n=0;n<r;n++)if(u===G)u(this[n],t);else for(i in t)u(this[n],i,t[i]);return this}for(var o=u.$dv,a=o===tt?Math.min(r,1):r,s=0;s<a;s++){var l=u(this[s],t,e);o=o?o+l:l}return o}for(n=0;n<r;n++)u(this[n],t,e);return this}}),et({removeData:U,dealoc:X,on:function e(r,t,o,n){if(F(n))throw pn("onargs","jqLite#on() does not support the `selector` or `eventData` parameters");var a=W(r,"events"),s=W(r,"handle");a||W(r,"events",a={}),s||W(r,"handle",s=function(i,r){function t(e,t){if(e.preventDefault||(e.preventDefault=function(){e.returnValue=!1}),e.stopPropagation||(e.stopPropagation=function(){e.cancelBubble=!0}),e.target||(e.target=e.srcElement||J),S(e.defaultPrevented)){var n=e.preventDefault;e.preventDefault=function(){e.defaultPrevented=!0,n.call(e)},e.defaultPrevented=!1}e.isDefaultPrevented=function(){return e.defaultPrevented||!1===e.returnValue},et(ot(r[t||e.type]||[]),function(t){t.call(i,e)}),He<=8?(e.preventDefault=null,e.stopPropagation=null,e.isDefaultPrevented=null):(delete e.preventDefault,delete e.stopPropagation,delete e.isDefaultPrevented)}return t.elem=i,t}(r,a)),et(t.split(" "),function(n){var t=a[n];if(!t){if("mouseenter"==n||"mouseleave"==n){var i=J.body.contains||J.body.compareDocumentPosition?function(t,e){var n=9===t.nodeType?t.documentElement:t,i=e&&e.parentNode;return t===i||!(!i||1!==i.nodeType||!(n.contains?n.contains(i):t.compareDocumentPosition&&16&t.compareDocumentPosition(i)))}:function(t,e){if(e)for(;e=e.parentNode;)if(e===t)return!0;return!1};a[n]=[];e(r,{mouseleave:"mouseout",mouseenter:"mouseover"}[n],function(t){var e=t.relatedTarget;e&&(e===this||i(this,e))||s(t,n)})}else un(r,n,s),a[n]=[];t=a[n]}t.push(o)})},off:Y,one:function(e,n,i){(e=Xe(e)).on(n,function t(){e.off(n,i),e.off(n,t)}),e.on(n,i)},replaceWith:function(e,t){var n,i=e.parentNode;X(e),et(new H(t),function(t){n?i.insertBefore(t,n.nextSibling):i.replaceChild(t,e),n=t})},children:function(t){var e=[];return et(t.childNodes,function(t){1===t.nodeType&&e.push(t)}),e},contents:function(t){return t.contentDocument||t.childNodes||[]},append:function(e,t){et(new H(t),function(t){1!==e.nodeType&&11!==e.nodeType||e.appendChild(t)})},prepend:function(e,t){if(1===e.nodeType){var n=e.firstChild;et(new H(t),function(t){e.insertBefore(t,n)})}},wrap:function(t,e){e=Xe(e)[0];var n=t.parentNode;n&&n.replaceChild(e,t),e.appendChild(t)},remove:function(t){X(t);var e=t.parentNode;e&&e.removeChild(t)},after:function(t,e){var n=t,i=t.parentNode;et(new H(e),function(t){i.insertBefore(t,n.nextSibling),n=t})},addClass:Q,removeClass:K,toggleClass:function(n,t,i){t&&et(t.split(" "),function(t){var e=i;S(e)&&(e=!Z(n,t)),(e?Q:K)(n,t)})},parent:function(t){var e=t.parentNode;return e&&11!==e.nodeType?e:null},next:function(t){if(t.nextElementSibling)return t.nextElementSibling;for(var e=t.nextSibling;null!=e&&1!==e.nodeType;)e=e.nextSibling;return e},find:function(t,e){return t.getElementsByTagName?t.getElementsByTagName(e):[]},clone:dt,triggerHandler:function(e,t,n){var i,r,o,a=t.type||t,s=(W(e,"events")||{})[a];s&&(i={preventDefault:function(){this.defaultPrevented=!0},isDefaultPrevented:function(){return!0===this.defaultPrevented},stopPropagation:E,type:a,target:e},t.type&&(i=M(i,t)),r=ot(s),o=n?[i].concat(n):[i],et(r,function(t){t.apply(e,o)}))}},function(o,t){H.prototype[t]=function(t,e,n){for(var i,r=0;r<this.length;r++)S(i)?F(i=o(this[r],t,e,n))&&(i=Xe(i)):pt(i,o(this[r],t,e,n));return F(i)?i:this},H.prototype.bind=H.prototype.on,H.prototype.unbind=H.prototype.off}),bt.prototype={put:function(t,e){this[yt(t,this.nextUid)]=e},get:function(t){return this[yt(t,this.nextUid)]},remove:function(t){var e=this[t=yt(t,this.nextUid)];return delete this[t],e}};var Cn=/^function\s*[^\(]*\(\s*([^\)]*)\)/m,En=/,/,Sn=/^\s*(_?)(\S+?)\1\s*$/,Tn=/((\/\/.*$)|(\/\*[\s\S]*?\*\/))/gm,Dn=k("$injector"),_n=k("$animate"),Pn=["$provide",function(i){this.$$selectors={},this.register=function(t,e){var n=t+"-animation";if(t&&"."!=t.charAt(0))throw _n("notcsel","Expecting class selector starting with '.' got '{0}'.",t);this.$$selectors[t.substr(1)]=n,i.factory(n,e)},this.classNameFilter=function(t){return 1===arguments.length&&(this.$$classNameFilter=t instanceof RegExp?t:null),this.$$classNameFilter},this.$get=["$timeout","$$asyncCallback",function(t,e){function r(t){t&&e(t)}return{enter:function(t,e,n,i){n?n.after(t):(e&&e[0]||(e=n.parent()),e.append(t)),r(i)},leave:function(t,e){t.remove(),r(e)},move:function(t,e,n,i){this.enter(t,e,n,i)},addClass:function(t,e,n){e=it(e)?e:en(e)?e.join(" "):"",et(t,function(t){Q(t,e)}),r(n)},removeClass:function(t,e,n){e=it(e)?e:en(e)?e.join(" "):"",et(t,function(t){K(t,e)}),r(n)},setClass:function(t,e,n,i){et(t,function(t){Q(t,e),K(t,n)}),r(i)},enabled:E}}]}],Mn=k("$compile");Dt.$inject=["$provide","$$sanitizeUriProvider"];var An=/^(x[\:\-_]|data[\:\-_])/i,Nn=k("$interpolate"),On=/^([^\?#]*)(\?([^#]*))?(#(.*))?$/,In={http:80,https:443,ftp:21},Bn=k("$location");Qt.prototype=Kt.prototype=Zt.prototype={$$html5:!1,$$replace:!1,absUrl:Jt("$$absUrl"),url:function(t){if(S(t))return this.$$url;var e=On.exec(t);return e[1]&&this.path(decodeURIComponent(e[1])),(e[2]||e[1])&&this.search(e[3]||""),this.hash(e[5]||""),this},protocol:Jt("$$protocol"),host:Jt("$$host"),port:Jt("$$port"),path:te("$$path",function(t){return"/"==(t=null!==t?t.toString():"").charAt(0)?t:"/"+t}),search:function(n,t){switch(arguments.length){case 0:return this.$$search;case 1:if(it(n)||l(n))n=n.toString(),this.$$search=s(n);else{if(!nt(n))throw Bn("isrcharg","The first argument of the `$location#search()` call must be a string or an object.");et(n,function(t,e){null==t&&delete n[e]}),this.$$search=n}break;default:S(t)||null===t?delete this.$$search[n]:this.$$search[n]=t}return this.$$compose(),this},hash:te("$$hash",function(t){return null!==t?t.toString():""}),replace:function(){return this.$$replace=!0,this}};var Ln,jn=k("$parse"),Rn={},Fn=Function.prototype.call,Vn=Function.prototype.apply,qn=Function.prototype.bind,zn={null:function(){return null},true:function(){return!0},false:function(){return!1},undefined:E,"+":function(t,e,n,i){return n=n(t,e),i=i(t,e),F(n)?F(i)?n+i:n:F(i)?i:tt},"-":function(t,e,n,i){return n=n(t,e),i=i(t,e),(F(n)?n:0)-(F(i)?i:0)},"*":function(t,e,n,i){return n(t,e)*i(t,e)},"/":function(t,e,n,i){return n(t,e)/i(t,e)},"%":function(t,e,n,i){return n(t,e)%i(t,e)},"^":function(t,e,n,i){return n(t,e)^i(t,e)},"=":E,"===":function(t,e,n,i){return n(t,e)===i(t,e)},"!==":function(t,e,n,i){return n(t,e)!==i(t,e)},"==":function(t,e,n,i){return n(t,e)==i(t,e)},"!=":function(t,e,n,i){return n(t,e)!=i(t,e)},"<":function(t,e,n,i){return n(t,e)<i(t,e)},">":function(t,e,n,i){return n(t,e)>i(t,e)},"<=":function(t,e,n,i){return n(t,e)<=i(t,e)},">=":function(t,e,n,i){return n(t,e)>=i(t,e)},"&&":function(t,e,n,i){return n(t,e)&&i(t,e)},"||":function(t,e,n,i){return n(t,e)||i(t,e)},"&":function(t,e,n,i){return n(t,e)&i(t,e)},"|":function(t,e,n,i){return i(t,e)(t,e,n(t,e))},"!":function(t,e,n){return!n(t,e)}},Hn={n:"\n",f:"\f",r:"\r",t:"\t",v:"\v","'":"'",'"':'"'},Xn=function(t){this.options=t};Xn.prototype={constructor:Xn,lex:function(t){for(this.text=t,this.index=0,this.ch=tt,this.lastCh=":",this.tokens=[];this.index<this.text.length;){if(this.ch=this.text.charAt(this.index),this.is("\"'"))this.readString(this.ch);else if(this.isNumber(this.ch)||this.is(".")&&this.isNumber(this.peek()))this.readNumber();else if(this.isIdent(this.ch))this.readIdent();else if(this.is("(){}[].,;:?"))this.tokens.push({index:this.index,text:this.ch}),this.index++;else{if(this.isWhitespace(this.ch)){this.index++;continue}var e=this.ch+this.peek(),n=e+this.peek(2),i=zn[this.ch],r=zn[e],o=zn[n];o?(this.tokens.push({index:this.index,text:n,fn:o}),this.index+=3):r?(this.tokens.push({index:this.index,text:e,fn:r}),this.index+=2):i?(this.tokens.push({index:this.index,text:this.ch,fn:i}),this.index+=1):this.throwError("Unexpected next character ",this.index,this.index+1)}this.lastCh=this.ch}return this.tokens},is:function(t){return-1!==t.indexOf(this.ch)},was:function(t){return-1!==t.indexOf(this.lastCh)},peek:function(t){var e=t||1;return this.index+e<this.text.length&&this.text.charAt(this.index+e)},isNumber:function(t){return"0"<=t&&t<="9"},isWhitespace:function(t){return" "===t||"\r"===t||"\t"===t||"\n"===t||"\v"===t||" "===t},isIdent:function(t){return"a"<=t&&t<="z"||"A"<=t&&t<="Z"||"_"===t||"$"===t},isExpOperator:function(t){return"-"===t||"+"===t||this.isNumber(t)},throwError:function(t,e,n){n=n||this.index;var i=F(e)?"s "+e+"-"+this.index+" ["+this.text.substring(e,n)+"]":" "+n;throw jn("lexerr","Lexer Error: {0} at column{1} in expression [{2}].",t,i,this.text)},readNumber:function(){for(var t="",e=this.index;this.index<this.text.length;){var n=Ve(this.text.charAt(this.index));if("."==n||this.isNumber(n))t+=n;else{var i=this.peek();if("e"==n&&this.isExpOperator(i))t+=n;else if(this.isExpOperator(n)&&i&&this.isNumber(i)&&"e"==t.charAt(t.length-1))t+=n;else{if(!this.isExpOperator(n)||i&&this.isNumber(i)||"e"!=t.charAt(t.length-1))break;this.throwError("Invalid exponent")}}this.index++}t*=1,this.tokens.push({index:e,text:t,literal:!0,constant:!0,fn:function(){return t}})},readIdent:function(){for(var t,e,n,i,r=this,o="",a=this.index;this.index<this.text.length&&("."===(i=this.text.charAt(this.index))||this.isIdent(i)||this.isNumber(i));)"."===i&&(t=this.index),o+=i,this.index++;if(t)for(e=this.index;e<this.text.length;){if("("===(i=this.text.charAt(e))){n=o.substr(t-a+1),o=o.substr(0,t-a),this.index=e;break}if(!this.isWhitespace(i))break;e++}var s={index:a,text:o};if(zn.hasOwnProperty(o))s.fn=zn[o],s.literal=!0,s.constant=!0;else{var l=le(o,this.options,this.text);s.fn=M(function(t,e){return l(t,e)},{assign:function(t,e){return oe(t,o,e,r.text,r.options)}})}this.tokens.push(s),n&&(this.tokens.push({index:t,text:"."}),this.tokens.push({index:t+1,text:n}))},readString:function(t){var e=this.index;this.index++;for(var n="",i=t,r=!1;this.index<this.text.length;){var o=this.text.charAt(this.index);if(i+=o,r){if("u"===o){var a=this.text.substring(this.index+1,this.index+5);a.match(/[\da-f]{4}/i)||this.throwError("Invalid unicode escape [\\u"+a+"]"),this.index+=4,n+=String.fromCharCode(parseInt(a,16))}else{n+=Hn[o]||o}r=!1}else if("\\"===o)r=!0;else{if(o===t)return this.index++,void this.tokens.push({index:e,text:i,string:n,literal:!0,constant:!0,fn:function(){return n}});n+=o}this.index++}this.throwError("Unterminated quote",e)}};var Yn=function(t,e,n){this.lexer=t,this.$filter=e,this.options=n};Yn.ZERO=M(function(){return 0},{constant:!0}),Yn.prototype={constructor:Yn,parse:function(t){this.text=t,this.tokens=this.lexer.lex(t);var e=this.statements();return 0!==this.tokens.length&&this.throwError("is an unexpected token",this.tokens[0]),e.literal=!!e.literal,e.constant=!!e.constant,e},primary:function(){var t,e,n;if(this.expect("("))t=this.filterChain(),this.consume(")");else if(this.expect("["))t=this.arrayDeclaration();else if(this.expect("{"))t=this.object();else{var i=this.expect();(t=i.fn)||this.throwError("not a primary expression",i),t.literal=!!i.literal,t.constant=!!i.constant}for(;e=this.expect("(","[",".");)"("===e.text?(t=this.functionCall(t,n),n=null):"["===e.text?(n=t,t=this.objectIndex(t)):"."===e.text?(n=t,t=this.fieldAccess(t)):this.throwError("IMPOSSIBLE");return t},throwError:function(t,e){throw jn("syntax","Syntax Error: Token '{0}' {1} at column {2} of the expression [{3}] starting at [{4}].",e.text,t,e.index+1,this.text,this.text.substring(e.index))},peekToken:function(){if(0===this.tokens.length)throw jn("ueoe","Unexpected end of expression: {0}",this.text);return this.tokens[0]},peek:function(t,e,n,i){if(0<this.tokens.length){var r=this.tokens[0],o=r.text;if(o===t||o===e||o===n||o===i||!t&&!e&&!n&&!i)return r}return!1},expect:function(t,e,n,i){var r=this.peek(t,e,n,i);return!!r&&(this.tokens.shift(),r)},consume:function(t){this.expect(t)||this.throwError("is unexpected, expecting ["+t+"]",this.peek())},unaryFn:function(n,i){return M(function(t,e){return n(t,e,i)},{constant:i.constant})},ternaryFn:function(n,i,r){return M(function(t,e){return n(t,e)?i(t,e):r(t,e)},{constant:n.constant&&i.constant&&r.constant})},binaryFn:function(n,i,r){return M(function(t,e){return i(t,e,n,r)},{constant:n.constant&&r.constant})},statements:function(){for(var o=[];;)if(0<this.tokens.length&&!this.peek("}",")",";","]")&&o.push(this.filterChain()),!this.expect(";"))return 1===o.length?o[0]:function(t,e){for(var n,i=0;i<o.length;i++){var r=o[i];r&&(n=r(t,e))}return n}},filterChain:function(){for(var t,e=this.expression();;){if(!(t=this.expect("|")))return e;e=this.binaryFn(e,t.fn,this.filter())}},filter:function(){for(var t=this.expect(),o=this.$filter(t.text),a=[];;){if(!(t=this.expect(":"))){var e=function(t,e,n){for(var i=[n],r=0;r<a.length;r++)i.push(a[r](t,e));return o.apply(t,i)};return function(){return e}}a.push(this.expression())}},expression:function(){return this.assignment()},assignment:function(){var n,t,i=this.ternary();return(t=this.expect("="))?(i.assign||this.throwError("implies assignment but ["+this.text.substring(0,t.index)+"] can not be assigned to",t),n=this.ternary(),function(t,e){return i.assign(t,n(t,e),e)}):i},ternary:function(){var t,e,n=this.logicalOR();return(e=this.expect("?"))?(t=this.assignment(),(e=this.expect(":"))?this.ternaryFn(n,t,this.assignment()):void this.throwError("expected :",e)):n},logicalOR:function(){for(var t,e=this.logicalAND();;){if(!(t=this.expect("||")))return e;e=this.binaryFn(e,t.fn,this.logicalAND())}},logicalAND:function(){var t,e=this.equality();return(t=this.expect("&&"))&&(e=this.binaryFn(e,t.fn,this.logicalAND())),e},equality:function(){var t,e=this.relational();return(t=this.expect("==","!=","===","!=="))&&(e=this.binaryFn(e,t.fn,this.equality())),e},relational:function(){var t,e=this.additive();return(t=this.expect("<",">","<=",">="))&&(e=this.binaryFn(e,t.fn,this.relational())),e},additive:function(){for(var t,e=this.multiplicative();t=this.expect("+","-");)e=this.binaryFn(e,t.fn,this.multiplicative());return e},multiplicative:function(){for(var t,e=this.unary();t=this.expect("*","/","%");)e=this.binaryFn(e,t.fn,this.unary());return e},unary:function(){var t;return this.expect("+")?this.primary():(t=this.expect("-"))?this.binaryFn(Yn.ZERO,t.fn,this.unary()):(t=this.expect("!"))?this.unaryFn(t.fn,this.unary()):this.primary()},fieldAccess:function(r){var o=this,a=this.expect().text,i=le(a,this.options,this.text);return M(function(t,e,n){return i(n||r(t,e))},{assign:function(t,e,n){var i=r(t,n);return i||r.assign(t,i={}),oe(i,a,e,o.text,o.options)}})},objectIndex:function(a){var s=this,l=this.expression();return this.consume("]"),M(function(t,e){var n,i,r=a(t,e),o=l(t,e);return ie(o,s.text),r?((n=re(r[o],s.text))&&n.then&&s.options.unwrapPromises&&("$$v"in(i=n)||(i.$$v=tt,i.then(function(t){i.$$v=t})),n=n.$$v),n):tt},{assign:function(t,e,n){var i=ie(l(t,n),s.text),r=re(a(t,n),s.text);return r||a.assign(t,r={}),r[i]=e}})},functionCall:function(a,s){var l=[];if(")"!==this.peekToken().text)for(;l.push(this.expression()),this.expect(","););this.consume(")");var u=this;return function(t,e){for(var n=[],i=s?s(t,e):t,r=0;r<l.length;r++)n.push(re(l[r](t,e),u.text));var o=a(t,e,i)||E;return re(i,u.text),function(t,e){if(t){if(t.constructor===t)throw jn("isecfn","Referencing Function in Angular expressions is disallowed! Expression: {0}",e);if(t===Fn||t===Vn||qn&&t===qn)throw jn("isecff","Referencing call, apply or bind in Angular expressions is disallowed! Expression: {0}",e)}}(o,u.text),re(o.apply?o.apply(i,n):o(n[0],n[1],n[2],n[3],n[4]),u.text)}},arrayDeclaration:function(){var r=[],t=!0;if("]"!==this.peekToken().text)do{if(this.peek("]"))break;var e=this.expression();r.push(e),e.constant||(t=!1)}while(this.expect(","));return this.consume("]"),M(function(t,e){for(var n=[],i=0;i<r.length;i++)n.push(r[i](t,e));return n},{literal:!0,constant:t})},object:function(){var o=[],t=!0;if("}"!==this.peekToken().text)do{if(this.peek("}"))break;var e=this.expect(),n=e.string||e.text;this.consume(":");var i=this.expression();o.push({key:n,value:i}),i.constant||(t=!1)}while(this.expect(","));return this.consume("}"),M(function(t,e){for(var n={},i=0;i<o.length;i++){var r=o[i];n[r.key]=r.value(t,e)}return n},{literal:!0,constant:t})}};var Un={},Wn={},Gn=k("$sce"),Zn={HTML:"html",CSS:"css",URL:"url",RESOURCE_URL:"resourceUrl",JS:"js"},Kn=J.createElement("a"),Qn=be(f.location.href);$e.$inject=["$provide"],Ce.$inject=["$locale"],Ee.$inject=["$locale"];var Jn=".",ti={yyyy:De("FullYear",4),yy:De("FullYear",2,0,!0),y:De("FullYear",1),MMMM:_e("Month"),MMM:_e("Month",!0),MM:De("Month",2,1),M:De("Month",1,1),dd:De("Date",2),d:De("Date",1),HH:De("Hours",2),H:De("Hours",1),hh:De("Hours",2,-12),h:De("Hours",1,-12),mm:De("Minutes",2),m:De("Minutes",1),ss:De("Seconds",2),s:De("Seconds",1),sss:De("Milliseconds",3),EEEE:_e("Day"),EEE:_e("Day",!0),a:function(t,e){return t.getHours()<12?e.AMPMS[0]:e.AMPMS[1]},Z:function(t){var e=-1*t.getTimezoneOffset();return(0<=e?"+":"")+(Te(Math[0<e?"floor":"ceil"](e/60),2)+Te(Math.abs(e%60),2))}},ei=/((?:[^yMdHhmsaZE']+)|(?:'(?:[^']|'')*')|(?:E+|y+|M+|d+|H+|h+|m+|s+|a|Z))(.*)/,ni=/^\-?\d+$/;Pe.$inject=["$locale"];var ii=y(Ve),ri=y(ze);Ne.$inject=["$parse"];var oi=y({restrict:"E",compile:function(t,e){return He<=8&&(e.href||e.name||e.$set("href",""),t.append(J.createComment("IE fix"))),e.href||e.xlinkHref||e.name?void 0:function(t,e){var n="[object SVGAnimatedString]"===Ke.call(e.prop("href"))?"xlink:href":"href";e.on("click",function(t){e.attr(n)||t.preventDefault()})}}}),ai={};et(xn,function(t,i){if("multiple"!=t){var r=_t("ng-"+i);ai[r]=function(){return{priority:100,link:function(t,e,n){t.$watch(n[r],function(t){n.$set(i,!!t)})}}}}}),et(["src","srcset","href"],function(o){var a=_t("ng-"+o);ai[a]=function(){return{priority:99,link:function(t,e,n){var i=o,r=o;"href"===o&&"[object SVGAnimatedString]"===Ke.call(e.prop("href"))&&(r="xlinkHref",n.$attr[r]="xlink:href",i=null),n.$observe(a,function(t){return t?(n.$set(r,t),void(He&&i&&e.prop(i,n[r]))):void("href"===o&&n.$set(r,null))})}}}});var si={$addControl:E,$removeControl:E,$setValidity:E,$setDirty:E,$setPristine:E};Ie.$inject=["$element","$attrs","$scope","$animate"];function li(t){return["$timeout",function(s){return{name:"form",restrict:t?"EAC":"E",controller:Ie,compile:function(){return{pre:function(t,e,n,i){if(!n.action){var r=function(t){t.preventDefault?t.preventDefault():t.returnValue=!1};un(e[0],"submit",r),e.on("$destroy",function(){s(function(){cn(e[0],"submit",r)},0,!1)})}var o=e.parent().controller("form"),a=n.name||n.ngForm;a&&oe(t,a,i,a),o&&e.on("$destroy",function(){o.$removeControl(i),a&&oe(t,a,tt,a),M(i,si)})}}}}}]}function ui(){return{require:["ngModel","^?form"],controller:Ei,link:function(t,e,n,i){var r=i[0],o=i[1]||si;o.$addControl(r),t.$on("$destroy",function(){o.$removeControl(r)})}}}function ci(){return{require:"?ngModel",link:function(t,e,n,i){if(i){n.required=!0;var r=function(t){return n.required&&i.$isEmpty(t)?void i.$setValidity("required",!1):(i.$setValidity("required",!0),t)};i.$formatters.push(r),i.$parsers.unshift(r),n.$observe("required",function(){r(i.$viewValue)})}}}}function hi(){return{require:"ngModel",link:function(t,e,n,i){var r=/\/(.*)\//.exec(n.ngList),o=r&&new RegExp(r[1])||n.ngList||",";i.$parsers.push(function(t){if(!S(t)){var e=[];return t&&et(t.split(o),function(t){t&&e.push(nn(t))}),e}}),i.$formatters.push(function(t){return en(t)?t.join(", "):tt}),i.$isEmpty=function(t){return!t||!t.length}}}}function di(){return{priority:100,compile:function(t,e){return Ti.test(e.ngValue)?function(t,e,n){n.$set("value",t.$eval(n.ngValue))}:function(t,e,n){t.$watch(n.ngValue,function(t){n.$set("value",t)})}}}}var pi=li(),fi=li(!0),gi=/^(ftp|http|https):\/\/(\w+:{0,1}\w*@)?(\S+)(:[0-9]+)?(\/|\/([\w#!:.?+=&%@!\-\/]))?$/,vi=/^[a-z0-9!#$%&'*+\/=?^_`{|}~.-]+@[a-z0-9]([a-z0-9-]*[a-z0-9])?(\.[a-z0-9]([a-z0-9-]*[a-z0-9])?)*$/i,mi=/^\s*(\-|\+)?(\d+|(\d*(\.\d*)))\s*$/,yi={text:je,number:function(t,e,n,i,r,o){if(je(t,e,n,i,r,o),i.$parsers.push(function(t){var e=i.$isEmpty(t);return e||mi.test(t)?(i.$setValidity("number",!0),""===t?null:e?t:parseFloat(t)):(i.$setValidity("number",!1),tt)}),function(e,n,i,r,o){if(nt(o)){e.$$hasNativeValidators=!0;e.$parsers.push(function(t){return e.$error[n]||Le(o,r)||!Le(o,i)?t:void e.$setValidity(n,!1)})}}(i,"number",bi,null,i.$$validityState),i.$formatters.push(function(t){return i.$isEmpty(t)?"":""+t}),n.min){var a=function(t){var e=parseFloat(n.min);return Be(i,"min",i.$isEmpty(t)||e<=t,t)};i.$parsers.push(a),i.$formatters.push(a)}if(n.max){var s=function(t){var e=parseFloat(n.max);return Be(i,"max",i.$isEmpty(t)||t<=e,t)};i.$parsers.push(s),i.$formatters.push(s)}i.$formatters.push(function(t){return Be(i,"number",i.$isEmpty(t)||l(t),t)})},url:function(t,e,n,i,r,o){function a(t){return Be(i,"url",i.$isEmpty(t)||gi.test(t),t)}je(t,e,n,i,r,o),i.$formatters.push(a),i.$parsers.push(a)},email:function(t,e,n,i,r,o){function a(t){return Be(i,"email",i.$isEmpty(t)||vi.test(t),t)}je(t,e,n,i,r,o),i.$formatters.push(a),i.$parsers.push(a)},radio:function(t,e,n,i){S(n.name)&&e.attr("name",r()),e.on("click",function(){e[0].checked&&t.$apply(function(){i.$setViewValue(n.value)})}),i.$render=function(){var t=n.value;e[0].checked=t==i.$viewValue},n.$observe("value",i.$render)},checkbox:function(t,e,n,i){var r=n.ngTrueValue,o=n.ngFalseValue;it(r)||(r=!0),it(o)||(o=!1),e.on("click",function(){t.$apply(function(){i.$setViewValue(e[0].checked)})}),i.$render=function(){e[0].checked=i.$viewValue},i.$isEmpty=function(t){return t!==r},i.$formatters.push(function(t){return t===r}),i.$parsers.push(function(t){return t?r:o})},hidden:E,button:E,submit:E,reset:E,file:E},bi=["badInput"],xi=["$browser","$sniffer",function(r,o){return{restrict:"E",require:"?ngModel",link:function(t,e,n,i){i&&(yi[Ve(n.type)]||yi.text)(t,e,n,i,o,r)}}}],wi="ng-valid",$i="ng-invalid",ki="ng-pristine",Ci="ng-dirty",Ei=["$scope","$exceptionHandler","$attrs","$element","$parse","$animate",function(i,n,t,r,e,o){function a(t,e){e=e?"-"+ut(e,"-"):"",o.removeClass(r,(t?$i:wi)+e),o.addClass(r,(t?wi:$i)+e)}this.$viewValue=Number.NaN,this.$modelValue=Number.NaN,this.$parsers=[],this.$formatters=[],this.$viewChangeListeners=[],this.$pristine=!0,this.$dirty=!1,this.$valid=!0,this.$invalid=!1,this.$name=t.name;var s=e(t.ngModel),l=s.assign;if(!l)throw k("ngModel")("nonassign","Expression '{0}' is non-assignable. Element: {1}",t.ngModel,lt(r));this.$render=E,this.$isEmpty=function(t){return S(t)||""===t||null===t||t!=t};var u=r.inheritedData("$formController")||si,c=0,h=this.$error={};r.addClass(ki),a(!0),this.$setValidity=function(t,e){h[t]!==!e&&(e?(h[t]&&c--,c||(a(!0),this.$valid=!0,this.$invalid=!1)):(a(!1),this.$invalid=!0,this.$valid=!1,c++),h[t]=!e,a(e,t),u.$setValidity(t,e,this))},this.$setPristine=function(){this.$dirty=!1,this.$pristine=!0,o.removeClass(r,Ci),o.addClass(r,ki)},this.$setViewValue=function(e){this.$viewValue=e,this.$pristine&&(this.$dirty=!0,this.$pristine=!1,o.removeClass(r,ki),o.addClass(r,Ci),u.$setDirty()),et(this.$parsers,function(t){e=t(e)}),this.$modelValue!==e&&(this.$modelValue=e,l(i,e),et(this.$viewChangeListeners,function(t){try{t()}catch(t){n(t)}}))};var d=this;i.$watch(function(){var t=s(i);if(d.$modelValue!==t){var e=d.$formatters,n=e.length;for(d.$modelValue=t;n--;)t=e[n](t);d.$viewValue!==t&&(d.$viewValue=t,d.$render())}return t})}],Si=y({require:"ngModel",link:function(t,e,n,i){i.$viewChangeListeners.push(function(){t.$eval(n.ngChange)})}}),Ti=/^(true|false|\d+)$/,Di=Oe({compile:function(t){return t.addClass("ng-binding"),function(t,e,n){e.data("$binding",n.ngBind),t.$watch(n.ngBind,function(t){e.text(t==tt?"":t)})}}}),_i=["$interpolate",function(r){return function(t,e,n){var i=r(e.attr(n.$attr.ngBindTemplate));e.addClass("ng-binding").data("$binding",i),n.$observe("ngBindTemplate",function(t){e.text(t)})}}],Pi=["$sce","$parse",function(r,o){return{compile:function(t){return t.addClass("ng-binding"),function(e,n,t){n.data("$binding",t.ngBindHtml);var i=o(t.ngBindHtml);e.$watch(function(){return(i(e)||"").toString()},function(t){n.html(r.getTrustedHtml(i(e))||"")})}}}}],Mi=Re("",!0),Ai=Re("Odd",0),Ni=Re("Even",1),Oi=Oe({compile:function(t,e){e.$set("ngCloak",tt),t.removeClass("ng-cloak")}}),Ii=[function(){return{scope:!0,controller:"@",priority:500}}],Bi={},Li={blur:!0,focus:!0};et("click dblclick mousedown mouseup mouseover mouseout mousemove mouseenter mouseleave keydown keyup keypress submit focus blur copy cut paste".split(" "),function(o){var a=_t("ng-"+o);Bi[a]=["$parse","$rootScope",function(n,r){return{compile:function(t,e){var i=n(e[a],!0);return function(n,t){t.on(o,function(t){function e(){i(n,{$event:t})}Li[o]&&r.$$phase?n.$evalAsync(e):n.$apply(e)})}}}}]});var ji=["$animate",function(l){return{transclude:"element",priority:600,terminal:!0,restrict:"A",$$tlb:!0,link:function(e,n,i,t,r){var o,a,s;e.$watch(i.ngIf,function(t){N(t)?a||(a=e.$new(),r(a,function(t){t[t.length++]=J.createComment(" end ngIf: "+i.ngIf+" "),o={clone:t},l.enter(t,n.parent(),n)})):(s&&(s.remove(),s=null),a&&(a.$destroy(),a=null),o&&(s=V(o.clone),l.leave(s,function(){s=null}),o=null))})}}}],Ri=["$http","$templateCache","$anchorScroll","$animate","$sce",function(g,v,m,y,i){return{restrict:"ECA",priority:400,terminal:!0,transclude:"element",controller:Je.noop,compile:function(t,e){var n=e.ngInclude||e.src,p=e.onload||"",f=e.autoscroll;return function(o,a,t,s,l){function u(){e&&(e.remove(),e=null),c&&(c.$destroy(),c=null),h&&(y.leave(h,function(){e=null}),e=h,h=null)}var c,e,h,d=0;o.$watch(i.parseAsResourceUrl(n),function(t){function i(){!F(f)||f&&!o.$eval(f)||m()}var r=++d;t?(g.get(t,{cache:v}).success(function(t){if(r===d){var e=o.$new();s.template=t;var n=l(e,function(t){u(),y.enter(t,null,a,i)});h=n,(c=e).$emit("$includeContentLoaded"),o.$eval(p)}}).error(function(){r===d&&u()}),o.$emit("$includeContentRequested")):(u(),s.template=null)})}}}}],Fi=["$compile",function(r){return{restrict:"ECA",priority:-400,require:"ngInclude",link:function(t,e,n,i){e.html(i.template),r(e.contents())(t)}}}],Vi=Oe({priority:450,compile:function(){return{pre:function(t,e,n){t.$eval(n.ngInit)}}}}),qi=Oe({terminal:!0,priority:1e3}),zi=["$locale","$interpolate",function(h,d){var p=/{}/g;return{restrict:"EA",link:function(e,n,i){var r=i.count,t=i.$attr.when&&n.attr(i.$attr.when),o=i.offset||0,a=e.$eval(t)||{},s={},l=d.startSymbol(),u=d.endSymbol(),c=/^when(Minus)?(.+)$/;et(i,function(t,e){c.test(e)&&(a[Ve(e.replace("when","").replace("Minus","-"))]=n.attr(i.$attr[e]))}),et(a,function(t,e){s[e]=d(t.replace(p,l+r+"-"+o+u))}),e.$watch(function(){var t=parseFloat(e.$eval(r));return isNaN(t)?"":(t in a||(t=h.pluralCat(t-o)),s[t](e,n,!0))},function(t){n.text(t)})}}}],Hi=["$parse","$animate",function(l,S){function T(t){return t.clone[t.clone.length-1]}var D="$$NG_REMOVED",_=k("ngRepeat");return{transclude:"element",priority:1e3,terminal:!0,$$tlb:!0,link:function(v,m,t,e,y){var n,i,b,x,w,r,o,$,k,C=t.ngRepeat,a=C.match(/^\s*([\s\S]+?)\s+in\s+([\s\S]+?)(?:\s+track\s+by\s+([\s\S]+?))?\s*$/),s={$id:yt};if(!a)throw _("iexp","Expected expression in form of '_item_ in _collection_[ track by _id_]' but got '{0}'.",C);if(r=a[1],o=a[2],(n=a[3])?(i=l(n),b=function(t,e,n){return k&&(s[k]=t),s[$]=e,s.$index=n,i(v,s)}):(x=function(t,e){return yt(e)},w=function(t){return t}),!(a=r.match(/^(?:([\$\w]+)|\(([\$\w]+)\s*,\s*([\$\w]+)\))$/)))throw _("iidexp","'_item_' in '_item_ in _collection_' should be an identifier or '(_key_, _value_)' expression, but got '{0}'.",r);$=a[3]||a[1],k=a[2];var E={};v.$watchCollection(o,function(t){var e,n,i,r,o,a,s,l,u,c,h,d,p=m[0],f={},g=[];if(P(t))c=t,u=b||x;else{for(a in u=b||w,c=[],t)t.hasOwnProperty(a)&&"$"!=a.charAt(0)&&c.push(a);c.sort()}for(r=c.length,n=g.length=c.length,e=0;e<n;e++)if(L(l=u(a=t===c?e:c[e],s=t[a],e),"`track by` id"),E.hasOwnProperty(l))h=E[l],delete E[l],f[l]=h,g[e]=h;else{if(f.hasOwnProperty(l))throw et(g,function(t){t&&t.scope&&(E[t.id]=t)}),_("dupes","Duplicates in a repeater are not allowed. Use 'track by' expression to specify unique keys. Repeater: {0}, Duplicate key: {1}, Duplicate value: {2}",C,l,A(s));g[e]={id:l},f[l]=!1}for(a in E)E.hasOwnProperty(a)&&(d=V((h=E[a]).clone),S.leave(d),et(d,function(t){t[D]=!0}),h.scope.$destroy());for(e=0,n=c.length;e<n;e++){if(s=t[a=t===c?e:c[e]],h=g[e],g[e-1]&&(p=T(g[e-1])),h.scope){for(o=h.scope,i=p;(i=i.nextSibling)&&i[D];);h.clone[0]!=i&&S.move(V(h.clone),null,Xe(p)),p=T(h)}else o=v.$new();o[$]=s,k&&(o[k]=a),o.$index=e,o.$first=0===e,o.$last=e===r-1,o.$middle=!(o.$first||o.$last),o.$odd=!(o.$even=0==(1&e)),h.scope||y(o,function(t){t[t.length++]=J.createComment(" end ngRepeat: "+C+" "),S.enter(t,null,Xe(p)),p=t,h.scope=o,h.clone=t,f[h.id]=h})}E=f})}}}],Xi=["$animate",function(i){return function(t,e,n){t.$watch(n.ngShow,function(t){i[N(t)?"removeClass":"addClass"](e,"ng-hide")})}}],Yi=["$animate",function(i){return function(t,e,n){t.$watch(n.ngHide,function(t){i[N(t)?"addClass":"removeClass"](e,"ng-hide")})}}],Ui=Oe(function(t,n,e){t.$watch(e.ngStyle,function(t,e){e&&t!==e&&et(e,function(t,e){n.css(e,"")}),t&&n.css(t)},!0)}),Wi=["$animate",function(h){return{restrict:"EA",require:"ngSwitch",controller:["$scope",function(){this.cases={}}],link:function(r,t,o,a){var e=o.ngSwitch||o.on,s=[],l=[],u=[],c=[];r.$watch(e,function(t){var e,n;for(e=0,n=u.length;e<n;++e)u[e].remove();for(u.length=0,e=0,n=c.length;e<n;++e){var i=l[e];c[e].$destroy(),u[e]=i,h.leave(i,function(){u.splice(e,1)})}l.length=0,c.length=0,(s=a.cases["!"+t]||a.cases["?"])&&(r.$eval(o.change),et(s,function(n){var t=r.$new();c.push(t),n.transclude(t,function(t){var e=n.element;l.push(t),h.enter(t,e.parent(),e)})}))})}}}],Gi=Oe({transclude:"element",priority:800,require:"^ngSwitch",link:function(t,e,n,i,r){i.cases["!"+n.ngSwitchWhen]=i.cases["!"+n.ngSwitchWhen]||[],i.cases["!"+n.ngSwitchWhen].push({transclude:r,element:e})}}),Zi=Oe({transclude:"element",priority:800,require:"^ngSwitch",link:function(t,e,n,i,r){i.cases["?"]=i.cases["?"]||[],i.cases["?"].push({transclude:r,element:e})}}),Ki=Oe({link:function(t,e,n,i,r){if(!r)throw k("ngTransclude")("orphan","Illegal use of ngTransclude directive in the template! No parent directive that requires a transclusion found. Element: {0}",lt(e));r(function(t){e.empty(),e.append(t)})}}),Qi=["$templateCache",function(r){return{restrict:"E",terminal:!0,compile:function(t,e){if("text/ng-template"==e.type){var n=e.id,i=t[0].text;r.put(n,i)}}}}],Ji=k("ngOptions"),tr=y({terminal:!0}),er=["$compile","$parse",function(b,x){var w=/^\s*([\s\S]+?)(?:\s+as\s+([\s\S]+?))?(?:\s+group\s+by\s+([\s\S]+?))?\s+for\s+(?:([\$\w][\$\w]*)|(?:\(\s*([\$\w][\$\w]*)\s*,\s*([\$\w][\$\w]*)\s*\)))\s+in\s+([\s\S]+?)(?:\s+track\s+by\s+([\s\S]+?))?$/,s={$setViewValue:E};return{restrict:"E",require:["select","?ngModel"],controller:["$element","$scope","$attrs",function(n,t,e){var i,r=this,o={},a=s;r.databound=e.ngModel,r.init=function(t,e,n){a=t,e,i=n},r.addOption=function(t){L(t,'"option value"'),o[t]=!0,a.$viewValue==t&&(n.val(t),i.parent()&&i.remove())},r.removeOption=function(t){this.hasOption(t)&&(delete o[t],a.$viewValue==t&&this.renderUnknownOption(t))},r.renderUnknownOption=function(t){var e="? "+yt(t)+" ?";i.val(e),n.prepend(i),n.val(e),i.prop("selected",!0)},r.hasOption=function(t){return o.hasOwnProperty(t)},t.$on("$destroy",function(){r.renderUnknownOption=E})}],link:function(t,e,n,i){if(i[1]){for(var r,O=i[0],o=i[1],I=n.multiple,a=n.ngOptions,B=!1,L=Xe(J.createElement("option")),j=Xe(J.createElement("optgroup")),s=L.clone(),l=0,u=e.children(),c=u.length;l<c;l++)if(""===u[l].value){r=B=u.eq(l);break}O.init(o,B,s),I&&(o.$isEmpty=function(t){return!t||0===t.length}),a?function(k,C,E){function h(){var t,e,n,i,r,o,a,s,l,u,c,h,d,p,f,g={"":[]},v=[""],m=E.$modelValue,y=M(k)||[],b=D?R(y):y,x={},w=function(){var t=!1;if(I){var e=E.$modelValue;if(A&&en(e)){t=new bt([]);for(var n={},i=0;i<e.length;i++)n[T]=e[i],t.put(A(k,n),e[i])}else t=new bt(e)}return t}();for(c=0;c<(l=b.length);c++){if(a=c,D){if("$"===(a=b[c]).charAt(0))continue;x[D]=a}if(x[T]=y[a],(e=g[t=_(k,x)||""])||(e=g[t]=[],v.push(t)),I)h=F(w.remove(A?A(k,x):P(k,x)));else{if(A){var $={};$[T]=m,h=A(k,$)===A(k,x)}else h=m===P(k,x);w=w||h}f=F(f=S(k,x))?f:"",e.push({id:A?A(k,x):D?b[c]:c,label:f,selected:h})}for(I||(B||null===m?g[""].unshift({id:"",label:"",selected:!w}):w||g[""].unshift({id:"?",label:"",selected:!0})),u=0,s=v.length;u<s;u++){for(e=g[t=v[u]],N.length<=u?(r=[i={element:j.clone().attr("label",t),label:e.label}],N.push(r),C.append(i.element)):(i=(r=N[u])[0]).label!=t&&i.element.attr("label",i.label=t),d=null,c=0,l=e.length;c<l;c++)n=e[c],(o=r[c+1])?(d=o.element,o.label!==n.label&&(d.text(o.label=n.label),d.prop("label",o.label)),o.id!==n.id&&d.val(o.id=n.id),d[0].selected!==n.selected&&(d.prop("selected",o.selected=n.selected),He&&d.prop("selected",o.selected))):(""===n.id&&B?p=B:(p=L.clone()).val(n.id).prop("selected",n.selected).attr("selected",n.selected).prop("label",n.label).text(n.label),r.push(o={element:p,label:n.label,id:n.id,selected:n.selected}),O.addOption(n.label,p),d?d.after(p):i.element.append(p),d=p);for(c++;r.length>c;)n=r.pop(),O.removeOption(n.label),n.element.remove()}for(;N.length>u;)N.pop()[0].element.remove()}var t;if(!(t=a.match(w)))throw Ji("iexp","Expected expression in form of '_select_ (as _label_)? for (_key_,)?_value_ in _collection_' but got '{0}'. Element: {1}",a,lt(C));var S=x(t[2]||t[1]),T=t[4]||t[6],D=t[5],_=x(t[3]||""),P=x(t[2]?t[1]:T),M=x(t[7]),A=t[8]?x(t[8]):null,N=[[{element:C,label:""}]];B&&(b(B)(k),B.removeClass("ng-scope"),B.remove()),C.empty(),C.on("change",function(){k.$apply(function(){var t,e,n,i,r,o,a,s,l,u=M(k)||[],c={};if(I){for(n=[],o=0,s=N.length;o<s;o++)for(r=1,a=(t=N[o]).length;r<a;r++)if((i=t[r].element)[0].selected){if(e=i.val(),D&&(c[D]=e),A)for(l=0;l<u.length&&(c[T]=u[l],A(k,c)!=e);l++);else c[T]=u[e];n.push(P(k,c))}}else if("?"==(e=C.val()))n=tt;else if(""===e)n=null;else if(A){for(l=0;l<u.length;l++)if(c[T]=u[l],A(k,c)==e){n=P(k,c);break}}else c[T]=u[e],D&&(c[D]=e),n=P(k,c);E.$setViewValue(n),h()})}),E.$render=h,k.$watchCollection(M,h),k.$watchCollection(function(){var t={},e=M(k);if(e){for(var n=new Array(e.length),i=0,r=e.length;i<r;i++)t[T]=e[i],n[i]=S(k,t);return n}},h),I&&k.$watchCollection(function(){return E.$modelValue},h)}(t,e,o):I?(g=t,v=e,(m=o).$render=function(){var e=new bt(m.$viewValue);et(v.find("option"),function(t){t.selected=F(e.get(t.value))})},g.$watch(function(){at(y,m.$viewValue)||(y=ot(m.$viewValue),m.$render())}),v.on("change",function(){g.$apply(function(){var e=[];et(v.find("option"),function(t){t.selected&&e.push(t.value)}),m.$setViewValue(e)})})):(h=t,d=e,f=O,(p=o).$render=function(){var t=p.$viewValue;f.hasOption(t)?(s.parent()&&s.remove(),d.val(t),""===t&&r.prop("selected",!0)):S(t)&&r?d.val(""):f.renderUnknownOption(t)},d.on("change",function(){h.$apply(function(){s.parent()&&s.remove(),p.$setViewValue(d.val())})}))}var h,d,p,f,g,v,m,y}}}],nr=["$interpolate",function(n){var s={addOption:E,removeOption:E};return{restrict:"E",priority:100,compile:function(t,e){if(S(e.value)){var a=n(t.text(),!0);a||e.$set("value",t.text())}return function(t,e,n){var i="$selectController",r=e.parent(),o=r.data(i)||r.parent().data(i);o&&o.databound?e.prop("selected",!1):o=s,a?t.$watch(a,function(t,e){n.$set("value",t),t!==e&&o.removeOption(e),o.addOption(t)}):o.addOption(n.value),e.on("$destroy",function(){o.removeOption(n.value)})}}}}],ir=y({restrict:"E",terminal:!0});f.angular.bootstrap?console.log("WARNING: Tried to load angular more than once."):((Ye=f.jQuery)&&Ye.fn.on?(M((Xe=Ye).fn,{scope:bn.scope,isolateScope:bn.isolateScope,controller:bn.controller,injector:bn.injector,inheritedData:bn.inheritedData}),t("remove",!0,!0,!1),t("empty",!1,!1,!1),t("html",!1,!1,!0)):Xe=H,Je.element=Xe,function(t){M(t,{bootstrap:I,copy:T,extend:M,equals:at,element:Xe,forEach:et,injector:wt,noop:E,bind:_,toJson:A,fromJson:a,identity:m,isUndefined:S,isDefined:F,isString:it,isFunction:rt,isObject:nt,isNumber:l,isElement:e,isArray:en,version:an,isDate:b,lowercase:Ve,uppercase:ze,callbacks:{counter:0},$$minErr:k,$$csp:rn}),Ue=function(t){function e(t,e,n){return t[e]||(t[e]=n())}var l=k("$injector"),n=k("ng"),i=e(t,"angular",Object);return i.$$minErr=i.$$minErr||k,e(i,"module",function(){var t={};return function(o,a,s){return function(t,e){if("hasOwnProperty"===t)throw n("badname","hasOwnProperty is not a valid {0} name",e)}(o,"module"),a&&t.hasOwnProperty(o)&&(t[o]=null),e(t,o,function(){function t(t,e,n){return function(){return i[n||"push"]([t,e,arguments]),r}}if(!a)throw l("nomod","Module '{0}' is not available! You either misspelled the module name or forgot to load it. If registering a module ensure that you specify the dependencies as the second argument.",o);var i=[],e=[],n=t("$injector","invoke"),r={_invokeQueue:i,_runBlocks:e,requires:a,name:o,provider:t("$provide","provider"),factory:t("$provide","factory"),service:t("$provide","service"),value:t("$provide","value"),constant:t("$provide","constant","unshift"),animation:t("$animateProvider","register"),filter:t("$filterProvider","register"),controller:t("$controllerProvider","register"),directive:t("$compileProvider","directive"),config:n,run:function(t){return e.push(t),this}};return s&&n(s),r})}})}(f);try{Ue("ngLocale")}catch(t){Ue("ngLocale",[]).provider("$locale",zt)}Ue("ng",["ngLocale"],["$provide",function(t){t.provider({$$sanitizeUri:pe}),t.provider("$compile",Dt).directive({a:oi,input:xi,textarea:xi,form:pi,script:Qi,select:er,style:ir,option:nr,ngBind:Di,ngBindHtml:Pi,ngBindTemplate:_i,ngClass:Mi,ngClassEven:Ni,ngClassOdd:Ai,ngCloak:Oi,ngController:Ii,ngForm:fi,ngHide:Yi,ngIf:ji,ngInclude:Ri,ngInit:Vi,ngNonBindable:qi,ngPluralize:zi,ngRepeat:Hi,ngShow:Xi,ngStyle:Ui,ngSwitch:Wi,ngSwitchWhen:Gi,ngSwitchDefault:Zi,ngOptions:tr,ngTransclude:Ki,ngModel:ui,ngList:hi,ngChange:Si,required:ci,ngRequired:ci,ngValue:di}).directive({ngInclude:Fi}).directive(ai).directive(Bi),t.provider({$anchorScroll:$t,$animate:Pn,$browser:Et,$cacheFactory:St,$controller:Mt,$document:At,$exceptionHandler:Nt,$filter:$e,$interpolate:Vt,$interval:qt,$http:jt,$httpBackend:Ft,$location:ee,$log:ne,$parse:ue,$rootScope:de,$q:ce,$sce:ve,$sceDelegate:ge,$sniffer:me,$templateCache:Tt,$timeout:ye,$window:we,$$rAF:he,$$asyncCallback:kt})}])}(Je),Xe(J).ready(function(){!function(e,t){function n(t){t&&o.push(t)}var i,r,o=[e],a=["ng:app","ng-app","x-ng-app","data-ng-app"],s=/\sng[:\-]app(:\s*([\w\d_]+);?)?\s/;et(a,function(t){a[t]=!0,n(J.getElementById(t)),t=t.replace(":","\\:"),e.querySelectorAll&&(et(e.querySelectorAll("."+t),n),et(e.querySelectorAll("."+t+"\\:"),n),et(e.querySelectorAll("["+t+"]"),n))}),et(o,function(e){if(!i){var t=" "+e.className+" ",n=s.exec(t);n?(i=e,r=(n[2]||"").replace(/\s+/g,",")):et(e.attributes,function(t){!i&&a[t.name]&&(i=e,r=t.value)})}}),i&&t(i,r?[r]:[])}(J,I)}))}(window,document),window.angular.$$csp()||window.angular.element(document).find("head").prepend('<style type="text/css">@charset "UTF-8";[ng\\:cloak],[ng-cloak],[data-ng-cloak],[x-ng-cloak],.ng-cloak,.x-ng-cloak,.ng-hide{display:none !important;}ng\\:form{display:block;}.ng-animate-block-transitions{transition:0s all!important;-webkit-transition:0s all!important;}.ng-hide-add-active,.ng-hide-remove{display:block!important;}</style>'),function(e,X,Y){"use strict";X.module("ngAnimate",["ng"]).directive("ngAnimateChildren",function(){var r="$$ngAnimateChildren";return function(t,e,n){var i=n.ngAnimateChildren;X.isString(i)&&0===i.length?e.data(r,!0):t.$watch(i,function(t){e.data(r,!!t)})}}).factory("$$animateReflow",["$$rAF","$document",function(e,t){var n=t[0].body;return function(t){return e(function(){n.offsetWidth,t()})}}]).config(["$provide","$animateProvider",function(t,n){function V(t){for(var e=0;e<t.length;e++){var n=t[e];if(n.nodeType==H)return n}}function u(t){return t&&X.element(t)}function c(t){return X.element(V(t))}function P(t,e){return V(t)==V(e)}var q=X.noop,z=X.forEach,w=n.$$selectors,H=1,M="$$ngAnimateState",A="ng-animate",N={running:!0};t.decorator("$animate",["$delegate","$injector","$sniffer","$rootElement","$$asyncCallback","$rootScope","$document",function(r,y,b,E,S,o,t){function a(t){var e=t.data(M)||{};e.running=!0,t.data(M,e)}function T(s,e,l){function n(t,e){var n=t[e],i=t["before"+e.charAt(0).toUpperCase()+e.substr(1)];return n||i?("leave"==e&&(i=n,n=null),v.push({event:e,fn:n}),p.push({event:e,fn:i}),!0):void 0}function i(t,i,r){var o=[];z(t,function(t){t.fn&&o.push(t)});var a=0;z(o,function(t,e){function n(){!function(t){if(i){if((i[t]||q)(),++a<o.length)return;i=null}r()}(e)}switch(t.event){case"setClass":i.push(t.fn(s,u,c,n));break;case"addClass":i.push(t.fn(s,u||l,n));break;case"removeClass":i.push(t.fn(s,c||l,n));break;default:i.push(t.fn(s,n))}}),i&&0===i.length&&r()}var t=s[0];if(t){var u,c,r="setClass"==e,o=r||"addClass"==e||"removeClass"==e;X.isArray(l)&&(u=l[0],c=l[1],l=u+" "+c);var a=s.attr("class")+" "+l;if(x(a)){var h=q,d=[],p=[],f=q,g=[],v=[],m=(" "+a).replace(/\s+/g,".");return z(function(t){if(t){var e=[],n={},i=t.substr(1).split(".");(b.transitions||b.animations)&&e.push(y.get(w[""]));for(var r=0;r<i.length;r++){var o=i[r],a=w[o];a&&!n[o]&&(e.push(y.get(a)),n[o]=!0)}return e}}(m),function(t){!n(t,e)&&r&&(n(t,"addClass"),n(t,"removeClass"))}),{node:t,event:e,className:l,isClassBased:o,isSetClassOperation:r,before:function(t){h=t,i(p,d,function(){h=q,t()})},after:function(t){f=t,i(v,g,function(){f=q,t()})},cancel:function(){z(d,function(t){(t||q)(!0)}),h(!0),z(g,function(t){(t||q)(!0)}),f(!0)}}}}}function s(n,i,r,t,e,o,a){function s(t){var e="$animate:"+t;f&&f[e]&&0<f[e].length&&S(function(){r.triggerHandler(e,{event:n,className:i})})}function l(){s("before")}function u(){s("after")}function c(){s("close"),a&&S(function(){a()})}function h(){h.hasBeenRun||(h.hasBeenRun=!0,o())}function d(){if(!d.hasBeenRun){d.hasBeenRun=!0;var t=r.data(M);t&&(p&&p.isClassBased?D(r,i):(S(function(){var t=r.data(M)||{};C==t.index&&D(r,i)}),r.data(M,t))),c()}}var p=T(r,n,i);if(!p)return h(),l(),u(),void d();i=p.className;var f=X.element._data(p.node);f=f&&f.events,t||(t=e?e.parent():r.parent());var g,v=r.data(M)||{},m=v.active||{},y=v.totalActive||0,b=v.last;if(p.isClassBased&&(g=v.running||v.disabled||b&&!b.isClassBased),g||function(t,e){if(N.disabled)return!0;if(P(t,E))return N.running;var n,i,r;do{if(0===e.length)break;var o=P(e,E),a=o?N:e.data(M)||{};if(a.disabled)return!0;if(o&&(r=!0),!1!==n){var s=e.data("$$ngAnimateChildren");X.isDefined(s)&&(n=s)}i=i||a.running||a.last&&!a.last.isClassBased}while(e=e.parent());return!r||!n&&i}(r,t))return h(),l(),u(),void d();var x=!1;if(0<y){var w=[];if(p.isClassBased){if("setClass"==b.event)w.push(b),D(r,i);else if(m[i]){var $=m[i];$.event==n?x=!0:(w.push($),D(r,i))}}else if("leave"==n&&m["ng-leave"])x=!0;else{for(var k in m)w.push(m[k]),D(r,k);m={},y=0}0<w.length&&z(w,function(t){t.cancel()})}if(!p.isClassBased||p.isSetClassOperation||x||(x="addClass"==n==r.hasClass(i)),x)return h(),l(),u(),void c();"leave"==n&&r.one("$destroy",function(t){var e=X.element(this),n=e.data(M);if(n){var i=n.active["ng-leave"];i&&(i.cancel(),D(e,"ng-leave"))}}),r.addClass(A);var C=_++;y++,m[i]=p,r.data(M,{last:p,active:m,index:C,totalActive:y}),l(),p.before(function(t){var e=r.data(M);t=t||!e||!e.active[i]||p.isClassBased&&e.active[i].event!=n,h(),!0===t?d():(u(),p.after(d))})}function l(t){var e=V(t);if(e){var n=X.isFunction(e.getElementsByClassName)?e.getElementsByClassName(A):e.querySelectorAll("."+A);z(n,function(t){var e=(t=X.element(t)).data(M);e&&e.active&&z(e.active,function(t){t.cancel()})})}}function D(t,e){if(P(t,E))N.disabled||(N.running=!1,N.structural=!1);else if(e){var n=t.data(M)||{},i=!0===e;!i&&n.active&&n.active[e]&&(n.totalActive--,delete n.active[e]),!i&&n.totalActive||(t.removeClass(A),t.removeData(M))}}var _=0;E.data(M,N),o.$$postDigest(function(){o.$$postDigest(function(){N.running=!1})});var e=n.classNameFilter(),x=e?function(t){return e.test(t)}:function(){return!0};return{enter:function(t,e,n,i){t=X.element(t),e=u(e),n=u(n),a(t),r.enter(t,e,n),o.$$postDigest(function(){s("enter","ng-enter",t=c(t),e,n,q,i)})},leave:function(t,e){l(t=X.element(t)),a(t),o.$$postDigest(function(){s("leave","ng-leave",c(t),null,null,function(){r.leave(t)},e)})},move:function(t,e,n,i){t=X.element(t),e=u(e),n=u(n),l(t),a(t),r.move(t,e,n),o.$$postDigest(function(){s("move","ng-move",t=c(t),e,n,q,i)})},addClass:function(t,e,n){t=c(t=X.element(t)),s("addClass",e,t,null,null,function(){r.addClass(t,e)},n)},removeClass:function(t,e,n){t=c(t=X.element(t)),s("removeClass",e,t,null,null,function(){r.removeClass(t,e)},n)},setClass:function(t,e,n,i){t=c(t=X.element(t)),s("setClass",[e,n],t,null,null,function(){r.setClass(t,e,n)},i)},enabled:function(t,e){switch(arguments.length){case 2:if(t)D(e);else{var n=e.data(M)||{};n.disabled=!0,e.data(M,n)}break;case 1:N.disabled=!t;break;default:t=!N.disabled}return!!t}}}]),n.register("",["$window","$sniffer","$timeout","$$animateReflow",function(h,t,r,n){function a(){g||(g=n(function(){L=[],g=null,I={}}))}function s(t,e){g&&g(),L.push(e),g=n(function(){z(L,function(t){t()}),L=[],g=null,I={}})}function k(t,e){var n=V(t);t=X.element(n),F.push(t);var i=Date.now()+e;i<=R||(r.cancel(j),R=i,j=r(function(){(function(t){z(t,function(t){var e=t.data(P);e&&(e.closeAnimationFn||q)()})})(F),F=[]},e,!1))}function v(t,e){var n=e?I[e]:null;if(!n){var i,r,o,a,s=0,l=0,u=0,c=0;z(t,function(t){if(t.nodeType==H){var e=h.getComputedStyle(t)||{};o=e[m+b],s=Math.max(d(o),s),a=e[m+x],i=e[m+w],l=Math.max(d(i),l),r=e[y+w],c=Math.max(d(r),c);var n=d(e[y+b]);0<n&&(n*=parseInt(e[y+$],10)||1),u=Math.max(n,u)}}),n={total:0,transitionPropertyStyle:a,transitionDurationStyle:o,transitionDelayStyle:i,transitionDelay:l,transitionDuration:s,animationDelayStyle:r,animationDelay:c,animationDuration:u},e&&(I[e]=n)}return n}function d(t){var e=0,n=X.isString(t)?t.split(/\s*,\s*/):[];return z(n,function(t){e=Math.max(parseFloat(t)||0,e)}),e}function o(t,e,n,i){var r=function(t){var e=t.parent(),n=e.data(_);return n||(e.data(_,++B),n=B),n+"-"+V(t).getAttribute("class")}(e),o=r+" "+n,a=I[o]?++I[o].total:0,s={};if(0<a){var l=n+"-stagger",u=r+" "+l,c=!I[u];c&&e.addClass(l),s=v(e,u),c&&e.removeClass(l)}i=i||function(t){return t()},e.addClass(n);var h=e.data(P)||{},d=i(function(){return v(e,o)}),p=d.transitionDuration,f=d.animationDuration;if(0===p&&0===f)return e.removeClass(n),!1;e.data(P,{running:h.running||0,itemIndex:a,stagger:s,timings:d,closeAnimationFn:q});var g=0<h.running||"setClass"==t;return 0<p&&function(t,e,n){!function(t){return"ng-enter"==t||"ng-move"==t||"ng-leave"==t}(e)&&n?t.addClass(M):V(t).style[m+x]="none"}(e,n,g),0<f&&0<s.animationDelay&&0===s.animationDuration&&function(t){V(t).style[y]="none 0s"}(e),!0}function l(t){var e=m+x,n=V(t);n.style[e]&&0<n.style[e].length&&(n.style[e]=""),t.removeClass(M)}function u(t){var e=y,n=V(t);n.style[e]&&0<n.style[e].length&&(n.style[e]="")}function C(t,n,i){var r="";return z(t.split(","),function(t,e){r+=(0<e?",":"")+(i*n+parseInt(t,10))+"s"}),r}function c(t,e,n,i){return o(t,e,n,i)?function(t){t&&E(e,n)}:void 0}function p(t,e,n,i){return e.data(P)?function(t,i,r,o){function e(t){i.off(v,a),i.removeClass(l),E(i,r);var e=V(i);for(var n in y)e.style.removeProperty(y[n])}function a(t){t.stopPropagation();var e=t.originalEvent||t,n=e.$manualTimeStamp||e.timeStamp||Date.now(),i=parseFloat(e.elapsedTime.toFixed(A));Math.max(n-g,0)>=f&&d<=i&&o()}var n=V(i),s=i.data(P);if(-1!=n.getAttribute("class").indexOf(r)&&s){var l="";z(r.split(" "),function(t,e){l+=(0<e?" ":"")+t+"-active"});var u=s.stagger,c=s.timings,h=s.itemIndex,d=Math.max(c.transitionDuration,c.animationDuration),p=Math.max(c.transitionDelay,c.animationDelay),f=p*O,g=Date.now(),v=T+" "+S,m="",y=[];if(0<c.transitionDuration){var b=c.transitionPropertyStyle;-1==b.indexOf("all")&&(m+=D+"transition-property: "+b+";",m+=D+"transition-duration: "+c.transitionDurationStyle+";",y.push(D+"transition-property"),y.push(D+"transition-duration"))}if(0<h){if(0<u.transitionDelay&&0===u.transitionDuration){var x=c.transitionDelayStyle;m+=D+"transition-delay: "+C(x,u.transitionDelay,h)+"; ",y.push(D+"transition-delay")}0<u.animationDelay&&0===u.animationDuration&&(m+=D+"animation-delay: "+C(c.animationDelayStyle,u.animationDelay,h)+"; ",y.push(D+"animation-delay"))}if(0<y.length){var w=n.getAttribute("style")||"";n.setAttribute("style",w+"; "+m)}i.on(v,a),i.addClass(l),s.closeAnimationFn=function(){e(),o()};var $=(h*(Math.max(u.animationDelay,u.transitionDelay)||0)+(p+d)*N)*O;return s.running++,k(i,$),e}o()}(0,e,n,i):(E(e,n),void i())}function i(t,e,n,i){var r=c(t,e,n);if(!r)return a(),void i();var o=r;return s(0,function(){l(e),u(e),o=p(0,e,n,i)}),function(t){(o||q)(t)}}function E(t,e){t.removeClass(e);var n=t.data(P);n&&(n.running&&n.running--,n.running&&0!==n.running||t.removeData(P))}function f(t,n){var i="";return t=X.isArray(t)?t:t.split(/\s+/),z(t,function(t,e){t&&0<t.length&&(i+=(0<e?" ":"")+t+n)}),i}var m,S,y,T,D="";S=e.ontransitionend===Y&&e.onwebkittransitionend!==Y?(D="-webkit-",m="WebkitTransition","webkitTransitionEnd transitionend"):(m="transition","transitionend"),T=e.onanimationend===Y&&e.onwebkitanimationend!==Y?(D="-webkit-",y="WebkitAnimation","webkitAnimationEnd animationend"):(y="animation","animationend");var g,b="Duration",x="Property",w="Delay",$="IterationCount",_="$$ngAnimateKey",P="$$ngAnimateCSS3Data",M="ng-animate-block-transitions",A=3,N=1.5,O=1e3,I={},B=0,L=[],j=null,R=0,F=[];return{enter:function(t,e){return i("enter",t,"ng-enter",e)},leave:function(t,e){return i("leave",t,"ng-leave",e)},move:function(t,e){return i("move",t,"ng-move",e)},beforeSetClass:function(i,r,o,t){var e=f(o,"-remove")+" "+f(r,"-add"),n=c("setClass",i,e,function(t){var e=i.attr("class");i.removeClass(o),i.addClass(r);var n=t();return i.attr("class",e),n});return n?(s(0,function(){l(i),u(i),t()}),n):(a(),void t())},beforeAddClass:function(n,i,t){var e=c("addClass",n,f(i,"-add"),function(t){n.addClass(i);var e=t();return n.removeClass(i),e});return e?(s(0,function(){l(n),u(n),t()}),e):(a(),void t())},setClass:function(t,e,n,i){return p(0,t,(n=f(n,"-remove"))+" "+(e=f(e,"-add")),i)},addClass:function(t,e,n){return p(0,t,f(e,"-add"),n)},beforeRemoveClass:function(i,r,t){var e=c("removeClass",i,f(r,"-remove"),function(t){var e=i.attr("class");i.removeClass(r);var n=t();return i.attr("class",e),n});return e?(s(0,function(){l(i),u(i),t()}),e):void t()},removeClass:function(t,e,n){return p(0,t,f(e,"-remove"),n)}}}])}])}(window,window.angular),angular.module("ui.bootstrap",["ui.bootstrap.tpls","ui.bootstrap.transition","ui.bootstrap.collapse","ui.bootstrap.accordion","ui.bootstrap.alert","ui.bootstrap.bindHtml","ui.bootstrap.buttons","ui.bootstrap.carousel","ui.bootstrap.dateparser","ui.bootstrap.position","ui.bootstrap.datepicker","ui.bootstrap.dropdown","ui.bootstrap.modal","ui.bootstrap.pagination","ui.bootstrap.tooltip","ui.bootstrap.popover","ui.bootstrap.progressbar","ui.bootstrap.rating","ui.bootstrap.tabs","ui.bootstrap.timepicker","ui.bootstrap.typeahead"]),angular.module("ui.bootstrap.tpls",["template/accordion/accordion-group.html","template/accordion/accordion.html","template/alert/alert.html","template/carousel/carousel.html","template/carousel/slide.html","template/datepicker/datepicker.html","template/datepicker/day.html","template/datepicker/month.html","template/datepicker/popup.html","template/datepicker/year.html","template/modal/backdrop.html","template/modal/window.html","template/pagination/pager.html","template/pagination/pagination.html","template/tooltip/tooltip-html-unsafe-popup.html","template/tooltip/tooltip-popup.html","template/popover/popover.html","template/progressbar/bar.html","template/progressbar/progress.html","template/progressbar/progressbar.html","template/rating/rating.html","template/tabs/tab.html","template/tabs/tabset.html","template/timepicker/timepicker.html","template/typeahead/typeahead-match.html","template/typeahead/typeahead-popup.html"]),angular.module("ui.bootstrap.transition",[]).factory("$transition",["$q","$timeout","$rootScope",function(a,s,l){function t(t){for(var e in t)if(void 0!==n.style[e])return t[e]}var u=function(e,t,n){n=n||{};var i=a.defer(),r=u[n.animation?"animationEndEventName":"transitionEndEventName"],o=function(t){l.$apply(function(){e.unbind(r,o),i.resolve(e)})};return r&&e.bind(r,o),s(function(){angular.isString(t)?e.addClass(t):angular.isFunction(t)?t(e):angular.isObject(t)&&e.css(t),r||i.resolve(e)}),i.promise.cancel=function(){r&&e.unbind(r,o),i.reject("Transition cancelled")},i.promise},n=document.createElement("trans");return u.transitionEndEventName=t({WebkitTransition:"webkitTransitionEnd",MozTransition:"transitionend",OTransition:"oTransitionEnd",transition:"transitionend"}),u.animationEndEventName=t({WebkitTransition:"webkitAnimationEnd",MozTransition:"animationend",OTransition:"oAnimationEnd",transition:"animationend"}),u}]),angular.module("ui.bootstrap.collapse",["ui.bootstrap.transition"]).directive("collapse",["$transition",function(l){return{link:function(t,i,e){function n(t){function e(){a===n&&(a=void 0)}var n=l(i,t);return a&&a.cancel(),(a=n).then(e,e),n}function r(){i.removeClass("collapsing"),i.addClass("collapse in"),i.css({height:"auto"})}function o(){i.removeClass("collapsing"),i.addClass("collapse")}var a,s=!0;t.$watch(e.collapse,function(t){t?s?(s=!1,o(),i.css({height:0})):(i.css({height:i[0].scrollHeight+"px"}),i[0].offsetWidth,i.removeClass("collapse in").addClass("collapsing"),n({height:0}).then(o)):s?(s=!1,r()):(i.removeClass("collapse").addClass("collapsing"),n({height:i[0].scrollHeight+"px"}).then(r))})}}}]),angular.module("ui.bootstrap.accordion",["ui.bootstrap.collapse"]).constant("accordionConfig",{closeOthers:!0}).controller("AccordionController",["$scope","$attrs","accordionConfig",function(t,n,i){this.groups=[],this.closeOthers=function(e){(angular.isDefined(n.closeOthers)?t.$eval(n.closeOthers):i.closeOthers)&&angular.forEach(this.groups,function(t){t!==e&&(t.isOpen=!1)})},this.addGroup=function(e){var n=this;this.groups.push(e),e.$on("$destroy",function(t){n.removeGroup(e)})},this.removeGroup=function(t){var e=this.groups.indexOf(t);-1!==e&&this.groups.splice(e,1)}}]).directive("accordion",function(){return{restrict:"EA",controller:"AccordionController",transclude:!0,replace:!1,templateUrl:"template/accordion/accordion.html"}}).directive("accordionGroup",function(){return{require:"^accordion",restrict:"EA",transclude:!0,replace:!0,templateUrl:"template/accordion/accordion-group.html",scope:{heading:"@",isOpen:"=?",isDisabled:"=?"},controller:function(){this.setHeading=function(t){this.heading=t}},link:function(e,t,n,i){i.addGroup(e),e.$watch("isOpen",function(t){t&&i.closeOthers(e)}),e.toggleOpen=function(){e.isDisabled||(e.isOpen=!e.isOpen)}}}}).directive("accordionHeading",function(){return{restrict:"EA",transclude:!0,template:"",replace:!0,require:"^accordionGroup",link:function(t,e,n,i,r){i.setHeading(r(t,function(){}))}}}).directive("accordionTransclude",function(){return{require:"^accordionGroup",link:function(t,e,n,i){t.$watch(function(){return i[n.accordionTransclude]},function(t){t&&(e.html(""),e.append(t))})}}}),angular.module("ui.bootstrap.alert",[]).controller("AlertController",["$scope","$attrs",function(t,e){t.closeable="close"in e,this.close=t.close}]).directive("alert",function(){return{restrict:"EA",controller:"AlertController",templateUrl:"template/alert/alert.html",transclude:!0,replace:!0,scope:{type:"@",close:"&"}}}).directive("dismissOnTimeout",["$timeout",function(r){return{require:"alert",link:function(t,e,n,i){r(function(){i.close()},parseInt(n.dismissOnTimeout,10))}}}]),angular.module("ui.bootstrap.bindHtml",[]).directive("bindHtmlUnsafe",function(){return function(t,e,n){e.addClass("ng-binding").data("$binding",n.bindHtmlUnsafe),t.$watch(n.bindHtmlUnsafe,function(t){e.html(t||"")})}}),angular.module("ui.bootstrap.buttons",[]).constant("buttonConfig",{activeClass:"active",toggleEvent:"click"}).controller("ButtonsController",["buttonConfig",function(t){this.activeClass=t.activeClass||"active",this.toggleEvent=t.toggleEvent||"click"}]).directive("btnRadio",function(){return{require:["btnRadio","ngModel"],controller:"ButtonsController",link:function(e,n,i,t){var r=t[0],o=t[1];o.$render=function(){n.toggleClass(r.activeClass,angular.equals(o.$modelValue,e.$eval(i.btnRadio)))},n.bind(r.toggleEvent,function(){var t=n.hasClass(r.activeClass);t&&!angular.isDefined(i.uncheckable)||e.$apply(function(){o.$setViewValue(t?null:e.$eval(i.btnRadio)),o.$render()})})}}}).directive("btnCheckbox",function(){return{require:["btnCheckbox","ngModel"],controller:"ButtonsController",link:function(i,t,e,n){function r(){return o(e.btnCheckboxTrue,!0)}function o(t,e){var n=i.$eval(t);return angular.isDefined(n)?n:e}var a=n[0],s=n[1];s.$render=function(){t.toggleClass(a.activeClass,angular.equals(s.$modelValue,r()))},t.bind(a.toggleEvent,function(){i.$apply(function(){s.$setViewValue(t.hasClass(a.activeClass)?o(e.btnCheckboxFalse,!1):r()),s.$render()})})}}}),angular.module("ui.bootstrap.carousel",["ui.bootstrap.transition"]).controller("CarouselController",["$scope","$timeout","$interval","$transition",function(a,e,n,s){function l(){i();var t=+a.interval;!isNaN(t)&&0<t&&(o=n(r,t))}function i(){o&&(n.cancel(o),o=null)}function r(){var t=+a.interval;u&&!isNaN(t)&&0<t?a.next():a.pause()}var o,u,c=this,h=c.slides=a.slides=[],d=-1;c.currentSlide=null;var p=!1;c.select=a.select=function(n,i){function t(){var t,e;p||(c.currentSlide&&angular.isString(i)&&!a.noTransition&&n.$element?(n.$element.addClass(i),n.$element[0].offsetWidth,angular.forEach(h,function(t){angular.extend(t,{direction:"",entering:!1,leaving:!1,active:!1})}),angular.extend(n,{direction:i,active:!0,entering:!0}),angular.extend(c.currentSlide||{},{direction:i,leaving:!0}),a.$currentTransition=s(n.$element,{}),t=n,e=c.currentSlide,a.$currentTransition.then(function(){r(t,e)},function(){r(t,e)})):r(n,c.currentSlide),c.currentSlide=n,d=o,l())}function r(t,e){angular.extend(t,{direction:"",active:!0,leaving:!1,entering:!1}),angular.extend(e||{},{direction:"",active:!1,leaving:!1,entering:!1}),a.$currentTransition=null}var o=h.indexOf(n);void 0===i&&(i=d<o?"next":"prev"),n&&n!==c.currentSlide&&(a.$currentTransition?(a.$currentTransition.cancel(),e(t)):t())},a.$on("$destroy",function(){p=!0}),c.indexOfSlide=function(t){return h.indexOf(t)},a.next=function(){var t=(d+1)%h.length;return a.$currentTransition?void 0:c.select(h[t],"next")},a.prev=function(){var t=d-1<0?h.length-1:d-1;return a.$currentTransition?void 0:c.select(h[t],"prev")},a.isActive=function(t){return c.currentSlide===t},a.$watch("interval",l),a.$on("$destroy",i),a.play=function(){u||(u=!0,l())},a.pause=function(){a.noPause||(u=!1,i())},c.addSlide=function(t,e){t.$element=e,h.push(t),1===h.length||t.active?(c.select(h[h.length-1]),1==h.length&&a.play()):t.active=!1},c.removeSlide=function(t){var e=h.indexOf(t);h.splice(e,1),0<h.length&&t.active?e>=h.length?c.select(h[e-1]):c.select(h[e]):e<d&&d--}}]).directive("carousel",[function(){return{restrict:"EA",transclude:!0,replace:!0,controller:"CarouselController",require:"carousel",templateUrl:"template/carousel/carousel.html",scope:{interval:"=",noTransition:"=",noPause:"="}}}]).directive("slide",function(){return{require:"^carousel",restrict:"EA",transclude:!0,replace:!0,templateUrl:"template/carousel/slide.html",scope:{active:"=?"},link:function(e,t,n,i){i.addSlide(e,t),e.$on("$destroy",function(){i.removeSlide(e)}),e.$watch("active",function(t){t&&i.select(e)})}}}),angular.module("ui.bootstrap.dateparser",[]).service("dateParser",["$locale","orderByFilter",function(h,d){this.parsers={};var p={yyyy:{regex:"\\d{4}",apply:function(t){this.year=+t}},yy:{regex:"\\d{2}",apply:function(t){this.year=+t+2e3}},y:{regex:"\\d{1,4}",apply:function(t){this.year=+t}},MMMM:{regex:h.DATETIME_FORMATS.MONTH.join("|"),apply:function(t){this.month=h.DATETIME_FORMATS.MONTH.indexOf(t)}},MMM:{regex:h.DATETIME_FORMATS.SHORTMONTH.join("|"),apply:function(t){this.month=h.DATETIME_FORMATS.SHORTMONTH.indexOf(t)}},MM:{regex:"0[1-9]|1[0-2]",apply:function(t){this.month=t-1}},M:{regex:"[1-9]|1[0-2]",apply:function(t){this.month=t-1}},dd:{regex:"[0-2][0-9]{1}|3[0-1]{1}",apply:function(t){this.date=+t}},d:{regex:"[1-2]?[0-9]{1}|3[0-1]{1}",apply:function(t){this.date=+t}},EEEE:{regex:h.DATETIME_FORMATS.DAY.join("|")},EEE:{regex:h.DATETIME_FORMATS.SHORTDAY.join("|")}};this.parse=function(t,e){if(!angular.isString(t)||!e)return t;e=h.DATETIME_FORMATS[e]||e,this.parsers[e]||(this.parsers[e]=function(o){var a=[],s=o.split("");return angular.forEach(p,function(t,e){var n=o.indexOf(e);if(-1<n){o=o.split(""),s[n]="("+t.regex+")",o[n]="$";for(var i=n+1,r=n+e.length;i<r;i++)s[i]="",o[i]="$";o=o.join(""),a.push({index:n,apply:t.apply})}}),{regex:new RegExp("^"+s.join("")+"$"),map:d(a,"index")}}(e));var n=this.parsers[e],i=n.regex,r=n.map,o=t.match(i);if(o&&o.length){for(var a,s={year:1900,month:0,date:1,hours:0},l=1,u=o.length;l<u;l++){var c=r[l-1];c.apply&&c.apply.call(s,o[l])}return function(t,e,n){return 1===e&&28<n?29===n&&(t%4==0&&t%100!=0||t%400==0):3!==e&&5!==e&&8!==e&&10!==e||n<31}(s.year,s.month,s.date)&&(a=new Date(s.year,s.month,s.date,s.hours)),a}}}]),angular.module("ui.bootstrap.position",[]).factory("$position",["$document","$window",function(o,n){function a(t){return"static"===(function(t,e){return t.currentStyle?t.currentStyle[e]:n.getComputedStyle?n.getComputedStyle(t)[e]:t.style[e]}(t,"position")||"static")}return{position:function(t){var e=this.offset(t),n={top:0,left:0},i=function(t){for(var e=o[0],n=t.offsetParent||e;n&&n!==e&&a(n);)n=n.offsetParent;return n||e}(t[0]);i!=o[0]&&((n=this.offset(angular.element(i))).top+=i.clientTop-i.scrollTop,n.left+=i.clientLeft-i.scrollLeft);var r=t[0].getBoundingClientRect();return{width:r.width||t.prop("offsetWidth"),height:r.height||t.prop("offsetHeight"),top:e.top-n.top,left:e.left-n.left}},offset:function(t){var e=t[0].getBoundingClientRect();return{width:e.width||t.prop("offsetWidth"),height:e.height||t.prop("offsetHeight"),top:e.top+(n.pageYOffset||o[0].documentElement.scrollTop),left:e.left+(n.pageXOffset||o[0].documentElement.scrollLeft)}},positionElements:function(t,e,n,i){var r,o,a,s,l=n.split("-"),u=l[0],c=l[1]||"center";r=i?this.offset(t):this.position(t),o=e.prop("offsetWidth"),a=e.prop("offsetHeight");var h={center:function(){return r.left+r.width/2-o/2},left:function(){return r.left},right:function(){return r.left+r.width}},d={center:function(){return r.top+r.height/2-a/2},top:function(){return r.top},bottom:function(){return r.top+r.height}};switch(u){case"right":s={top:d[c](),left:h[u]()};break;case"left":s={top:d[c](),left:r.left-o};break;case"bottom":s={top:d[u](),left:h[c]()};break;default:s={top:r.top-a,left:h[c]()}}return s}}}]),angular.module("ui.bootstrap.datepicker",["ui.bootstrap.dateparser","ui.bootstrap.position"]).constant("datepickerConfig",{formatDay:"dd",formatMonth:"MMMM",formatYear:"yyyy",formatDayHeader:"EEE",formatDayTitle:"MMMM yyyy",formatMonthTitle:"yyyy",datepickerMode:"day",minMode:"day",maxMode:"year",showWeeks:!0,startingDay:0,yearRange:20,minDate:null,maxDate:null}).controller("DatepickerController",["$scope","$attrs","$parse","$interpolate","$timeout","$log","dateFilter","datepickerConfig",function(n,i,t,r,e,o,a,s){var l=this,u={$setViewValue:angular.noop};this.modes=["day","month","year"],angular.forEach(["formatDay","formatMonth","formatYear","formatDayHeader","formatDayTitle","formatMonthTitle","minMode","maxMode","showWeeks","startingDay","yearRange"],function(t,e){l[t]=angular.isDefined(i[t])?e<8?r(i[t])(n.$parent):n.$parent.$eval(i[t]):s[t]}),angular.forEach(["minDate","maxDate"],function(e){i[e]?n.$parent.$watch(t(i[e]),function(t){l[e]=t?new Date(t):null,l.refreshView()}):l[e]=s[e]?new Date(s[e]):null}),n.datepickerMode=n.datepickerMode||s.datepickerMode,n.uniqueId="datepicker-"+n.$id+"-"+Math.floor(1e4*Math.random()),this.activeDate=angular.isDefined(i.initDate)?n.$parent.$eval(i.initDate):new Date,n.isActive=function(t){return 0===l.compare(t.date,l.activeDate)&&(n.activeDateId=t.uid,!0)},this.init=function(t){(u=t).$render=function(){l.render()}},this.render=function(){if(u.$modelValue){var t=new Date(u.$modelValue),e=!isNaN(t);e?this.activeDate=t:o.error('Datepicker directive: "ng-model" value must be a Date object, a number of milliseconds since 01.01.1970 or a string representing an RFC2822 or ISO 8601 date.'),u.$setValidity("date",e)}this.refreshView()},this.refreshView=function(){if(this.element){this._refreshView();var t=u.$modelValue?new Date(u.$modelValue):null;u.$setValidity("date-disabled",!t||this.element&&!this.isDisabled(t))}},this.createDateObject=function(t,e){var n=u.$modelValue?new Date(u.$modelValue):null;return{date:t,label:a(t,e),selected:n&&0===this.compare(t,n),disabled:this.isDisabled(t),current:0===this.compare(t,new Date)}},this.isDisabled=function(t){return this.minDate&&this.compare(t,this.minDate)<0||this.maxDate&&0<this.compare(t,this.maxDate)||i.dateDisabled&&n.dateDisabled({date:t,mode:n.datepickerMode})},this.split=function(t,e){for(var n=[];0<t.length;)n.push(t.splice(0,e));return n},n.select=function(t){if(n.datepickerMode===l.minMode){var e=u.$modelValue?new Date(u.$modelValue):new Date(0,0,0,0,0,0,0);e.setFullYear(t.getFullYear(),t.getMonth(),t.getDate()),u.$setViewValue(e),u.$render()}else l.activeDate=t,n.datepickerMode=l.modes[l.modes.indexOf(n.datepickerMode)-1]},n.move=function(t){var e=l.activeDate.getFullYear()+t*(l.step.years||0),n=l.activeDate.getMonth()+t*(l.step.months||0);l.activeDate.setFullYear(e,n,1),l.refreshView()},n.toggleMode=function(t){t=t||1,n.datepickerMode===l.maxMode&&1===t||n.datepickerMode===l.minMode&&-1===t||(n.datepickerMode=l.modes[l.modes.indexOf(n.datepickerMode)+t])},n.keys={13:"enter",32:"space",33:"pageup",34:"pagedown",35:"end",36:"home",37:"left",38:"up",39:"right",40:"down"};function c(){e(function(){l.element[0].focus()},0,!1)}n.$on("datepicker.focus",c),n.keydown=function(t){var e=n.keys[t.which];if(e&&!t.shiftKey&&!t.altKey)if(t.preventDefault(),t.stopPropagation(),"enter"===e||"space"===e){if(l.isDisabled(l.activeDate))return;n.select(l.activeDate),c()}else!t.ctrlKey||"up"!==e&&"down"!==e?(l.handleKeyDown(e,t),l.refreshView()):(n.toggleMode("up"===e?1:-1),c())}}]).directive("datepicker",function(){return{restrict:"EA",replace:!0,templateUrl:"template/datepicker/datepicker.html",scope:{datepickerMode:"=?",dateDisabled:"&"},require:["datepicker","?^ngModel"],controller:"DatepickerController",link:function(t,e,n,i){var r=i[0],o=i[1];o&&r.init(o)}}}).directive("daypicker",["dateFilter",function(p){return{restrict:"EA",replace:!0,templateUrl:"template/datepicker/day.html",require:"^datepicker",link:function(h,t,e,d){function r(t,e){return 1!==e||t%4!=0||t%100==0&&t%400!=0?n[e]:29}h.showWeeks=d.showWeeks,d.step={months:1},d.element=t;var n=[31,28,31,30,31,30,31,31,30,31,30,31];d._refreshView=function(){var t=d.activeDate.getFullYear(),e=d.activeDate.getMonth(),n=new Date(t,e,1),i=d.startingDay-n.getDay(),r=0<i?7-i:-i,o=new Date(n);0<r&&o.setDate(1-r);for(var a=function(t,e){var n=new Array(e),i=new Date(t),r=0;for(i.setHours(12);r<e;)n[r++]=new Date(i),i.setDate(i.getDate()+1);return n}(o,42),s=0;s<42;s++)a[s]=angular.extend(d.createDateObject(a[s],d.formatDay),{secondary:a[s].getMonth()!==e,uid:h.uniqueId+"-"+s});h.labels=new Array(7);for(var l=0;l<7;l++)h.labels[l]={abbr:p(a[l].date,d.formatDayHeader),full:p(a[l].date,"EEEE")};if(h.title=p(d.activeDate,d.formatDayTitle),h.rows=d.split(a,7),h.showWeeks){h.weekNumbers=[];for(var u=function(t){var e=new Date(t);e.setDate(e.getDate()+4-(e.getDay()||7));var n=e.getTime();return e.setMonth(0),e.setDate(1),Math.floor(Math.round((n-e)/864e5)/7)+1}(h.rows[0][0].date),c=h.rows.length;h.weekNumbers.push(u++)<c;);}},d.compare=function(t,e){return new Date(t.getFullYear(),t.getMonth(),t.getDate())-new Date(e.getFullYear(),e.getMonth(),e.getDate())},d.handleKeyDown=function(t,e){var n=d.activeDate.getDate();if("left"===t)n-=1;else if("up"===t)n-=7;else if("right"===t)n+=1;else if("down"===t)n+=7;else if("pageup"===t||"pagedown"===t){var i=d.activeDate.getMonth()+("pageup"===t?-1:1);d.activeDate.setMonth(i,1),n=Math.min(r(d.activeDate.getFullYear(),d.activeDate.getMonth()),n)}else"home"===t?n=1:"end"===t&&(n=r(d.activeDate.getFullYear(),d.activeDate.getMonth()));d.activeDate.setDate(n)},d.refreshView()}}}]).directive("monthpicker",["dateFilter",function(o){return{restrict:"EA",replace:!0,templateUrl:"template/datepicker/month.html",require:"^datepicker",link:function(i,t,e,r){r.step={years:1},r.element=t,r._refreshView=function(){for(var t=new Array(12),e=r.activeDate.getFullYear(),n=0;n<12;n++)t[n]=angular.extend(r.createDateObject(new Date(e,n,1),r.formatMonth),{uid:i.uniqueId+"-"+n});i.title=o(r.activeDate,r.formatMonthTitle),i.rows=r.split(t,3)},r.compare=function(t,e){return new Date(t.getFullYear(),t.getMonth())-new Date(e.getFullYear(),e.getMonth())},r.handleKeyDown=function(t,e){var n=r.activeDate.getMonth();if("left"===t)n-=1;else if("up"===t)n-=3;else if("right"===t)n+=1;else if("down"===t)n+=3;else if("pageup"===t||"pagedown"===t){var i=r.activeDate.getFullYear()+("pageup"===t?-1:1);r.activeDate.setFullYear(i)}else"home"===t?n=0:"end"===t&&(n=11);r.activeDate.setMonth(n)},r.refreshView()}}}]).directive("yearpicker",["dateFilter",function(t){return{restrict:"EA",replace:!0,templateUrl:"template/datepicker/year.html",require:"^datepicker",link:function(i,t,e,r){function o(t){return parseInt((t-1)/a,10)*a+1}var a=r.yearRange;r.step={years:a},r.element=t,r._refreshView=function(){for(var t=new Array(a),e=0,n=o(r.activeDate.getFullYear());e<a;e++)t[e]=angular.extend(r.createDateObject(new Date(n+e,0,1),r.formatYear),{uid:i.uniqueId+"-"+e});i.title=[t[0].label,t[a-1].label].join(" - "),i.rows=r.split(t,5)},r.compare=function(t,e){return t.getFullYear()-e.getFullYear()},r.handleKeyDown=function(t,e){var n=r.activeDate.getFullYear();"left"===t?n-=1:"up"===t?n-=5:"right"===t?n+=1:"down"===t?n+=5:"pageup"===t||"pagedown"===t?n+=("pageup"===t?-1:1)*r.step.years:"home"===t?n=o(r.activeDate.getFullYear()):"end"===t&&(n=o(r.activeDate.getFullYear())+a-1),r.activeDate.setFullYear(n)},r.refreshView()}}}]).constant("datepickerPopupConfig",{datepickerPopup:"yyyy-MM-dd",currentText:"Today",clearText:"Clear",closeText:"Done",closeOnDateSelection:!0,appendToBody:!1,showButtonBar:!0}).directive("datepickerPopup",["$compile","$parse","$document","$position","dateFilter","dateParser","datepickerPopupConfig",function(f,g,v,m,y,b,x){return{restrict:"EA",require:"ngModel",scope:{isOpen:"=?",currentText:"@",clearText:"@",closeText:"@",dateDisabled:"&"},link:function(i,e,r,n){function o(t){return t.replace(/([A-Z])/g,function(t){return"-"+t.toLowerCase()})}function a(t){if(!t)return n.$setValidity("date",!0),null;if(angular.isDate(t)&&!isNaN(t))return n.$setValidity("date",!0),t;if(angular.isString(t)){var e=b.parse(t,s)||new Date(t);return isNaN(e)?void n.$setValidity("date",!1):(n.$setValidity("date",!0),e)}n.$setValidity("date",!1)}var s,l=angular.isDefined(r.closeOnDateSelection)?i.$parent.$eval(r.closeOnDateSelection):x.closeOnDateSelection,u=angular.isDefined(r.datepickerAppendToBody)?i.$parent.$eval(r.datepickerAppendToBody):x.appendToBody;i.showButtonBar=angular.isDefined(r.showButtonBar)?i.$parent.$eval(r.showButtonBar):x.showButtonBar,i.getText=function(t){return i[t+"Text"]||x[t+"Text"]},r.$observe("datepickerPopup",function(t){s=t||x.datepickerPopup,n.$render()});var t=angular.element("<div datepicker-popup-wrap><div datepicker></div></div>");t.attr({"ng-model":"date","ng-change":"dateSelection()"});var c=angular.element(t.children()[0]);r.datepickerOptions&&angular.forEach(i.$parent.$eval(r.datepickerOptions),function(t,e){c.attr(o(e),t)}),i.watchData={},angular.forEach(["minDate","maxDate","datepickerMode"],function(e){if(r[e]){var t=g(r[e]);if(i.$parent.$watch(t,function(t){i.watchData[e]=t}),c.attr(o(e),"watchData."+e),"datepickerMode"===e){var n=t.assign;i.$watch("watchData."+e,function(t,e){t!==e&&n(i.$parent,t)})}}}),r.dateDisabled&&c.attr("date-disabled","dateDisabled({ date: date, mode: mode })"),n.$parsers.unshift(a),i.dateSelection=function(t){angular.isDefined(t)&&(i.date=t),n.$setViewValue(i.date),n.$render(),l&&(i.isOpen=!1,e[0].focus())},e.bind("input change keyup",function(){i.$apply(function(){i.date=n.$modelValue})}),n.$render=function(){var t=n.$viewValue?y(n.$viewValue,s):"";e.val(t),i.date=a(n.$modelValue)};function h(t){i.isOpen&&t.target!==e[0]&&i.$apply(function(){i.isOpen=!1})}function d(t,e){i.keydown(t)}e.bind("keydown",d),i.keydown=function(t){27===t.which?(t.preventDefault(),t.stopPropagation(),i.close()):40!==t.which||i.isOpen||(i.isOpen=!0)},i.$watch("isOpen",function(t){t?(i.$broadcast("datepicker.focus"),i.position=u?m.offset(e):m.position(e),i.position.top=i.position.top+e.prop("offsetHeight"),v.bind("click",h)):v.unbind("click",h)}),i.select=function(t){if("today"===t){var e=new Date;angular.isDate(n.$modelValue)?(t=new Date(n.$modelValue)).setFullYear(e.getFullYear(),e.getMonth(),e.getDate()):t=new Date(e.setHours(0,0,0,0))}i.dateSelection(t)},i.close=function(){i.isOpen=!1,e[0].focus()};var p=f(t)(i);t.remove(),u?v.find("body").append(p):e.after(p),i.$on("$destroy",function(){p.remove(),e.unbind("keydown",d),v.unbind("click",h)})}}}]).directive("datepickerPopupWrap",function(){return{restrict:"EA",replace:!0,transclude:!0,templateUrl:"template/datepicker/popup.html",link:function(t,e,n){e.bind("click",function(t){t.preventDefault(),t.stopPropagation()})}}}),angular.module("ui.bootstrap.dropdown",[]).constant("dropdownConfig",{openClass:"open"}).service("dropdownService",["$document",function(e){var n=null;this.open=function(t){n||(e.bind("click",i),e.bind("keydown",r)),n&&n!==t&&(n.isOpen=!1),n=t},this.close=function(t){n===t&&(n=null,e.unbind("click",i),e.unbind("keydown",r))};var i=function(t){if(n){var e=n.getToggleElement();t&&e&&e[0].contains(t.target)||n.$apply(function(){n.isOpen=!1})}},r=function(t){27===t.which&&(n.focusToggleElement(),i())}}]).controller("DropdownController",["$scope","$attrs","$parse","dropdownConfig","dropdownService","$animate",function(n,e,i,t,r,o){var a,s=this,l=n.$new(),u=t.openClass,c=angular.noop,h=e.onToggle?i(e.onToggle):angular.noop;this.init=function(t){s.$element=t,e.isOpen&&(a=i(e.isOpen),c=a.assign,n.$watch(a,function(t){l.isOpen=!!t}))},this.toggle=function(t){return l.isOpen=arguments.length?!!t:!l.isOpen},this.isOpen=function(){return l.isOpen},l.getToggleElement=function(){return s.toggleElement},l.focusToggleElement=function(){s.toggleElement&&s.toggleElement[0].focus()},l.$watch("isOpen",function(t,e){o[t?"addClass":"removeClass"](s.$element,u),t?(l.focusToggleElement(),r.open(l)):r.close(l),c(n,t),angular.isDefined(t)&&t!==e&&h(n,{open:!!t})}),n.$on("$locationChangeSuccess",function(){l.isOpen=!1}),n.$on("$destroy",function(){l.$destroy()})}]).directive("dropdown",function(){return{controller:"DropdownController",link:function(t,e,n,i){i.init(e)}}}).directive("dropdownToggle",function(){return{require:"?^dropdown",link:function(e,n,i,r){if(r){r.toggleElement=n;function t(t){t.preventDefault(),n.hasClass("disabled")||i.disabled||e.$apply(function(){r.toggle()})}n.bind("click",t),n.attr({"aria-haspopup":!0,"aria-expanded":!1}),e.$watch(r.isOpen,function(t){n.attr("aria-expanded",!!t)}),e.$on("$destroy",function(){n.unbind("click",t)})}}}}),angular.module("ui.bootstrap.modal",["ui.bootstrap.transition"]).factory("$$stackedMap",function(){return{createNew:function(){var i=[];return{add:function(t,e){i.push({key:t,value:e})},get:function(t){for(var e=0;e<i.length;e++)if(t==i[e].key)return i[e]},keys:function(){for(var t=[],e=0;e<i.length;e++)t.push(i[e].key);return t},top:function(){return i[i.length-1]},remove:function(t){for(var e=-1,n=0;n<i.length;n++)if(t==i[n].key){e=n;break}return i.splice(e,1)[0]},removeTop:function(){return i.splice(i.length-1,1)[0]},length:function(){return i.length}}}}}).directive("modalBackdrop",["$timeout",function(i){return{restrict:"EA",replace:!0,templateUrl:"template/modal/backdrop.html",link:function(t,e,n){t.backdropClass=n.backdropClass||"",t.animate=!1,i(function(){t.animate=!0})}}}]).directive("modalWindow",["$modalStack","$timeout",function(i,r){return{restrict:"EA",scope:{index:"@",animate:"="},replace:!0,transclude:!0,templateUrl:function(t,e){return e.templateUrl||"template/modal/window.html"},link:function(t,e,n){e.addClass(n.windowClass||""),t.size=n.size,r(function(){t.animate=!0,e[0].querySelectorAll("[autofocus]").length||e[0].focus()}),t.close=function(t){var e=i.getTop();e&&e.value.backdrop&&"static"!=e.value.backdrop&&t.target===t.currentTarget&&(t.preventDefault(),t.stopPropagation(),i.dismiss(e.key,"backdrop click"))}}}}]).directive("modalTransclude",function(){return{link:function(t,e,n,i,r){r(t.$parent,function(t){e.empty(),e.append(t)})}}}).factory("$modalStack",["$transition","$timeout","$document","$compile","$rootScope","$$stackedMap",function(s,l,u,c,h,t){function d(){for(var t=-1,e=v.keys(),n=0;n<e.length;n++)v.get(e[n]).value.backdrop&&(t=n);return t}function i(t){var e=u.find("body").eq(0),n=v.get(t).value;v.remove(t),r(n.modalDomEl,n.modalScope,300,function(){n.modalScope.$destroy(),e.toggleClass(g,0<v.length()),function(){if(p&&-1==d()){var t=f;r(p,f,150,function(){t.$destroy(),t=null}),f=p=void 0}}()})}function r(t,e,n,i){function r(){r.done||(r.done=!0,t.remove(),i&&i())}e.animate=!1;var o=s.transitionEndEventName;if(o){var a=l(r,n);t.bind(o,function(){l.cancel(a),r(),e.$apply()})}else l(r)}var p,f,g="modal-open",v=t.createNew(),n={};return h.$watch(d,function(t){f&&(f.index=t)}),u.bind("keydown",function(t){var e;27!==t.which||(e=v.top())&&e.value.keyboard&&(t.preventDefault(),h.$apply(function(){n.dismiss(e.key,"escape key press")}))}),n.open=function(t,e){v.add(t,{deferred:e.deferred,modalScope:e.scope,backdrop:e.backdrop,keyboard:e.keyboard});var n=u.find("body").eq(0),i=d();if(0<=i&&!p){(f=h.$new(!0)).index=i;var r=angular.element("<div modal-backdrop></div>");r.attr("backdrop-class",e.backdropClass),p=c(r)(f),n.append(p)}var o=angular.element("<div modal-window></div>");o.attr({"template-url":e.windowTemplateUrl,"window-class":e.windowClass,size:e.size,index:v.length()-1,animate:"animate"}).html(e.content);var a=c(o)(e.scope);v.top().value.modalDomEl=a,n.append(a),n.addClass(g)},n.close=function(t,e){var n=v.get(t);n&&(n.value.deferred.resolve(e),i(t))},n.dismiss=function(t,e){var n=v.get(t);n&&(n.value.deferred.reject(e),i(t))},n.dismissAll=function(t){for(var e=this.getTop();e;)this.dismiss(e.key,t),e=this.getTop()},n.getTop=function(){return v.top()},n}]).provider("$modal",function(){var d={options:{backdrop:!0,keyboard:!0},$get:["$injector","$rootScope","$q","$http","$templateCache","$controller","$modalStack",function(n,l,i,r,u,c,h){var t={};return t.open=function(o){var a=i.defer(),t=i.defer(),s={result:a.promise,opened:t.promise,close:function(t){h.close(s,t)},dismiss:function(t){h.dismiss(s,t)}};if((o=angular.extend({},d.options,o)).resolve=o.resolve||{},!o.template&&!o.templateUrl)throw new Error("One of template or templateUrl options is required.");var e=i.all([function(t){return t.template?i.when(t.template):r.get(angular.isFunction(t.templateUrl)?t.templateUrl():t.templateUrl,{cache:u}).then(function(t){return t.data})}(o)].concat(function(t){var e=[];return angular.forEach(t,function(t){(angular.isFunction(t)||angular.isArray(t))&&e.push(i.when(n.invoke(t)))}),e}(o.resolve)));return e.then(function(n){var t=(o.scope||l).$new();t.$close=s.close,t.$dismiss=s.dismiss;var e,i={},r=1;o.controller&&(i.$scope=t,i.$modalInstance=s,angular.forEach(o.resolve,function(t,e){i[e]=n[r++]}),e=c(o.controller,i),o.controllerAs&&(t[o.controllerAs]=e)),h.open(s,{scope:t,deferred:a,content:n[0],backdrop:o.backdrop,keyboard:o.keyboard,backdropClass:o.backdropClass,windowClass:o.windowClass,windowTemplateUrl:o.windowTemplateUrl,size:o.size})},function(t){a.reject(t)}),e.then(function(){t.resolve(!0)},function(){t.reject(!1)}),s},t}]};return d}),angular.module("ui.bootstrap.pagination",[]).controller("PaginationController",["$scope","$attrs","$parse",function(n,i,r){var o=this,a={$setViewValue:angular.noop},e=i.numPages?r(i.numPages).assign:angular.noop;this.init=function(t,e){a=t,this.config=e,a.$render=function(){o.render()},i.itemsPerPage?n.$parent.$watch(r(i.itemsPerPage),function(t){o.itemsPerPage=parseInt(t,10),n.totalPages=o.calculateTotalPages()}):this.itemsPerPage=e.itemsPerPage},this.calculateTotalPages=function(){var t=this.itemsPerPage<1?1:Math.ceil(n.totalItems/this.itemsPerPage);return Math.max(t||0,1)},this.render=function(){n.page=parseInt(a.$viewValue,10)||1},n.selectPage=function(t){n.page!==t&&0<t&&t<=n.totalPages&&(a.$setViewValue(t),a.$render())},n.getText=function(t){return n[t+"Text"]||o.config[t+"Text"]},n.noPrevious=function(){return 1===n.page},n.noNext=function(){return n.page===n.totalPages},n.$watch("totalItems",function(){n.totalPages=o.calculateTotalPages()}),n.$watch("totalPages",function(t){e(n.$parent,t),n.page>t?n.selectPage(t):a.$render()})}]).constant("paginationConfig",{itemsPerPage:10,boundaryLinks:!1,directionLinks:!0,firstText:"First",previousText:"Previous",nextText:"Next",lastText:"Last",rotate:!0}).directive("pagination",["$parse","paginationConfig",function(s,l){return{restrict:"EA",scope:{totalItems:"=",firstText:"@",previousText:"@",nextText:"@",lastText:"@"},require:["pagination","?ngModel"],controller:"PaginationController",templateUrl:"template/pagination/pagination.html",replace:!0,link:function(t,e,n,i){function c(t,e,n){return{number:t,text:e,active:n}}var r=i[0],o=i[1];if(o){var h=angular.isDefined(n.maxSize)?t.$parent.$eval(n.maxSize):l.maxSize,d=angular.isDefined(n.rotate)?t.$parent.$eval(n.rotate):l.rotate;t.boundaryLinks=angular.isDefined(n.boundaryLinks)?t.$parent.$eval(n.boundaryLinks):l.boundaryLinks,t.directionLinks=angular.isDefined(n.directionLinks)?t.$parent.$eval(n.directionLinks):l.directionLinks,r.init(o,l),n.maxSize&&t.$parent.$watch(s(n.maxSize),function(t){h=parseInt(t,10),r.render()});var a=r.render;r.render=function(){a(),0<t.page&&t.page<=t.totalPages&&(t.pages=function(t,e){var n=[],i=1,r=e,o=angular.isDefined(h)&&h<e;o&&(d?e<(r=(i=Math.max(t-Math.floor(h/2),1))+h-1)&&(i=(r=e)-h+1):(i=(Math.ceil(t/h)-1)*h+1,r=Math.min(i+h-1,e)));for(var a=i;a<=r;a++){var s=c(a,a,a===t);n.push(s)}if(o&&!d){if(1<i){var l=c(i-1,"...",!1);n.unshift(l)}if(r<e){var u=c(r+1,"...",!1);n.push(u)}}return n}(t.page,t.totalPages))}}}}}]).constant("pagerConfig",{itemsPerPage:10,previousText:"« Previous",nextText:"Next »",align:!0}).directive("pager",["pagerConfig",function(a){return{restrict:"EA",scope:{totalItems:"=",previousText:"@",nextText:"@"},require:["pager","?ngModel"],controller:"PaginationController",templateUrl:"template/pagination/pager.html",replace:!0,link:function(t,e,n,i){var r=i[0],o=i[1];o&&(t.align=angular.isDefined(n.align)?t.$parent.$eval(n.align):a.align,r.init(o,a))}}}]),angular.module("ui.bootstrap.tooltip",["ui.bootstrap.position","ui.bootstrap.bindHtml"]).provider("$tooltip",function(){var s={placement:"top",animation:!0,popupDelay:0},l={mouseenter:"mouseleave",click:"click",focus:"blur"},u={};this.options=function(t){angular.extend(u,t)},this.setTriggers=function(t){angular.extend(l,t)},this.$get=["$window","$compile","$timeout","$document","$position","$interpolate",function(t,o,T,D,_,a){return function(k,C,n){function E(t){var e=t||S.trigger||n;return{show:e,hide:l[e]||e}}var S=angular.extend({},s,u),t=function(t){return t.replace(/[A-Z]/g,function(t,e){return(e?"-":"")+t.toLowerCase()})}(k),e=a.startSymbol(),i=a.endSymbol(),r="<div "+t+'-popup title="'+e+"title"+i+'" content="'+e+"content"+i+'" placement="'+e+"placement"+i+'" animation="animation" is-open="isOpen"></div>';return{restrict:"EA",compile:function(t,e){var $=o(r);return function(t,e,n){function i(){v.isOpen?o():r()}function r(){g&&!t.$eval(n[C+"Enable"])||(function(){var t=n[C+"Placement"];v.placement=angular.isDefined(t)?t:S.placement}(),function(){var t=n[C+"PopupDelay"],e=parseInt(t,10);v.popupDelay=isNaN(e)?S.popupDelay:e}(),v.popupDelay?d||(d=T(a,v.popupDelay,!1)).then(function(t){t()}):a()())}function o(){t.$apply(function(){s()})}function a(){return d=null,h&&(T.cancel(h),h=null),v.content?(u&&l(),c=v.$new(),(u=$(c,angular.noop)).css({top:0,left:0,display:"block"}),p?D.find("body").append(u):e.after(u),m(),v.isOpen=!0,v.$digest(),m):angular.noop}function s(){v.isOpen=!1,T.cancel(d),d=null,v.animation?h||(h=T(l,500)):l()}function l(){h=null,u&&(u.remove(),u=null),c&&(c.$destroy(),c=null)}var u,c,h,d,p=!!angular.isDefined(S.appendToBody)&&S.appendToBody,f=E(void 0),g=angular.isDefined(n[C+"Enable"]),v=t.$new(!0),m=function(){var t=_.positionElements(e,u,v.placement,p);t.top+="px",t.left+="px",u.css(t)};v.isOpen=!1,n.$observe(k,function(t){!(v.content=t)&&v.isOpen&&s()}),n.$observe(C+"Title",function(t){v.title=t});function y(){e.unbind(f.show,r),e.unbind(f.hide,o)}var b;b=n[C+"Trigger"],y(),(f=E(b)).show===f.hide?e.bind(f.show,i):(e.bind(f.show,r),e.bind(f.hide,o));var x=t.$eval(n[C+"Animation"]);v.animation=angular.isDefined(x)?!!x:S.animation;var w=t.$eval(n[C+"AppendToBody"]);(p=angular.isDefined(w)?w:p)&&t.$on("$locationChangeSuccess",function(){v.isOpen&&s()}),t.$on("$destroy",function(){T.cancel(h),T.cancel(d),y(),l(),v=null})}}}}}]}).directive("tooltipPopup",function(){return{restrict:"EA",replace:!0,scope:{content:"@",placement:"@",animation:"&",isOpen:"&"},templateUrl:"template/tooltip/tooltip-popup.html"}}).directive("tooltip",["$tooltip",function(t){return t("tooltip","tooltip","mouseenter")}]).directive("tooltipHtmlUnsafePopup",function(){return{restrict:"EA",replace:!0,scope:{content:"@",placement:"@",animation:"&",isOpen:"&"},templateUrl:"template/tooltip/tooltip-html-unsafe-popup.html"}}).directive("tooltipHtmlUnsafe",["$tooltip",function(t){return t("tooltipHtmlUnsafe","tooltip","mouseenter")}]),angular.module("ui.bootstrap.popover",["ui.bootstrap.tooltip"]).directive("popoverPopup",function(){return{restrict:"EA",replace:!0,scope:{title:"@",content:"@",placement:"@",animation:"&",isOpen:"&"},templateUrl:"template/popover/popover.html"}}).directive("popover",["$tooltip",function(t){return t("popover","popover","click")}]),angular.module("ui.bootstrap.progressbar",[]).constant("progressConfig",{animate:!0,max:100}).controller("ProgressController",["$scope","$attrs","progressConfig",function(n,t,e){var i=this,r=angular.isDefined(t.animate)?n.$parent.$eval(t.animate):e.animate;this.bars=[],n.max=angular.isDefined(t.max)?n.$parent.$eval(t.max):e.max,this.addBar=function(e,t){r||t.css({transition:"none"}),this.bars.push(e),e.$watch("value",function(t){e.percent=+(100*t/n.max).toFixed(2)}),e.$on("$destroy",function(){t=null,i.removeBar(e)})},this.removeBar=function(t){this.bars.splice(this.bars.indexOf(t),1)}}]).directive("progress",function(){return{restrict:"EA",replace:!0,transclude:!0,controller:"ProgressController",require:"progress",scope:{},templateUrl:"template/progressbar/progress.html"}}).directive("bar",function(){return{restrict:"EA",replace:!0,transclude:!0,require:"^progress",scope:{value:"=",type:"@"},templateUrl:"template/progressbar/bar.html",link:function(t,e,n,i){i.addBar(t,e)}}}).directive("progressbar",function(){return{restrict:"EA",replace:!0,transclude:!0,controller:"ProgressController",scope:{value:"=",type:"@"},templateUrl:"template/progressbar/progressbar.html",link:function(t,e,n,i){i.addBar(t,angular.element(e.children()[0]))}}}),angular.module("ui.bootstrap.rating",[]).constant("ratingConfig",{max:5,stateOn:null,stateOff:null}).controller("RatingController",["$scope","$attrs","ratingConfig",function(n,i,r){var o={$setViewValue:angular.noop};this.init=function(t){(o=t).$render=this.render,this.stateOn=angular.isDefined(i.stateOn)?n.$parent.$eval(i.stateOn):r.stateOn,this.stateOff=angular.isDefined(i.stateOff)?n.$parent.$eval(i.stateOff):r.stateOff;var e=angular.isDefined(i.ratingStates)?n.$parent.$eval(i.ratingStates):new Array(angular.isDefined(i.max)?n.$parent.$eval(i.max):r.max);n.range=this.buildTemplateObjects(e)},this.buildTemplateObjects=function(t){for(var e=0,n=t.length;e<n;e++)t[e]=angular.extend({index:e},{stateOn:this.stateOn,stateOff:this.stateOff},t[e]);return t},n.rate=function(t){!n.readonly&&0<=t&&t<=n.range.length&&(o.$setViewValue(t),o.$render())},n.enter=function(t){n.readonly||(n.value=t),n.onHover({value:t})},n.reset=function(){n.value=o.$viewValue,n.onLeave()},n.onKeydown=function(t){/(37|38|39|40)/.test(t.which)&&(t.preventDefault(),t.stopPropagation(),n.rate(n.value+(38===t.which||39===t.which?1:-1)))},this.render=function(){n.value=o.$viewValue}}]).directive("rating",function(){return{restrict:"EA",require:["rating","ngModel"],scope:{readonly:"=?",onHover:"&",onLeave:"&"},controller:"RatingController",templateUrl:"template/rating/rating.html",replace:!0,link:function(t,e,n,i){var r=i[0],o=i[1];o&&r.init(o)}}}),angular.module("ui.bootstrap.tabs",[]).controller("TabsetController",["$scope",function(t){var i,r=this,o=r.tabs=t.tabs=[];r.select=function(e){angular.forEach(o,function(t){t.active&&t!==e&&(t.active=!1,t.onDeselect())}),e.active=!0,e.onSelect()},r.addTab=function(t){o.push(t),1===o.length?t.active=!0:t.active&&r.select(t)},r.removeTab=function(t){var e=o.indexOf(t);if(t.active&&1<o.length&&!i){var n=e==o.length-1?e-1:e+1;r.select(o[n])}o.splice(e,1)},t.$on("$destroy",function(){i=!0})}]).directive("tabset",function(){return{restrict:"EA",transclude:!0,replace:!0,scope:{type:"@"},controller:"TabsetController",templateUrl:"template/tabs/tabset.html",link:function(t,e,n){t.vertical=!!angular.isDefined(n.vertical)&&t.$parent.$eval(n.vertical),t.justified=!!angular.isDefined(n.justified)&&t.$parent.$eval(n.justified)}}}).directive("tab",["$parse",function(o){return{require:"^tabset",restrict:"EA",replace:!0,templateUrl:"template/tabs/tab.html",transclude:!0,scope:{active:"=?",heading:"@",onSelect:"&select",onDeselect:"&deselect"},controller:function(){},compile:function(t,e,r){return function(e,t,n,i){e.$watch("active",function(t){t&&i.select(e)}),e.disabled=!1,n.disabled&&e.$parent.$watch(o(n.disabled),function(t){e.disabled=!!t}),e.select=function(){e.disabled||(e.active=!0)},i.addTab(e),e.$on("$destroy",function(){i.removeTab(e)}),e.$transcludeFn=r}}}}]).directive("tabHeadingTransclude",[function(){return{restrict:"A",require:"^tab",link:function(t,e,n,i){t.$watch("headingElement",function(t){t&&(e.html(""),e.append(t))})}}}]).directive("tabContentTransclude",function(){return{restrict:"A",require:"^tabset",link:function(t,e,n){var i=t.$eval(n.tabContentTransclude);i.$transcludeFn(i.$parent,function(t){angular.forEach(t,function(t){!function(t){return t.tagName&&(t.hasAttribute("tab-heading")||t.hasAttribute("data-tab-heading")||"tab-heading"===t.tagName.toLowerCase()||"data-tab-heading"===t.tagName.toLowerCase())}(t)?e.append(t):i.headingElement=t})})}}}),angular.module("ui.bootstrap.timepicker",[]).constant("timepickerConfig",{hourStep:1,minuteStep:1,showMeridian:!0,meridians:null,readonlyInput:!1,mousewheel:!0}).controller("TimepickerController",["$scope","$attrs","$parse","$log","$locale","timepickerConfig",function(r,o,t,e,n,a){function i(){var t=parseInt(r.hours,10);return(r.showMeridian?0<t&&t<13:0<=t&&t<24)?(r.showMeridian&&(12===t&&(t=0),r.meridian===g[1]&&(t+=12)),t):void 0}function s(){var t=parseInt(r.minutes,10);return 0<=t&&t<60?t:void 0}function l(t){return angular.isDefined(t)&&t.toString().length<2?"0"+t:t}function u(t){c(),f.$setViewValue(new Date(p)),h(t)}function c(){f.$setValidity("time",!0),r.invalidHours=!1,r.invalidMinutes=!1}function h(t){var e=p.getHours(),n=p.getMinutes();r.showMeridian&&(e=0===e||12===e?12:e%12),r.hours="h"===t?e:l(e),r.minutes="m"===t?n:l(n),r.meridian=p.getHours()<12?g[0]:g[1]}function d(t){var e=new Date(p.getTime()+6e4*t);p.setHours(e.getHours(),e.getMinutes()),u()}var p=new Date,f={$setViewValue:angular.noop},g=angular.isDefined(o.meridians)?r.$parent.$eval(o.meridians):a.meridians||n.DATETIME_FORMATS.AMPMS;this.init=function(t,e){(f=t).$render=this.render;var n=e.eq(0),i=e.eq(1);(angular.isDefined(o.mousewheel)?r.$parent.$eval(o.mousewheel):a.mousewheel)&&this.setupMousewheelEvents(n,i),r.readonlyInput=angular.isDefined(o.readonlyInput)?r.$parent.$eval(o.readonlyInput):a.readonlyInput,this.setupInputEvents(n,i)};var v=a.hourStep;o.hourStep&&r.$parent.$watch(t(o.hourStep),function(t){v=parseInt(t,10)});var m=a.minuteStep;o.minuteStep&&r.$parent.$watch(t(o.minuteStep),function(t){m=parseInt(t,10)}),r.showMeridian=a.showMeridian,o.showMeridian&&r.$parent.$watch(t(o.showMeridian),function(t){if(r.showMeridian=!!t,f.$error.time){var e=i(),n=s();angular.isDefined(e)&&angular.isDefined(n)&&(p.setHours(e),u())}else h()}),this.setupMousewheelEvents=function(t,e){function n(t){t.originalEvent&&(t=t.originalEvent);var e=t.wheelDelta?t.wheelDelta:-t.deltaY;return t.detail||0<e}t.bind("mousewheel wheel",function(t){r.$apply(n(t)?r.incrementHours():r.decrementHours()),t.preventDefault()}),e.bind("mousewheel wheel",function(t){r.$apply(n(t)?r.incrementMinutes():r.decrementMinutes()),t.preventDefault()})},this.setupInputEvents=function(t,e){if(r.readonlyInput)return r.updateHours=angular.noop,void(r.updateMinutes=angular.noop);function n(t,e){f.$setViewValue(null),f.$setValidity("time",!1),angular.isDefined(t)&&(r.invalidHours=t),angular.isDefined(e)&&(r.invalidMinutes=e)}r.updateHours=function(){var t=i();angular.isDefined(t)?(p.setHours(t),u("h")):n(!0)},t.bind("blur",function(t){!r.invalidHours&&r.hours<10&&r.$apply(function(){r.hours=l(r.hours)})}),r.updateMinutes=function(){var t=s();angular.isDefined(t)?(p.setMinutes(t),u("m")):n(void 0,!0)},e.bind("blur",function(t){!r.invalidMinutes&&r.minutes<10&&r.$apply(function(){r.minutes=l(r.minutes)})})},this.render=function(){var t=f.$modelValue?new Date(f.$modelValue):null;isNaN(t)?(f.$setValidity("time",!1),e.error('Timepicker directive: "ng-model" value must be a Date object, a number of milliseconds since 01.01.1970 or a string representing an RFC2822 or ISO 8601 date.')):(t&&(p=t),c(),h())},r.incrementHours=function(){d(60*v)},r.decrementHours=function(){d(60*-v)},r.incrementMinutes=function(){d(m)},r.decrementMinutes=function(){d(-m)},r.toggleMeridian=function(){d(720*(p.getHours()<12?1:-1))}}]).directive("timepicker",function(){return{restrict:"EA",require:["timepicker","?^ngModel"],controller:"TimepickerController",replace:!0,scope:{},templateUrl:"template/timepicker/timepicker.html",link:function(t,e,n,i){var r=i[0],o=i[1];o&&r.init(o,e.find("input"))}}}),angular.module("ui.bootstrap.typeahead",["ui.bootstrap.position","ui.bootstrap.bindHtml"]).factory("typeaheadParser",["$parse",function(n){var i=/^\s*([\s\S]+?)(?:\s+as\s+([\s\S]+?))?\s+for\s+(?:([\$\w][\$\w\d]*))\s+in\s+([\s\S]+?)$/;return{parse:function(t){var e=t.match(i);if(!e)throw new Error('Expected typeahead specification in form of "_modelValue_ (as _label_)? for _item_ in _collection_" but got "'+t+'".');return{itemName:e[3],source:n(e[4]),viewMapper:n(e[2]||e[1]),modelMapper:n(e[1])}}}}]).directive("typeahead",["$compile","$parse","$q","$timeout","$document","$position","typeaheadParser",function(E,S,T,D,_,P,M){var A=[9,13,27,38,40];return{require:"ngModel",link:function(o,a,t,s){var l,e=o.$eval(t.typeaheadMinLength)||1,n=o.$eval(t.typeaheadWaitMs)||0,i=!1!==o.$eval(t.typeaheadEditable),u=S(t.typeaheadLoading).assign||angular.noop,r=S(t.typeaheadOnSelect),c=t.typeaheadInputFormatter?S(t.typeaheadInputFormatter):void 0,h=!!t.typeaheadAppendToBody&&o.$eval(t.typeaheadAppendToBody),d=!1!==o.$eval(t.typeaheadFocusFirst),p=S(t.ngModel).assign,f=M.parse(t.typeahead),g=o.$new();o.$on("$destroy",function(){g.$destroy()});var v="typeahead-"+g.$id+"-"+Math.floor(1e4*Math.random());a.attr({"aria-autocomplete":"list","aria-expanded":!1,"aria-owns":v});var m=angular.element("<div typeahead-popup></div>");m.attr({id:v,matches:"matches",active:"activeIdx",select:"select(activeIdx)",query:"query",position:"position"}),angular.isDefined(t.typeaheadTemplateUrl)&&m.attr("template-url",t.typeaheadTemplateUrl);function y(){g.matches=[],g.activeIdx=-1,a.attr("aria-expanded",!1)}function b(t){return v+"-option-"+t}g.$watch("activeIdx",function(t){t<0?a.removeAttr("aria-activedescendant"):a.attr("aria-activedescendant",b(t))});function x(i){var r={$viewValue:i};u(o,!0),T.when(f.source(o,r)).then(function(t){var e=i===s.$viewValue;if(e&&l)if(0<t.length){g.activeIdx=d?0:-1;for(var n=g.matches.length=0;n<t.length;n++)r[f.itemName]=t[n],g.matches.push({id:b(n),label:f.viewMapper(g,r),model:t[n]});g.query=i,g.position=h?P.offset(a):P.position(a),g.position.top=g.position.top+a.prop("offsetHeight"),a.attr("aria-expanded",!0)}else y();e&&u(o,!1)},function(){y(),u(o,!1)})}y(),g.query=void 0;function w(){$&&D.cancel($)}var $;s.$parsers.unshift(function(t){return l=!0,t&&t.length>=e?0<n?(w(),function(t){$=D(function(){x(t)},n)}(t)):x(t):(u(o,!1),w(),y()),i?t:t?void s.$setValidity("editable",!1):(s.$setValidity("editable",!0),t)}),s.$formatters.push(function(t){var e,n={};return c?(n.$model=t,c(o,n)):(n[f.itemName]=t,e=f.viewMapper(o,n),n[f.itemName]=void 0,e!==f.viewMapper(o,n)?e:t)}),g.select=function(t){var e,n,i={};i[f.itemName]=n=g.matches[t].model,e=f.modelMapper(o,i),p(o,e),s.$setValidity("editable",!0),r(o,{$item:n,$model:e,$label:f.viewMapper(o,i)}),y(),D(function(){a[0].focus()},0,!1)},a.bind("keydown",function(t){0!==g.matches.length&&-1!==A.indexOf(t.which)&&(-1!=g.activeIdx||13!==t.which&&9!==t.which)&&(t.preventDefault(),40===t.which?(g.activeIdx=(g.activeIdx+1)%g.matches.length,g.$digest()):38===t.which?(g.activeIdx=(0<g.activeIdx?g.activeIdx:g.matches.length)-1,g.$digest()):13===t.which||9===t.which?g.$apply(function(){g.select(g.activeIdx)}):27===t.which&&(t.stopPropagation(),y(),g.$digest()))}),a.bind("blur",function(t){l=!1});function k(t){a[0]!==t.target&&(y(),g.$digest())}_.bind("click",k),o.$on("$destroy",function(){_.unbind("click",k),h&&C.remove()});var C=E(m)(g);h?_.find("body").append(C):a.after(C)}}}]).directive("typeaheadPopup",function(){return{restrict:"EA",scope:{matches:"=",query:"=",active:"=",position:"=",select:"&"},replace:!0,templateUrl:"template/typeahead/typeahead-popup.html",link:function(e,t,n){e.templateUrl=n.templateUrl,e.isOpen=function(){return 0<e.matches.length},e.isActive=function(t){return e.active==t},e.selectActive=function(t){e.active=t},e.selectMatch=function(t){e.select({activeIdx:t})}}}}).directive("typeaheadMatch",["$http","$templateCache","$compile","$parse",function(r,o,a,s){return{restrict:"EA",scope:{index:"=",match:"=",query:"="},link:function(e,n,t){var i=s(t.templateUrl)(e.$parent)||"template/typeahead/typeahead-match.html";r.get(i,{cache:o}).success(function(t){n.replaceWith(a(t.trim())(e))})}}}]).filter("typeaheadHighlight",function(){return function(t,e){return e?(""+t).replace(new RegExp(function(t){return t.replace(/([.?*+^$[\]\\(){}|-])/g,"\\$1")}(e),"gi"),"<strong>$&</strong>"):t}}),angular.module("template/accordion/accordion-group.html",[]).run(["$templateCache",function(t){t.put("template/accordion/accordion-group.html",'<div class="panel panel-default">\n  <div class="panel-heading">\n    <h4 class="panel-title">\n      <a href class="accordion-toggle" ng-click="toggleOpen()" accordion-transclude="heading"><span ng-class="{\'text-muted\': isDisabled}">{{heading}}</span></a>\n    </h4>\n  </div>\n  <div class="panel-collapse" collapse="!isOpen">\n\t  <div class="panel-body" ng-transclude></div>\n  </div>\n</div>\n')}]),angular.module("template/accordion/accordion.html",[]).run(["$templateCache",function(t){t.put("template/accordion/accordion.html",'<div class="panel-group" ng-transclude></div>')}]),angular.module("template/alert/alert.html",[]).run(["$templateCache",function(t){t.put("template/alert/alert.html",'<div class="alert" ng-class="[\'alert-\' + (type || \'warning\'), closeable ? \'alert-dismissable\' : null]" role="alert">\n    <button ng-show="closeable" type="button" class="close" ng-click="close()">\n        <span aria-hidden="true">&times;</span>\n        <span class="sr-only">Close</span>\n    </button>\n    <div ng-transclude></div>\n</div>\n')}]),angular.module("template/carousel/carousel.html",[]).run(["$templateCache",function(t){t.put("template/carousel/carousel.html",'<div ng-mouseenter="pause()" ng-mouseleave="play()" class="carousel" ng-swipe-right="prev()" ng-swipe-left="next()">\n    <ol class="carousel-indicators" ng-show="slides.length > 1">\n        <li ng-repeat="slide in slides track by $index" ng-class="{active: isActive(slide)}" ng-click="select(slide)"></li>\n    </ol>\n    <div class="carousel-inner" ng-transclude></div>\n    <a class="left carousel-control" ng-click="prev()" ng-show="slides.length > 1"><span class="glyphicon glyphicon-chevron-left"></span></a>\n    <a class="right carousel-control" ng-click="next()" ng-show="slides.length > 1"><span class="glyphicon glyphicon-chevron-right"></span></a>\n</div>\n')}]),angular.module("template/carousel/slide.html",[]).run(["$templateCache",function(t){t.put("template/carousel/slide.html","<div ng-class=\"{\n    'active': leaving || (active && !entering),\n    'prev': (next || active) && direction=='prev',\n    'next': (next || active) && direction=='next',\n    'right': direction=='prev',\n    'left': direction=='next'\n  }\" class=\"item text-center\" ng-transclude></div>\n")}]),angular.module("template/datepicker/datepicker.html",[]).run(["$templateCache",function(t){t.put("template/datepicker/datepicker.html",'<div ng-switch="datepickerMode" role="application" ng-keydown="keydown($event)">\n  <daypicker ng-switch-when="day" tabindex="0"></daypicker>\n  <monthpicker ng-switch-when="month" tabindex="0"></monthpicker>\n  <yearpicker ng-switch-when="year" tabindex="0"></yearpicker>\n</div>')}]),angular.module("template/datepicker/day.html",[]).run(["$templateCache",function(t){t.put("template/datepicker/day.html",'<table role="grid" aria-labelledby="{{uniqueId}}-title" aria-activedescendant="{{activeDateId}}">\n  <thead>\n    <tr>\n      <th><button type="button" class="btn btn-default btn-sm pull-left" ng-click="move(-1)" tabindex="-1"><i class="glyphicon glyphicon-chevron-left"></i></button></th>\n      <th colspan="{{5 + showWeeks}}"><button id="{{uniqueId}}-title" role="heading" aria-live="assertive" aria-atomic="true" type="button" class="btn btn-default btn-sm" ng-click="toggleMode()" tabindex="-1" style="width:100%;"><strong>{{title}}</strong></button></th>\n      <th><button type="button" class="btn btn-default btn-sm pull-right" ng-click="move(1)" tabindex="-1"><i class="glyphicon glyphicon-chevron-right"></i></button></th>\n    </tr>\n    <tr>\n      <th ng-show="showWeeks" class="text-center"></th>\n      <th ng-repeat="label in labels track by $index" class="text-center"><small aria-label="{{label.full}}">{{label.abbr}}</small></th>\n    </tr>\n  </thead>\n  <tbody>\n    <tr ng-repeat="row in rows track by $index">\n      <td ng-show="showWeeks" class="text-center h6"><em>{{ weekNumbers[$index] }}</em></td>\n      <td ng-repeat="dt in row track by dt.date" class="text-center" role="gridcell" id="{{dt.uid}}" aria-disabled="{{!!dt.disabled}}">\n        <button type="button" style="width:100%;" class="btn btn-default btn-sm" ng-class="{\'btn-info\': dt.selected, active: isActive(dt)}" ng-click="select(dt.date)" ng-disabled="dt.disabled" tabindex="-1"><span ng-class="{\'text-muted\': dt.secondary, \'text-info\': dt.current}">{{dt.label}}</span></button>\n      </td>\n    </tr>\n  </tbody>\n</table>\n')}]),angular.module("template/datepicker/month.html",[]).run(["$templateCache",function(t){t.put("template/datepicker/month.html",'<table role="grid" aria-labelledby="{{uniqueId}}-title" aria-activedescendant="{{activeDateId}}">\n  <thead>\n    <tr>\n      <th><button type="button" class="btn btn-default btn-sm pull-left" ng-click="move(-1)" tabindex="-1"><i class="glyphicon glyphicon-chevron-left"></i></button></th>\n      <th><button id="{{uniqueId}}-title" role="heading" aria-live="assertive" aria-atomic="true" type="button" class="btn btn-default btn-sm" ng-click="toggleMode()" tabindex="-1" style="width:100%;"><strong>{{title}}</strong></button></th>\n      <th><button type="button" class="btn btn-default btn-sm pull-right" ng-click="move(1)" tabindex="-1"><i class="glyphicon glyphicon-chevron-right"></i></button></th>\n    </tr>\n  </thead>\n  <tbody>\n    <tr ng-repeat="row in rows track by $index">\n      <td ng-repeat="dt in row track by dt.date" class="text-center" role="gridcell" id="{{dt.uid}}" aria-disabled="{{!!dt.disabled}}">\n        <button type="button" style="width:100%;" class="btn btn-default" ng-class="{\'btn-info\': dt.selected, active: isActive(dt)}" ng-click="select(dt.date)" ng-disabled="dt.disabled" tabindex="-1"><span ng-class="{\'text-info\': dt.current}">{{dt.label}}</span></button>\n      </td>\n    </tr>\n  </tbody>\n</table>\n')}]),angular.module("template/datepicker/popup.html",[]).run(["$templateCache",function(t){t.put("template/datepicker/popup.html",'<ul class="dropdown-menu" ng-style="{display: (isOpen && \'block\') || \'none\', top: position.top+\'px\', left: position.left+\'px\'}" ng-keydown="keydown($event)">\n\t<li ng-transclude></li>\n\t<li ng-if="showButtonBar" style="padding:10px 9px 2px">\n\t\t<span class="btn-group pull-left">\n\t\t\t<button type="button" class="btn btn-sm btn-info" ng-click="select(\'today\')">{{ getText(\'current\') }}</button>\n\t\t\t<button type="button" class="btn btn-sm btn-danger" ng-click="select(null)">{{ getText(\'clear\') }}</button>\n\t\t</span>\n\t\t<button type="button" class="btn btn-sm btn-success pull-right" ng-click="close()">{{ getText(\'close\') }}</button>\n\t</li>\n</ul>\n')}]),angular.module("template/datepicker/year.html",[]).run(["$templateCache",function(t){t.put("template/datepicker/year.html",'<table role="grid" aria-labelledby="{{uniqueId}}-title" aria-activedescendant="{{activeDateId}}">\n  <thead>\n    <tr>\n      <th><button type="button" class="btn btn-default btn-sm pull-left" ng-click="move(-1)" tabindex="-1"><i class="glyphicon glyphicon-chevron-left"></i></button></th>\n      <th colspan="3"><button id="{{uniqueId}}-title" role="heading" aria-live="assertive" aria-atomic="true" type="button" class="btn btn-default btn-sm" ng-click="toggleMode()" tabindex="-1" style="width:100%;"><strong>{{title}}</strong></button></th>\n      <th><button type="button" class="btn btn-default btn-sm pull-right" ng-click="move(1)" tabindex="-1"><i class="glyphicon glyphicon-chevron-right"></i></button></th>\n    </tr>\n  </thead>\n  <tbody>\n    <tr ng-repeat="row in rows track by $index">\n      <td ng-repeat="dt in row track by dt.date" class="text-center" role="gridcell" id="{{dt.uid}}" aria-disabled="{{!!dt.disabled}}">\n        <button type="button" style="width:100%;" class="btn btn-default" ng-class="{\'btn-info\': dt.selected, active: isActive(dt)}" ng-click="select(dt.date)" ng-disabled="dt.disabled" tabindex="-1"><span ng-class="{\'text-info\': dt.current}">{{dt.label}}</span></button>\n      </td>\n    </tr>\n  </tbody>\n</table>\n')}]),angular.module("template/modal/backdrop.html",[]).run(["$templateCache",function(t){t.put("template/modal/backdrop.html",'<div class="modal-backdrop fade {{ backdropClass }}"\n     ng-class="{in: animate}"\n     ng-style="{\'z-index\': 1040 + (index && 1 || 0) + index*10}"\n></div>\n')}]),angular.module("template/modal/window.html",[]).run(["$templateCache",function(t){t.put("template/modal/window.html",'<div tabindex="-1" role="dialog" class="modal fade" ng-class="{in: animate}" ng-style="{\'z-index\': 1050 + index*10, display: \'block\'}" ng-click="close($event)">\n    <div class="modal-dialog" ng-class="{\'modal-sm\': size == \'sm\', \'modal-lg\': size == \'lg\'}"><div class="modal-content" modal-transclude></div></div>\n</div>')}]),angular.module("template/pagination/pager.html",[]).run(["$templateCache",function(t){t.put("template/pagination/pager.html",'<ul class="pager">\n  <li ng-class="{disabled: noPrevious(), previous: align}"><a href ng-click="selectPage(page - 1)">{{getText(\'previous\')}}</a></li>\n  <li ng-class="{disabled: noNext(), next: align}"><a href ng-click="selectPage(page + 1)">{{getText(\'next\')}}</a></li>\n</ul>')}]),angular.module("template/pagination/pagination.html",[]).run(["$templateCache",function(t){t.put("template/pagination/pagination.html",'<ul class="pagination">\n  <li ng-if="boundaryLinks" ng-class="{disabled: noPrevious()}"><a href ng-click="selectPage(1)">{{getText(\'first\')}}</a></li>\n  <li ng-if="directionLinks" ng-class="{disabled: noPrevious()}"><a href ng-click="selectPage(page - 1)">{{getText(\'previous\')}}</a></li>\n  <li ng-repeat="page in pages track by $index" ng-class="{active: page.active}"><a href ng-click="selectPage(page.number)">{{page.text}}</a></li>\n  <li ng-if="directionLinks" ng-class="{disabled: noNext()}"><a href ng-click="selectPage(page + 1)">{{getText(\'next\')}}</a></li>\n  <li ng-if="boundaryLinks" ng-class="{disabled: noNext()}"><a href ng-click="selectPage(totalPages)">{{getText(\'last\')}}</a></li>\n</ul>')}]),angular.module("template/tooltip/tooltip-html-unsafe-popup.html",[]).run(["$templateCache",function(t){t.put("template/tooltip/tooltip-html-unsafe-popup.html",'<div class="tooltip {{placement}}" ng-class="{ in: isOpen(), fade: animation() }">\n  <div class="tooltip-arrow"></div>\n  <div class="tooltip-inner" bind-html-unsafe="content"></div>\n</div>\n')}]),angular.module("template/tooltip/tooltip-popup.html",[]).run(["$templateCache",function(t){t.put("template/tooltip/tooltip-popup.html",'<div class="tooltip {{placement}}" ng-class="{ in: isOpen(), fade: animation() }">\n  <div class="tooltip-arrow"></div>\n  <div class="tooltip-inner" ng-bind="content"></div>\n</div>\n')}]),angular.module("template/popover/popover.html",[]).run(["$templateCache",function(t){t.put("template/popover/popover.html",'<div class="popover {{placement}}" ng-class="{ in: isOpen(), fade: animation() }">\n  <div class="arrow"></div>\n\n  <div class="popover-inner">\n      <h3 class="popover-title" ng-bind="title" ng-show="title"></h3>\n      <div class="popover-content" ng-bind="content"></div>\n  </div>\n</div>\n')}]),angular.module("template/progressbar/bar.html",[]).run(["$templateCache",function(t){t.put("template/progressbar/bar.html",'<div class="progress-bar" ng-class="type && \'progress-bar-\' + type" role="progressbar" aria-valuenow="{{value}}" aria-valuemin="0" aria-valuemax="{{max}}" ng-style="{width: percent + \'%\'}" aria-valuetext="{{percent | number:0}}%" ng-transclude></div>')}]),angular.module("template/progressbar/progress.html",[]).run(["$templateCache",function(t){t.put("template/progressbar/progress.html",'<div class="progress" ng-transclude></div>')}]),angular.module("template/progressbar/progressbar.html",[]).run(["$templateCache",function(t){t.put("template/progressbar/progressbar.html",'<div class="progress">\n  <div class="progress-bar" ng-class="type && \'progress-bar-\' + type" role="progressbar" aria-valuenow="{{value}}" aria-valuemin="0" aria-valuemax="{{max}}" ng-style="{width: percent + \'%\'}" aria-valuetext="{{percent | number:0}}%" ng-transclude></div>\n</div>')}]),angular.module("template/rating/rating.html",[]).run(["$templateCache",function(t){t.put("template/rating/rating.html",'<span ng-mouseleave="reset()" ng-keydown="onKeydown($event)" tabindex="0" role="slider" aria-valuemin="0" aria-valuemax="{{range.length}}" aria-valuenow="{{value}}">\n    <i ng-repeat="r in range track by $index" ng-mouseenter="enter($index + 1)" ng-click="rate($index + 1)" class="glyphicon" ng-class="$index < value && (r.stateOn || \'glyphicon-star\') || (r.stateOff || \'glyphicon-star-empty\')">\n        <span class="sr-only">({{ $index < value ? \'*\' : \' \' }})</span>\n    </i>\n</span>')}]),angular.module("template/tabs/tab.html",[]).run(["$templateCache",function(t){t.put("template/tabs/tab.html",'<li ng-class="{active: active, disabled: disabled}">\n  <a href ng-click="select()" tab-heading-transclude>{{heading}}</a>\n</li>\n')}]),angular.module("template/tabs/tabset.html",[]).run(["$templateCache",function(t){t.put("template/tabs/tabset.html",'<div>\n  <ul class="nav nav-{{type || \'tabs\'}}" ng-class="{\'nav-stacked\': vertical, \'nav-justified\': justified}" ng-transclude></ul>\n  <div class="tab-content">\n    <div class="tab-pane" \n         ng-repeat="tab in tabs" \n         ng-class="{active: tab.active}"\n         tab-content-transclude="tab">\n    </div>\n  </div>\n</div>\n')}]),angular.module("template/timepicker/timepicker.html",[]).run(["$templateCache",function(t){t.put("template/timepicker/timepicker.html",'<table>\n\t<tbody>\n\t\t<tr class="text-center">\n\t\t\t<td><a ng-click="incrementHours()" class="btn btn-link"><span class="glyphicon glyphicon-chevron-up"></span></a></td>\n\t\t\t<td>&nbsp;</td>\n\t\t\t<td><a ng-click="incrementMinutes()" class="btn btn-link"><span class="glyphicon glyphicon-chevron-up"></span></a></td>\n\t\t\t<td ng-show="showMeridian"></td>\n\t\t</tr>\n\t\t<tr>\n\t\t\t<td style="width:50px;" class="form-group" ng-class="{\'has-error\': invalidHours}">\n\t\t\t\t<input type="text" ng-model="hours" ng-change="updateHours()" class="form-control text-center" ng-mousewheel="incrementHours()" ng-readonly="readonlyInput" maxlength="2">\n\t\t\t</td>\n\t\t\t<td>:</td>\n\t\t\t<td style="width:50px;" class="form-group" ng-class="{\'has-error\': invalidMinutes}">\n\t\t\t\t<input type="text" ng-model="minutes" ng-change="updateMinutes()" class="form-control text-center" ng-readonly="readonlyInput" maxlength="2">\n\t\t\t</td>\n\t\t\t<td ng-show="showMeridian"><button type="button" class="btn btn-default text-center" ng-click="toggleMeridian()">{{meridian}}</button></td>\n\t\t</tr>\n\t\t<tr class="text-center">\n\t\t\t<td><a ng-click="decrementHours()" class="btn btn-link"><span class="glyphicon glyphicon-chevron-down"></span></a></td>\n\t\t\t<td>&nbsp;</td>\n\t\t\t<td><a ng-click="decrementMinutes()" class="btn btn-link"><span class="glyphicon glyphicon-chevron-down"></span></a></td>\n\t\t\t<td ng-show="showMeridian"></td>\n\t\t</tr>\n\t</tbody>\n</table>\n')}]),angular.module("template/typeahead/typeahead-match.html",[]).run(["$templateCache",function(t){t.put("template/typeahead/typeahead-match.html",'<a tabindex="-1" bind-html-unsafe="match.label | typeaheadHighlight:query"></a>')}]),angular.module("template/typeahead/typeahead-popup.html",[]).run(["$templateCache",function(t){t.put("template/typeahead/typeahead-popup.html",'<ul class="dropdown-menu" ng-show="isOpen()" ng-style="{top: position.top+\'px\', left: position.left+\'px\'}" style="display: block;" role="listbox" aria-hidden="{{!isOpen()}}">\n    <li ng-repeat="match in matches track by $index" ng-class="{active: isActive($index) }" ng-mouseenter="selectActive($index)" ng-click="selectMatch($index)" role="option" id="{{match.id}}">\n        <div typeahead-match index="$index" match="match" query="query" template-url="templateUrl"></div>\n    </li>\n</ul>\n')}]),angular.module("colorpicker.module",[]).factory("Helper",function(){"use strict";return{closestSlider:function(t){return(t.matches||t.webkitMatchesSelector||t.mozMatchesSelector||t.msMatchesSelector).bind(t)("I")?t.parentNode:t},getOffset:function(t,e){for(var n=0,i=0,r=0,o=0;t&&!isNaN(t.offsetLeft)&&!isNaN(t.offsetTop);)n+=t.offsetLeft,i+=t.offsetTop,e||"BODY"!==t.tagName?(r+=t.scrollLeft,o+=t.scrollTop):(r+=document.documentElement.scrollLeft||t.scrollLeft,o+=document.documentElement.scrollTop||t.scrollTop),t=t.offsetParent;return{top:i,left:n,scrollX:r,scrollY:o}},stringParsers:[{re:/rgba?\(\s*(\d{1,3})\s*,\s*(\d{1,3})\s*,\s*(\d{1,3})\s*(?:,\s*(\d+(?:\.\d+)?)\s*)?\)/,parse:function(t){return[t[1],t[2],t[3],t[4]]}},{re:/rgba?\(\s*(\d+(?:\.\d+)?)\%\s*,\s*(\d+(?:\.\d+)?)\%\s*,\s*(\d+(?:\.\d+)?)\%\s*(?:,\s*(\d+(?:\.\d+)?)\s*)?\)/,parse:function(t){return[2.55*t[1],2.55*t[2],2.55*t[3],t[4]]}},{re:/#([a-fA-F0-9]{2})([a-fA-F0-9]{2})([a-fA-F0-9]{2})/,parse:function(t){return[parseInt(t[1],16),parseInt(t[2],16),parseInt(t[3],16)]}},{re:/#([a-fA-F0-9])([a-fA-F0-9])([a-fA-F0-9])/,parse:function(t){return[parseInt(t[1]+t[1],16),parseInt(t[2]+t[2],16),parseInt(t[3]+t[3],16)]}}]}}).factory("Color",["Helper",function(o){"use strict";return{value:{h:1,s:1,b:1,a:1},rgb:function(){var t=this.toRGB();return"rgb("+t.r+","+t.g+","+t.b+")"},rgba:function(){var t=this.toRGB();return"rgba("+t.r+","+t.g+","+t.b+","+t.a+")"},hex:function(){return this.toHex()},RGBtoHSB:function(t,e,n,i){var r,o;return t/=255,e/=255,n/=255,{h:((0==(o=(r=Math.max(t,e,n))-Math.min(t,e,n))?null:r===t?(e-n)/o:r===e?(n-t)/o+2:(t-e)/o+4)+360)%6*60/360||1,s:0==o?0:o/r,b:r,a:i||1}},setColor:function(t){for(var e in t=t.toLowerCase(),o.stringParsers)if(o.stringParsers.hasOwnProperty(e)){var n=o.stringParsers[e],i=n.re.exec(t),r=i&&n.parse(i);if(r)return this.value=this.RGBtoHSB.apply(null,r),!1}},setHue:function(t){this.value.h=1-t},setSaturation:function(t){this.value.s=t},setLightness:function(t){this.value.b=1-t},setAlpha:function(t){this.value.a=parseInt(100*(1-t),10)/100},toRGB:function(t,e,n,i){var r,o,a,s,l;return t||(t=this.value.h,e=this.value.s,n=this.value.b),t=(t*=360)%360/60,r=o=a=n-(l=n*e),r+=[l,s=l*(1-Math.abs(t%2-1)),0,0,s,l][t=~~t],o+=[s,l,l,s,0,0][t],a+=[0,0,s,l,l,s][t],{r:Math.round(255*r),g:Math.round(255*o),b:Math.round(255*a),a:i||this.value.a}},toHex:function(t,e,n,i){var r=this.toRGB(t,e,n,i);return"#"+(1<<24|parseInt(r.r,10)<<16|parseInt(r.g,10)<<8|parseInt(r.b,10)).toString(16).substr(1)}}}]).factory("Slider",["Helper",function(s){"use strict";var l={maxLeft:0,maxTop:0,callLeft:null,callTop:null,knob:{top:0,left:0}},u={};return{getSlider:function(){return l},getLeftPosition:function(t){return Math.max(0,Math.min(l.maxLeft,l.left+((t.pageX||u.left)-u.left)))},getTopPosition:function(t){return Math.max(0,Math.min(l.maxTop,l.top+((t.pageY||u.top)-u.top)))},setSlider:function(t,e){var n=s.closestSlider(t.target),i=s.getOffset(n,e),r=n.getBoundingClientRect(),o=t.clientX-r.left,a=t.clientY-r.top;l.knob=n.children[0].style,l.left=t.pageX-i.left-window.pageXOffset+i.scrollX,l.top=t.pageY-i.top-window.pageYOffset+i.scrollY,u={left:t.pageX-(o-l.left),top:t.pageY-(a-l.top)}},setSaturation:function(t,e){l={maxLeft:100,maxTop:100,callLeft:"setSaturation",callTop:"setLightness"},this.setSlider(t,e)},setHue:function(t,e){l={maxLeft:0,maxTop:100,callLeft:!1,callTop:"setHue"},this.setSlider(t,e)},setAlpha:function(t,e){l={maxLeft:0,maxTop:100,callLeft:!1,callTop:"setAlpha"},this.setSlider(t,e)},setKnob:function(t,e){l.knob.top=t+"px",l.knob.left=e+"px"}}}]).directive("colorpicker",["$document","$compile","Color","Slider","Helper",function(D,_,P,M,A){"use strict";return{require:"?ngModel",restrict:"A",link:function(o,a,n,s){var t,l=n.colorpicker?n.colorpicker:"hex",i=angular.isDefined(n.colorpickerPosition)?n.colorpickerPosition:"bottom",e=!!angular.isDefined(n.colorpickerInline)&&n.colorpickerInline,r=!!angular.isDefined(n.colorpickerFixedPosition)&&n.colorpickerFixedPosition,u=angular.isDefined(n.colorpickerParent)?a.parent():angular.element(document.body),c=!!angular.isDefined(n.colorpickerWithInput)&&n.colorpickerWithInput,h='<div class="colorpicker dropdown"><div class="dropdown-menu"><colorpicker-saturation><i></i></colorpicker-saturation><colorpicker-hue><i></i></colorpicker-hue><colorpicker-alpha><i></i></colorpicker-alpha><colorpicker-preview></colorpicker-preview>'+(c?'<input type="text" name="colorpicker-input">':"")+(e?"":'<button type="button" class="close close-colorpicker">&times;</button>')+"</div></div>",d=angular.element(h),p=P,f=d.find("colorpicker-hue"),g=d.find("colorpicker-saturation"),v=d.find("colorpicker-preview"),m=d.find("i");if(_(d)(o),c){var y=d.find("input");y.on("mousedown",function(t){t.stopPropagation()}).on("keyup",function(t){var e=this.value;a.val(e),s&&o.$apply(s.$setViewValue(e)),t.stopPropagation(),t.preventDefault()}),a.on("keyup",function(){y.val(a.val())})}function b(){D.on("mousemove",k),D.on("mouseup",C)}"rgba"===l&&(d.addClass("alpha"),(t=d.find("colorpicker-alpha")).on("click",function(t){M.setAlpha(t,r),k(t)}).on("mousedown",function(t){M.setAlpha(t,r),b()}).on("mouseup",function(t){S("colorpicker-selected-alpha")})),f.on("click",function(t){M.setHue(t,r),k(t)}).on("mousedown",function(t){M.setHue(t,r),b()}).on("mouseup",function(t){S("colorpicker-selected-hue")}),g.on("click",function(t){M.setSaturation(t,r),k(t),angular.isDefined(n.colorpickerCloseOnSelect)&&T()}).on("mousedown",function(t){M.setSaturation(t,r),b()}).on("mouseup",function(t){S("colorpicker-selected-saturation")}),r&&d.addClass("colorpicker-fixed-position"),d.addClass("colorpicker-position-"+i),"true"===e&&d.addClass("colorpicker-inline"),u.append(d),s&&(s.$render=function(){a.val(s.$viewValue)},o.$watch(n.ngModel,function(t){E(),c&&y.val(t)})),a.on("$destroy",function(){d.remove()});function x(){try{v.css("backgroundColor",p[l]())}catch(t){v.css("backgroundColor",p.toHex())}g.css("backgroundColor",p.toHex(p.value.h,1,1,1)),"rgba"===l&&(t.css.backgroundColor=p.toHex())}function w(){T()}function $(){d.hasClass("colorpicker-visible")||(E(),d.addClass("colorpicker-visible").css(function(){var t,e=A.getOffset(a[0]);return angular.isDefined(n.colorpickerParent)&&(e.left=0,e.top=0),"top"===i?t={top:e.top-147,left:e.left}:"right"===i?t={top:e.top,left:e.left+126}:"bottom"===i?t={top:e.top+a[0].offsetHeight+2,left:e.left}:"left"===i&&(t={top:e.top,left:e.left-150}),{top:t.top+"px",left:t.left+"px"}}()),S("colorpicker-shown"),!1===e&&D.on("mousedown",w),n.colorpickerIsOpen&&(o[n.colorpickerIsOpen]=!0,o.$$phase||o.$digest()))}var k=function(t){var e=M.getLeftPosition(t),n=M.getTopPosition(t),i=M.getSlider();M.setKnob(n,e),i.callLeft&&p[i.callLeft].call(p,e/100),i.callTop&&p[i.callTop].call(p,n/100),x();var r=p[l]();return a.val(r),s&&o.$apply(s.$setViewValue(r)),c&&y.val(r),!1},C=function(){S("colorpicker-selected"),D.off("mousemove",k),D.off("mouseup",C)},E=function(){p.setColor(a.val()),m.eq(0).css({left:100*p.value.s+"px",top:100-100*p.value.b+"px"}),m.eq(1).css("top",100*(1-p.value.h)+"px"),m.eq(2).css("top",100*(1-p.value.a)+"px"),x()};!1===e?a.on("click",$):$(),d.on("mousedown",function(t){t.stopPropagation(),t.preventDefault()});var S=function(t){s&&o.$emit(t,{name:n.ngModel,value:s.$modelValue})},T=function(){d.hasClass("colorpicker-visible")&&(d.removeClass("colorpicker-visible"),S("colorpicker-closed"),D.off("mousedown",w),n.colorpickerIsOpen&&(o[n.colorpickerIsOpen]=!1,o.$$phase||o.$digest()))};d.find("button").on("click",function(){T()}),n.colorpickerIsOpen&&o.$watch(n.colorpickerIsOpen,function(t){!0===t?$():!1===t&&T()})}}}]),function(t,u){"use strict";u.module("ngCookies",["ng"]).factory("$cookies",["$rootScope","$browser",function(e,r){var n,o={},a={},i=!1,s=u.copy,l=u.isUndefined;return r.addPollFn(function(){var t=r.cookies();n!=t&&(s(n=t,a),s(t,o),i&&e.$apply())})(),i=!0,e.$watch(function(){var t,e,n,i;for(t in a)l(o[t])&&r.cookies(t,void 0);for(t in o)e=o[t],u.isString(e)||(e=""+e,o[t]=e),e!==a[t]&&(r.cookies(t,e),i=!0);if(i)for(t in i=!1,n=r.cookies(),o)o[t]!==n[t]&&(l(n[t])?delete o[t]:o[t]=n[t],i=!0)}),o}]).factory("$cookieStore",["$cookies",function(n){return{get:function(t){var e=n[t];return e?u.fromJson(e):e},put:function(t,e){n[t]=u.toJson(e)},remove:function(t){delete n[t]}}}])}(window,window.angular),function(t,T,D){"use strict";function _(t,e){if(!function(t){return null!=t&&""!==t&&"hasOwnProperty"!==t&&a.test("."+t)}(e))throw M("badmember",'Dotted member path "@{0}" is invalid.',e);for(var n=e.split("."),i=0,r=n.length;i<r&&t!==D;i++){var o=n[i];t=null!==t?t[o]:D}return t}function P(t,n){for(var e in n=n||{},T.forEach(n,function(t,e){delete n[e]}),t)!t.hasOwnProperty(e)||"$"===e.charAt(0)&&"$"===e.charAt(1)||(n[e]=t[e]);return n}var M=T.$$minErr("$resource"),a=/^(\.[a-zA-Z_$][0-9a-zA-Z_$]*)+$/;T.module("ngResource",["ng"]).factory("$resource",["$http","$q",function(x,w){function l(t){return function(t,e){return encodeURIComponent(t).replace(/%40/gi,"@").replace(/%3A/gi,":").replace(/%24/g,"$").replace(/%2C/gi,",").replace(/%20/g,e?"%20":"+")}(t,!0).replace(/%26/gi,"&").replace(/%3D/gi,"=").replace(/%2B/gi,"+")}function t(t,e){this.template=t,this.defaults=e||{},this.urlParams={}}var r={get:{method:"GET"},save:{method:"POST"},query:{method:"GET",isArray:!0},remove:{method:"DELETE"},delete:{method:"DELETE"}},$=T.noop,k=T.forEach,C=T.extend,E=T.copy,S=T.isFunction;return t.prototype={setUrlParams:function(n,i,t){var r,o,a=this,s=t||a.template,e=a.urlParams={};k(s.split(/\W/),function(t){if("hasOwnProperty"===t)throw M("badname","hasOwnProperty is not a valid parameter name.");!new RegExp("^\\d+$").test(t)&&t&&new RegExp("(^|[^\\\\]):"+t+"(\\W|$)").test(s)&&(e[t]=!0)}),s=s.replace(/\\:/g,":"),i=i||{},k(a.urlParams,function(t,e){r=i.hasOwnProperty(e)?i[e]:a.defaults[e],s=T.isDefined(r)&&null!==r?(o=l(r),s.replace(new RegExp(":"+e+"(\\W|$)","g"),function(t,e){return o+e})):s.replace(new RegExp("(/?):"+e+"(\\W|$)","g"),function(t,e,n){return"/"==n.charAt(0)?n:e+n})}),s=(s=s.replace(/\/+$/,"")||"/").replace(/\/\.(?=\w+($|\?))/,"."),n.url=s.replace(/\/\\\./,"/."),k(i,function(t,e){a.urlParams[e]||(n.params=n.params||{},n.params[e]=t)})}},function e(n,v,i){function m(t){return t.resource}function y(t){P(t||{},this)}var b=new t(n);return i=C({},r,i),k(i,function(f,r){var g=/^(POST|PUT|PATCH)$/i.test(f.method);y[r]=function(t,e,n,i){var r,o,a,s={};switch(arguments.length){case 4:a=i,o=n;case 3:case 2:if(!S(e)){s=t,r=e,o=n;break}if(S(t)){o=t,a=e;break}o=e,a=n;case 1:S(t)?o=t:g?r=t:s=t;break;case 0:break;default:throw M("badargs","Expected up to 4 arguments [params, data, success, error], got {0} arguments",arguments.length)}var l=this instanceof y,u=l?r:f.isArray?[]:new y(r),c={},h=f.interceptor&&f.interceptor.response||m,d=f.interceptor&&f.interceptor.responseError||D;k(f,function(t,e){"params"!=e&&"isArray"!=e&&"interceptor"!=e&&(c[e]=E(t))}),g&&(c.data=r),b.setUrlParams(c,C({},function(n,t){var i={};return t=C({},v,t),k(t,function(t,e){S(t)&&(t=t()),i[e]=t&&t.charAt&&"@"==t.charAt(0)?_(n,t.substr(1)):t}),i}(r,f.params||{}),s),f.url);var p=x(c).then(function(t){var e=t.data,n=u.$promise;if(e){if(T.isArray(e)!==!!f.isArray)throw M("badcfg","Error in resource configuration. Expected response to contain an {0} but got an {1}",f.isArray?"array":"object",T.isArray(e)?"array":"object");f.isArray?(u.length=0,k(e,function(t){"object"==typeof t?u.push(new y(t)):u.push(t)})):(P(e,u),u.$promise=n)}return u.$resolved=!0,t.resource=u,t},function(t){return u.$resolved=!0,(a||$)(t),w.reject(t)});return p=p.then(function(t){var e=h(t);return(o||$)(e,t.headers),e},d),l?p:(u.$promise=p,u.$resolved=!1,u)},y.prototype["$"+r]=function(t,e,n){S(t)&&(n=e,e=t,t={});var i=y[r].call(this,t,this,e,n);return i.$promise||i}}),y.bind=function(t){return e(n,C({},v,t),i)},y}}])}(window,window.angular),function(t,v){"use strict";function e(p,f,g){return{restrict:"ECA",terminal:!0,priority:400,transclude:"element",link:function(o,a,t,e,s){function l(){i&&(i.remove(),i=null),u&&(u.$destroy(),u=null),c&&(g.leave(c,function(){i=null}),i=c,c=null)}function n(){var t=p.current&&p.current.locals,e=t&&t.$template;if(v.isDefined(e)){var n=o.$new(),i=p.current,r=s(n,function(t){g.enter(t,null,c||a,function(){!v.isDefined(h)||h&&!o.$eval(h)||f()}),l()});c=r,(u=i.scope=n).$emit("$viewContentLoaded"),u.$eval(d)}else l()}var u,c,i,h=t.autoscroll,d=t.onload||"";o.$on("$routeChangeSuccess",n),n()}}}function n(a,s,l){return{restrict:"ECA",priority:-400,link:function(t,e){var n=l.current,i=n.locals;e.html(i.$template);var r=a(e.contents());if(n.controller){i.$scope=t;var o=s(n.controller,i);n.controllerAs&&(t[n.controllerAs]=o),e.data("$ngControllerController",o),e.children().data("$ngControllerController",o)}r(t)}}}var i=v.module("ngRoute",["ng"]).provider("$route",function(){function p(t,e){return v.extend(new(v.extend(function(){},{prototype:t})),e)}function i(t,e){var n=e.caseInsensitiveMatch,i={originalPath:t,regexp:t},a=i.keys=[];return t=t.replace(/([().])/g,"\\$1").replace(/(\/)?:(\w+)([\?\*])?/g,function(t,e,n,i){var r="?"===i?i:null,o="*"===i?i:null;return a.push({name:n,optional:!!r}),e=e||"",(r?"":e)+"(?:"+(r?e:"")+(o?"(.+?)":"([^/]+)")+(r||"")+")"+(r||"")}).replace(/([\/$\*])/g,"\\$1"),i.regexp=new RegExp("^"+t+"$",n?"i":""),i}var f={};this.when=function(t,e){if(f[t]=v.extend({reloadOnSearch:!0},e,t&&i(t,e)),t){var n="/"==t[t.length-1]?t.substr(0,t.length-1):t+"/";f[n]=v.extend({redirectTo:t},i(n,e))}return this},this.otherwise=function(t){return this.when(null,t),this},this.$get=["$rootScope","$location","$routeParams","$q","$injector","$http","$templateCache","$sce",function(n,r,o,a,s,l,u,c){function t(){var i=function(){var n,i;return v.forEach(f,function(t,e){!i&&(n=function(t,e){var n=e.keys,i={};if(!e.regexp)return null;var r=e.regexp.exec(t);if(!r)return null;for(var o=1,a=r.length;o<a;++o){var s=n[o-1],l=r[o];s&&l&&(i[s.name]=l)}return i}(r.path(),t))&&((i=p(t,{params:v.extend({},r.search(),n),pathParams:n})).$$route=t)}),i||f.null&&p(f.null,{params:{},pathParams:{}})}(),e=d.current;i&&e&&i.$$route===e.$$route&&v.equals(i.pathParams,e.pathParams)&&!i.reloadOnSearch&&!h?(e.params=i.params,v.copy(e.params,o),n.$broadcast("$routeUpdate",e)):(i||e)&&(h=!1,n.$broadcast("$routeChangeStart",i,e),(d.current=i)&&i.redirectTo&&(v.isString(i.redirectTo)?r.path(function(t,r){var o=[];return v.forEach((t||"").split(":"),function(t,e){if(0===e)o.push(t);else{var n=t.match(/(\w+)(?:[?*])?(.*)/),i=n[1];o.push(r[i]),o.push(n[2]||""),delete r[i]}}),o.join("")}(i.redirectTo,i.params)).search(i.params).replace():r.url(i.redirectTo(i.pathParams,r.path(),r.search())).replace()),a.when(i).then(function(){if(i){var t,e,n=v.extend({},i.resolve);return v.forEach(n,function(t,e){n[e]=v.isString(t)?s.get(t):s.invoke(t)}),v.isDefined(t=i.template)?v.isFunction(t)&&(t=t(i.params)):v.isDefined(e=i.templateUrl)&&(v.isFunction(e)&&(e=e(i.params)),e=c.getTrustedResourceUrl(e),v.isDefined(e)&&(i.loadedTemplateUrl=e,t=l.get(e,{cache:u}).then(function(t){return t.data}))),v.isDefined(t)&&(n.$template=t),a.all(n)}}).then(function(t){i==d.current&&(i&&(i.locals=t,v.copy(i.params,o)),n.$broadcast("$routeChangeSuccess",i,e))},function(t){i==d.current&&n.$broadcast("$routeChangeError",i,e,t)}))}var h=!1,d={routes:f,reload:function(){h=!0,n.$evalAsync(t)}};return n.$on("$locationChangeSuccess",t),d}]});i.provider("$routeParams",function(){this.$get=function(){return{}}}),i.directive("ngView",e),i.directive("ngView",n),e.$inject=["$route","$anchorScroll","$animate"],n.$inject=["$compile","$controller","$route"]}(window,window.angular),function(t,p){"use strict";function e(t){var e,n={},i=t.split(",");for(e=0;e<i.length;e++)n[i[e]]=!0;return n}function c(t){if(!t)return"";var e=D.exec(t),n=e[1],i=e[3],r=e[2];return r&&(a.innerHTML=r.replace(/</g,"&lt;"),r="textContent"in a?a.textContent:a.innerText),n+r+i}function s(t){return t.replace(/&/g,"&amp;").replace(n,function(t){return"&#"+(1024*(t.charCodeAt(0)-55296)+(t.charCodeAt(1)-56320)+65536)+";"}).replace(i,function(t){return"&#"+t.charCodeAt(0)+";"}).replace(/</g,"&lt;").replace(/>/g,"&gt;")}function f(t,o){var n=!1,a=p.bind(t,t.push);return{start:function(r,t,e){r=p.lowercase(r),!n&&S[r]&&(n=r),n||!0!==l[r]||(a("<"),a(r),p.forEach(t,function(t,e){var n=p.lowercase(e),i="img"===r&&"src"===n||"background"===n;!0!==T[n]||!0===u[n]&&!o(t,i)||(a(" "),a(e),a('="'),a(s(t)),a('"'))}),a(e?"/>":">"))},end:function(t){t=p.lowercase(t),n||!0!==l[t]||(a("</"),a(t),a(">")),t==n&&(n=!1)},chars:function(t){n||a(s(t))}}}var h=p.$$minErr("$sanitize"),d=/^<((?:[a-zA-Z])[\w:-]*)((?:\s+[\w:-]+(?:\s*=\s*(?:(?:"[^"]*")|(?:'[^']*')|[^>\s]+))?)*)\s*(\/?)\s*(>?)/,g=/^<\/\s*([\w:-]+)[^>]*>/,v=/([\w:-]+)(?:\s*=\s*(?:(?:"((?:[^"])*)")|(?:'((?:[^'])*)')|([^>\s]+)))?/g,m=/^</,y=/^<\//,b=/<!--(.*?)-->/g,x=/<!DOCTYPE([^>]*?)>/i,w=/<!\[CDATA\[(.*?)]]>/g,n=/[\uD800-\uDBFF][\uDC00-\uDFFF]/g,i=/([^\#-~| |!])/g,$=e("area,br,col,hr,img,wbr"),r=e("colgroup,dd,dt,li,p,tbody,td,tfoot,th,thead,tr"),o=e("rp,rt"),k=p.extend({},o,r),C=p.extend({},r,e("address,article,aside,blockquote,caption,center,del,dir,div,dl,figure,figcaption,footer,h1,h2,h3,h4,h5,h6,header,hgroup,hr,ins,map,menu,nav,ol,pre,script,section,table,ul")),E=p.extend({},o,e("a,abbr,acronym,b,bdi,bdo,big,br,cite,code,del,dfn,em,font,i,img,ins,kbd,label,map,mark,q,ruby,rp,rt,s,samp,small,span,strike,strong,sub,sup,time,tt,u,var")),S=e("script,style"),l=p.extend({},$,C,E,k),u=e("background,cite,href,longdesc,src,usemap"),T=p.extend({},u,e("abbr,align,alt,axis,bgcolor,border,cellpadding,cellspacing,class,clear,color,cols,colspan,compact,coords,dir,face,headers,height,hreflang,hspace,ismap,lang,language,nohref,nowrap,rel,rev,rows,rowspan,rules,scope,scrolling,shape,size,span,start,summary,target,title,type,valign,value,vspace,width")),a=document.createElement("pre"),D=/^(\s*)([\s\S]*?)(\s*)$/;p.module("ngSanitize",[]).provider("$sanitize",function(){this.$get=["$$sanitizeUri",function(n){return function(t){var e=[];return function(t,r){function e(t,e,n,i){if(e=p.lowercase(e),C[e])for(;l.last()&&E[l.last()];)o("",l.last());k[e]&&l.last()==e&&o("",e),(i=$[e]||!!i)||l.push(e);var a={};n.replace(v,function(t,e,n,i,r){var o=n||i||r||"";a[e]=c(o)}),r.start&&r.start(e,a,i)}function o(t,e){var n,i=0;if(e=p.lowercase(e))for(i=l.length-1;0<=i&&l[i]!=e;i--);if(0<=i){for(n=l.length-1;i<=n;n--)r.end&&r.end(l[n]);l.length=i}}"string"!=typeof t&&(t=null==t?"":""+t);var n,i,a,s,l=[],u=t;for(l.last=function(){return l[l.length-1]};t;){if(i=!(s=""),l.last()&&S[l.last()]?(t=t.replace(new RegExp("(.*)<\\s*\\/\\s*"+l.last()+"[^>]*>","i"),function(t,e){return e=e.replace(b,"$1").replace(w,"$1"),r.chars&&r.chars(c(e)),""}),o(0,l.last())):(0===t.indexOf("\x3c!--")?0<=(n=t.indexOf("--",4))&&t.lastIndexOf("--\x3e",n)===n&&(r.comment&&r.comment(t.substring(4,n)),t=t.substring(n+3),i=!1):x.test(t)?(a=t.match(x))&&(t=t.replace(a[0],""),i=!1):y.test(t)?(a=t.match(g))&&(t=t.substring(a[0].length),a[0].replace(g,o),i=!1):m.test(t)&&((a=t.match(d))?(a[4]&&(t=t.substring(a[0].length),a[0].replace(d,e)),i=!1):(s+="<",t=t.substring(1))),i&&(n=t.indexOf("<"),s+=n<0?t:t.substring(0,n),t=n<0?"":t.substring(n),r.chars&&r.chars(c(s)))),t==u)throw h("badparse","The sanitizer was unable to parse the following block of html: {0}",t);u=t}o()}(t,f(e,function(t,e){return!/^unsafe/.test(n(t,e))})),e.join("")}}]}),p.module("ngSanitize").filter("linky",["$sanitize",function(c){var h=/((ftp|https?):\/\/|(mailto:)?[A-Za-z0-9._%+-]+@)\S*[^\s.;,(){}<>"]/,d=/^mailto:/;return function(t,e){function n(t){t&&s.push(function(t){var e=[];return f(e,p.noop).chars(t),e.join("")}(t))}if(!t)return t;for(var i,r,o,a=t,s=[];i=a.match(h);)r=i[0],i[2]==i[3]&&(r="mailto:"+r),o=i.index,n(a.substr(0,o)),l=r,u=i[0].replace(d,""),s.push("<a "),p.isDefined(e)&&(s.push('target="'),s.push(e),s.push('" ')),s.push('href="',l.replace('"',"&quot;"),'">'),n(u),s.push("</a>"),a=a.substring(o+i[0].length);var l,u;return n(a),c(s.join(""))}}])}(window,window.angular),function(t,e){"object"==typeof exports?module.exports=e():"function"==typeof define&&define.amd?define(e):t.Spinner=e()}(this,function(){"use strict";function g(t,e){var n,i=document.createElement(t||"div");for(n in e)i[n]=e[n];return i}function v(t){for(var e=1,n=arguments.length;e<n;e++)t.appendChild(arguments[e]);return t}function i(t,e){var n,i,r=t.style;for(e=e.charAt(0).toUpperCase()+e.slice(1),i=0;i<o.length;i++)if(void 0!==r[n=o[i]+e])return n;return void 0!==r[e]?e:void 0}function m(t,e){for(var n in e)t.style[i(t,n)||n]=e[n];return t}function e(t){for(var e=1;e<arguments.length;e++){var n=arguments[e];for(var i in n)void 0===t[i]&&(t[i]=n[i])}return t}function y(t,e){return"string"==typeof t?t:t[e%t.length]}function n(t){this.opts=e(t||{},n.defaults,r)}var b,t,o=["webkit","Moz","ms","O"],x={},w=(t=g("style",{type:"text/css"}),v(document.getElementsByTagName("head")[0],t),t.sheet||t.styleSheet),r={lines:12,length:7,width:5,radius:10,rotate:0,corners:1,color:"#000",direction:1,speed:1,trail:100,opacity:.25,fps:20,zIndex:2e9,className:"spinner",top:"50%",left:"50%",position:"absolute"};n.defaults={},e(n.prototype,{spin:function(t){this.stop();var n=this,i=n.opts,r=n.el=m(g(0,{className:i.className}),{position:i.position,width:0,zIndex:i.zIndex});if(m(r,{left:i.left,top:i.top}),t&&t.insertBefore(r,t.firstChild||null),r.setAttribute("role","progressbar"),n.lines(r,n.opts),!b){var o,a=0,s=(i.lines-1)*(1-i.direction)/2,l=i.fps,u=l/i.speed,c=(1-i.opacity)/(u*i.trail/100),h=u/i.lines;!function t(){a++;for(var e=0;e<i.lines;e++)o=Math.max(1-(a+(i.lines-e)*h)%u*c,i.opacity),n.opacity(r,e*i.direction+s,o,i);n.timeout=n.el&&setTimeout(t,~~(1e3/l))}()}return n},stop:function(){var t=this.el;return t&&(clearTimeout(this.timeout),t.parentNode&&t.parentNode.removeChild(t),this.el=void 0),this},lines:function(t,n){function e(t,e){return m(g(),{position:"absolute",width:n.length+n.width+"px",height:n.width+"px",background:t,boxShadow:e,transformOrigin:"left",transform:"rotate("+~~(360/n.lines*r+n.rotate)+"deg) translate("+n.radius+"px,0)",borderRadius:(n.corners*n.width>>1)+"px"})}for(var i,r=0,o=(n.lines-1)*(1-n.direction)/2;r<n.lines;r++)i=m(g(),{position:"absolute",top:1+~(n.width/2)+"px",transform:n.hwaccel?"translate3d(0,0,0)":"",opacity:n.opacity,animation:b&&(a=n.opacity,s=n.trail,l=o+r*n.direction,u=n.lines,void 0,void 0,void 0,void 0,void 0,c=["opacity",s,~~(100*a),l,u].join("-"),h=.01+l/u*100,d=Math.max(1-(1-a)/s*(100-h),a),p=b.substring(0,b.indexOf("Animation")).toLowerCase(),f=p&&"-"+p+"-"||"",x[c]||(w.insertRule("@"+f+"keyframes "+c+"{0%{opacity:"+d+"}"+h+"%{opacity:"+a+"}"+(.01+h)+"%{opacity:1}"+(h+s)%100+"%{opacity:"+a+"}100%{opacity:"+d+"}}",w.cssRules.length),x[c]=1),c+" "+1/n.speed+"s linear infinite")}),n.shadow&&v(i,m(e("#000","0 0 4px #000"),{top:"2px"})),v(t,v(i,e(y(n.color,r),"0 0 1px rgba(0,0,0,.1)")));var a,s,l,u,c,h,d,p,f;return t},opacity:function(t,e,n){e<t.childNodes.length&&(t.childNodes[e].style.opacity=n)}});var a=m(g("group"),{behavior:"url(#default#VML)"});return!i(a,"transform")&&a.adj?(w.addRule(".spin-vml","behavior:url(#default#VML)"),n.prototype.lines=function(t,i){function r(){return m(u("group",{coordsize:a+" "+a,coordorigin:-o+" "+-o}),{width:a,height:a})}function e(t,e,n){v(l,v(m(r(),{rotation:360/i.lines*t+"deg",left:~~e}),v(m(u("roundrect",{arcsize:i.corners}),{width:o,height:i.width,left:i.radius,top:-i.width>>1,filter:n}),u("fill",{color:y(i.color,t),opacity:i.opacity}),u("stroke",{opacity:0}))))}var n,o=i.length+i.width,a=2*o,s=2*-(i.width+i.length)+"px",l=m(r(),{position:"absolute",top:s,left:s});if(i.shadow)for(n=1;n<=i.lines;n++)e(n,-2,"progid:DXImageTransform.Microsoft.Blur(pixelradius=2,makeshadow=1,shadowopacity=.3)");for(n=1;n<=i.lines;n++)e(n);return v(t,l)},n.prototype.opacity=function(t,e,n,i){var r=t.firstChild;i=i.shadow&&i.lines||0,r&&e+i<r.childNodes.length&&(r=(r=(r=r.childNodes[e+i])&&r.firstChild)&&r.firstChild)&&(r.opacity=n)}):b=i(a,"animation"),n;function u(t,e){return g("<"+t+' xmlns="urn:schemas-microsoft.com:vml" class="spin-vml">',e)}}),function(t){"use strict";function e(a,s){a.module("angularSpinner",[]).factory("usSpinnerService",["$rootScope",function(e){var t={spin:function(t){e.$broadcast("us-spinner:spin",t)},stop:function(t){e.$broadcast("us-spinner:stop",t)}};return t}]).directive("usSpinner",["$window",function(o){return{scope:!0,link:function(n,e,t){function i(){n.spinner&&n.spinner.stop()}var r=s||o.Spinner;n.spinner=null,n.key=!!a.isDefined(t.spinnerKey)&&t.spinnerKey,n.startActive=a.isDefined(t.spinnerStartActive)?t.spinnerStartActive:!n.key,n.spin=function(){n.spinner&&n.spinner.spin(e[0])},n.stop=function(){n.startActive=!1,i()},n.$watch(t.usSpinner,function(t){i(),n.spinner=new r(t),n.key&&!n.startActive||n.spinner.spin(e[0])},!0),n.$on("us-spinner:spin",function(t,e){e===n.key&&n.spin()}),n.$on("us-spinner:stop",function(t,e){e===n.key&&n.stop()}),n.$on("$destroy",function(){n.stop(),n.spinner=null})}}}])}"function"==typeof define&&define.amd?define(["angular","spin"],e):e(t.angular)}(window),function(i,r){"use strict";function n(e){return function(t){return t[e]}}var o=i.module("angular-underscore",[]),t=i.module("angular-underscore/utils",[]),a=i.module("angular-underscore/filters",[]);(r._=r).each(["min","max","sortedIndex"],function(t){r[t]=r.wrap(r[t],function(t){var e=r.toArray(arguments).slice(1);return r.isString(e[2])?e[2]=n(e[2]):r.isString(e[1])&&(e[1]=n(e[1])),t.apply(r,e)})}),i.injector(["ng"]).invoke(["$filter",function(t){r.filter=r.select=r.wrap(t("filter"),function(t,e,n,i){return r.isArray(e)||(e=r.toArray(e)),t(e,n,i)}),r.reject=function(t,e){return r.isString(e)?r.filter(t,"!"+e):r.bind(r.difference,r,t)(r.filter(t,e))}}]),r.each(r.methods(r),function(e){function n(t){t[e]=r.bind(r[e],r)}r.each([o,t,i.module("angular-underscore/utils/"+e,[])],function(t){t.run(["$rootScope",n])})});r.each([["map","collect"],["reduce","inject","foldl"],["reduceRight","foldr"],["find","detect"],["filter","select"],"where","findWhere","reject","invoke","pluck","max","min","sortBy","groupBy","indexBy","countBy","shuffle","sample","toArray","size",["first","head","take"],"initial","last",["rest","tail","drop"],"compact","flatten","without","partition","union","intersection","difference",["uniq","unique"],"zip","object","indexOf","lastIndexOf","sortedIndex","keys","values","pairs","invert",["functions","methods"],"pick","omit","tap","identity","uniqueId","escape","unescape","result","template"],function(t){r.isArray(t)||(t=[t]);function n(){return e}var e=r.bind(r[t[0]],r);r.each(t,function(e){r.each([o,a,i.module("angular-underscore/filters/"+e,[])],function(t){t.filter(e,n)})})})}(angular,_),"undefined"==typeof jQuery)throw new Error("Bootstrap requires jQuery");var cola;!function(i){"use strict";i.fn.emulateTransitionEnd=function(t){var e=!1,n=this;i(this).one(i.support.transition.end,function(){e=!0});return setTimeout(function(){e||i(n).trigger(i.support.transition.end)},t),this},i(function(){i.support.transition=function(){var t=document.createElement("bootstrap"),e={WebkitTransition:"webkitTransitionEnd",MozTransition:"transitionend",OTransition:"oTransitionEnd otransitionend",transition:"transitionend"};for(var n in e)if(void 0!==t.style[n])return{end:e[n]}}()})}(jQuery),function(o){"use strict";function i(t){o(t).on("click",e,this.close)}var e='[data-dismiss="alert"]';i.prototype.close=function(t){function e(){r.trigger("closed.bs.alert").remove()}var n=o(this),i=n.attr("data-target");i||(i=(i=n.attr("href"))&&i.replace(/.*(?=#[^\s]*$)/,""));var r=o(i);t&&t.preventDefault(),r.length||(r=n.hasClass("alert")?n:n.parent()),r.trigger(t=o.Event("close.bs.alert")),t.isDefaultPrevented()||(r.removeClass("in"),o.support.transition&&r.hasClass("fade")?r.one(o.support.transition.end,e).emulateTransitionEnd(150):e())};var t=o.fn.alert;o.fn.alert=function(n){return this.each(function(){var t=o(this),e=t.data("bs.alert");e||t.data("bs.alert",e=new i(this)),"string"==typeof n&&e[n].call(t)})},o.fn.alert.Constructor=i,o.fn.alert.noConflict=function(){return o.fn.alert=t,this},o(document).on("click.bs.alert.data-api",e,i.prototype.close)}(jQuery),function(r){"use strict";var o=function(t,e){this.$element=r(t),this.options=r.extend({},o.DEFAULTS,e)};o.DEFAULTS={loadingText:"loading..."},o.prototype.setState=function(t){var e="disabled",n=this.$element,i=n.is("input")?"val":"html",r=n.data();t+="Text",r.resetText||n.data("resetText",n[i]()),n[i](r[t]||this.options[t]),setTimeout(function(){"loadingText"==t?n.addClass(e).attr(e,e):n.removeClass(e).removeAttr(e)},0)},o.prototype.toggle=function(){var t=this.$element.closest('[data-toggle="buttons"]'),e=!0;if(t.length){var n=this.$element.find("input");"radio"===n.prop("type")&&(n.prop("checked")&&this.$element.hasClass("active")?e=!1:t.find(".active").removeClass("active")),e&&n.prop("checked",!this.$element.hasClass("active")).trigger("change")}e&&this.$element.toggleClass("active")};var t=r.fn.button;r.fn.button=function(i){return this.each(function(){var t=r(this),e=t.data("bs.button"),n="object"==typeof i&&i;e||t.data("bs.button",e=new o(this,n)),"toggle"==i?e.toggle():i&&e.setState(i)})},r.fn.button.Constructor=o,r.fn.button.noConflict=function(){return r.fn.button=t,this},r(document).on("click.bs.button.data-api","[data-toggle^=button]",function(t){var e=r(t.target);e.hasClass("btn")||(e=e.closest(".btn")),e.button("toggle"),t.preventDefault()})}(jQuery),function(u){"use strict";function o(t,e){this.$element=u(t),this.$indicators=this.$element.find(".carousel-indicators"),this.options=e,this.paused=this.sliding=this.interval=this.$active=this.$items=null,"hover"==this.options.pause&&this.$element.on("mouseenter",u.proxy(this.pause,this)).on("mouseleave",u.proxy(this.cycle,this))}o.DEFAULTS={interval:5e3,pause:"hover",wrap:!0},o.prototype.cycle=function(t){return t||(this.paused=!1),this.interval&&clearInterval(this.interval),this.options.interval&&!this.paused&&(this.interval=setInterval(u.proxy(this.next,this),this.options.interval)),this},o.prototype.getActiveIndex=function(){return this.$active=this.$element.find(".item.active"),this.$items=this.$active.parent().children(),this.$items.index(this.$active)},o.prototype.to=function(t){var e=this,n=this.getActiveIndex();return t>this.$items.length-1||t<0?void 0:this.sliding?this.$element.one("slid.bs.carousel",function(){e.to(t)}):n==t?this.pause().cycle():this.slide(n<t?"next":"prev",u(this.$items[t]))},o.prototype.pause=function(t){return t||(this.paused=!0),this.$element.find(".next, .prev").length&&u.support.transition.end&&(this.$element.trigger(u.support.transition.end),this.cycle(!0)),this.interval=clearInterval(this.interval),this},o.prototype.next=function(){return this.sliding?void 0:this.slide("next")},o.prototype.prev=function(){return this.sliding?void 0:this.slide("prev")},o.prototype.slide=function(t,e){var n=this.$element.find(".item.active"),i=e||n[t](),r=this.interval,o="next"==t?"left":"right",a="next"==t?"first":"last",s=this;if(!i.length){if(!this.options.wrap)return;i=this.$element.find(".item")[a]()}this.sliding=!0,r&&this.pause();var l=u.Event("slide.bs.carousel",{relatedTarget:i[0],direction:o});if(!i.hasClass("active")){if(this.$indicators.length&&(this.$indicators.find(".active").removeClass("active"),this.$element.one("slid.bs.carousel",function(){var t=u(s.$indicators.children()[s.getActiveIndex()]);t&&t.addClass("active")})),u.support.transition&&this.$element.hasClass("slide")){if(this.$element.trigger(l),l.isDefaultPrevented())return;i.addClass(t),i[0].offsetWidth,n.addClass(o),i.addClass(o),n.one(u.support.transition.end,function(){i.removeClass([t,o].join(" ")).addClass("active"),n.removeClass(["active",o].join(" ")),s.sliding=!1,setTimeout(function(){s.$element.trigger("slid.bs.carousel")},0)}).emulateTransitionEnd(600)}else{if(this.$element.trigger(l),l.isDefaultPrevented())return;n.removeClass("active"),i.addClass("active"),this.sliding=!1,this.$element.trigger("slid.bs.carousel")}return r&&this.cycle(),this}};var t=u.fn.carousel;u.fn.carousel=function(r){return this.each(function(){var t=u(this),e=t.data("bs.carousel"),n=u.extend({},o.DEFAULTS,t.data(),"object"==typeof r&&r),i="string"==typeof r?r:n.slide;e||t.data("bs.carousel",e=new o(this,n)),"number"==typeof r?e.to(r):i?e[i]():n.interval&&e.pause().cycle()})},u.fn.carousel.Constructor=o,u.fn.carousel.noConflict=function(){return u.fn.carousel=t,this},u(document).on("click.bs.carousel.data-api","[data-slide], [data-slide-to]",function(t){var e,n=u(this),i=u(n.attr("data-target")||(e=n.attr("href"))&&e.replace(/.*(?=#[^\s]+$)/,"")),r=u.extend({},i.data(),n.data()),o=n.attr("data-slide-to");o&&(r.interval=!1),i.carousel(r),(o=n.attr("data-slide-to"))&&i.data("bs.carousel").to(o),t.preventDefault()}),u(window).on("load",function(){u('[data-ride="carousel"]').each(function(){var t=u(this);t.carousel(t.data())})})}(jQuery),function(u){"use strict";var r=function(t,e){this.$element=u(t),this.options=u.extend({},r.DEFAULTS,e),this.transitioning=null,this.options.parent&&(this.$parent=u(this.options.parent)),this.options.toggle&&this.toggle()};r.DEFAULTS={toggle:!0},r.prototype.dimension=function(){return this.$element.hasClass("width")?"width":"height"},r.prototype.show=function(){if(!this.transitioning&&!this.$element.hasClass("in")){var t=u.Event("show.bs.collapse");if(this.$element.trigger(t),!t.isDefaultPrevented()){var e=this.$parent&&this.$parent.find("> .panel > .in");if(e&&e.length){var n=e.data("bs.collapse");if(n&&n.transitioning)return;e.collapse("hide"),n||e.data("bs.collapse",null)}var i=this.dimension();this.$element.removeClass("collapse").addClass("collapsing")[i](0),this.transitioning=1;var r=function(){this.$element.removeClass("collapsing").addClass("in")[i]("auto"),this.transitioning=0,this.$element.trigger("shown.bs.collapse")};if(!u.support.transition)return r.call(this);var o=u.camelCase(["scroll",i].join("-"));this.$element.one(u.support.transition.end,u.proxy(r,this)).emulateTransitionEnd(350)[i](this.$element[0][o])}}},r.prototype.hide=function(){if(!this.transitioning&&this.$element.hasClass("in")){var t=u.Event("hide.bs.collapse");if(this.$element.trigger(t),!t.isDefaultPrevented()){var e=this.dimension();this.$element[e](this.$element[e]())[0].offsetHeight,this.$element.addClass("collapsing").removeClass("collapse").removeClass("in"),this.transitioning=1;var n=function(){this.transitioning=0,this.$element.trigger("hidden.bs.collapse").removeClass("collapsing").addClass("collapse")};return u.support.transition?void this.$element[e](0).one(u.support.transition.end,u.proxy(n,this)).emulateTransitionEnd(350):n.call(this)}}},r.prototype.toggle=function(){this[this.$element.hasClass("in")?"hide":"show"]()};var t=u.fn.collapse;u.fn.collapse=function(i){return this.each(function(){var t=u(this),e=t.data("bs.collapse"),n=u.extend({},r.DEFAULTS,t.data(),"object"==typeof i&&i);e||t.data("bs.collapse",e=new r(this,n)),"string"==typeof i&&e[i]()})},u.fn.collapse.Constructor=r,u.fn.collapse.noConflict=function(){return u.fn.collapse=t,this},u(document).on("click.bs.collapse.data-api","[data-toggle=collapse]",function(t){var e,n=u(this),i=n.attr("data-target")||t.preventDefault()||(e=n.attr("href"))&&e.replace(/.*(?=#[^\s]+$)/,""),r=u(i),o=r.data("bs.collapse"),a=o?"toggle":n.data(),s=n.attr("data-parent"),l=s&&u(s);o&&o.transitioning||(l&&l.find('[data-toggle=collapse][data-parent="'+s+'"]').not(n).addClass("collapsed"),n[r.hasClass("in")?"addClass":"removeClass"]("collapsed")),r.collapse(a)})}(jQuery),function(a){"use strict";function r(){a(".dropdown-backdrop").remove(),a(l).each(function(t){var e=s(a(this));e.hasClass("open")&&(e.trigger(t=a.Event("hide.bs.dropdown")),t.isDefaultPrevented()||e.removeClass("open").trigger("hidden.bs.dropdown"))})}function s(t){var e=t.attr("data-target");e||(e=(e=t.attr("href"))&&/#/.test(e)&&e.replace(/.*(?=#[^\s]*$)/,""));var n=e&&a(e);return n&&n.length?n:t.parent()}function i(t){a(t).on("click.bs.dropdown",this.toggle)}var l="[data-toggle=dropdown]";i.prototype.toggle=function(t){var e=a(this);if(!e.is(".disabled, :disabled")){var n=s(e),i=n.hasClass("open");if(r(),!i){if("ontouchstart"in document.documentElement&&!n.closest(".navbar-nav").length&&a('<div class="dropdown-backdrop"/>').insertAfter(a(this)).on("click",r),n.trigger(t=a.Event("show.bs.dropdown")),t.isDefaultPrevented())return;n.toggleClass("open").trigger("shown.bs.dropdown"),e.focus()}return!1}},i.prototype.keydown=function(t){if(/(38|40|27)/.test(t.keyCode)){var e=a(this);if(t.preventDefault(),t.stopPropagation(),!e.is(".disabled, :disabled")){var n=s(e),i=n.hasClass("open");if(!i||i&&27==t.keyCode)return 27==t.which&&n.find(l).focus(),e.click();var r=a("[role=menu] li:not(.divider):visible a",n);if(r.length){var o=r.index(r.filter(":focus"));38==t.keyCode&&0<o&&o--,40==t.keyCode&&o<r.length-1&&o++,~o||(o=0),r.eq(o).focus()}}}};var t=a.fn.dropdown;a.fn.dropdown=function(n){return this.each(function(){var t=a(this),e=t.data("bs.dropdown");e||t.data("bs.dropdown",e=new i(this)),"string"==typeof n&&e[n].call(t)})},a.fn.dropdown.Constructor=i,a.fn.dropdown.noConflict=function(){return a.fn.dropdown=t,this},a(document).on("click.bs.dropdown.data-api",r).on("click.bs.dropdown.data-api",".dropdown form",function(t){t.stopPropagation()}).on("click.bs.dropdown.data-api",l,i.prototype.toggle).on("keydown.bs.dropdown.data-api",l+", [role=menu]",i.prototype.keydown)}(jQuery),function(o){"use strict";function a(t,e){this.options=e,this.$element=o(t),this.$backdrop=this.isShown=null,this.options.remote&&this.$element.load(this.options.remote)}a.DEFAULTS={backdrop:!0,keyboard:!0,show:!0},a.prototype.toggle=function(t){return this[this.isShown?"hide":"show"](t)},a.prototype.show=function(n){var i=this,t=o.Event("show.bs.modal",{relatedTarget:n});this.$element.trigger(t),this.isShown||t.isDefaultPrevented()||(this.isShown=!0,this.escape(),this.$element.on("click.dismiss.modal",'[data-dismiss="modal"]',o.proxy(this.hide,this)),this.backdrop(function(){var t=o.support.transition&&i.$element.hasClass("fade");i.$element.parent().length||i.$element.appendTo(document.body),i.$element.show(),t&&i.$element[0].offsetWidth,i.$element.addClass("in").attr("aria-hidden",!1),i.enforceFocus();var e=o.Event("shown.bs.modal",{relatedTarget:n});t?i.$element.find(".modal-dialog").one(o.support.transition.end,function(){i.$element.focus().trigger(e)}).emulateTransitionEnd(300):i.$element.focus().trigger(e)}))},a.prototype.hide=function(t){t&&t.preventDefault(),t=o.Event("hide.bs.modal"),this.$element.trigger(t),this.isShown&&!t.isDefaultPrevented()&&(this.isShown=!1,this.escape(),o(document).off("focusin.bs.modal"),this.$element.removeClass("in").attr("aria-hidden",!0).off("click.dismiss.modal"),o.support.transition&&this.$element.hasClass("fade")?this.$element.one(o.support.transition.end,o.proxy(this.hideModal,this)).emulateTransitionEnd(300):this.hideModal())},a.prototype.enforceFocus=function(){o(document).off("focusin.bs.modal").on("focusin.bs.modal",o.proxy(function(t){this.$element[0]===t.target||this.$element.has(t.target).length||this.$element.focus()},this))},a.prototype.escape=function(){this.isShown&&this.options.keyboard?this.$element.on("keyup.dismiss.bs.modal",o.proxy(function(t){27==t.which&&this.hide()},this)):this.isShown||this.$element.off("keyup.dismiss.bs.modal")},a.prototype.hideModal=function(){var t=this;this.$element.hide(),this.backdrop(function(){t.removeBackdrop(),t.$element.trigger("hidden.bs.modal")})},a.prototype.removeBackdrop=function(){this.$backdrop&&this.$backdrop.remove(),this.$backdrop=null},a.prototype.backdrop=function(t){var e=this.$element.hasClass("fade")?"fade":"";if(this.isShown&&this.options.backdrop){var n=o.support.transition&&e;if(this.$backdrop=o('<div class="modal-backdrop '+e+'" />').appendTo(document.body),this.$element.on("click.dismiss.modal",o.proxy(function(t){t.target===t.currentTarget&&("static"==this.options.backdrop?this.$element[0].focus.call(this.$element[0]):this.hide.call(this))},this)),n&&this.$backdrop[0].offsetWidth,this.$backdrop.addClass("in"),!t)return;n?this.$backdrop.one(o.support.transition.end,t).emulateTransitionEnd(150):t()}else!this.isShown&&this.$backdrop?(this.$backdrop.removeClass("in"),o.support.transition&&this.$element.hasClass("fade")?this.$backdrop.one(o.support.transition.end,t).emulateTransitionEnd(150):t()):t&&t()};var t=o.fn.modal;o.fn.modal=function(i,r){return this.each(function(){var t=o(this),e=t.data("bs.modal"),n=o.extend({},a.DEFAULTS,t.data(),"object"==typeof i&&i);e||t.data("bs.modal",e=new a(this,n)),"string"==typeof i?e[i](r):n.show&&e.show(r)})},o.fn.modal.Constructor=a,o.fn.modal.noConflict=function(){return o.fn.modal=t,this},o(document).on("click.bs.modal.data-api",'[data-toggle="modal"]',function(t){var e=o(this),n=e.attr("href"),i=o(e.attr("data-target")||n&&n.replace(/.*(?=#[^\s]+$)/,"")),r=i.data("modal")?"toggle":o.extend({remote:!/#/.test(n)&&n},i.data(),e.data());t.preventDefault(),i.modal(r,this).one("hide",function(){e.is(":visible")&&e.focus()})}),o(document).on("show.bs.modal",".modal",function(){o(document.body).addClass("modal-open")}).on("hidden.bs.modal",".modal",function(){o(document.body).removeClass("modal-open")})}(jQuery),function(g){"use strict";function r(t,e){this.type=this.options=this.enabled=this.timeout=this.hoverState=this.$element=null,this.init("tooltip",t,e)}r.DEFAULTS={animation:!0,placement:"top",selector:!1,template:'<div class="tooltip"><div class="tooltip-arrow"></div><div class="tooltip-inner"></div></div>',trigger:"hover focus",title:"",delay:0,html:!1,container:!1},r.prototype.init=function(t,e,n){this.enabled=!0,this.type=t,this.$element=g(e),this.options=this.getOptions(n);for(var i=this.options.trigger.split(" "),r=i.length;r--;){var o=i[r];if("click"==o)this.$element.on("click."+this.type,this.options.selector,g.proxy(this.toggle,this));else if("manual"!=o){var a="hover"==o?"mouseenter":"focus",s="hover"==o?"mouseleave":"blur";this.$element.on(a+"."+this.type,this.options.selector,g.proxy(this.enter,this)),this.$element.on(s+"."+this.type,this.options.selector,g.proxy(this.leave,this))}}this.options.selector?this._options=g.extend({},this.options,{trigger:"manual",selector:""}):this.fixTitle()},r.prototype.getDefaults=function(){return r.DEFAULTS},r.prototype.getOptions=function(t){return(t=g.extend({},this.getDefaults(),this.$element.data(),t)).delay&&"number"==typeof t.delay&&(t.delay={show:t.delay,hide:t.delay}),t},r.prototype.getDelegateOptions=function(){var n={},i=this.getDefaults();return this._options&&g.each(this._options,function(t,e){i[t]!=e&&(n[t]=e)}),n},r.prototype.enter=function(t){var e=t instanceof this.constructor?t:g(t.currentTarget)[this.type](this.getDelegateOptions()).data("bs."+this.type);return clearTimeout(e.timeout),e.hoverState="in",e.options.delay&&e.options.delay.show?void(e.timeout=setTimeout(function(){"in"==e.hoverState&&e.show()},e.options.delay.show)):e.show()},r.prototype.leave=function(t){var e=t instanceof this.constructor?t:g(t.currentTarget)[this.type](this.getDelegateOptions()).data("bs."+this.type);return clearTimeout(e.timeout),e.hoverState="out",e.options.delay&&e.options.delay.hide?void(e.timeout=setTimeout(function(){"out"==e.hoverState&&e.hide()},e.options.delay.hide)):e.hide()},r.prototype.show=function(){var t=g.Event("show.bs."+this.type);if(this.hasContent()&&this.enabled){if(this.$element.trigger(t),t.isDefaultPrevented())return;var e=this.tip();this.setContent(),this.options.animation&&e.addClass("fade");var n="function"==typeof this.options.placement?this.options.placement.call(this,e[0],this.$element[0]):this.options.placement,i=/\s?auto?\s?/i,r=i.test(n);r&&(n=n.replace(i,"")||"top"),e.detach().css({top:0,left:0,display:"block"}).addClass(n),this.options.container?e.appendTo(this.options.container):e.insertAfter(this.$element);var o=this.getPosition(),a=e[0].offsetWidth,s=e[0].offsetHeight;if(r){var l=this.$element.parent(),u=n,c=document.documentElement.scrollTop||document.body.scrollTop,h="body"==this.options.container?window.innerWidth:l.outerWidth(),d="body"==this.options.container?window.innerHeight:l.outerHeight(),p="body"==this.options.container?0:l.offset().left;n="bottom"==n&&o.top+o.height+s-c>d?"top":"top"==n&&o.top-c-s<0?"bottom":"right"==n&&o.right+a>h?"left":"left"==n&&o.left-a<p?"right":n,e.removeClass(u).addClass(n)}var f=this.getCalculatedOffset(n,o,a,s);this.applyPlacement(f,n),this.$element.trigger("shown.bs."+this.type)}},r.prototype.applyPlacement=function(t,e){var n,i=this.tip(),r=i[0].offsetWidth,o=i[0].offsetHeight,a=parseInt(i.css("margin-top"),10),s=parseInt(i.css("margin-left"),10);isNaN(a)&&(a=0),isNaN(s)&&(s=0),t.top=t.top+a,t.left=t.left+s,i.offset(t).addClass("in");var l=i[0].offsetWidth,u=i[0].offsetHeight;if("top"==e&&u!=o&&(n=!0,t.top=t.top+o-u),/bottom|top/.test(e)){var c=0;t.left<0&&(c=-2*t.left,t.left=0,i.offset(t),l=i[0].offsetWidth,u=i[0].offsetHeight),this.replaceArrow(c-r+l,l,"left")}else this.replaceArrow(u-o,u,"top");n&&i.offset(t)},r.prototype.replaceArrow=function(t,e,n){this.arrow().css(n,t?50*(1-t/e)+"%":"")},r.prototype.setContent=function(){var t=this.tip(),e=this.getTitle();t.find(".tooltip-inner")[this.options.html?"html":"text"](e),t.removeClass("fade in top bottom left right")},r.prototype.hide=function(){function t(){"in"!=e.hoverState&&n.detach()}var e=this,n=this.tip(),i=g.Event("hide.bs."+this.type);return this.$element.trigger(i),i.isDefaultPrevented()?void 0:(n.removeClass("in"),g.support.transition&&this.$tip.hasClass("fade")?n.one(g.support.transition.end,t).emulateTransitionEnd(150):t(),this.$element.trigger("hidden.bs."+this.type),this)},r.prototype.fixTitle=function(){var t=this.$element;!t.attr("title")&&"string"==typeof t.attr("data-original-title")||t.attr("data-original-title",t.attr("title")||"").attr("title","")},r.prototype.hasContent=function(){return this.getTitle()},r.prototype.getPosition=function(){var t=this.$element[0];return g.extend({},"function"==typeof t.getBoundingClientRect?t.getBoundingClientRect():{width:t.offsetWidth,height:t.offsetHeight},this.$element.offset())},r.prototype.getCalculatedOffset=function(t,e,n,i){return"bottom"==t?{top:e.top+e.height,left:e.left+e.width/2-n/2}:"top"==t?{top:e.top-i,left:e.left+e.width/2-n/2}:"left"==t?{top:e.top+e.height/2-i/2,left:e.left-n}:{top:e.top+e.height/2-i/2,left:e.left+e.width}},r.prototype.getTitle=function(){var t=this.$element,e=this.options;return t.attr("data-original-title")||("function"==typeof e.title?e.title.call(t[0]):e.title)},r.prototype.tip=function(){return this.$tip=this.$tip||g(this.options.template)},r.prototype.arrow=function(){return this.$arrow=this.$arrow||this.tip().find(".tooltip-arrow")},r.prototype.validate=function(){this.$element[0].parentNode||(this.hide(),this.$element=null,this.options=null)},r.prototype.enable=function(){this.enabled=!0},r.prototype.disable=function(){this.enabled=!1},r.prototype.toggleEnabled=function(){this.enabled=!this.enabled},r.prototype.toggle=function(t){var e=t?g(t.currentTarget)[this.type](this.getDelegateOptions()).data("bs."+this.type):this;e.tip().hasClass("in")?e.leave(e):e.enter(e)},r.prototype.destroy=function(){this.hide().$element.off("."+this.type).removeData("bs."+this.type)};var t=g.fn.tooltip;g.fn.tooltip=function(i){return this.each(function(){var t=g(this),e=t.data("bs.tooltip"),n="object"==typeof i&&i;e||t.data("bs.tooltip",e=new r(this,n)),"string"==typeof i&&e[i]()})},g.fn.tooltip.Constructor=r,g.fn.tooltip.noConflict=function(){return g.fn.tooltip=t,this}}(jQuery),function(r){"use strict";function o(t,e){this.init("popover",t,e)}if(!r.fn.tooltip)throw new Error("Popover requires tooltip.js");o.DEFAULTS=r.extend({},r.fn.tooltip.Constructor.DEFAULTS,{placement:"right",trigger:"click",content:"",template:'<div class="popover"><div class="arrow"></div><h3 class="popover-title"></h3><div class="popover-content"></div></div>'}),((o.prototype=r.extend({},r.fn.tooltip.Constructor.prototype)).constructor=o).prototype.getDefaults=function(){return o.DEFAULTS},o.prototype.setContent=function(){var t=this.tip(),e=this.getTitle(),n=this.getContent();t.find(".popover-title")[this.options.html?"html":"text"](e),t.find(".popover-content")[this.options.html?"html":"text"](n),t.removeClass("fade top bottom left right in"),t.find(".popover-title").html()||t.find(".popover-title").hide()},o.prototype.hasContent=function(){return this.getTitle()||this.getContent()},o.prototype.getContent=function(){var t=this.$element,e=this.options;return t.attr("data-content")||("function"==typeof e.content?e.content.call(t[0]):e.content)},o.prototype.arrow=function(){return this.$arrow=this.$arrow||this.tip().find(".arrow")},o.prototype.tip=function(){return this.$tip||(this.$tip=r(this.options.template)),this.$tip};var t=r.fn.popover;r.fn.popover=function(i){return this.each(function(){var t=r(this),e=t.data("bs.popover"),n="object"==typeof i&&i;e||t.data("bs.popover",e=new o(this,n)),"string"==typeof i&&e[i]()})},r.fn.popover.Constructor=o,r.fn.popover.noConflict=function(){return r.fn.popover=t,this}}(jQuery),function(o){"use strict";function r(t,e){var n,i=o.proxy(this.process,this);this.$element=o(o(t).is("body")?window:t),this.$body=o("body"),this.$scrollElement=this.$element.on("scroll.bs.scroll-spy.data-api",i),this.options=o.extend({},r.DEFAULTS,e),this.selector=(this.options.target||(n=o(t).attr("href"))&&n.replace(/.*(?=#[^\s]+$)/,"")||"")+" .nav li > a",this.offsets=o([]),this.targets=o([]),this.activeTarget=null,this.refresh(),this.process()}r.DEFAULTS={offset:10},r.prototype.refresh=function(){var i=this.$element[0]==window?"offset":"position";this.offsets=o([]),this.targets=o([]);var r=this;this.$body.find(this.selector).map(function(){var t=o(this),e=t.data("target")||t.attr("href"),n=/^#\w/.test(e)&&o(e);return n&&n.length&&[[n[i]().top+(!o.isWindow(r.$scrollElement.get(0))&&r.$scrollElement.scrollTop()),e]]||null}).sort(function(t,e){return t[0]-e[0]}).each(function(){r.offsets.push(this[0]),r.targets.push(this[1])})},r.prototype.process=function(){var t,e=this.$scrollElement.scrollTop()+this.options.offset,n=(this.$scrollElement[0].scrollHeight||this.$body[0].scrollHeight)-this.$scrollElement.height(),i=this.offsets,r=this.targets,o=this.activeTarget;if(n<=e)return o!=(t=r.last()[0])&&this.activate(t);for(t=i.length;t--;)o!=r[t]&&e>=i[t]&&(!i[t+1]||e<=i[t+1])&&this.activate(r[t])},r.prototype.activate=function(t){this.activeTarget=t,o(this.selector).parents(".active").removeClass("active");var e=this.selector+'[data-target="'+t+'"],'+this.selector+'[href="'+t+'"]',n=o(e).parents("li").addClass("active");n.parent(".dropdown-menu").length&&(n=n.closest("li.dropdown").addClass("active")),n.trigger("activate.bs.scrollspy")};var t=o.fn.scrollspy;o.fn.scrollspy=function(i){return this.each(function(){var t=o(this),e=t.data("bs.scrollspy"),n="object"==typeof i&&i;e||t.data("bs.scrollspy",e=new r(this,n)),"string"==typeof i&&e[i]()})},o.fn.scrollspy.Constructor=r,o.fn.scrollspy.noConflict=function(){return o.fn.scrollspy=t,this},o(window).on("load",function(){o('[data-spy="scroll"]').each(function(){var t=o(this);t.scrollspy(t.data())})})}(jQuery),function(a){"use strict";function i(t){this.element=a(t)}i.prototype.show=function(){var t=this.element,e=t.closest("ul:not(.dropdown-menu)"),n=t.data("target");if(n||(n=(n=t.attr("href"))&&n.replace(/.*(?=#[^\s]*$)/,"")),!t.parent("li").hasClass("active")){var i=e.find(".active:last a")[0],r=a.Event("show.bs.tab",{relatedTarget:i});if(t.trigger(r),!r.isDefaultPrevented()){var o=a(n);this.activate(t.parent("li"),e),this.activate(o,o.parent(),function(){t.trigger({type:"shown.bs.tab",relatedTarget:i})})}}},i.prototype.activate=function(t,e,n){function i(){r.removeClass("active").find("> .dropdown-menu > .active").removeClass("active"),t.addClass("active"),o?(t[0].offsetWidth,t.addClass("in")):t.removeClass("fade"),t.parent(".dropdown-menu")&&t.closest("li.dropdown").addClass("active"),n&&n()}var r=e.find("> .active"),o=n&&a.support.transition&&r.hasClass("fade");o?r.one(a.support.transition.end,i).emulateTransitionEnd(150):i(),r.removeClass("in")};var t=a.fn.tab;a.fn.tab=function(n){return this.each(function(){var t=a(this),e=t.data("bs.tab");e||t.data("bs.tab",e=new i(this)),"string"==typeof n&&e[n]()})},a.fn.tab.Constructor=i,a.fn.tab.noConflict=function(){return a.fn.tab=t,this},a(document).on("click.bs.tab.data-api",'[data-toggle="tab"], [data-toggle="pill"]',function(t){t.preventDefault(),a(this).tab("show")})}(jQuery),function(s){"use strict";var l=function(t,e){this.options=s.extend({},l.DEFAULTS,e),this.$window=s(window).on("scroll.bs.affix.data-api",s.proxy(this.checkPosition,this)).on("click.bs.affix.data-api",s.proxy(this.checkPositionWithEventLoop,this)),this.$element=s(t),this.affixed=this.unpin=null,this.checkPosition()};l.RESET="affix affix-top affix-bottom",l.DEFAULTS={offset:0},l.prototype.checkPositionWithEventLoop=function(){setTimeout(s.proxy(this.checkPosition,this),1)},l.prototype.checkPosition=function(){if(this.$element.is(":visible")){var t=s(document).height(),e=this.$window.scrollTop(),n=this.$element.offset(),i=this.options.offset,r=i.top,o=i.bottom;"object"!=typeof i&&(o=r=i),"function"==typeof r&&(r=i.top()),"function"==typeof o&&(o=i.bottom());var a=!(null!=this.unpin&&e+this.unpin<=n.top)&&(null!=o&&n.top+this.$element.height()>=t-o?"bottom":null!=r&&e<=r&&"top");this.affixed!==a&&(this.unpin&&this.$element.css("top",""),this.affixed=a,this.unpin="bottom"==a?n.top-e:null,this.$element.removeClass(l.RESET).addClass("affix"+(a?"-"+a:"")),"bottom"==a&&this.$element.offset({top:document.body.offsetHeight-o-this.$element.height()}))}};var t=s.fn.affix;s.fn.affix=function(i){return this.each(function(){var t=s(this),e=t.data("bs.affix"),n="object"==typeof i&&i;e||t.data("bs.affix",e=new l(this,n)),"string"==typeof i&&e[i]()})},s.fn.affix.Constructor=l,s.fn.affix.noConflict=function(){return s.fn.affix=t,this},s(window).on("load",function(){s('[data-spy="affix"]').each(function(){var t=s(this),e=t.data();e.offset=e.offset||{},e.offsetBottom&&(e.offset.bottom=e.offsetBottom),e.offsetTop&&(e.offset.top=e.offsetTop),t.affix(e)})})}(jQuery),function(t){var s=(e.prototype.add=function(t,e){isNaN(e[0])||isNaN(e[1]),this.locks[t]=e},e.prototype.clear=function(){this.locks={}},e.prototype.isEmpty=function(){for(var t in this.locks)return!1;return!0},e.prototype.apply=function(t){for(var e in this.locks)t(e,this.locks[e])},e);function e(){this.locks={}}t.Locks=s;var n=(a.createSquareMatrix=function(t,e){for(var n=new Array(t),i=0;i<t;++i){n[i]=new Array(t);for(var r=0;r<t;++r)n[i][r]=e(i,r)}return n},a.prototype.offsetDir=function(){for(var e=this,t=new Array(this.k),n=0,i=0;i<this.k;++i){var r=t[i]=this.random.getNextBetween(.01,1)-.5;n+=r*r}return n=Math.sqrt(n),t.map(function(t){return t*(e.minD/n)})},a.prototype.computeDerivatives=function(n){var i=this,t=this.n;if(!(t<1)){for(var r,e=new Array(this.k),o=new Array(this.k),a=new Array(this.k),s=0,l=0;l<t;++l){for(r=0;r<this.k;++r)a[r]=this.g[r][l]=0;for(var u=0;u<t;++u)if(l!==u){for(var c=t;c--;){var h=0;for(r=0;r<this.k;++r){var d=e[r]=n[r][l]-n[r][u];h+=o[r]=d*d}if(1e-9<h)break;var p=this.offsetDir();for(r=0;r<this.k;++r)n[r][u]+=p[r]}var f=Math.sqrt(h),g=this.D[l][u],v=null!=this.G?this.G[l][u]:1;if(1<v&&g<f||!isFinite(g))for(r=0;r<this.k;++r)this.H[r][l][u]=0;else{1<v&&(v=1);var m=g*g,y=v*(f-g)/(m*f),b=-v/(m*f*f*f);for(isFinite(y)||console.log(y),r=0;r<this.k;++r)this.g[r][l]+=e[r]*y,a[r]-=this.H[r][l][u]=b*(g*(o[r]-h)+f*h)}}for(r=0;r<this.k;++r)s=Math.max(s,this.H[r][l][l]=a[r])}this.locks.isEmpty()||this.locks.apply(function(t,e){for(r=0;r<i.k;++r)i.H[r][t][t]+=s,i.g[r][t]-=s*(e[r]-n[r][t])})}},a.dotProd=function(t,e){for(var n=0,i=t.length;i--;)n+=t[i]*e[i];return n},a.rightMultiply=function(t,e,n){for(var i=t.length;i--;)n[i]=a.dotProd(t[i],e)},a.prototype.computeStepSize=function(t){for(var e=0,n=0,i=0;i<2;++i)e+=a.dotProd(this.g[i],t[i]),a.rightMultiply(this.H[i],t[i],this.Hd[i]),n+=a.dotProd(t[i],this.Hd[i]);return 0!==n&&isFinite(n)?e/n:0},a.prototype.reduceStress=function(){this.computeDerivatives(this.x);for(var t=this.computeStepSize(this.g),e=0;e<this.k;++e)this.takeDescentStep(this.x[e],this.g[e],t);return this.computeStress()},a.copy=function(t,e){for(var n=t.length,i=e[0].length,r=0;r<n;++r)for(var o=0;o<i;++o)e[r][o]=t[r][o]},a.prototype.stepAndProject=function(t,e,n,i){a.copy(t,e),this.takeDescentStep(e[0],n[0],i),this.project&&this.project[0](t[0],t[1],e[0]),this.takeDescentStep(e[1],n[1],i),this.project&&this.project[1](e[0],t[1],e[1])},a.mApply=function(t,e,n){for(var i=t;0<i--;)for(var r=e;0<r--;)n(i,r)},a.prototype.matrixApply=function(t){a.mApply(this.k,this.n,t)},a.prototype.computeNextPosition=function(n,i){var r=this;this.computeDerivatives(n);var t=this.computeStepSize(this.g);this.stepAndProject(n,i,this.g,t);for(var e=0;e<this.n;++e)for(var o=0;o<this.k;++o)isNaN(i[o][e]);if(this.project){this.matrixApply(function(t,e){return r.e[t][e]=n[t][e]-i[t][e]});var a=this.computeStepSize(this.e);a=Math.max(.2,Math.min(a,1)),this.stepAndProject(n,i,this.e,a)}},a.prototype.run=function(t){for(var e=Number.MAX_VALUE,n=!1;!n&&0<t--;){var i=this.rungeKutta();n=Math.abs(e/i-1)<this.threshold,e=i}return e},a.prototype.rungeKutta=function(){var r=this;this.computeNextPosition(this.x,this.a),a.mid(this.x,this.a,this.ia),this.computeNextPosition(this.ia,this.b),a.mid(this.x,this.b,this.ib),this.computeNextPosition(this.ib,this.c),this.computeNextPosition(this.c,this.d);var o=0;return this.matrixApply(function(t,e){var n=(r.a[t][e]+2*r.b[t][e]+2*r.c[t][e]+r.d[t][e])/6,i=r.x[t][e]-n;o+=i*i,r.x[t][e]=n}),o},a.mid=function(n,i,r){a.mApply(n.length,n[0].length,function(t,e){return r[t][e]=n[t][e]+(i[t][e]-n[t][e])/2})},a.prototype.takeDescentStep=function(t,e,n){for(var i=0;i<this.n;++i)t[i]=t[i]-n*e[i]},a.prototype.computeStress=function(){for(var t=0,e=0,n=this.n-1;e<n;++e)for(var i=e+1,r=this.n;i<r;++i){for(var o=0,a=0;a<this.k;++a){var s=this.x[a][e]-this.x[a][i];o+=s*s}o=Math.sqrt(o);var l=this.D[e][i];if(isFinite(l)){var u=l-o;t+=u*u/(l*l)}}return t},a.zeroDistance=1e-10,a);function a(t,e,n){void 0===n&&(n=null),this.D=e,this.G=n,this.threshold=1e-4,this.random=new l,this.project=null,this.x=t,this.k=t.length;var i=this.n=t[0].length;this.H=new Array(this.k),this.g=new Array(this.k),this.Hd=new Array(this.k),this.a=new Array(this.k),this.b=new Array(this.k),this.c=new Array(this.k),this.d=new Array(this.k),this.e=new Array(this.k),this.ia=new Array(this.k),this.ib=new Array(this.k),this.xtmp=new Array(this.k),this.locks=new s,this.minD=Number.MAX_VALUE;for(var r,o=i;o--;)for(r=i;--r>o;){var a=e[o][r];0<a&&a<this.minD&&(this.minD=a)}for(this.minD===Number.MAX_VALUE&&(this.minD=1),o=this.k;o--;){for(this.g[o]=new Array(i),this.H[o]=new Array(i),r=i;r--;)this.H[o][r]=new Array(i);this.Hd[o]=new Array(i),this.a[o]=new Array(i),this.b[o]=new Array(i),this.c[o]=new Array(i),this.d[o]=new Array(i),this.e[o]=new Array(i),this.ia[o]=new Array(i),this.ib[o]=new Array(i),this.xtmp[o]=new Array(i)}}t.Descent=n;var l=(i.prototype.getNext=function(){return this.seed=(this.seed*this.a+this.c)%this.m,(this.seed>>16)/this.range},i.prototype.getNextBetween=function(t,e){return t+this.getNext()*(e-t)},i);function i(t){void 0===t&&(t=1),this.seed=t,this.a=214013,this.c=2531011,this.m=2147483648,this.range=32767}t.PseudoRandom=l}(cola||(cola={}));var __extends=this.__extends||function(t,e){function n(){this.constructor=t}for(var i in e)e.hasOwnProperty(i)&&(t[i]=e[i]);n.prototype=e.prototype,t.prototype=new n},cola,cola,cola;!function(E){!function(g){function b(t,e,n){return(e.x-t.x)*(n.y-t.y)-(n.x-t.x)*(e.y-t.y)}function s(t,e,n){return 0<b(t,e,n)}function l(t,e,n){return b(t,e,n)<0}function u(t,e){var n,i,r,o,a=e.length-1;if(l(t,e[1],e[0])&&!s(t,e[a-1],e[0]))return 0;for(n=0,i=a;;){if(i-n==1)return s(t,e[n],e[i])?n:i;if((o=l(t,e[(r=Math.floor((n+i)/2))+1],e[r]))&&!s(t,e[r-1],e[r]))return r;s(t,e[n+1],e[n])?o?i=r:s(t,e[n],e[r])?i=r:n=r:o&&l(t,e[n],e[r])?i=r:n=r}}function c(t,e){var n,i,r,o,a=e.length-1;if(s(t,e[a-1],e[0])&&!l(t,e[1],e[0]))return 0;for(n=0,i=a;;){if(i-n==1)return l(t,e[n],e[i])?n:i;if(o=l(t,e[(r=Math.floor((n+i)/2))+1],e[r]),s(t,e[r-1],e[r])&&!o)return r;l(t,e[n+1],e[n])?o?l(t,e[n],e[r])?i=r:n=r:i=r:o?n=r:s(t,e[n],e[r])?i=r:n=r}}function n(t,e,n,i,r,o){var a,s;s=i(t[a=n(e[0],t)],e);for(var l=!1;!l;){for(l=!0;a===t.length-1&&(a=0),!r(e[s],t[a],t[a+1]);)++a;for(;0===s&&(s=e.length-1),!o(t[a],e[s],e[s-1]);)--s,l=!1}return{t1:a,t2:s}}function i(t,e){return n(t,e,u,c,s,l)}function a(t,e){for(var n=[],i=1,r=e.length;i<r;++i){var o=E.vpsc.Rectangle.lineIntersection(t.x1,t.y1,t.x2,t.y2,e[i-1].x,e[i-1].y,e[i].x,e[i].y);o&&n.push(o)}return n}function h(t,e){return!t.every(function(t){return!function(t,e){for(var n=1,i=e.length;n<i;++n)if(l(e[n-1],e[n],t))return!1;return!0}(t,e)})}function t(){}g.Point=t;var d=function(t,e,n,i){this.x1=t,this.y1=e,this.x2=n,this.y2=i};g.LineSegment=d;var e,r=(__extends(o,e=t),o);function o(){e.apply(this,arguments)}g.PolyPoint=r,g.isLeft=b,g.ConvexHull=function(t){var e,n=t.slice(0).sort(function(t,e){return t.x!==e.x?e.x-t.x:e.y-t.y}),i=t.length,r=n[0].x;for(e=1;e<i&&n[e].x===r;++e);var o=e-1,a=[];if(a.push(n[0]),o==i-1)n[o].y!==n[0].y&&a.push(n[o]);else{var s,l=i-1,u=n[i-1].x;for(e=i-2;0<=e&&n[e].x===u;e--);for(s=e+1,e=o;++e<=s;)if(!(0<=b(n[0],n[s],n[e])&&e<s)){for(;1<a.length&&!(0<b(a[a.length-2],a[a.length-1],n[e]));)a.length-=1;0!=e&&a.push(n[e])}l!=s&&a.push(n[l]);var c=a.length;for(e=s;--e>=o;)if(!(0<=b(n[l],n[o],n[e])&&o<e)){for(;a.length>c&&!(0<b(a[a.length-2],a[a.length-1],n[e]));)a.length-=1;0!=e&&a.push(n[e])}}return a},g.clockwiseRadialSweep=function(n,t,e){t.slice(0).sort(function(t,e){return Math.atan2(t.y-n.y,t.x-n.x)-Math.atan2(e.y-n.y,e.x-n.x)}).forEach(e)},g.tangent_PolyPolyC=n,g.LRtangent_PolyPolyC=function(t,e){var n=i(e,t);return{t1:n.t2,t2:n.t1}},g.RLtangent_PolyPolyC=i,g.LLtangent_PolyPolyC=function(t,e){return n(t,e,c,c,l,l)},g.RRtangent_PolyPolyC=function(t,e){return n(t,e,u,u,s,s)};var x=function(t,e){this.t1=t,this.t2=e};g.BiTangent=x;var w=function(){};g.BiTangents=w;var p,f=(__extends(v,p=t),v);function v(){p.apply(this,arguments)}g.TVGPoint=f;var m=function(t,e,n,i){this.id=t,this.polyid=e,this.polyvertid=n,(this.p=i).vv=this};g.VisibilityVertex=m;var y=($.prototype.length=function(){var t=this.source.p.x-this.target.p.x,e=this.source.p.y-this.target.p.y;return Math.sqrt(t*t+e*e)},$);function $(t,e){this.source=t,this.target=e}g.VisibilityEdge=y;var k=(C.prototype.addEdgeIfVisible=function(t,e,n,i){this.intersectsPolys(new d(t.x,t.y,e.x,e.y),n,i)||this.E.push(new y(t.vv,e.vv))},C.prototype.addPoint=function(t,e){var n,i,r=this.P.length;this.V.push(new m(this.V.length,r,0,t));for(var o=0;o<r;++o)if(o!==e){var a=this.P[o],s={rtan:u(n=t,i=a),ltan:c(n,i)};this.addEdgeIfVisible(t,a[s.ltan],e,o),this.addEdgeIfVisible(t,a[s.rtan],e,o)}return t.vv},C.prototype.intersectsPolys=function(t,e,n){for(var i=0,r=this.P.length;i<r;++i)if(i!=e&&i!=n&&0<a(t,this.P[i]).length)return!0;return!1},C);function C(t,e){if(this.P=t,this.V=[],this.E=[],e)this.V=e.V.slice(0),this.E=e.E.slice(0);else{for(var n=t.length,i=0;i<n;i++)for(var r=t[i],o=0;o<r.length;++o){var a=r[o],s=new m(this.V.length,i,o,a);this.V.push(s),0<o&&this.E.push(new y(r[o-1].vv,s))}for(i=0;i<n-1;i++){var l=t[i];for(o=i+1;o<n;o++){var u=t[o],c=g.tangents(l,u);for(var h in c){var d=c[h],p=l[d.t1],f=u[d.t2];this.addEdgeIfVisible(p,f,i,o)}}}}}g.TangentVisibilityGraph=k,g.tangents=function(t,e){for(var n=t.length-1,i=e.length-1,r=new w,o=0;o<n;++o)for(var a=0;a<i;++a){var s=t[0==o?n-1:o-1],l=t[o],u=t[o+1],c=e[0==a?i-1:a-1],h=e[a],d=e[a+1],p=b(s,l,h),f=b(l,c,h),g=b(l,h,d),v=b(c,h,l),m=b(h,s,l),y=b(h,l,u);0<=p&&0<=f&&g<0&&0<=v&&0<=m&&y<0?r.ll=new x(o,a):p<=0&&f<=0&&0<g&&v<=0&&m<=0&&0<y?r.rr=new x(o,a):p<=0&&0<f&&g<=0&&0<=v&&m<0&&0<=y?r.rl=new x(o,a):0<=p&&f<0&&0<=g&&v<=0&&0<m&&y<=0&&(r.lr=new x(o,a))}return r},g.polysOverlap=function(t,e){if(h(t,e))return!0;if(h(e,t))return!0;for(var n=1,i=t.length;n<i;++n){var r=t[n],o=t[n-1];if(0<a(new d(o.x,o.y,r.x,r.y),e).length)return!0}return!1}}(E.geom||(E.geom={}))}(cola||(cola={})),function(t){!function(t){var e=(n.prototype.addVariable=function(t){var e=this.scale/t.scale,n=t.offset/t.scale,i=t.weight;this.AB+=i*e*n,this.AD+=i*e*t.desiredPosition,this.A2+=i*e*e},n.prototype.getPosn=function(){return(this.AD-this.AB)/this.A2},n);function n(t){this.scale=t,this.AB=0,this.AD=0,this.A2=0}t.PositionStats=e;var i=(r.prototype.slack=function(){return this.unsatisfiable?Number.MAX_VALUE:this.right.scale*this.right.position()-this.gap-this.left.scale*this.left.position()},r);function r(t,e,n,i){void 0===i&&(i=!1),this.left=t,this.right=e,this.gap=n,this.equality=i,this.active=!1,this.unsatisfiable=!1,this.left=t,this.right=e,this.gap=n,this.equality=i}t.Constraint=i;var o=(a.prototype.dfdv=function(){return 2*this.weight*(this.position()-this.desiredPosition)},a.prototype.position=function(){return(this.block.ps.scale*this.block.posn+this.offset)/this.scale},a.prototype.visitNeighbours=function(n,i){function e(t,e){return t.active&&n!==e&&i(t,e)}this.cOut.forEach(function(t){return e(t,t.right)}),this.cIn.forEach(function(t){return e(t,t.left)})},a);function a(t,e,n){void 0===e&&(e=1),void 0===n&&(n=1),this.desiredPosition=t,this.weight=e,this.scale=n,this.offset=0}t.Variable=o;var s=(l.prototype.addVariable=function(t){(t.block=this).vars.push(t),this.ps.addVariable(t),this.posn=this.ps.getPosn()},l.prototype.updateWeightedPosition=function(){this.ps.AB=this.ps.AD=this.ps.A2=0;for(var t=0,e=this.vars.length;t<e;++t)this.ps.addVariable(this.vars[t]);this.posn=this.ps.getPosn()},l.prototype.compute_lm=function(i,t,r){var o=this,a=i.dfdv();return i.visitNeighbours(t,function(t,e){var n=o.compute_lm(e,i,r);e===t.right?(a+=n*t.left.scale,t.lm=n):(a+=n*t.right.scale,t.lm=-n),r(t)}),a/i.scale},l.prototype.populateSplitBlock=function(n,t){var i=this;n.visitNeighbours(t,function(t,e){e.offset=n.offset+(e===t.right?t.gap:-t.gap),i.addVariable(e),i.populateSplitBlock(e,n)})},l.prototype.traverse=function(n,i,r,t){var o=this;void 0===r&&(r=this.vars[0]),void 0===t&&(t=null),r.visitNeighbours(t,function(t,e){i.push(n(t)),o.traverse(n,i,e,r)})},l.prototype.findMinLM=function(){var e=null;return this.compute_lm(this.vars[0],null,function(t){!t.equality&&(null===e||t.lm<e.lm)&&(e=t)}),e},l.prototype.findMinLMBetween=function(t,e){this.compute_lm(t,null,function(){});var n=null;return this.findPath(t,null,e,function(t,e){!t.equality&&t.right===e&&(null===n||t.lm<n.lm)&&(n=t)}),n},l.prototype.findPath=function(n,t,i,r){var o=this,a=!1;return n.visitNeighbours(t,function(t,e){a||e!==i&&!o.findPath(e,n,i,r)||(a=!0,r(t,e))}),a},l.prototype.isActiveDirectedPathBetween=function(t,e){if(t===e)return!0;for(var n=t.cOut.length;n--;){var i=t.cOut[n];if(i.active&&this.isActiveDirectedPathBetween(i.right,e))return!0}return!1},l.split=function(t){return t.active=!1,[l.createSplitBlock(t.left),l.createSplitBlock(t.right)]},l.createSplitBlock=function(t){var e=new l(t);return e.populateSplitBlock(t,null),e},l.prototype.splitBetween=function(t,e){var n=this.findMinLMBetween(t,e);if(null===n)return null;var i=l.split(n);return{constraint:n,lb:i[0],rb:i[1]}},l.prototype.mergeAcross=function(t,e,n){e.active=!0;for(var i=0,r=t.vars.length;i<r;++i){var o=t.vars[i];o.offset+=n,this.addVariable(o)}this.posn=this.ps.getPosn()},l.prototype.cost=function(){for(var t=0,e=this.vars.length;e--;){var n=this.vars[e],i=n.position()-n.desiredPosition;t+=i*i*n.weight}return t},l);function l(t){this.vars=[],t.offset=0,this.ps=new e(t.scale),this.addVariable(t)}t.Block=s;var u=(c.prototype.cost=function(){for(var t=0,e=this.list.length;e--;)t+=this.list[e].cost();return t},c.prototype.insert=function(t){t.blockInd=this.list.length,this.list.push(t)},c.prototype.remove=function(t){var e=this.list.length-1,n=this.list[e];this.list.length=e,t!==n&&((this.list[t.blockInd]=n).blockInd=t.blockInd)},c.prototype.merge=function(t){var e=t.left.block,n=t.right.block,i=t.right.offset-t.left.offset-t.gap;e.vars.length<n.vars.length?(n.mergeAcross(e,t,i),this.remove(e)):(e.mergeAcross(n,t,-i),this.remove(n))},c.prototype.forEach=function(t){this.list.forEach(t)},c.prototype.updateBlockPositions=function(){this.list.forEach(function(t){return t.updateWeightedPosition()})},c.prototype.split=function(n){var i=this;this.updateBlockPositions(),this.list.forEach(function(t){var e=t.findMinLM();null!==e&&e.lm<h.LAGRANGIAN_TOLERANCE&&(t=e.left.block,s.split(e).forEach(function(t){return i.insert(t)}),i.remove(t),n.push(e))})},c);function c(t){var e=(this.vs=t).length;for(this.list=new Array(e);e--;){var n=new s(t[e]);(this.list[e]=n).blockInd=e}}t.Blocks=u;var h=(d.prototype.cost=function(){return this.bs.cost()},d.prototype.setStartingPositions=function(n){this.inactive=this.cs.map(function(t){return t.active=!1,t}),this.bs=new u(this.vs),this.bs.forEach(function(t,e){return t.posn=n[e]})},d.prototype.setDesiredPositions=function(n){this.vs.forEach(function(t,e){return t.desiredPosition=n[e]})},d.prototype.mostViolated=function(){for(var t=Number.MAX_VALUE,e=null,n=this.inactive,i=n.length,r=i,o=0;o<i;++o){var a=n[o];if(!a.unsatisfiable){var s=a.slack();if((a.equality||s<t)&&(t=s,r=o,(e=a).equality))break}}return r!==i&&(t<d.ZERO_UPPERBOUND&&!e.active||e.equality)&&(n[r]=n[i-1],n.length=i-1),e},d.prototype.satisfy=function(){null==this.bs&&(this.bs=new u(this.vs)),this.bs.split(this.inactive);for(var t=null;(t=this.mostViolated())&&(t.equality||t.slack()<d.ZERO_UPPERBOUND&&!t.active);){var e=t.left.block;if(e!==t.right.block)this.bs.merge(t);else{if(e.isActiveDirectedPathBetween(t.right,t.left)){t.unsatisfiable=!0;continue}var n=e.splitBetween(t.left,t.right);if(null===n){t.unsatisfiable=!0;continue}this.bs.insert(n.lb),this.bs.insert(n.rb),this.bs.remove(e),this.inactive.push(n.constraint),0<=t.slack()?this.inactive.push(t):this.bs.merge(t)}}},d.prototype.solve=function(){this.satisfy();for(var t=Number.MAX_VALUE,e=this.bs.cost();1e-4<Math.abs(t-e);)this.satisfy(),t=e,e=this.bs.cost();return e},d.LAGRANGIAN_TOLERANCE=-1e-4,d.ZERO_UPPERBOUND=-1e-10,d);function d(t,e){this.vs=t,this.cs=e,(this.vs=t).forEach(function(t){t.cIn=[],t.cOut=[]}),(this.cs=e).forEach(function(t){t.left.cOut.push(t),t.right.cIn.push(t)}),this.inactive=e.map(function(t){return t.active=!1,t}),this.bs=null}t.Solver=h}(t.vpsc||(t.vpsc={}))}(cola||(cola={})),function(t){!function(g){function u(t){return t.bounds=void 0!==t.leaves?t.leaves.reduce(function(t,e){return e.bounds.union(t)},c.empty()):c.empty(),void 0!==t.groups&&(t.bounds=t.groups.reduce(function(t,e){return u(e).union(t)},t.bounds)),t.bounds=t.bounds.inflate(t.padding),t.bounds}function v(t,e){return t.pos>e.pos?1:t.pos<e.pos?-1:t.isOpen?-1:0}function m(){return new RBTree(function(t,e){return t.pos-e.pos})}function w(t,i,n,e){void 0===e&&(e=!1);function r(t,e){h[d]=t,c[d++]=e}var o=t.padding,a=void 0!==t.groups?t.groups.length:0,s=void 0!==t.leaves?t.leaves.length:0,l=a?t.groups.reduce(function(t,e){return t.concat(w(e,i,n,!0))},[]):[],u=(e?2:0)+s+a,c=new Array(u),h=new Array(u),d=0;if(e){var p=t.bounds,f=i.getCentre(p),g=i.getSize(p)/2,v=i.getOpen(p),m=i.getClose(p),y=f-g+o/2,b=f+g-o/2;t.minVar.desiredPosition=y,r(i.makeRect(v,m,y,o),t.minVar),t.maxVar.desiredPosition=b,r(i.makeRect(v,m,b,o),t.maxVar)}s&&t.leaves.forEach(function(t){return r(t.bounds,t.variable)}),a&&t.groups.forEach(function(t){var e=t.bounds;r(i.makeRect(i.getOpen(e),i.getClose(e),i.getCentre(e),i.getSize(e)),t.minVar)});var x=$(h,c,i,n);return a&&(c.forEach(function(t){t.cOut=[],t.cIn=[]}),x.forEach(function(t){t.left.cOut.push(t),t.right.cIn.push(t)}),t.groups.forEach(function(e){var n=(e.padding-i.getSize(e.bounds))/2;e.minVar.cIn.forEach(function(t){return t.gap+=n}),e.minVar.cOut.forEach(function(t){t.left=e.maxVar,t.gap+=n})})),l.concat(x)}function $(t,e,i,r){var n,o=t.length,a=2*o;console.assert(e.length>=o);var s=new Array(a);for(n=0;n<o;++n){var l=t[n],u=new y(e[n],l,i.getCentre(l));s[n]=new b(!0,u,i.getOpen(l)),s[n+o]=new b(!1,u,i.getClose(l))}s.sort(v);var c=new Array,h=m();for(n=0;n<a;++n){var d=s[n];u=d.v;if(d.isOpen)h.insert(u),i.findNeighbours(u,h);else{h.remove(u);function p(t,e){var n=(i.getSize(t.r)+i.getSize(e.r))/2+r;c.push(new g.Constraint(t.v,e.v,n))}function f(t,e,n){for(var i,r=u[t].iterator();null!==(i=r[t]());)n(i,u),i[e].remove(u)}f("prev","next",function(t,e){return p(t,e)}),f("next","prev",function(t,e){return p(e,t)})}}return console.assert(0===h.size),c}function n(t){return w(t,r,1e-6)}function i(t){return w(t,o,1e-6)}g.computeGroupBounds=u;var c=(l.empty=function(){return new l(Number.POSITIVE_INFINITY,Number.NEGATIVE_INFINITY,Number.POSITIVE_INFINITY,Number.NEGATIVE_INFINITY)},l.prototype.cx=function(){return(this.x+this.X)/2},l.prototype.cy=function(){return(this.y+this.Y)/2},l.prototype.overlapX=function(t){var e=this.cx(),n=t.cx();return e<=n&&t.x<this.X?this.X-t.x:n<=e&&this.x<t.X?t.X-this.x:0},l.prototype.overlapY=function(t){var e=this.cy(),n=t.cy();return e<=n&&t.y<this.Y?this.Y-t.y:n<=e&&this.y<t.Y?t.Y-this.y:0},l.prototype.setXCentre=function(t){var e=t-this.cx();this.x+=e,this.X+=e},l.prototype.setYCentre=function(t){var e=t-this.cy();this.y+=e,this.Y+=e},l.prototype.width=function(){return this.X-this.x},l.prototype.height=function(){return this.Y-this.y},l.prototype.union=function(t){return new l(Math.min(this.x,t.x),Math.max(this.X,t.X),Math.min(this.y,t.y),Math.max(this.Y,t.Y))},l.prototype.lineIntersections=function(t,e,n,i){for(var r=[[this.x,this.y,this.X,this.y],[this.X,this.y,this.X,this.Y],[this.X,this.Y,this.x,this.Y],[this.x,this.Y,this.x,this.y]],o=[],a=0;a<4;++a){var s=l.lineIntersection(t,e,n,i,r[a][0],r[a][1],r[a][2],r[a][3]);null!==s&&o.push({x:s.x,y:s.y})}return o},l.prototype.rayIntersection=function(t,e){var n=this.lineIntersections(this.cx(),this.cy(),t,e);return 0<n.length?n[0]:null},l.prototype.vertices=function(){return[{x:this.x,y:this.y},{x:this.X,y:this.y},{x:this.X,y:this.Y},{x:this.x,y:this.Y},{x:this.x,y:this.y}]},l.lineIntersection=function(t,e,n,i,r,o,a,s){var l=n-t,u=a-r,c=i-e,h=s-o,d=h*l-u*c;if(0==d)return null;var p=t-r,f=e-o,g=(u*f-h*p)/d,v=(l*f-c*p)/d;return 0<=g&&g<=1&&0<=v&&v<=1?{x:t+g*l,y:e+g*c}:null},l.prototype.inflate=function(t){return new l(this.x-t,this.X+t,this.y-t,this.Y+t)},l);function l(t,e,n,i){this.x=t,this.X=e,this.y=n,this.Y=i}g.Rectangle=c,g.makeEdgeBetween=function(t,e,n,i){var r=e.rayIntersection(n.cx(),n.cy());r||(r={x:e.cx(),y:e.cy()});var o=n.rayIntersection(e.cx(),e.cy());o||(o={x:n.cx(),y:n.cy()});var a=o.x-r.x,s=o.y-r.y,l=Math.sqrt(a*a+s*s),u=l-i;t.sourceIntersection=r,t.targetIntersection=o,t.arrowStart={x:r.x+u*a/l,y:r.y+u*s/l}},g.makeEdgeTo=function(t,e,n){var i=e.rayIntersection(t.x,t.y);i||(i={x:e.cx(),y:e.cy()});var r=i.x-t.x,o=i.y-t.y,a=Math.sqrt(r*r+o*o);return{x:i.x-n*r/a,y:i.y-n*o/a}};var y=function(t,e,n){this.v=t,this.r=e,this.pos=n,this.prev=m(),this.next=m()},b=function(t,e,n){this.isOpen=t,this.v=e,this.pos=n},r={getCentre:function(t){return t.cx()},getOpen:function(t){return t.y},getClose:function(t){return t.Y},getSize:function(t){return t.width()},makeRect:function(t,e,n,i){return new c(n-i/2,n+i/2,t,e)},findNeighbours:function(o,a){function t(t,e){for(var n,i=a.findIter(o);null!==(n=i[t]());){var r=n.r.overlapX(o.r);if((r<=0||r<=n.r.overlapY(o.r))&&(o[t].insert(n),n[e].insert(o)),r<=0)break}}t("next","prev"),t("prev","next")}},o={getCentre:function(t){return t.cy()},getOpen:function(t){return t.x},getClose:function(t){return t.X},getSize:function(t){return t.height()},makeRect:function(t,e,n,i){return new c(t,e,n-i/2,n+i/2)},findNeighbours:function(i,r){function t(t,e){var n=r.findIter(i)[t]();null!==n&&0<n.r.overlapX(i.r)&&(i[t].insert(n),n[e].insert(i))}t("next","prev"),t("prev","next")}};g.generateXConstraints=function(t,e){return $(t,e,r,1e-6)},g.generateYConstraints=function(t,e){return $(t,e,o,1e-6)},g.generateXGroupConstraints=n,g.generateYGroupConstraints=i,g.removeOverlaps=function(n){var t=n.map(function(t){return new g.Variable(t.cx())}),e=g.generateXConstraints(n,t),i=new g.Solver(t,e);i.solve(),t.forEach(function(t,e){return n[e].setXCentre(t.position())}),t=n.map(function(t){return new g.Variable(t.cy())}),e=g.generateYConstraints(n,t),(i=new g.Solver(t,e)).solve(),t.forEach(function(t,e){return n[e].setYCentre(t.position())})};var a,s=(a=g.Variable,__extends(h,a),h),t=(e.prototype.createSeparation=function(t){return new g.Constraint(this.nodes[t.left].variable,this.nodes[t.right].variable,t.gap,void 0!==t.equality&&t.equality)},e.prototype.makeFeasible=function(t){var e=this;if(this.avoidOverlaps){var n="x",i="width";"x"===t.axis&&(n="y",i="height");var r=t.offsets.map(function(t){return e.nodes[t.node]}).sort(function(t,e){return t[n]-e[n]}),o=null;r.forEach(function(t){o&&(t[n]=o[n]+o[i]+1),o=t})}},e.prototype.createAlignment=function(t){var n=this,i=this.nodes[t.offsets[0].node].variable;this.makeFeasible(t);var r="x"===t.axis?this.xConstraints:this.yConstraints;t.offsets.slice(1).forEach(function(t){var e=n.nodes[t.node].variable;r.push(new g.Constraint(i,e,t.offset,!0))})},e.prototype.createConstraints=function(t){function e(t){return void 0===t.type||"separation"===t.type}var n=this;this.xConstraints=t.filter(function(t){return"x"===t.axis&&e(t)}).map(function(t){return n.createSeparation(t)}),this.yConstraints=t.filter(function(t){return"y"===t.axis&&e(t)}).map(function(t){return n.createSeparation(t)}),t.filter(function(t){return"alignment"===t.type}).forEach(function(t){return n.createAlignment(t)})},e.prototype.setupVariablesAndBounds=function(a,s,l,u){this.nodes.forEach(function(t,e){t.fixed?(t.variable.weight=1e3,l[e]=u(t)):t.variable.weight=1;var n=(t.width||0)/2,i=(t.height||0)/2,r=a[e],o=s[e];t.bounds=new c(r-n,r+n,o-i,o+i)})},e.prototype.xProject=function(t,e,r){(this.rootGroup||this.avoidOverlaps||this.xConstraints)&&this.project(t,e,t,r,function(t){return t.px},this.xConstraints,n,function(t){return t.bounds.setXCentre(r[t.variable.index]=t.variable.position())},function(t){var e=r[t.minVar.index]=t.minVar.position(),n=r[t.maxVar.index]=t.maxVar.position(),i=t.padding/2;t.bounds.x=e-i,t.bounds.X=n+i})},e.prototype.yProject=function(t,e,r){(this.rootGroup||this.yConstraints)&&this.project(t,e,e,r,function(t){return t.py},this.yConstraints,i,function(t){return t.bounds.setYCentre(r[t.variable.index]=t.variable.position())},function(t){var e=r[t.minVar.index]=t.minVar.position(),n=r[t.maxVar.index]=t.maxVar.position(),i=t.padding/2;t.bounds.y=e-i,t.bounds.Y=n+i})},e.prototype.projectFunctions=function(){var i=this;return[function(t,e,n){return i.xProject(t,e,n)},function(t,e,n){return i.yProject(t,e,n)}]},e.prototype.project=function(t,e,n,i,r,o,a,s,l){this.setupVariablesAndBounds(t,e,i,r),this.rootGroup&&this.avoidOverlaps&&(u(this.rootGroup),o=o.concat(a(this.rootGroup))),this.solve(this.variables,o,n,i),this.nodes.forEach(s),this.rootGroup&&this.avoidOverlaps&&this.groups.forEach(l)},e.prototype.solve=function(t,e,n,i){var r=new g.Solver(t,e);r.setStartingPositions(n),r.setDesiredPositions(i),r.solve()},e);function e(t,e,n,i,r){var o=this;if(void 0===n&&(n=null),void 0===i&&(i=null),void 0===r&&(r=!1),this.nodes=t,this.groups=e,this.rootGroup=n,this.avoidOverlaps=r,this.variables=t.map(function(t,e){return t.variable=new s(e,1)}),i&&this.createConstraints(i),r&&n&&void 0!==n.groups){t.forEach(function(t){if(t.width&&t.height){var e=t.width/2,n=t.height/2;t.bounds=new g.Rectangle(t.x-e,t.x+e,t.y-n,t.y+n)}else t.bounds=new g.Rectangle(t.x,t.x,t.y,t.y)}),u(n);var a=t.length;e.forEach(function(t){o.variables[a]=t.minVar=new s(a++,.01),o.variables[a]=t.maxVar=new s(a++,.01)})}}function h(t,e){a.call(this,0,e),this.index=t}g.Projection=t}(t.vpsc||(t.vpsc={}))}(cola||(cola={}));var PairingHeap=function(){function a(t){this.elem=t,this.subheaps=[]}return a.prototype.toString=function(t){for(var e="",n=!1,i=0;i<this.subheaps.length;++i){var r=this.subheaps[i];n=!!r.elem&&(n&&(e+=","),e+=r.toString(t),!0)}return""!==e&&(e="("+e+")"),(this.elem?t(this.elem):"")+e},a.prototype.forEach=function(e){this.empty()||(e(this.elem,this),this.subheaps.forEach(function(t){return t.forEach(e)}))},a.prototype.count=function(){return this.empty()?0:1+this.subheaps.reduce(function(t,e){return t+e.count()},0)},a.prototype.min=function(){return this.elem},a.prototype.empty=function(){return null==this.elem},a.prototype.contains=function(t){if(this===t)return!0;for(var e=0;e<this.subheaps.length;e++)if(this.subheaps[e].contains(t))return!0;return!1},a.prototype.isHeap=function(e){var n=this;return this.subheaps.every(function(t){return e(n.elem,t.elem)&&t.isHeap(e)})},a.prototype.insert=function(t,e){return this.merge(new a(t),e)},a.prototype.merge=function(t,e){return this.empty()?t:t.empty()?this:e(this.elem,t.elem)?(this.subheaps.push(t),this):(t.subheaps.push(this),t)},a.prototype.removeMin=function(t){return this.empty()?null:this.mergePairs(t)},a.prototype.mergePairs=function(t){if(0==this.subheaps.length)return new a(null);if(1==this.subheaps.length)return this.subheaps[0];var e=this.subheaps.pop().merge(this.subheaps.pop(),t),n=this.mergePairs(t);return e.merge(n,t)},a.prototype.decreaseKey=function(t,e,n,i){var r=t.removeMin(i);t.elem=r.elem,t.subheaps=r.subheaps,null!==n&&null!==r.elem&&n(t.elem,t);var o=new a(e);return null!==n&&n(e,o),this.merge(o,i)},a}(),PriorityQueue=function(){function t(t){this.lessThan=t}return t.prototype.top=function(){return this.empty()?null:this.root.elem},t.prototype.push=function(){for(var t=[],e=0;e<arguments.length;e++)t[e-0]=arguments[e];for(var n,i,r=0;i=t[r];++r)n=new PairingHeap(i),this.root=this.empty()?n:this.root.merge(n,this.lessThan);return n},t.prototype.empty=function(){return!this.root||!this.root.elem},t.prototype.isHeap=function(){return this.root.isHeap(this.lessThan)},t.prototype.forEach=function(t){this.root.forEach(t)},t.prototype.pop=function(){if(this.empty())return null;var t=this.root.min();return this.root=this.root.removeMin(this.lessThan),t},t.prototype.reduceKey=function(t,e,n){void 0===n&&(n=null),this.root=this.root.decreaseKey(t,e,n,this.lessThan)},t.prototype.toString=function(t){return this.root.toString(t)},t.prototype.count=function(){return this.root.count()},t}(),cola,cola,cola,cola,cola,cola,cytoscape;if(function(t){var e,c,h,f,n;function i(t,e,n,i,r){this.n=t,this.es=e,this.neighbours=new Array(this.n);for(var o=this.n;o--;)this.neighbours[o]=new h(o);for(o=this.es.length;o--;){var a=this.es[o],s=n(a),l=i(a),u=r(a);this.neighbours[s].neighbours.push(new c(l,u)),this.neighbours[l].neighbours.push(new c(s,u))}}e=t.shortestpaths||(t.shortestpaths={}),c=function(t,e){this.id=t,this.distance=e},h=function(t){this.id=t,this.neighbours=[]},f=function(t,e,n){this.node=t,this.prev=e,this.d=n},i.prototype.DistanceMatrix=function(){for(var t=new Array(this.n),e=0;e<this.n;++e)t[e]=this.dijkstraNeighbours(e);return t},i.prototype.DistancesFromNode=function(t){return this.dijkstraNeighbours(t)},i.prototype.PathFromNodeToNode=function(t,e){return this.dijkstraNeighbours(t,e)},i.prototype.PathFromNodeToNodeWithPrevCost=function(t,e,n){var i=new PriorityQueue(function(t,e){return t.d<=e.d}),r=this.neighbours[t],o=new f(r,null,0),a={};for(i.push(o);!i.empty()&&(r=(o=i.pop()).node).id!==e;)for(var s=r.neighbours.length;s--;){var l=r.neighbours[s],u=this.neighbours[l.id];if(!o.prev||u.id!==o.prev.node.id){var c=u.id+","+r.id;if(!(c in a&&a[c]<=o.d)){var h=o.prev?n(o.prev.node.id,r.id,u.id):0,d=o.d+l.distance+h;a[c]=d,i.push(new f(u,o,d))}}}for(var p=[];o.prev;)o=o.prev,p.push(o.node.id);return p},i.prototype.dijkstraNeighbours=function(t,e){void 0===e&&(e=-1);for(var n=new PriorityQueue(function(t,e){return t.d<=e.d}),i=this.neighbours.length,r=new Array(i);i--;){var o=this.neighbours[i];o.d=i===t?0:Number.POSITIVE_INFINITY,o.q=n.push(o)}for(;!n.empty();){var a=n.pop();if(r[a.id]=a.d,a.id===e){for(var s=[],l=a;void 0!==l.prev;)s.push(l.prev.id),l=l.prev;return s}for(i=a.neighbours.length;i--;){var u=a.neighbours[i],c=(l=this.neighbours[u.id],a.d+u.distance);a.d!==Number.MAX_VALUE&&l.d>c&&(l.d=c,l.prev=a,n.reduceKey(l.q,l,function(t,e){return t.q=e}))}}return r},n=i,e.Calculator=n}(cola||(cola={})),function(m){var p=function(t,e,n){this.id=t,this.rect=e,this.children=n,this.leaf=void 0===n||0===n.length};m.NodeWrapper=p;var f=function(t,e,n,i,r){void 0===i&&(i=null),void 0===r&&(r=null),this.id=t,this.x=e,this.y=n,this.node=i,this.line=r};m.Vert=f;var t=(o.findMatch=function(t,e){for(var n=t.length,i=e.length,r={length:0,si:-1,ti:-1},o=new Array(n),a=0;a<n;a++){o[a]=new Array(i);for(var s=0;s<i;s++)if(t[a]===e[s]){var l=o[a][s]=0===a||0===s?1:o[a-1][s-1]+1;l>r.length&&(r.length=l,r.si=a-l+1,r.ti=s-l+1)}else o[a][s]=0}return r},o.prototype.getSequence=function(){return 0<=this.length?this.s.slice(this.si,this.si+this.length):[]},o);function o(t,e){this.s=t,this.t=e;var n=o.findMatch(t,e),i=e.slice(0).reverse(),r=o.findMatch(t,i);n.length>=r.length?(this.length=n.length,this.si=n.si,this.ti=n.ti,this.reversed=!1):(this.length=r.length,this.si=r.si,this.ti=e.length-r.ti-r.length,this.reversed=!0)}m.LongestCommonSubsequence=t;var e=(g.prototype.avg=function(t){return t.reduce(function(t,e){return t+e})/t.length},g.prototype.getGridDim=function(n){for(var t=[],e=this.leaves.slice(0,this.leaves.length);0<e.length;){var i=e[0].rect,r=e.filter(function(t){return t.rect["overlap"+n.toUpperCase()](i)});t.push(r),r.forEach(function(t){return e.splice(e.indexOf(t),1)}),r[n]=this.avg(r.map(function(t){return t.rect["c"+n]()}))}return t.sort(function(t,e){return t[n]-e[n]}),t},g.prototype.getDepth=function(t){for(var e=0;t.parent!==this.root;)e++,t=t.parent;return e},g.prototype.midPoints=function(t){for(var e=t[1]-t[0],n=[t[0]-e/2],i=1;i<t.length;i++)n.push((t[i]+t[i-1])/2);return n.push(t[t.length-1]+e/2),n},g.prototype.findLineage=function(t){for(var e=[t];t=t.parent,e.push(t),t!==this.root;);return e.reverse()},g.prototype.findAncestorPathBetween=function(t,e){for(var n=this.findLineage(t),i=this.findLineage(e),r=0;n[r]===i[r];)r++;return{commonAncestor:n[r-1],lineages:n.slice(r).concat(i.slice(r))}},g.prototype.siblingObstacles=function(t,e){var n=this,i=this.findAncestorPathBetween(t,e),r={};i.lineages.forEach(function(t){return r[t.id]={}});var o=i.commonAncestor.children.filter(function(t){return!(t in r)});return i.lineages.filter(function(t){return t.parent!==i.commonAncestor}).forEach(function(e){return o=o.concat(e.parent.children.filter(function(t){return t!==e.id}))}),o.map(function(t){return n.nodes[t]})},g.getSegmentSets=function(t,n){for(var e=[],i=0;i<t.length;i++)for(var r=t[i],o=0;o<r.length;o++){(c=r[o]).edgeid=i,c.i=o;var a=c[1][n]-c[0][n];Math.abs(a)<.1&&e.push(c)}e.sort(function(t,e){return t[0][n]-e[0][n]});for(var s=[],l=null,u=0;u<e.length;u++){var c=e[u];(!l||.1<Math.abs(c[0][n]-l.pos))&&(l={pos:c[0][n],segments:[]},s.push(l)),l.segments.push(c)}return s},g.nudgeSegs=function(o,t,a,s,e,n){var i=s.length;if(!(i<=1)){for(var r=s.map(function(t){return new m.vpsc.Variable(t[0][o])}),l=[],u=0;u<i;u++)for(var c=0;c<i;c++)if(u!==c){var h=s[u],d=s[c],p=h.edgeid,f=d.edgeid,g=-1,v=-1;"x"==o?e(p,f)&&(v=h[0][t]<h[1][t]?(g=c,u):(g=u,c)):e(p,f)&&(v=h[0][t]<h[1][t]?(g=u,c):(g=c,u)),0<=g&&l.push(new m.vpsc.Constraint(r[g],r[v],n))}new m.vpsc.Solver(r,l).solve(),r.forEach(function(t,e){var n=s[e],i=t.position();n[0][o]=n[1][o]=i;var r=a[n.edgeid];0<n.i&&(r[n.i-1][1][o]=i),n.i<r.length-1&&(r[n.i+1][0][o]=i)})}},g.nudgeSegments=function(e,n,i,r,o){for(var t=g.getSegmentSets(e,n,i),a=0;a<t.length;a++){for(var s=t[a],l=[],u=0;u<s.segments.length;u++){var c=s.segments[u];l.push({type:0,s:c,pos:Math.min(c[0][i],c[1][i])}),l.push({type:1,s:c,pos:Math.max(c[0][i],c[1][i])})}l.sort(function(t,e){return t.pos-e.pos+t.type-e.type});var h=[],d=0;l.forEach(function(t){0===t.type?(h.push(t.s),d++):d--,0==d&&(g.nudgeSegs(n,i,e,h,r,o),h=[])})}},g.prototype.routeEdges=function(t,e,n,i){var r=this,o=t.map(function(t){return r.route(n(t),i(t))}),a=m.GridRouter.orderEdges(o),s=o.map(function(t){return m.GridRouter.makeSegments(t)});return m.GridRouter.nudgeSegments(s,"x","y",a,e),m.GridRouter.nudgeSegments(s,"y","x",a,e),s},g.angleBetween2Lines=function(t,e){var n=Math.atan2(t[0].y-t[1].y,t[0].x-t[1].x),i=Math.atan2(e[0].y-e[1].y,e[0].x-e[1].x),r=n-i;return(r>Math.PI||r<-Math.PI)&&(r=i-n),r},g.isLeft=function(t,e,n){return(e.x-t.x)*(n.y-t.y)-(e.y-t.y)*(n.x-t.x)<=0},g.getOrder=function(t){for(var n={},e=0;e<t.length;e++){var i=t[e];void 0===n[i.l]&&(n[i.l]={}),n[i.l][i.r]=!0}return function(t,e){return void 0!==n[t]&&n[t][e]}},g.orderEdges=function(t){for(var e=[],n=0;n<t.length-1;n++)for(var i=n+1;i<t.length;i++){var r,o,a,s=t[n],l=t[i],u=new m.LongestCommonSubsequence(s,l);0!==u.length&&(u.reversed&&(l.reverse(),l.reversed=!0,u=new m.LongestCommonSubsequence(s,l)),u.length!==s.length&&u.length!==l.length?(u.si+u.length>=s.length||u.ti+u.length>=l.length?(r=s[u.si+1],a=s[u.si-1],o=l[u.ti-1]):(r=s[u.si+u.length-2],o=s[u.si+u.length],a=l[u.ti+u.length]),e.push(g.isLeft(r,o,a)?{l:i,r:n}:{l:n,r:i})):e.push({l:n,r:i}))}return m.GridRouter.getOrder(e)},g.makeSegments=function(t){function e(t){return{x:t.x,y:t.y}}for(var n=[],i=e(t[0]),r=1;r<t.length;r++){var o=e(t[r]),a=r<t.length-1?t[r+1]:null;a&&(s=i,l=o,u=a,Math.abs((l.x-s.x)*(u.y-s.y)-(l.y-s.y)*(u.x-s.x))<.001)||(n.push([i,o]),i=o)}var s,l,u;return n},g.prototype.route=function(t,e){var l=this,u=this.nodes[t],c=this.nodes[e];this.obstacles=this.siblingObstacles(u,c);var i={};this.obstacles.forEach(function(t){return i[t.id]=t}),this.passableEdges=this.edges.filter(function(t){var e=l.verts[t.source],n=l.verts[t.target];return!(e.node&&e.node.id in i||n.node&&n.node.id in i)});for(var n=1;n<u.ports.length;n++){var r=u.ports[0].id,o=u.ports[n].id;this.passableEdges.push({source:r,target:o,length:0})}for(n=1;n<c.ports.length;n++)r=c.ports[0].id,o=c.ports[n].id,this.passableEdges.push({source:r,target:o,length:0});var a=new m.shortestpaths.Calculator(this.verts.length,this.passableEdges,function(t){return t.source},function(t){return t.target},function(t){return t.length}).PathFromNodeToNodeWithPrevCost(u.ports[0].id,c.ports[0].id,function(t,e,n){var i=l.verts[t],r=l.verts[e],o=l.verts[n],a=Math.abs(o.x-i.x),s=Math.abs(o.y-i.y);return i.node===u&&i.node===r.node||r.node===c&&r.node===o.node?0:1<a&&1<s?1e3:0}).reverse().map(function(t){return l.verts[t]});return a.push(this.nodes[c.id].ports[0]),a.filter(function(t,e){return!(e<a.length-1&&a[e+1].node===u&&t.node===u||0<e&&t.node===c&&a[e-1].node===c)})},g);function g(t,n,e){var l=this;void 0===e&&(e=12),this.originalnodes=t,this.groupPadding=e,this.leaves=null,this.nodes=t.map(function(t,e){return new p(e,n.getBounds(t),n.getChildren(t))}),this.leaves=this.nodes.filter(function(t){return t.leaf}),this.groups=this.nodes.filter(function(t){return!t.leaf}),this.cols=this.getGridDim("x"),this.rows=this.getGridDim("y"),this.groups.forEach(function(e){return e.children.forEach(function(t){return l.nodes[t].parent=e})}),this.root={children:[]},this.nodes.forEach(function(t){void 0===t.parent&&(t.parent=l.root,l.root.children.push(t.id)),t.ports=[]}),this.backToFront=this.nodes.slice(0),this.backToFront.sort(function(t,e){return l.getDepth(t)-l.getDepth(e)}),this.backToFront.slice(0).reverse().filter(function(t){return!t.leaf}).forEach(function(t){var e=m.vpsc.Rectangle.empty();t.children.forEach(function(t){return e=e.union(l.nodes[t].rect)}),t.rect=e.inflate(l.groupPadding)});var i=this.midPoints(this.cols.map(function(t){return t.x})),r=this.midPoints(this.rows.map(function(t){return t.y})),o=i[0],a=i[i.length-1],s=r[0],u=r[r.length-1],c=this.rows.map(function(t){return{x1:o,x2:a,y1:t.y,y2:t.y}}).concat(r.map(function(t){return{x1:o,x2:a,y1:t,y2:t}})),h=this.cols.map(function(t){return{x1:t.x,x2:t.x,y1:s,y2:u}}).concat(i.map(function(t){return{x1:t,x2:t,y1:s,y2:u}})),d=c.concat(h);d.forEach(function(t){return t.verts=[]}),this.verts=[],this.edges=[],c.forEach(function(s){return h.forEach(function(t){var e=new f(l.verts.length,t.x1,s.y1);s.verts.push(e),t.verts.push(e),l.verts.push(e);for(var n=l.backToFront.length;0<n--;){var i=l.backToFront[n],r=i.rect,o=Math.abs(e.x-r.cx()),a=Math.abs(e.y-r.cy());if(o<r.width()/2&&a<r.height()/2){e.node=i;break}}})}),d.forEach(function(i){l.nodes.forEach(function(n){n.rect.lineIntersections(i.x1,i.y1,i.x2,i.y2).forEach(function(t){var e=new f(l.verts.length,t.x,t.y,n,i);l.verts.push(e),i.verts.push(e),n.ports.push(e)})});function t(t,e){return n?e.x-t.x:e.y-t.y}var n=Math.abs(i.y1-i.y2)<.1;i.verts.sort(t);for(var e=1;e<i.verts.length;e++){var r=i.verts[e-1],o=i.verts[e];r.node&&r.node===o.node&&r.node.leaf||l.edges.push({source:r.id,target:o.id,length:Math.abs(t(r,o))})}})}m.GridRouter=e}(cola||(cola={})),function(t){function i(t,e){var n={};for(var i in t)n[i]={};for(var i in e)n[i]={};return Object.keys(n).length}function r(t,e){var n=0;for(var i in t)void 0!==e[i]&&++n;return n}function o(t,i,r,o){var a=function(t,i){function r(t,e){void 0===n[t]&&(n[t]={}),n[t][e]={}}var n={};return t.forEach(function(t){var e=i.getSourceIndex(t),n=i.getTargetIndex(t);r(e,n),r(n,e)}),n}(t,o);t.forEach(function(t){var e=a[o.getSourceIndex(t)],n=a[o.getTargetIndex(t)];o.setLength(t,1+i*r(e,n))})}t.symmetricDiffLinkLengths=function(t,e,n){void 0===n&&(n=1),o(t,n,function(t,e){return Math.sqrt(i(t,e)-r(t,e))},e)},t.jaccardLinkLengths=function(t,e,n){void 0===n&&(n=1),o(t,n,function(t,e){return Math.min(Object.keys(t).length,Object.keys(e).length)<1.1?0:r(t,e)/i(t,e)},e)},t.generateDirectedEdgeConstraints=function(t,e,o,a){var n=function(t,e,n){function a(t){l[t]=h,u[t]=h,c[t]=!0,h+=1,d.push(t);for(var e=s[t],n=0;n<e.length;++n){var i=e[n];l[i]<0?(a(i),u[t]=0|Math.min(u[t],u[i])):c[i]&&(u[t]=Math.min(u[t],u[i]))}if(u[t]===l[t]){var r=[];for(n=d.length-1;0<=n;--n){var o=d[n];if(c[o]=!1,r.push(o),o===t){d.length=n;break}}p.push(r)}}for(var s=new Array(t),l=new Array(t),u=new Array(t),c=new Array(t),i=0;i<t;++i)s[i]=[],l[i]=-1,u[i]=0,c[i]=!1;for(i=0;i<e.length;++i)s[n.getSourceIndex(e[i])].push(n.getTargetIndex(e[i]));var h=0,d=[],p=[];for(i=0;i<t;++i)l[i]<0&&a(i);return p}(t,e,a),s={};n.filter(function(t){return 1<t.length}).forEach(function(e){return e.forEach(function(t){return s[t]=e})});var l=[];return e.forEach(function(t){var e=a.getSourceIndex(t),n=a.getTargetIndex(t),i=s[e],r=s[n];i&&r&&i.component===r.component||l.push({axis:o,left:e,right:n,gap:a.getMinSeparation(t)})}),l}}(cola||(cola={})),function(t){!function(l){var a=function(t,e,n){this.source=t,this.target=e,this.type=n};l.PowerEdge=a;var t=(n.prototype.initModulesFromGroup=function(t){var e=new c;this.roots.push(e);for(var n=0;n<t.leaves.length;++n){var i=t.leaves[n],r=new u(i.id);this.modules[i.id]=r,e.add(r)}if(t.groups)for(var o=0;o<t.groups.length;++o){var a=t.groups[o];e.add(new u(-1-o,new s,new s,this.initModulesFromGroup(a),!0))}return e},n.prototype.merge=function(o,a,t){void 0===t&&(t=0);var e=o.incoming.intersection(a.incoming),n=o.outgoing.intersection(a.outgoing),i=new c;i.add(o),i.add(a);var s=new u(this.modules.length,n,e,i);function r(t,i,r){t.forAll(function(t,n){t.forAll(function(t){var e=t[i];e.add(n,s),e.remove(n,o),e.remove(n,a),o[r].remove(n,t),a[r].remove(n,t)})})}return this.modules.push(s),r(n,"incoming","outgoing"),r(e,"outgoing","incoming"),this.R-=e.count()+n.count(),this.roots[t].remove(o),this.roots[t].remove(a),this.roots[t].add(s),s},n.prototype.rootMerges=function(t){void 0===t&&(t=0);for(var e=this.roots[t].modules(),n=e.length,i=new Array(n*(n-1)),r=0,o=0,a=n-1;o<a;++o)for(var s=o+1;s<n;++s){var l=e[o],u=e[s];i[r++]={nEdges:this.nEdges(l,u),a:l,b:u}}return i},n.prototype.greedyMerge=function(){for(var t=0;t<this.roots.length;++t)if(!(this.roots[t].modules().length<2)){var e=this.rootMerges(t).sort(function(t,e){return t.nEdges-e.nEdges})[0];if(!(e.nEdges>=this.R))return this.merge(e.a,e.b,t),!0}},n.prototype.nEdges=function(t,e){var n=t.incoming.intersection(e.incoming),i=t.outgoing.intersection(e.outgoing);return this.R-n.count()-i.count()},n.prototype.getGroupHierarchy=function(i){var r=this,o=[];return function n(t,i,r){t.forAll(function(t){if(t.isLeaf())i.leaves||(i.leaves=[]),i.leaves.push(t.id);else{var e=i;t.gid=r.length,t.isIsland()&&!t.predefined||(e={id:t.gid},i.groups||(i.groups=[]),i.groups.push(t.gid),r.push(e)),n(t.children,e,r)}})}(this.roots[0],{},o),this.allEdges().forEach(function(t){var e=r.modules[t.source],n=r.modules[t.target];i.push(new a(void 0===e.gid?t.source:o[e.gid],void 0===n.gid?t.target:o[n.gid],t.type))}),o},n.prototype.allEdges=function(){var t=[];return n.getEdges(this.roots[0],t),t},n.getEdges=function(t,e){t.forAll(function(t){t.getEdges(e),n.getEdges(t.children,e)})},n);function n(t,e,r,n){var o=this;if(this.linkAccessor=r,this.modules=new Array(t),this.roots=[],n)this.initModulesFromGroup(n);else{this.roots.push(new c);for(var i=0;i<t;++i)this.roots[0].add(this.modules[i]=new u(i))}this.R=e.length,e.forEach(function(t){var e=o.modules[r.getSourceIndex(t)],n=o.modules[r.getTargetIndex(t)],i=r.getType(t);e.outgoing.add(i,n),n.incoming.add(i,e)})}l.Configuration=t;var u=(e.prototype.getEdges=function(n){var i=this;this.outgoing.forAll(function(t,e){t.forAll(function(t){n.push(new a(i.id,t.id,e))})})},e.prototype.isLeaf=function(){return 0===this.children.count()},e.prototype.isIsland=function(){return 0===this.outgoing.count()&&0===this.incoming.count()},e);function e(t,e,n,i,r){void 0===e&&(e=new s),void 0===n&&(n=new s),void 0===i&&(i=new c),void 0===r&&(r=!1),this.id=t,this.outgoing=e,this.incoming=n,this.children=i,this.predefined=r}l.Module=u;var c=(i.prototype.count=function(){return Object.keys(this.table).length},i.prototype.intersection=function(t){var e=new i;return e.table=function(t,e){var n={};for(var i in t)i in e&&(n[i]=t[i]);return n}(this.table,t.table),e},i.prototype.intersectionCount=function(t){return this.intersection(t).count()},i.prototype.contains=function(t){return t in this.table},i.prototype.add=function(t){this.table[t.id]=t},i.prototype.remove=function(t){delete this.table[t.id]},i.prototype.forAll=function(t){for(var e in this.table)t(this.table[e])},i.prototype.modules=function(){var e=[];return this.forAll(function(t){t.predefined||e.push(t)}),e},i);function i(){this.table={}}l.ModuleSet=c;var s=(h.prototype.count=function(){return this.n},h.prototype.contains=function(e){var n=!1;return this.forAllModules(function(t){n||t.id!=e||(n=!0)}),n},h.prototype.add=function(t,e){(t in this.sets?this.sets[t]:this.sets[t]=new c).add(e),++this.n},h.prototype.remove=function(t,e){var n=this.sets[t];n.remove(e),0===n.count()&&delete this.sets[t],--this.n},h.prototype.forAll=function(t){for(var e in this.sets)t(this.sets[e],e)},h.prototype.forAllModules=function(e){this.forAll(function(t){return t.forAll(e)})},h.prototype.intersection=function(r){var o=new h;return this.forAll(function(t,e){if(e in r.sets){var n=t.intersection(r.sets[e]),i=n.count();0<i&&(o.sets[e]=n,o.n+=i)}}),o},h);function h(){this.sets={},this.n=0}l.LinkSets=s,l.getGroups=function(i,t,e,n){for(var r=i.length,o=new l.Configuration(r,t,e,n);o.greedyMerge(););var a=[],s=o.getGroupHierarchy(a);return a.forEach(function(n){function t(t){var e=n[t];"number"==typeof e&&(n[t]=i[e])}t("source"),t("target")}),{groups:s,powerEdges:a}}}(t.powergraph||(t.powergraph={}))}(cola||(cola={})),function(I){function u(t){t.fixed|=2,t.px=t.x,t.py=t.y}function c(t){t.fixed&=-7}function h(t){t.fixed|=4,t.px=t.x,t.py=t.y}function d(t){t.fixed&=-5}I.d3adaptor=function(){var n=d3.dispatch("start","tick","end"),i=I.adaptor({trigger:function(t){n[t.type](t)},on:function(t,e){return n.on(t,e),i},kick:function(t){d3.timer(t)},drag:function(){var t=d3.behavior.drag().origin(function(t){return t}).on("dragstart.d3adaptor",u).on("drag.d3adaptor",function(t){t.px=d3.event.x,t.py=d3.event.y,i.resume()}).on("dragend.d3adaptor",c);return arguments.length?void this.call(t):t}});return i},I.adaptor=function(t){function m(t){return"function"==typeof n?+n.call(null,t):n}function y(t){return"number"==typeof t.source?t.source:t.source.index}function b(t){return"number"==typeof t.target?t.target:t.target.index}var o,a,x={},s=t.trigger,e=t.kick,w=[1,1],n=20,$=null,i=null,r=!1,k=!0,l=!1,C=[],E=[],S=null,T=[],D=[],_=null,P=null,M=null,A=.01,N=10,p=null;x.on=t.on,x.drag=t.drag,x.dragstart=u,x.dragend=c,x.mouseover=h,x.mouseout=d,x.tick=function(){if(o<A)return l=!1,s({type:"end",alpha:o=0,stress:a}),delete a,!0;var t,e=C.length;T.length,P.locks.clear();for(var n=0;n<e;++n)if((t=C[n]).fixed){void 0!==t.px&&void 0!==t.py||(t.px=t.x,t.py=t.y);var i=[t.px,t.py];P.locks.add(n,i)}var r=P.rungeKutta();0===r?o=0:void 0!==a&&(o=Math.abs(Math.abs(a/r)-1)),a=r;for(n=0;n<e;++n)(t=C[n]).fixed?(t.x=t.px,t.y=t.py):(t.x=P.x[0][n],t.y=P.x[1][n]);s({type:"tick",alpha:o,stress:a})},x.nodes=function(t){if(arguments.length)return C=t,x;if(0===C.length&&0<T.length){var e=0;T.forEach(function(t){e=Math.max(e,t.source,t.target)}),C=new Array(++e);for(var n=0;n<e;++n)C[n]={}}return C},x.groups=function(t){return arguments.length?(S={},(E=t).forEach(function(n){void 0===n.padding&&(n.padding=1),void 0!==n.leaves&&n.leaves.forEach(function(t,e){(n.leaves[e]=C[t]).parent=n}),void 0!==n.groups&&n.groups.forEach(function(t,e){(n.groups[e]=E[t]).parent=n})}),S.leaves=C.filter(function(t){return void 0===t.parent}),S.groups=E.filter(function(t){return void 0===t.parent}),x):E},x.powerGraphGroups=function(t){var e=I.powergraph.getGroups(C,T,O,S);return this.groups(e.groups),t(e),x},x.avoidOverlaps=function(t){return arguments.length?(r=t,x):r},x.handleDisconnected=function(t){return arguments.length?(k=t,x):k},x.flowLayout=function(t,e){return arguments.length||(t="y"),M={axis:t,getMinSeparation:"number"==typeof e?function(){return e}:e},x},x.links=function(t){return arguments.length?(T=t,x):T},x.constraints=function(t){return arguments.length?(D=t,x):D},x.distanceMatrix=function(t){return arguments.length?(_=t,x):_},x.size=function(t){return arguments.length?(w=t,x):w},x.defaultNodeSize=function(t){return arguments.length?(N=t,x):N},x.linkDistance=function(t){return arguments.length?(n="function"==typeof t?t:+t,$=null,x):"function"==typeof n?n():n},x.linkType=function(t){return i=t,x},x.convergenceThreshold=function(t){return arguments.length?(A="function"==typeof t?t:+t,x):A},x.alpha=function(t){return arguments.length?(t=+t,o?o=0<t?t:0:0<t&&(l||(l=!0,s({type:"start",alpha:o=t}),e(x.tick))),x):o};var O={getSourceIndex:y,getTargetIndex:b,setLength:function(t,e){t.length=e},getType:function(t){return"function"==typeof i?i(t):0}};return x.symmetricDiffLinkLengths=function(e,t){return this.linkDistance(function(t){return e*t.length}),$=function(){I.symmetricDiffLinkLengths(T,O,t)},x},x.jaccardLinkLengths=function(e,t){return this.linkDistance(function(t){return e*t.length}),$=function(){I.jaccardLinkLengths(T,O,t)},x},x.start=function(){var t=this.nodes().length,e=t+2*E.length,n=(T.length,w[0]),i=w[1];$&&$();var r=new Array(e),o=new Array(e);new Array(e);var a,s=null,l=this.avoidOverlaps();C.forEach(function(t,e){t.index=e,void 0===t.x&&(t.x=n/2,t.y=i/2),r[e]=t.x,o[e]=t.y}),_?a=_:(a=new I.shortestpaths.Calculator(e,T,y,b,m).DistanceMatrix(),s=I.Descent.createSquareMatrix(e,function(){return 2}),T.forEach(function(t){var e=y(t),n=b(t);s[e][n]=s[n][e]=1}));var u=I.Descent.createSquareMatrix(e,function(t,e){return a[t][e]});if(S&&void 0!==S.groups){var c=t;E.forEach(function(){s[c][c+1]=s[c+1][c]=1e-6,u[c][c+1]=u[c+1][c]=.1,r[c]=0,o[c++]=0,r[c]=0,o[c++]=0})}else S={leaves:C,groups:[]};var h=D||[];M&&(O.getMinSeparation=M.getMinSeparation,h=h.concat(I.generateDirectedEdgeConstraints(t,T,M.axis,O)));var d=0<arguments.length?arguments[0]:0,p=1<arguments.length?arguments[1]:0,f=2<arguments.length?arguments[2]:0;this.avoidOverlaps(!1),(P=new I.Descent([r,o],u)).locks.clear();for(c=0;c<t;++c){var g=C[c];if(g.fixed){g.px=g.x,g.py=g.y;var v=[g.x,g.y];P.locks.add(c,v)}}return P.threshold=A,P.run(d),0<h.length&&(P.project=new I.vpsc.Projection(C,E,S,h).projectFunctions()),P.run(p),this.avoidOverlaps(l),l&&(C.forEach(function(t,e){t.x=r[e],t.y=o[e]}),P.project=new I.vpsc.Projection(C,E,S,h,!0).projectFunctions(),C.forEach(function(t,e){r[e]=t.x,o[e]=t.y})),P.G=s,P.run(f),T.forEach(function(t){"number"==typeof t.source&&(t.source=C[t.source]),"number"==typeof t.target&&(t.target=C[t.target])}),C.forEach(function(t,e){t.x=r[e],t.y=o[e]}),!_&&k&&(I.applyPacking(I.separateGraphs(C,T),n,i,N),C.forEach(function(t,e){P.x[0][e]=t.x,P.x[1][e]=t.y})),x.resume()},x.resume=function(){return x.alpha(.1)},x.stop=function(){return x.alpha(0)},x.prepareEdgeRouting=function(e){p=new I.geom.TangentVisibilityGraph(C.map(function(t){return t.bounds.inflate(-e).vertices()}))},x.routeEdge=function(t,e){var n=[],i=new I.geom.TangentVisibilityGraph(p.P,{V:p.V,E:p.E}),r={x:t.source.x,y:t.source.y},o={x:t.target.x,y:t.target.y},a=i.addPoint(r,t.source.id),s=i.addPoint(o,t.target.id);i.addEdgeIfVisible(r,o,t.source.id,t.target.id),void 0!==e&&e(i);var l=new I.shortestpaths.Calculator(i.V.length,i.E,function(t){return t.source.id},function(t){return t.target.id},function(t){return t.length()}).PathFromNodeToNode(a.id,s.id);if(1===l.length||l.length===i.V.length)I.vpsc.makeEdgeBetween(t,t.source.innerBounds,t.target.innerBounds,5),n=[{x:t.sourceIntersection.x,y:t.sourceIntersection.y},{x:t.arrowStart.x,y:t.arrowStart.y}];else{for(var u=l.length-2,c=i.V[l[u]].p,h=i.V[l[0]].p,d=(n=[t.source.innerBounds.rayIntersection(c.x,c.y)],u);0<=d;--d)n.push(i.V[l[d]].p);n.push(I.vpsc.makeEdgeTo(h,t.target.innerBounds,5))}return n},x.linkId=function(t){return y(t)+"-"+b(t)},x}}(cola||(cola={})),RBTree=function(){var i=function(t){var e=i.m[t];if(e.mod)return e.mod.exports;var n=e.mod={exports:{}};return e(n,n.exports),n.exports};return i.m={},i.m["./treebase"]=function(t){function e(){}function n(t){this._tree=t,this._ancestors=[],this._cursor=null}e.prototype.clear=function(){this._root=null,this.size=0},e.prototype.find=function(t){for(var e=this._root;null!==e;){var n=this._comparator(t,e.data);if(0===n)return e.data;e=e.get_child(0<n)}return null},e.prototype.findIter=function(t){for(var e=this._root,n=this.iterator();null!==e;){var i=this._comparator(t,e.data);if(0===i)return n._cursor=e,n;n._ancestors.push(e),e=e.get_child(0<i)}return null},e.prototype.lowerBound=function(t){return this._bound(t,this._comparator)},e.prototype.upperBound=function(t){var n=this._comparator;return this._bound(t,function(t,e){return n(e,t)})},e.prototype.min=function(){var t=this._root;if(null===t)return null;for(;null!==t.left;)t=t.left;return t.data},e.prototype.max=function(){var t=this._root;if(null===t)return null;for(;null!==t.right;)t=t.right;return t.data},e.prototype.iterator=function(){return new n(this)},e.prototype.each=function(t){for(var e,n=this.iterator();null!==(e=n.next());)t(e)},e.prototype.reach=function(t){for(var e,n=this.iterator();null!==(e=n.prev());)t(e)},e.prototype._bound=function(t,e){for(var n=this._root,i=this.iterator();null!==n;){var r=this._comparator(t,n.data);if(0===r)return i._cursor=n,i;i._ancestors.push(n),n=n.get_child(0<r)}for(var o=i._ancestors.length-1;0<=o;--o)if(0<e(t,(n=i._ancestors[o]).data))return i._cursor=n,i._ancestors.length=o,i;return i._ancestors.length=0,i},n.prototype.data=function(){return null!==this._cursor?this._cursor.data:null},n.prototype.next=function(){if(null===this._cursor){var t=this._tree._root;null!==t&&this._minNode(t)}else if(null===this._cursor.right){var e;do{if(e=this._cursor,!this._ancestors.length){this._cursor=null;break}this._cursor=this._ancestors.pop()}while(this._cursor.right===e)}else this._ancestors.push(this._cursor),this._minNode(this._cursor.right);return null!==this._cursor?this._cursor.data:null},n.prototype.prev=function(){if(null===this._cursor){var t=this._tree._root;null!==t&&this._maxNode(t)}else if(null===this._cursor.left){var e;do{if(e=this._cursor,!this._ancestors.length){this._cursor=null;break}this._cursor=this._ancestors.pop()}while(this._cursor.left===e)}else this._ancestors.push(this._cursor),this._maxNode(this._cursor.left);return null!==this._cursor?this._cursor.data:null},n.prototype._minNode=function(t){for(;null!==t.left;)this._ancestors.push(t),t=t.left;this._cursor=t},n.prototype._maxNode=function(t){for(;null!==t.right;)this._ancestors.push(t),t=t.right;this._cursor=t},t.exports=e},i.m.__main__=function(t){function p(t){this.data=t,this.left=null,this.right=null,this.red=!0}function e(t){this._root=null,this._comparator=t,this.size=0}function f(t){return null!==t&&t.red}function g(t,e){var n=t.get_child(!e);return t.set_child(!e,n.get_child(e)),n.set_child(e,t),t.red=!0,n.red=!1,n}function v(t,e){return t.set_child(!e,g(t.get_child(!e),!e)),g(t,e)}var n=i("./treebase");p.prototype.get_child=function(t){return t?this.right:this.left},p.prototype.set_child=function(t,e){t?this.right=e:this.left=e},(e.prototype=new n).insert=function(t){var e=!1;if(null===this._root)this._root=new p(t),e=!0,this.size++;else{var n=new p(void 0),i=0,r=0,o=null,a=n,s=null,l=this._root;for(a.right=this._root;;){if(null===l?(l=new p(t),s.set_child(i,l),e=!0,this.size++):f(l.left)&&f(l.right)&&(l.red=!0,l.left.red=!1,l.right.red=!1),f(l)&&f(s)){var u=a.right===o;l===s.get_child(r)?a.set_child(u,g(o,!r)):a.set_child(u,v(o,!r))}var c=this._comparator(l.data,t);if(0===c)break;r=i,i=c<0,null!==o&&(a=o),o=s,l=(s=l).get_child(i)}this._root=n.right}return this._root.red=!1,e},e.prototype.remove=function(t){if(null===this._root)return!1;var e=new p(void 0),n=e;n.right=this._root;for(var i=null,r=null,o=null,a=1;null!==n.get_child(a);){var s=a;r=i,n=(i=n).get_child(a);var l=this._comparator(t,n.data);if(a=0<l,0===l&&(o=n),!f(n)&&!f(n.get_child(a)))if(f(n.get_child(!a))){var u=g(n,a);i.set_child(s,u),i=u}else if(!f(n.get_child(!a))){var c=i.get_child(!s);if(null!==c)if(f(c.get_child(!s))||f(c.get_child(s))){var h=r.right===i;f(c.get_child(s))?r.set_child(h,v(i,s)):f(c.get_child(!s))&&r.set_child(h,g(i,s));var d=r.get_child(h);d.red=!0,n.red=!0,d.left.red=!1,d.right.red=!1}else i.red=!1,c.red=!0,n.red=!0}}return null!==o&&(o.data=n.data,i.set_child(i.right===n,n.get_child(null===n.left)),this.size--),this._root=e.right,null!==this._root&&(this._root.red=!1),null!==o},t.exports=e},i("__main__")}(window),function(t){var v={PADDING:10};v.GOLDEN_SECTION=(1+Math.sqrt(5))/2,v.FLOAT_EPSILON=1e-4,v.MAX_INERATIONS=100,t.applyPacking=function(t,e,n,s,i){function p(t,e){g=[],d=a;for(var n=h=c=0;n<t.length;n++){r(t[n],e)}return Math.abs(c/h-i)}function r(t,e){for(var n=void 0,i=0;i<g.length;i++)if(g[i].space_left>=t.height&&g[i].x+g[i].width+t.width+v.PADDING-e<=v.FLOAT_EPSILON){n=g[i];break}g.push(t),void 0!==n?(t.x=n.x+n.width+v.PADDING,t.y=n.bottom,t.space_left=t.height,t.bottom=t.y,n.space_left-=t.height+v.PADDING,n.bottom+=t.height+v.PADDING):
```
