## Supplementary material for "Tensor-decomposition-based unsupervised feature extraction applied to prostate cancer multiomics data": output from metascape: custom.js

```
$( document ).ready(function(){
  // Custom Cytoscape.JS code goes here.
  
  // Example: add linkouts to nodes that opens the "href" node attribute on click
  // cy.on('tap', 'node', function(){
  //   try { // your browser may block popups
  //     window.open( this.data('href') );
  //   } catch(e){ // fall back on url change
  //     window.location.href = this.data('href');
  //   }
  // });
  
  // For more options, check out http://js.cytoscape.org/

cy.on('select', 'node', function(e) {
    var node = e.cyTarget;
    if (IS_PPI) {
        node.css({ 'content' : node.data('Symbol')});
    } else {
        node.css({ 'content' : node.data('Description')});
    }
});

cy.on('unselect', 'node', function(e) {
    var node = e.cyTarget;
    node.css({ 'content' : ''});
});

});
```
