## Supplementary material for "Tensor-decomposition-based unsupervised feature extraction applied to prostate cancer multiomics data": output from metascape: initialize.js

```
var gup = function(name) {
               name = name.replace(/[\[]/, "\\[").replace(/[\]]/, "\\]");
               var regexS = "[\?&]" + name + "=([^&#]*)";
               var regex = new RegExp(regexS);
               var results = regex.exec(decodeURI(window.location.href));
               return results == null ? null : results[1];
}

var my_loadjs = function (url) {
    jQuery.ajax({
        url: url,
        dataType: 'script',
        //success: callback,
        async: false 
    });
}
var DEFAULT_NETWORK_NAME=null;
var DEFAULT_STYLE_NAME=null;
var IS_PPI=null;
(function init() {
    var session_id=gup('session_id');
    if (session_id!=null) {
        //Metascape online browsing
        //load network and styles
        var IS_PPI=gup("isPPI");
        if (IS_PPI) {
            my_loadjs("/gp_server/get_file?session_id="+session_id+"&file_name=Enrichment_PPI/PPINetwork.js");
            my_loadjs("/gp_server/get_file?session_id="+session_id+"&file_name=Enrichment_PPI/PPINetwork.style.js");
        } else {
            my_loadjs("/gp_server/get_file?session_id="+session_id+"&file_name=Enrichment_GO/GONetwork.js");
            my_loadjs("/gp_server/get_file?session_id="+session_id+"&file_name=Enrichment_GO/GONetwork.style.js");
        }
    }

    var network_name=gup('Network')
    DEFAULT_NETWORK_NAME = (network_name==null || !(network_name in networks))? null : network_name;

    var style_name=gup('Style');
    var style_names={};
    styles.forEach(function(b) {
        style_names[b.title]=true;
    });
    DEFAULT_STYLE_NAME= (style_name==null || !(style_name in style_names))? styles[0].title : style_name;

})();
```
