## Supplementary material for "Tensor-decomposition-based unsupervised feature extraction applied to prostate cancer multiomics data": output from metascape: ColorByCluster.pdf

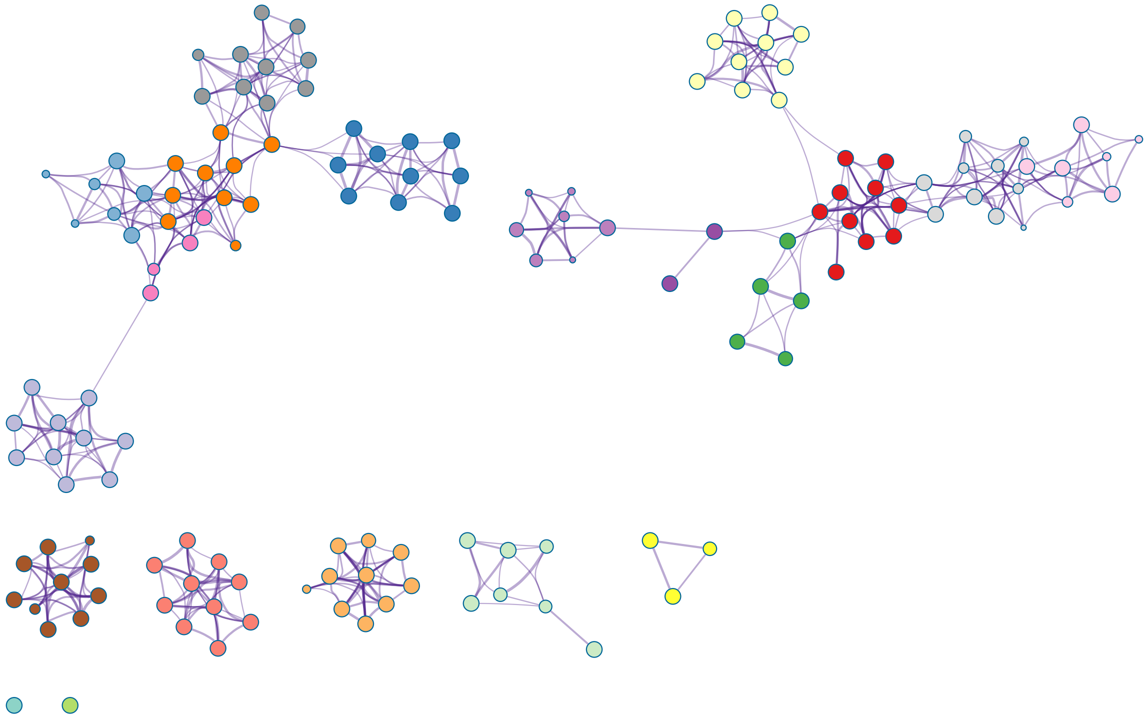

chordate embryonic development
blood vessel development
gland development
epithelial cell differentiation
leukocyte differentiation
fat cell differentiation
response to wounding
negative regulation of cell differentiation
cell junction assembly
Transcriptional misregulation in cancer
muscle structure development
regulation of neuron differentiation
regulation of cellular amide metabolic process
myeloid cell differentiation
cellular response to steroid hormone stimulus
negative regulation of cell proliferation
ossification
connective tissue development
regulation of epithelial cell differentiation
Signaling by Nuclear Receptors

created by
http://metascape.org
