## Supplementary material for "Tensor-decomposition-based unsupervised feature extraction applied to prostate cancer multiomics data": output from metascape: GONetwork.js

```
var networks={"GONetwork": {"format_version": "1.0", "generated_by": "cytoscape-3.8.0", "target_cytoscapejs_version": "~2.1", "data": {"shared_name": "GONetwork", "name": "GONetwork", "SUID": 579188, "__Annotations": [], "selected": true}, "elements": {"nodes": [{"data": {"id": "579341", "Enrichment": 2.161428042515774, "BestLogPInGroup": -11.981654863057864, "_PATTERN_": "M1", "_GeneInGOAndHitList": 92.0, "Hits": "ABL1|AKT1|BCL2L1|ZFP36L1|RUNX2|CEBPB|CFL1|COL1A1|DLX1|DLX2|DVL3|ECE1|EGFR|EPHA2|ELF3|FOXF1|GATA2|FOXA1|HOXA3|HOXA5|HOXA6|HOXA7|HOXA9|HOXA11|HOXC4|HOXC6|HOXD10|ISL1|ITPK1|JUNB|KRT8|LFNG|SMAD3|MBNL1|MGAT1|MMP14|MYH9|NKX3-1|NOTCH1|ROR2|PBX1|PDGFB|PTCH1|PTK7|PTPRR|RARA|RARG|RXRA|SDC4|STIL|SOX4|SP3|SRF|TBX3|HNF1B|TRIP6|PCGF2|ZFAND5|FOSL1|YBX3|DCHS1|SH3PXD2A|ZEB2|SEC24D|SCO2|LPAR6|PEMT|EMG1|GNA13|COBL|USP22|HEY2|MTHFD1L|SLC39A1|SEMA4C|MESP1|XAB2|RTN4|SHROOM3|CAMSAP3|NXN|GRHL2|CMIP|EGFL8|HES7|UNK|PLCD3|MESP2|RUNDC1|ARL13B|SMIM14|ST8SIA6", "Description": "embryo development ending in birth or egg hatching", "_MEMBER_Input_ID": 1.0, "_GeneInHitList": 1562.0, "Category": "GO Biological Processes", "Log_q_value_": -7.675226902393069, "Symbol": "GO:0009792", "LogP": -11.693302966038864, "_RANK_": 1.0, "FirstInGroupByEnrichment": 0.0, "_TotalGeneInLibrary": 24367.0, "_GeneInGO": 664.0, "shared_name": "144", "_InGO": 5.8898847631242, "Gene": "GO:0009792", "_COUNT_001": 92.0, "SUID": 579341, "id_original": "144", "selected": false, "CategoryID": 19.0, "STDV_InGO": 0.5957047138803491, "EvidenceCutoff": 0.0, "PARENT_GO": "19_GO:0032502 developmental process", "FirstInGroupByLogP": 0.0, "GROUP_ID": 1.0, "name": "144", "BestEnrichmentInGroup": 2.5999786598378147, "CLUSTER_LABEL": " ", "GeneID": "25|207|598|677|860|1051|1072|1277|1745|1746|1857|1889|1956|1969|1999|2294|2624|3169|3200|3202|3203|3204|3205|3207|3221|3223|3236|3670|3705|3726|3856|3955|4088|4154|4245|4323|4627|4824|4851|4920|5087|5155|5727|5754|5801|5914|5916|6256|6385|6491|6659|6670|6722|6926|6928|7205|7703|7763|8061|8531|8642|9644|9839|9871|9997|10161|10400|10436|10672|23242|23326|23493|25902|27173|54910|55897|56949|57142|57619|57662|64359|79977|80790|80864|84667|85451|113026|145873|146923|200894|201895|338596", "_LogP_Input_ID": -11.693302966038864, "Z_score": 7.941312487036475, "GiniIndex": 0.0, "_COUNT_001_PCT": 100.0}, "position": {"x": 1006.6826779717153, "y": 145.7528796738128}, "selected": false}, {"data": {"id": "579340", "Enrichment": 2.0279833546734953, "BestLogPInGroup": -11.981654863057864, "_PATTERN_": "M1", "_GeneInGOAndHitList": 78.0, "Hits": "ABL1|ALDH1A3|ZFP36L1|RUNX2|CFL1|COL4A2|DLX2|DVL3|ECE1|EPHA2|ETS2|EXT2|EYA2|FOXF1|GATA2|HOXA3|HOXA5|HOXA6|HOXA7|HOXA9|HOXA10|HOXA11|HOXC4|HOXD10|HOXD12|HOXD13|HPN|ID2|ITGA3|ITGA7|AFF3|LAMA3|SMAD3|MBNL1|MMP14|MYO6|NOTCH1|ROR2|OTX1|PBX1|PBX2|PRKAR1A|PTCH1|PTK7|RARA|RARB|RARG|SDC4|STIL|SOX4|SP3|SRF|TBX3|HNF1B|UGDH|PCGF2|ZBTB16|ZEB2|TRIOBP|KDM6B|PHLDB1|COBL|MTHFD1L|LRIG1|SLC39A1|PAF1|SEMA4C|CCDC40|MESP1|SHROOM3|PDZD7|GRHL2|SLC44A4|MESP2|ARL13B|SP8|TPRN|MIR200C", "Description": "embryonic morphogenesis", "_MEMBER_Input_ID": 1.0, "_GeneInHitList": 1562.0, "Category": "GO Biological Processes", "Log_q_value_": -5.607011768762616, "Symbol": "GO:0048598", "LogP": -8.695668906694118, "_RANK_": 1.0, "FirstInGroupByEnrichment": 0.0, "_TotalGeneInLibrary": 24367.0, "_GeneInGO": 600.0, "shared_name": "143", "_InGO": 4.99359795134443, "Gene": "GO:0048598", "_COUNT_001": 78.0, "SUID": 579340, "id_original": "143", "selected": false, "CategoryID": 19.0, "STDV_InGO": 0.5511156038954877, "EvidenceCutoff": 0.0, "PARENT_GO": "19_GO:0032502 developmental process", "FirstInGroupByLogP": 0.0, "GROUP_ID": 1.0, "name": "143", "BestEnrichmentInGroup": 2.5999786598378147, "CLUSTER_LABEL": " ", "GeneID": "25|220|677|860|1072|1284|1746|1857|1889|1969|2114|2132|2139|2294|2624|3200|3202|3203|3204|3205|3206|3207|3221|3236|3238|3239|3249|3398|3675|3679|3899|3909|4088|4154|4323|4646|4851|4920|5013|5087|5089|5573|5727|5754|5914|5915|5916|6385|6491|6659|6670|6722|6926|6928|7358|7703|7704|9839|11078|23135|23187|23242|25902|26018|27173|54623|54910|55036|55897|57619|79955|79977|80736|145873|200894|221833|286262|406985", "_LogP_Input_ID": -8.695668906694118, "Z_score": 6.672554367117394, "GiniIndex": 0.0, "_COUNT_001_PCT": 100.0}, "position": {"x": 1071.071689354772, "y": 39.55081515965264}, "selected": false}, {"data": {"id": "579339", "Enrichment": 2.0986374832771597, "BestLogPInGroup": -11.981654863057864, "_PATTERN_": "M1", "_GeneInGOAndHitList": 60.0, "Hits": "BMPR1B|DLX1|DLX2|DNAH5|ETS2|FOXF1|FOXA1|HOXA3|HOXA5|HOXA6|HOXA7|HOXA9|HOXA10|HOXA11|HOXC4|HOXC6|HOXC8|HOXD10|HOXD11|HOXD12|HOXD13|ISL1|LFNG|SMAD3|NBL1|NKX3-1|NOTCH1|ROR2|OTX1|PBX1|PBX2|PBX3|PTCH1|RARG|RFNG|STIL|SIM2|SMARCD3|SRF|TBX3|HNF1B|PCGF2|ZBTB16|BTG2|BHLHE40|DCHS1|SYNGAP1|ZEB2|CITED2|COBL|HEY2|IRX4|CCDC40|MESP1|SHROOM3|HES7|MESP2|ARL13B|SP8|NEK8", "Description": "pattern specification process", "_MEMBER_Input_ID": 1.0, "_GeneInHitList": 1562.0, "Category": "GO Biological Processes", "Log_q_value_": -4.626523481148543, "Symbol": "GO:0007389", "LogP": -7.365845943841509, "_RANK_": 1.0, "FirstInGroupByEnrichment": 0.0, "_TotalGeneInLibrary": 24367.0, "_GeneInGO": 446.0, "shared_name": "142", "_InGO": 3.8412291933418694, "Gene": "GO:0007389", "_COUNT_001": 60.0, "SUID": 579339, "id_original": "142", "selected": false, "CategoryID": 19.0, "STDV_InGO": 0.4862829551112782, "EvidenceCutoff": 0.0, "PARENT_GO": "19_GO:0032502 developmental process", "FirstInGroupByLogP": 0.0, "GROUP_ID": 1.0, "name": "142", "BestEnrichmentInGroup": 2.5999786598378147, "CLUSTER_LABEL": " ", "GeneID": "658|1745|1746|1767|2114|2294|3169|3200|3202|3203|3204|3205|3206|3207|3221|3223|3224|3236|3237|3238|3239|3670|3955|4088|4681|4824|4851|4920|5013|5087|5089|5090|5727|5916|5986|6491|6493|6604|6722|6926|6928|7703|7704|7832|8553|8642|8831|9839|10370|23242|23493|50805|55036|55897|57619|84667|145873|200894|221833|284086", "_LogP_Input_ID": -7.365845943841509, "Z_score": 6.128437656096849, "GiniIndex": 0.0, "_COUNT_001_PCT": 100.0}, "position": {"x": 999.2244527214712, "y": 33.44446139988702}, "selected": false}, {"data": {"id": "579338", "Enrichment": 2.576125644609945, "BestLogPInGroup": -11.981654863057864, "_PATTERN_": "M1", "_GeneInGOAndHitList": 36.0, "Hits": "ETS2|FOXF1|HOXA3|HOXA5|HOXA6|HOXA7|HOXA9|HOXA10|HOXA11|HOXC4|HOXC6|HOXC8|HOXD10|HOXD13|LFNG|SMAD3|NKX3-1|NOTCH1|ROR2|OTX1|PBX1|PBX3|RARG|SRF|TBX3|HNF1B|PCGF2|ZBTB16|BTG2|BHLHE40|ZEB2|COBL|HEY2|MESP1|HES7|MESP2", "Description": "anterior/posterior pattern specification", "_MEMBER_Input_ID": 1.0, "_GeneInHitList": 1562.0, "Category": "GO Biological Processes", "Log_q_value_": -4.210600614899919, "Symbol": "GO:0009952", "LogP": -6.800641497674141, "_RANK_": 1.0, "FirstInGroupByEnrichment": 0.0, "_TotalGeneInLibrary": 24367.0, "_GeneInGO": 218.0, "shared_name": "141", "_InGO": 2.3047375160051216, "Gene": "GO:0009952", "_COUNT_001": 36.0, "SUID": 579338, "id_original": "141", "selected": false, "CategoryID": 19.0, "STDV_InGO": 0.37967060373244593, "EvidenceCutoff": 0.0, "PARENT_GO": "19_GO:0032502 developmental process", "FirstInGroupByLogP": 0.0, "GROUP_ID": 1.0, "name": "141", "BestEnrichmentInGroup": 2.5999786598378147, "CLUSTER_LABEL": " ", "GeneID": "2114|2294|3200|3202|3203|3204|3205|3206|3207|3221|3223|3224|3236|3239|3955|4088|4824|4851|4920|5013|5087|5090|5916|6722|6926|6928|7703|7704|7832|8553|9839|23242|23493|55897|84667|145873", "_LogP_Input_ID": -6.800641497674141, "Z_score": 6.117684879523703, "GiniIndex": 0.0, "_COUNT_001_PCT": 100.0}, "position": {"x": 1052.7113454216665, "y": 86.5597644394378}, "selected": false}, {"data": {"id": "579337", "Enrichment": 2.0332417384799086, "BestLogPInGroup": -11.981654863057864, "_PATTERN_": "M1", "_GeneInGOAndHitList": 58.0, "Hits": "AKT1|ALDH1A3|ZFP36L1|RUNX2|CEBPB|DLX2|EGFR|EPHA2|FOXF1|GATA2|HOXA3|HOXA5|HOXA6|HOXA7|HOXA9|HOXA11|HOXC4|HOXD10|HPN|ID2|ID3|JUNB|KRT8|SMAD3|MMP14|MYO6|NOTCH1|ROR2|OTX1|PBX1|PDGFB|PTCH1|PTK7|RARA|RARB|RARG|STIL|SP3|SRF|TBX3|HNF1B|TEAD3|PCGF2|CITED2|TRIOBP|COBL|HEY2|MTHFD1L|LRIG1|SLC39A1|CCDC40|MESP1|PDZD7|GRHL2|SLC44A4|PLCD3|ARL13B|TPRN", "Description": "embryonic organ development", "_MEMBER_Input_ID": 1.0, "_GeneInHitList": 1562.0, "Category": "GO Biological Processes", "Log_q_value_": -4.11736101884013, "Symbol": "GO:0048568", "LogP": -6.667051135715807, "_RANK_": 1.0, "FirstInGroupByEnrichment": 0.0, "_TotalGeneInLibrary": 24367.0, "_GeneInGO": 445.0, "shared_name": "140", "_InGO": 3.7131882202304736, "Gene": "GO:0048568", "_COUNT_001": 58.0, "SUID": 579337, "id_original": "140", "selected": false, "CategoryID": 19.0, "STDV_InGO": 0.47842775961251943, "EvidenceCutoff": 0.0, "PARENT_GO": "19_GO:0032502 developmental process", "FirstInGroupByLogP": 0.0, "GROUP_ID": 1.0, "name": "140", "BestEnrichmentInGroup": 2.5999786598378147, "CLUSTER_LABEL": " ", "GeneID": "207|220|677|860|1051|1746|1956|1969|2294|2624|3200|3202|3203|3204|3205|3207|3221|3236|3249|3398|3399|3726|3856|4088|4323|4646|4851|4920|5013|5087|5155|5727|5754|5914|5915|5916|6491|6670|6722|6926|6928|7005|7703|10370|11078|23242|23493|25902|26018|27173|55036|55897|79955|79977|80736|113026|200894|286262", "_LogP_Input_ID": -6.667051135715807, "Z_score": 5.757060592066372, "GiniIndex": 0.0, "_COUNT_001_PCT": 100.0}, "position": {"x": 953.3718716973012, "y": 129.0922351425628}, "selected": false}, {"data": {"id": "579336", "Enrichment": 2.1593043107127614, "BestLogPInGroup": -11.981654863057864, "_PATTERN_": "M1", "_GeneInGOAndHitList": 49.0, "Hits": "BMPR1B|DLX1|DLX2|ETS2|FOXF1|FOXA1|HOXA3|HOXA5|HOXA6|HOXA7|HOXA9|HOXA10|HOXA11|HOXC4|HOXC6|HOXC8|HOXD10|HOXD11|HOXD13|ISL1|LFNG|SMAD3|NBL1|NKX3-1|NOTCH1|ROR2|OTX1|PBX1|PBX2|PBX3|PTCH1|RARG|SMARCD3|SRF|TBX3|HNF1B|PCGF2|ZBTB16|BTG2|BHLHE40|ZEB2|CITED2|COBL|HEY2|MESP1|HES7|MESP2|ARL13B|SP8", "Description": "regionalization", "_MEMBER_Input_ID": 1.0, "_GeneInHitList": 1562.0, "Category": "GO Biological Processes", "Log_q_value_": -3.98861494291392, "Symbol": "GO:0003002", "LogP": -6.508380452770115, "_RANK_": 1.0, "FirstInGroupByEnrichment": 0.0, "_TotalGeneInLibrary": 24367.0, "_GeneInGO": 354.0, "shared_name": "139", "_InGO": 3.1370038412291934, "Gene": "GO:0003002", "_COUNT_001": 49.0, "SUID": 579336, "id_original": "139", "selected": false, "CategoryID": 19.0, "STDV_InGO": 0.44105825989215863, "EvidenceCutoff": 0.0, "PARENT_GO": "19_GO:0032502 developmental process", "FirstInGroupByLogP": 0.0, "GROUP_ID": 1.0, "name": "139", "BestEnrichmentInGroup": 2.5999786598378147, "CLUSTER_LABEL": " ", "GeneID": "658|1745|1746|2114|2294|3169|3200|3202|3203|3204|3205|3206|3207|3221|3223|3224|3236|3237|3239|3670|3955|4088|4681|4824|4851|4920|5013|5087|5089|5090|5727|5916|6604|6722|6926|6928|7703|7704|7832|8553|9839|10370|23242|23493|55897|84667|145873|200894|221833", "_LogP_Input_ID": -6.508380452770115, "Z_score": 5.750339255582647, "GiniIndex": 0.0, "_COUNT_001_PCT": 100.0}, "position": {"x": 1085.2232320183462, "y": 172.65429263768}, "selected": false}, {"data": {"id": "579335", "Enrichment": 2.1839820742637643, "BestLogPInGroup": -11.981654863057864, "_PATTERN_": "M1", "_GeneInGOAndHitList": 42.0, "Hits": "ALDH1A3|RUNX2|DLX2|EPHA2|FOXF1|GATA2|HOXA3|HOXA5|HOXA6|HOXA7|HOXA9|HOXA11|HOXC4|HOXD10|HPN|ID2|SMAD3|MMP14|MYO6|NOTCH1|ROR2|OTX1|PTK7|RARB|RARG|STIL|SP3|SRF|TBX3|HNF1B|PCGF2|TRIOBP|MTHFD1L|LRIG1|SLC39A1|CCDC40|MESP1|PDZD7|GRHL2|SLC44A4|ARL13B|TPRN", "Description": "embryonic organ morphogenesis", "_MEMBER_Input_ID": 1.0, "_GeneInHitList": 1562.0, "Category": "GO Biological Processes", "Log_q_value_": -3.462221847143521, "Symbol": "GO:0048562", "LogP": -5.80360430116445, "_RANK_": 1.0, "FirstInGroupByEnrichment": 0.0, "_TotalGeneInLibrary": 24367.0, "_GeneInGO": 300.0, "shared_name": "138", "_InGO": 2.6888604353393086, "Gene": "GO:0048562", "_COUNT_001": 42.0, "SUID": 579335, "id_original": "138", "selected": false, "CategoryID": 19.0, "STDV_InGO": 0.40928412025975003, "EvidenceCutoff": 0.0, "PARENT_GO": "19_GO:0032502 developmental process", "FirstInGroupByLogP": 0.0, "GROUP_ID": 1.0, "name": "138", "BestEnrichmentInGroup": 2.5999786598378147, "CLUSTER_LABEL": " ", "GeneID": "220|860|1746|1969|2294|2624|3200|3202|3203|3204|3205|3207|3221|3236|3249|3398|4088|4323|4646|4851|4920|5013|5754|5915|5916|6491|6670|6722|6926|6928|7703|11078|25902|26018|27173|55036|55897|79955|79977|80736|200894|286262", "_LogP_Input_ID": -5.80360430116445, "Z_score": 5.400231427051371, "GiniIndex": 0.0, "_COUNT_001_PCT": 100.0}, "position": {"x": 1036.0821795338338, "y": 182.3758044785003}, "selected": false}, {"data": {"id": "579334", "Enrichment": 1.9854382493306948, "BestLogPInGroup": -11.981654863057864, "_PATTERN_": "M1", "_GeneInGOAndHitList": 49.0, "Hits": "AKT1|BCL2L1|ZFP36L1|CEBPB|EGFR|ELF3|FOXF1|GATA2|JUNB|KRT8|SMAD3|MBNL1|MGAT1|MYH9|NOTCH1|PDGFB|PTCH1|PTPRR|RXRA|STIL|SP3|SRF|TBX3|HNF1B|PCGF2|ZFAND5|FOSL1|YBX3|SH3PXD2A|SEC24D|SCO2|LPAR6|PEMT|EMG1|GNA13|HEY2|SLC39A1|XAB2|RTN4|CAMSAP3|NXN|GRHL2|CMIP|EGFL8|UNK|PLCD3|RUNDC1|SMIM14|ST8SIA6", "Description": "in utero embryonic development", "_MEMBER_Input_ID": 1.0, "_GeneInHitList": 1562.0, "Category": "GO Biological Processes", "Log_q_value_": -3.2169893165144963, "Symbol": "GO:0001701", "LogP": -5.440595943313896, "_RANK_": 1.0, "FirstInGroupByEnrichment": 0.0, "_TotalGeneInLibrary": 24367.0, "_GeneInGO": 385.0, "shared_name": "137", "_InGO": 3.1370038412291934, "Gene": "GO:0001701", "_COUNT_001": 49.0, "SUID": 579334, "id_original": "137", "selected": false, "CategoryID": 19.0, "STDV_InGO": 0.44105825989215863, "EvidenceCutoff": 0.0, "PARENT_GO": "19_GO:0032502 developmental process", "FirstInGroupByLogP": 0.0, "GROUP_ID": 1.0, "name": "137", "BestEnrichmentInGroup": 2.5999786598378147, "CLUSTER_LABEL": " ", "GeneID": "207|598|677|1051|1956|1999|2294|2624|3726|3856|4088|4154|4245|4627|4851|5155|5727|5801|6256|6491|6670|6722|6926|6928|7703|7763|8061|8531|9644|9871|9997|10161|10400|10436|10672|23493|27173|56949|57142|57662|64359|79977|80790|80864|85451|113026|146923|201895|338596", "_LogP_Input_ID": -5.440595943313896, "Z_score": 5.10075750288195, "GiniIndex": 0.0, "_COUNT_001_PCT": 100.0}, "position": {"x": 982.3598172539419, "y": 236.818110997055}, "selected": false}, {"data": {"id": "579333", "Enrichment": 2.559353993277849, "BestLogPInGroup": -11.981654863057864, "_PATTERN_": "M1", "_GeneInGOAndHitList": 21.0, "Hits": "RUNX2|COL1A1|DLX1|DLX2|HOXA3|HOXA5|HOXA6|HOXA7|HOXA9|HOXA11|HOXC4|HOXC6|HOXD10|SMAD3|MMP14|PBX1|SP3|PCGF2|MTHFD1L|SLC39A1|GRHL2", "Description": "embryonic skeletal system development", "_MEMBER_Input_ID": 1.0, "_GeneInHitList": 1562.0, "Category": "GO Biological Processes", "Log_q_value_": -2.3034096595169222, "Symbol": "GO:0048706", "LogP": -4.19767807447811, "_RANK_": 1.0, "FirstInGroupByEnrichment": 0.0, "_TotalGeneInLibrary": 24367.0, "_GeneInGO": 128.0, "shared_name": "136", "_InGO": 1.3444302176696543, "Gene": "GO:0048706", "_COUNT_001": 21.0, "SUID": 579333, "id_original": "136", "selected": false, "CategoryID": 19.0, "STDV_InGO": 0.29139991611511556, "EvidenceCutoff": 0.0, "PARENT_GO": "19_GO:0032502 developmental process", "FirstInGroupByLogP": 0.0, "GROUP_ID": 1.0, "name": "136", "BestEnrichmentInGroup": 2.5999786598378147, "CLUSTER_LABEL": " ", "GeneID": "860|1277|1745|1746|3200|3202|3203|3204|3205|3207|3221|3223|3236|4088|4323|5087|6670|7703|25902|27173|79977", "_LogP_Input_ID": -4.19767807447811, "Z_score": 4.629242681749554, "GiniIndex": 0.0, "_COUNT_001_PCT": 100.0}, "position": {"x": 1094.5582654350942, "y": 117.5448718613128}, "selected": false}, {"data": {"id": "579332", "Enrichment": 2.0347659076991595, "BestLogPInGroup": -11.458053982125241, "_PATTERN_": "M1", "_GeneInGOAndHitList": 90.0, "Hits": "ABL1|AKT1|ANPEP|ANXA2|KLK3|ZFP36L1|BTG1|RUNX1|COL4A1|COL4A2|TYMP|EPHA2|EPHB4|ERBB2|FOXF1|GATA2|B4GALT1|NR4A1|HOXA3|HOXA5|HOXA7|HSPB1|HSPG2|ID1|ISL1|JUNB|JUP|LRP1|SMAD7|MEIS1|MMP14|MYH9|NFE2L2|NOTCH1|NOTCH4|SERPINF1|PKM|SERPINF2|PTPRB|PTPRM|RRAS|CX3CL1|SHB|SHC1|SOX4|SP1|SPARC|SRF|TJP1|XBP1|YWHAZ|PIK3R3|HGS|QKI|RAMP1|HOXB13|GNA13|JMJD6|HEY2|TSPAN12|TIPARP|TNFRSF12A|HDAC7|RIN2|PARVA|ADAMTS9|SEMA6A|NIBAN2|MMRN2|ZC3H12A|COL18A1|APOLD1|ADTRP|CREB3L1|PLCD3|DAB2IP|MIA3|MIRLET7G|MIR106B|MIR125A|MIR125B1|MIR132|MIR21|MIR212|MIR27B|MIR29A|MIR29B1|MIR30B|MIR99B|MIR375", "Description": "blood vessel morphogenesis", "_MEMBER_Input_ID": 1.0, "_GeneInHitList": 1562.0, "Category": "GO Biological Processes", "Log_q_value_": -6.605623527795833, "Symbol": "GO:0048514", "LogP": -10.021639600113666, "_RANK_": 1.0, "FirstInGroupByEnrichment": 0.0, "_TotalGeneInLibrary": 24367.0, "_GeneInGO": 690.0, "shared_name": "135", "_InGO": 5.761843790012804, "Gene": "GO:0048514", "_COUNT_001": 90.0, "SUID": 579332, "id_original": "135", "selected": false, "CategoryID": 19.0, "STDV_InGO": 0.589594760428026, "EvidenceCutoff": 0.0, "PARENT_GO": "19_GO:0032502 developmental process", "FirstInGroupByLogP": 0.0, "GROUP_ID": 2.0, "name": "135", "BestEnrichmentInGroup": 2.262576849019167, "CLUSTER_LABEL": " ", "GeneID": "25|207|290|302|354|677|694|861|1282|1284|1890|1969|2050|2064|2294|2624|2683|3164|3200|3202|3204|3315|3339|3397|3670|3726|3728|4035|4092|4211|4323|4627|4780|4851|4855|5176|5315|5345|5787|5797|6237|6376|6461|6464|6659|6667|6678|6722|7082|7494|7534|8503|9146|9444|10267|10481|10672|23210|23493|23554|25976|51330|51564|54453|55742|56999|57556|64855|79812|80149|80781|81575|84830|90993|113026|153090|375056|406890|406900|406910|406911|406921|406991|406994|407019|407021|407024|407030|407056|494324", "_LogP_Input_ID": -10.021639600113666, "Z_score": 7.216405292446978, "GiniIndex": 0.0, "_COUNT_001_PCT": 100.0}, "position": {"x": 310.70873998343404, "y": 64.91212039646905}, "selected": false}, {"data": {"id": "579331", "Enrichment": 2.0484680350237325, "BestLogPInGroup": -11.458053982125241, "_PATTERN_": "M1", "_GeneInGOAndHitList": 78.0, "Hits": "ABL1|AKT1|ANPEP|ANXA2|KLK3|BTG1|RUNX1|COL4A1|COL4A2|TYMP|EPHA2|EPHB4|ERBB2|GATA2|B4GALT1|NR4A1|HOXA3|HOXA5|HOXA7|HSPB1|HSPG2|ID1|ISL1|JUP|MEIS1|MMP14|MYH9|NFE2L2|NOTCH1|NOTCH4|SERPINF1|PKM|PTPRB|PTPRM|RRAS|CX3CL1|SHB|SHC1|SP1|SPARC|SRF|TJP1|XBP1|YWHAZ|PIK3R3|HGS|RAMP1|HOXB13|GNA13|JMJD6|TSPAN12|TNFRSF12A|HDAC7|PARVA|ADAMTS9|SEMA6A|NIBAN2|MMRN2|ZC3H12A|COL18A1|APOLD1|ADTRP|CREB3L1|PLCD3|DAB2IP|MIA3|MIRLET7G|MIR106B|MIR125A|MIR125B1|MIR132|MIR21|MIR212|MIR27B|MIR29A|MIR30B|MIR99B|MIR375", "Description": "angiogenesis", "_MEMBER_Input_ID": 1.0, "_GeneInHitList": 1562.0, "Category": "GO Biological Processes", "Log_q_value_": -5.7810708527999815, "Symbol": "GO:0001525", "LogP": -8.896056929453833, "_RANK_": 1.0, "FirstInGroupByEnrichment": 0.0, "_TotalGeneInLibrary": 24367.0, "_GeneInGO": 594.0, "shared_name": "134", "_InGO": 4.99359795134443, "Gene": "GO:0001525", "_COUNT_001": 78.0, "SUID": 579331, "id_original": "134", "selected": false, "CategoryID": 19.0, "STDV_InGO": 0.5511156038954877, "EvidenceCutoff": 0.0, "PARENT_GO": "19_GO:0032502 developmental process", "FirstInGroupByLogP": 0.0, "GROUP_ID": 2.0, "name": "134", "BestEnrichmentInGroup": 2.262576849019167, "CLUSTER_LABEL": " ", "GeneID": "25|207|290|302|354|694|861|1282|1284|1890|1969|2050|2064|2624|2683|3164|3200|3202|3204|3315|3339|3397|3670|3728|4211|4323|4627|4780|4851|4855|5176|5315|5787|5797|6237|6376|6461|6464|6667|6678|6722|7082|7494|7534|8503|9146|10267|10481|10672|23210|23554|51330|51564|55742|56999|57556|64855|79812|80149|80781|81575|84830|90993|113026|153090|375056|406890|406900|406910|406911|406921|406991|406994|407019|407021|407030|407056|494324", "_LogP_Input_ID": -8.896056929453833, "Z_score": 6.770551069289629, "GiniIndex": 0.0, "_COUNT_001_PCT": 100.0}, "position": {"x": 295.9089047783559, "y": 131.70192294773858}, "selected": false}, {"data": {"id": "579330", "Enrichment": 1.9207829034409145, "BestLogPInGroup": -11.458053982125241, "_PATTERN_": "M1", "_GeneInGOAndHitList": 74.0, "Hits": "ABL1|ACTN4|AGER|AKT1|BCL6|CCR7|COL1A1|CLDN4|CLDN3|CLDN7|EGFR|FGR|FOXF1|MTOR|GATA2|HMGB1|HSPB1|IGF1R|IL1R1|ILK|ITGA3|LRP1|SMAD3|MMP14|NEDD9|NFE2L2|NOTCH1|ROR2|P2RY6|PDGFB|PPP3CA|LGMN|RRAS|RREB1|CX3CL1|SP1|SPARC|TJP1|TRIP6|XBP1|PTP4A1|PIK3R3|IRS2|CCN4|SPAG9|ARHGEF2|DOCK4|ELP5|FAM89B|FOXP1|HDAC7|SSH1|RIN2|TRPM4|SEMA4C|RTN4|S100A14|SEMA6A|GPSM3|VSIR|FBXO31|CFAP69|ZC3H12A|COL18A1|WNT5B|ARHGEF39|DAB2IP|CAVIN1|MIA3|MIR132|MIR21|MIR27B|MIR29A|MIR29B1", "Description": "positive regulation of cellular component movement", "_MEMBER_Input_ID": 1.0, "_GeneInHitList": 1562.0, "Category": "GO Biological Processes", "Log_q_value_": -4.56887222107189, "Symbol": "GO:0051272", "LogP": -7.285918289053704, "_RANK_": 1.0, "FirstInGroupByEnrichment": 0.0, "_TotalGeneInLibrary": 24367.0, "_GeneInGO": 601.0, "shared_name": "133", "_InGO": 4.7375160051216385, "Gene": "GO:0051272", "_COUNT_001": 74.0, "SUID": 579330, "id_original": "133", "selected": false, "CategoryID": 19.0, "STDV_InGO": 0.5375214271483969, "EvidenceCutoff": 0.0, "PARENT_GO": "19_GO:0048518 positive regulation of biological process", "FirstInGroupByLogP": 0.0, "GROUP_ID": 2.0, "name": "133", "BestEnrichmentInGroup": 2.262576849019167, "CLUSTER_LABEL": " ", "GeneID": "25|81|177|207|604|1236|1277|1364|1365|1366|1956|2268|2294|2475|2624|3146|3315|3480|3554|3611|3675|4035|4088|4323|4739|4780|4851|4920|5031|5155|5530|5641|6237|6239|6376|6667|6678|7082|7205|7494|7803|8503|8660|8840|9043|9181|9732|23587|23625|27086|51564|54434|54453|54795|54910|57142|57402|57556|63940|64115|79791|79846|80149|80781|81029|84904|153090|284119|375056|406921|406991|407019|407021|407024", "_LogP_Input_ID": -7.285918289053704, "Z_score": 5.981829538764541, "GiniIndex": 0.0, "_COUNT_001_PCT": 100.0}, "position": {"x": 110.68191503226217, "y": 100.90262180027764}, "selected": false}, {"data": {"id": "579329", "Enrichment": 1.9232718853594795, "BestLogPInGroup": -11.458053982125241, "_PATTERN_": "M1", "_GeneInGOAndHitList": 72.0, "Hits": "ABL1|ACTN4|AGER|AKT1|CCR7|COL1A1|CLDN4|CLDN3|CLDN7|EGFR|FGR|FOXF1|MTOR|GATA2|HMGB1|HSPB1|IGF1R|IL1R1|ILK|ITGA3|LRP1|SMAD3|MMP14|NEDD9|NFE2L2|NOTCH1|ROR2|P2RY6|PDGFB|PPP3CA|LGMN|RRAS|RREB1|CX3CL1|SP1|SPARC|TJP1|TRIP6|XBP1|PTP4A1|PIK3R3|IRS2|CCN4|SPAG9|ARHGEF2|DOCK4|ELP5|FAM89B|FOXP1|HDAC7|SSH1|RIN2|SEMA4C|RTN4|S100A14|SEMA6A|GPSM3|VSIR|FBXO31|CFAP69|ZC3H12A|COL18A1|WNT5B|ARHGEF39|DAB2IP|CAVIN1|MIA3|MIR132|MIR21|MIR27B|MIR29A|MIR29B1", "Description": "positive regulation of cell motility", "_MEMBER_Input_ID": 1.0, "_GeneInHitList": 1562.0, "Category": "GO Biological Processes", "Log_q_value_": -4.443958839207675, "Symbol": "GO:2000147", "LogP": -7.128280321523706, "_RANK_": 1.0, "FirstInGroupByEnrichment": 0.0, "_TotalGeneInLibrary": 24367.0, "_GeneInGO": 584.0, "shared_name": "132", "_InGO": 4.609475032010243, "Gene": "GO:2000147", "_COUNT_001": 72.0, "SUID": 579329, "id_original": "132", "selected": false, "CategoryID": 19.0, "STDV_InGO": 0.5305640725101949, "EvidenceCutoff": 0.0, "PARENT_GO": "19_GO:0040011 locomotion", "FirstInGroupByLogP": 0.0, "GROUP_ID": 2.0, "name": "132", "BestEnrichmentInGroup": 2.262576849019167, "CLUSTER_LABEL": " ", "GeneID": "25|81|177|207|1236|1277|1364|1365|1366|1956|2268|2294|2475|2624|3146|3315|3480|3554|3611|3675|4035|4088|4323|4739|4780|4851|4920|5031|5155|5530|5641|6237|6239|6376|6667|6678|7082|7205|7494|7803|8503|8660|8840|9043|9181|9732|23587|23625|27086|51564|54434|54453|54910|57142|57402|57556|63940|64115|79791|79846|80149|80781|81029|84904|153090|284119|375056|406921|406991|407019|407021|407024", "_LogP_Input_ID": -7.128280321523706, "Z_score": 5.910446806291715, "GiniIndex": 0.0, "_COUNT_001_PCT": 100.0}, "position": {"x": 119.86486791312154, "y": -19.838413661148138}, "selected": false}, {"data": {"id": "579328", "Enrichment": 1.9221270806658133, "BestLogPInGroup": -11.458053982125241, "_PATTERN_": "M1", "_GeneInGOAndHitList": 69.0, "Hits": "ABL1|ACTN4|AGER|AKT1|CCR7|COL1A1|CLDN4|CLDN3|EGFR|FGR|FOXF1|MTOR|GATA2|HMGB1|HSPB1|IGF1R|IL1R1|ILK|ITGA3|LRP1|SMAD3|MMP14|NEDD9|NFE2L2|NOTCH1|ROR2|P2RY6|PDGFB|PPP3CA|LGMN|RRAS|RREB1|CX3CL1|SP1|SPARC|TJP1|TRIP6|XBP1|PTP4A1|PIK3R3|IRS2|CCN4|SPAG9|ARHGEF2|DOCK4|ELP5|FAM89B|FOXP1|HDAC7|SSH1|RIN2|SEMA4C|RTN4|S100A14|SEMA6A|GPSM3|VSIR|FBXO31|ZC3H12A|COL18A1|WNT5B|ARHGEF39|DAB2IP|MIA3|MIR132|MIR21|MIR27B|MIR29A|MIR29B1", "Description": "positive regulation of cell migration", "_MEMBER_Input_ID": 1.0, "_GeneInHitList": 1562.0, "Category": "GO Biological Processes", "Log_q_value_": -4.236027627458415, "Symbol": "GO:0030335", "LogP": -6.847563510670255, "_RANK_": 1.0, "FirstInGroupByEnrichment": 0.0, "_TotalGeneInLibrary": 24367.0, "_GeneInGO": 560.0, "shared_name": "131", "_InGO": 4.41741357234315, "Gene": "GO:0030335", "_COUNT_001": 69.0, "SUID": 579328, "id_original": "131", "selected": false, "CategoryID": 19.0, "STDV_InGO": 0.5199156674482185, "EvidenceCutoff": 0.0, "PARENT_GO": "19_GO:0040011 locomotion", "FirstInGroupByLogP": 0.0, "GROUP_ID": 2.0, "name": "131", "BestEnrichmentInGroup": 2.262576849019167, "CLUSTER_LABEL": " ", "GeneID": "25|81|177|207|1236|1277|1364|1365|1956|2268|2294|2475|2624|3146|3315|3480|3554|3611|3675|4035|4088|4323|4739|4780|4851|4920|5031|5155|5530|5641|6237|6239|6376|6667|6678|7082|7205|7494|7803|8503|8660|8840|9043|9181|9732|23587|23625|27086|51564|54434|54453|54910|57142|57402|57556|63940|64115|79791|80149|80781|81029|84904|153090|375056|406921|406991|407019|407021|407024", "_LogP_Input_ID": -6.847563510670255, "Z_score": 5.777634313139373, "GiniIndex": 0.0, "_COUNT_001_PCT": 100.0}, "position": {"x": 162.2745511406606, "y": 25.623317295883112}, "selected": false}, {"data": {"id": "579327", "Enrichment": 2.161938776930357, "BestLogPInGroup": -11.458053982125241, "_PATTERN_": "M1", "_GeneInGOAndHitList": 51.0, "Hits": "ABL1|ACTA2|ACTG2|AKT1|DPP4|EPHA2|EPHB4|FOXF1|MTOR|GATA2|HMGB1|NR4A1|HSPB1|ID1|ITGA3|JUP|MYH9|NFE2L2|NOTCH1|PDGFB|SERPINF1|LGMN|PTPRM|PTPRR|RRAS|RREB1|S100A2|SP1|SPARC|SRF|PIK3R3|ZEB2|KANK1|MACF1|CORO1C|FOXP1|HDAC7|RIN2|ADAMTS9|RTN4|MMRN2|GRHL2|ZC3H12A|ADTRP|DAB2IP|MIA3|MIR132|MIR21|MIR212|MIR27B|MIR29A", "Description": "tissue migration", "_MEMBER_Input_ID": 1.0, "_GeneInHitList": 1562.0, "Category": "GO Biological Processes", "Log_q_value_": -4.184765854218302, "Symbol": "GO:0090130", "LogP": -6.763509224033834, "_RANK_": 1.0, "FirstInGroupByEnrichment": 0.0, "_TotalGeneInLibrary": 24367.0, "_GeneInGO": 368.0, "shared_name": "130", "_InGO": 3.265044814340589, "Gene": "GO:0090130", "_COUNT_001": 51.0, "SUID": 579327, "id_original": "130", "selected": false, "CategoryID": 19.0, "STDV_InGO": 0.4496719274960147, "EvidenceCutoff": 0.0, "PARENT_GO": "19_GO:0032501 multicellular organismal process", "FirstInGroupByLogP": 0.0, "GROUP_ID": 2.0, "name": "130", "BestEnrichmentInGroup": 2.262576849019167, "CLUSTER_LABEL": " ", "GeneID": "25|59|72|207|1803|1969|2050|2294|2475|2624|3146|3164|3315|3397|3675|3728|4627|4780|4851|5155|5176|5641|5797|5801|6237|6239|6273|6667|6678|6722|8503|9839|23189|23499|23603|27086|51564|54453|56999|57142|79812|79977|80149|84830|153090|375056|406921|406991|406994|407019|407021", "_LogP_Input_ID": -6.763509224033834, "Z_score": 5.877981006145582, "GiniIndex": 0.0, "_COUNT_001_PCT": 100.0}, "position": {"x": 220.49414037405904, "y": 3.6025500839690494}, "selected": false}, {"data": {"id": "579326", "Enrichment": 1.8352790540031634, "BestLogPInGroup": -11.458053982125241, "_PATTERN_": "M1", "_GeneInGOAndHitList": 72.0, "Hits": "ABL1|ACTN4|AGER|AKT1|CCR7|COL1A1|CLDN4|CLDN3|CLDN7|EGFR|FGR|FOXF1|MTOR|GATA2|HMGB1|HSPB1|IGF1R|IL1R1|ILK|ITGA3|LRP1|SMAD3|MMP14|NEDD9|NFE2L2|NOTCH1|ROR2|P2RY6|PDGFB|PPP3CA|LGMN|RRAS|RREB1|CX3CL1|SP1|SPARC|TJP1|TRIP6|XBP1|PTP4A1|PIK3R3|IRS2|CCN4|SPAG9|ARHGEF2|DOCK4|ELP5|FAM89B|FOXP1|HDAC7|SSH1|RIN2|SEMA4C|RTN4|S100A14|SEMA6A|GPSM3|VSIR|FBXO31|CFAP69|ZC3H12A|COL18A1|WNT5B|ARHGEF39|DAB2IP|CAVIN1|MIA3|MIR132|MIR21|MIR27B|MIR29A|MIR29B1", "Description": "positive regulation of locomotion", "_MEMBER_Input_ID": 1.0, "_GeneInHitList": 1562.0, "Category": "GO Biological Processes", "Log_q_value_": -3.842561890756183, "Symbol": "GO:0040017", "LogP": -6.335593147365133, "_RANK_": 1.0, "FirstInGroupByEnrichment": 0.0, "_TotalGeneInLibrary": 24367.0, "_GeneInGO": 612.0, "shared_name": "129", "_InGO": 4.609475032010243, "Gene": "GO:0040017", "_COUNT_001": 72.0, "SUID": 579326, "id_original": "129", "selected": false, "CategoryID": 19.0, "STDV_InGO": 0.5305640725101949, "EvidenceCutoff": 0.0, "PARENT_GO": "19_GO:0040011 locomotion", "FirstInGroupByLogP": 0.0, "GROUP_ID": 2.0, "name": "129", "BestEnrichmentInGroup": 2.262576849019167, "CLUSTER_LABEL": " ", "GeneID": "25|81|177|207|1236|1277|1364|1365|1366|1956|2268|2294|2475|2624|3146|3315|3480|3554|3611|3675|4035|4088|4323|4739|4780|4851|4920|5031|5155|5530|5641|6237|6239|6376|6667|6678|7082|7205|7494|7803|8503|8660|8840|9043|9181|9732|23587|23625|27086|51564|54434|54453|54910|57142|57402|57556|63940|64115|79791|79846|80149|80781|81029|84904|153090|284119|375056|406921|406991|407019|407021|407024", "_LogP_Input_ID": -6.335593147365133, "Z_score": 5.477058517548229, "GiniIndex": 0.0, "_COUNT_001_PCT": 100.0}, "position": {"x": 91.62100194632467, "y": 45.443217808578424}, "selected": false}, {"data": {"id": "579325", "Enrichment": 1.9499839948783613, "BestLogPInGroup": -11.458053982125241, "_PATTERN_": "M1", "_GeneInGOAndHitList": 59.0, "Hits": "ABL1|AKT1|CFL1|DPP4|EPHA2|EPHB4|MTOR|GATA2|HMGB1|NR4A1|HSPB1|ID1|ILK|ISL1|ITGA3|ITGB7|JUP|MYH9|NFE2L2|NOTCH1|PDGFB|SERPINF1|LGMN|PTPRM|PTPRR|RRAS|RREB1|S100A2|SDC4|SP1|SPARC|SRF|TNS1|ZFAND5|PIK3R3|ZEB2|KANK1|MACF1|CORO1C|FOXP1|HDAC7|RIN2|SEMA4C|APPL2|MESP1|ADAMTS9|RTN4|SEMA6A|MMRN2|ZC3H12A|ARID5B|ADTRP|DAB2IP|MIA3|MIR132|MIR21|MIR212|MIR27B|MIR29A", "Description": "ameboidal-type cell migration", "_MEMBER_Input_ID": 1.0, "_GeneInHitList": 1562.0, "Category": "GO Biological Processes", "Log_q_value_": -3.718509269595985, "Symbol": "GO:0001667", "LogP": -6.154431885609997, "_RANK_": 1.0, "FirstInGroupByEnrichment": 0.0, "_TotalGeneInLibrary": 24367.0, "_GeneInGO": 472.0, "shared_name": "128", "_InGO": 3.7772087067861717, "Gene": "GO:0001667", "_COUNT_001": 59.0, "SUID": 579325, "id_original": "128", "selected": false, "CategoryID": 19.0, "STDV_InGO": 0.48237406725279086, "EvidenceCutoff": 0.0, "PARENT_GO": "19_GO:0040011 locomotion", "FirstInGroupByLogP": 0.0, "GROUP_ID": 2.0, "name": "128", "BestEnrichmentInGroup": 2.262576849019167, "CLUSTER_LABEL": " ", "GeneID": "25|207|1072|1803|1969|2050|2475|2624|3146|3164|3315|3397|3611|3670|3675|3695|3728|4627|4780|4851|5155|5176|5641|5797|5801|6237|6239|6273|6385|6667|6678|6722|7145|7763|8503|9839|23189|23499|23603|27086|51564|54453|54910|55198|55897|56999|57142|57556|79812|80149|84159|84830|153090|375056|406921|406991|406994|407019|407021", "_LogP_Input_ID": -6.154431885609997, "Z_score": 5.454455045773322, "GiniIndex": 0.0, "_COUNT_001_PCT": 100.0}, "position": {"x": 221.48583959280904, "y": 65.25806766209405}, "selected": false}, {"data": {"id": "579324", "Enrichment": 2.0684913094842283, "BestLogPInGroup": -11.458053982125241, "_PATTERN_": "M1", "_GeneInGOAndHitList": 48.0, "Hits": "ABL1|AKT1|DPP4|EPHA2|EPHB4|MTOR|GATA2|HMGB1|NR4A1|HSPB1|ID1|ITGA3|JUP|MYH9|NFE2L2|NOTCH1|PDGFB|SERPINF1|LGMN|PTPRM|PTPRR|RRAS|RREB1|S100A2|SP1|SPARC|SRF|PIK3R3|ZEB2|KANK1|MACF1|CORO1C|FOXP1|HDAC7|RIN2|ADAMTS9|RTN4|MMRN2|GRHL2|ZC3H12A|ADTRP|DAB2IP|MIA3|MIR132|MIR21|MIR212|MIR27B|MIR29A", "Description": "epithelium migration", "_MEMBER_Input_ID": 1.0, "_GeneInHitList": 1562.0, "Category": "GO Biological Processes", "Log_q_value_": -3.4836403560467692, "Symbol": "GO:0090132", "LogP": -5.843705023035452, "_RANK_": 1.0, "FirstInGroupByEnrichment": 0.0, "_TotalGeneInLibrary": 24367.0, "_GeneInGO": 362.0, "shared_name": "127", "_InGO": 3.0729833546734957, "Gene": "GO:0090132", "_COUNT_001": 48.0, "SUID": 579324, "id_original": "127", "selected": false, "CategoryID": 19.0, "STDV_InGO": 0.43667870324195607, "EvidenceCutoff": 0.0, "PARENT_GO": "19_GO:0032501 multicellular organismal process", "FirstInGroupByLogP": 0.0, "GROUP_ID": 2.0, "name": "127", "BestEnrichmentInGroup": 2.262576849019167, "CLUSTER_LABEL": " ", "GeneID": "25|207|1803|1969|2050|2475|2624|3146|3164|3315|3397|3675|3728|4627|4780|4851|5155|5176|5641|5797|5801|6237|6239|6273|6667|6678|6722|8503|9839|23189|23499|23603|27086|51564|54453|56999|57142|79812|79977|80149|84830|153090|375056|406921|406991|406994|407019|407021", "_LogP_Input_ID": -5.843705023035452, "Z_score": 5.3603360921656265, "GiniIndex": 0.0, "_COUNT_001_PCT": 100.0}, "position": {"x": 199.6452786797231, "y": 112.40643649754327}, "selected": false}, {"data": {"id": "579323", "Enrichment": 2.693503072062197, "BestLogPInGroup": -10.799037371088092, "_PATTERN_": "M1", "_GeneInGOAndHitList": 24.0, "Hits": "CCND1|CEBPB|DBP|EGFR|HPN|SMAD3|NOTCH1|PKM|PTCH1|RARA|RARB|REG1A|SP3|TAF10|HNF1B|XBP1|CFLAR|RB1CC1|ZMPSTE24|CITED2|COBL|CCDC40|MESP1|ARID5B", "Description": "liver development", "_MEMBER_Input_ID": 1.0, "_GeneInHitList": 1562.0, "Category": "GO Biological Processes", "Log_q_value_": -2.9604580736572177, "Symbol": "GO:0001889", "LogP": -5.092043412130531, "_RANK_": 1.0, "FirstInGroupByEnrichment": 0.0, "_TotalGeneInLibrary": 24367.0, "_GeneInGO": 139.0, "shared_name": "126", "_InGO": 1.5364916773367479, "Gene": "GO:0001889", "_COUNT_001": 24.0, "SUID": 579323, "id_original": "126", "selected": false, "CategoryID": 19.0, "STDV_InGO": 0.3112162348789158, "EvidenceCutoff": 0.0, "PARENT_GO": "19_GO:0032502 developmental process", "FirstInGroupByLogP": 0.0, "GROUP_ID": 3.0, "name": "126", "BestEnrichmentInGroup": 2.693503072062197, "CLUSTER_LABEL": " ", "GeneID": "595|1051|1628|1956|3249|4088|4851|5315|5727|5914|5915|5967|6670|6881|6928|7494|8837|9821|10269|10370|23242|55036|55897|84159", "_LogP_Input_ID": -5.092043412130531, "Z_score": 5.240252367471818, "GiniIndex": 0.0, "_COUNT_001_PCT": 100.0}, "position": {"x": 920.0171582573598, "y": 288.2905536240081}, "selected": false}, {"data": {"id": "579322", "Enrichment": 2.6365980775820095, "BestLogPInGroup": -10.799037371088092, "_PATTERN_": "M1", "_GeneInGOAndHitList": 24.0, "Hits": "CCND1|CEBPB|DBP|EGFR|HPN|SMAD3|NOTCH1|PKM|PTCH1|RARA|RARB|REG1A|SP3|TAF10|HNF1B|XBP1|CFLAR|RB1CC1|ZMPSTE24|CITED2|COBL|CCDC40|MESP1|ARID5B", "Description": "hepaticobiliary system development", "_MEMBER_Input_ID": 1.0, "_GeneInHitList": 1562.0, "Category": "GO Biological Processes", "Log_q_value_": -2.853399113573478, "Symbol": "GO:0061008", "LogP": -4.92946712419696, "_RANK_": 1.0, "FirstInGroupByEnrichment": 0.0, "_TotalGeneInLibrary": 24367.0, "_GeneInGO": 142.0, "shared_name": "125", "_InGO": 1.5364916773367479, "Gene": "GO:0061008", "_COUNT_001": 24.0, "SUID": 579322, "id_original": "125", "selected": false, "CategoryID": 19.0, "STDV_InGO": 0.3112162348789158, "EvidenceCutoff": 0.0, "PARENT_GO": "19_GO:0032502 developmental process", "FirstInGroupByLogP": 0.0, "GROUP_ID": 3.0, "name": "125", "BestEnrichmentInGroup": 2.693503072062197, "CLUSTER_LABEL": " ", "GeneID": "595|1051|1628|1956|3249|4088|4851|5315|5727|5914|5915|5967|6670|6881|6928|7494|8837|9821|10269|10370|23242|55036|55897|84159", "_LogP_Input_ID": -4.92946712419696, "Z_score": 5.118844213278565, "GiniIndex": 0.0, "_COUNT_001_PCT": 100.0}, "position": {"x": 847.1867368096059, "y": 262.46868472752374}, "selected": false}, {"data": {"id": "579321", "Enrichment": 2.1272552671400304, "BestLogPInGroup": -10.799037371088092, "_PATTERN_": "M1", "_GeneInGOAndHitList": 18.0, "Hits": "EGFR|FOXF1|HOXA5|HOXD13|ID2|SMAD3|NOTCH1|RARB|RB1|REG1A|HNF1B|DCHS1|CBFA2T2|AGR2|COBL|SPDEF|CCDC40|MIR29B1", "Description": "digestive tract development", "_MEMBER_Input_ID": 1.0, "_GeneInHitList": 1562.0, "Category": "GO Biological Processes", "Log_q_value_": -1.2402490058758742, "Symbol": "GO:0048565", "LogP": -2.713017953055845, "_RANK_": 1.0, "FirstInGroupByEnrichment": 0.0, "_TotalGeneInLibrary": 24367.0, "_GeneInGO": 132.0, "shared_name": "124", "_InGO": 1.1523687580025608, "Gene": "GO:0048565", "_COUNT_001": 18.0, "SUID": 579321, "id_original": "124", "selected": false, "CategoryID": 19.0, "STDV_InGO": 0.2700463777281498, "EvidenceCutoff": 0.0, "PARENT_GO": "19_GO:0032502 developmental process", "FirstInGroupByLogP": 0.0, "GROUP_ID": 3.0, "name": "124", "BestEnrichmentInGroup": 2.693503072062197, "CLUSTER_LABEL": " ", "GeneID": "1956|2294|3202|3239|3398|4088|4851|5915|5925|5967|6928|8642|9139|10551|23242|25803|55036|407024", "_LogP_Input_ID": -2.713017953055845, "Z_score": 3.398641242210073, "GiniIndex": 0.0, "_COUNT_001_PCT": 100.0}, "position": {"x": 891.7064359062856, "y": 391.6527362411956}, "selected": false}, {"data": {"id": "579320", "Enrichment": 2.072710260290286, "BestLogPInGroup": -10.799037371088092, "_PATTERN_": "M1", "_GeneInGOAndHitList": 19.0, "Hits": "EGFR|FOXF1|HOXA5|HOXD13|ID2|SMAD3|NOTCH1|RARB|RB1|REG1A|HNF1B|XBP1|DCHS1|CBFA2T2|AGR2|COBL|SPDEF|CCDC40|MIR29B1", "Description": "digestive system development", "_MEMBER_Input_ID": 1.0, "_GeneInHitList": 1562.0, "Category": "GO Biological Processes", "Log_q_value_": -1.2279724245766641, "Symbol": "GO:0055123", "LogP": -2.6988893668950418, "_RANK_": 1.0, "FirstInGroupByEnrichment": 0.0, "_TotalGeneInLibrary": 24367.0, "_GeneInGO": 143.0, "shared_name": "123", "_InGO": 1.2163892445582587, "Gene": "GO:0055123", "_COUNT_001": 19.0, "SUID": 579320, "id_original": "123", "selected": false, "CategoryID": 19.0, "STDV_InGO": 0.2773564175744805, "EvidenceCutoff": 0.0, "PARENT_GO": "19_GO:0032502 developmental process", "FirstInGroupByLogP": 0.0, "GROUP_ID": 3.0, "name": "123", "BestEnrichmentInGroup": 2.693503072062197, "CLUSTER_LABEL": " ", "GeneID": "1956|2294|3202|3239|3398|4088|4851|5915|5925|5967|6928|7494|8642|9139|10551|23242|25803|55036|407024", "_LogP_Input_ID": -2.6988893668950418, "Z_score": 3.367015540130033, "GiniIndex": 0.0, "_COUNT_001_PCT": 100.0}, "position": {"x": 805.5274045342153, "y": 361.30022769627374}, "selected": false}, {"data": {"id": "579319", "Enrichment": 2.907248865091375, "BestLogPInGroup": -10.448191879197235, "_PATTERN_": "M1", "_GeneInGOAndHitList": 41.0, "Hits": "ACTA2|CLDN3|EPHA2|FASN|FAT1|FLNB|FOSL2|B4GALT1|SFN|RAPGEF1|FOXA1|HOXA5|ID1|NOTCH1|NOTCH4|PDGFB|RARA|RARB|RARG|SDC1|TJP1|TNFRSF1A|EZR|XBP1|FEM1B|HOXB13|TRIOBP|PALLD|SPDEF|ARHGEF26|ADAMTSL4|HYDIN|SHROOM3|PDZD7|GRHL2|COL18A1|WNT5B|TMEM79|TPRN|MIR29B1|CCDC88C", "Description": "epithelial cell development", "_MEMBER_Input_ID": 1.0, "_GeneInHitList": 1562.0, "Category": "GO Biological Processes", "Log_q_value_": -6.008590390060213, "Symbol": "GO:0002064", "LogP": -9.240482026055455, "_RANK_": 1.0, "FirstInGroupByEnrichment": 0.0, "_TotalGeneInLibrary": 24367.0, "_GeneInGO": 220.0, "shared_name": "122", "_InGO": 2.624839948783611, "Gene": "GO:0002064", "_COUNT_001": 41.0, "SUID": 579319, "id_original": "122", "selected": false, "CategoryID": 19.0, "STDV_InGO": 0.40451533583791444, "EvidenceCutoff": 0.0, "PARENT_GO": "19_GO:0032502 developmental process", "FirstInGroupByLogP": 0.0, "GROUP_ID": 4.0, "name": "122", "BestEnrichmentInGroup": 2.907248865091375, "CLUSTER_LABEL": " ", "GeneID": "59|1365|1969|2194|2195|2317|2355|2683|2810|2889|3169|3202|3397|4851|4855|5155|5914|5915|5916|6382|7082|7132|7430|7494|10116|10481|11078|23022|25803|26084|54507|54768|57619|79955|79977|80781|81029|84283|286262|407024|440193", "_LogP_Input_ID": -9.240482026055455, "Z_score": 7.4371186627175945, "GiniIndex": 0.0, "_COUNT_001_PCT": 100.0}, "position": {"x": 684.998046624059, "y": 257.4757342880706}, "selected": false}, {"data": {"id": "579318", "Enrichment": 2.004452821550941, "BestLogPInGroup": -10.38272655985909, "_PATTERN_": "M1", "_GeneInGOAndHitList": 92.0, "Hits": "ABL1|ACTN4|AGER|AKT1|ANK3|BCL6|RUNX1|CBFB|CD4|CD9|CEBPB|CCR7|COL1A1|CLDN7|DPP4|EPHA2|ERBB2|ERBB3|FBLN1|FOXF1|FLOT2|GSN|HLA-A|HMGB1|FOXA1|HOXA7|IL4R|ILK|IRF1|ITGA3|ITPKB|JUP|LAMA3|LIMS1|LRP1|SMAD3|SMAD7|MMP14|NEDD9|NINJ1|NOTCH1|P4HB|PIK3CB|SERPINF2|PPP1CB|PPP3CA|PRKAR1A|PTPN6|PTPRJ|PTPRR|RAC3|RARA|RRAS|RREB1|CX3CL1|SDC4|SPOCK1|SRF|TBCD|PRDX2|TJP1|UTRN|XBP1|ZBTB16|DLG5|PIEZO1|FLOT1|CITED2|AGR2|TRIOBP|ZBTB1|LIMCH1|KANK1|MACF1|CORO1C|ZBTB7B|RIN2|SEMA6A|CAMSAP3|PLEKHA2|VSIR|ZC3H12A|ADTRP|PLXNA4|CCDC88B|ACER2|MIA3|MIRLET7G|MIR141|MIR21|MIR27B|MIR30B", "Description": "regulation of cell adhesion", "_MEMBER_Input_ID": 1.0, "_GeneInHitList": 1562.0, "Category": "GO Biological Processes", "Log_q_value_": -6.517089133355425, "Symbol": "GO:0030155", "LogP": -9.881952683225876, "_RANK_": 1.0, "FirstInGroupByEnrichment": 0.0, "_TotalGeneInLibrary": 24367.0, "_GeneInGO": 716.0, "shared_name": "121", "_InGO": 5.8898847631242, "Gene": "GO:0030155", "_COUNT_001": 92.0, "SUID": 579318, "id_original": "121", "selected": false, "CategoryID": 19.0, "STDV_InGO": 0.5957047138803491, "EvidenceCutoff": 0.0, "PARENT_GO": "19_GO:0022610 biological adhesion", "FirstInGroupByLogP": 0.0, "GROUP_ID": 5.0, "name": "121", "BestEnrichmentInGroup": 6.239948783610755, "CLUSTER_LABEL": " ", "GeneID": "25|81|177|207|288|604|861|865|920|928|1051|1236|1277|1366|1803|1969|2064|2065|2192|2294|2319|2934|3105|3146|3169|3204|3566|3611|3659|3675|3707|3728|3909|3987|4035|4088|4092|4323|4739|4814|4851|5034|5291|5345|5500|5530|5573|5777|5795|5801|5881|5914|6237|6239|6376|6385|6695|6722|6904|7001|7082|7402|7494|7704|9231|9780|10211|10370|10551|11078|22890|22998|23189|23499|23603|51043|54453|57556|57662|59339|64115|80149|84830|91584|283234|340485|375056|406890|406933|406991|407019|407030", "_LogP_Input_ID": -9.881952683225876, "Z_score": 7.139682456443422, "GiniIndex": 0.0, "_COUNT_001_PCT": 100.0}, "position": {"x": -26.809149039393105, "y": 8.704341465804987}, "selected": false}, {"data": {"id": "579317", "Enrichment": 2.082118433956361, "BestLogPInGroup": -10.38272655985909, "_PATTERN_": "M1", "_GeneInGOAndHitList": 65.0, "Hits": "ABL1|AGER|B2M|BCL6|ZFP36L1|ZFP36L2|RUNX1|CBFB|CD4|CEBPB|ERBB2|FOXO3|FOS|MTOR|GATA2|H3-3B|HMGB1|HOXA5|HOXA7|HOXA9|HSPA1A|HSPA1B|ID2|IL4R|IRF1|ITPKB|SMAD7|MEIS1|MEIS2|MMP14|NFE2L2|NOTCH1|ROR2|PSMB3|PSMB8|PSMB9|PSMD1|PSMD2|PTPN6|RARA|RARG|RB1|PRDX2|KLF10|XBP1|ZFP36|ZBTB16|H3C8|H3C7|RNF41|TRIB1|N4BP2L2|ZBTB1|FOXP1|PRELID1|ZBTB7B|KLF13|PAF1|KMT2E|SH3RF1|VSIR|ZC3H12A|MIR125B1|MIR21|MIR30B", "Description": "regulation of hemopoiesis", "_MEMBER_Input_ID": 1.0, "_GeneInHitList": 1562.0, "Category": "GO Biological Processes", "Log_q_value_": -4.963555118146418, "Symbol": "GO:1903706", "LogP": -7.791299483621922, "_RANK_": 1.0, "FirstInGroupByEnrichment": 0.0, "_TotalGeneInLibrary": 24367.0, "_GeneInGO": 487.0, "shared_name": "120", "_InGO": 4.161331626120359, "Gene": "GO:1903706", "_COUNT_001": 65.0, "SUID": 579317, "id_original": "120", "selected": false, "CategoryID": 19.0, "STDV_InGO": 0.5052962022959689, "EvidenceCutoff": 0.0, "PARENT_GO": "19_GO:0002376 immune system process", "FirstInGroupByLogP": 0.0, "GROUP_ID": 5.0, "name": "120", "BestEnrichmentInGroup": 6.239948783610755, "CLUSTER_LABEL": " ", "GeneID": "25|177|567|604|677|678|861|865|920|1051|2064|2309|2353|2475|2624|3021|3146|3202|3204|3205|3303|3304|3398|3566|3659|3707|4092|4211|4212|4323|4780|4851|4920|5691|5696|5698|5707|5708|5777|5914|5916|5925|7001|7071|7494|7538|7704|8355|8968|10193|10221|10443|22890|27086|27166|51043|51621|54623|55904|57630|64115|80149|406911|406991|407030", "_LogP_Input_ID": -7.791299483621922, "Z_score": 6.313057008483269, "GiniIndex": 0.0, "_COUNT_001_PCT": 100.0}, "position": {"x": -212.10916162789408, "y": 146.49200778660577}, "selected": false}, {"data": {"id": "579316", "Enrichment": 2.5678801578645083, "BestLogPInGroup": -10.38272655985909, "_PATTERN_": "M1", "_GeneInGOAndHitList": 40.0, "Hits": "ABL1|B2M|BCL6|ZFP36L1|ZFP36L2|RUNX2|RUNX1|CBFB|CD4|CCR7|ERBB2|MTOR|HMGB1|IL4R|IRF1|ITPKB|LFNG|SMAD7|RARA|RORC|SOX4|SP3|SRF|STAT6|STK11|PRDX2|XBP1|ZBTB16|ZBTB1|JMJD6|FOXP1|PRELID1|NCAPH2|ZBTB7B|SH3RF1|VSIR|NHEJ1|ZC3H12A|MIR21|MIR30B", "Description": "T cell differentiation", "_MEMBER_Input_ID": 1.0, "_GeneInHitList": 1562.0, "Category": "GO Biological Processes", "Log_q_value_": -4.689810061262917, "Symbol": "GO:0030217", "LogP": -7.440714396505698, "_RANK_": 1.0, "FirstInGroupByEnrichment": 0.0, "_TotalGeneInLibrary": 24367.0, "_GeneInGO": 243.0, "shared_name": "119", "_InGO": 2.560819462227913, "Gene": "GO:0030217", "_COUNT_001": 40.0, "SUID": 579316, "id_original": "119", "selected": false, "CategoryID": 19.0, "STDV_InGO": 0.3996830930161245, "EvidenceCutoff": 0.0, "PARENT_GO": "19_GO:0002376 immune system process", "FirstInGroupByLogP": 0.0, "GROUP_ID": 5.0, "name": "119", "BestEnrichmentInGroup": 6.239948783610755, "CLUSTER_LABEL": " ", "GeneID": "25|567|604|677|678|860|861|865|920|1236|2064|2475|3146|3566|3659|3707|3955|4092|5914|6097|6659|6670|6722|6778|6794|7001|7494|7704|22890|23210|27086|27166|29781|51043|57630|64115|79840|80149|406991|407030", "_LogP_Input_ID": -7.440714396505698, "Z_score": 6.428489476558687, "GiniIndex": 0.0, "_COUNT_001_PCT": 100.0}, "position": {"x": -64.32071329415385, "y": 115.94578128514092}, "selected": false}, {"data": {"id": "579315", "Enrichment": 2.241108365944708, "BestLogPInGroup": -10.38272655985909, "_PATTERN_": "M1", "_GeneInGOAndHitList": 51.0, "Hits": "ABL1|B2M|BCL6|ZFP36L1|ZFP36L2|RUNX2|RUNX1|CBFB|CD4|CCR7|KLF6|ERBB2|MTOR|HMGB1|ID2|IL4R|IRF1|ITPKB|LFNG|SMAD7|MMP14|POU2F2|PTPN6|PTPRJ|RARA|RORC|SOX4|SP3|SRF|STAT6|STK11|PRDX2|TPD52|XBP1|ZBTB16|RNF41|ZBTB1|JMJD6|FOXP1|PRELID1|NCAPH2|ZBTB7B|ZBTB7A|LRRC8A|SH3RF1|VSIR|NHEJ1|ZC3H12A|IRF2BP2|MIR21|MIR30B", "Description": "lymphocyte differentiation", "_MEMBER_Input_ID": 1.0, "_GeneInHitList": 1562.0, "Category": "GO Biological Processes", "Log_q_value_": -4.55939158243124, "Symbol": "GO:0030098", "LogP": -7.265713785021281, "_RANK_": 1.0, "FirstInGroupByEnrichment": 0.0, "_TotalGeneInLibrary": 24367.0, "_GeneInGO": 355.0, "shared_name": "118", "_InGO": 3.265044814340589, "Gene": "GO:0030098", "_COUNT_001": 51.0, "SUID": 579315, "id_original": "118", "selected": false, "CategoryID": 19.0, "STDV_InGO": 0.4496719274960147, "EvidenceCutoff": 0.0, "PARENT_GO": "19_GO:0002376 immune system process", "FirstInGroupByLogP": 0.0, "GROUP_ID": 5.0, "name": "118", "BestEnrichmentInGroup": 6.239948783610755, "CLUSTER_LABEL": " ", "GeneID": "25|567|604|677|678|860|861|865|920|1236|1316|2064|2475|3146|3398|3566|3659|3707|3955|4092|4323|5452|5777|5795|5914|6097|6659|6670|6722|6778|6794|7001|7163|7494|7704|10193|22890|23210|27086|27166|29781|51043|51341|56262|57630|64115|79840|80149|359948|406991|407030", "_LogP_Input_ID": -7.265713785021281, "Z_score": 6.164918060913958, "GiniIndex": 0.0, "_COUNT_001_PCT": 100.0}, "position": {"x": -112.06080271065775, "y": 103.97247653660577}, "selected": false}, {"data": {"id": "579314", "Enrichment": 2.062483071505959, "BestLogPInGroup": -10.38272655985909, "_PATTERN_": "M1", "_GeneInGOAndHitList": 55.0, "Hits": "ABL1|AGER|AKT1|ANK3|BCL6|RUNX1|CBFB|CD4|CCR7|DPP4|ERBB2|FOXF1|FLOT2|HLA-A|HMGB1|FOXA1|IL4R|ILK|ITGA3|ITPKB|JUP|LIMS1|SMAD3|SMAD7|NEDD9|NINJ1|P4HB|SERPINF2|PPP3CA|PTPN6|PTPRJ|RAC3|RARA|RRAS|RREB1|CX3CL1|SDC4|TJP1|UTRN|XBP1|ZBTB16|PIEZO1|FLOT1|CITED2|AGR2|TRIOBP|ZBTB1|ZBTB7B|RIN2|PLEKHA2|VSIR|CCDC88B|MIR21|MIR27B|MIR30B", "Description": "positive regulation of cell adhesion", "_MEMBER_Input_ID": 1.0, "_GeneInHitList": 1562.0, "Category": "GO Biological Processes", "Log_q_value_": -4.024058214047317, "Symbol": "GO:0045785", "LogP": -6.557834438346326, "_RANK_": 1.0, "FirstInGroupByEnrichment": 0.0, "_TotalGeneInLibrary": 24367.0, "_GeneInGO": 416.0, "shared_name": "117", "_InGO": 3.5211267605633805, "Gene": "GO:0045785", "_COUNT_001": 55.0, "SUID": 579314, "id_original": "117", "selected": false, "CategoryID": 19.0, "STDV_InGO": 0.46635477364244116, "EvidenceCutoff": 0.0, "PARENT_GO": "19_GO:0022610 biological adhesion", "FirstInGroupByLogP": 0.0, "GROUP_ID": 5.0, "name": "117", "BestEnrichmentInGroup": 6.239948783610755, "CLUSTER_LABEL": " ", "GeneID": "25|177|207|288|604|861|865|920|1236|1803|2064|2294|2319|3105|3146|3169|3566|3611|3675|3707|3728|3987|4088|4092|4739|4814|5034|5345|5530|5777|5795|5881|5914|6237|6239|6376|6385|7082|7402|7494|7704|9780|10211|10370|10551|11078|22890|51043|54453|59339|64115|283234|406991|407019|407030", "_LogP_Input_ID": -6.557834438346326, "Z_score": 5.720374414863983, "GiniIndex": 0.0, "_COUNT_001_PCT": 100.0}, "position": {"x": -118.10121179879252, "y": -12.623630946304388}, "selected": false}, {"data": {"id": "579313", "Enrichment": 2.281613925980123, "BestLogPInGroup": -10.38272655985909, "_PATTERN_": "M1", "_GeneInGOAndHitList": 43.0, "Hits": "ABL1|ACTN4|AKT1|BCL6|RUNX1|CBFB|CD9|CEBPB|COL1A1|CLDN7|ERBB2|ERBB3|FBLN1|HMGB1|HOXA7|IL4R|IRF1|LRP1|SMAD7|MMP14|NOTCH1|PRKAR1A|PTPN6|CX3CL1|SDC4|SPOCK1|TBCD|PRDX2|DLG5|KANK1|CORO1C|ZBTB7B|SEMA6A|VSIR|ZC3H12A|ADTRP|PLXNA4|ACER2|MIA3|MIRLET7G|MIR141|MIR21|MIR30B", "Description": "negative regulation of cell adhesion", "_MEMBER_Input_ID": 1.0, "_GeneInHitList": 1562.0, "Category": "GO Biological Processes", "Log_q_value_": -3.941225542013802, "Symbol": "GO:0007162", "LogP": -6.447418244680723, "_RANK_": 1.0, "FirstInGroupByEnrichment": 0.0, "_TotalGeneInLibrary": 24367.0, "_GeneInGO": 294.0, "shared_name": "116", "_InGO": 2.7528809218950063, "Gene": "GO:0007162", "_COUNT_001": 43.0, "SUID": 579313, "id_original": "116", "selected": false, "CategoryID": 19.0, "STDV_InGO": 0.41399163922614024, "EvidenceCutoff": 0.0, "PARENT_GO": "19_GO:0022610 biological adhesion", "FirstInGroupByLogP": 0.0, "GROUP_ID": 5.0, "name": "116", "BestEnrichmentInGroup": 6.239948783610755, "CLUSTER_LABEL": " ", "GeneID": "25|81|207|604|861|865|928|1051|1277|1366|2064|2065|2192|3146|3204|3566|3659|4035|4092|4323|4851|5573|5777|6376|6385|6695|6904|7001|9231|23189|23603|51043|57556|64115|80149|84830|91584|340485|375056|406890|406933|406991|407030", "_LogP_Input_ID": -6.447418244680723, "Z_score": 5.7860658440014845, "GiniIndex": 0.0, "_COUNT_001_PCT": 100.0}, "position": {"x": -94.32050348580424, "y": 46.396586949203424}, "selected": false}, {"data": {"id": "579312", "Enrichment": 2.276137901495027, "BestLogPInGroup": -10.38272655985909, "_PATTERN_": "M1", "_GeneInGOAndHitList": 41.0, "Hits": "ABL1|AGER|BCL6|ZFP36L1|ZFP36L2|RUNX1|CBFB|CD4|CEBPB|ERBB2|FOS|MTOR|GATA2|HMGB1|HOXA7|ID2|IL4R|IRF1|ITPKB|SMAD7|MMP14|ROR2|PTPN6|RARA|RB1|PRDX2|KLF10|XBP1|ZBTB16|RNF41|TRIB1|ZBTB1|FOXP1|PRELID1|ZBTB7B|SH3RF1|VSIR|ZC3H12A|MIR125B1|MIR21|MIR30B", "Description": "regulation of leukocyte differentiation", "_MEMBER_Input_ID": 1.0, "_GeneInHitList": 1562.0, "Category": "GO Biological Processes", "Log_q_value_": -3.718509269595985, "Symbol": "GO:1902105", "LogP": -6.15112460373328, "_RANK_": 1.0, "FirstInGroupByEnrichment": 0.0, "_TotalGeneInLibrary": 24367.0, "_GeneInGO": 281.0, "shared_name": "115", "_InGO": 2.624839948783611, "Gene": "GO:1902105", "_COUNT_001": 41.0, "SUID": 579312, "id_original": "115", "selected": false, "CategoryID": 19.0, "STDV_InGO": 0.40451533583791444, "EvidenceCutoff": 0.0, "PARENT_GO": "19_GO:0002376 immune system process", "FirstInGroupByLogP": 0.0, "GROUP_ID": 5.0, "name": "115", "BestEnrichmentInGroup": 6.239948783610755, "CLUSTER_LABEL": " ", "GeneID": "25|177|604|677|678|861|865|920|1051|2064|2353|2475|2624|3146|3204|3398|3566|3659|3707|4092|4323|4920|5777|5914|5925|7001|7071|7494|7704|10193|10221|22890|27086|27166|51043|57630|64115|80149|406911|406991|407030", "_LogP_Input_ID": -6.15112460373328, "Z_score": 5.631006424542979, "GiniIndex": 0.0, "_COUNT_001_PCT": 100.0}, "position": {"x": -204.04350305855814, "y": 99.50052219090264}, "selected": false}, {"data": {"id": "579311", "Enrichment": 2.8579918092873693, "BestLogPInGroup": -10.38272655985909, "_PATTERN_": "M1", "_GeneInGOAndHitList": 24.0, "Hits": "AKT1|BCL6|RUNX1|CBFB|CEBPB|ERBB2|HMGB1|IL4R|IRF1|SMAD7|PRKAR1A|PTPN6|SDC4|PRDX2|DLG5|ZBTB7B|VSIR|ZC3H12A|ADTRP|MIA3|MIRLET7G|MIR141|MIR21|MIR30B", "Description": "negative regulation of leukocyte cell-cell adhesion", "_MEMBER_Input_ID": 1.0, "_GeneInHitList": 1562.0, "Category": "GO Biological Processes", "Log_q_value_": -3.3000110727214644, "Symbol": "GO:1903038", "LogP": -5.554659142804322, "_RANK_": 1.0, "FirstInGroupByEnrichment": 0.0, "_TotalGeneInLibrary": 24367.0, "_GeneInGO": 131.0, "shared_name": "114", "_InGO": 1.5364916773367479, "Gene": "GO:1903038", "_COUNT_001": 24.0, "SUID": 579311, "id_original": "114", "selected": false, "CategoryID": 19.0, "STDV_InGO": 0.3112162348789158, "EvidenceCutoff": 0.0, "PARENT_GO": "19_GO:0022610 biological adhesion", "FirstInGroupByLogP": 0.0, "GROUP_ID": 5.0, "name": "114", "BestEnrichmentInGroup": 6.239948783610755, "CLUSTER_LABEL": " ", "GeneID": "207|604|861|865|1051|2064|3146|3566|3659|4092|5573|5777|6385|7001|9231|51043|64115|80149|84830|375056|406890|406933|406991|407030", "_LogP_Input_ID": -5.554659142804322, "Z_score": 5.58041590010704, "GiniIndex": 0.0, "_COUNT_001_PCT": 100.0}, "position": {"x": -145.69694544137064, "y": 59.62374454197686}, "selected": false}, {"data": {"id": "579310", "Enrichment": 4.506629677052212, "BestLogPInGroup": -10.38272655985909, "_PATTERN_": "M1", "_GeneInGOAndHitList": 13.0, "Hits": "BCL6|RUNX1|CBFB|ERBB2|HMGB1|IL4R|IRF1|SMAD7|PRDX2|ZBTB7B|ZC3H12A|MIR21|MIR30B", "Description": "negative regulation of T cell differentiation", "_MEMBER_Input_ID": 1.0, "_GeneInHitList": 1562.0, "Category": "GO Biological Processes", "Log_q_value_": -3.265852058684192, "Symbol": "GO:0045581", "LogP": -5.509411156601438, "_RANK_": 1.0, "FirstInGroupByEnrichment": 0.0, "_TotalGeneInLibrary": 24367.0, "_GeneInGO": 45.0, "shared_name": "113", "_InGO": 0.8322663252240717, "Gene": "GO:0045581", "_COUNT_001": 13.0, "SUID": 579310, "id_original": "113", "selected": false, "CategoryID": 19.0, "STDV_InGO": 0.22986658336957838, "EvidenceCutoff": 0.0, "PARENT_GO": "19_GO:0022610 biological adhesion", "FirstInGroupByLogP": 0.0, "GROUP_ID": 5.0, "name": "113", "BestEnrichmentInGroup": 6.239948783610755, "CLUSTER_LABEL": " ", "GeneID": "604|861|865|2064|3146|3566|3659|4092|7001|51043|80149|406991|407030", "_LogP_Input_ID": -5.509411156601438, "Z_score": 6.161892469361177, "GiniIndex": 0.0, "_COUNT_001_PCT": 100.0}, "position": {"x": -91.57901407540385, "y": 189.42228656468194}, "selected": false}, {"data": {"id": "579309", "Enrichment": 3.278233962404201, "BestLogPInGroup": -10.25219876997185, "_PATTERN_": "M1", "_GeneInGOAndHitList": 29.0, "Hits": "AKT1|ZFP36L1|ZFP36L2|CEBPB|MTOR|GATA2|GNB3|HNRNPU|ID2|SMAD3|RORC|RREB1|TRIO|XBP1|ZFP36|ZBTB16|CCN4|NR1D1|ADIRF|CCDC85B|ZBTB7B|TRPM4|ATAT1|ZC3H12A|WNT5B|SH3PXD2B|MIR21|MIR27B|MIR29B1", "Description": "regulation of fat cell differentiation", "_MEMBER_Input_ID": 1.0, "_GeneInHitList": 1562.0, "Category": "GO Biological Processes", "Log_q_value_": -5.082626852088901, "Symbol": "GO:0045598", "LogP": -7.939334913499722, "_RANK_": 1.0, "FirstInGroupByEnrichment": 0.0, "_TotalGeneInLibrary": 24367.0, "_GeneInGO": 138.0, "shared_name": "112", "_InGO": 1.8565941101152368, "Gene": "GO:0045598", "_COUNT_001": 29.0, "SUID": 579309, "id_original": "112", "selected": false, "CategoryID": 19.0, "STDV_InGO": 0.341545471972582, "EvidenceCutoff": 0.0, "PARENT_GO": "19_GO:0032502 developmental process", "FirstInGroupByLogP": 0.0, "GROUP_ID": 6.0, "name": "112", "BestEnrichmentInGroup": 3.843446714542856, "CLUSTER_LABEL": " ", "GeneID": "207|677|678|1051|2475|2624|2784|3192|3398|4088|6097|6239|7204|7494|7538|7704|8840|9572|10974|11007|51043|54795|79969|80149|81029|285590|406991|407019|407024", "_LogP_Input_ID": -7.939334913499722, "Z_score": 7.024051973291059, "GiniIndex": 0.0, "_COUNT_001_PCT": 100.0}, "position": {"x": 649.7948305481618, "y": 716.9538837021331}, "selected": false}, {"data": {"id": "579308", "Enrichment": 3.843446714542856, "BestLogPInGroup": -10.25219876997185, "_PATTERN_": "M1", "_GeneInGOAndHitList": 17.0, "Hits": "AKT1|ZFP36L1|CEBPB|HNRNPU|ID2|RREB1|XBP1|ZFP36|ZBTB16|ADIRF|ZBTB7B|TRPM4|ZC3H12A|WNT5B|SH3PXD2B|MIR21|MIR29B1", "Description": "positive regulation of fat cell differentiation", "_MEMBER_Input_ID": 1.0, "_GeneInHitList": 1562.0, "Category": "GO Biological Processes", "Log_q_value_": -3.5480980256750203, "Symbol": "GO:0045600", "LogP": -5.9318844396665895, "_RANK_": 1.0, "FirstInGroupByEnrichment": 0.0, "_TotalGeneInLibrary": 24367.0, "_GeneInGO": 69.0, "shared_name": "111", "_InGO": 1.088348271446863, "Gene": "GO:0045600", "_COUNT_001": 17.0, "SUID": 579308, "id_original": "111", "selected": false, "CategoryID": 19.0, "STDV_InGO": 0.2625228789306787, "EvidenceCutoff": 0.0, "PARENT_GO": "19_GO:0048518 positive regulation of biological process", "FirstInGroupByLogP": 0.0, "GROUP_ID": 6.0, "name": "111", "BestEnrichmentInGroup": 3.843446714542856, "CLUSTER_LABEL": " ", "GeneID": "207|677|1051|3192|3398|6239|7494|7538|7704|10974|51043|54795|80149|81029|285590|406991|407024", "_LogP_Input_ID": -5.9318844396665895, "Z_score": 6.190161388882072, "GiniIndex": 0.0, "_COUNT_001_PCT": 100.0}, "position": {"x": 756.555151211614, "y": 731.7010150497894}, "selected": false}, {"data": {"id": "579307", "Enrichment": 2.0312333279982924, "BestLogPInGroup": -9.631064746783506, "_PATTERN_": "M1", "_GeneInGOAndHitList": 75.0, "Hits": "ACTB|ACTN1|AGER|DST|CAPZB|CD9|CDKN1B|COL1A1|CLDN4|CLDN3|CSRP1|DGKA|DGKQ|EGFR|ERBB2|ERBB3|FBLN1|MTOR|GATA2|B4GALT1|GSN|H3-3B|HSPB1|ILK|IRF1|ITPK1|ITPR1|SMAD3|MYH9|NFE2L2|NINJ1|NOTCH1|NOTCH4|PDGFB|PIK3CB|PKM|PLEC|SERPINF2|PPP3CA|PRKAR1A|PRKCH|PTK7|PTPN6|RAD51B|REG1A|RREB1|CX3CL1|SDC1|SDC4|SPARC|SRF|TARBP2|PRDX2|TLN1|XBP1|YWHAZ|H3C8|CFLAR|CCN4|H3C7|SYT7|FIBP|GNA13|KANK1|MACF1|MYOF|EHD2|TNFRSF12A|WFDC1|WNT5B|ADTRP|CADM4|JMJD1C|MIA3|MIR29B1", "Description": "wound healing", "_MEMBER_Input_ID": 1.0, "_GeneInHitList": 1562.0, "Category": "GO Biological Processes", "Log_q_value_": -5.438437835501816, "Symbol": "GO:0042060", "LogP": -8.415121213989385, "_RANK_": 1.0, "FirstInGroupByEnrichment": 0.0, "_TotalGeneInLibrary": 24367.0, "_GeneInGO": 576.0, "shared_name": "110", "_InGO": 4.801536491677337, "Gene": "GO:0042060", "_COUNT_001": 75.0, "SUID": 579307, "id_original": "110", "selected": false, "CategoryID": 19.0, "STDV_InGO": 0.5409592753355752, "EvidenceCutoff": 0.0, "PARENT_GO": "19_GO:0050896 response to stimulus", "FirstInGroupByLogP": 0.0, "GROUP_ID": 7.0, "name": "110", "BestEnrichmentInGroup": 2.8599765258215957, "CLUSTER_LABEL": " ", "GeneID": "60|87|177|667|832|928|1027|1277|1364|1365|1465|1606|1609|1956|2064|2065|2192|2475|2624|2683|2934|3021|3315|3611|3659|3705|3708|4088|4627|4780|4814|4851|4855|5155|5291|5315|5339|5345|5530|5573|5583|5754|5777|5890|5967|6239|6376|6382|6385|6678|6722|6895|7001|7094|7494|7534|8355|8837|8840|8968|9066|9158|10672|23189|23499|26509|30846|51330|58189|81029|84830|199731|221037|375056|407024", "_LogP_Input_ID": -8.415121213989385, "Z_score": 6.555101639484613, "GiniIndex": 0.0, "_COUNT_001_PCT": 100.0}, "position": {"x": -426.78987146676127, "y": 876.0394702499847}, "selected": false}, {"data": {"id": "579306", "Enrichment": 1.630755845815434, "BestLogPInGroup": -9.631064746783506, "_PATTERN_": "M1", "_GeneInGOAndHitList": 53.0, "Hits": "ACTB|ACTN1|ANXA2|CCND1|CAPZB|CD9|CHRM1|COL1A1|CLDN4|CSRP1|DGKA|DGKQ|SLC29A1|FBLN1|GATA2|SFN|H3-3B|HSPB1|ILK|IRF1|ITPK1|ITPR1|MYH9|NFE2L2|NPR3|PDGFB|PIK3CB|SERPINF2|PRKAR1A|PRKCH|PTPN6|RAD51B|SCNN1A|SRF|VAMP2|PRDX2|TLN1|UCN|XBP1|YWHAZ|H3C8|SOCS2|H3C7|FIBP|NCOR2|AGR2|GNA13|EHD2|ZBTB7B|PDSS2|TMEM79|ADTRP|JMJD1C", "Description": "regulation of body fluid levels", "_MEMBER_Input_ID": 1.0, "_GeneInHitList": 1562.0, "Category": "GO Biological Processes", "Log_q_value_": -1.7969940818144057, "Symbol": "GO:0050878", "LogP": -3.4928508507262817, "_RANK_": 1.0, "FirstInGroupByEnrichment": 0.0, "_TotalGeneInLibrary": 24367.0, "_GeneInGO": 507.0, "shared_name": "109", "_InGO": 3.3930857874519846, "Gene": "GO:0050878", "_COUNT_001": 53.0, "SUID": 579306, "id_original": "109", "selected": false, "CategoryID": 19.0, "STDV_InGO": 0.4581007579903272, "EvidenceCutoff": 0.0, "PARENT_GO": "19_GO:0065007 biological regulation", "FirstInGroupByLogP": 0.0, "GROUP_ID": 7.0, "name": "109", "BestEnrichmentInGroup": 2.8599765258215957, "CLUSTER_LABEL": " ", "GeneID": "60|87|302|595|832|928|1128|1277|1364|1465|1606|1609|2030|2192|2624|2810|3021|3315|3611|3659|3705|3708|4627|4780|4883|5155|5291|5345|5573|5583|5777|5890|6337|6722|6844|7001|7094|7349|7494|7534|8355|8835|8968|9158|9612|10551|10672|30846|51043|57107|84283|84830|221037", "_LogP_Input_ID": -3.4928508507262817, "Z_score": 3.7561907654832365, "GiniIndex": 0.0, "_COUNT_001_PCT": 100.0}, "position": {"x": -403.33332086861674, "y": 791.087153965805}, "selected": false}, {"data": {"id": "579305", "Enrichment": 2.8599765258215957, "BestLogPInGroup": -9.631064746783506, "_PATTERN_": "M1", "_GeneInGOAndHitList": 11.0, "Hits": "ACTB|ACTN1|CD9|CSRP1|HSPB1|ILK|MYH9|PIK3CB|PTPN6|TLN1|FIBP", "Description": "platelet aggregation", "_MEMBER_Input_ID": 1.0, "_GeneInHitList": 1562.0, "Category": "GO Biological Processes", "Log_q_value_": -1.3501258675186305, "Symbol": "GO:0070527", "LogP": -2.8692025675842725, "_RANK_": 1.0, "FirstInGroupByEnrichment": 0.0, "_TotalGeneInLibrary": 24367.0, "_GeneInGO": 60.0, "shared_name": "108", "_InGO": 0.704225352112676, "Gene": "GO:0070527", "_COUNT_001": 11.0, "SUID": 579305, "id_original": "108", "selected": false, "CategoryID": 19.0, "STDV_InGO": 0.21158296421416498, "EvidenceCutoff": 0.0, "PARENT_GO": "19_GO:0022610 biological adhesion", "FirstInGroupByLogP": 0.0, "GROUP_ID": 7.0, "name": "108", "BestEnrichmentInGroup": 2.8599765258215957, "CLUSTER_LABEL": " ", "GeneID": "60|87|928|1465|3315|3611|4627|5291|5777|7094|9158", "_LogP_Input_ID": -2.8692025675842725, "Z_score": 3.7751568838377088, "GiniIndex": 0.0, "_COUNT_001_PCT": 100.0}, "position": {"x": -352.30928446114604, "y": 716.9538837021331}, "selected": false}, {"data": {"id": "579304", "Enrichment": 1.671414852752881, "BestLogPInGroup": -9.631064746783506, "_PATTERN_": "M1", "_GeneInGOAndHitList": 36.0, "Hits": "ACTB|ACTN1|CAPZB|CD9|COL1A1|CSRP1|DGKA|DGKQ|FBLN1|GATA2|H3-3B|HSPB1|ILK|IRF1|ITPK1|ITPR1|MYH9|NFE2L2|PDGFB|PIK3CB|SERPINF2|PRKAR1A|PRKCH|PTPN6|RAD51B|SRF|PRDX2|TLN1|YWHAZ|H3C8|H3C7|FIBP|GNA13|EHD2|ADTRP|JMJD1C", "Description": "blood coagulation", "_MEMBER_Input_ID": 1.0, "_GeneInHitList": 1562.0, "Category": "GO Biological Processes", "Log_q_value_": -1.2662225633999553, "Symbol": "GO:0007596", "LogP": -2.7471094008021057, "_RANK_": 1.0, "FirstInGroupByEnrichment": 0.0, "_TotalGeneInLibrary": 24367.0, "_GeneInGO": 336.0, "shared_name": "107", "_InGO": 2.3047375160051216, "Gene": "GO:0007596", "_COUNT_001": 36.0, "SUID": 579304, "id_original": "107", "selected": false, "CategoryID": 19.0, "STDV_InGO": 0.37967060373244593, "EvidenceCutoff": 0.0, "PARENT_GO": "19_GO:0032501 multicellular organismal process", "FirstInGroupByLogP": 0.0, "GROUP_ID": 7.0, "name": "107", "BestEnrichmentInGroup": 2.8599765258215957, "CLUSTER_LABEL": " ", "GeneID": "60|87|832|928|1277|1465|1606|1609|2192|2624|3021|3315|3611|3659|3705|3708|4627|4780|5155|5291|5345|5573|5583|5777|5890|6722|7001|7094|7534|8355|8968|9158|10672|30846|84830|221037", "_LogP_Input_ID": -2.7471094008021057, "Z_score": 3.243338352918285, "GiniIndex": 0.0, "_COUNT_001_PCT": 100.0}, "position": {"x": -336.7590339309031, "y": 815.6308856552581}, "selected": false}, {"data": {"id": "579303", "Enrichment": 2.025957397276219, "BestLogPInGroup": -9.631064746783506, "_PATTERN_": "M1", "_GeneInGOAndHitList": 20.0, "Hits": "ACTB|ACTN1|CD9|COL1A1|CSRP1|DGKA|DGKQ|HSPB1|ILK|ITPR1|MYH9|PDGFB|PIK3CB|PRKCH|PTPN6|SRF|TLN1|YWHAZ|FIBP|GNA13", "Description": "platelet activation", "_MEMBER_Input_ID": 1.0, "_GeneInHitList": 1562.0, "Category": "GO Biological Processes", "Log_q_value_": -1.224398854713646, "Symbol": "GO:0030168", "LogP": -2.6904153841715566, "_RANK_": 1.0, "FirstInGroupByEnrichment": 0.0, "_TotalGeneInLibrary": 24367.0, "_GeneInGO": 154.0, "shared_name": "106", "_InGO": 1.2804097311139564, "Gene": "GO:0030168", "_COUNT_001": 20.0, "SUID": 579303, "id_original": "106", "selected": false, "CategoryID": 19.0, "STDV_InGO": 0.284469454773297, "EvidenceCutoff": 0.0, "PARENT_GO": "19_GO:0032501 multicellular organismal process", "FirstInGroupByLogP": 0.0, "GROUP_ID": 7.0, "name": "106", "BestEnrichmentInGroup": 2.8599765258215957, "CLUSTER_LABEL": " ", "GeneID": "60|87|928|1277|1465|1606|1609|3315|3611|3708|4627|5155|5291|5583|5777|6722|7094|7534|9158|10672", "_LogP_Input_ID": -2.6904153841715566, "Z_score": 3.3425890255464092, "GiniIndex": 0.0, "_COUNT_001_PCT": 100.0}, "position": {"x": -469.7723238593394, "y": 758.5205951279144}, "selected": false}, {"data": {"id": "579302", "Enrichment": 1.6469073035922817, "BestLogPInGroup": -9.631064746783506, "_PATTERN_": "M1", "_GeneInGOAndHitList": 36.0, "Hits": "ACTB|ACTN1|CAPZB|CD9|COL1A1|CSRP1|DGKA|DGKQ|FBLN1|GATA2|H3-3B|HSPB1|ILK|IRF1|ITPK1|ITPR1|MYH9|NFE2L2|PDGFB|PIK3CB|SERPINF2|PRKAR1A|PRKCH|PTPN6|RAD51B|SRF|PRDX2|TLN1|YWHAZ|H3C8|H3C7|FIBP|GNA13|EHD2|ADTRP|JMJD1C", "Description": "hemostasis", "_MEMBER_Input_ID": 1.0, "_GeneInHitList": 1562.0, "Category": "GO Biological Processes", "Log_q_value_": -1.1828036209589823, "Symbol": "GO:0007599", "LogP": -2.6378246843593196, "_RANK_": 1.0, "FirstInGroupByEnrichment": 0.0, "_TotalGeneInLibrary": 24367.0, "_GeneInGO": 341.0, "shared_name": "105", "_InGO": 2.3047375160051216, "Gene": "GO:0007599", "_COUNT_001": 36.0, "SUID": 579302, "id_original": "105", "selected": false, "CategoryID": 19.0, "STDV_InGO": 0.37967060373244593, "EvidenceCutoff": 0.0, "PARENT_GO": "19_GO:0065007 biological regulation", "FirstInGroupByLogP": 0.0, "GROUP_ID": 7.0, "name": "105", "BestEnrichmentInGroup": 2.8599765258215957, "CLUSTER_LABEL": " ", "GeneID": "60|87|832|928|1277|1465|1606|1609|2192|2624|3021|3315|3611|3659|3705|3708|4627|4780|5155|5291|5345|5573|5583|5777|5890|6722|7001|7094|7534|8355|8968|9158|10672|30846|84830|221037", "_LogP_Input_ID": -2.6378246843593196, "Z_score": 3.1484449482777923, "GiniIndex": 0.0, "_COUNT_001_PCT": 100.0}, "position": {"x": -427.11313654244486, "y": 728.5873370712737}, "selected": false}, {"data": {"id": "579301", "Enrichment": 1.6420917851607253, "BestLogPInGroup": -9.631064746783506, "_PATTERN_": "M1", "_GeneInGOAndHitList": 36.0, "Hits": "ACTB|ACTN1|CAPZB|CD9|COL1A1|CSRP1|DGKA|DGKQ|FBLN1|GATA2|H3-3B|HSPB1|ILK|IRF1|ITPK1|ITPR1|MYH9|NFE2L2|PDGFB|PIK3CB|SERPINF2|PRKAR1A|PRKCH|PTPN6|RAD51B|SRF|PRDX2|TLN1|YWHAZ|H3C8|H3C7|FIBP|GNA13|EHD2|ADTRP|JMJD1C", "Description": "coagulation", "_MEMBER_Input_ID": 1.0, "_GeneInHitList": 1562.0, "Category": "GO Biological Processes", "Log_q_value_": -1.1757824654387388, "Symbol": "GO:0050817", "LogP": -2.6164099499285847, "_RANK_": 1.0, "FirstInGroupByEnrichment": 0.0, "_TotalGeneInLibrary": 24367.0, "_GeneInGO": 342.0, "shared_name": "104", "_InGO": 2.3047375160051216, "Gene": "GO:0050817", "_COUNT_001": 36.0, "SUID": 579301, "id_original": "104", "selected": false, "CategoryID": 19.0, "STDV_InGO": 0.37967060373244593, "EvidenceCutoff": 0.0, "PARENT_GO": "19_GO:0032501 multicellular organismal process", "FirstInGroupByLogP": 0.0, "GROUP_ID": 7.0, "name": "104", "BestEnrichmentInGroup": 2.8599765258215957, "CLUSTER_LABEL": " ", "GeneID": "60|87|832|928|1277|1465|1606|1609|2192|2624|3021|3315|3611|3659|3705|3708|4627|4780|5155|5291|5345|5573|5583|5777|5890|6722|7001|7094|7534|8355|8968|9158|10672|30846|84830|221037", "_LogP_Input_ID": -2.6164099499285847, "Z_score": 3.1296521141162224, "GiniIndex": 0.0, "_COUNT_001_PCT": 100.0}, "position": {"x": -368.3075335151011, "y": 856.4522967880706}, "selected": false}, {"data": {"id": "579300", "Enrichment": 2.414265898420828, "BestLogPInGroup": -9.631064746783506, "_PATTERN_": "M1", "_GeneInGOAndHitList": 13.0, "Hits": "ACTB|ACTN1|ANK3|CD9|CSRP1|HSPB1|ILK|JUP|MYH9|PIK3CB|PTPN6|TLN1|FIBP", "Description": "homotypic cell-cell adhesion", "_MEMBER_Input_ID": 1.0, "_GeneInHitList": 1562.0, "Category": "GO Biological Processes", "Log_q_value_": -1.1612993096991469, "Symbol": "GO:0034109", "LogP": -2.588007346137889, "_RANK_": 1.0, "FirstInGroupByEnrichment": 0.0, "_TotalGeneInLibrary": 24367.0, "_GeneInGO": 84.0, "shared_name": "103", "_InGO": 0.8322663252240717, "Gene": "GO:0034109", "_COUNT_001": 13.0, "SUID": 579300, "id_original": "103", "selected": false, "CategoryID": 19.0, "STDV_InGO": 0.22986658336957838, "EvidenceCutoff": 0.0, "PARENT_GO": "19_GO:0022610 biological adhesion", "FirstInGroupByLogP": 0.0, "GROUP_ID": 7.0, "name": "103", "BestEnrichmentInGroup": 2.8599765258215957, "CLUSTER_LABEL": " ", "GeneID": "60|87|288|928|1465|3315|3611|3728|4627|5291|5777|7094|9158", "_LogP_Input_ID": -2.588007346137889, "Z_score": 3.398107697962696, "GiniIndex": 0.0, "_COUNT_001_PCT": 100.0}, "position": {"x": -450.4270327216441, "y": 839.4347796982269}, "selected": false}, {"data": {"id": "579299", "Enrichment": 1.4546504399092595, "BestLogPInGroup": -9.631064746783506, "_PATTERN_": "M1", "_GeneInGOAndHitList": 58.0, "Hits": "ABL1|ACTB|ACTN4|ACTN1|AKT1|ANXA2|ATP2A2|CALM1|CAPZB|CD9|CD63|CFL1|COL1A1|CTSW|DGKA|DGKQ|FGR|GATA2|GNB3|GPC1|GUCY1A1|H3-3B|IRF1|ITGA3|ITPK1|ITPR1|KIFC1|PDE9A|PDGFB|PIK3CB|SERPINF2|PPP2R5C|PRKAR1A|PRKCH|PTPN6|RAD51B|SDC1|SDC4|SHC1|SPARC|TLN1|YWHAZ|TUBA1A|H3C8|PIK3R3|H3C7|SLC16A3|PICK1|DOCK4|ABCC4|GNA13|ENDOD1|SLC16A8|EHD2|TUBA1C|JMJD1C|DAGLB|NHLRC2", "Description": "Hemostasis", "_MEMBER_Input_ID": 1.0, "_GeneInHitList": 1562.0, "Category": "Reactome Gene Sets", "Log_q_value_": -1.145509114443904, "Symbol": "R-HSA-109582", "LogP": -2.56424804509721, "_RANK_": 1.0, "FirstInGroupByEnrichment": 0.0, "_TotalGeneInLibrary": 24367.0, "_GeneInGO": 622.0, "shared_name": "102", "_InGO": 3.7131882202304736, "Gene": "R-HSA-109582", "_COUNT_001": 58.0, "SUID": 579299, "id_original": "102", "selected": false, "CategoryID": 6.0, "STDV_InGO": 0.47842775961251943, "EvidenceCutoff": 0.0, "PARENT_GO": "", "FirstInGroupByLogP": 0.0, "GROUP_ID": 7.0, "name": "102", "BestEnrichmentInGroup": 2.8599765258215957, "CLUSTER_LABEL": " ", "GeneID": "25|60|81|87|207|302|488|801|832|928|967|1072|1277|1521|1606|1609|2268|2624|2784|2817|2982|3021|3659|3675|3705|3708|3833|5152|5155|5291|5345|5527|5573|5583|5777|5890|6382|6385|6464|6678|7094|7534|7846|8355|8503|8968|9123|9463|9732|10257|10672|23052|23539|30846|84790|221037|221955|374354", "_LogP_Input_ID": -2.56424804509721, "Z_score": 3.006106187017174, "GiniIndex": 0.0, "_COUNT_001_PCT": 100.0}, "position": {"x": -487.60092188180033, "y": 822.9748797958831}, "selected": false}, {"data": {"id": "579298", "Enrichment": 3.1795917368717226, "BestLogPInGroup": -9.567032373961917, "_PATTERN_": "M1", "_GeneInGOAndHitList": 32.0, "Hits": "BCL6|ZFP36L1|RUNX1|CBFB|ERBB2|GATA2|HMGB1|HOXA5|HOXA7|HOXA9|ID2|IL4R|IRF1|ITPKB|SMAD7|MEIS1|MEIS2|NFE2L2|NOTCH1|RARA|PRDX2|ZFP36|ZBTB16|TRIB1|N4BP2L2|ZBTB7B|KLF13|PAF1|ZC3H12A|MIR125B1|MIR21|MIR30B", "Description": "negative regulation of hemopoiesis", "_MEMBER_Input_ID": 1.0, "_GeneInHitList": 1562.0, "Category": "GO Biological Processes", "Log_q_value_": -5.393878257702022, "Symbol": "GO:1903707", "LogP": -8.3436993759719, "_RANK_": 1.0, "FirstInGroupByEnrichment": 0.0, "_TotalGeneInLibrary": 24367.0, "_GeneInGO": 157.0, "shared_name": "101", "_InGO": 2.0486555697823303, "Gene": "GO:1903707", "_COUNT_001": 32.0, "SUID": 579298, "id_original": "101", "selected": false, "CategoryID": 19.0, "STDV_InGO": 0.35842571499821285, "EvidenceCutoff": 0.0, "PARENT_GO": "19_GO:0002376 immune system process", "FirstInGroupByLogP": 0.0, "GROUP_ID": 8.0, "name": "101", "BestEnrichmentInGroup": 3.1795917368717226, "CLUSTER_LABEL": " ", "GeneID": "604|677|861|865|2064|2624|3146|3202|3204|3205|3398|3566|3659|3707|4092|4211|4212|4780|4851|5914|7001|7538|7704|10221|10443|51043|51621|54623|80149|406911|406991|407030", "_LogP_Input_ID": -8.3436993759719, "Z_score": 7.170429673538275, "GiniIndex": 0.0, "_COUNT_001_PCT": 100.0}, "position": {"x": -173.08385492623393, "y": 184.77954631504815}, "selected": false}, {"data": {"id": "579297", "Enrichment": 1.8732765499452575, "BestLogPInGroup": -9.567032373961917, "_PATTERN_": "M1", "_GeneInGOAndHitList": 58.0, "Hits": "AKT1|BCL6|ZFP36L1|RUNX1|CBFB|CEBPB|DPP4|ELF1|ERBB2|FGR|FOXF1|GATA2|HMGB1|HOXA5|HOXA7|HOXA9|ID2|IFI16|IL4R|IRF1|ITPKB|SMAD7|MEIS1|MEIS2|NBL1|NFE2L2|NOTCH1|PRKAR1A|PTPN6|PTPRJ|RARA|CX3CL1|SDC4|STAT6|TARBP2|PRDX2|EZR|ZFP36|ZBTB16|DLG5|NR1D1|LRRC14|TRIB1|N4BP2L2|ZBTB7B|KLF13|PAF1|BANK1|VSIR|ILRUN|ZC3H12A|ADTRP|IL31RA|DAB2IP|MIA3|MIR125B1|MIR21|MIR30B", "Description": "negative regulation of immune system process", "_MEMBER_Input_ID": 1.0, "_GeneInHitList": 1562.0, "Category": "GO Biological Processes", "Log_q_value_": -3.2633690818392007, "Symbol": "GO:0002683", "LogP": -5.503293895101352, "_RANK_": 1.0, "FirstInGroupByEnrichment": 0.0, "_TotalGeneInLibrary": 24367.0, "_GeneInGO": 483.0, "shared_name": "100", "_InGO": 3.7131882202304736, "Gene": "GO:0002683", "_COUNT_001": 58.0, "SUID": 579297, "id_original": "100", "selected": false, "CategoryID": 19.0, "STDV_InGO": 0.47842775961251943, "EvidenceCutoff": 0.0, "PARENT_GO": "19_GO:0002376 immune system process", "FirstInGroupByLogP": 0.0, "GROUP_ID": 8.0, "name": "100", "BestEnrichmentInGroup": 3.1795917368717226, "CLUSTER_LABEL": " ", "GeneID": "207|604|677|861|865|1051|1803|1997|2064|2268|2294|2624|3146|3202|3204|3205|3398|3428|3566|3659|3707|4092|4211|4212|4681|4780|4851|5573|5777|5795|5914|6376|6385|6778|6895|7001|7430|7538|7704|9231|9572|9684|10221|10443|51043|51621|54623|55024|64115|64771|80149|84830|133396|153090|375056|406911|406991|407030", "_LogP_Input_ID": -5.503293895101352, "Z_score": 5.073286969142864, "GiniIndex": 0.0, "_COUNT_001_PCT": 100.0}, "position": {"x": -148.19892908394877, "y": 139.34817844090264}, "selected": false}, {"data": {"id": "579296", "Enrichment": 2.489341270057482, "BestLogPInGroup": -9.567032373961917, "_PATTERN_": "M1", "_GeneInGOAndHitList": 15.0, "Hits": "ZFP36L1|GATA2|HOXA5|HOXA7|HOXA9|ITPKB|MEIS1|MEIS2|RARA|ZFP36|ZBTB16|TRIB1|KLF13|PAF1|MIR125B1", "Description": "negative regulation of myeloid cell differentiation", "_MEMBER_Input_ID": 1.0, "_GeneInHitList": 1562.0, "Category": "GO Biological Processes", "Log_q_value_": -1.4742927997790294, "Symbol": "GO:0045638", "LogP": -3.0405824279005342, "_RANK_": 1.0, "FirstInGroupByEnrichment": 0.0, "_TotalGeneInLibrary": 24367.0, "_GeneInGO": 94.0, "shared_name": "99", "_InGO": 0.9603072983354674, "Gene": "GO:0045638", "_COUNT_001": 15.0, "SUID": 579296, "id_original": "99", "selected": false, "CategoryID": 19.0, "STDV_InGO": 0.24675686391426488, "EvidenceCutoff": 0.0, "PARENT_GO": "19_GO:0002376 immune system process", "FirstInGroupByLogP": 0.0, "GROUP_ID": 8.0, "name": "99", "BestEnrichmentInGroup": 3.1795917368717226, "CLUSTER_LABEL": " ", "GeneID": "677|2624|3202|3204|3205|3707|4211|4212|5914|7538|7704|10221|51621|54623|406911", "_LogP_Input_ID": -3.0405824279005342, "Z_score": 3.786291263501722, "GiniIndex": 0.0, "_COUNT_001_PCT": 100.0}, "position": {"x": -238.02709986031596, "y": 231.78586536106866}, "selected": false}, {"data": {"id": "579295", "Enrichment": 2.8297442158234816, "BestLogPInGroup": -9.213753097063153, "_PATTERN_": "M1", "_GeneInGOAndHitList": 39.0, "Hits": "ABL1|ACTN4|BCL6|CCR7|COL1A1|FBLN1|FOXF1|HOXA7|ILK|ITGA3|JUP|LIMS1|LRP1|SMAD3|MMP14|NEDD9|NINJ1|NOTCH1|P4HB|PIK3CB|PTPRJ|RAC3|RRAS|RREB1|CX3CL1|SDC4|SPOCK1|TBCD|UTRN|AGR2|TRIOBP|LIMCH1|KANK1|MACF1|CORO1C|RIN2|CAMSAP3|PLEKHA2|ACER2", "Description": "regulation of cell-substrate adhesion", "_MEMBER_Input_ID": 1.0, "_GeneInHitList": 1562.0, "Category": "GO Biological Processes", "Log_q_value_": -5.47857610632771, "Symbol": "GO:0010810", "LogP": -8.475462870903566, "_RANK_": 1.0, "FirstInGroupByEnrichment": 0.0, "_TotalGeneInLibrary": 24367.0, "_GeneInGO": 215.0, "shared_name": "98", "_InGO": 2.496798975672215, "Gene": "GO:0010810", "_COUNT_001": 39.0, "SUID": 579295, "id_original": "98", "selected": false, "CategoryID": 19.0, "STDV_InGO": 0.3947850615732865, "EvidenceCutoff": 0.0, "PARENT_GO": "19_GO:0022610 biological adhesion", "FirstInGroupByLogP": 0.0, "GROUP_ID": 9.0, "name": "98", "BestEnrichmentInGroup": 4.120720894837292, "CLUSTER_LABEL": " ", "GeneID": "25|81|604|1236|1277|2192|2294|3204|3611|3675|3728|3987|4035|4088|4323|4739|4814|4851|5034|5291|5795|5881|6237|6239|6376|6385|6695|6904|7402|10551|11078|22998|23189|23499|23603|54453|57662|59339|340485", "_LogP_Input_ID": -8.475462870903566, "Z_score": 7.052622840603081, "GiniIndex": 0.0, "_COUNT_001_PCT": 100.0}, "position": {"x": -77.20679308236186, "y": -94.04764217677314}, "selected": false}, {"data": {"id": "579294", "Enrichment": 2.3421904074459636, "BestLogPInGroup": -9.213753097063153, "_PATTERN_": "M1", "_GeneInGOAndHitList": 53.0, "Hits": "ABL1|ACTN4|ACTN1|BCL6|CD63|CCR7|COL1A1|FBLN1|FOXF1|HOXA7|ID1|ILK|ITGA3|ITGA7|ITGB7|JUP|LIMS1|LRP1|BCAM|SMAD3|MMP14|NEDD9|NINJ1|NOTCH1|P4HB|PIK3CB|PTPRJ|RAC3|RRAS|RREB1|CX3CL1|SDC4|SPOCK1|SRF|TBCD|TRIP6|UTRN|AGR2|SORBS1|TRIOBP|LIMCH1|KANK1|MACF1|CORO1C|SNED1|TNFRSF12A|RIN2|PARVA|ADAMTS9|CAMSAP3|PLEKHA2|MICALL2|ACER2", "Description": "cell-substrate adhesion", "_MEMBER_Input_ID": 1.0, "_GeneInHitList": 1562.0, "Category": "GO Biological Processes", "Log_q_value_": -5.281503596852938, "Symbol": "GO:0031589", "LogP": -8.185636308191896, "_RANK_": 1.0, "FirstInGroupByEnrichment": 0.0, "_TotalGeneInLibrary": 24367.0, "_GeneInGO": 353.0, "shared_name": "97", "_InGO": 3.3930857874519846, "Gene": "GO:0031589", "_COUNT_001": 53.0, "SUID": 579294, "id_original": "97", "selected": false, "CategoryID": 19.0, "STDV_InGO": 0.4581007579903272, "EvidenceCutoff": 0.0, "PARENT_GO": "19_GO:0022610 biological adhesion", "FirstInGroupByLogP": 0.0, "GROUP_ID": 9.0, "name": "97", "BestEnrichmentInGroup": 4.120720894837292, "CLUSTER_LABEL": " ", "GeneID": "25|81|87|604|967|1236|1277|2192|2294|3204|3397|3611|3675|3679|3695|3728|3987|4035|4059|4088|4323|4739|4814|4851|5034|5291|5795|5881|6237|6239|6376|6385|6695|6722|6904|7205|7402|10551|10580|11078|22998|23189|23499|23603|25992|51330|54453|55742|56999|57662|59339|79778|340485", "_LogP_Input_ID": -8.185636308191896, "Z_score": 6.647935950352027, "GiniIndex": 0.0, "_COUNT_001_PCT": 100.0}, "position": {"x": -35.26066423714701, "y": -65.30893368067939}, "selected": false}, {"data": {"id": "579293", "Enrichment": 2.452153786201551, "BestLogPInGroup": -9.213753097063153, "_PATTERN_": "M1", "_GeneInGOAndHitList": 47.0, "Hits": "ABL1|ACTB|ACTN4|ACTN1|DST|RUNX1|CBFB|CD9|CLDN3|EPHA2|RAPGEF1|ILK|JUP|LAMA3|LIMS1|LRP1|SMAD3|SMAD7|MMP14|CLDN11|PLEC|PRKCH|PTPRJ|NECTIN1|SDC4|SRF|TBCD|TJP1|TLN1|TNS1|TRIP6|CNTNAP1|CLDN12|CLDN6|CLDN9|DLG5|FLOT1|SORBS1|LIMCH1|MACF1|CORO1C|HDAC7|CAMSAP3|CDH23|MICALL2|GRHL2|VMP1", "Description": "cell junction organization", "_MEMBER_Input_ID": 1.0, "_GeneInHitList": 1562.0, "Category": "GO Biological Processes", "Log_q_value_": -5.094886003491915, "Symbol": "GO:0034330", "LogP": -7.966834031459472, "_RANK_": 1.0, "FirstInGroupByEnrichment": 0.0, "_TotalGeneInLibrary": 24367.0, "_GeneInGO": 299.0, "shared_name": "96", "_InGO": 3.0089628681177976, "Gene": "GO:0034330", "_COUNT_001": 47.0, "SUID": 579293, "id_original": "96", "selected": false, "CategoryID": 19.0, "STDV_InGO": 0.43224870525504855, "EvidenceCutoff": 0.0, "PARENT_GO": "19_GO:0071840 cellular component organization or biogenesis", "FirstInGroupByLogP": 0.0, "GROUP_ID": 9.0, "name": "96", "BestEnrichmentInGroup": 4.120720894837292, "CLUSTER_LABEL": " ", "GeneID": "25|60|81|87|667|861|865|928|1365|1969|2889|3611|3728|3909|3987|4035|4088|4092|4323|5010|5339|5583|5795|5818|6385|6722|6904|7082|7094|7145|7205|8506|9069|9074|9080|9231|10211|10580|22998|23499|23603|51564|57662|64072|79778|79977|81671", "_LogP_Input_ID": -7.966834031459472, "Z_score": 6.612197250847835, "GiniIndex": 0.0, "_COUNT_001_PCT": 100.0}, "position": {"x": 38.69114659964498, "y": -142.0643047744294}, "selected": false}, {"data": {"id": "579292", "Enrichment": 3.043877455419881, "BestLogPInGroup": -9.213753097063153, "_PATTERN_": "M1", "_GeneInGOAndHitList": 24.0, "Hits": "ABL1|BCL6|CCR7|HOXA7|ILK|JUP|LIMS1|LRP1|SMAD3|MMP14|NINJ1|PIK3CB|PTPRJ|RRAS|CX3CL1|SDC4|UTRN|LIMCH1|MACF1|CORO1C|RIN2|CAMSAP3|PLEKHA2|ACER2", "Description": "regulation of cell-matrix adhesion", "_MEMBER_Input_ID": 1.0, "_GeneInHitList": 1562.0, "Category": "GO Biological Processes", "Log_q_value_": -3.6483039689004197, "Symbol": "GO:0001952", "LogP": -6.064320041218252, "_RANK_": 1.0, "FirstInGroupByEnrichment": 0.0, "_TotalGeneInLibrary": 24367.0, "_GeneInGO": 123.0, "shared_name": "95", "_InGO": 1.5364916773367479, "Gene": "GO:0001952", "_COUNT_001": 24.0, "SUID": 579292, "id_original": "95", "selected": false, "CategoryID": 19.0, "STDV_InGO": 0.3112162348789158, "EvidenceCutoff": 0.0, "PARENT_GO": "19_GO:0022610 biological adhesion", "FirstInGroupByLogP": 0.0, "GROUP_ID": 9.0, "name": "95", "BestEnrichmentInGroup": 4.120720894837292, "CLUSTER_LABEL": " ", "GeneID": "25|604|1236|3204|3611|3728|3987|4035|4088|4323|4814|5291|5795|6237|6376|6385|7402|22998|23499|23603|54453|57662|59339|340485", "_LogP_Input_ID": -6.064320041218252, "Z_score": 5.947341377929819, "GiniIndex": 0.0, "_COUNT_001_PCT": 100.0}, "position": {"x": -37.19458700478617, "y": -129.9298138076325}, "selected": false}, {"data": {"id": "579291", "Enrichment": 2.3842599064014895, "BestLogPInGroup": -9.213753097063153, "_PATTERN_": "M1", "_GeneInGOAndHitList": 35.0, "Hits": "ABL1|ACTN1|BCL6|CD63|CCR7|HOXA7|ILK|ITGA3|ITGA7|ITGB7|JUP|LIMS1|LRP1|BCAM|SMAD3|MMP14|NINJ1|PIK3CB|PTPRJ|RRAS|CX3CL1|SDC4|SRF|TRIP6|UTRN|SORBS1|LIMCH1|MACF1|CORO1C|SNED1|RIN2|ADAMTS9|CAMSAP3|PLEKHA2|ACER2", "Description": "cell-matrix adhesion", "_MEMBER_Input_ID": 1.0, "_GeneInHitList": 1562.0, "Category": "GO Biological Processes", "Log_q_value_": -3.465377983798246, "Symbol": "GO:0007160", "LogP": -5.816001094554087, "_RANK_": 1.0, "FirstInGroupByEnrichment": 0.0, "_TotalGeneInLibrary": 24367.0, "_GeneInGO": 229.0, "shared_name": "94", "_InGO": 2.240717029449424, "Gene": "GO:0007160", "_COUNT_001": 35.0, "SUID": 579291, "id_original": "94", "selected": false, "CategoryID": 19.0, "STDV_InGO": 0.3744829043781526, "EvidenceCutoff": 0.0, "PARENT_GO": "19_GO:0022610 biological adhesion", "FirstInGroupByLogP": 0.0, "GROUP_ID": 9.0, "name": "94", "BestEnrichmentInGroup": 4.120720894837292, "CLUSTER_LABEL": " ", "GeneID": "25|87|604|967|1236|3204|3611|3675|3679|3695|3728|3987|4035|4059|4088|4323|4814|5291|5795|6237|6376|6385|6722|7205|7402|10580|22998|23499|23603|25992|54453|56999|57662|59339|340485", "_LogP_Input_ID": -5.816001094554087, "Z_score": 5.508106595224043, "GiniIndex": 0.0, "_COUNT_001_PCT": 100.0}, "position": {"x": -82.92107034981791, "y": -152.5463604384919}, "selected": false}, {"data": {"id": "579290", "Enrichment": 4.120720894837292, "BestLogPInGroup": -9.213753097063153, "_PATTERN_": "M1", "_GeneInGOAndHitList": 14.0, "Hits": "ABL1|CCR7|ILK|JUP|LIMS1|SMAD3|NINJ1|PTPRJ|RRAS|CX3CL1|SDC4|UTRN|RIN2|PLEKHA2", "Description": "positive regulation of cell-matrix adhesion", "_MEMBER_Input_ID": 1.0, "_GeneInHitList": 1562.0, "Category": "GO Biological Processes", "Log_q_value_": -3.1784057418545855, "Symbol": "GO:0001954", "LogP": -5.376937869958512, "_RANK_": 1.0, "FirstInGroupByEnrichment": 0.0, "_TotalGeneInLibrary": 24367.0, "_GeneInGO": 53.0, "shared_name": "93", "_InGO": 0.8962868117797695, "Gene": "GO:0001954", "_COUNT_001": 14.0, "SUID": 579290, "id_original": "93", "selected": false, "CategoryID": 19.0, "STDV_InGO": 0.23846681525444388, "EvidenceCutoff": 0.0, "PARENT_GO": "19_GO:0022610 biological adhesion", "FirstInGroupByLogP": 0.0, "GROUP_ID": 9.0, "name": "93", "BestEnrichmentInGroup": 4.120720894837292, "CLUSTER_LABEL": " ", "GeneID": "25|1236|3611|3728|3987|4088|4814|5795|6237|6376|6385|7402|54453|59339", "_LogP_Input_ID": -5.376937869958512, "Z_score": 5.952264940970148, "GiniIndex": 0.0, "_COUNT_001_PCT": 100.0}, "position": {"x": -158.53537012398783, "y": -151.58304256739814}, "selected": false}, {"data": {"id": "579289", "Enrichment": 2.8599765258215957, "BestLogPInGroup": -9.213753097063153, "_PATTERN_": "M1", "_GeneInGOAndHitList": 22.0, "Hits": "ABL1|CCR7|FOXF1|ILK|ITGA3|JUP|LIMS1|SMAD3|NEDD9|NINJ1|P4HB|PTPRJ|RAC3|RRAS|RREB1|CX3CL1|SDC4|UTRN|AGR2|TRIOBP|RIN2|PLEKHA2", "Description": "positive regulation of cell-substrate adhesion", "_MEMBER_Input_ID": 1.0, "_GeneInHitList": 1562.0, "Category": "GO Biological Processes", "Log_q_value_": -3.0051670699759185, "Symbol": "GO:0010811", "LogP": -5.151086860873421, "_RANK_": 1.0, "FirstInGroupByEnrichment": 0.0, "_TotalGeneInLibrary": 24367.0, "_GeneInGO": 120.0, "shared_name": "92", "_InGO": 1.408450704225352, "Gene": "GO:0010811", "_COUNT_001": 22.0, "SUID": 579289, "id_original": "92", "selected": false, "CategoryID": 19.0, "STDV_InGO": 0.29816053328431674, "EvidenceCutoff": 0.0, "PARENT_GO": "19_GO:0022610 biological adhesion", "FirstInGroupByLogP": 0.0, "GROUP_ID": 9.0, "name": "92", "BestEnrichmentInGroup": 4.120720894837292, "CLUSTER_LABEL": " ", "GeneID": "25|1236|2294|3611|3675|3728|3987|4088|4739|4814|5034|5795|5881|6237|6239|6376|6385|7402|10551|11078|54453|59339", "_LogP_Input_ID": -5.151086860873421, "Z_score": 5.345479598600731, "GiniIndex": 0.0, "_COUNT_001_PCT": 100.0}, "position": {"x": -151.3410494452769, "y": -77.48938412013251}, "selected": false}, {"data": {"id": "579288", "Enrichment": 3.087474658557405, "BestLogPInGroup": -9.213753097063153, "_PATTERN_": "M1", "_GeneInGOAndHitList": 19.0, "Hits": "ABL1|RUNX1|CBFB|CLDN3|EPHA2|RAPGEF1|LIMS1|LRP1|SMAD3|MMP14|PRKCH|PTPRJ|SDC4|TJP1|FLOT1|LIMCH1|MACF1|CORO1C|CAMSAP3", "Description": "regulation of cell junction assembly", "_MEMBER_Input_ID": 1.0, "_GeneInHitList": 1562.0, "Category": "GO Biological Processes", "Log_q_value_": -2.911276951118425, "Symbol": "GO:1901888", "LogP": -5.028985886107749, "_RANK_": 1.0, "FirstInGroupByEnrichment": 0.0, "_TotalGeneInLibrary": 24367.0, "_GeneInGO": 96.0, "shared_name": "91", "_InGO": 1.2163892445582587, "Gene": "GO:1901888", "_COUNT_001": 19.0, "SUID": 579288, "id_original": "91", "selected": false, "CategoryID": 19.0, "STDV_InGO": 0.2773564175744805, "EvidenceCutoff": 0.0, "PARENT_GO": "19_GO:0071840 cellular component organization or biogenesis", "FirstInGroupByLogP": 0.0, "GROUP_ID": 9.0, "name": "91", "BestEnrichmentInGroup": 4.120720894837292, "CLUSTER_LABEL": " ", "GeneID": "25|861|865|1365|1969|2889|3987|4035|4088|4323|5583|5795|6385|7082|10211|22998|23499|23603|57662", "_LogP_Input_ID": -5.028985886107749, "Z_score": 5.363281470344731, "GiniIndex": 0.0, "_COUNT_001_PCT": 100.0}, "position": {"x": -44.8193721896464, "y": -227.15039486232}, "selected": false}, {"data": {"id": "579287", "Enrichment": 2.993914820419302, "BestLogPInGroup": -9.213753097063153, "_PATTERN_": "M1", "_GeneInGOAndHitList": 19.0, "Hits": "ABL1|ACTN1|DST|LAMA3|LIMS1|LRP1|SMAD3|MMP14|PLEC|PTPRJ|SDC4|TLN1|TNS1|TRIP6|SORBS1|LIMCH1|MACF1|CORO1C|CAMSAP3", "Description": "cell-substrate junction assembly", "_MEMBER_Input_ID": 1.0, "_GeneInHitList": 1562.0, "Category": "GO Biological Processes", "Log_q_value_": -2.772230134994353, "Symbol": "GO:0007044", "LogP": -4.827991014206236, "_RANK_": 1.0, "FirstInGroupByEnrichment": 0.0, "_TotalGeneInLibrary": 24367.0, "_GeneInGO": 99.0, "shared_name": "90", "_InGO": 1.2163892445582587, "Gene": "GO:0007044", "_COUNT_001": 19.0, "SUID": 579287, "id_original": "90", "selected": false, "CategoryID": 19.0, "STDV_InGO": 0.2773564175744805, "EvidenceCutoff": 0.0, "PARENT_GO": "19_GO:0071840 cellular component organization or biogenesis", "FirstInGroupByLogP": 0.0, "GROUP_ID": 9.0, "name": "90", "BestEnrichmentInGroup": 4.120720894837292, "CLUSTER_LABEL": " ", "GeneID": "25|87|667|3909|3987|4035|4088|4323|5339|5795|6385|7094|7145|7205|10580|22998|23499|23603|57662", "_LogP_Input_ID": -4.827991014206236, "Z_score": 5.202652295103474, "GiniIndex": 0.0, "_COUNT_001_PCT": 100.0}, "position": {"x": 19.101035820836387, "y": -202.1852764541169}, "selected": false}, {"data": {"id": "579286", "Enrichment": 1.959916001643446, "BestLogPInGroup": -9.059097575411174, "_PATTERN_": "M1", "_GeneInGOAndHitList": 74.0, "Hits": "ZFP36L1|CDKN1B|DNAH5|ECE1|EGFR|EPHB4|ERBB2|ERBB3|FGF12|FHL2|FOXF1|MTOR|HNRNPU|ID1|ID2|ID3|ILK|ISL1|ITGA3|LMNA|SMAD3|SMAD7|MEF2D|MEIS1|NKX3-1|NOTCH1|OXCT1|PDGFB|PRKAR1A|PTCH1|PTK7|RARA|RARB|RB1|RXRA|SHC1|STIL|SMARCD3|SOX4|SPARC|SRF|TBX3|TNFRSF1A|SORBS2|PDLIM4|DCHS1|DHRS3|SNX17|RB1CC1|ZMPSTE24|CITED2|PDLIM5|HEXIM1|NCOA6|KDM6B|JMJD6|HEY2|IRX4|CCDC40|PARVA|MESP1|ADAMTS9|RTN4|ADGRG6|GRHL2|ALPK2|MTERF4|MESP2|SIK1|ARL13B|NEK8|SH3PXD2B|MIR21|MIR25", "Description": "heart development", "_MEMBER_Input_ID": 1.0, "_GeneInHitList": 1562.0, "Category": "GO Biological Processes", "Log_q_value_": -4.835123316708915, "Symbol": "GO:0007507", "LogP": -7.649079397698785, "_RANK_": 1.0, "FirstInGroupByEnrichment": 0.0, "_TotalGeneInLibrary": 24367.0, "_GeneInGO": 589.0, "shared_name": "89", "_InGO": 4.7375160051216385, "Gene": "GO:0007507", "_COUNT_001": 74.0, "SUID": 579286, "id_original": "89", "selected": false, "CategoryID": 19.0, "STDV_InGO": 0.5375214271483969, "EvidenceCutoff": 0.0, "PARENT_GO": "19_GO:0032502 developmental process", "FirstInGroupByLogP": 0.0, "GROUP_ID": 11.0, "name": "89", "BestEnrichmentInGroup": 2.5527063205680363, "CLUSTER_LABEL": " ", "GeneID": "677|1027|1767|1889|1956|2050|2064|2065|2257|2274|2294|2475|3192|3397|3398|3399|3611|3670|3675|4000|4088|4092|4209|4211|4824|4851|5019|5155|5573|5727|5754|5914|5915|5925|6256|6464|6491|6604|6659|6678|6722|6926|7132|8470|8572|8642|9249|9784|9821|10269|10370|10611|10614|23054|23135|23210|23493|50805|55036|55742|55897|56999|57142|57211|79977|115701|130916|145873|150094|200894|284086|285590|406991|407014", "_LogP_Input_ID": -7.649079397698785, "Z_score": 6.171927470558202, "GiniIndex": 0.0, "_COUNT_001_PCT": 100.0}, "position": {"x": 800.0211789482778, "y": -216.9142803603669}, "selected": false}, {"data": {"id": "579285", "Enrichment": 2.1309103449186213, "BestLogPInGroup": -9.059097575411174, "_PATTERN_": "M1", "_GeneInGOAndHitList": 53.0, "Hits": "ABL1|AKT1|ATP2A2|CD9|CSRP1|FHL2|FOXF1|MTOR|H3-3B|HNRNPU|IL4R|KRT8|LMNA|MMP14|MYH9|NFATC3|NOTCH1|PDGFB|PPP3CA|PRKAR1A|RARA|RARB|RB1|RXRA|SDC1|SMARCD3|SRF|TARBP2|TBX3|XBP1|SORBS2|CFLAR|SPAG9|QKI|GDF15|FLOT1|ZMPSTE24|SPEG|PDLIM5|KDM6B|HEY2|MYOF|EHD2|PLEKHO1|SEMA4C|MESP1|NIBAN2|TANC1|ALPK2|SIK1|BHLHA15|MIR125B1|MIR21", "Description": "muscle cell differentiation", "_MEMBER_Input_ID": 1.0, "_GeneInHitList": 1562.0, "Category": "GO Biological Processes", "Log_q_value_": -4.210600614899919, "Symbol": "GO:0042692", "LogP": -6.797312914386727, "_RANK_": 1.0, "FirstInGroupByEnrichment": 0.0, "_TotalGeneInLibrary": 24367.0, "_GeneInGO": 388.0, "shared_name": "88", "_InGO": 3.3930857874519846, "Gene": "GO:0042692", "_COUNT_001": 53.0, "SUID": 579285, "id_original": "88", "selected": false, "CategoryID": 19.0, "STDV_InGO": 0.4581007579903272, "EvidenceCutoff": 0.0, "PARENT_GO": "19_GO:0032502 developmental process", "FirstInGroupByLogP": 0.0, "GROUP_ID": 11.0, "name": "88", "BestEnrichmentInGroup": 2.5527063205680363, "CLUSTER_LABEL": " ", "GeneID": "25|207|488|928|1465|2274|2294|2475|3021|3192|3566|3856|4000|4323|4627|4775|4851|5155|5530|5573|5914|5915|5925|6256|6382|6604|6722|6895|6926|7494|8470|8837|9043|9444|9518|10211|10269|10290|10611|23135|23493|26509|30846|51177|54910|55897|64855|85461|115701|150094|168620|406911|406991", "_LogP_Input_ID": -6.797312914386727, "Z_score": 5.87687024597664, "GiniIndex": 0.0, "_COUNT_001_PCT": 100.0}, "position": {"x": 863.6219174736684, "y": -227.15039486232}, "selected": false}, {"data": {"id": "579284", "Enrichment": 2.1152368758002558, "BestLogPInGroup": -9.059097575411174, "_PATTERN_": "M1", "_GeneInGOAndHitList": 40.0, "Hits": "AKT1|CD9|CSRP1|FHL2|MTOR|HNRNPU|IL4R|KRT8|LMNA|MMP14|MYH9|NOTCH1|PPP3CA|PRKAR1A|RARA|RARB|RB1|RXRA|SDC1|SRF|TBX3|XBP1|SORBS2|CFLAR|SPAG9|GDF15|FLOT1|ZMPSTE24|PDLIM5|KDM6B|HEY2|MYOF|EHD2|PLEKHO1|MESP1|NIBAN2|TANC1|ALPK2|SIK1|BHLHA15", "Description": "striated muscle cell differentiation", "_MEMBER_Input_ID": 1.0, "_GeneInHitList": 1562.0, "Category": "GO Biological Processes", "Log_q_value_": -3.0545246690799845, "Symbol": "GO:0051146", "LogP": -5.214926268065006, "_RANK_": 1.0, "FirstInGroupByEnrichment": 0.0, "_TotalGeneInLibrary": 24367.0, "_GeneInGO": 295.0, "shared_name": "87", "_InGO": 2.560819462227913, "Gene": "GO:0051146", "_COUNT_001": 40.0, "SUID": 579284, "id_original": "87", "selected": false, "CategoryID": 19.0, "STDV_InGO": 0.3996830930161245, "EvidenceCutoff": 0.0, "PARENT_GO": "19_GO:0032502 developmental process", "FirstInGroupByLogP": 0.0, "GROUP_ID": 11.0, "name": "87", "BestEnrichmentInGroup": 2.5527063205680363, "CLUSTER_LABEL": " ", "GeneID": "207|928|1465|2274|2475|3192|3566|3856|4000|4323|4627|4851|5530|5573|5914|5915|5925|6256|6382|6722|6926|7494|8470|8837|9043|9518|10211|10269|10611|23135|23493|26509|30846|51177|55897|64855|85461|115701|150094|168620", "_LogP_Input_ID": -5.214926268065006, "Z_score": 5.043588411308091, "GiniIndex": 0.0, "_COUNT_001_PCT": 100.0}, "position": {"x": 919.9378507012075, "y": -188.40428585353095}, "selected": false}, {"data": {"id": "579283", "Enrichment": 1.8689333153846395, "BestLogPInGroup": -9.059097575411174, "_PATTERN_": "M1", "_GeneInGOAndHitList": 49.0, "Hits": "CSRP1|DDX5|ERBB3|EYA2|FHL2|FLNB|FOS|MTOR|GPC1|HNRNPU|HOXD10|ID2|ISL1|LMNA|SMAD3|SMAD7|MEF2D|MEIS1|NOTCH1|PPP3CA|PRKAR1A|PTCH1|RARA|RARB|RB1|RXRA|SRF|TBX3|ZFAND5|BTG2|SORBS2|YBX3|CFLAR|FLOT1|ZMPSTE24|DDX17|PDLIM5|KDM6B|HEY2|TIPARP|MYLPF|MESP1|ADAMTS9|NIBAN2|UQCC2|ALPK2|SIK1|BCL9L|MIR25", "Description": "muscle tissue development", "_MEMBER_Input_ID": 1.0, "_GeneInHitList": 1562.0, "Category": "GO Biological Processes", "Log_q_value_": -2.6860983876845537, "Symbol": "GO:0060537", "LogP": -4.723299851753482, "_RANK_": 1.0, "FirstInGroupByEnrichment": 0.0, "_TotalGeneInLibrary": 24367.0, "_GeneInGO": 409.0, "shared_name": "86", "_InGO": 3.1370038412291934, "Gene": "GO:0060537", "_COUNT_001": 49.0, "SUID": 579283, "id_original": "86", "selected": false, "CategoryID": 19.0, "STDV_InGO": 0.44105825989215863, "EvidenceCutoff": 0.0, "PARENT_GO": "19_GO:0032502 developmental process", "FirstInGroupByLogP": 0.0, "GROUP_ID": 11.0, "name": "86", "BestEnrichmentInGroup": 2.5527063205680363, "CLUSTER_LABEL": " ", "GeneID": "1465|1655|2065|2139|2274|2317|2353|2475|2817|3192|3236|3398|3670|4000|4088|4092|4209|4211|4851|5530|5573|5727|5914|5915|5925|6256|6722|6926|7763|7832|8470|8531|8837|10211|10269|10521|10611|23135|23493|25976|29895|55897|56999|64855|84300|115701|150094|283149|407014", "_LogP_Input_ID": -4.723299851753482, "Z_score": 4.638103318530384, "GiniIndex": 0.0, "_COUNT_001_PCT": 100.0}, "position": {"x": 814.7728726738637, "y": -88.68295711817939}, "selected": false}, {"data": {"id": "579282", "Enrichment": 2.0321672283789813, "BestLogPInGroup": -9.059097575411174, "_PATTERN_": "M1", "_GeneInGOAndHitList": 34.0, "Hits": "EPHB4|FHL2|FOXF1|MTOR|ID2|ILK|ISL1|SMAD3|SMAD7|NOTCH1|PTCH1|RARA|RARB|RXRA|STIL|SMARCD3|SOX4|SRF|TBX3|DCHS1|DHRS3|ZMPSTE24|CITED2|HEY2|CCDC40|PARVA|MESP1|RTN4|ADGRG6|GRHL2|ALPK2|MESP2|ARL13B|MIR21", "Description": "heart morphogenesis", "_MEMBER_Input_ID": 1.0, "_GeneInHitList": 1562.0, "Category": "GO Biological Processes", "Log_q_value_": -2.2861831011788287, "Symbol": "GO:0003007", "LogP": -4.174998528148335, "_RANK_": 1.0, "FirstInGroupByEnrichment": 0.0, "_TotalGeneInLibrary": 24367.0, "_GeneInGO": 261.0, "shared_name": "85", "_InGO": 2.176696542893726, "Gene": "GO:0003007", "_COUNT_001": 34.0, "SUID": 579282, "id_original": "85", "selected": false, "CategoryID": 19.0, "STDV_InGO": 0.3692152164903899, "EvidenceCutoff": 0.0, "PARENT_GO": "19_GO:0032502 developmental process", "FirstInGroupByLogP": 0.0, "GROUP_ID": 11.0, "name": "85", "BestEnrichmentInGroup": 2.5527063205680363, "CLUSTER_LABEL": " ", "GeneID": "2050|2274|2294|2475|3398|3611|3670|4088|4092|4851|5727|5914|5915|6256|6491|6604|6659|6722|6926|8642|9249|10269|10370|23493|55036|55742|55897|57142|57211|79977|115701|145873|200894|406991", "_LogP_Input_ID": -4.174998528148335, "Z_score": 4.38758294784934, "GiniIndex": 0.0, "_COUNT_001_PCT": 100.0}, "position": {"x": 880.4598386162465, "y": -70.5856213027497}, "selected": false}, {"data": {"id": "579281", "Enrichment": 1.799985226041564, "BestLogPInGroup": -9.059097575411174, "_PATTERN_": "M1", "_GeneInGOAndHitList": 45.0, "Hits": "DDX5|ERBB3|EYA2|FHL2|FLNB|FOS|MTOR|GPC1|HNRNPU|HOXD10|ID2|ISL1|LMNA|SMAD3|SMAD7|MEF2D|MEIS1|NOTCH1|PPP3CA|PRKAR1A|RARA|RARB|RB1|RXRA|SRF|TBX3|BTG2|SORBS2|YBX3|CFLAR|FLOT1|ZMPSTE24|DDX17|PDLIM5|KDM6B|HEY2|MYLPF|MESP1|ADAMTS9|NIBAN2|UQCC2|ALPK2|SIK1|BCL9L|MIR25", "Description": "striated muscle tissue development", "_MEMBER_Input_ID": 1.0, "_GeneInHitList": 1562.0, "Category": "GO Biological Processes", "Log_q_value_": -2.165046738457108, "Symbol": "GO:0014706", "LogP": -4.001279214158131, "_RANK_": 1.0, "FirstInGroupByEnrichment": 0.0, "_TotalGeneInLibrary": 24367.0, "_GeneInGO": 390.0, "shared_name": "84", "_InGO": 2.880921895006402, "Gene": "GO:0014706", "_COUNT_001": 45.0, "SUID": 579281, "id_original": "84", "selected": false, "CategoryID": 19.0, "STDV_InGO": 0.423231031594662, "EvidenceCutoff": 0.0, "PARENT_GO": "19_GO:0032502 developmental process", "FirstInGroupByLogP": 0.0, "GROUP_ID": 11.0, "name": "84", "BestEnrichmentInGroup": 2.5527063205680363, "CLUSTER_LABEL": " ", "GeneID": "1655|2065|2139|2274|2317|2353|2475|2817|3192|3236|3398|3670|4000|4088|4092|4209|4211|4851|5530|5573|5914|5915|5925|6256|6722|6926|7832|8470|8531|8837|10211|10269|10521|10611|23135|23493|29895|55897|56999|64855|84300|115701|150094|283149|407014", "_LogP_Input_ID": -4.001279214158131, "Z_score": 4.168064633723845, "GiniIndex": 0.0, "_COUNT_001_PCT": 100.0}, "position": {"x": 808.4472348564809, "y": -139.2421383315583}, "selected": false}, {"data": {"id": "579280", "Enrichment": 2.3816598410728074, "BestLogPInGroup": -9.059097575411174, "_PATTERN_": "M1", "_GeneInGOAndHitList": 20.0, "Hits": "FHL2|FOXF1|ID2|ISL1|SMAD7|NOTCH1|RARA|RARB|RXRA|SMARCD3|SOX4|SRF|TBX3|DHRS3|CITED2|HEY2|PARVA|MESP1|ADGRG6|GRHL2", "Description": "cardiac chamber morphogenesis", "_MEMBER_Input_ID": 1.0, "_GeneInHitList": 1562.0, "Category": "GO Biological Processes", "Log_q_value_": -1.8665932225946078, "Symbol": "GO:0003206", "LogP": -3.5913067315289573, "_RANK_": 1.0, "FirstInGroupByEnrichment": 0.0, "_TotalGeneInLibrary": 24367.0, "_GeneInGO": 131.0, "shared_name": "83", "_InGO": 1.2804097311139564, "Gene": "GO:0003206", "_COUNT_001": 20.0, "SUID": 579280, "id_original": "83", "selected": false, "CategoryID": 19.0, "STDV_InGO": 0.284469454773297, "EvidenceCutoff": 0.0, "PARENT_GO": "19_GO:0032502 developmental process", "FirstInGroupByLogP": 0.0, "GROUP_ID": 11.0, "name": "83", "BestEnrichmentInGroup": 2.5527063205680363, "CLUSTER_LABEL": " ", "GeneID": "2274|2294|3398|3670|4092|4851|5914|5915|6256|6604|6659|6722|6926|9249|10370|23493|55742|55897|57211|79977", "_LogP_Input_ID": -3.5913067315289573, "Z_score": 4.149768856418864, "GiniIndex": 0.0, "_COUNT_001_PCT": 100.0}, "position": {"x": 733.8901059502309, "y": -104.04777187648017}, "selected": false}, {"data": {"id": "579279", "Enrichment": 2.073971416518026, "BestLogPInGroup": -9.059097575411174, "_PATTERN_": "M1", "_GeneInGOAndHitList": 23.0, "Hits": "FHL2|FOXF1|ID2|ISL1|SMAD7|NOTCH1|PTK7|RARA|RARB|RXRA|SMARCD3|SOX4|SRF|TBX3|DHRS3|SNX17|ZMPSTE24|CITED2|HEY2|PARVA|MESP1|ADGRG6|GRHL2", "Description": "cardiac chamber development", "_MEMBER_Input_ID": 1.0, "_GeneInHitList": 1562.0, "Category": "GO Biological Processes", "Log_q_value_": -1.5483694780681447, "Symbol": "GO:0003205", "LogP": -3.147316233971964, "_RANK_": 1.0, "FirstInGroupByEnrichment": 0.0, "_TotalGeneInLibrary": 24367.0, "_GeneInGO": 173.0, "shared_name": "82", "_InGO": 1.47247119078105, "Gene": "GO:0003205", "_COUNT_001": 23.0, "SUID": 579279, "id_original": "82", "selected": false, "CategoryID": 19.0, "STDV_InGO": 0.30476260956438245, "EvidenceCutoff": 0.0, "PARENT_GO": "19_GO:0032502 developmental process", "FirstInGroupByLogP": 0.0, "GROUP_ID": 11.0, "name": "82", "BestEnrichmentInGroup": 2.5527063205680363, "CLUSTER_LABEL": " ", "GeneID": "2274|2294|3398|3670|4092|4851|5754|5914|5915|6256|6604|6659|6722|6926|9249|9784|10269|10370|23493|55742|55897|57211|79977", "_LogP_Input_ID": -3.147316233971964, "Z_score": 3.7100480360681645, "GiniIndex": 0.0, "_COUNT_001_PCT": 100.0}, "position": {"x": 856.648925530309, "y": -173.48244137110908}, "selected": false}, {"data": {"id": "579278", "Enrichment": 2.0222056243183006, "BestLogPInGroup": -9.059097575411174, "_PATTERN_": "M1", "_GeneInGOAndHitList": 21.0, "Hits": "EGFR|FHL2|MTOR|HNRNPU|ISL1|LMNA|NOTCH1|RARA|RARB|RXRA|SRF|TBX3|SORBS2|ZMPSTE24|PDLIM5|KDM6B|HEY2|MESP1|ALPK2|SIK1|MIR21", "Description": "cardiocyte differentiation", "_MEMBER_Input_ID": 1.0, "_GeneInHitList": 1562.0, "Category": "GO Biological Processes", "Log_q_value_": -1.2907261509923595, "Symbol": "GO:0035051", "LogP": -2.785868388512291, "_RANK_": 1.0, "FirstInGroupByEnrichment": 0.0, "_TotalGeneInLibrary": 24367.0, "_GeneInGO": 162.0, "shared_name": "81", "_InGO": 1.3444302176696543, "Gene": "GO:0035051", "_COUNT_001": 21.0, "SUID": 579278, "id_original": "81", "selected": false, "CategoryID": 19.0, "STDV_InGO": 0.29139991611511556, "EvidenceCutoff": 0.0, "PARENT_GO": "19_GO:0032502 developmental process", "FirstInGroupByLogP": 0.0, "GROUP_ID": 11.0, "name": "81", "BestEnrichmentInGroup": 2.5527063205680363, "CLUSTER_LABEL": " ", "GeneID": "1956|2274|2475|3192|3670|4000|4851|5914|5915|6256|6722|6926|8470|10269|10611|23135|23493|55897|115701|150094|406991", "_LogP_Input_ID": -2.785868388512291, "Z_score": 3.4163379585021056, "GiniIndex": 0.0, "_COUNT_001_PCT": 100.0}, "position": {"x": 765.6105649346059, "y": -175.49507564845283}, "selected": false}, {"data": {"id": "579277", "Enrichment": 1.8777623654384215, "BestLogPInGroup": -8.650869564251822, "_PATTERN_": "M1", "_GeneInGOAndHitList": 91.0, "Hits": "ABL1|ACTB|ACTN4|ACTN1|ALCAM|ANK3|RERE|BMPR1B|CFL1|CLDN3|CUX1|EPHA2|ERBB2|FBLN1|FAT1|FLNB|GPC1|GRIN1|HSP90AB1|ID1|ID2|ILK|ISL1|ITGB7|KIF5C|LIMS1|LRP1|MYH9|NEDD9|NOTCH1|NOTCH4|NR4A2|P4HB|PIK3CB|PPP3CA|PTCH1|PTPN6|PTPRM|NECTIN1|RAC3|RAP1GAP|RB1|RREB1|S100A6|SCN1B|SHC1|SRF|STK11|TBCD|TRIO|EZR|KLF7|IRS2|SYNGAP1|ZEB2|FARP1|FLOT1|BAIAP2|PDLIM5|TRIOBP|SHANK2|PALLD|MYCBP2|SPART|KANK1|COBL|CUX2|NEDD4L|MACF1|CORO1C|ARHGEF26|ANAPC2|TNFRSF12A|SEMA4C|ENAH|PARVA|RTN4|SEMA6A|NIBAN2|VWA1|MICALL2|FBXO31|PDZD7|GRHL2|COL18A1|UNK|PLXNA4|C12orf57|ARL13B|TPRN|CCDC88C", "Description": "cell morphogenesis involved in differentiation", "_MEMBER_Input_ID": 1.0, "_GeneInHitList": 1562.0, "Category": "GO Biological Processes", "Log_q_value_": -5.393878257702022, "Symbol": "GO:0000904", "LogP": -8.332773075300192, "_RANK_": 1.0, "FirstInGroupByEnrichment": 0.0, "_TotalGeneInLibrary": 24367.0, "_GeneInGO": 756.0, "shared_name": "80", "_InGO": 5.825864276568502, "Gene": "GO:0000904", "_COUNT_001": 91.0, "SUID": 579277, "id_original": "80", "selected": false, "CategoryID": 19.0, "STDV_InGO": 0.5926598246801607, "EvidenceCutoff": 0.0, "PARENT_GO": "19_GO:0032502 developmental process", "FirstInGroupByLogP": 0.0, "GROUP_ID": 12.0, "name": "80", "BestEnrichmentInGroup": 2.1347193207089425, "CLUSTER_LABEL": " ", "GeneID": "25|60|81|87|214|288|473|658|1072|1365|1523|1969|2064|2192|2195|2317|2817|2902|3326|3397|3398|3611|3670|3695|3800|3987|4035|4627|4739|4851|4855|4929|5034|5291|5530|5727|5777|5797|5818|5881|5909|5925|6239|6277|6324|6464|6722|6794|6904|7204|7430|8609|8660|8831|9839|10160|10211|10458|10611|11078|22941|23022|23077|23111|23189|23242|23316|23327|23499|23603|26084|29882|51330|54910|55740|55742|57142|57556|64855|64856|79778|79791|79955|79977|80781|85451|91584|113246|200894|286262|440193", "_LogP_Input_ID": -8.332773075300192, "Z_score": 6.416500904946731, "GiniIndex": 0.0, "_COUNT_001_PCT": 100.0}, "position": {"x": -487.60092188180033, "y": 506.92721133885186}, "selected": false}, {"data": {"id": "579276", "Enrichment": 1.9259101183983813, "BestLogPInGroup": -8.650869564251822, "_PATTERN_": "M1", "_GeneInGOAndHitList": 70.0, "Hits": "ABL1|BCL6|CSNK1D|CUX1|DLX1|DLX2|DVL3|EIF4G1|MTOR|GATA2|RAPGEF1|FOXA1|HOXA11|ID2|ILK|ITGA3|LIMS1|LRP1|SMAD7|NBL1|NEDD9|NFE2L2|NOTCH1|ROR2|P4HB|SERPINF1|PRKCH|PTK7|RAC3|RARA|RARB|REG1A|RREB1|SCN1B|CX3CL1|SMARCD3|SRF|STK11|SOCS2|CFLAR|SPAG9|CBFA2T2|ARHGEF2|DLG5|ZEB2|TRIB1|BAIAP2|TRIOBP|CPEB3|SHANK2|COBL|CUX2|NEDD4L|MACF1|ZMYND8|ANAPC2|PACSIN1|TNFRSF12A|SRRT|STYXL1|SEMA6A|NIBAN2|FBXO31|PLXNA4|RNF157|DAB2IP|BRINP3|MIR125B1|MIR200C|FOXO6", "Description": "positive regulation of cell development", "_MEMBER_Input_ID": 1.0, "_GeneInHitList": 1562.0, "Category": "GO Biological Processes", "Log_q_value_": -4.332987204708262, "Symbol": "GO:0010720", "LogP": -6.970852026642451, "_RANK_": 1.0, "FirstInGroupByEnrichment": 0.0, "_TotalGeneInLibrary": 24367.0, "_GeneInGO": 567.0, "shared_name": "79", "_InGO": 4.481434058898848, "Gene": "GO:0010720", "_COUNT_001": 70.0, "SUID": 579276, "id_original": "79", "selected": false, "CategoryID": 19.0, "STDV_InGO": 0.5234942156012867, "EvidenceCutoff": 0.0, "PARENT_GO": "19_GO:0048518 positive regulation of biological process", "FirstInGroupByLogP": 0.0, "GROUP_ID": 12.0, "name": "79", "BestEnrichmentInGroup": 2.1347193207089425, "CLUSTER_LABEL": " ", "GeneID": "25|604|1453|1523|1745|1746|1857|1981|2475|2624|2889|3169|3207|3398|3611|3675|3987|4035|4092|4681|4739|4780|4851|4920|5034|5176|5583|5754|5881|5914|5915|5967|6239|6324|6376|6604|6722|6794|8835|8837|9043|9139|9181|9231|9839|10221|10458|11078|22849|22941|23242|23316|23327|23499|23613|29882|29993|51330|51593|51657|57556|64855|79791|91584|114804|153090|339479|406911|406985|100132074", "_LogP_Input_ID": -6.970852026642451, "Z_score": 5.838341271867907, "GiniIndex": 0.0, "_COUNT_001_PCT": 100.0}, "position": {"x": -394.5131228095347, "y": 616.9538837021331}, "selected": false}, {"data": {"id": "579275", "Enrichment": 2.1347193207089425, "BestLogPInGroup": -8.650869564251822, "_PATTERN_": "M1", "_GeneInGOAndHitList": 52.0, "Hits": "BCL6|CSNK1D|CUX1|DLX1|DLX2|DVL3|EIF4G1|MTOR|GATA2|RAPGEF1|FOXA1|ILK|ITGA3|LRP1|NBL1|NFE2L2|ROR2|SERPINF1|PTK7|RARA|RARB|REG1A|SCN1B|CX3CL1|SRF|STK11|SOCS2|CFLAR|SPAG9|CBFA2T2|ARHGEF2|DLG5|ZEB2|BAIAP2|CPEB3|SHANK2|COBL|CUX2|NEDD4L|MACF1|ZMYND8|ANAPC2|PACSIN1|TNFRSF12A|STYXL1|FBXO31|PLXNA4|RNF157|DAB2IP|BRINP3|MIR200C|FOXO6", "Description": "positive regulation of neuron differentiation", "_MEMBER_Input_ID": 1.0, "_GeneInHitList": 1562.0, "Category": "GO Biological Processes", "Log_q_value_": -4.1354428848374765, "Symbol": "GO:0045666", "LogP": -6.706360917141052, "_RANK_": 1.0, "FirstInGroupByEnrichment": 0.0, "_TotalGeneInLibrary": 24367.0, "_GeneInGO": 380.0, "shared_name": "78", "_InGO": 3.329065300896287, "Gene": "GO:0045666", "_COUNT_001": 52.0, "SUID": 579275, "id_original": "78", "selected": false, "CategoryID": 19.0, "STDV_InGO": 0.45390879852988153, "EvidenceCutoff": 0.0, "PARENT_GO": "19_GO:0048518 positive regulation of biological process", "FirstInGroupByLogP": 0.0, "GROUP_ID": 12.0, "name": "78", "BestEnrichmentInGroup": 2.1347193207089425, "CLUSTER_LABEL": " ", "GeneID": "604|1453|1523|1745|1746|1857|1981|2475|2624|2889|3169|3611|3675|4035|4681|4780|4920|5176|5754|5914|5915|5967|6324|6376|6722|6794|8835|8837|9043|9139|9181|9231|9839|10458|22849|22941|23242|23316|23327|23499|23613|29882|29993|51330|51657|79791|91584|114804|153090|339479|406985|100132074", "_LogP_Input_ID": -6.706360917141052, "Z_score": 5.834583731785901, "GiniIndex": 0.0, "_COUNT_001_PCT": 100.0}, "position": {"x": -288.4714663153941, "y": 539.2592425888519}, "selected": false}, {"data": {"id": "579274", "Enrichment": 1.9228470406752827, "BestLogPInGroup": -8.650869564251822, "_PATTERN_": "M1", "_GeneInGOAndHitList": 62.0, "Hits": "ABL1|ACTN4|RND3|CAPZB|CFL1|CLDN4|CLDN3|CUX1|DVL3|FBLN1|FGR|GRIN1|HPN|ID1|ILK|ITGA7|LIMS1|LRP1|MYH9|NEDD9|P4HB|PPP3CA|RAC3|REG1A|RREB1|S100A13|SPARC|SRF|STK11|EZR|SYNGAP1|ZEB2|ZMPSTE24|BAIAP2|PDLIM5|GNA13|TRIOBP|MYCBP2|SPART|KANK1|DNMBP|CUX2|NEDD4L|RHOQ|MACF1|CORO1C|ANAPC2|PLEKHO1|TNFRSF12A|RHOF|SEMA4C|PARVA|RTN4|SEMA6A|SHROOM3|RHOU|FBXO31|PLXNA4|RNF157|FGD2|BCL9L|MIR21", "Description": "regulation of cell morphogenesis", "_MEMBER_Input_ID": 1.0, "_GeneInHitList": 1562.0, "Category": "GO Biological Processes", "Log_q_value_": -3.7590423347686484, "Symbol": "GO:0022604", "LogP": -6.223370200603977, "_RANK_": 1.0, "FirstInGroupByEnrichment": 0.0, "_TotalGeneInLibrary": 24367.0, "_GeneInGO": 503.0, "shared_name": "77", "_InGO": 3.9692701664532652, "Gene": "GO:0022604", "_COUNT_001": 62.0, "SUID": 579274, "id_original": "77", "selected": false, "CategoryID": 19.0, "STDV_InGO": 0.49399201616029104, "EvidenceCutoff": 0.0, "PARENT_GO": "19_GO:0032502 developmental process", "FirstInGroupByLogP": 0.0, "GROUP_ID": 12.0, "name": "77", "BestEnrichmentInGroup": 2.1347193207089425, "CLUSTER_LABEL": " ", "GeneID": "25|81|390|832|1072|1364|1365|1523|1857|2192|2268|2902|3249|3397|3611|3679|3987|4035|4627|4739|5034|5530|5881|5967|6239|6284|6678|6722|6794|7430|8831|9839|10269|10458|10611|10672|11078|23077|23111|23189|23268|23316|23327|23433|23499|23603|29882|51177|51330|54509|54910|55742|57142|57556|57619|58480|79791|91584|114804|221472|283149|406991", "_LogP_Input_ID": -6.223370200603977, "Z_score": 5.473431568085911, "GiniIndex": 0.0, "_COUNT_001_PCT": 100.0}, "position": {"x": -483.3367312079722, "y": 568.8312640732269}, "selected": false}, {"data": {"id": "579273", "Enrichment": 1.9016372842615425, "BestLogPInGroup": -8.650869564251822, "_PATTERN_": "M1", "_GeneInGOAndHitList": 59.0, "Hits": "BCL6|CSNK1D|CUX1|DLX1|DLX2|DVL3|EIF4G1|MTOR|GATA2|RAPGEF1|FOXA1|ID2|ILK|ITGA3|LRP1|NBL1|NFE2L2|NOTCH1|ROR2|SERPINF1|PRKCH|PTK7|RARA|RARB|REG1A|SCN1B|CX3CL1|SMARCD3|SRF|STK11|SOCS2|CFLAR|SPAG9|CBFA2T2|ARHGEF2|DLG5|ZEB2|BAIAP2|CPEB3|SHANK2|COBL|CUX2|NEDD4L|MACF1|ZMYND8|ANAPC2|PACSIN1|TNFRSF12A|SRRT|STYXL1|SEMA6A|FBXO31|PLXNA4|RNF157|DAB2IP|BRINP3|MIR125B1|MIR200C|FOXO6", "Description": "positive regulation of neurogenesis", "_MEMBER_Input_ID": 1.0, "_GeneInHitList": 1562.0, "Category": "GO Biological Processes", "Log_q_value_": -3.459826882526509, "Symbol": "GO:0050769", "LogP": -5.796661708796717, "_RANK_": 1.0, "FirstInGroupByEnrichment": 0.0, "_TotalGeneInLibrary": 24367.0, "_GeneInGO": 484.0, "shared_name": "76", "_InGO": 3.7772087067861717, "Gene": "GO:0050769", "_COUNT_001": 59.0, "SUID": 579273, "id_original": "76", "selected": false, "CategoryID": 19.0, "STDV_InGO": 0.48237406725279086, "EvidenceCutoff": 0.0, "PARENT_GO": "19_GO:0048518 positive regulation of biological process", "FirstInGroupByLogP": 0.0, "GROUP_ID": 12.0, "name": "76", "BestEnrichmentInGroup": 2.1347193207089425, "CLUSTER_LABEL": " ", "GeneID": "604|1453|1523|1745|1746|1857|1981|2475|2624|2889|3169|3398|3611|3675|4035|4681|4780|4851|4920|5176|5583|5754|5914|5915|5967|6324|6376|6604|6722|6794|8835|8837|9043|9139|9181|9231|9839|10458|22849|22941|23242|23316|23327|23499|23613|29882|29993|51330|51593|51657|57556|79791|91584|114804|153090|339479|406911|406985|100132074", "_LogP_Input_ID": -5.796661708796717, "Z_score": 5.243577607521376, "GiniIndex": 0.0, "_COUNT_001_PCT": 100.0}, "position": {"x": -316.74450708687846, "y": 608.12157779393}, "selected": false}, {"data": {"id": "579272", "Enrichment": 1.7307550250399653, "BestLogPInGroup": -8.650869564251822, "_PATTERN_": "M1", "_GeneInGOAndHitList": 75.0, "Hits": "ABL1|ACTB|ALCAM|ANK3|RERE|BMPR1B|CFL1|CUX1|DVL3|EGFR|EPHA2|ERBB2|GPC1|GPM6A|GRIN1|MNX1|HSP90AB1|ID1|ILK|ISL1|KIF5C|LRP1|NBL1|NEK3|NOTCH1|NR4A2|PIK3CB|PPP3CA|PTCH1|PTPRM|NECTIN1|RAP1GAP|REG1A|RREB1|S100A6|SCN1B|SHC1|SRF|STK11|TRIO|EZR|CNTNAP1|KLF7|IRS2|SYNGAP1|ZEB2|FARP1|FLOT1|BAIAP2|PDLIM5|PACSIN2|SHANK2|PALLD|MYCBP2|SPART|KANK1|COBL|CUX2|NEDD4L|MACF1|CORO1C|ANAPC2|PACSIN1|PLEKHO1|TNFRSF12A|SEMA4C|ENAH|RTN4|SEMA6A|NIBAN2|FBXO31|PLXNA4|C12orf57|RNF157|DAB2IP", "Description": "cell projection morphogenesis", "_MEMBER_Input_ID": 1.0, "_GeneInHitList": 1562.0, "Category": "GO Biological Processes", "Log_q_value_": -3.322656432492372, "Symbol": "GO:0048858", "LogP": -5.600776845543783, "_RANK_": 1.0, "FirstInGroupByEnrichment": 0.0, "_TotalGeneInLibrary": 24367.0, "_GeneInGO": 676.0, "shared_name": "75", "_InGO": 4.801536491677337, "Gene": "GO:0048858", "_COUNT_001": 75.0, "SUID": 579272, "id_original": "75", "selected": false, "CategoryID": 19.0, "STDV_InGO": 0.5409592753355752, "EvidenceCutoff": 0.0, "PARENT_GO": "19_GO:0032502 developmental process", "FirstInGroupByLogP": 0.0, "GROUP_ID": 12.0, "name": "75", "BestEnrichmentInGroup": 2.1347193207089425, "CLUSTER_LABEL": " ", "GeneID": "25|60|214|288|473|658|1072|1523|1857|1956|1969|2064|2817|2823|2902|3110|3326|3397|3611|3670|3800|4035|4681|4752|4851|4929|5291|5530|5727|5797|5818|5909|5967|6239|6277|6324|6464|6722|6794|7204|7430|8506|8609|8660|8831|9839|10160|10211|10458|10611|11252|22941|23022|23077|23111|23189|23242|23316|23327|23499|23603|29882|29993|51177|51330|54910|55740|57142|57556|64855|79791|91584|113246|114804|153090", "_LogP_Input_ID": -5.600776845543783, "Z_score": 5.042792038356827, "GiniIndex": 0.0, "_COUNT_001_PCT": 100.0}, "position": {"x": -416.8386538153941, "y": 567.4450030868987}, "selected": false}, {"data": {"id": "579271", "Enrichment": 1.7178430431071274, "BestLogPInGroup": -8.650869564251822, "_PATTERN_": "M1", "_GeneInGOAndHitList": 74.0, "Hits": "ABL1|ACTB|ALCAM|ANK3|RERE|BMPR1B|CFL1|CUX1|DVL3|EGFR|EPHA2|ERBB2|GPC1|GPM6A|GRIN1|MNX1|HSP90AB1|ID1|ILK|ISL1|KIF5C|LRP1|NBL1|NEK3|NOTCH1|NR4A2|PIK3CB|PPP3CA|PTCH1|PTPRM|NECTIN1|RAP1GAP|REG1A|RREB1|S100A6|SCN1B|SHC1|SRF|STK11|TRIO|EZR|CNTNAP1|KLF7|IRS2|SYNGAP1|ZEB2|FARP1|FLOT1|BAIAP2|PDLIM5|SHANK2|PALLD|MYCBP2|SPART|KANK1|COBL|CUX2|NEDD4L|MACF1|CORO1C|ANAPC2|PACSIN1|PLEKHO1|TNFRSF12A|SEMA4C|ENAH|RTN4|SEMA6A|NIBAN2|FBXO31|PLXNA4|C12orf57|RNF157|DAB2IP", "Description": "plasma membrane bounded cell projection morphogenesis", "_MEMBER_Input_ID": 1.0, "_GeneInHitList": 1562.0, "Category": "GO Biological Processes", "Log_q_value_": -3.2014082371305945, "Symbol": "GO:0120039", "LogP": -5.416710575484414, "_RANK_": 1.0, "FirstInGroupByEnrichment": 0.0, "_TotalGeneInLibrary": 24367.0, "_GeneInGO": 672.0, "shared_name": "74", "_InGO": 4.7375160051216385, "Gene": "GO:0120039", "_COUNT_001": 74.0, "SUID": 579271, "id_original": "74", "selected": false, "CategoryID": 19.0, "STDV_InGO": 0.5375214271483969, "EvidenceCutoff": 0.0, "PARENT_GO": "19_GO:0032502 developmental process", "FirstInGroupByLogP": 0.0, "GROUP_ID": 12.0, "name": "74", "BestEnrichmentInGroup": 2.1347193207089425, "CLUSTER_LABEL": " ", "GeneID": "25|60|214|288|473|658|1072|1523|1857|1956|1969|2064|2817|2823|2902|3110|3326|3397|3611|3670|3800|4035|4681|4752|4851|4929|5291|5530|5727|5797|5818|5909|5967|6239|6277|6324|6464|6722|6794|7204|7430|8506|8609|8660|8831|9839|10160|10211|10458|10611|22941|23022|23077|23111|23189|23242|23316|23327|23499|23603|29882|29993|51177|51330|54910|55740|57142|57556|64855|79791|91584|113246|114804|153090", "_LogP_Input_ID": -5.416710575484414, "Z_score": 4.9385945437114485, "GiniIndex": 0.0, "_COUNT_001_PCT": 100.0}, "position": {"x": -408.95224024117533, "y": 506.201411778305}, "selected": false}, {"data": {"id": "579270", "Enrichment": 1.7034342943764993, "BestLogPInGroup": -8.650869564251822, "_PATTERN_": "M1", "_GeneInGOAndHitList": 76.0, "Hits": "ABL1|ACTB|ALCAM|ANK3|RERE|BCL2L1|BMPR1B|CFL1|CUX1|DVL3|EGFR|EPHA2|ERBB2|GPC1|GPM6A|GRIN1|MNX1|HSP90AB1|ID1|ILK|ISL1|KIF5C|LRP1|NBL1|NEK3|NOTCH1|NR4A2|PIK3CB|PPP3CA|PTCH1|PTPRM|NECTIN1|RAP1GAP|REG1A|RREB1|S100A6|SCN1B|SHC1|SRF|STK11|TRIO|EZR|CNTNAP1|KLF7|IRS2|SYNGAP1|ZEB2|FARP1|FLOT1|BAIAP2|PDLIM5|PACSIN2|SHANK2|PALLD|MYCBP2|SPART|KANK1|COBL|CUX2|NEDD4L|MACF1|CORO1C|ANAPC2|PACSIN1|PLEKHO1|TNFRSF12A|SEMA4C|ENAH|RTN4|SEMA6A|NIBAN2|FBXO31|PLXNA4|C12orf57|RNF157|DAB2IP", "Description": "cell part morphogenesis", "_MEMBER_Input_ID": 1.0, "_GeneInHitList": 1562.0, "Category": "GO Biological Processes", "Log_q_value_": -3.2000512336719136, "Symbol": "GO:0032990", "LogP": -5.411947323333822, "_RANK_": 1.0, "FirstInGroupByEnrichment": 0.0, "_TotalGeneInLibrary": 24367.0, "_GeneInGO": 696.0, "shared_name": "73", "_InGO": 4.865556978233035, "Gene": "GO:0032990", "_COUNT_001": 76.0, "SUID": 579270, "id_original": "73", "selected": false, "CategoryID": 19.0, "STDV_InGO": 0.5443705930550844, "EvidenceCutoff": 0.0, "PARENT_GO": "19_GO:0032502 developmental process", "FirstInGroupByLogP": 0.0, "GROUP_ID": 12.0, "name": "73", "BestEnrichmentInGroup": 2.1347193207089425, "CLUSTER_LABEL": " ", "GeneID": "25|60|214|288|473|598|658|1072|1523|1857|1956|1969|2064|2817|2823|2902|3110|3326|3397|3611|3670|3800|4035|4681|4752|4851|4929|5291|5530|5727|5797|5818|5909|5967|6239|6277|6324|6464|6722|6794|7204|7430|8506|8609|8660|8831|9839|10160|10211|10458|10611|11252|22941|23022|23077|23111|23189|23242|23316|23327|23499|23603|29882|29993|51177|51330|54910|55740|57142|57556|64855|79791|91584|113246|114804|153090", "_LogP_Input_ID": -5.411947323333822, "Z_score": 4.927622743771421, "GiniIndex": 0.0, "_COUNT_001_PCT": 100.0}, "position": {"x": -363.20129419625346, "y": 533.6563067978362}, "selected": false}, {"data": {"id": "579269", "Enrichment": 1.7217249988223657, "BestLogPInGroup": -8.650869564251822, "_PATTERN_": "M1", "_GeneInGOAndHitList": 66.0, "Hits": "ABL1|ACTB|ALCAM|ANK3|RERE|BMPR1B|CFL1|CUX1|EPHA2|ERBB2|GPC1|GRIN1|HSP90AB1|ID1|ID2|ILK|ISL1|KIF5C|LRP1|NOTCH1|NR4A2|PIK3CB|PPP3CA|PTCH1|PTPRM|NECTIN1|RAP1GAP|RB1|S100A6|SCN1B|SHC1|SRF|STK11|TBCD|TRIO|EZR|KLF7|IRS2|SYNGAP1|ZEB2|FARP1|FLOT1|BAIAP2|PDLIM5|TRIOBP|SHANK2|PALLD|MYCBP2|SPART|COBL|CUX2|NEDD4L|MACF1|ANAPC2|TNFRSF12A|SEMA4C|ENAH|RTN4|SEMA6A|NIBAN2|FBXO31|PDZD7|UNK|PLXNA4|C12orf57|TPRN", "Description": "cell morphogenesis involved in neuron differentiation", "_MEMBER_Input_ID": 1.0, "_GeneInHitList": 1562.0, "Category": "GO Biological Processes", "Log_q_value_": -2.853399113573478, "Symbol": "GO:0048667", "LogP": -4.930200777987243, "_RANK_": 1.0, "FirstInGroupByEnrichment": 0.0, "_TotalGeneInLibrary": 24367.0, "_GeneInGO": 598.0, "shared_name": "72", "_InGO": 4.225352112676056, "Gene": "GO:0048667", "_COUNT_001": 66.0, "SUID": 579269, "id_original": "72", "selected": false, "CategoryID": 19.0, "STDV_InGO": 0.5089981691815402, "EvidenceCutoff": 0.0, "PARENT_GO": "19_GO:0032502 developmental process", "FirstInGroupByLogP": 0.0, "GROUP_ID": 12.0, "name": "72", "BestEnrichmentInGroup": 2.1347193207089425, "CLUSTER_LABEL": " ", "GeneID": "25|60|214|288|473|658|1072|1523|1969|2064|2817|2902|3326|3397|3398|3611|3670|3800|4035|4851|4929|5291|5530|5727|5797|5818|5909|5925|6277|6324|6464|6722|6794|6904|7204|7430|8609|8660|8831|9839|10160|10211|10458|10611|11078|22941|23022|23077|23111|23242|23316|23327|23499|29882|51330|54910|55740|57142|57556|64855|79791|79955|85451|91584|113246|286262", "_LogP_Input_ID": -4.930200777987243, "Z_score": 4.676645832599008, "GiniIndex": 0.0, "_COUNT_001_PCT": 100.0}, "position": {"x": -455.81774927437846, "y": 442.9584155624847}, "selected": false}, {"data": {"id": "579268", "Enrichment": 2.213495345537599, "BestLogPInGroup": -8.621070944395578, "_PATTERN_": "M1", "_GeneInGOAndHitList": 63.0, "Hits": "AKT1|ATF4|ZFP36L1|ZFP36L2|CIRBP|DAPK1|DHFR|EIF4G1|ERBB2|FOXO3|MTOR|MKNK2|HSPB1|NOTCH4|PA2G4|PER1|RARA|UPF1|RXRA|SOX4|TARBP2|UCN|ZFP36|BTG2|YBX3|EIF3H|PER2|EIF2B4|EIF2B5|QKI|TOB1|CASC3|CPEB3|SAMD4A|PASK|PPP1R15A|EIF2AK1|EIF3K|COA3|DDX25|BANK1|SAMD4B|CNOT11|EEFSEC|METTL16|ZC3H12A|UQCC2|PYM1|UNK|DPH6|PATL2|CNOT6L|NANOS3|MIR106B|MIR125B1|MIR132|MIR148A|MIR200C|MIR21|MIR212|MIR27B|MIR29A|MIR29B1", "Description": "regulation of translation", "_MEMBER_Input_ID": 1.0, "_GeneInHitList": 1562.0, "Category": "GO Biological Processes", "Log_q_value_": -5.585758129620247, "Symbol": "GO:0006417", "LogP": -8.603834193266042, "_RANK_": 1.0, "FirstInGroupByEnrichment": 0.0, "_TotalGeneInLibrary": 24367.0, "_GeneInGO": 444.0, "shared_name": "71", "_InGO": 4.0332906530089625, "Gene": "GO:0006417", "_COUNT_001": 63.0, "SUID": 579268, "id_original": "71", "selected": false, "CategoryID": 19.0, "STDV_InGO": 0.4977938729581788, "EvidenceCutoff": 0.0, "PARENT_GO": "19_GO:0008152 metabolic process", "FirstInGroupByLogP": 0.0, "GROUP_ID": 13.0, "name": "71", "BestEnrichmentInGroup": 2.455535400957936, "CLUSTER_LABEL": " ", "GeneID": "207|468|677|678|1153|1612|1719|1981|2064|2309|2475|2872|3315|4855|5036|5187|5914|5976|6256|6659|6895|7349|7538|7832|8531|8667|8864|8890|8893|9444|10140|22794|22849|23034|23178|23645|27102|27335|28958|29118|55024|55095|55571|60678|79066|80149|84300|84305|85451|89978|197135|246175|342977|406900|406911|406921|406940|406985|406991|406994|407019|407021|407024", "_LogP_Input_ID": -8.603834193266042, "Z_score": 6.7536628474871465, "GiniIndex": 0.0, "_COUNT_001_PCT": 100.0}, "position": {"x": -85.15127636254374, "y": 790.9972491806487}, "selected": false}, {"data": {"id": "579267", "Enrichment": 2.455535400957936, "BestLogPInGroup": -8.621070944395578, "_PATTERN_": "M1", "_GeneInGOAndHitList": 34.0, "Hits": "ATF4|ZFP36L1|ZFP36L2|CIRBP|DAPK1|DHFR|EIF4G1|NOTCH4|RARA|UPF1|ZFP36|BTG2|TOB1|CPEB3|SAMD4A|EIF2AK1|BANK1|SAMD4B|METTL16|ZC3H12A|UNK|PATL2|CNOT6L|NANOS3|MIR106B|MIR125B1|MIR132|MIR148A|MIR200C|MIR21|MIR212|MIR27B|MIR29A|MIR29B1", "Description": "negative regulation of translation", "_MEMBER_Input_ID": 1.0, "_GeneInHitList": 1562.0, "Category": "GO Biological Processes", "Log_q_value_": -3.566429835951556, "Symbol": "GO:0017148", "LogP": -5.96522769121098, "_RANK_": 1.0, "FirstInGroupByEnrichment": 0.0, "_TotalGeneInLibrary": 24367.0, "_GeneInGO": 216.0, "shared_name": "70", "_InGO": 2.176696542893726, "Gene": "GO:0017148", "_COUNT_001": 34.0, "SUID": 579267, "id_original": "70", "selected": false, "CategoryID": 19.0, "STDV_InGO": 0.3692152164903899, "EvidenceCutoff": 0.0, "PARENT_GO": "19_GO:0048519 negative regulation of biological process", "FirstInGroupByLogP": 0.0, "GROUP_ID": 13.0, "name": "70", "BestEnrichmentInGroup": 2.455535400957936, "CLUSTER_LABEL": " ", "GeneID": "468|677|678|1153|1612|1719|1981|4855|5914|5976|7538|7832|10140|22849|23034|27102|55024|55095|79066|80149|85451|197135|246175|342977|406900|406911|406921|406940|406985|406991|406994|407019|407021|407024", "_LogP_Input_ID": -5.96522769121098, "Z_score": 5.623409181903807, "GiniIndex": 0.0, "_COUNT_001_PCT": 100.0}, "position": {"x": -218.46656872094218, "y": 832.6443439072112}, "selected": false}, {"data": {"id": "579266", "Enrichment": 2.1402263358421036, "BestLogPInGroup": -8.621070944395578, "_PATTERN_": "M1", "_GeneInGOAndHitList": 45.0, "Hits": "AKT1|ZFP36L1|ZFP36L2|CIRBP|DDX5|DYRK1A|EIF4G1|MTOR|HNRNPA2B1|HNRNPU|HSPA1A|HSPB1|MBNL1|PSMB3|PSMB8|PSMB9|PSMD1|PSMD2|UPF1|SRSF3|UBC|YWHAZ|ZFP36|BTG2|ANP32A|YBX3|THRAP3|TOB1|MBNL2|DDX17|KHDRBS3|SRSF8|CPEB3|SAMD4A|JMJD6|DAZAP1|ZBTB7A|PAF1|SAMD4B|METTL16|ZC3H12A|SRRM4|CNOT6L|SLC39A5|NANOS3", "Description": "regulation of mRNA metabolic process", "_MEMBER_Input_ID": 1.0, "_GeneInHitList": 1562.0, "Category": "GO Biological Processes", "Log_q_value_": -3.5480980256750203, "Symbol": "GO:1903311", "LogP": -5.924435541500853, "_RANK_": 1.0, "FirstInGroupByEnrichment": 0.0, "_TotalGeneInLibrary": 24367.0, "_GeneInGO": 328.0, "shared_name": "69", "_InGO": 2.880921895006402, "Gene": "GO:1903311", "_COUNT_001": 45.0, "SUID": 579266, "id_original": "69", "selected": false, "CategoryID": 19.0, "STDV_InGO": 0.423231031594662, "EvidenceCutoff": 0.0, "PARENT_GO": "19_GO:0008152 metabolic process", "FirstInGroupByLogP": 0.0, "GROUP_ID": 13.0, "name": "69", "BestEnrichmentInGroup": 2.455535400957936, "CLUSTER_LABEL": " ", "GeneID": "207|677|678|1153|1655|1859|1981|2475|3181|3192|3303|3315|4154|5691|5696|5698|5707|5708|5976|6428|7316|7534|7538|7832|8125|8531|9967|10140|10150|10521|10656|10929|22849|23034|23210|26528|51341|54623|55095|79066|80149|84530|246175|283375|342977", "_LogP_Input_ID": -5.924435541500853, "Z_score": 5.441108513456418, "GiniIndex": 0.0, "_COUNT_001_PCT": 100.0}, "position": {"x": -64.61292911828104, "y": 862.9146991318206}, "selected": false}, {"data": {"id": "579265", "Enrichment": 1.7045503247069491, "BestLogPInGroup": -8.621070944395578, "_PATTERN_": "M1", "_GeneInGOAndHitList": 79.0, "Hits": "AKT1|ATF4|ZFP36L1|ZFP36L2|MRPL49|CIRBP|DAPK1|DHFR|EEF1A1|EGFR|EIF4G1|ERBB2|FOXO3|MTOR|MKNK2|HSPB1|NOTCH4|PA2G4|PER1|RARA|UPF1|RPL18|RPL23A|RPL27A|MRPL12|RXRA|SOX4|TARBP2|UCN|ZFP36|BTG2|AIMP2|YBX3|DENR|YARS1|EIF3H|PER2|EIF2B4|EIF2B5|QKI|TOB1|CASC3|CPEB3|SAMD4A|PASK|PPP1R15A|EIF2AK1|EIF3K|COA3|MRPS18B|DDX25|MRPL30|BANK1|SAMD4B|MRPS18A|CNOT11|EEFSEC|METTL16|ZC3H12A|UQCC2|PYM1|UNK|DPH6|MRPL52|PATL2|RPL22L1|LTO1|CNOT6L|NANOS3|MIR106B|MIR125B1|MIR132|MIR148A|MIR200C|MIR21|MIR212|MIR27B|MIR29A|MIR29B1", "Description": "translation", "_MEMBER_Input_ID": 1.0, "_GeneInHitList": 1562.0, "Category": "GO Biological Processes", "Log_q_value_": -3.3265428815774203, "Symbol": "GO:0006412", "LogP": -5.612225185400247, "_RANK_": 1.0, "FirstInGroupByEnrichment": 0.0, "_TotalGeneInLibrary": 24367.0, "_GeneInGO": 723.0, "shared_name": "68", "_InGO": 5.057618437900128, "Gene": "GO:0006412", "_COUNT_001": 79.0, "SUID": 579265, "id_original": "68", "selected": false, "CategoryID": 19.0, "STDV_InGO": 0.5544502413377405, "EvidenceCutoff": 0.0, "PARENT_GO": "19_GO:0008152 metabolic process", "FirstInGroupByLogP": 0.0, "GROUP_ID": 13.0, "name": "68", "BestEnrichmentInGroup": 2.455535400957936, "CLUSTER_LABEL": " ", "GeneID": "207|468|677|678|740|1153|1612|1719|1915|1956|1981|2064|2309|2475|2872|3315|4855|5036|5187|5914|5976|6141|6147|6157|6182|6256|6659|6895|7349|7538|7832|7965|8531|8562|8565|8667|8864|8890|8893|9444|10140|22794|22849|23034|23178|23645|27102|27335|28958|28973|29118|51263|55024|55095|55168|55571|60678|79066|80149|84300|84305|85451|89978|122704|197135|200916|220064|246175|342977|406900|406911|406921|406940|406985|406991|406994|407019|407021|407024", "_LogP_Input_ID": -5.612225185400247, "Z_score": 5.033131636954529, "GiniIndex": 0.0, "_COUNT_001_PCT": 100.0}, "position": {"x": -236.75903393090312, "y": 761.0486102646331}, "selected": false}, {"data": {"id": "579264", "Enrichment": 1.687035552311319, "BestLogPInGroup": -8.621070944395578, "_PATTERN_": "M1", "_GeneInGOAndHitList": 81.0, "Hits": "AKT1|ATF4|ZFP36L1|ZFP36L2|MRPL49|CIRBP|DAPK1|DHFR|EEF1A1|EGFR|EIF4G1|ERBB2|FOXO3|MTOR|MKNK2|HSPB1|NFE2L2|NOTCH4|PA2G4|PER1|RARA|UPF1|RPL18|RPL23A|RPL27A|MRPL12|RXRA|SOX4|TARBP2|UCN|ZFP36|BTG2|AIMP2|YBX3|DENR|YARS1|EIF3H|PER2|EIF2B4|EIF2B5|QKI|TOB1|CASC3|CPEB3|SAMD4A|PASK|PPP1R15A|EIF2AK1|EIF3K|COA3|MRPS18B|DDX25|MRPL30|BANK1|SAMD4B|MRPS18A|CNOT11|EEFSEC|GGCT|METTL16|ZC3H12A|UQCC2|PYM1|UNK|DPH6|MRPL52|PATL2|RPL22L1|LTO1|CNOT6L|NANOS3|MIR106B|MIR125B1|MIR132|MIR148A|MIR200C|MIR21|MIR212|MIR27B|MIR29A|MIR29B1", "Description": "peptide biosynthetic process", "_MEMBER_Input_ID": 1.0, "_GeneInHitList": 1562.0, "Category": "GO Biological Processes", "Log_q_value_": -3.306817106661465, "Symbol": "GO:0043043", "LogP": -5.56522532561763, "_RANK_": 1.0, "FirstInGroupByEnrichment": 0.0, "_TotalGeneInLibrary": 24367.0, "_GeneInGO": 749.0, "shared_name": "67", "_InGO": 5.185659411011524, "Gene": "GO:0043043", "_COUNT_001": 81.0, "SUID": 579264, "id_original": "67", "selected": false, "CategoryID": 19.0, "STDV_InGO": 0.5610460309948695, "EvidenceCutoff": 0.0, "PARENT_GO": "19_GO:0008152 metabolic process", "FirstInGroupByLogP": 0.0, "GROUP_ID": 13.0, "name": "67", "BestEnrichmentInGroup": 2.455535400957936, "CLUSTER_LABEL": " ", "GeneID": "207|468|677|678|740|1153|1612|1719|1915|1956|1981|2064|2309|2475|2872|3315|4780|4855|5036|5187|5914|5976|6141|6147|6157|6182|6256|6659|6895|7349|7538|7832|7965|8531|8562|8565|8667|8864|8890|8893|9444|10140|22794|22849|23034|23178|23645|27102|27335|28958|28973|29118|51263|55024|55095|55168|55571|60678|79017|79066|80149|84300|84305|85451|89978|122704|197135|200916|220064|246175|342977|406900|406911|406921|406940|406985|406991|406994|407019|407021|407024", "_LogP_Input_ID": -5.56522532561763, "Z_score": 4.998229085280297, "GiniIndex": 0.0, "_COUNT_001_PCT": 100.0}, "position": {"x": -177.71852184594218, "y": 716.9538837021331}, "selected": false}, {"data": {"id": "579263", "Enrichment": 2.31354033290653, "BestLogPInGroup": -8.621070944395578, "_PATTERN_": "M1", "_GeneInGOAndHitList": 35.0, "Hits": "ATF4|ZFP36L1|ZFP36L2|CIRBP|DAPK1|DHFR|EIF4G1|NOTCH4|RARA|UPF1|ZFP36|BTG2|TOB1|ABCA7|CPEB3|SAMD4A|EIF2AK1|BANK1|SAMD4B|METTL16|ZC3H12A|UNK|PATL2|CNOT6L|NANOS3|MIR106B|MIR125B1|MIR132|MIR148A|MIR200C|MIR21|MIR212|MIR27B|MIR29A|MIR29B1", "Description": "negative regulation of cellular amide metabolic process", "_MEMBER_Input_ID": 1.0, "_GeneInHitList": 1562.0, "Category": "GO Biological Processes", "Log_q_value_": -3.265852058684192, "Symbol": "GO:0034249", "LogP": -5.51114154305111, "_RANK_": 1.0, "FirstInGroupByEnrichment": 0.0, "_TotalGeneInLibrary": 24367.0, "_GeneInGO": 236.0, "shared_name": "66", "_InGO": 2.240717029449424, "Gene": "GO:0034249", "_COUNT_001": 35.0, "SUID": 579263, "id_original": "66", "selected": false, "CategoryID": 19.0, "STDV_InGO": 0.3744829043781526, "EvidenceCutoff": 0.0, "PARENT_GO": "19_GO:0048519 negative regulation of biological process", "FirstInGroupByLogP": 0.0, "GROUP_ID": 13.0, "name": "66", "BestEnrichmentInGroup": 2.455535400957936, "CLUSTER_LABEL": " ", "GeneID": "468|677|678|1153|1612|1719|1981|4855|5914|5976|7538|7832|10140|10347|22849|23034|27102|55024|55095|79066|80149|85451|197135|246175|342977|406900|406911|406921|406940|406985|406991|406994|407019|407021|407024", "_LogP_Input_ID": -5.51114154305111, "Z_score": 5.30675882295899, "GiniIndex": 0.0, "_COUNT_001_PCT": 100.0}, "position": {"x": -130.2983634597117, "y": 835.3411212509612}, "selected": false}, {"data": {"id": "579262", "Enrichment": 2.3399807938540333, "BestLogPInGroup": -8.621070944395578, "_PATTERN_": "M1", "_GeneInGOAndHitList": 30.0, "Hits": "AKT1|ZFP36L1|ZFP36L2|CIRBP|EIF4G1|MTOR|HNRNPU|HSPA1A|HSPB1|PSMB3|PSMB8|PSMB9|PSMD1|PSMD2|UPF1|UBC|YWHAZ|ZFP36|BTG2|ANP32A|YBX3|THRAP3|TOB1|CPEB3|SAMD4A|SAMD4B|METTL16|ZC3H12A|CNOT6L|NANOS3", "Description": "regulation of mRNA catabolic process", "_MEMBER_Input_ID": 1.0, "_GeneInHitList": 1562.0, "Category": "GO Biological Processes", "Log_q_value_": -2.842843789567818, "Symbol": "GO:0061013", "LogP": -4.916437181063444, "_RANK_": 1.0, "FirstInGroupByEnrichment": 0.0, "_TotalGeneInLibrary": 24367.0, "_GeneInGO": 200.0, "shared_name": "65", "_InGO": 1.9206145966709347, "Gene": "GO:0061013", "_COUNT_001": 30.0, "SUID": 579262, "id_original": "65", "selected": false, "CategoryID": 19.0, "STDV_InGO": 0.3472709584609166, "EvidenceCutoff": 0.0, "PARENT_GO": "19_GO:0048519 negative regulation of biological process", "FirstInGroupByLogP": 0.0, "GROUP_ID": 13.0, "name": "65", "BestEnrichmentInGroup": 2.455535400957936, "CLUSTER_LABEL": " ", "GeneID": "207|677|678|1153|1981|2475|3192|3303|3315|5691|5696|5698|5707|5708|5976|7316|7534|7538|7832|8125|8531|9967|10140|22849|23034|55095|79066|80149|246175|342977", "_LogP_Input_ID": -4.916437181063444, "Z_score": 4.979890539037137, "GiniIndex": 0.0, "_COUNT_001_PCT": 100.0}, "position": {"x": -121.22519756615702, "y": 754.8796039169769}, "selected": false}, {"data": {"id": "579261", "Enrichment": 2.366272712886101, "BestLogPInGroup": -8.621070944395578, "_PATTERN_": "M1", "_GeneInGOAndHitList": 27.0, "Hits": "AKT1|ZFP36L1|ZFP36L2|CIRBP|EIF4G1|MTOR|HNRNPU|HSPA1A|HSPB1|PSMB3|PSMB8|PSMB9|PSMD1|PSMD2|UPF1|UBC|YWHAZ|ZFP36|ANP32A|YBX3|THRAP3|CPEB3|SAMD4A|SAMD4B|METTL16|ZC3H12A|CNOT6L", "Description": "regulation of mRNA stability", "_MEMBER_Input_ID": 1.0, "_GeneInHitList": 1562.0, "Category": "GO Biological Processes", "Log_q_value_": -2.5786213420744057, "Symbol": "GO:0043488", "LogP": -4.577581115273128, "_RANK_": 1.0, "FirstInGroupByEnrichment": 0.0, "_TotalGeneInLibrary": 24367.0, "_GeneInGO": 178.0, "shared_name": "64", "_InGO": 1.7285531370038412, "Gene": "GO:0043488", "_COUNT_001": 27.0, "SUID": 579261, "id_original": "64", "selected": false, "CategoryID": 19.0, "STDV_InGO": 0.3297725691215553, "EvidenceCutoff": 0.0, "PARENT_GO": "19_GO:0048519 negative regulation of biological process", "FirstInGroupByLogP": 0.0, "GROUP_ID": 13.0, "name": "64", "BestEnrichmentInGroup": 2.455535400957936, "CLUSTER_LABEL": " ", "GeneID": "207|677|678|1153|1981|2475|3192|3303|3315|5691|5696|5698|5707|5708|5976|7316|7534|7538|8125|8531|9967|22849|23034|55095|79066|80149|246175", "_LogP_Input_ID": -4.577581115273128, "Z_score": 4.788020873243368, "GiniIndex": 0.0, "_COUNT_001_PCT": 100.0}, "position": {"x": -123.06399608910624, "y": 909.6545978134612}, "selected": false}, {"data": {"id": "579260", "Enrichment": 1.855119908641035, "BestLogPInGroup": -8.621070944395578, "_PATTERN_": "M1", "_GeneInGOAndHitList": 44.0, "Hits": "AKT1|ZFP36L1|ZFP36L2|CIRBP|DDX5|DDX6|EIF4G1|MTOR|HNRNPU|HSPA1A|HSPA1B|HSPB1|PSMB3|PSMB8|PSMB9|PSMD1|PSMD2|UPF1|RPL18|RPL23A|RPL27A|UBC|YWHAZ|ZFP36|BTG2|ANP32A|YBX3|THRAP3|TOB1|CASC3|CPEB3|SAMD4A|SMG6|SMG5|LSM4|SAMD4B|CNOT11|METTL16|TUT7|ZC3H12A|PYM1|PATL2|CNOT6L|NANOS3", "Description": "mRNA catabolic process", "_MEMBER_Input_ID": 1.0, "_GeneInHitList": 1562.0, "Category": "GO Biological Processes", "Log_q_value_": -2.3173070855236997, "Symbol": "GO:0006402", "LogP": -4.230855522682361, "_RANK_": 1.0, "FirstInGroupByEnrichment": 0.0, "_TotalGeneInLibrary": 24367.0, "_GeneInGO": 370.0, "shared_name": "63", "_InGO": 2.816901408450704, "Gene": "GO:0006402", "_COUNT_001": 44.0, "SUID": 579260, "id_original": "63", "selected": false, "CategoryID": 19.0, "STDV_InGO": 0.4186399594939643, "EvidenceCutoff": 0.0, "PARENT_GO": "19_GO:0048519 negative regulation of biological process", "FirstInGroupByLogP": 0.0, "GROUP_ID": 13.0, "name": "63", "BestEnrichmentInGroup": 2.455535400957936, "CLUSTER_LABEL": " ", "GeneID": "207|677|678|1153|1655|1656|1981|2475|3192|3303|3304|3315|5691|5696|5698|5707|5708|5976|6141|6147|6157|7316|7534|7538|7832|8125|8531|9967|10140|22794|22849|23034|23293|23381|25804|55095|55571|79066|79670|80149|84305|197135|246175|342977", "_LogP_Input_ID": -4.230855522682361, "Z_score": 4.337774789405793, "GiniIndex": 0.0, "_COUNT_001_PCT": 100.0}, "position": {"x": -183.92495800316874, "y": 871.3175311630706}, "selected": false}, {"data": {"id": "579259", "Enrichment": 2.1767263198642173, "BestLogPInGroup": -8.219544186232241, "_PATTERN_": "M1", "_GeneInGOAndHitList": 36.0, "Hits": "B2M|ZFP36L1|RUNX1|CBFB|CD4|CEBPB|FOXO3|FOS|MTOR|GATA2|H3-3B|HOXA5|HOXA7|HOXA9|HSPA1A|HSPA1B|ID2|ITPKB|MEIS1|MEIS2|ROR2|RARA|RARG|RB1|KLF10|ZFP36|ZBTB16|H3C8|H3C7|RNF41|TRIB1|FOXP1|KLF13|PAF1|KMT2E|MIR125B1", "Description": "regulation of myeloid cell differentiation", "_MEMBER_Input_ID": 1.0, "_GeneInHitList": 1562.0, "Category": "GO Biological Processes", "Log_q_value_": -2.9164461488779456, "Symbol": "GO:0045637", "LogP": -5.036895121233299, "_RANK_": 1.0, "FirstInGroupByEnrichment": 0.0, "_TotalGeneInLibrary": 24367.0, "_GeneInGO": 258.0, "shared_name": "62", "_InGO": 2.3047375160051216, "Gene": "GO:0045637", "_COUNT_001": 36.0, "SUID": 579259, "id_original": "62", "selected": false, "CategoryID": 19.0, "STDV_InGO": 0.37967060373244593, "EvidenceCutoff": 0.0, "PARENT_GO": "19_GO:0002376 immune system process", "FirstInGroupByLogP": 0.0, "GROUP_ID": 14.0, "name": "62", "BestEnrichmentInGroup": 3.1908829007100454, "CLUSTER_LABEL": " ", "GeneID": "567|677|861|865|920|1051|2309|2353|2475|2624|3021|3202|3204|3205|3303|3304|3398|3707|4211|4212|4920|5914|5916|5925|7071|7538|7704|8355|8968|10193|10221|27086|51621|54623|55904|406911", "_LogP_Input_ID": -5.036895121233299, "Z_score": 4.972941133446377, "GiniIndex": 0.0, "_COUNT_001_PCT": 100.0}, "position": {"x": -277.37988306344096, "y": 171.09616809544366}, "selected": false}, {"data": {"id": "579258", "Enrichment": 2.324980916970354, "BestLogPInGroup": -8.219544186232241, "_PATTERN_": "M1", "_GeneInGOAndHitList": 31.0, "Hits": "ANXA2|ZFP36L1|RUNX1|CD4|CEBPB|CCR7|EPHA2|FASN|FOS|MTOR|GATA2|HOXA7|ID2|IFI16|JUNB|LTBR|MYH9|ROR2|RARA|RB1|SP3|KLF10|CCN4|SH3PXD2A|TSPAN2|TRIB1|FOXP1|EIF2AK1|IL25|IL31RA|MIR125B1", "Description": "myeloid leukocyte differentiation", "_MEMBER_Input_ID": 1.0, "_GeneInHitList": 1562.0, "Category": "GO Biological Processes", "Log_q_value_": -2.891959501297568, "Symbol": "GO:0002573", "LogP": -5.00092250581151, "_RANK_": 1.0, "FirstInGroupByEnrichment": 0.0, "_TotalGeneInLibrary": 24367.0, "_GeneInGO": 208.0, "shared_name": "61", "_InGO": 1.9846350832266326, "Gene": "GO:0002573", "_COUNT_001": 31.0, "SUID": 579258, "id_original": "61", "selected": false, "CategoryID": 19.0, "STDV_InGO": 0.3528961317999472, "EvidenceCutoff": 0.0, "PARENT_GO": "19_GO:0002376 immune system process", "FirstInGroupByLogP": 0.0, "GROUP_ID": 14.0, "name": "61", "BestEnrichmentInGroup": 3.1908829007100454, "CLUSTER_LABEL": " ", "GeneID": "302|677|861|920|1051|1236|1969|2194|2353|2475|2624|3204|3398|3428|3726|4055|4627|4920|5914|5925|6670|7071|8840|9644|10100|10221|27086|27102|64806|133396|406911", "_LogP_Input_ID": -5.00092250581151, "Z_score": 5.022493812504384, "GiniIndex": 0.0, "_COUNT_001_PCT": 100.0}, "position": {"x": -304.1570131903941, "y": 38.11419254002374}, "selected": false}, {"data": {"id": "579257", "Enrichment": 3.1908829007100454, "BestLogPInGroup": -8.219544186232241, "_PATTERN_": "M1", "_GeneInGOAndHitList": 9.0, "Hits": "CD4|GATA2|ID2|ROR2|RB1|TSPAN2|TRIB1|EIF2AK1|IL31RA", "Description": "macrophage differentiation", "_MEMBER_Input_ID": 1.0, "_GeneInHitList": 1562.0, "Category": "GO Biological Processes", "Log_q_value_": -1.2907261509923595, "Symbol": "GO:0030225", "LogP": -2.784969516437186, "_RANK_": 1.0, "FirstInGroupByEnrichment": 0.0, "_TotalGeneInLibrary": 24367.0, "_GeneInGO": 44.0, "shared_name": "60", "_InGO": 0.5761843790012804, "Gene": "GO:0030225", "_COUNT_001": 9.0, "SUID": 579257, "id_original": "60", "selected": false, "CategoryID": 19.0, "STDV_InGO": 0.19150734627097452, "EvidenceCutoff": 0.0, "PARENT_GO": "19_GO:0002376 immune system process", "FirstInGroupByLogP": 0.0, "GROUP_ID": 14.0, "name": "60", "BestEnrichmentInGroup": 3.1908829007100454, "CLUSTER_LABEL": " ", "GeneID": "920|2624|3398|4920|5925|10100|10221|27102|133396", "_LogP_Input_ID": -2.784969516437186, "Z_score": 3.8067515266240517, "GiniIndex": 0.0, "_COUNT_001_PCT": 100.0}, "position": {"x": -430.98107935250346, "y": 61.62023502049249}, "selected": false}, {"data": {"id": "579256", "Enrichment": 2.2515279116121283, "BestLogPInGroup": -8.219544186232241, "_PATTERN_": "M1", "_GeneInGOAndHitList": 14.0, "Hits": "ZFP36L1|RUNX1|CD4|FOXO3|FOS|GATA2|HOXA5|HSPA1A|HSPA1B|ID2|ROR2|RB1|KLF10|TRIB1", "Description": "positive regulation of myeloid cell differentiation", "_MEMBER_Input_ID": 1.0, "_GeneInHitList": 1562.0, "Category": "GO Biological Processes", "Log_q_value_": -1.0624154238333992, "Symbol": "GO:0045639", "LogP": -2.456708905779763, "_RANK_": 1.0, "FirstInGroupByEnrichment": 0.0, "_TotalGeneInLibrary": 24367.0, "_GeneInGO": 97.0, "shared_name": "59", "_InGO": 0.8962868117797695, "Gene": "GO:0045639", "_COUNT_001": 14.0, "SUID": 579256, "id_original": "59", "selected": false, "CategoryID": 19.0, "STDV_InGO": 0.23846681525444388, "EvidenceCutoff": 0.0, "PARENT_GO": "19_GO:0002376 immune system process", "FirstInGroupByLogP": 0.0, "GROUP_ID": 14.0, "name": "59", "BestEnrichmentInGroup": 3.1908829007100454, "CLUSTER_LABEL": " ", "GeneID": "677|861|920|2309|2353|2624|3202|3303|3304|3398|4920|5925|7071|10221", "_LogP_Input_ID": -2.456708905779763, "Z_score": 3.232281042647679, "GiniIndex": 0.0, "_COUNT_001_PCT": 100.0}, "position": {"x": -343.8961489325816, "y": 79.39794498142999}, "selected": false}, {"data": {"id": "579255", "Enrichment": 2.079982927870252, "BestLogPInGroup": -8.219544186232241, "_PATTERN_": "M1", "_GeneInGOAndHitList": 16.0, "Hits": "ZFP36L1|RUNX1|CD4|CEBPB|FOS|MTOR|GATA2|HOXA7|ID2|ROR2|RARA|RB1|KLF10|TRIB1|FOXP1|MIR125B1", "Description": "regulation of myeloid leukocyte differentiation", "_MEMBER_Input_ID": 1.0, "_GeneInHitList": 1562.0, "Category": "GO Biological Processes", "Log_q_value_": -1.0055782424841617, "Symbol": "GO:0002761", "LogP": -2.376271336015337, "_RANK_": 1.0, "FirstInGroupByEnrichment": 0.0, "_TotalGeneInLibrary": 24367.0, "_GeneInGO": 120.0, "shared_name": "58", "_InGO": 1.0243277848911652, "Gene": "GO:0002761", "_COUNT_001": 16.0, "SUID": 579255, "id_original": "58", "selected": false, "CategoryID": 19.0, "STDV_InGO": 0.25476701097987803, "EvidenceCutoff": 0.0, "PARENT_GO": "19_GO:0002376 immune system process", "FirstInGroupByLogP": 0.0, "GROUP_ID": 14.0, "name": "58", "BestEnrichmentInGroup": 3.1908829007100454, "CLUSTER_LABEL": " ", "GeneID": "677|861|920|1051|2353|2475|2624|3204|3398|4920|5914|5925|7071|10221|27086|406911", "_LogP_Input_ID": -2.376271336015337, "Z_score": 3.1038169716778716, "GiniIndex": 0.0, "_COUNT_001_PCT": 100.0}, "position": {"x": -308.94747949898783, "y": 132.94756656346124}, "selected": false}, {"data": {"id": "579254", "Enrichment": 2.5071222791293217, "BestLogPInGroup": -8.219544186232241, "_PATTERN_": "M1", "_GeneInGOAndHitList": 9.0, "Hits": "ZFP36L1|RUNX1|CD4|FOS|ID2|ROR2|RB1|KLF10|TRIB1", "Description": "positive regulation of myeloid leukocyte differentiation", "_MEMBER_Input_ID": 1.0, "_GeneInHitList": 1562.0, "Category": "GO Biological Processes", "Log_q_value_": -0.7855360641132059, "Symbol": "GO:0002763", "LogP": -2.055036510828009, "_RANK_": 1.0, "FirstInGroupByEnrichment": 0.0, "_TotalGeneInLibrary": 24367.0, "_GeneInGO": 56.0, "shared_name": "57", "_InGO": 0.5761843790012804, "Gene": "GO:0002763", "_COUNT_001": 9.0, "SUID": 579254, "id_original": "57", "selected": false, "CategoryID": 19.0, "STDV_InGO": 0.19150734627097452, "EvidenceCutoff": 0.0, "PARENT_GO": "19_GO:0002376 immune system process", "FirstInGroupByLogP": 0.0, "GROUP_ID": 14.0, "name": "57", "BestEnrichmentInGroup": 3.1908829007100454, "CLUSTER_LABEL": " ", "GeneID": "677|861|920|2353|3398|4920|5925|7071|10221", "_LogP_Input_ID": -2.055036510828009, "Z_score": 2.9550081847974945, "GiniIndex": 0.0, "_COUNT_001_PCT": 100.0}, "position": {"x": -378.7042544013316, "y": 150.03431277928155}, "selected": false}, {"data": {"id": "579253", "Enrichment": 1.8404343322447454, "BestLogPInGroup": -8.096413799222795, "_PATTERN_": "M1", "_GeneInGOAndHitList": 84.0, "Hits": "ACACA|AKT1|ATP1A1|ATP6V0A1|BCL2L2|ZFP36L1|ZFP36L2|KLF9|RUNX1|CBFB|CHRM1|CTSB|DDX5|EGFR|FHL2|FOXO3|FOS|GPT|NR4A1|FOXA1|HNRNPU|IGF1R|INHBB|ISL1|JUNB|JUND|NFE2L2|NKX3-1|NOTCH1|NR4A2|P2RY6|SERPINF1|PER1|PKM|PRKAR1A|RARA|RARB|RARG|RB1|REG1A|RORC|RXRA|SHC1|SP1|SREBF1|STAT6|VAMP2|THRA|THRB|XBP1|ZFP36|NRIP1|PIK3R3|IRS2|SOCS2|CFLAR|GDF15|NR1D1|NCOR2|RAMP1|BAIAP2|DDX17|SORBS1|ABHD2|PHB2|CBX3|KANK1|RHOQ|ATP6V0A2|WBP2|FOXP1|ZBTB7B|ZBTB7A|APPL2|BAIAP2L1|PMEPA1|ARID1B|NUCKS1|OR51E2|SLC25A33|ADTRP|ATP6V1E2|EGLN2|FOXO6", "Description": "cellular response to hormone stimulus", "_MEMBER_Input_ID": 1.0, "_GeneInHitList": 1562.0, "Category": "GO Biological Processes", "Log_q_value_": -4.615828549812055, "Symbol": "GO:0032870", "LogP": -7.343870002095332, "_RANK_": 1.0, "FirstInGroupByEnrichment": 0.0, "_TotalGeneInLibrary": 24367.0, "_GeneInGO": 712.0, "shared_name": "56", "_InGO": 5.377720870678617, "Gene": "GO:0032870", "_COUNT_001": 84.0, "SUID": 579253, "id_original": "56", "selected": false, "CategoryID": 19.0, "STDV_InGO": 0.570762347817035, "EvidenceCutoff": 0.0, "PARENT_GO": "19_GO:0050896 response to stimulus", "FirstInGroupByLogP": 0.0, "GROUP_ID": 15.0, "name": "56", "BestEnrichmentInGroup": 2.631303703932246, "CLUSTER_LABEL": " ", "GeneID": "31|207|476|535|599|677|678|687|861|865|1128|1508|1655|1956|2274|2309|2353|2875|3164|3169|3192|3480|3625|3670|3726|3727|4780|4824|4851|4929|5031|5176|5187|5315|5573|5914|5915|5916|5925|5967|6097|6256|6464|6667|6720|6778|6844|7067|7068|7494|7538|8204|8503|8660|8835|8837|9518|9572|9612|10267|10458|10521|10580|11057|11331|11335|23189|23433|23545|23558|27086|51043|51341|55198|55971|56937|57492|64710|81285|84275|84830|90423|112398|100132074", "_LogP_Input_ID": -7.343870002095332, "Z_score": 5.956620095692685, "GiniIndex": 0.0, "_COUNT_001_PCT": 100.0}, "position": {"x": 76.52084468542989, "y": 780.8180804794769}, "selected": false}, {"data": {"id": "579252", "Enrichment": 1.9499839948783608, "BestLogPInGroup": -8.096413799222795, "_PATTERN_": "M1", "_GeneInGOAndHitList": 68.0, "Hits": "ABL1|ACTB|AKT1|ATF4|ATP1A1|BCL2L2|ZFP36L1|ZFP36L2|KLF9|RUNX1|CBFB|CDKN1B|CEBPB|COL1A1|DDX5|SLC26A3|EGFR|FHL2|FOXF1|FOXO3|RAPGEF1|NR4A1|FOXA1|HNRNPU|HSP90AB1|ID1|ID3|ISL1|JUP|NKX3-1|NR4A2|P2RY6|PDGFB|SERPINF1|PER1|PTCH1|RARA|RARB|RARG|RB1|RORC|RXRA|THRA|THRB|EZR|ZFP36|NRIP1|CFLAR|NR1D1|NCOR2|FLOT1|DDX17|ABHD2|PHB2|CBX3|WBP2|TIPARP|FOXP1|ZBTB7A|SSH1|TRPM4|PMEPA1|ZC3H12A|OR51E2|ADTRP|EGLN2|MIR125B1|MIR21", "Description": "cellular response to organic cyclic compound", "_MEMBER_Input_ID": 1.0, "_GeneInHitList": 1562.0, "Category": "GO Biological Processes", "Log_q_value_": -4.348012921678118, "Symbol": "GO:0071407", "LogP": -6.995021123052176, "_RANK_": 1.0, "FirstInGroupByEnrichment": 0.0, "_TotalGeneInLibrary": 24367.0, "_GeneInGO": 544.0, "shared_name": "55", "_InGO": 4.353393085787452, "Gene": "GO:0071407", "_COUNT_001": 68.0, "SUID": 579252, "id_original": "55", "selected": false, "CategoryID": 19.0, "STDV_InGO": 0.5163072349210871, "EvidenceCutoff": 0.0, "PARENT_GO": "19_GO:0050896 response to stimulus", "FirstInGroupByLogP": 0.0, "GROUP_ID": 15.0, "name": "55", "BestEnrichmentInGroup": 2.631303703932246, "CLUSTER_LABEL": " ", "GeneID": "25|60|207|468|476|599|677|678|687|861|865|1027|1051|1277|1655|1811|1956|2274|2294|2309|2889|3164|3169|3192|3326|3397|3399|3670|3728|4824|4929|5031|5155|5176|5187|5727|5914|5915|5916|5925|6097|6256|7067|7068|7430|7538|8204|8837|9572|9612|10211|10521|11057|11331|11335|23558|25976|27086|51341|54434|54795|56937|80149|81285|84830|112398|406911|406991", "_LogP_Input_ID": -6.995021123052176, "Z_score": 5.864555617569266, "GiniIndex": 0.0, "_COUNT_001_PCT": 100.0}, "position": {"x": 204.29694865881856, "y": 738.03808170018}, "selected": false}, {"data": {"id": "579251", "Enrichment": 2.1015371551020676, "BestLogPInGroup": -8.096413799222795, "_PATTERN_": "M1", "_GeneInGOAndHitList": 52.0, "Hits": "ATP1A1|CCND1|ZFP36L1|ZFP36L2|KLF9|RUNX1|CBFB|COL1A1|CLDN4|DDX5|EGFR|FHL2|FOXO3|FOS|NR4A1|FOXA1|HNRNPU|ISL1|NKX3-1|NOTCH1|NR4A2|SERPINF1|PER1|RARA|RARB|RARG|RB1|RORC|RXRA|SDC1|SPARC|SREBF1|THRA|THRB|UCN|ZFP36|FOSL1|NRIP1|CFLAR|NR1D1|NCOR2|DDX17|ABHD2|PHB2|CBX3|WBP2|FOXP1|ZBTB7A|PMEPA1|OR51E2|ADTRP|EGLN2", "Description": "response to steroid hormone", "_MEMBER_Input_ID": 1.0, "_GeneInHitList": 1562.0, "Category": "GO Biological Processes", "Log_q_value_": -3.9776739665613756, "Symbol": "GO:0048545", "LogP": -6.490600051887265, "_RANK_": 1.0, "FirstInGroupByEnrichment": 0.0, "_TotalGeneInLibrary": 24367.0, "_GeneInGO": 386.0, "shared_name": "54", "_InGO": 3.329065300896287, "Gene": "GO:0048545", "_COUNT_001": 52.0, "SUID": 579251, "id_original": "54", "selected": false, "CategoryID": 19.0, "STDV_InGO": 0.45390879852988153, "EvidenceCutoff": 0.0, "PARENT_GO": "19_GO:0050896 response to stimulus", "FirstInGroupByLogP": 0.0, "GROUP_ID": 15.0, "name": "54", "BestEnrichmentInGroup": 2.631303703932246, "CLUSTER_LABEL": " ", "GeneID": "476|595|677|678|687|861|865|1277|1364|1655|1956|2274|2309|2353|3164|3169|3192|3670|4824|4851|4929|5176|5187|5914|5915|5916|5925|6097|6256|6382|6678|6720|7067|7068|7349|7538|8061|8204|8837|9572|9612|10521|11057|11331|11335|23558|27086|51341|56937|81285|84830|112398", "_LogP_Input_ID": -6.490600051887265, "Z_score": 5.709219688425162, "GiniIndex": 0.0, "_COUNT_001_PCT": 100.0}, "position": {"x": 98.36489986121114, "y": 839.4090838974456}, "selected": false}, {"data": {"id": "579250", "Enrichment": 1.8233616575485976, "BestLogPInGroup": -8.096413799222795, "_PATTERN_": "M1", "_GeneInGOAndHitList": 72.0, "Hits": "ABL1|ACACA|AKT1|ATP1A1|BCL2L2|ZFP36L1|ZFP36L2|KLF9|RUNX1|CBFB|CEBPB|COL1A1|DDX5|EGFR|FHL2|FOXO3|HMGB1|NR4A1|FOXA1|HNRNPU|ID3|ISL1|NKX3-1|NR4A2|P2RY6|SERPINF1|PER1|PTCH1|PTK7|RARA|RARB|RARG|RB1|UPF1|RORC|RXRA|CX3CL1|SREBF1|PRDX2|THRA|THRB|XBP1|ZFP36|NRIP1|CFLAR|LITAF|NR1D1|NCOR2|TRIB1|SPON2|DDX17|ABHD2|PHB2|CBX3|CDK19|WBP2|FOXP1|ZBTB7A|PAF1|SPHK2|PMEPA1|ZC3H12A|WNT5B|OR51E2|ADTRP|TRIM5|EGLN2|DAB2IP|SGMS1|BRINP3|MIR125B1|MIR21", "Description": "cellular response to lipid", "_MEMBER_Input_ID": 1.0, "_GeneInHitList": 1562.0, "Category": "GO Biological Processes", "Log_q_value_": -3.7590423347686484, "Symbol": "GO:0071396", "LogP": -6.228569962153308, "_RANK_": 1.0, "FirstInGroupByEnrichment": 0.0, "_TotalGeneInLibrary": 24367.0, "_GeneInGO": 616.0, "shared_name": "53", "_InGO": 4.609475032010243, "Gene": "GO:0071396", "_COUNT_001": 72.0, "SUID": 579250, "id_original": "53", "selected": false, "CategoryID": 19.0, "STDV_InGO": 0.5305640725101949, "EvidenceCutoff": 0.0, "PARENT_GO": "19_GO:0050896 response to stimulus", "FirstInGroupByLogP": 0.0, "GROUP_ID": 15.0, "name": "53", "BestEnrichmentInGroup": 2.631303703932246, "CLUSTER_LABEL": " ", "GeneID": "25|31|207|476|599|677|678|687|861|865|1051|1277|1655|1956|2274|2309|3146|3164|3169|3192|3399|3670|4824|4929|5031|5176|5187|5727|5754|5914|5915|5916|5925|5976|6097|6256|6376|6720|7001|7067|7068|7494|7538|8204|8837|9516|9572|9612|10221|10417|10521|11057|11331|11335|23097|23558|27086|51341|54623|56848|56937|80149|81029|81285|84830|85363|112398|153090|259230|339479|406911|406991", "_LogP_Input_ID": -6.228569962153308, "Z_score": 5.416985106067004, "GiniIndex": 0.0, "_COUNT_001_PCT": 100.0}, "position": {"x": 223.03952907263692, "y": 797.92187076268}, "selected": false}, {"data": {"id": "579249", "Enrichment": 2.571407465773663, "BestLogPInGroup": -8.096413799222795, "_PATTERN_": "M1", "_GeneInGOAndHitList": 30.0, "Hits": "RUNX1|CBFB|DDX5|FHL2|NR4A1|FOXA1|ISL1|NKX3-1|NR4A2|PER1|RARA|RARB|RARG|RB1|RORC|RXRA|THRA|THRB|NRIP1|NR1D1|NCOR2|DDX17|ABHD2|PHB2|WBP2|FOXP1|ZBTB7A|PMEPA1|OR51E2|EGLN2", "Description": "steroid hormone mediated signaling pathway", "_MEMBER_Input_ID": 1.0, "_GeneInHitList": 1562.0, "Category": "GO Biological Processes", "Log_q_value_": -3.4350157147153833, "Symbol": "GO:0043401", "LogP": -5.762895698332665, "_RANK_": 1.0, "FirstInGroupByEnrichment": 0.0, "_TotalGeneInLibrary": 24367.0, "_GeneInGO": 182.0, "shared_name": "52", "_InGO": 1.9206145966709347, "Gene": "GO:0043401", "_COUNT_001": 30.0, "SUID": 579249, "id_original": "52", "selected": false, "CategoryID": 19.0, "STDV_InGO": 0.3472709584609166, "EvidenceCutoff": 0.0, "PARENT_GO": "19_GO:0023052 signaling", "FirstInGroupByLogP": 0.0, "GROUP_ID": 15.0, "name": "52", "BestEnrichmentInGroup": 2.631303703932246, "CLUSTER_LABEL": " ", "GeneID": "861|865|1655|2274|3164|3169|3670|4824|4929|5187|5914|5915|5916|5925|6097|6256|7067|7068|8204|9572|9612|10521|11057|11331|23558|27086|51341|56937|81285|112398", "_LogP_Input_ID": -5.762895698332665, "Z_score": 5.568895349270733, "GiniIndex": 0.0, "_COUNT_001_PCT": 100.0}, "position": {"x": 92.04515193640646, "y": 726.5351825302581}, "selected": false}, {"data": {"id": "579248", "Enrichment": 1.991473016045986, "BestLogPInGroup": -8.096413799222795, "_PATTERN_": "M1", "_GeneInGOAndHitList": 36.0, "Hits": "ACTN4|ALOX15B|RUNX1|CBFB|DDX5|FHL2|NR4A1|FOXA1|HSPA1A|HSPA1B|ISL1|NKX3-1|NR4A2|PER1|RARA|RARB|RARG|RB1|RORC|RXRA|SREBF1|THRA|THRB|UBC|NRIP1|DHRS3|NR1D1|NCOR2|CITED2|DDX17|PHB2|WBP2|FOXP1|ZBTB7A|PMEPA1|EGLN2", "Description": "intracellular receptor signaling pathway", "_MEMBER_Input_ID": 1.0, "_GeneInHitList": 1562.0, "Category": "GO Biological Processes", "Log_q_value_": -2.3034096595169222, "Symbol": "GO:0030522", "LogP": -4.1976340821956315, "_RANK_": 1.0, "FirstInGroupByEnrichment": 0.0, "_TotalGeneInLibrary": 24367.0, "_GeneInGO": 282.0, "shared_name": "51", "_InGO": 2.3047375160051216, "Gene": "GO:0030522", "_COUNT_001": 36.0, "SUID": 579248, "id_original": "51", "selected": false, "CategoryID": 19.0, "STDV_InGO": 0.37967060373244593, "EvidenceCutoff": 0.0, "PARENT_GO": "19_GO:0023052 signaling", "FirstInGroupByLogP": 0.0, "GROUP_ID": 15.0, "name": "51", "BestEnrichmentInGroup": 2.631303703932246, "CLUSTER_LABEL": " ", "GeneID": "81|247|861|865|1655|2274|3164|3169|3303|3304|3670|4824|4929|5187|5914|5915|5916|5925|6097|6256|6720|7067|7068|7316|8204|9249|9572|9612|10370|10521|11331|23558|27086|51341|56937|112398", "_LogP_Input_ID": -4.1976340821956315, "Z_score": 4.382780535131396, "GiniIndex": 0.0, "_COUNT_001_PCT": 100.0}, "position": {"x": 142.1281444901174, "y": 778.7357440536956}, "selected": false}, {"data": {"id": "579247", "Enrichment": 2.075519445192419, "BestLogPInGroup": -8.096413799222795, "_PATTERN_": "M1", "_GeneInGOAndHitList": 31.0, "Hits": "RUNX1|CBFB|DDX5|FHL2|NR4A1|FOXA1|ISL1|NKX3-1|NR4A2|PER1|RARA|RARB|RARG|RB1|RORC|RXRA|THRA|THRB|NRIP1|NR1D1|NCOR2|DDX17|ABHD2|PHB2|WBP2|FOXP1|ZBTB7A|APPL2|PMEPA1|OR51E2|EGLN2", "Description": "hormone-mediated signaling pathway", "_MEMBER_Input_ID": 1.0, "_GeneInHitList": 1562.0, "Category": "GO Biological Processes", "Log_q_value_": -2.190943234022998, "Symbol": "GO:0009755", "LogP": -4.032928038613112, "_RANK_": 1.0, "FirstInGroupByEnrichment": 0.0, "_TotalGeneInLibrary": 24367.0, "_GeneInGO": 233.0, "shared_name": "50", "_InGO": 1.9846350832266326, "Gene": "GO:0009755", "_COUNT_001": 31.0, "SUID": 579247, "id_original": "50", "selected": false, "CategoryID": 19.0, "STDV_InGO": 0.3528961317999472, "EvidenceCutoff": 0.0, "PARENT_GO": "19_GO:0023052 signaling", "FirstInGroupByLogP": 0.0, "GROUP_ID": 15.0, "name": "50", "BestEnrichmentInGroup": 2.631303703932246, "CLUSTER_LABEL": " ", "GeneID": "861|865|1655|2274|3164|3169|3670|4824|4929|5187|5914|5915|5916|5925|6097|6256|7067|7068|8204|9572|9612|10521|11057|11331|23558|27086|51341|55198|56937|81285|112398", "_LogP_Input_ID": -4.032928038613112, "Z_score": 4.317170188908889, "GiniIndex": 0.0, "_COUNT_001_PCT": 100.0}, "position": {"x": 177.86556361853536, "y": 830.5884051865081}, "selected": false}, {"data": {"id": "579246", "Enrichment": 2.159982271249877, "BestLogPInGroup": -8.096413799222795, "_PATTERN_": "M1", "_GeneInGOAndHitList": 18.0, "Hits": "RUNX1|CBFB|DDX5|FHL2|FOXA1|ISL1|NKX3-1|PER1|RB1|NRIP1|NCOR2|DDX17|PHB2|WBP2|FOXP1|ZBTB7A|PMEPA1|EGLN2", "Description": "intracellular steroid hormone receptor signaling pathway", "_MEMBER_Input_ID": 1.0, "_GeneInHitList": 1562.0, "Category": "GO Biological Processes", "Log_q_value_": -1.2907261509923595, "Symbol": "GO:0030518", "LogP": -2.7884364055118924, "_RANK_": 1.0, "FirstInGroupByEnrichment": 0.0, "_TotalGeneInLibrary": 24367.0, "_GeneInGO": 130.0, "shared_name": "49", "_InGO": 1.1523687580025608, "Gene": "GO:0030518", "_COUNT_001": 18.0, "SUID": 579246, "id_original": "49", "selected": false, "CategoryID": 19.0, "STDV_InGO": 0.2700463777281498, "EvidenceCutoff": 0.0, "PARENT_GO": "19_GO:0023052 signaling", "FirstInGroupByLogP": 0.0, "GROUP_ID": 15.0, "name": "49", "BestEnrichmentInGroup": 2.631303703932246, "CLUSTER_LABEL": " ", "GeneID": "861|865|1655|2274|3169|3670|4824|5187|5925|8204|9612|10521|11331|23558|27086|51341|56937|112398", "_LogP_Input_ID": -2.7884364055118924, "Z_score": 3.4705730722740467, "GiniIndex": 0.0, "_COUNT_001_PCT": 100.0}, "position": {"x": 146.1317837113088, "y": 716.9538837021331}, "selected": false}, {"data": {"id": "579245", "Enrichment": 2.557356058856867, "BestLogPInGroup": -8.096413799222795, "_PATTERN_": "M1", "_GeneInGOAndHitList": 10.0, "Hits": "DDX5|FHL2|NKX3-1|RB1|NRIP1|NCOR2|DDX17|FOXP1|ZBTB7A|PMEPA1", "Description": "androgen receptor signaling pathway", "_MEMBER_Input_ID": 1.0, "_GeneInHitList": 1562.0, "Category": "GO Biological Processes", "Log_q_value_": -0.9528189069327339, "Symbol": "GO:0030521", "LogP": -2.290559457163966, "_RANK_": 1.0, "FirstInGroupByEnrichment": 0.0, "_TotalGeneInLibrary": 24367.0, "_GeneInGO": 61.0, "shared_name": "48", "_InGO": 0.6402048655569782, "Gene": "GO:0030521", "_COUNT_001": 10.0, "SUID": 579245, "id_original": "48", "selected": false, "CategoryID": 19.0, "STDV_InGO": 0.20180146473426705, "EvidenceCutoff": 0.0, "PARENT_GO": "19_GO:0023052 signaling", "FirstInGroupByLogP": 0.0, "GROUP_ID": 15.0, "name": "48", "BestEnrichmentInGroup": 2.631303703932246, "CLUSTER_LABEL": " ", "GeneID": "1655|2274|4824|5925|8204|9612|10521|27086|51341|56937", "_LogP_Input_ID": -2.290559457163966, "Z_score": 3.187232050282483, "GiniIndex": 0.0, "_COUNT_001_PCT": 100.0}, "position": {"x": 35.387070881718955, "y": 803.9924579208831}, "selected": false}, {"data": {"id": "579244", "Enrichment": 2.4417190892389913, "BestLogPInGroup": -7.604995071485691, "_PATTERN_": "M1", "_GeneInGOAndHitList": 36.0, "Hits": "AKT1|BMPR1B|RUNX2|CBFB|CEBPB|COL1A1|DDX5|EPHA2|FASN|FHL2|H3-3B|HNRNPU|ID1|ID2|ID3|ILK|JUNB|JUND|SMAD3|MEF2D|NOTCH1|PTCH1|TPM4|IFITM1|CCN4|TOB1|HDAC7|WWOX|TRPM4|BCAP29|TP53INP2|CREB3L1|SH3PXD2B|MIR200C|MIR21|MIR29B1", "Description": "osteoblast differentiation", "_MEMBER_Input_ID": 1.0, "_GeneInHitList": 1562.0, "Category": "GO Biological Processes", "Log_q_value_": -3.7590423347686484, "Symbol": "GO:0001649", "LogP": -6.215837148854855, "_RANK_": 1.0, "FirstInGroupByEnrichment": 0.0, "_TotalGeneInLibrary": 24367.0, "_GeneInGO": 230.0, "shared_name": "47", "_InGO": 2.3047375160051216, "Gene": "GO:0001649", "_COUNT_001": 36.0, "SUID": 579244, "id_original": "47", "selected": false, "CategoryID": 19.0, "STDV_InGO": 0.37967060373244593, "EvidenceCutoff": 0.0, "PARENT_GO": "19_GO:0032502 developmental process", "FirstInGroupByLogP": 0.0, "GROUP_ID": 17.0, "name": "47", "BestEnrichmentInGroup": 2.6999778390623463, "CLUSTER_LABEL": " ", "GeneID": "207|658|860|865|1051|1277|1655|1969|2194|2274|3021|3192|3397|3398|3399|3611|3726|3727|4088|4209|4851|5727|7171|8519|8840|10140|51564|51741|54795|55973|58476|90993|285590|406985|406991|407024", "_LogP_Input_ID": -6.215837148854855, "Z_score": 5.749373435570277, "GiniIndex": 0.0, "_COUNT_001_PCT": 100.0}, "position": {"x": 1421.1374509209338, "y": -26.547692304279394}, "selected": false}, {"data": {"id": "579243", "Enrichment": 2.0899349992954686, "BestLogPInGroup": -7.604995071485691, "_PATTERN_": "M1", "_GeneInGOAndHitList": 28.0, "Hits": "ADRB2|BMPR1B|RUNX2|CEBPB|DDX5|ID1|ID2|ID3|ILK|JUND|LTBP3|SMAD3|SMAD7|NOTCH1|PBX1|PTCH1|ZBTB16|IFITM1|CCN4|DHRS3|TOB1|ZMPSTE24|HDAC7|TRPM4|ANKH|MIR200C|MIR21|MIR29B1", "Description": "regulation of ossification", "_MEMBER_Input_ID": 1.0, "_GeneInHitList": 1562.0, "Category": "GO Biological Processes", "Log_q_value_": -1.9896802026747114, "Symbol": "GO:0030278", "LogP": -3.7549032353406133, "_RANK_": 1.0, "FirstInGroupByEnrichment": 0.0, "_TotalGeneInLibrary": 24367.0, "_GeneInGO": 209.0, "shared_name": "46", "_InGO": 1.792573623559539, "Gene": "GO:0030278", "_COUNT_001": 28.0, "SUID": 579243, "id_original": "46", "selected": false, "CategoryID": 19.0, "STDV_InGO": 0.33571453996393463, "EvidenceCutoff": 0.0, "PARENT_GO": "19_GO:0032501 multicellular organismal process", "FirstInGroupByLogP": 0.0, "GROUP_ID": 17.0, "name": "46", "BestEnrichmentInGroup": 2.6999778390623463, "CLUSTER_LABEL": " ", "GeneID": "154|658|860|1051|1655|3397|3398|3399|3611|3727|4054|4088|4092|4851|5087|5727|7704|8519|8840|9249|10140|10269|51564|54795|56172|406985|406991|407024", "_LogP_Input_ID": -3.7549032353406133, "Z_score": 4.141530362345323, "GiniIndex": 0.0, "_COUNT_001_PCT": 100.0}, "position": {"x": 1387.2608640068715, "y": 50.90799289402764}, "selected": false}, {"data": {"id": "579242", "Enrichment": 2.4185847998491297, "BestLogPInGroup": -7.604995071485691, "_PATTERN_": "M1", "_GeneInGOAndHitList": 20.0, "Hits": "BMPR1B|RUNX2|CEBPB|DDX5|ID1|ID2|ID3|ILK|JUND|SMAD3|NOTCH1|PTCH1|IFITM1|CCN4|TOB1|HDAC7|TRPM4|MIR200C|MIR21|MIR29B1", "Description": "regulation of osteoblast differentiation", "_MEMBER_Input_ID": 1.0, "_GeneInHitList": 1562.0, "Category": "GO Biological Processes", "Log_q_value_": -1.9376981144954906, "Symbol": "GO:0045667", "LogP": -3.682772906077548, "_RANK_": 1.0, "FirstInGroupByEnrichment": 0.0, "_TotalGeneInLibrary": 24367.0, "_GeneInGO": 129.0, "shared_name": "45", "_InGO": 1.2804097311139564, "Gene": "GO:0045667", "_COUNT_001": 20.0, "SUID": 579242, "id_original": "45", "selected": false, "CategoryID": 19.0, "STDV_InGO": 0.284469454773297, "EvidenceCutoff": 0.0, "PARENT_GO": "19_GO:0032502 developmental process", "FirstInGroupByLogP": 0.0, "GROUP_ID": 17.0, "name": "45", "BestEnrichmentInGroup": 2.6999778390623463, "CLUSTER_LABEL": " ", "GeneID": "658|860|1051|1655|3397|3398|3399|3611|3727|4088|4851|5727|8519|8840|10140|51564|54795|406985|406991|407024", "_LogP_Input_ID": -3.682772906077548, "Z_score": 4.227847980027615, "GiniIndex": 0.0, "_COUNT_001_PCT": 100.0}, "position": {"x": 1476.3439938896838, "y": 97.41587405857842}, "selected": false}, {"data": {"id": "579241", "Enrichment": 2.6999778390623463, "BestLogPInGroup": -7.604995071485691, "_PATTERN_": "M1", "_GeneInGOAndHitList": 9.0, "Hits": "ID1|ID2|ID3|SMAD3|NOTCH1|PTCH1|TOB1|HDAC7|TRPM4", "Description": "negative regulation of osteoblast differentiation", "_MEMBER_Input_ID": 1.0, "_GeneInHitList": 1562.0, "Category": "GO Biological Processes", "Log_q_value_": -0.9329570220486093, "Symbol": "GO:0045668", "LogP": -2.2694651052370807, "_RANK_": 1.0, "FirstInGroupByEnrichment": 0.0, "_TotalGeneInLibrary": 24367.0, "_GeneInGO": 52.0, "shared_name": "44", "_InGO": 0.5761843790012804, "Gene": "GO:0045668", "_COUNT_001": 9.0, "SUID": 579241, "id_original": "44", "selected": false, "CategoryID": 19.0, "STDV_InGO": 0.19150734627097452, "EvidenceCutoff": 0.0, "PARENT_GO": "19_GO:0048519 negative regulation of biological process", "FirstInGroupByLogP": 0.0, "GROUP_ID": 17.0, "name": "44", "BestEnrichmentInGroup": 2.6999778390623463, "CLUSTER_LABEL": " ", "GeneID": "3397|3398|3399|4088|4851|5727|10140|51564|54795", "_LogP_Input_ID": -2.2694651052370807, "Z_score": 3.2116291510514854, "GiniIndex": 0.0, "_COUNT_001_PCT": 100.0}, "position": {"x": 1523.782500969762, "y": -0.5965141707916928}, "selected": false}, {"data": {"id": "579240", "Enrichment": 2.204329733340756, "BestLogPInGroup": -7.604995071485691, "_PATTERN_": "M1", "_GeneInGOAndHitList": 13.0, "Hits": "ADRB2|BMPR1B|RUNX2|CEBPB|ILK|JUND|SMAD3|ZBTB16|IFITM1|CCN4|MIR200C|MIR21|MIR29B1", "Description": "positive regulation of ossification", "_MEMBER_Input_ID": 1.0, "_GeneInHitList": 1562.0, "Category": "GO Biological Processes", "Log_q_value_": -0.9136993571601828, "Symbol": "GO:0045778", "LogP": -2.2406939286830094, "_RANK_": 1.0, "FirstInGroupByEnrichment": 0.0, "_TotalGeneInLibrary": 24367.0, "_GeneInGO": 92.0, "shared_name": "43", "_InGO": 0.8322663252240717, "Gene": "GO:0045778", "_COUNT_001": 13.0, "SUID": 579240, "id_original": "43", "selected": false, "CategoryID": 19.0, "STDV_InGO": 0.22986658336957838, "EvidenceCutoff": 0.0, "PARENT_GO": "19_GO:0048518 positive regulation of biological process", "FirstInGroupByLogP": 0.0, "GROUP_ID": 17.0, "name": "43", "BestEnrichmentInGroup": 2.6999778390623463, "CLUSTER_LABEL": " ", "GeneID": "154|658|860|1051|3611|3727|4088|7704|8519|8840|406985|406991|407024", "_LogP_Input_ID": -2.2406939286830094, "Z_score": 3.028846746403711, "GiniIndex": 0.0, "_COUNT_001_PCT": 100.0}, "position": {"x": 1396.1531674736684, "y": 111.3543201035003}, "selected": false}, {"data": {"id": "579239", "Enrichment": 2.3636169634889224, "BestLogPInGroup": -7.604995071485691, "_PATTERN_": "M1", "_GeneInGOAndHitList": 10.0, "Hits": "BMPR1B|RUNX2|CEBPB|ILK|JUND|IFITM1|CCN4|MIR200C|MIR21|MIR29B1", "Description": "positive regulation of osteoblast differentiation", "_MEMBER_Input_ID": 1.0, "_GeneInHitList": 1562.0, "Category": "GO Biological Processes", "Log_q_value_": -0.7798589685164289, "Symbol": "GO:0045669", "LogP": -2.0466559281788816, "_RANK_": 1.0, "FirstInGroupByEnrichment": 0.0, "_TotalGeneInLibrary": 24367.0, "_GeneInGO": 66.0, "shared_name": "42", "_InGO": 0.6402048655569782, "Gene": "GO:0045669", "_COUNT_001": 10.0, "SUID": 579239, "id_original": "42", "selected": false, "CategoryID": 19.0, "STDV_InGO": 0.20180146473426705, "EvidenceCutoff": 0.0, "PARENT_GO": "19_GO:0048518 positive regulation of biological process", "FirstInGroupByLogP": 0.0, "GROUP_ID": 17.0, "name": "42", "BestEnrichmentInGroup": 2.6999778390623463, "CLUSTER_LABEL": " ", "GeneID": "658|860|1051|3611|3727|8519|8840|406985|406991|407024", "_LogP_Input_ID": -2.0466559281788816, "Z_score": 2.903152748585865, "GiniIndex": 0.0, "_COUNT_001_PCT": 100.0}, "position": {"x": 1466.226592766637, "y": 30.46535068211358}, "selected": false}, {"data": {"id": "579238", "Enrichment": 1.975787185736865, "BestLogPInGroup": -7.592008542189207, "_PATTERN_": "M1", "_GeneInGOAndHitList": 67.0, "Hits": "TLE5|ANXA2|BMPR1B|RUNX2|RUNX1|SERPINH1|COL1A1|DLX1|DLX2|EPHA2|ETS2|FGR|HOXA3|HOXA5|HOXA6|HOXA7|HOXA9|HOXA10|HOXA11|HOXC4|HOXC6|HOXC8|HOXD10|HOXD12|HOXD13|LTBP3|SMAD3|SMAD7|MEF2D|MEIS1|MMP14|NAB2|NPR3|ROR2|PBX1|PTPN6|RARA|RARB|RARG|SOX4|SP3|SPARC|SRF|TBX3|THRA|TRPS1|PCGF2|ZBTB16|ZFAND5|DCHS1|CCN4|DHRS3|ZMPSTE24|RAI1|MTHFD1L|TIPARP|FOXP1|SLC39A1|WWOX|ANKH|VWA1|GRHL2|WNT5B|ARID5B|HES7|SH3PXD2B|MIR21", "Description": "skeletal system development", "_MEMBER_Input_ID": 1.0, "_GeneInHitList": 1562.0, "Category": "GO Biological Processes", "Log_q_value_": -4.443958839207675, "Symbol": "GO:0001501", "LogP": -7.118351905579556, "_RANK_": 1.0, "FirstInGroupByEnrichment": 0.0, "_TotalGeneInLibrary": 24367.0, "_GeneInGO": 529.0, "shared_name": "41", "_InGO": 4.289372599231754, "Gene": "GO:0001501", "_COUNT_001": 67.0, "SUID": 579238, "id_original": "41", "selected": false, "CategoryID": 19.0, "STDV_InGO": 0.5126682869938446, "EvidenceCutoff": 0.0, "PARENT_GO": "19_GO:0032502 developmental process", "FirstInGroupByLogP": 0.0, "GROUP_ID": 18.0, "name": "41", "BestEnrichmentInGroup": 4.679961587708067, "CLUSTER_LABEL": " ", "GeneID": "166|302|658|860|861|871|1277|1745|1746|1969|2114|2268|3200|3202|3203|3204|3205|3206|3207|3221|3223|3224|3236|3238|3239|4054|4088|4092|4209|4211|4323|4665|4883|4920|5087|5777|5914|5915|5916|6659|6670|6678|6722|6926|7067|7227|7703|7704|7763|8642|8840|9249|10269|10743|25902|25976|27086|27173|51741|56172|64856|79977|81029|84159|84667|285590|406991", "_LogP_Input_ID": -7.118351905579556, "Z_score": 5.938347872508957, "GiniIndex": 0.0, "_COUNT_001_PCT": 100.0}, "position": {"x": 1139.605712639684, "y": 76.91036563572686}, "selected": false}, {"data": {"id": "579237", "Enrichment": 2.4049802603499786, "BestLogPInGroup": -7.592008542189207, "_PATTERN_": "M1", "_GeneInGOAndHitList": 37.0, "Hits": "BMPR1B|RUNX2|SERPINH1|COL1A1|DLX2|FGR|HOXA3|HOXA5|HOXA6|HOXA7|HOXA9|HOXA11|HOXC4|HOXC8|HOXD10|LTBP3|SMAD3|MEF2D|MMP14|NAB2|ROR2|RARA|RARB|RARG|THRA|PCGF2|ZFAND5|DHRS3|ZMPSTE24|MTHFD1L|TIPARP|SLC39A1|WWOX|VWA1|GRHL2|ARID5B|SH3PXD2B", "Description": "skeletal system morphogenesis", "_MEMBER_Input_ID": 1.0, "_GeneInHitList": 1562.0, "Category": "GO Biological Processes", "Log_q_value_": -3.753875562308351, "Symbol": "GO:0048705", "LogP": -6.203749901887151, "_RANK_": 1.0, "FirstInGroupByEnrichment": 0.0, "_TotalGeneInLibrary": 24367.0, "_GeneInGO": 240.0, "shared_name": "40", "_InGO": 2.3687580025608193, "Gene": "GO:0048705", "_COUNT_001": 37.0, "SUID": 579237, "id_original": "40", "selected": false, "CategoryID": 19.0, "STDV_InGO": 0.38478154982955665, "EvidenceCutoff": 0.0, "PARENT_GO": "19_GO:0032502 developmental process", "FirstInGroupByLogP": 0.0, "GROUP_ID": 18.0, "name": "40", "BestEnrichmentInGroup": 4.679961587708067, "CLUSTER_LABEL": " ", "GeneID": "658|860|871|1277|1746|2268|3200|3202|3203|3204|3205|3207|3221|3224|3236|4054|4088|4209|4323|4665|4920|5914|5915|5916|7067|7703|7763|9249|10269|25902|25976|27173|51741|64856|79977|84159|285590", "_LogP_Input_ID": -6.203749901887151, "Z_score": 5.724555598472613, "GiniIndex": 0.0, "_COUNT_001_PCT": 100.0}, "position": {"x": 1160.181167351598, "y": 133.51812320408624}, "selected": false}, {"data": {"id": "579236", "Enrichment": 3.3646782656724654, "BestLogPInGroup": -7.592008542189207, "_PATTERN_": "M1", "_GeneInGOAndHitList": 11.0, "Hits": "BMPR1B|RUNX2|HOXA11|LTBP3|SMAD3|SMAD7|RARB|RARG|TRPS1|ZBTB16|CCN4", "Description": "regulation of chondrocyte differentiation", "_MEMBER_Input_ID": 1.0, "_GeneInHitList": 1562.0, "Category": "GO Biological Processes", "Log_q_value_": -1.7984117274904843, "Symbol": "GO:0032330", "LogP": -3.4963415050252253, "_RANK_": 1.0, "FirstInGroupByEnrichment": 0.0, "_TotalGeneInLibrary": 24367.0, "_GeneInGO": 51.0, "shared_name": "39", "_InGO": 0.704225352112676, "Gene": "GO:0032330", "_COUNT_001": 11.0, "SUID": 579236, "id_original": "39", "selected": false, "CategoryID": 19.0, "STDV_InGO": 0.21158296421416498, "EvidenceCutoff": 0.0, "PARENT_GO": "19_GO:0032502 developmental process", "FirstInGroupByLogP": 0.0, "GROUP_ID": 18.0, "name": "39", "BestEnrichmentInGroup": 4.679961587708067, "CLUSTER_LABEL": " ", "GeneID": "658|860|3207|4054|4088|4092|5915|5916|7227|7704|8840", "_LogP_Input_ID": -3.4963415050252253, "Z_score": 4.424138117652016, "GiniIndex": 0.0, "_COUNT_001_PCT": 100.0}, "position": {"x": 1318.2811581963247, "y": 3.6254172867278385}, "selected": false}, {"data": {"id": "579235", "Enrichment": 2.8971190781049936, "BestLogPInGroup": -7.592008542189207, "_PATTERN_": "M1", "_GeneInGOAndHitList": 13.0, "Hits": "BMPR1B|RUNX2|SERPINH1|COL1A1|HOXA11|MEF2D|MMP14|NAB2|RARA|RARB|RARG|ZMPSTE24|VWA1", "Description": "endochondral bone morphogenesis", "_MEMBER_Input_ID": 1.0, "_GeneInHitList": 1562.0, "Category": "GO Biological Processes", "Log_q_value_": -1.6906718863138661, "Symbol": "GO:0060350", "LogP": -3.3470201139420683, "_RANK_": 1.0, "FirstInGroupByEnrichment": 0.0, "_TotalGeneInLibrary": 24367.0, "_GeneInGO": 70.0, "shared_name": "38", "_InGO": 0.8322663252240717, "Gene": "GO:0060350", "_COUNT_001": 13.0, "SUID": 579235, "id_original": "38", "selected": false, "CategoryID": 19.0, "STDV_InGO": 0.22986658336957838, "EvidenceCutoff": 0.0, "PARENT_GO": "19_GO:0032502 developmental process", "FirstInGroupByLogP": 0.0, "GROUP_ID": 18.0, "name": "38", "BestEnrichmentInGroup": 4.679961587708067, "CLUSTER_LABEL": " ", "GeneID": "658|860|871|1277|3207|4209|4323|4665|5914|5915|5916|10269|64856", "_LogP_Input_ID": -3.3470201139420683, "Z_score": 4.159919064493216, "GiniIndex": 0.0, "_COUNT_001_PCT": 100.0}, "position": {"x": 1210.3931271904653, "y": 50.84979587254327}, "selected": false}, {"data": {"id": "579234", "Enrichment": 1.9867761457251227, "BestLogPInGroup": -7.592008542189207, "_PATTERN_": "M1", "_GeneInGOAndHitList": 27.0, "Hits": "BMPR1B|RUNX2|RUNX1|SERPINH1|COL1A1|DLX2|HOXA3|HOXA5|HOXA11|HOXC4|LTBP3|SMAD3|SMAD7|MEF2D|ROR2|RARA|RARB|RARG|SRF|THRA|TRPS1|ZBTB16|CCN4|ZMPSTE24|VWA1|WNT5B|MIR21", "Description": "cartilage development", "_MEMBER_Input_ID": 1.0, "_GeneInHitList": 1562.0, "Category": "GO Biological Processes", "Log_q_value_": -1.649004455204928, "Symbol": "GO:0051216", "LogP": -3.2890053102429238, "_RANK_": 1.0, "FirstInGroupByEnrichment": 0.0, "_TotalGeneInLibrary": 24367.0, "_GeneInGO": 212.0, "shared_name": "37", "_InGO": 1.7285531370038412, "Gene": "GO:0051216", "_COUNT_001": 27.0, "SUID": 579234, "id_original": "37", "selected": false, "CategoryID": 19.0, "STDV_InGO": 0.3297725691215553, "EvidenceCutoff": 0.0, "PARENT_GO": "19_GO:0032502 developmental process", "FirstInGroupByLogP": 0.0, "GROUP_ID": 18.0, "name": "37", "BestEnrichmentInGroup": 4.679961587708067, "CLUSTER_LABEL": " ", "GeneID": "658|860|861|871|1277|1746|3200|3202|3207|3221|4054|4088|4092|4209|4920|5914|5915|5916|6722|7067|7227|7704|8840|10269|64856|81029|406991", "_LogP_Input_ID": -3.2890053102429238, "Z_score": 3.7765970390641113, "GiniIndex": 0.0, "_COUNT_001_PCT": 100.0}, "position": {"x": 1229.5634000176137, "y": 102.44727664646905}, "selected": false}, {"data": {"id": "579233", "Enrichment": 2.8563145840471766, "BestLogPInGroup": -7.592008542189207, "_PATTERN_": "M1", "_GeneInGOAndHitList": 13.0, "Hits": "BMPR1B|RUNX2|HOXA11|LTBP3|SMAD3|SMAD7|RARA|RARB|RARG|TRPS1|ZBTB16|CCN4|MIR21", "Description": "regulation of cartilage development", "_MEMBER_Input_ID": 1.0, "_GeneInHitList": 1562.0, "Category": "GO Biological Processes", "Log_q_value_": -1.649004455204928, "Symbol": "GO:0061035", "LogP": -3.285063476275855, "_RANK_": 1.0, "FirstInGroupByEnrichment": 0.0, "_TotalGeneInLibrary": 24367.0, "_GeneInGO": 71.0, "shared_name": "36", "_InGO": 0.8322663252240717, "Gene": "GO:0061035", "_COUNT_001": 13.0, "SUID": 579233, "id_original": "36", "selected": false, "CategoryID": 19.0, "STDV_InGO": 0.22986658336957838, "EvidenceCutoff": 0.0, "PARENT_GO": "19_GO:0032502 developmental process", "FirstInGroupByLogP": 0.0, "GROUP_ID": 18.0, "name": "36", "BestEnrichmentInGroup": 4.679961587708067, "CLUSTER_LABEL": " ", "GeneID": "658|860|3207|4054|4088|4092|5914|5915|5916|7227|7704|8840|406991", "_LogP_Input_ID": -3.285063476275855, "Z_score": 4.099500635029687, "GiniIndex": 0.0, "_COUNT_001_PCT": 100.0}, "position": {"x": 1307.9404904717153, "y": 87.5564456528167}, "selected": false}, {"data": {"id": "579232", "Enrichment": 4.679961587708067, "BestLogPInGroup": -7.592008542189207, "_PATTERN_": "M1", "_GeneInGOAndHitList": 6.0, "Hits": "BMPR1B|RUNX2|HOXA11|SMAD3|SMAD7|ZBTB16", "Description": "positive regulation of chondrocyte differentiation", "_MEMBER_Input_ID": 1.0, "_GeneInHitList": 1562.0, "Category": "GO Biological Processes", "Log_q_value_": -1.3856638777324064, "Symbol": "GO:0032332", "LogP": -2.913781461953367, "_RANK_": 1.0, "FirstInGroupByEnrichment": 0.0, "_TotalGeneInLibrary": 24367.0, "_GeneInGO": 20.0, "shared_name": "35", "_InGO": 0.38412291933418696, "Gene": "GO:0032332", "_COUNT_001": 6.0, "SUID": 579232, "id_original": "35", "selected": false, "CategoryID": 19.0, "STDV_InGO": 0.15651604933038119, "EvidenceCutoff": 0.0, "PARENT_GO": "19_GO:0048518 positive regulation of biological process", "FirstInGroupByLogP": 0.0, "GROUP_ID": 18.0, "name": "35", "BestEnrichmentInGroup": 4.679961587708067, "CLUSTER_LABEL": " ", "GeneID": "658|860|3207|4088|4092|7704", "_LogP_Input_ID": -2.913781461953367, "Z_score": 4.308768407872969, "GiniIndex": 0.0, "_COUNT_001_PCT": 100.0}, "position": {"x": 1317.571014153356, "y": 157.45487552342217}, "selected": false}, {"data": {"id": "579231", "Enrichment": 2.12725526714003, "BestLogPInGroup": -7.592008542189207, "_PATTERN_": "M1", "_GeneInGOAndHitList": 15.0, "Hits": "BMPR1B|RUNX2|SERPINH1|COL1A1|HOXA11|LTBP3|MEF2D|MMP14|NAB2|RARA|RARB|RARG|DHRS3|ZMPSTE24|VWA1", "Description": "bone morphogenesis", "_MEMBER_Input_ID": 1.0, "_GeneInHitList": 1562.0, "Category": "GO Biological Processes", "Log_q_value_": -0.9926178670826462, "Symbol": "GO:0060349", "LogP": -2.3536380778713375, "_RANK_": 1.0, "FirstInGroupByEnrichment": 0.0, "_TotalGeneInLibrary": 24367.0, "_GeneInGO": 110.0, "shared_name": "34", "_InGO": 0.9603072983354674, "Gene": "GO:0060349", "_COUNT_001": 15.0, "SUID": 579231, "id_original": "34", "selected": false, "CategoryID": 19.0, "STDV_InGO": 0.24675686391426488, "EvidenceCutoff": 0.0, "PARENT_GO": "19_GO:0032502 developmental process", "FirstInGroupByLogP": 0.0, "GROUP_ID": 18.0, "name": "34", "BestEnrichmentInGroup": 4.679961587708067, "CLUSTER_LABEL": " ", "GeneID": "658|860|871|1277|3207|4054|4209|4323|4665|5914|5915|5916|9249|10269|64856", "_LogP_Input_ID": -2.3536380778713375, "Z_score": 3.1011135466656667, "GiniIndex": 0.0, "_COUNT_001_PCT": 100.0}, "position": {"x": 1213.8271939629262, "y": -5.685400431778021}, "selected": false}, {"data": {"id": "579230", "Enrichment": 2.0128867043905663, "BestLogPInGroup": -7.592008542189207, "_PATTERN_": "M1", "_GeneInGOAndHitList": 16.0, "Hits": "BMPR1B|RUNX2|RUNX1|SERPINH1|HOXA11|LTBP3|SMAD3|SMAD7|MEF2D|RARB|RARG|TRPS1|ZBTB16|CCN4|VWA1|WNT5B", "Description": "chondrocyte differentiation", "_MEMBER_Input_ID": 1.0, "_GeneInHitList": 1562.0, "Category": "GO Biological Processes", "Log_q_value_": -0.9111599186721123, "Symbol": "GO:0002062", "LogP": -2.236829747484277, "_RANK_": 1.0, "FirstInGroupByEnrichment": 0.0, "_TotalGeneInLibrary": 24367.0, "_GeneInGO": 124.0, "shared_name": "33", "_InGO": 1.0243277848911652, "Gene": "GO:0002062", "_COUNT_001": 16.0, "SUID": 579230, "id_original": "33", "selected": false, "CategoryID": 19.0, "STDV_InGO": 0.25476701097987803, "EvidenceCutoff": 0.0, "PARENT_GO": "19_GO:0032502 developmental process", "FirstInGroupByLogP": 0.0, "GROUP_ID": 18.0, "name": "33", "BestEnrichmentInGroup": 4.679961587708067, "CLUSTER_LABEL": " ", "GeneID": "658|860|861|871|3207|4054|4088|4092|4209|5915|5916|7227|7704|8840|64856|81029", "_LogP_Input_ID": -2.236829747484277, "Z_score": 2.959348615477092, "GiniIndex": 0.0, "_COUNT_001_PCT": 100.0}, "position": {"x": 1271.5015561455434, "y": 46.72810702976983}, "selected": false}, {"data": {"id": "579229", "Enrichment": 4.091769694170987, "BestLogPInGroup": -7.5613451204042255, "_PATTERN_": "M1", "_GeneInGOAndHitList": 16.0, "Hits": "ALOX15B|ZFP36L1|RUNX1|CBFB|SFN|HOXA7|NOTCH1|PRKCH|PTCH1|SULT2B1|ZFP36|KLF7|PTCH2|HEY2|ESRP1|GRHL2", "Description": "regulation of epidermal cell differentiation", "_MEMBER_Input_ID": 1.0, "_GeneInHitList": 1562.0, "Category": "GO Biological Processes", "Log_q_value_": -3.6074470770164018, "Symbol": "GO:0045604", "LogP": -6.012739283942461, "_RANK_": 1.0, "FirstInGroupByEnrichment": 0.0, "_TotalGeneInLibrary": 24367.0, "_GeneInGO": 61.0, "shared_name": "32", "_InGO": 1.0243277848911652, "Gene": "GO:0045604", "_COUNT_001": 16.0, "SUID": 579229, "id_original": "32", "selected": false, "CategoryID": 19.0, "STDV_InGO": 0.25476701097987803, "EvidenceCutoff": 0.0, "PARENT_GO": "19_GO:0032502 developmental process", "FirstInGroupByLogP": 0.0, "GROUP_ID": 19.0, "name": "32", "BestEnrichmentInGroup": 4.549962654716176, "CLUSTER_LABEL": " ", "GeneID": "247|677|861|865|2810|3204|4851|5583|5727|6820|7538|8609|8643|23493|54845|79977", "_LogP_Input_ID": -6.012739283942461, "Z_score": 6.327510915254041, "GiniIndex": 0.0, "_COUNT_001_PCT": 100.0}, "position": {"x": 445.7692258232778, "y": 215.9824481552581}, "selected": false}, {"data": {"id": "579228", "Enrichment": 3.265089479796326, "BestLogPInGroup": -7.5613451204042255, "_PATTERN_": "M1", "_GeneInGOAndHitList": 18.0, "Hits": "ALOX15B|ZFP36L1|RUNX1|CBFB|SFN|HOXA7|NAB2|NOTCH1|PRKCH|PTCH1|SULT2B1|ZFP36|KLF7|PTCH2|HEY2|ESRP1|GRHL2|TMEM79", "Description": "regulation of epidermis development", "_MEMBER_Input_ID": 1.0, "_GeneInHitList": 1562.0, "Category": "GO Biological Processes", "Log_q_value_": -3.0051670699759185, "Symbol": "GO:0045682", "LogP": -5.151831402271893, "_RANK_": 1.0, "FirstInGroupByEnrichment": 0.0, "_TotalGeneInLibrary": 24367.0, "_GeneInGO": 86.0, "shared_name": "31", "_InGO": 1.1523687580025608, "Gene": "GO:0045682", "_COUNT_001": 18.0, "SUID": 579228, "id_original": "31", "selected": false, "CategoryID": 19.0, "STDV_InGO": 0.2700463777281498, "EvidenceCutoff": 0.0, "PARENT_GO": "19_GO:0032502 developmental process", "FirstInGroupByLogP": 0.0, "GROUP_ID": 19.0, "name": "31", "BestEnrichmentInGroup": 4.549962654716176, "CLUSTER_LABEL": " ", "GeneID": "247|677|861|865|2810|3204|4665|4851|5583|5727|6820|7538|8609|8643|23493|54845|79977|84283", "_LogP_Input_ID": -5.151831402271893, "Z_score": 5.507048490838912, "GiniIndex": 0.0, "_COUNT_001_PCT": 100.0}, "position": {"x": 410.70873998343404, "y": 161.12504153904717}, "selected": false}, {"data": {"id": "579227", "Enrichment": 2.9823284627551403, "BestLogPInGroup": -7.5613451204042255, "_PATTERN_": "M1", "_GeneInGOAndHitList": 13.0, "Hits": "ALOX15B|BTG1|SFN|NOTCH1|PRKCH|PTCH1|SULT2B1|PTCH2|ZEB2|MESP1|MIR200C|MIR21|MIR99B", "Description": "positive regulation of epithelial cell differentiation", "_MEMBER_Input_ID": 1.0, "_GeneInHitList": 1562.0, "Category": "GO Biological Processes", "Log_q_value_": -1.7823539286407861, "Symbol": "GO:0030858", "LogP": -3.47511962057552, "_RANK_": 1.0, "FirstInGroupByEnrichment": 0.0, "_TotalGeneInLibrary": 24367.0, "_GeneInGO": 68.0, "shared_name": "30", "_InGO": 0.8322663252240717, "Gene": "GO:0030858", "_COUNT_001": 13.0, "SUID": 579227, "id_original": "30", "selected": false, "CategoryID": 19.0, "STDV_InGO": 0.22986658336957838, "EvidenceCutoff": 0.0, "PARENT_GO": "19_GO:0048518 positive regulation of biological process", "FirstInGroupByLogP": 0.0, "GROUP_ID": 19.0, "name": "30", "BestEnrichmentInGroup": 4.549962654716176, "CLUSTER_LABEL": " ", "GeneID": "247|694|2810|4851|5583|5727|6820|8643|9839|55897|406985|406991|407056", "_LogP_Input_ID": -3.47511962057552, "Z_score": 4.2840395245299625, "GiniIndex": 0.0, "_COUNT_001_PCT": 100.0}, "position": {"x": 495.80017064749654, "y": 137.2006335800628}, "selected": false}, {"data": {"id": "579226", "Enrichment": 4.549962654716176, "BestLogPInGroup": -7.5613451204042255, "_PATTERN_": "M1", "_GeneInGOAndHitList": 7.0, "Hits": "ALOX15B|SFN|NOTCH1|PRKCH|PTCH1|SULT2B1|PTCH2", "Description": "positive regulation of epidermal cell differentiation", "_MEMBER_Input_ID": 1.0, "_GeneInHitList": 1562.0, "Category": "GO Biological Processes", "Log_q_value_": -1.6207242506102761, "Symbol": "GO:0045606", "LogP": -3.2391265927750332, "_RANK_": 1.0, "FirstInGroupByEnrichment": 0.0, "_TotalGeneInLibrary": 24367.0, "_GeneInGO": 24.0, "shared_name": "29", "_InGO": 0.44814340588988477, "Gene": "GO:0045606", "_COUNT_001": 7.0, "SUID": 579226, "id_original": "29", "selected": false, "CategoryID": 19.0, "STDV_InGO": 0.169002322292619, "EvidenceCutoff": 0.0, "PARENT_GO": "19_GO:0048518 positive regulation of biological process", "FirstInGroupByLogP": 0.0, "GROUP_ID": 19.0, "name": "29", "BestEnrichmentInGroup": 4.549962654716176, "CLUSTER_LABEL": " ", "GeneID": "247|2810|4851|5583|5727|6820|8643", "_LogP_Input_ID": -3.2391265927750332, "Z_score": 4.553653197987221, "GiniIndex": 0.0, "_COUNT_001_PCT": 100.0}, "position": {"x": 511.06030248343404, "y": 215.02939944920342}, "selected": false}, {"data": {"id": "579225", "Enrichment": 3.424362137347366, "BestLogPInGroup": -7.5613451204042255, "_PATTERN_": "M1", "_GeneInGOAndHitList": 9.0, "Hits": "ALOX15B|ZFP36L1|RUNX1|CBFB|HOXA7|NOTCH1|PRKCH|ZFP36|GRHL2", "Description": "regulation of keratinocyte differentiation", "_MEMBER_Input_ID": 1.0, "_GeneInHitList": 1562.0, "Category": "GO Biological Processes", "Log_q_value_": -1.4594334981148884, "Symbol": "GO:0045616", "LogP": -3.0158609936972285, "_RANK_": 1.0, "FirstInGroupByEnrichment": 0.0, "_TotalGeneInLibrary": 24367.0, "_GeneInGO": 41.0, "shared_name": "28", "_InGO": 0.5761843790012804, "Gene": "GO:0045616", "_COUNT_001": 9.0, "SUID": 579225, "id_original": "28", "selected": false, "CategoryID": 19.0, "STDV_InGO": 0.19150734627097452, "EvidenceCutoff": 0.0, "PARENT_GO": "19_GO:0032502 developmental process", "FirstInGroupByLogP": 0.0, "GROUP_ID": 19.0, "name": "28", "BestEnrichmentInGroup": 4.549962654716176, "CLUSTER_LABEL": " ", "GeneID": "247|677|861|865|3204|4851|5583|7538|79977", "_LogP_Input_ID": -3.0158609936972285, "Z_score": 4.066040183601785, "GiniIndex": 0.0, "_COUNT_001_PCT": 100.0}, "position": {"x": 509.0646359795278, "y": 92.42682223973077}, "selected": false}, {"data": {"id": "579224", "Enrichment": 3.1999737351850026, "BestLogPInGroup": -7.5613451204042255, "_PATTERN_": "M1", "_GeneInGOAndHitList": 8.0, "Hits": "ALOX15B|SFN|NOTCH1|PRKCH|PTCH1|SULT2B1|PTCH2|TMEM79", "Description": "positive regulation of epidermis development", "_MEMBER_Input_ID": 1.0, "_GeneInHitList": 1562.0, "Category": "GO Biological Processes", "Log_q_value_": -1.1262807051997246, "Symbol": "GO:0045684", "LogP": -2.5395908841416324, "_RANK_": 1.0, "FirstInGroupByEnrichment": 0.0, "_TotalGeneInLibrary": 24367.0, "_GeneInGO": 39.0, "shared_name": "27", "_InGO": 0.5121638924455826, "Gene": "GO:0045684", "_COUNT_001": 8.0, "SUID": 579224, "id_original": "27", "selected": false, "CategoryID": 19.0, "STDV_InGO": 0.18061297922967348, "EvidenceCutoff": 0.0, "PARENT_GO": "19_GO:0048518 positive regulation of biological process", "FirstInGroupByLogP": 0.0, "GROUP_ID": 19.0, "name": "27", "BestEnrichmentInGroup": 4.549962654716176, "CLUSTER_LABEL": " ", "GeneID": "247|2810|4851|5583|5727|6820|8643|84283", "_LogP_Input_ID": -2.5395908841416324, "Z_score": 3.5984393918577124, "GiniIndex": 0.0, "_COUNT_001_PCT": 100.0}, "position": {"x": 432.76818822562154, "y": 94.93265109715264}, "selected": false}, {"data": {"id": "579223", "Enrichment": 2.4484103971566866, "BestLogPInGroup": -7.519436038415945, "_PATTERN_": "M1", "_GeneInGOAndHitList": 35.0, "Hits": "AKT1|CCND1|CALM1|RUNX1|CBFB|CDKN1B|CHD1|DDX5|EGFR|FKBP5|FOXO3|FOS|GNB3|H2BC5|H3-3B|FOXA1|HSPB1|HSP90AB1|IGF1R|JUND|POLR2A|SHC1|SP1|SRF|TLE3|NRIP1|H2BC9|H2BC10|H3C8|PIK3R3|H3C7|GREB1|PTGES3|KDM4B|KANK1", "Description": "ESR-mediated signaling", "_MEMBER_Input_ID": 1.0, "_GeneInHitList": 1562.0, "Category": "Reactome Gene Sets", "Log_q_value_": -3.6689989857093113, "Symbol": "R-HSA-8939211", "LogP": -6.090477953728646, "_RANK_": 1.0, "FirstInGroupByEnrichment": 0.0, "_TotalGeneInLibrary": 24367.0, "_GeneInGO": 223.0, "shared_name": "26", "_InGO": 2.240717029449424, "Gene": "R-HSA-8939211", "_COUNT_001": 35.0, "SUID": 579223, "id_original": "26", "selected": false, "CategoryID": 6.0, "STDV_InGO": 0.3744829043781526, "EvidenceCutoff": 0.0, "PARENT_GO": "", "FirstInGroupByLogP": 0.0, "GROUP_ID": 20.0, "name": "26", "BestEnrichmentInGroup": 2.4484103971566866, "CLUSTER_LABEL": " ", "GeneID": "207|595|801|861|865|1027|1105|1655|1956|2289|2309|2353|2784|3017|3021|3169|3315|3326|3480|3727|5430|6464|6667|6722|7090|8204|8345|8346|8355|8503|8968|9687|10728|23030|23189", "_LogP_Input_ID": -6.090477953728646, "Z_score": 5.686657216414608, "GiniIndex": 0.0, "_COUNT_001_PCT": 100.0}, "position": {"x": 329.666497456426, "y": 829.3525958603362}, "selected": false}, {"data": {"id": "579222", "Enrichment": 2.247439180537772, "BestLogPInGroup": -7.519436038415945, "_PATTERN_": "M1", "_GeneInGOAndHitList": 34.0, "Hits": "ACTA2|AKT1|ATP2A2|CCND1|RUNX1|EGFR|ELF3|B4GALT1|H2BC5|H3-3B|LFNG|SMAD3|NOTCH1|NOTCH4|PBX1|PSMB3|PSMB8|PSMB9|PSMD1|PSMD2|RFNG|TLE1|TLE3|UBC|YWHAZ|H2BC9|H2BC10|H3C8|H3C7|NCOR2|WWC1|HEY2|HDAC7|MAML3", "Description": "Signaling by NOTCH", "_MEMBER_Input_ID": 1.0, "_GeneInHitList": 1562.0, "Category": "Reactome Gene Sets", "Log_q_value_": -2.9604580736572177, "Symbol": "R-HSA-157118", "LogP": -5.093842225734982, "_RANK_": 1.0, "FirstInGroupByEnrichment": 0.0, "_TotalGeneInLibrary": 24367.0, "_GeneInGO": 236.0, "shared_name": "25", "_InGO": 2.176696542893726, "Gene": "R-HSA-157118", "_COUNT_001": 34.0, "SUID": 579222, "id_original": "25", "selected": false, "CategoryID": 6.0, "STDV_InGO": 0.3692152164903899, "EvidenceCutoff": 0.0, "PARENT_GO": "", "FirstInGroupByLogP": 0.0, "GROUP_ID": 20.0, "name": "25", "BestEnrichmentInGroup": 2.4484103971566866, "CLUSTER_LABEL": " ", "GeneID": "59|207|488|595|861|1956|1999|2683|3017|3021|3955|4088|4851|4855|5087|5691|5696|5698|5707|5708|5986|7088|7090|7316|7534|8345|8346|8355|8968|9612|23286|23493|51564|55534", "_LogP_Input_ID": -5.093842225734982, "Z_score": 5.039707355445641, "GiniIndex": 0.0, "_COUNT_001_PCT": 100.0}, "position": {"x": 549.7948305481618, "y": 911.7290912216644}, "selected": false}, {"data": {"id": "579221", "Enrichment": 2.39198036705079, "BestLogPInGroup": -7.519436038415945, "_PATTERN_": "M1", "_GeneInGOAndHitList": 23.0, "Hits": "CCND1|RUNX1|CBFB|CHD1|DDX5|FOS|H2BC5|H3-3B|FOXA1|HSP90AB1|JUND|POLR2A|SP1|TLE3|NRIP1|H2BC9|H2BC10|H3C8|H3C7|GREB1|PTGES3|KDM4B|KANK1", "Description": "Estrogen-dependent gene expression", "_MEMBER_Input_ID": 1.0, "_GeneInHitList": 1562.0, "Category": "Reactome Gene Sets", "Log_q_value_": -2.2160275766145303, "Symbol": "R-HSA-9018519", "LogP": -4.066786305512149, "_RANK_": 1.0, "FirstInGroupByEnrichment": 0.0, "_TotalGeneInLibrary": 24367.0, "_GeneInGO": 150.0, "shared_name": "24", "_InGO": 1.47247119078105, "Gene": "R-HSA-9018519", "_COUNT_001": 23.0, "SUID": 579221, "id_original": "24", "selected": false, "CategoryID": 6.0, "STDV_InGO": 0.30476260956438245, "EvidenceCutoff": 0.0, "PARENT_GO": "", "FirstInGroupByLogP": 0.0, "GROUP_ID": 20.0, "name": "24", "BestEnrichmentInGroup": 2.4484103971566866, "CLUSTER_LABEL": " ", "GeneID": "595|861|865|1105|1655|2353|3017|3021|3169|3326|3727|5430|6667|7090|8204|8345|8346|8355|8968|9687|10728|23030|23189", "_LogP_Input_ID": -4.066786305512149, "Z_score": 4.475444388031784, "GiniIndex": 0.0, "_COUNT_001_PCT": 100.0}, "position": {"x": 395.8407223099416, "y": 734.3594928329925}, "selected": false}, {"data": {"id": "579220", "Enrichment": 2.2898894618755063, "BestLogPInGroup": -7.519436038415945, "_PATTERN_": "M1", "_GeneInGOAndHitList": 16.0, "Hits": "ATP2A2|CCND1|RUNX1|ELF3|B4GALT1|H2BC5|H3-3B|LFNG|NOTCH1|NOTCH4|RFNG|H2BC9|H2BC10|H3C8|H3C7|MAML3", "Description": "Pre-NOTCH Expression and Processing", "_MEMBER_Input_ID": 1.0, "_GeneInHitList": 1562.0, "Category": "Reactome Gene Sets", "Log_q_value_": -1.302883418246244, "Symbol": "R-HSA-1912422", "LogP": -2.8064141675908596, "_RANK_": 1.0, "FirstInGroupByEnrichment": 0.0, "_TotalGeneInLibrary": 24367.0, "_GeneInGO": 109.0, "shared_name": "23", "_InGO": 1.0243277848911652, "Gene": "R-HSA-1912422", "_COUNT_001": 16.0, "SUID": 579220, "id_original": "23", "selected": false, "CategoryID": 6.0, "STDV_InGO": 0.25476701097987803, "EvidenceCutoff": 0.0, "PARENT_GO": "", "FirstInGroupByLogP": 0.0, "GROUP_ID": 20.0, "name": "23", "BestEnrichmentInGroup": 2.4484103971566866, "CLUSTER_LABEL": " ", "GeneID": "488|595|861|1999|2683|3017|3021|3955|4851|4855|5986|8345|8346|8355|8968|55534", "_LogP_Input_ID": -2.8064141675908596, "Z_score": 3.532285871360611, "GiniIndex": 0.0, "_COUNT_001_PCT": 100.0}, "position": {"x": 462.64605067908224, "y": 834.964595372055}, "selected": false}, {"data": {"id": "579219", "Enrichment": 2.191717547962455, "BestLogPInGroup": -7.519436038415945, "_PATTERN_": "M1", "_GeneInGOAndHitList": 17.0, "Hits": "H2BC5|H3-3B|HOXA3|HOXC4|MEIS1|PBX1|POLR2A|RARA|RARB|RARG|RXRA|PCGF2|H2BC9|H2BC10|H3C8|H3C7|NCOA6", "Description": "Activation of anterior HOX genes in hindbrain development during early embryogenesis", "_MEMBER_Input_ID": 1.0, "_GeneInHitList": 1562.0, "Category": "Reactome Gene Sets", "Log_q_value_": -1.2551740957821733, "Symbol": "R-HSA-5617472", "LogP": -2.734174060635192, "_RANK_": 1.0, "FirstInGroupByEnrichment": 0.0, "_TotalGeneInLibrary": 24367.0, "_GeneInGO": 121.0, "shared_name": "22", "_InGO": 1.088348271446863, "Gene": "R-HSA-5617472", "_COUNT_001": 17.0, "SUID": 579219, "id_original": "22", "selected": false, "CategoryID": 6.0, "STDV_InGO": 0.2625228789306787, "EvidenceCutoff": 0.0, "PARENT_GO": "", "FirstInGroupByLogP": 0.0, "GROUP_ID": 20.0, "name": "22", "BestEnrichmentInGroup": 2.4484103971566866, "CLUSTER_LABEL": " ", "GeneID": "3017|3021|3200|3221|4211|5087|5430|5914|5915|5916|6256|7703|8345|8346|8355|8968|23054", "_LogP_Input_ID": -2.734174060635192, "Z_score": 3.439248571086318, "GiniIndex": 0.0, "_COUNT_001_PCT": 100.0}, "position": {"x": 464.46904872595724, "y": 727.6708026474456}, "selected": false}, {"data": {"id": "579218", "Enrichment": 2.191717547962455, "BestLogPInGroup": -7.519436038415945, "_PATTERN_": "M1", "_GeneInGOAndHitList": 17.0, "Hits": "H2BC5|H3-3B|HOXA3|HOXC4|MEIS1|PBX1|POLR2A|RARA|RARB|RARG|RXRA|PCGF2|H2BC9|H2BC10|H3C8|H3C7|NCOA6", "Description": "Activation of HOX genes during differentiation", "_MEMBER_Input_ID": 1.0, "_GeneInHitList": 1562.0, "Category": "Reactome Gene Sets", "Log_q_value_": -1.2551740957821733, "Symbol": "R-HSA-5619507", "LogP": -2.734174060635192, "_RANK_": 1.0, "FirstInGroupByEnrichment": 0.0, "_TotalGeneInLibrary": 24367.0, "_GeneInGO": 121.0, "shared_name": "21", "_InGO": 1.088348271446863, "Gene": "R-HSA-5619507", "_COUNT_001": 17.0, "SUID": 579218, "id_original": "21", "selected": false, "CategoryID": 6.0, "STDV_InGO": 0.2625228789306787, "EvidenceCutoff": 0.0, "PARENT_GO": "", "FirstInGroupByLogP": 0.0, "GROUP_ID": 20.0, "name": "21", "BestEnrichmentInGroup": 2.4484103971566866, "CLUSTER_LABEL": " ", "GeneID": "3017|3021|3200|3221|4211|5087|5430|5914|5915|5916|6256|7703|8345|8346|8355|8968|23054", "_LogP_Input_ID": -2.734174060635192, "Z_score": 3.439248571086318, "GiniIndex": 0.0, "_COUNT_001_PCT": 100.0}, "position": {"x": 381.93364833533224, "y": 813.8188129013519}, "selected": false}, {"data": {"id": "579217", "Enrichment": 2.3999803013887524, "BestLogPInGroup": -7.519436038415945, "_PATTERN_": "M1", "_GeneInGOAndHitList": 46.0, "Hits": "AKT1|ALDH1A3|CCND1|CALM1|RUNX1|CBFB|CDKN1B|CHD1|DDX5|EGFR|FASN|FKBP5|FOXO3|FOS|GNB3|H2BC5|H3-3B|FOXA1|HSPB1|HSP90AB1|IGF1R|JUND|POLR2A|RARA|RARB|RARG|RDH5|RXRA|SHC1|SP1|SREBF1|SRF|TLE3|NRIP1|H2BC9|H2BC10|H3C8|PIK3R3|H3C7|DHRS3|NCOR2|GREB1|PTGES3|KDM4B|KANK1|RDH11", "Description": "Signaling by Nuclear Receptors", "_MEMBER_Input_ID": 1.0, "_GeneInHitList": 1562.0, "Category": "Reactome Gene Sets", "Log_q_value_": -4.756632479873455, "Symbol": "R-HSA-9006931", "LogP": -7.519436038415945, "_RANK_": 1.0, "FirstInGroupByEnrichment": 1.0, "_TotalGeneInLibrary": 24367.0, "_GeneInGO": 299.0, "shared_name": "20", "_InGO": 2.9449423815621, "Gene": "R-HSA-9006931", "_COUNT_001": 46.0, "SUID": 579217, "id_original": "20", "selected": false, "CategoryID": 6.0, "STDV_InGO": 0.4277666988065664, "EvidenceCutoff": 0.0, "PARENT_GO": "", "FirstInGroupByLogP": 1.0, "GROUP_ID": 20.0, "name": "20", "BestEnrichmentInGroup": 2.4484103971566866, "CLUSTER_LABEL": "Signaling by Nuclear Receptors", "GeneID": "207|220|595|801|861|865|1027|1105|1655|1956|2194|2289|2309|2353|2784|3017|3021|3169|3315|3326|3480|3727|5430|5914|5915|5916|5959|6256|6464|6667|6720|6722|7090|8204|8345|8346|8355|8503|8968|9249|9612|9687|10728|23030|23189|51109", "_LogP_Input_ID": -7.519436038415945, "Z_score": 6.374631935022215, "GiniIndex": 0.0, "_COUNT_001_PCT": 100.0}, "position": {"x": 323.0395290726369, "y": 716.9538837021331}, "selected": false}, {"data": {"id": "579216", "Enrichment": 3.119974391805378, "BestLogPInGroup": -9.062335119268083, "_PATTERN_": "M1", "_GeneInGOAndHitList": 36.0, "Hits": "BCL2L1|BCL6|RUNX2|RUNX1|CCND2|CDKN1B|CEBPB|CCR7|DDX5|ETV6|H3-3B|HOXA9|HOXA10|HOXA11|ID2|IGF1R|ITGB7|JUP|MEIS1|PBX1|PBX3|CDK14|RARA|RXRA|SP1|TMPRSS2|TRAF1|ZBTB16|H3C8|PER2|H3C7|KLF3|FEV|ASPSCR1|SLC45A3|JMJD1C", "Description": "Transcriptional misregulation in cancer", "_MEMBER_Input_ID": 1.0, "_GeneInHitList": 1562.0, "Category": "KEGG Pathway", "Log_q_value_": -5.916082775157079, "Symbol": "hsa05202", "LogP": -9.062335119268083, "_RANK_": 1.0, "FirstInGroupByEnrichment": 1.0, "_TotalGeneInLibrary": 24367.0, "_GeneInGO": 180.0, "shared_name": "19", "_InGO": 2.3047375160051216, "Gene": "hsa05202", "_COUNT_001": 36.0, "SUID": 579216, "id_original": "19", "selected": false, "CategoryID": 24.0, "STDV_InGO": 0.37967060373244593, "EvidenceCutoff": 0.0, "PARENT_GO": "", "FirstInGroupByLogP": 1.0, "GROUP_ID": 10.0, "name": "19", "BestEnrichmentInGroup": 3.119974391805378, "CLUSTER_LABEL": "Transcriptional misregulation in cancer", "GeneID": "598|604|860|861|894|1027|1051|1236|1655|2120|3021|3205|3206|3207|3398|3480|3695|3728|4211|5087|5090|5218|5914|6256|6667|7113|7185|7704|8355|8864|8968|51274|54738|79058|85414|221037", "_LogP_Input_ID": -9.062335119268083, "Z_score": 7.471253315628116, "GiniIndex": 0.0, "_COUNT_001_PCT": 100.0}, "position": {"x": -487.60092188180033, "y": 1011.7290912216644}, "selected": false}, {"data": {"id": "579215", "Enrichment": 2.0478596394089355, "BestLogPInGroup": -11.458053982125241, "_PATTERN_": "M1", "_GeneInGOAndHitList": 102.0, "Hits": "ABL1|ACTA2|AKT1|ANPEP|ANXA2|KLK3|ZFP36L1|BTG1|RUNX1|COL1A1|COL4A1|COL4A2|TYMP|EPHA2|EPHB4|ERBB2|FOXF1|GATA2|B4GALT1|RAPGEF1|NR4A1|HOXA3|HOXA5|HOXA7|HSPB1|HSPG2|ID1|ISL1|JUNB|JUP|LRP1|SMAD7|MEIS1|MMP14|MYH9|NFE2L2|NKX3-1|NOTCH1|NOTCH4|PDGFB|SERPINF1|PKM|SERPINF2|PTK7|PTPRB|PTPRM|RRAS|CX3CL1|SHB|SHC1|SOX4|SP1|SPARC|SRF|TBX3|TJP1|XBP1|YWHAZ|FOSL1|PIK3R3|CFLAR|HGS|QKI|SNX17|RAMP1|HOXB13|GNA13|JMJD6|HEY2|TSPAN12|TIPARP|TNFRSF12A|HDAC7|RIN2|PARVA|MESP1|SPHK2|ADAMTS9|SEMA6A|NIBAN2|MMRN2|ZC3H12A|COL18A1|APOLD1|ADTRP|CREB3L1|PLCD3|DAB2IP|MIA3|MIRLET7G|MIR106B|MIR125A|MIR125B1|MIR132|MIR21|MIR212|MIR27B|MIR29A|MIR29B1|MIR30B|MIR99B|MIR375", "Description": "blood vessel development", "_MEMBER_Input_ID": 1.0, "_GeneInHitList": 1562.0, "Category": "GO Biological Processes", "Log_q_value_": -7.6160691775351275, "Symbol": "GO:0001568", "LogP": -11.458053982125241, "_RANK_": 1.0, "FirstInGroupByEnrichment": 1.0, "_TotalGeneInLibrary": 24367.0, "_GeneInGO": 777.0, "shared_name": "18", "_InGO": 6.530089628681178, "Gene": "GO:0001568", "_COUNT_001": 102.0, "SUID": 579215, "id_original": "18", "selected": false, "CategoryID": 19.0, "STDV_InGO": 0.6251078740756986, "EvidenceCutoff": 0.0, "PARENT_GO": "19_GO:0032502 developmental process", "FirstInGroupByLogP": 1.0, "GROUP_ID": 2.0, "name": "18", "BestEnrichmentInGroup": 2.262576849019167, "CLUSTER_LABEL": "blood vessel development", "GeneID": "25|59|207|290|302|354|677|694|861|1277|1282|1284|1890|1969|2050|2064|2294|2624|2683|2889|3164|3200|3202|3204|3315|3339|3397|3670|3726|3728|4035|4092|4211|4323|4627|4780|4824|4851|4855|5155|5176|5315|5345|5754|5787|5797|6237|6376|6461|6464|6659|6667|6678|6722|6926|7082|7494|7534|8061|8503|8837|9146|9444|9784|10267|10481|10672|23210|23493|23554|25976|51330|51564|54453|55742|55897|56848|56999|57556|64855|79812|80149|80781|81575|84830|90993|113026|153090|375056|406890|406900|406910|406911|406921|406991|406994|407019|407021|407024|407030|407056|494324", "_LogP_Input_ID": -11.458053982125241, "Z_score": 7.769038046177723, "GiniIndex": 0.0, "_COUNT_001_PCT": 100.0}, "position": {"x": 294.93850682913717, "y": 1.964778477523737}, "selected": false}, {"data": {"id": "579214", "Enrichment": 2.778924559502765, "BestLogPInGroup": -9.213753097063153, "_PATTERN_": "M1", "_GeneInGOAndHitList": 44.0, "Hits": "ABL1|ACTB|ACTN4|ACTN1|DST|RUNX1|CBFB|CD9|CLDN3|EPHA2|RAPGEF1|ILK|JUP|LAMA3|LIMS1|LRP1|SMAD3|SMAD7|MMP14|CLDN11|PLEC|PRKCH|PTPRJ|SDC4|SRF|TBCD|TJP1|TLN1|TNS1|TRIP6|CNTNAP1|CLDN12|DLG5|FLOT1|SORBS1|LIMCH1|MACF1|CORO1C|HDAC7|CAMSAP3|CDH23|MICALL2|GRHL2|VMP1", "Description": "cell junction assembly", "_MEMBER_Input_ID": 1.0, "_GeneInHitList": 1562.0, "Category": "GO Biological Processes", "Log_q_value_": -6.008590390060213, "Symbol": "GO:0034329", "LogP": -9.213753097063153, "_RANK_": 1.0, "FirstInGroupByEnrichment": 1.0, "_TotalGeneInLibrary": 24367.0, "_GeneInGO": 247.0, "shared_name": "17", "_InGO": 2.816901408450704, "Gene": "GO:0034329", "_COUNT_001": 44.0, "SUID": 579214, "id_original": "17", "selected": false, "CategoryID": 19.0, "STDV_InGO": 0.4186399594939643, "EvidenceCutoff": 0.0, "PARENT_GO": "19_GO:0071840 cellular component organization or biogenesis", "FirstInGroupByLogP": 1.0, "GROUP_ID": 9.0, "name": "17", "BestEnrichmentInGroup": 4.120720894837292, "CLUSTER_LABEL": "cell junction assembly", "GeneID": "25|60|81|87|667|861|865|928|1365|1969|2889|3611|3728|3909|3987|4035|4088|4092|4323|5010|5339|5583|5795|6385|6722|6904|7082|7094|7145|7205|8506|9069|9231|10211|10580|22998|23499|23603|51564|57662|64072|79778|79977|81671", "_LogP_Input_ID": -9.213753097063153, "Z_score": 7.354191800391712, "GiniIndex": 0.0, "_COUNT_001_PCT": 100.0}, "position": {"x": 33.885574089879356, "y": -91.42410702052314}, "selected": false}, {"data": {"id": "579213", "Enrichment": 2.631303703932246, "BestLogPInGroup": -8.096413799222795, "_PATTERN_": "M1", "_GeneInGOAndHitList": 42.0, "Hits": "ATP1A1|ZFP36L1|ZFP36L2|KLF9|RUNX1|CBFB|DDX5|EGFR|FHL2|FOXO3|NR4A1|FOXA1|HNRNPU|ISL1|NKX3-1|NR4A2|SERPINF1|PER1|RARA|RARB|RARG|RB1|RORC|RXRA|THRA|THRB|ZFP36|NRIP1|CFLAR|NR1D1|NCOR2|DDX17|ABHD2|PHB2|CBX3|WBP2|FOXP1|ZBTB7A|PMEPA1|OR51E2|ADTRP|EGLN2", "Description": "cellular response to steroid hormone stimulus", "_MEMBER_Input_ID": 1.0, "_GeneInHitList": 1562.0, "Category": "GO Biological Processes", "Log_q_value_": -5.208671504072005, "Symbol": "GO:0071383", "LogP": -8.096413799222795, "_RANK_": 1.0, "FirstInGroupByEnrichment": 1.0, "_TotalGeneInLibrary": 24367.0, "_GeneInGO": 249.0, "shared_name": "16", "_InGO": 2.6888604353393086, "Gene": "GO:0071383", "_COUNT_001": 42.0, "SUID": 579213, "id_original": "16", "selected": false, "CategoryID": 19.0, "STDV_InGO": 0.40928412025975003, "EvidenceCutoff": 0.0, "PARENT_GO": "19_GO:0050896 response to stimulus", "FirstInGroupByLogP": 1.0, "GROUP_ID": 15.0, "name": "16", "BestEnrichmentInGroup": 2.631303703932246, "CLUSTER_LABEL": "cellular response to steroid hormone stimulus", "GeneID": "476|677|678|687|861|865|1655|1956|2274|2309|3164|3169|3192|3670|4824|4929|5176|5187|5914|5915|5916|5925|6097|6256|7067|7068|7538|8204|8837|9572|9612|10521|11057|11331|11335|23558|27086|51341|56937|81285|84830|112398", "_LogP_Input_ID": -8.096413799222795, "Z_score": 6.771446426603564, "GiniIndex": 0.0, "_COUNT_001_PCT": 100.0}, "position": {"x": 140.99559138953146, "y": 865.979991490219}, "selected": false}, {"data": {"id": "579212", "Enrichment": 2.197505183082735, "BestLogPInGroup": -11.981654863057864, "_PATTERN_": "M1", "_GeneInGOAndHitList": 91.0, "Hits": "ABL1|AKT1|BCL2L1|ZFP36L1|RUNX2|CEBPB|CFL1|COL1A1|DLX1|DLX2|DVL3|ECE1|EGFR|EPHA2|ELF3|FOXF1|GATA2|FOXA1|HOXA3|HOXA5|HOXA6|HOXA7|HOXA9|HOXA11|HOXC4|HOXC6|HOXD10|ISL1|ITPK1|JUNB|KRT8|LFNG|SMAD3|MBNL1|MGAT1|MMP14|MYH9|NKX3-1|NOTCH1|ROR2|PBX1|PDGFB|PTCH1|PTK7|PTPRR|RARA|RARG|RXRA|SDC4|STIL|SOX4|SP3|SRF|TBX3|HNF1B|TRIP6|PCGF2|ZFAND5|FOSL1|YBX3|DCHS1|SH3PXD2A|ZEB2|SEC24D|SCO2|LPAR6|PEMT|EMG1|GNA13|COBL|HEY2|MTHFD1L|SLC39A1|SEMA4C|MESP1|XAB2|RTN4|SHROOM3|CAMSAP3|NXN|GRHL2|CMIP|EGFL8|HES7|UNK|PLCD3|MESP2|RUNDC1|ARL13B|SMIM14|ST8SIA6", "Description": "chordate embryonic development", "_MEMBER_Input_ID": 1.0, "_GeneInHitList": 1562.0, "Category": "GO Biological Processes", "Log_q_value_": -7.675226902393069, "Symbol": "GO:0043009", "LogP": -11.981654863057864, "_RANK_": 1.0, "FirstInGroupByEnrichment": 1.0, "_TotalGeneInLibrary": 24367.0, "_GeneInGO": 646.0, "shared_name": "15", "_InGO": 5.825864276568502, "Gene": "GO:0043009", "_COUNT_001": 91.0, "SUID": 579212, "id_original": "15", "selected": false, "CategoryID": 19.0, "STDV_InGO": 0.5926598246801607, "EvidenceCutoff": 0.0, "PARENT_GO": "19_GO:0032502 developmental process", "FirstInGroupByLogP": 1.0, "GROUP_ID": 1.0, "name": "15", "BestEnrichmentInGroup": 2.5999786598378147, "CLUSTER_LABEL": "chordate embryonic development", "GeneID": "25|207|598|677|860|1051|1072|1277|1745|1746|1857|1889|1956|1969|1999|2294|2624|3169|3200|3202|3203|3204|3205|3207|3221|3223|3236|3670|3705|3726|3856|3955|4088|4154|4245|4323|4627|4824|4851|4920|5087|5155|5727|5754|5801|5914|5916|6256|6385|6491|6659|6670|6722|6926|6928|7205|7703|7763|8061|8531|8642|9644|9839|9871|9997|10161|10400|10436|10672|23242|23493|25902|27173|54910|55897|56949|57142|57619|57662|64359|79977|80790|80864|84667|85451|113026|145873|146923|200894|201895|338596", "_LogP_Input_ID": -11.981654863057864, "Z_score": 8.07318232058665, "GiniIndex": 0.0, "_COUNT_001_PCT": 100.0}, "position": {"x": 989.2212979668325, "y": 94.76131778416436}, "selected": false}, {"data": {"id": "579211", "Enrichment": 2.46904448272368, "BestLogPInGroup": -7.592008542189207, "_PATTERN_": "M1", "_GeneInGOAndHitList": 44.0, "Hits": "ACTA2|BMPR1B|RUNX2|RUNX1|SERPINH1|COL1A1|DLX2|FOXA1|HOXA3|HOXA5|HOXA11|HOXC4|ID2|LTBP3|SMAD3|SMAD7|MEF2D|NOTCH1|ROR2|OXCT1|PDGFB|RARA|RARB|RARG|RORC|SRF|THRA|TRPS1|XBP1|ZBTB16|KLF7|CFLAR|CCN4|ZNF516|ZMPSTE24|SPART|TRPM4|ARRDC3|VWA1|WNT5B|ARID5B|SH3PXD2B|MIR125A|MIR21", "Description": "connective tissue development", "_MEMBER_Input_ID": 1.0, "_GeneInHitList": 1562.0, "Category": "GO Biological Processes", "Log_q_value_": -4.804381399921686, "Symbol": "GO:0061448", "LogP": -7.592008542189207, "_RANK_": 1.0, "FirstInGroupByEnrichment": 1.0, "_TotalGeneInLibrary": 24367.0, "_GeneInGO": 278.0, "shared_name": "14", "_InGO": 2.816901408450704, "Gene": "GO:0061448", "_COUNT_001": 44.0, "SUID": 579211, "id_original": "14", "selected": false, "CategoryID": 19.0, "STDV_InGO": 0.4186399594939643, "EvidenceCutoff": 0.0, "PARENT_GO": "19_GO:0032502 developmental process", "FirstInGroupByLogP": 1.0, "GROUP_ID": 18.0, "name": "14", "BestEnrichmentInGroup": 4.679961587708067, "CLUSTER_LABEL": "connective tissue development", "GeneID": "59|658|860|861|871|1277|1746|3169|3200|3202|3207|3221|3398|4054|4088|4092|4209|4851|4920|5019|5155|5914|5915|5916|6097|6722|7067|7227|7494|7704|8609|8837|8840|9658|10269|23111|54795|57561|64856|81029|84159|285590|406910|406991", "_LogP_Input_ID": -7.592008542189207, "Z_score": 6.4471170490313865, "GiniIndex": 0.0, "_COUNT_001_PCT": 100.0}, "position": {"x": 1268.745849358434, "y": 137.17045169529717}, "selected": false}, {"data": {"id": "579210", "Enrichment": 1.9847165342273396, "BestLogPInGroup": -10.448191879197235, "_PATTERN_": "M1", "_GeneInGOAndHitList": 100.0, "Hits": "ABL1|ACTA2|AKT1|ALOX15B|ANXA4|CCND1|ZFP36L1|BTG1|RUNX1|CBFB|CDKN1B|CEBPB|TPP1|CNN3|COL4A1|CLDN3|CTSB|EPHA2|ELF3|FASN|FAT1|FOXF1|FLNB|FOSL2|B4GALT1|SFN|RAPGEF1|FOXA1|HOXA5|HOXA7|ID1|ID2|ID3|JUP|KRT7|KRT8|KRT86|SMAD3|MYO6|NOTCH1|NOTCH4|PDGFB|PRKCH|PRSS8|PTCH1|PTK7|RARA|RARB|RARG|RXRA|SDC1|SOX4|SULT2B1|TAF10|TBX3|HNF1B|THRA|THRB|TJP1|TNFRSF1A|UPK2|EZR|XBP1|ZFP36|AIMP2|KLF7|PTCH2|CBFA2T2|DLG5|ZEB2|FEM1B|HOXB13|AGR2|TRIOBP|PALLD|KDM6B|HEY2|SPDEF|ARHGEF26|EHF|ADAMTSL4|HYDIN|ESRP1|MESP1|SHROOM3|CAMSAP3|PDZD7|GRHL2|SLC44A4|COL18A1|WNT5B|APOLD1|TMEM79|IL31RA|TPRN|MIR200C|MIR21|MIR29B1|MIR99B|CCDC88C", "Description": "epithelial cell differentiation", "_MEMBER_Input_ID": 1.0, "_GeneInHitList": 1562.0, "Category": "GO Biological Processes", "Log_q_value_": -6.841771750932958, "Symbol": "GO:0030855", "LogP": -10.448191879197235, "_RANK_": 1.0, "FirstInGroupByEnrichment": 1.0, "_TotalGeneInLibrary": 24367.0, "_GeneInGO": 786.0, "shared_name": "13", "_InGO": 6.402048655569782, "Gene": "GO:0030855", "_COUNT_001": 100.0, "SUID": 579210, "id_original": "13", "selected": false, "CategoryID": 19.0, "STDV_InGO": 0.6193728182154163, "EvidenceCutoff": 0.0, "PARENT_GO": "19_GO:0032502 developmental process", "FirstInGroupByLogP": 1.0, "GROUP_ID": 4.0, "name": "13", "BestEnrichmentInGroup": 2.907248865091375, "CLUSTER_LABEL": "epithelial cell differentiation", "GeneID": "25|59|207|247|307|595|677|694|861|865|1027|1051|1200|1266|1282|1365|1508|1969|1999|2194|2195|2294|2317|2355|2683|2810|2889|3169|3202|3204|3397|3398|3399|3728|3855|3856|3892|4088|4646|4851|4855|5155|5583|5652|5727|5754|5914|5915|5916|6256|6382|6659|6820|6881|6926|6928|7067|7068|7082|7132|7379|7430|7494|7538|7965|8609|8643|9139|9231|9839|10116|10481|10551|11078|23022|23135|23493|25803|26084|26298|54507|54768|54845|55897|57619|57662|79955|79977|80736|80781|81029|81575|84283|133396|286262|406985|406991|407024|407056|440193", "_LogP_Input_ID": -10.448191879197235, "Z_score": 7.344445127720126, "GiniIndex": 0.0, "_COUNT_001_PCT": 100.0}, "position": {"x": 764.8268124931997, "y": 164.3221393173675}, "selected": false}, {"data": {"id": "579209", "Enrichment": 2.9843233312921, "BestLogPInGroup": -10.25219876997185, "_PATTERN_": "M1", "_GeneInGOAndHitList": 44.0, "Hits": "ADRB1|ADRB2|AKT1|CCND1|ZFP36L1|ZFP36L2|CEBPB|CTBP1|MTOR|GATA2|GNB3|NR4A1|HNRNPU|ID2|INHBB|SMAD3|NR4A2|PSMB8|RORC|RREB1|SREBF1|TRIO|XBP1|ZFP36|ZBTB16|CCN4|PER2|NR1D1|ZNF516|ADIRF|CCDC85B|ZBTB7B|ZBTB7A|TRPM4|STEAP4|ATAT1|ZC3H12A|WNT5B|ARID5B|C1QTNF3|SH3PXD2B|MIR21|MIR27B|MIR29B1", "Description": "fat cell differentiation", "_MEMBER_Input_ID": 1.0, "_GeneInHitList": 1562.0, "Category": "GO Biological Processes", "Log_q_value_": -6.77819075067633, "Symbol": "GO:0045444", "LogP": -10.25219876997185, "_RANK_": 1.0, "FirstInGroupByEnrichment": 1.0, "_TotalGeneInLibrary": 24367.0, "_GeneInGO": 230.0, "shared_name": "12", "_InGO": 2.816901408450704, "Gene": "GO:0045444", "_COUNT_001": 44.0, "SUID": 579209, "id_original": "12", "selected": false, "CategoryID": 19.0, "STDV_InGO": 0.4186399594939643, "EvidenceCutoff": 0.0, "PARENT_GO": "19_GO:0032502 developmental process", "FirstInGroupByLogP": 1.0, "GROUP_ID": 6.0, "name": "12", "BestEnrichmentInGroup": 3.843446714542856, "CLUSTER_LABEL": "fat cell differentiation", "GeneID": "153|154|207|595|677|678|1051|1487|2475|2624|2784|3164|3192|3398|3625|4088|4929|5696|6097|6239|6720|7204|7494|7538|7704|8840|8864|9572|9658|10974|11007|51043|51341|54795|79689|79969|80149|81029|84159|114899|285590|406991|407019|407024", "_LogP_Input_ID": -10.25219876997185, "Z_score": 7.913203018304444, "GiniIndex": 0.0, "_COUNT_001_PCT": 100.0}, "position": {"x": 690.3884718292898, "y": 816.8079486435394}, "selected": false}, {"data": {"id": "579208", "Enrichment": 2.399980301388752, "BestLogPInGroup": -10.799037371088092, "_PATTERN_": "M1", "_GeneInGOAndHitList": 68.0, "Hits": "ABL1|AKT1|ALDH1A3|ALOX15B|CCND1|CDKN1B|CEBPB|DBP|EGFR|EPHA2|ELF3|SLC29A1|FASN|FOXF1|GATA2|FOXA1|HOXA3|HOXA5|HOXA9|HOXD13|HPN|ID2|ISL1|SMAD3|NKX3-1|NOTCH1|NOTCH4|PBX1|SERPINF1|PKM|PTCH1|RARA|RARB|RARG|REG1A|RREB1|RXRA|SP3|SRF|STAT6|STK11|TAF10|TBX3|HNF1B|PRDX2|THRA|THRB|EZR|XBP1|IRS2|SOCS2|CFLAR|NCOR2|RB1CC1|FEM1B|ZMPSTE24|CITED2|HOXB13|PHB2|ZBTB1|COBL|ZBTB7B|CCDC40|MESP1|RTN4|ESRP2|ARID5B|PYGO2", "Description": "gland development", "_MEMBER_Input_ID": 1.0, "_GeneInHitList": 1562.0, "Category": "GO Biological Processes", "Log_q_value_": -7.081991303106278, "Symbol": "GO:0048732", "LogP": -10.799037371088092, "_RANK_": 1.0, "FirstInGroupByEnrichment": 1.0, "_TotalGeneInLibrary": 24367.0, "_GeneInGO": 442.0, "shared_name": "11", "_InGO": 4.353393085787452, "Gene": "GO:0048732", "_COUNT_001": 68.0, "SUID": 579208, "id_original": "11", "selected": false, "CategoryID": 19.0, "STDV_InGO": 0.5163072349210871, "EvidenceCutoff": 0.0, "PARENT_GO": "19_GO:0032502 developmental process", "FirstInGroupByLogP": 1.0, "GROUP_ID": 3.0, "name": "11", "BestEnrichmentInGroup": 2.693503072062197, "CLUSTER_LABEL": "gland development", "GeneID": "25|207|220|247|595|1027|1051|1628|1956|1969|1999|2030|2194|2294|2624|3169|3200|3202|3205|3239|3249|3398|3670|4088|4824|4851|4855|5087|5176|5315|5727|5914|5915|5916|5967|6239|6256|6670|6722|6778|6794|6881|6926|6928|7001|7067|7068|7430|7494|8660|8835|8837|9612|9821|10116|10269|10370|10481|11331|22890|23242|51043|55036|55897|57142|80004|84159|90780", "_LogP_Input_ID": -10.799037371088092, "Z_score": 7.773644624516887, "GiniIndex": 0.0, "_COUNT_001_PCT": 100.0}, "position": {"x": 895.5674435967153, "y": 181.41636233982842}, "selected": false}, {"data": {"id": "579207", "Enrichment": 2.2413609136532884, "BestLogPInGroup": -10.38272655985909, "_PATTERN_": "M1", "_GeneInGOAndHitList": 75.0, "Hits": "ABL1|AGER|ANXA2|B2M|BCL6|ZFP36L1|ZFP36L2|RUNX2|RUNX1|CBFB|CD4|CEBPB|CCR7|KLF6|EPHA2|ERBB2|FASN|FOS|MTOR|GATA2|HMGB1|HOXA7|ID2|IFI16|IL4R|IRF1|ITPKB|JUNB|LFNG|LTBR|SMAD7|MMP14|MYH9|ROR2|POU2F2|PTPN6|PTPRJ|RARA|RB1|RORC|RRAS|SOX4|SP3|SRF|STAT6|STK11|PRDX2|KLF10|TPD52|XBP1|ZBTB16|CCN4|SH3PXD2A|TSPAN2|RNF41|TRIB1|ZBTB1|JMJD6|FOXP1|EIF2AK1|PRELID1|NCAPH2|ZBTB7B|ZBTB7A|LRRC8A|SH3RF1|VSIR|IL25|NHEJ1|ZC3H12A|IL31RA|IRF2BP2|MIR125B1|MIR21|MIR30B", "Description": "leukocyte differentiation", "_MEMBER_Input_ID": 1.0, "_GeneInHitList": 1562.0, "Category": "GO Biological Processes", "Log_q_value_": -6.841771750932958, "Symbol": "GO:0002521", "LogP": -10.38272655985909, "_RANK_": 1.0, "FirstInGroupByEnrichment": 1.0, "_TotalGeneInLibrary": 24367.0, "_GeneInGO": 522.0, "shared_name": "10", "_InGO": 4.801536491677337, "Gene": "GO:0002521", "_COUNT_001": 75.0, "SUID": 579207, "id_original": "10", "selected": false, "CategoryID": 19.0, "STDV_InGO": 0.5409592753355752, "EvidenceCutoff": 0.0, "PARENT_GO": "19_GO:0002376 immune system process", "FirstInGroupByLogP": 1.0, "GROUP_ID": 5.0, "name": "10", "BestEnrichmentInGroup": 6.239948783610755, "CLUSTER_LABEL": "leukocyte differentiation", "GeneID": "25|177|302|567|604|677|678|860|861|865|920|1051|1236|1316|1969|2064|2194|2353|2475|2624|3146|3204|3398|3428|3566|3659|3707|3726|3955|4055|4092|4323|4627|4920|5452|5777|5795|5914|5925|6097|6237|6659|6670|6722|6778|6794|7001|7071|7163|7494|7704|8840|9644|10100|10193|10221|22890|23210|27086|27102|27166|29781|51043|51341|56262|57630|64115|64806|79840|80149|133396|359948|406911|406991|407030", "_LogP_Input_ID": -10.38272655985909, "Z_score": 7.503298969732083, "GiniIndex": 0.0, "_COUNT_001_PCT": 100.0}, "position": {"x": -199.14993311226908, "y": 42.58025699314874}, "selected": false}, {"data": {"id": "579206", "Enrichment": 1.9845990953791603, "BestLogPInGroup": -9.059097575411174, "_PATTERN_": "M1", "_GeneInGOAndHitList": 86.0, "Hits": "ABL1|AKT1|ATP2A2|ZFP36L1|BTG1|CD9|CHD2|CRYAB|CSRP1|CTF1|DDX5|ERBB3|FHL2|FOXF1|FLNB|FLOT2|FOS|MTOR|GPC1|H3-3B|HNRNPU|HOXD10|ID3|IL4R|ILK|ISL1|ITGA7|KRT8|LMNA|SMAD3|SMAD7|MBNL1|MEF2D|MEIS1|MMP14|MYH9|NFATC3|NOTCH1|PDGFB|PPP3CA|PRKAR1A|RARA|RARB|RB1|RXRA|SDC1|SMARCD3|SRF|TARBP2|TBX3|UTRN|XBP1|BTG2|SORBS2|YBX3|PDLIM4|CFLAR|SPAG9|QKI|GDF15|FLOT1|ZMPSTE24|SPEG|DDX17|PDLIM5|KDM6B|HEY2|MYOF|MYLPF|EHD2|PLEKHO1|SEMA4C|MESP1|NIBAN2|ARID5B|UQCC2|TANC1|ALPK2|CMTM5|SIK1|BHLHA15|BCL9L|MIR125B1|MIR21|MIR25|ZBTB42", "Description": "muscle structure development", "_MEMBER_Input_ID": 1.0, "_GeneInHitList": 1562.0, "Category": "GO Biological Processes", "Log_q_value_": -5.916082775157079, "Symbol": "GO:0061061", "LogP": -9.059097575411174, "_RANK_": 1.0, "FirstInGroupByEnrichment": 1.0, "_TotalGeneInLibrary": 24367.0, "_GeneInGO": 676.0, "shared_name": "9", "_InGO": 5.505761843790013, "Gene": "GO:0061061", "_COUNT_001": 86.0, "SUID": 579206, "id_original": "9", "selected": false, "CategoryID": 19.0, "STDV_InGO": 0.5771262923289444, "EvidenceCutoff": 0.0, "PARENT_GO": "19_GO:0032502 developmental process", "FirstInGroupByLogP": 1.0, "GROUP_ID": 11.0, "name": "9", "BestEnrichmentInGroup": 2.5527063205680363, "CLUSTER_LABEL": "muscle structure development", "GeneID": "25|207|488|677|694|928|1106|1410|1465|1489|1655|2065|2274|2294|2317|2319|2353|2475|2817|3021|3192|3236|3399|3566|3611|3670|3679|3856|4000|4088|4092|4154|4209|4211|4323|4627|4775|4851|5155|5530|5573|5914|5915|5925|6256|6382|6604|6722|6895|6926|7402|7494|7832|8470|8531|8572|8837|9043|9444|9518|10211|10269|10290|10521|10611|23135|23493|26509|29895|30846|51177|54910|55897|64855|84159|84300|85461|115701|116173|150094|168620|283149|406911|406991|407014|100128927", "_LogP_Input_ID": -9.059097575411174, "Z_score": 6.7945184213818015, "GiniIndex": 0.0, "_COUNT_001_PCT": 100.0}, "position": {"x": 891.5056302422231, "y": -129.58621639552314}, "selected": false}, {"data": {"id": "579205", "Enrichment": 2.2127477956066506, "BestLogPInGroup": -8.219544186232241, "_PATTERN_": "M1", "_GeneInGOAndHitList": 60.0, "Hits": "ACTN1|ANXA2|B2M|BCL6|ZFP36L1|RUNX1|CBFB|CD4|CEBPB|CCR7|EPHA2|FASN|FOXO3|FOS|MTOR|GATA2|H3-3B|HOXA5|HOXA7|HOXA9|HSPA1A|HSPA1B|ID2|IFI16|ITPKB|JUNB|LTBR|MEIS1|MEIS2|MYH9|ROR2|PTPN6|RARA|RARG|RB1|SP3|SRF|THRA|KLF10|ZFP36|ZBTB16|H3C8|CCN4|H3C7|SH3PXD2A|TSPAN2|RNF41|TRIB1|NCOA6|JMJD6|FOXP1|EIF2AK1|ZBTB7A|KLF13|PAF1|KMT2E|IL25|KAT8|IL31RA|MIR125B1", "Description": "myeloid cell differentiation", "_MEMBER_Input_ID": 1.0, "_GeneInHitList": 1562.0, "Category": "GO Biological Processes", "Log_q_value_": -5.298378135594502, "Symbol": "GO:0030099", "LogP": -8.219544186232241, "_RANK_": 1.0, "FirstInGroupByEnrichment": 1.0, "_TotalGeneInLibrary": 24367.0, "_GeneInGO": 423.0, "shared_name": "8", "_InGO": 3.8412291933418694, "Gene": "GO:0030099", "_COUNT_001": 60.0, "SUID": 579205, "id_original": "8", "selected": false, "CategoryID": 19.0, "STDV_InGO": 0.4862829551112782, "EvidenceCutoff": 0.0, "PARENT_GO": "19_GO:0002376 immune system process", "FirstInGroupByLogP": 1.0, "GROUP_ID": 14.0, "name": "8", "BestEnrichmentInGroup": 3.1908829007100454, "CLUSTER_LABEL": "myeloid cell differentiation", "GeneID": "87|302|567|604|677|861|865|920|1051|1236|1969|2194|2309|2353|2475|2624|3021|3202|3204|3205|3303|3304|3398|3428|3707|3726|4055|4211|4212|4627|4920|5777|5914|5916|5925|6670|6722|7067|7071|7538|7704|8355|8840|8968|9644|10100|10193|10221|23054|23210|27086|27102|51341|51621|54623|55904|64806|84148|133396|406911", "_LogP_Input_ID": -8.219544186232241, "Z_score": 6.585062815310404, "GiniIndex": 0.0, "_COUNT_001_PCT": 100.0}, "position": {"x": -255.00778223336283, "y": 96.11472659764092}, "selected": false}, {"data": {"id": "579204", "Enrichment": 1.9525531437517185, "BestLogPInGroup": -9.567032373961917, "_PATTERN_": "M1", "_GeneInGOAndHitList": 95.0, "Hits": "ACTN4|B2M|CCND1|BCL6|ZFP36L1|ZFP36L2|RUNX1|CBFB|DDX6|DLX1|DLX2|EGFR|ERBB2|FBLN1|FOXO3|GATA2|HMGB1|FOXA1|HNRNPU|HOXA5|HOXA7|HOXA9|HPN|ID1|ID2|ID3|IDH2|IL4R|IRF1|ISL1|ITPKB|LRP1|LTBP3|SMAD3|SMAD7|MEIS1|MEIS2|NFATC3|NFE2L2|NOTCH1|NOTCH4|PBX1|PDGFB|PPP3CA|PTCH1|RARA|RARB|RARG|RB1|CX3CL1|SPOCK1|TBX3|PRDX2|THRB|TRIO|XBP1|ZFP36|ZBTB16|SYNGAP1|CFLAR|CCN4|CBFA2T2|ARHGEF2|NR1D1|TOB1|TRIB1|N4BP2L2|CCDC85B|SPART|KANK1|HEY2|CORO1C|SPDEF|ZBTB7B|HDAC7|KLF13|PAF1|TRPM4|SEMA4C|APPL2|MESP1|RTN4|SEMA6A|GRHL2|ZC3H12A|DIXDC1|CMTM5|DAB2IP|BHLHA15|MIR125B1|MIR149|MIR21|MIR27B|MIR29B1|MIR30B", "Description": "negative regulation of cell differentiation", "_MEMBER_Input_ID": 1.0, "_GeneInHitList": 1562.0, "Category": "GO Biological Processes", "Log_q_value_": -6.289318999810366, "Symbol": "GO:0045596", "LogP": -9.567032373961917, "_RANK_": 1.0, "FirstInGroupByEnrichment": 1.0, "_TotalGeneInLibrary": 24367.0, "_GeneInGO": 759.0, "shared_name": "7", "_InGO": 6.081946222791293, "Gene": "GO:0045596", "_COUNT_001": 95.0, "SUID": 579204, "id_original": "7", "selected": false, "CategoryID": 19.0, "STDV_InGO": 0.6047213686393024, "EvidenceCutoff": 0.0, "PARENT_GO": "19_GO:0048519 negative regulation of biological process", "FirstInGroupByLogP": 1.0, "GROUP_ID": 8.0, "name": "7", "BestEnrichmentInGroup": 3.1795917368717226, "CLUSTER_LABEL": "negative regulation of cell differentiation", "GeneID": "81|567|595|604|677|678|861|865|1656|1745|1746|1956|2064|2192|2309|2624|3146|3169|3192|3202|3204|3205|3249|3397|3398|3399|3418|3566|3659|3670|3707|4035|4054|4088|4092|4211|4212|4775|4780|4851|4855|5087|5155|5530|5727|5914|5915|5916|5925|6376|6695|6926|7001|7068|7204|7494|7538|7704|8831|8837|8840|9139|9181|9572|10140|10221|10443|11007|23111|23189|23493|23603|25803|51043|51564|51621|54623|54795|54910|55198|55897|57142|57556|79977|80149|85458|116173|153090|168620|406911|406941|406991|407019|407024|407030", "_LogP_Input_ID": -9.567032373961917, "Z_score": 6.977471974714938, "GiniIndex": 0.0, "_COUNT_001_PCT": 100.0}, "position": {"x": -243.5862887030894, "y": 274.17887836155694}, "selected": false}, {"data": {"id": "579203", "Enrichment": 1.8364661685270978, "BestLogPInGroup": -7.862534304035942, "_PATTERN_": "M1", "_GeneInGOAndHitList": 91.0, "Hits": "ALOX15B|B2M|BCL6|BMPR1B|KLF9|BTG1|CD9|CDKN1B|CEBPB|CLDN3|CTBP1|DHCR24|ERBB2|FBLN1|FTH1|GATA2|B4GALT1|SFN|RAPGEF1|HPN|HSPA1A|HSPA1B|IDH2|ILK|IRF1|ISL1|LMNA|SMAD3|SMAD7|MEIS1|MNT|MXI1|NKX3-1|NOTCH1|NPR3|ROR2|SERPINF1|PPP2R5C|PRKAR1A|PTCH1|PTPN6|PTPRJ|PTPRM|RARA|RARB|RARG|RB1|REG1A|SDC4|SOX4|SPARC|SRF|STK11|SULT2B1|KLF10|ZBTB16|BTG2|AIMP2|FOSL1|IFITM1|PER2|HGS|DLG5|ST18|TOB1|RNF41|TRIB1|SPEG|NDRG1|PEMT|P3H3|PTGDR2|PHB2|SH3BP4|EIF2AK1|ZBTB7B|KLF13|APPL2|WFDC1|VSIR|NIBAN2|COL18A1|LZTS2|ZNF503|DAB2IP|MIR125A|MIR132|MIR21|MIR29A|MIR29B1|MIR30B", "Description": "negative regulation of cell proliferation", "_MEMBER_Input_ID": 1.0, "_GeneInHitList": 1562.0, "Category": "GO Biological Processes", "Log_q_value_": -5.020549499445829, "Symbol": "GO:0008285", "LogP": -7.862534304035942, "_RANK_": 1.0, "FirstInGroupByEnrichment": 1.0, "_TotalGeneInLibrary": 24367.0, "_GeneInGO": 773.0, "shared_name": "6", "_InGO": 5.825864276568502, "Gene": "GO:0008285", "_COUNT_001": 91.0, "SUID": 579203, "id_original": "6", "selected": false, "CategoryID": 19.0, "STDV_InGO": 0.5926598246801607, "EvidenceCutoff": 0.0, "PARENT_GO": "19_GO:0008283 cell proliferation", "FirstInGroupByLogP": 1.0, "GROUP_ID": 16.0, "name": "6", "BestEnrichmentInGroup": 1.8364661685270978, "CLUSTER_LABEL": "negative regulation of cell proliferation", "GeneID": "247|567|604|658|687|694|928|1027|1051|1365|1487|1718|2064|2192|2495|2624|2683|2810|2889|3249|3303|3304|3418|3611|3659|3670|4000|4088|4092|4211|4335|4601|4824|4851|4883|4920|5176|5527|5573|5727|5777|5795|5797|5914|5915|5916|5925|5967|6385|6659|6678|6722|6794|6820|7071|7704|7832|7965|8061|8519|8864|9146|9231|9705|10140|10193|10221|10290|10397|10400|10536|11251|11331|23677|27102|51043|51621|55198|58189|64115|64855|80781|84445|84858|153090|406910|406921|406991|407021|407024|407030", "_LogP_Input_ID": -7.862534304035942, "Z_score": 6.185216932655457, "GiniIndex": 0.0, "_COUNT_001_PCT": 100.0}, "position": {"x": -387.60092188180033, "y": 1011.7290912216644}, "selected": false}, {"data": {"id": "579202", "Enrichment": 2.179393876628757, "BestLogPInGroup": -7.604995071485691, "_PATTERN_": "M1", "_GeneInGOAndHitList": 57.0, "Hits": "ADRB2|AKT1|BMPR1B|RUNX2|RUNX1|CBFB|CEBPB|COL1A1|DDX5|EGFR|EPHA2|EXT2|FASN|FGR|FHL2|H3-3B|HNRNPU|ID1|ID2|ID3|ILK|JUNB|JUND|LTBP3|SMAD3|SMAD7|MEF2D|MMP14|NAB2|NOTCH1|ROR2|PBX1|PTCH1|SP3|SPARC|THRA|KLF10|TPM4|ZBTB16|IFITM1|DCHS1|CCN4|DHRS3|TOB1|ZMPSTE24|HDAC7|WWOX|TRPM4|BCAP29|ANKH|TP53INP2|KAZALD1|CREB3L1|SH3PXD2B|MIR200C|MIR21|MIR29B1", "Description": "ossification", "_MEMBER_Input_ID": 1.0, "_GeneInHitList": 1562.0, "Category": "GO Biological Processes", "Log_q_value_": -4.804402952053802, "Symbol": "GO:0001503", "LogP": -7.604995071485691, "_RANK_": 1.0, "FirstInGroupByEnrichment": 1.0, "_TotalGeneInLibrary": 24367.0, "_GeneInGO": 408.0, "shared_name": "5", "_InGO": 3.649167733674776, "Gene": "GO:0001503", "_COUNT_001": 57.0, "SUID": 579202, "id_original": "5", "selected": false, "CategoryID": 19.0, "STDV_InGO": 0.4744430984441882, "EvidenceCutoff": 0.0, "PARENT_GO": "19_GO:0032501 multicellular organismal process", "FirstInGroupByLogP": 1.0, "GROUP_ID": 17.0, "name": "5", "BestEnrichmentInGroup": 2.6999778390623463, "CLUSTER_LABEL": "ossification", "GeneID": "154|207|658|860|861|865|1051|1277|1655|1956|1969|2132|2194|2268|2274|3021|3192|3397|3398|3399|3611|3726|3727|4054|4088|4092|4209|4323|4665|4851|4920|5087|5727|6670|6678|7067|7071|7171|7704|8519|8642|8840|9249|10140|10269|51564|51741|54795|55973|56172|58476|81621|90993|285590|406985|406991|407024", "_LogP_Input_ID": -7.604995071485691, "Z_score": 6.287416099705775, "GiniIndex": 0.0, "_COUNT_001_PCT": 100.0}, "position": {"x": 1323.2499081963247, "y": 47.99004703221124}, "selected": false}, {"data": {"id": "579201", "Enrichment": 2.1272552671400304, "BestLogPInGroup": -8.621070944395578, "_PATTERN_": "M1", "_GeneInGOAndHitList": 69.0, "Hits": "AKT1|ATF4|ZFP36L1|ZFP36L2|CIRBP|PLK3|DAPK1|DHFR|EEF1A1|EIF4G1|ERBB2|FOXO3|MTOR|MKNK2|HSPB1|NFE2L2|NOTCH4|PA2G4|PER1|RARA|UPF1|RXRA|SOX4|TARBP2|TNFRSF1A|UCN|ZFP36|BTG2|YBX3|EIF3H|PER2|EIF2B4|EIF2B5|QKI|TOB1|ABCA7|CASC3|CPEB3|SAMD4A|PASK|PPP1R15A|EIF2AK1|EIF3K|COA3|DDX25|BANK1|SAMD4B|CNOT11|SPHK2|EEFSEC|METTL16|ZC3H12A|UQCC2|PYM1|UNK|DPH6|PATL2|CNOT6L|NANOS3|MIR106B|MIR125B1|MIR132|MIR148A|MIR200C|MIR21|MIR212|MIR27B|MIR29A|MIR29B1", "Description": "regulation of cellular amide metabolic process", "_MEMBER_Input_ID": 1.0, "_GeneInHitList": 1562.0, "Category": "GO Biological Processes", "Log_q_value_": -5.585758129620247, "Symbol": "GO:0034248", "LogP": -8.621070944395578, "_RANK_": 1.0, "FirstInGroupByEnrichment": 1.0, "_TotalGeneInLibrary": 24367.0, "_GeneInGO": 506.0, "shared_name": "4", "_InGO": 4.41741357234315, "Gene": "GO:0034248", "_COUNT_001": 69.0, "SUID": 579201, "id_original": "4", "selected": false, "CategoryID": 19.0, "STDV_InGO": 0.5199156674482185, "EvidenceCutoff": 0.0, "PARENT_GO": "19_GO:0008152 metabolic process", "FirstInGroupByLogP": 1.0, "GROUP_ID": 13.0, "name": "4", "BestEnrichmentInGroup": 2.455535400957936, "CLUSTER_LABEL": "regulation of cellular amide metabolic process", "GeneID": "207|468|677|678|1153|1263|1612|1719|1915|1981|2064|2309|2475|2872|3315|4780|4855|5036|5187|5914|5976|6256|6659|6895|7132|7349|7538|7832|8531|8667|8864|8890|8893|9444|10140|10347|22794|22849|23034|23178|23645|27102|27335|28958|29118|55024|55095|55571|56848|60678|79066|80149|84300|84305|85451|89978|197135|246175|342977|406900|406911|406921|406940|406985|406991|406994|407019|407021|407024", "_LogP_Input_ID": -8.621070944395578, "Z_score": 6.70611208225811, "GiniIndex": 0.0, "_COUNT_001_PCT": 100.0}, "position": {"x": -170.55138469994608, "y": 793.8471026962737}, "selected": false}, {"data": {"id": "579200", "Enrichment": 3.02247519206146, "BestLogPInGroup": -7.5613451204042255, "_PATTERN_": "M1", "_GeneInGOAndHitList": 31.0, "Hits": "ALOX15B|CCND1|ZFP36L1|BTG1|RUNX1|CBFB|CDKN1B|CEBPB|SFN|HOXA7|ID1|NOTCH1|NOTCH4|PRKCH|PTCH1|SULT2B1|TBX3|TNFRSF1A|ZFP36|KLF7|PTCH2|ZEB2|HEY2|ESRP1|MESP1|GRHL2|APOLD1|MIR200C|MIR21|MIR29B1|MIR99B", "Description": "regulation of epithelial cell differentiation", "_MEMBER_Input_ID": 1.0, "_GeneInHitList": 1562.0, "Category": "GO Biological Processes", "Log_q_value_": -4.786307105444725, "Symbol": "GO:0030856", "LogP": -7.5613451204042255, "_RANK_": 1.0, "FirstInGroupByEnrichment": 1.0, "_TotalGeneInLibrary": 24367.0, "_GeneInGO": 160.0, "shared_name": "3", "_InGO": 1.9846350832266326, "Gene": "GO:0030856", "_COUNT_001": 31.0, "SUID": 579200, "id_original": "3", "selected": false, "CategoryID": 19.0, "STDV_InGO": 0.3528961317999472, "EvidenceCutoff": 0.0, "PARENT_GO": "19_GO:0032502 developmental process", "FirstInGroupByLogP": 1.0, "GROUP_ID": 19.0, "name": "3", "BestEnrichmentInGroup": 4.549962654716176, "CLUSTER_LABEL": "regulation of epithelial cell differentiation", "GeneID": "247|595|677|694|861|865|1027|1051|2810|3204|3397|4851|4855|5583|5727|6820|6926|7132|7538|8609|8643|9839|23493|54845|55897|79977|81575|406985|406991|407024|407056", "_LogP_Input_ID": -7.5613451204042255, "Z_score": 6.717231344558502, "GiniIndex": 0.0, "_COUNT_001_PCT": 100.0}, "position": {"x": 573.544189202184, "y": 157.73606448826592}, "selected": false}, {"data": {"id": "579199", "Enrichment": 1.964601566054361, "BestLogPInGroup": -8.650869564251822, "_PATTERN_": "M1", "_GeneInGOAndHitList": 84.0, "Hits": "ABL1|AKT1|B2M|BCL6|CFL1|CSNK1D|CUX1|DDX6|DLX1|DLX2|DVL3|EIF4G1|FOXO3|MTOR|GATA2|RAPGEF1|GRIN1|FOXA1|ID1|ID2|ILK|ISL1|ITGA3|LRP1|MEIS1|NBL1|NFE2L2|NOTCH1|ROR2|PBX1|SERPINF1|PPP3CA|PTK7|RAC3|RARA|RARB|REG1A|SCN1B|CX3CL1|SPOCK1|SRF|STK11|THRB|STK24|SYNGAP1|SOCS2|CFLAR|SPAG9|CBFA2T2|ARHGEF2|DLG5|ZEB2|BAIAP2|PDLIM5|CPEB3|SHANK2|MYCBP2|SPART|KANK1|COBL|CUX2|NEDD4L|NCS1|HEY2|MACF1|ZMYND8|ANAPC2|PACSIN1|TNFRSF12A|STYXL1|ESRP1|CASZ1|SEMA4C|RTN4|SEMA6A|FBXO31|CDK5RAP3|DIXDC1|PLXNA4|RNF157|DAB2IP|BRINP3|MIR200C|FOXO6", "Description": "regulation of neuron differentiation", "_MEMBER_Input_ID": 1.0, "_GeneInHitList": 1562.0, "Category": "GO Biological Processes", "Log_q_value_": -5.5870360100453516, "Symbol": "GO:0045664", "LogP": -8.650869564251822, "_RANK_": 1.0, "FirstInGroupByEnrichment": 1.0, "_TotalGeneInLibrary": 24367.0, "_GeneInGO": 667.0, "shared_name": "2", "_InGO": 5.377720870678617, "Gene": "GO:0045664", "_COUNT_001": 84.0, "SUID": 579199, "id_original": "2", "selected": false, "CategoryID": 19.0, "STDV_InGO": 0.570762347817035, "EvidenceCutoff": 0.0, "PARENT_GO": "19_GO:0032502 developmental process", "FirstInGroupByLogP": 1.0, "GROUP_ID": 12.0, "name": "2", "BestEnrichmentInGroup": 2.1347193207089425, "CLUSTER_LABEL": "regulation of neuron differentiation", "GeneID": "25|207|567|604|1072|1453|1523|1656|1745|1746|1857|1981|2309|2475|2624|2889|2902|3169|3397|3398|3611|3670|3675|4035|4211|4681|4780|4851|4920|5087|5176|5530|5754|5881|5914|5915|5967|6324|6376|6695|6722|6794|7068|8428|8831|8835|8837|9043|9139|9181|9231|9839|10458|10611|22849|22941|23077|23111|23189|23242|23316|23327|23413|23493|23499|23613|29882|29993|51330|51657|54845|54897|54910|57142|57556|79791|80279|85458|91584|114804|153090|339479|406985|100132074", "_LogP_Input_ID": -8.650869564251822, "Z_score": 6.610804332550497, "GiniIndex": 0.0, "_COUNT_001_PCT": 100.0}, "position": {"x": -353.86630274117533, "y": 462.06051212010186}, "selected": false}, {"data": {"id": "579198", "Enrichment": 2.0092454476894255, "BestLogPInGroup": -9.631064746783506, "_PATTERN_": "M1", "_GeneInGOAndHitList": 89.0, "Hits": "ACTB|ACTN1|AGER|DST|ZFP36L1|ZFP36L2|CAPZB|CD9|CDKN1B|COL1A1|CLDN4|CLDN3|CSRP1|DGKA|DGKQ|DHFR|EGFR|ERBB2|ERBB3|FBLN1|FOXF1|MTOR|GATA2|B4GALT1|GSN|H3-3B|HSPB1|ID3|ILK|INHBB|IRF1|ISL1|ITPK1|ITPR1|LRP1|SMAD3|MYH9|NFE2L2|NINJ1|NOTCH1|NOTCH4|PDGFB|PIK3CB|PKM|PLEC|SERPINF2|PPP3CA|PRKAR1A|PRKCH|PTK7|PTPN6|RAD51B|REG1A|RREB1|CX3CL1|SDC1|SDC4|SPARC|SRF|TARBP2|PRDX2|TLN1|XBP1|YWHAZ|ZFP36|H3C8|STK24|CFLAR|CCN4|H3C7|SYT7|FIBP|HOXB13|GNA13|ABHD2|PHB2|KANK1|MACF1|MYOF|EHD2|TNFRSF12A|WFDC1|WNT5B|ADTRP|CADM4|JMJD1C|MIA3|MIR21|MIR29B1", "Description": "response to wounding", "_MEMBER_Input_ID": 1.0, "_GeneInHitList": 1562.0, "Category": "GO Biological Processes", "Log_q_value_": -6.31195868747373, "Symbol": "GO:0009611", "LogP": -9.631064746783506, "_RANK_": 1.0, "FirstInGroupByEnrichment": 1.0, "_TotalGeneInLibrary": 24367.0, "_GeneInGO": 691.0, "shared_name": "1", "_InGO": 5.697823303457106, "Gene": "GO:0009611", "_COUNT_001": 89.0, "SUID": 579198, "id_original": "1", "selected": false, "CategoryID": 19.0, "STDV_InGO": 0.586509204822956, "EvidenceCutoff": 0.0, "PARENT_GO": "19_GO:0050896 response to stimulus", "FirstInGroupByLogP": 1.0, "GROUP_ID": 7.0, "name": "1", "BestEnrichmentInGroup": 2.8599765258215957, "CLUSTER_LABEL": "response to wounding", "GeneID": "60|87|177|667|677|678|832|928|1027|1277|1364|1365|1465|1606|1609|1719|1956|2064|2065|2192|2294|2475|2624|2683|2934|3021|3315|3399|3611|3625|3659|3670|3705|3708|4035|4088|4627|4780|4814|4851|4855|5155|5291|5315|5339|5345|5530|5573|5583|5754|5777|5890|5967|6239|6376|6382|6385|6678|6722|6895|7001|7094|7494|7534|7538|8355|8428|8837|8840|8968|9066|9158|10481|10672|11057|11331|23189|23499|26509|30846|51330|58189|81029|84830|199731|221037|375056|406991|407024", "_LogP_Input_ID": -9.631064746783506, "Z_score": 7.04367409984526, "GiniIndex": 0.0, "_COUNT_001_PCT": 100.0}, "position": {"x": -350.2060282355601, "y": 759.3035846298675}, "selected": false}], "edges": [{"data": {"id": "579918", "source": "579341", "target": "579212", "SCORE": 0.994072785386284, "shared_interaction": "Direct", "source_original": "1", "Name_B": "GO:0043009", "Name_A": "GO:0009792", "target_original": "130", "Gene_B": "GO:0043009", "interaction": "Direct", "SUID": 579918, "TYPE": "Direct", "Gene_A": "GO:0009792", "BEND_MAP_ID": 579918, "selected": false}, "selected": false}, {"data": {"id": "579917", "source": "579341", "target": "579208", "SCORE": 0.30352898153273944, "shared_interaction": "Direct", "source_original": "1", "Name_B": "GO:0048732", "Name_A": "GO:0009792", "target_original": "134", "Gene_B": "GO:0048732", "interaction": "Direct", "SUID": 579917, "TYPE": "Direct", "Gene_A": "GO:0009792", "BEND_MAP_ID": 579917, "selected": false}, "selected": false}, {"data": {"id": "579916", "source": "579341", "target": "579340", "SCORE": 0.5055601228855197, "shared_interaction": "Direct", "source_original": "1", "Name_B": "GO:0048598", "Name_A": "GO:0009792", "target_original": "2", "Gene_B": "GO:0048598", "interaction": "Direct", "SUID": 579916, "TYPE": "Direct", "Gene_A": "GO:0009792", "BEND_MAP_ID": 579916, "selected": false}, "selected": false}, {"data": {"id": "579915", "source": "579341", "target": "579338", "SCORE": 0.39516848917701797, "shared_interaction": "Direct", "source_original": "1", "Name_B": "GO:0009952", "Name_A": "GO:0009792", "target_original": "4", "Gene_B": "GO:0009952", "interaction": "Direct", "SUID": 579915, "TYPE": "Direct", "Gene_A": "GO:0009792", "BEND_MAP_ID": 579915, "selected": false}, "selected": false}, {"data": {"id": "579914", "source": "579341", "target": "579337", "SCORE": 0.5457584419362754, "shared_interaction": "Direct", "source_original": "1", "Name_B": "GO:0048568", "Name_A": "GO:0009792", "target_original": "5", "Gene_B": "GO:0048568", "interaction": "Direct", "SUID": 579914, "TYPE": "Direct", "Gene_A": "GO:0009792", "BEND_MAP_ID": 579914, "selected": false}, "selected": false}, {"data": {"id": "579913", "source": "579341", "target": "579335", "SCORE": 0.41918028395437984, "shared_interaction": "Direct", "source_original": "1", "Name_B": "GO:0048562", "Name_A": "GO:0009792", "target_original": "7", "Gene_B": "GO:0048562", "interaction": "Direct", "SUID": 579913, "TYPE": "Direct", "Gene_A": "GO:0009792", "BEND_MAP_ID": 579913, "selected": false}, "selected": false}, {"data": {"id": "579912", "source": "579341", "target": "579333", "SCORE": 0.35279843200755917, "shared_interaction": "Direct", "source_original": "1", "Name_B": "GO:0048706", "Name_A": "GO:0009792", "target_original": "9", "Gene_B": "GO:0048706", "interaction": "Direct", "SUID": 579912, "TYPE": "Direct", "Gene_A": "GO:0009792", "BEND_MAP_ID": 579912, "selected": false}, "selected": false}, {"data": {"id": "579911", "source": "579340", "target": "579237", "SCORE": 0.3186469056188752, "shared_interaction": "Direct", "source_original": "2", "Name_B": "GO:0048705", "Name_A": "GO:0048598", "target_original": "105", "Gene_B": "GO:0048705", "interaction": "Direct", "SUID": 579911, "TYPE": "Direct", "Gene_A": "GO:0048598", "BEND_MAP_ID": 579911, "selected": false}, "selected": false}, {"data": {"id": "579910", "source": "579340", "target": "579333", "SCORE": 0.34514687100962044, "shared_interaction": "Direct", "source_original": "2", "Name_B": "GO:0048706", "Name_A": "GO:0048598", "target_original": "9", "Gene_B": "GO:0048706", "interaction": "Direct", "SUID": 579910, "TYPE": "Direct", "Gene_A": "GO:0048598", "BEND_MAP_ID": 579910, "selected": false}, "selected": false}, {"data": {"id": "579909", "source": "579339", "target": "579212", "SCORE": 0.4424156288395526, "shared_interaction": "Direct", "source_original": "3", "Name_B": "GO:0043009", "Name_A": "GO:0007389", "target_original": "130", "Gene_B": "GO:0043009", "interaction": "Direct", "SUID": 579909, "TYPE": "Direct", "Gene_A": "GO:0007389", "BEND_MAP_ID": 579909, "selected": false}, "selected": false}, {"data": {"id": "579908", "source": "579339", "target": "579341", "SCORE": 0.43891229414056954, "shared_interaction": "Direct", "source_original": "3", "Name_B": "GO:0009792", "Name_A": "GO:0007389", "target_original": "1", "Gene_B": "GO:0009792", "interaction": "Direct", "SUID": 579908, "TYPE": "Direct", "Gene_A": "GO:0007389", "BEND_MAP_ID": 579908, "selected": false}, "selected": false}, {"data": {"id": "579907", "source": "579339", "target": "579340", "SCORE": 0.49236100173294656, "shared_interaction": "Direct", "source_original": "3", "Name_B": "GO:0048598", "Name_A": "GO:0007389", "target_original": "2", "Gene_B": "GO:0048598", "interaction": "Direct", "SUID": 579907, "TYPE": "Direct", "Gene_A": "GO:0007389", "BEND_MAP_ID": 579907, "selected": false}, "selected": false}, {"data": {"id": "579906", "source": "579339", "target": "579338", "SCORE": 0.7400177462292078, "shared_interaction": "Direct", "source_original": "3", "Name_B": "GO:0009952", "Name_A": "GO:0007389", "target_original": "4", "Gene_B": "GO:0009952", "interaction": "Direct", "SUID": 579906, "TYPE": "Direct", "Gene_A": "GO:0007389", "BEND_MAP_ID": 579906, "selected": false}, "selected": false}, {"data": {"id": "579905", "source": "579339", "target": "579337", "SCORE": 0.4467320460503204, "shared_interaction": "Direct", "source_original": "3", "Name_B": "GO:0048568", "Name_A": "GO:0007389", "target_original": "5", "Gene_B": "GO:0048568", "interaction": "Direct", "SUID": 579905, "TYPE": "Direct", "Gene_A": "GO:0007389", "BEND_MAP_ID": 579905, "selected": false}, "selected": false}, {"data": {"id": "579904", "source": "579339", "target": "579335", "SCORE": 0.42681478376359944, "shared_interaction": "Direct", "source_original": "3", "Name_B": "GO:0048562", "Name_A": "GO:0007389", "target_original": "7", "Gene_B": "GO:0048562", "interaction": "Direct", "SUID": 579904, "TYPE": "Direct", "Gene_A": "GO:0007389", "BEND_MAP_ID": 579904, "selected": false}, "selected": false}, {"data": {"id": "579903", "source": "579339", "target": "579333", "SCORE": 0.3278362117482446, "shared_interaction": "Direct", "source_original": "3", "Name_B": "GO:0048706", "Name_A": "GO:0007389", "target_original": "9", "Gene_B": "GO:0048706", "interaction": "Direct", "SUID": 579903, "TYPE": "Direct", "Gene_A": "GO:0007389", "BEND_MAP_ID": 579903, "selected": false}, "selected": false}, {"data": {"id": "579902", "source": "579338", "target": "579212", "SCORE": 0.39872937340167, "shared_interaction": "Direct", "source_original": "4", "Name_B": "GO:0043009", "Name_A": "GO:0009952", "target_original": "130", "Gene_B": "GO:0043009", "interaction": "Direct", "SUID": 579902, "TYPE": "Direct", "Gene_A": "GO:0009952", "BEND_MAP_ID": 579902, "selected": false}, "selected": false}, {"data": {"id": "579901", "source": "579338", "target": "579340", "SCORE": 0.4505906619168729, "shared_interaction": "Direct", "source_original": "4", "Name_B": "GO:0048598", "Name_A": "GO:0009952", "target_original": "2", "Gene_B": "GO:0048598", "interaction": "Direct", "SUID": 579901, "TYPE": "Direct", "Gene_A": "GO:0009952", "BEND_MAP_ID": 579901, "selected": false}, "selected": false}, {"data": {"id": "579900", "source": "579338", "target": "579337", "SCORE": 0.44712808513911256, "shared_interaction": "Direct", "source_original": "4", "Name_B": "GO:0048568", "Name_A": "GO:0009952", "target_original": "5", "Gene_B": "GO:0048568", "interaction": "Direct", "SUID": 579900, "TYPE": "Direct", "Gene_A": "GO:0009952", "BEND_MAP_ID": 579900, "selected": false}, "selected": false}, {"data": {"id": "579899", "source": "579338", "target": "579237", "SCORE": 0.33547266322537517, "shared_interaction": "Direct", "source_original": "4", "Name_B": "GO:0048705", "Name_A": "GO:0009952", "target_original": "105", "Gene_B": "GO:0048705", "interaction": "Direct", "SUID": 579899, "TYPE": "Direct", "Gene_A": "GO:0009952", "BEND_MAP_ID": 579899, "selected": false}, "selected": false}, {"data": {"id": "579898", "source": "579338", "target": "579335", "SCORE": 0.4696352611103063, "shared_interaction": "Direct", "source_original": "4", "Name_B": "GO:0048562", "Name_A": "GO:0009952", "target_original": "7", "Gene_B": "GO:0048562", "interaction": "Direct", "SUID": 579898, "TYPE": "Direct", "Gene_A": "GO:0009952", "BEND_MAP_ID": 579898, "selected": false}, "selected": false}, {"data": {"id": "579897", "source": "579338", "target": "579333", "SCORE": 0.40764565337360414, "shared_interaction": "Direct", "source_original": "4", "Name_B": "GO:0048706", "Name_A": "GO:0009952", "target_original": "9", "Gene_B": "GO:0048706", "interaction": "Direct", "SUID": 579897, "TYPE": "Direct", "Gene_A": "GO:0009952", "BEND_MAP_ID": 579897, "selected": false}, "selected": false}, {"data": {"id": "579896", "source": "579337", "target": "579208", "SCORE": 0.3963257597441109, "shared_interaction": "Direct", "source_original": "5", "Name_B": "GO:0048732", "Name_A": "GO:0048568", "target_original": "134", "Gene_B": "GO:0048732", "interaction": "Direct", "SUID": 579896, "TYPE": "Direct", "Gene_A": "GO:0048568", "BEND_MAP_ID": 579896, "selected": false}, "selected": false}, {"data": {"id": "579895", "source": "579337", "target": "579340", "SCORE": 0.6725908582985578, "shared_interaction": "Direct", "source_original": "5", "Name_B": "GO:0048598", "Name_A": "GO:0048568", "target_original": "2", "Gene_B": "GO:0048598", "interaction": "Direct", "SUID": 579895, "TYPE": "Direct", "Gene_A": "GO:0048568", "BEND_MAP_ID": 579895, "selected": false}, "selected": false}, {"data": {"id": "579894", "source": "579337", "target": "579237", "SCORE": 0.3978402212098363, "shared_interaction": "Direct", "source_original": "5", "Name_B": "GO:0048705", "Name_A": "GO:0048568", "target_original": "105", "Gene_B": "GO:0048705", "interaction": "Direct", "SUID": 579894, "TYPE": "Direct", "Gene_A": "GO:0048568", "BEND_MAP_ID": 579894, "selected": false}, "selected": false}, {"data": {"id": "579893", "source": "579337", "target": "579322", "SCORE": 0.3218174024869213, "shared_interaction": "Direct", "source_original": "5", "Name_B": "GO:0061008", "Name_A": "GO:0048568", "target_original": "20", "Gene_B": "GO:0061008", "interaction": "Direct", "SUID": 579893, "TYPE": "Direct", "Gene_A": "GO:0048568", "BEND_MAP_ID": 579893, "selected": false}, "selected": false}, {"data": {"id": "579892", "source": "579337", "target": "579333", "SCORE": 0.440988552667334, "shared_interaction": "Direct", "source_original": "5", "Name_B": "GO:0048706", "Name_A": "GO:0048568", "target_original": "9", "Gene_B": "GO:0048706", "interaction": "Direct", "SUID": 579892, "TYPE": "Direct", "Gene_A": "GO:0048568", "BEND_MAP_ID": 579892, "selected": false}, "selected": false}, {"data": {"id": "579891", "source": "579336", "target": "579212", "SCORE": 0.4410487103805413, "shared_interaction": "Direct", "source_original": "6", "Name_B": "GO:0043009", "Name_A": "GO:0003002", "target_original": "130", "Gene_B": "GO:0043009", "interaction": "Direct", "SUID": 579891, "TYPE": "Direct", "Gene_A": "GO:0003002", "BEND_MAP_ID": 579891, "selected": false}, "selected": false}, {"data": {"id": "579890", "source": "579336", "target": "579341", "SCORE": 0.43738959010769596, "shared_interaction": "Direct", "source_original": "6", "Name_B": "GO:0009792", "Name_A": "GO:0003002", "target_original": "1", "Gene_B": "GO:0009792", "interaction": "Direct", "SUID": 579890, "TYPE": "Direct", "Gene_A": "GO:0003002", "BEND_MAP_ID": 579890, "selected": false}, "selected": false}, {"data": {"id": "579889", "source": "579336", "target": "579340", "SCORE": 0.4770821529749386, "shared_interaction": "Direct", "source_original": "6", "Name_B": "GO:0048598", "Name_A": "GO:0003002", "target_original": "2", "Gene_B": "GO:0048598", "interaction": "Direct", "SUID": 579889, "TYPE": "Direct", "Gene_A": "GO:0003002", "BEND_MAP_ID": 579889, "selected": false}, "selected": false}, {"data": {"id": "579888", "source": "579336", "target": "579339", "SCORE": 0.8942134112319263, "shared_interaction": "Direct", "source_original": "6", "Name_B": "GO:0007389", "Name_A": "GO:0003002", "target_original": "3", "Gene_B": "GO:0007389", "interaction": "Direct", "SUID": 579888, "TYPE": "Direct", "Gene_A": "GO:0003002", "BEND_MAP_ID": 579888, "selected": false}, "selected": false}, {"data": {"id": "579887", "source": "579336", "target": "579338", "SCORE": 0.8414436165343484, "shared_interaction": "Direct", "source_original": "6", "Name_B": "GO:0009952", "Name_A": "GO:0003002", "target_original": "4", "Gene_B": "GO:0009952", "interaction": "Direct", "SUID": 579887, "TYPE": "Direct", "Gene_A": "GO:0003002", "BEND_MAP_ID": 579887, "selected": false}, "selected": false}, {"data": {"id": "579886", "source": "579336", "target": "579337", "SCORE": 0.46157701303093135, "shared_interaction": "Direct", "source_original": "6", "Name_B": "GO:0048568", "Name_A": "GO:0003002", "target_original": "5", "Gene_B": "GO:0048568", "interaction": "Direct", "SUID": 579886, "TYPE": "Direct", "Gene_A": "GO:0003002", "BEND_MAP_ID": 579886, "selected": false}, "selected": false}, {"data": {"id": "579885", "source": "579336", "target": "579237", "SCORE": 0.324537389622698, "shared_interaction": "Direct", "source_original": "6", "Name_B": "GO:0048705", "Name_A": "GO:0003002", "target_original": "105", "Gene_B": "GO:0048705", "interaction": "Direct", "SUID": 579885, "TYPE": "Direct", "Gene_A": "GO:0003002", "BEND_MAP_ID": 579885, "selected": false}, "selected": false}, {"data": {"id": "579884", "source": "579336", "target": "579335", "SCORE": 0.4399235390503396, "shared_interaction": "Direct", "source_original": "6", "Name_B": "GO:0048562", "Name_A": "GO:0003002", "target_original": "7", "Gene_B": "GO:0048562", "interaction": "Direct", "SUID": 579884, "TYPE": "Direct", "Gene_A": "GO:0003002", "BEND_MAP_ID": 579884, "selected": false}, "selected": false}, {"data": {"id": "579883", "source": "579336", "target": "579333", "SCORE": 0.38456152634432994, "shared_interaction": "Direct", "source_original": "6", "Name_B": "GO:0048706", "Name_A": "GO:0003002", "target_original": "9", "Gene_B": "GO:0048706", "interaction": "Direct", "SUID": 579883, "TYPE": "Direct", "Gene_A": "GO:0003002", "BEND_MAP_ID": 579883, "selected": false}, "selected": false}, {"data": {"id": "579882", "source": "579335", "target": "579208", "SCORE": 0.31510940848061986, "shared_interaction": "Direct", "source_original": "7", "Name_B": "GO:0048732", "Name_A": "GO:0048562", "target_original": "134", "Gene_B": "GO:0048732", "interaction": "Direct", "SUID": 579882, "TYPE": "Direct", "Gene_A": "GO:0048562", "BEND_MAP_ID": 579882, "selected": false}, "selected": false}, {"data": {"id": "579881", "source": "579335", "target": "579340", "SCORE": 0.6853409701094568, "shared_interaction": "Direct", "source_original": "7", "Name_B": "GO:0048598", "Name_A": "GO:0048562", "target_original": "2", "Gene_B": "GO:0048598", "interaction": "Direct", "SUID": 579881, "TYPE": "Direct", "Gene_A": "GO:0048562", "BEND_MAP_ID": 579881, "selected": false}, "selected": false}, {"data": {"id": "579880", "source": "579335", "target": "579337", "SCORE": 0.8330603233389676, "shared_interaction": "Direct", "source_original": "7", "Name_B": "GO:0048568", "Name_A": "GO:0048562", "target_original": "5", "Gene_B": "GO:0048568", "interaction": "Direct", "SUID": 579880, "TYPE": "Direct", "Gene_A": "GO:0048562", "BEND_MAP_ID": 579880, "selected": false}, "selected": false}, {"data": {"id": "579879", "source": "579335", "target": "579237", "SCORE": 0.46298614215537554, "shared_interaction": "Direct", "source_original": "7", "Name_B": "GO:0048705", "Name_A": "GO:0048562", "target_original": "105", "Gene_B": "GO:0048705", "interaction": "Direct", "SUID": 579879, "TYPE": "Direct", "Gene_A": "GO:0048562", "BEND_MAP_ID": 579879, "selected": false}, "selected": false}, {"data": {"id": "579878", "source": "579335", "target": "579333", "SCORE": 0.5284160284167957, "shared_interaction": "Direct", "source_original": "7", "Name_B": "GO:0048706", "Name_A": "GO:0048562", "target_original": "9", "Gene_B": "GO:0048706", "interaction": "Direct", "SUID": 579878, "TYPE": "Direct", "Gene_A": "GO:0048562", "BEND_MAP_ID": 579878, "selected": false}, "selected": false}, {"data": {"id": "579877", "source": "579334", "target": "579212", "SCORE": 0.6827573761619289, "shared_interaction": "Direct", "source_original": "8", "Name_B": "GO:0043009", "Name_A": "GO:0001701", "target_original": "130", "Gene_B": "GO:0043009", "interaction": "Direct", "SUID": 579877, "TYPE": "Direct", "Gene_A": "GO:0001701", "BEND_MAP_ID": 579877, "selected": false}, "selected": false}, {"data": {"id": "579876", "source": "579334", "target": "579341", "SCORE": 0.6774366983284124, "shared_interaction": "Direct", "source_original": "8", "Name_B": "GO:0009792", "Name_A": "GO:0001701", "target_original": "1", "Gene_B": "GO:0009792", "interaction": "Direct", "SUID": 579876, "TYPE": "Direct", "Gene_A": "GO:0001701", "BEND_MAP_ID": 579876, "selected": false}, "selected": false}, {"data": {"id": "579875", "source": "579334", "target": "579337", "SCORE": 0.3832609421990668, "shared_interaction": "Direct", "source_original": "8", "Name_B": "GO:0048568", "Name_A": "GO:0001701", "target_original": "5", "Gene_B": "GO:0048568", "interaction": "Direct", "SUID": 579875, "TYPE": "Direct", "Gene_A": "GO:0001701", "BEND_MAP_ID": 579875, "selected": false}, "selected": false}, {"data": {"id": "579874", "source": "579333", "target": "579234", "SCORE": 0.3196183012241896, "shared_interaction": "Direct", "source_original": "9", "Name_B": "GO:0051216", "Name_A": "GO:0048706", "target_original": "108", "Gene_B": "GO:0051216", "interaction": "Direct", "SUID": 579874, "TYPE": "Direct", "Gene_A": "GO:0048706", "BEND_MAP_ID": 579874, "selected": false}, "selected": false}, {"data": {"id": "579873", "source": "579332", "target": "579327", "SCORE": 0.4367984007385437, "shared_interaction": "Direct", "source_original": "10", "Name_B": "GO:0090130", "Name_A": "GO:0048514", "target_original": "15", "Gene_B": "GO:0090130", "interaction": "Direct", "SUID": 579873, "TYPE": "Direct", "Gene_A": "GO:0048514", "BEND_MAP_ID": 579873, "selected": false}, "selected": false}, {"data": {"id": "579872", "source": "579332", "target": "579324", "SCORE": 0.4335057741552127, "shared_interaction": "Direct", "source_original": "10", "Name_B": "GO:0090132", "Name_A": "GO:0048514", "target_original": "18", "Gene_B": "GO:0090132", "interaction": "Direct", "SUID": 579872, "TYPE": "Direct", "Gene_A": "GO:0048514", "BEND_MAP_ID": 579872, "selected": false}, "selected": false}, {"data": {"id": "579871", "source": "579331", "target": "579215", "SCORE": 0.8557893441615239, "shared_interaction": "Direct", "source_original": "11", "Name_B": "GO:0001568", "Name_A": "GO:0001525", "target_original": "127", "Gene_B": "GO:0001568", "interaction": "Direct", "SUID": 579871, "TYPE": "Direct", "Gene_A": "GO:0001525", "BEND_MAP_ID": 579871, "selected": false}, "selected": false}, {"data": {"id": "579870", "source": "579331", "target": "579332", "SCORE": 0.9230870192939199, "shared_interaction": "Direct", "source_original": "11", "Name_B": "GO:0048514", "Name_A": "GO:0001525", "target_original": "10", "Gene_B": "GO:0048514", "interaction": "Direct", "SUID": 579870, "TYPE": "Direct", "Gene_A": "GO:0001525", "BEND_MAP_ID": 579870, "selected": false}, "selected": false}, {"data": {"id": "579869", "source": "579331", "target": "579327", "SCORE": 0.4517705540665793, "shared_interaction": "Direct", "source_original": "11", "Name_B": "GO:0090130", "Name_A": "GO:0001525", "target_original": "15", "Gene_B": "GO:0090130", "interaction": "Direct", "SUID": 579869, "TYPE": "Direct", "Gene_A": "GO:0001525", "BEND_MAP_ID": 579869, "selected": false}, "selected": false}, {"data": {"id": "579868", "source": "579331", "target": "579325", "SCORE": 0.45023784355222257, "shared_interaction": "Direct", "source_original": "11", "Name_B": "GO:0001667", "Name_A": "GO:0001525", "target_original": "17", "Gene_B": "GO:0001667", "interaction": "Direct", "SUID": 579868, "TYPE": "Direct", "Gene_A": "GO:0001525", "BEND_MAP_ID": 579868, "selected": false}, "selected": false}, {"data": {"id": "579867", "source": "579331", "target": "579324", "SCORE": 0.4649318039153025, "shared_interaction": "Direct", "source_original": "11", "Name_B": "GO:0090132", "Name_A": "GO:0001525", "target_original": "18", "Gene_B": "GO:0090132", "interaction": "Direct", "SUID": 579867, "TYPE": "Direct", "Gene_A": "GO:0001525", "BEND_MAP_ID": 579867, "selected": false}, "selected": false}, {"data": {"id": "579866", "source": "579330", "target": "579329", "SCORE": 0.9853916338435565, "shared_interaction": "Direct", "source_original": "12", "Name_B": "GO:2000147", "Name_A": "GO:0051272", "target_original": "13", "Gene_B": "GO:2000147", "interaction": "Direct", "SUID": 579866, "TYPE": "Direct", "Gene_A": "GO:0051272", "BEND_MAP_ID": 579866, "selected": false}, "selected": false}, {"data": {"id": "579865", "source": "579330", "target": "579327", "SCORE": 0.41801485405081007, "shared_interaction": "Direct", "source_original": "12", "Name_B": "GO:0090130", "Name_A": "GO:0051272", "target_original": "15", "Gene_B": "GO:0090130", "interaction": "Direct", "SUID": 579865, "TYPE": "Direct", "Gene_A": "GO:0051272", "BEND_MAP_ID": 579865, "selected": false}, "selected": false}, {"data": {"id": "579864", "source": "579330", "target": "579324", "SCORE": 0.41348248454569253, "shared_interaction": "Direct", "source_original": "12", "Name_B": "GO:0090132", "Name_A": "GO:0051272", "target_original": "18", "Gene_B": "GO:0090132", "interaction": "Direct", "SUID": 579864, "TYPE": "Direct", "Gene_A": "GO:0051272", "BEND_MAP_ID": 579864, "selected": false}, "selected": false}, {"data": {"id": "579863", "source": "579328", "target": "579332", "SCORE": 0.30594268892921966, "shared_interaction": "Direct", "source_original": "14", "Name_B": "GO:0048514", "Name_A": "GO:0030335", "target_original": "10", "Gene_B": "GO:0048514", "interaction": "Direct", "SUID": 579863, "TYPE": "Direct", "Gene_A": "GO:0030335", "BEND_MAP_ID": 579863, "selected": false}, "selected": false}, {"data": {"id": "579862", "source": "579328", "target": "579330", "SCORE": 0.9627662278249655, "shared_interaction": "Direct", "source_original": "14", "Name_B": "GO:0051272", "Name_A": "GO:0030335", "target_original": "12", "Gene_B": "GO:0051272", "interaction": "Direct", "SUID": 579862, "TYPE": "Direct", "Gene_A": "GO:0030335", "BEND_MAP_ID": 579862, "selected": false}, "selected": false}, {"data": {"id": "579861", "source": "579328", "target": "579329", "SCORE": 0.977362280769545, "shared_interaction": "Direct", "source_original": "14", "Name_B": "GO:2000147", "Name_A": "GO:0030335", "target_original": "13", "Gene_B": "GO:2000147", "interaction": "Direct", "SUID": 579861, "TYPE": "Direct", "Gene_A": "GO:0030335", "BEND_MAP_ID": 579861, "selected": false}, "selected": false}, {"data": {"id": "579860", "source": "579328", "target": "579327", "SCORE": 0.4385712788735635, "shared_interaction": "Direct", "source_original": "14", "Name_B": "GO:0090130", "Name_A": "GO:0030335", "target_original": "15", "Gene_B": "GO:0090130", "interaction": "Direct", "SUID": 579860, "TYPE": "Direct", "Gene_A": "GO:0030335", "BEND_MAP_ID": 579860, "selected": false}, "selected": false}, {"data": {"id": "579859", "source": "579328", "target": "579326", "SCORE": 0.977362280769545, "shared_interaction": "Direct", "source_original": "14", "Name_B": "GO:0040017", "Name_A": "GO:0030335", "target_original": "16", "Gene_B": "GO:0040017", "interaction": "Direct", "SUID": 579859, "TYPE": "Direct", "Gene_A": "GO:0030335", "BEND_MAP_ID": 579859, "selected": false}, "selected": false}, {"data": {"id": "579858", "source": "579328", "target": "579324", "SCORE": 0.4342068965522328, "shared_interaction": "Direct", "source_original": "14", "Name_B": "GO:0090132", "Name_A": "GO:0030335", "target_original": "18", "Gene_B": "GO:0090132", "interaction": "Direct", "SUID": 579858, "TYPE": "Direct", "Gene_A": "GO:0030335", "BEND_MAP_ID": 579858, "selected": false}, "selected": false}, {"data": {"id": "579857", "source": "579327", "target": "579329", "SCORE": 0.4260444990210453, "shared_interaction": "Direct", "source_original": "15", "Name_B": "GO:2000147", "Name_A": "GO:0090130", "target_original": "13", "Gene_B": "GO:2000147", "interaction": "Direct", "SUID": 579857, "TYPE": "Direct", "Gene_A": "GO:0090130", "BEND_MAP_ID": 579857, "selected": false}, "selected": false}, {"data": {"id": "579856", "source": "579327", "target": "579324", "SCORE": 0.968361947953818, "shared_interaction": "Direct", "source_original": "15", "Name_B": "GO:0090132", "Name_A": "GO:0090130", "target_original": "18", "Gene_B": "GO:0090132", "interaction": "Direct", "SUID": 579856, "TYPE": "Direct", "Gene_A": "GO:0090130", "BEND_MAP_ID": 579856, "selected": false}, "selected": false}, {"data": {"id": "579855", "source": "579326", "target": "579330", "SCORE": 0.9853916338435565, "shared_interaction": "Direct", "source_original": "16", "Name_B": "GO:0051272", "Name_A": "GO:0040017", "target_original": "12", "Gene_B": "GO:0051272", "interaction": "Direct", "SUID": 579855, "TYPE": "Direct", "Gene_A": "GO:0040017", "BEND_MAP_ID": 579855, "selected": false}, "selected": false}, {"data": {"id": "579854", "source": "579326", "target": "579329", "SCORE": 1.0, "shared_interaction": "Direct", "source_original": "16", "Name_B": "GO:2000147", "Name_A": "GO:0040017", "target_original": "13", "Gene_B": "GO:2000147", "interaction": "Direct", "SUID": 579854, "TYPE": "Direct", "Gene_A": "GO:0040017", "BEND_MAP_ID": 579854, "selected": false}, "selected": false}, {"data": {"id": "579853", "source": "579326", "target": "579327", "SCORE": 0.4260444990210453, "shared_interaction": "Direct", "source_original": "16", "Name_B": "GO:0090130", "Name_A": "GO:0040017", "target_original": "15", "Gene_B": "GO:0090130", "interaction": "Direct", "SUID": 579853, "TYPE": "Direct", "Gene_A": "GO:0040017", "BEND_MAP_ID": 579853, "selected": false}, "selected": false}, {"data": {"id": "579852", "source": "579326", "target": "579324", "SCORE": 0.42157214644702573, "shared_interaction": "Direct", "source_original": "16", "Name_B": "GO:0090132", "Name_A": "GO:0040017", "target_original": "18", "Gene_B": "GO:0090132", "interaction": "Direct", "SUID": 579852, "TYPE": "Direct", "Gene_A": "GO:0040017", "BEND_MAP_ID": 579852, "selected": false}, "selected": false}, {"data": {"id": "579851", "source": "579325", "target": "579332", "SCORE": 0.4211745768503232, "shared_interaction": "Direct", "source_original": "17", "Name_B": "GO:0048514", "Name_A": "GO:0001667", "target_original": "10", "Gene_B": "GO:0048514", "interaction": "Direct", "SUID": 579851, "TYPE": "Direct", "Gene_A": "GO:0001667", "BEND_MAP_ID": 579851, "selected": false}, "selected": false}, {"data": {"id": "579850", "source": "579325", "target": "579330", "SCORE": 0.4185559723812603, "shared_interaction": "Direct", "source_original": "17", "Name_B": "GO:0051272", "Name_A": "GO:0001667", "target_original": "12", "Gene_B": "GO:0051272", "interaction": "Direct", "SUID": 579850, "TYPE": "Direct", "Gene_A": "GO:0001667", "BEND_MAP_ID": 579850, "selected": false}, "selected": false}, {"data": {"id": "579849", "source": "579325", "target": "579329", "SCORE": 0.4262665820904273, "shared_interaction": "Direct", "source_original": "17", "Name_B": "GO:2000147", "Name_A": "GO:0001667", "target_original": "13", "Gene_B": "GO:2000147", "interaction": "Direct", "SUID": 579849, "TYPE": "Direct", "Gene_A": "GO:0001667", "BEND_MAP_ID": 579849, "selected": false}, "selected": false}, {"data": {"id": "579848", "source": "579325", "target": "579328", "SCORE": 0.4382621199096796, "shared_interaction": "Direct", "source_original": "17", "Name_B": "GO:0030335", "Name_A": "GO:0001667", "target_original": "14", "Gene_B": "GO:0030335", "interaction": "Direct", "SUID": 579848, "TYPE": "Direct", "Gene_A": "GO:0001667", "BEND_MAP_ID": 579848, "selected": false}, "selected": false}, {"data": {"id": "579847", "source": "579325", "target": "579327", "SCORE": 0.8474231501522994, "shared_interaction": "Direct", "source_original": "17", "Name_B": "GO:0090130", "Name_A": "GO:0001667", "target_original": "15", "Gene_B": "GO:0090130", "interaction": "Direct", "SUID": 579847, "TYPE": "Direct", "Gene_A": "GO:0001667", "BEND_MAP_ID": 579847, "selected": false}, "selected": false}, {"data": {"id": "579846", "source": "579325", "target": "579326", "SCORE": 0.4262665820904273, "shared_interaction": "Direct", "source_original": "17", "Name_B": "GO:0040017", "Name_A": "GO:0001667", "target_original": "16", "Gene_B": "GO:0040017", "interaction": "Direct", "SUID": 579846, "TYPE": "Direct", "Gene_A": "GO:0001667", "BEND_MAP_ID": 579846, "selected": false}, "selected": false}, {"data": {"id": "579845", "source": "579325", "target": "579324", "SCORE": 0.8727576415568307, "shared_interaction": "Direct", "source_original": "17", "Name_B": "GO:0090132", "Name_A": "GO:0001667", "target_original": "18", "Gene_B": "GO:0090132", "interaction": "Direct", "SUID": 579845, "TYPE": "Direct", "Gene_A": "GO:0001667", "BEND_MAP_ID": 579845, "selected": false}, "selected": false}, {"data": {"id": "579844", "source": "579324", "target": "579329", "SCORE": 0.42157214644702573, "shared_interaction": "Direct", "source_original": "18", "Name_B": "GO:2000147", "Name_A": "GO:0090132", "target_original": "13", "Gene_B": "GO:2000147", "interaction": "Direct", "SUID": 579844, "TYPE": "Direct", "Gene_A": "GO:0090132", "BEND_MAP_ID": 579844, "selected": false}, "selected": false}, {"data": {"id": "579843", "source": "579323", "target": "579208", "SCORE": 0.5068094873762228, "shared_interaction": "Direct", "source_original": "19", "Name_B": "GO:0048732", "Name_A": "GO:0001889", "target_original": "134", "Gene_B": "GO:0048732", "interaction": "Direct", "SUID": 579843, "TYPE": "Direct", "Gene_A": "GO:0001889", "BEND_MAP_ID": 579843, "selected": false}, "selected": false}, {"data": {"id": "579842", "source": "579323", "target": "579337", "SCORE": 0.3218174024869213, "shared_interaction": "Direct", "source_original": "19", "Name_B": "GO:0048568", "Name_A": "GO:0001889", "target_original": "5", "Gene_B": "GO:0048568", "interaction": "Direct", "SUID": 579842, "TYPE": "Direct", "Gene_A": "GO:0001889", "BEND_MAP_ID": 579842, "selected": false}, "selected": false}, {"data": {"id": "579841", "source": "579323", "target": "579322", "SCORE": 1.0, "shared_interaction": "Direct", "source_original": "19", "Name_B": "GO:0061008", "Name_A": "GO:0001889", "target_original": "20", "Gene_B": "GO:0061008", "interaction": "Direct", "SUID": 579841, "TYPE": "Direct", "Gene_A": "GO:0001889", "BEND_MAP_ID": 579841, "selected": false}, "selected": false}, {"data": {"id": "579840", "source": "579323", "target": "579321", "SCORE": 0.3698924731198064, "shared_interaction": "Direct", "source_original": "19", "Name_B": "GO:0048565", "Name_A": "GO:0001889", "target_original": "21", "Gene_B": "GO:0048565", "interaction": "Direct", "SUID": 579840, "TYPE": "Direct", "Gene_A": "GO:0001889", "BEND_MAP_ID": 579840, "selected": false}, "selected": false}, {"data": {"id": "579839", "source": "579323", "target": "579320", "SCORE": 0.4078894188034386, "shared_interaction": "Direct", "source_original": "19", "Name_B": "GO:0055123", "Name_A": "GO:0001889", "target_original": "22", "Gene_B": "GO:0055123", "interaction": "Direct", "SUID": 579839, "TYPE": "Direct", "Gene_A": "GO:0001889", "BEND_MAP_ID": 579839, "selected": false}, "selected": false}, {"data": {"id": "579838", "source": "579321", "target": "579322", "SCORE": 0.3698924731198064, "shared_interaction": "Direct", "source_original": "21", "Name_B": "GO:0061008", "Name_A": "GO:0048565", "target_original": "20", "Gene_B": "GO:0061008", "interaction": "Direct", "SUID": 579838, "TYPE": "Direct", "Gene_A": "GO:0048565", "BEND_MAP_ID": 579838, "selected": false}, "selected": false}, {"data": {"id": "579837", "source": "579321", "target": "579320", "SCORE": 0.9725398313027933, "shared_interaction": "Direct", "source_original": "21", "Name_B": "GO:0055123", "Name_A": "GO:0048565", "target_original": "22", "Gene_B": "GO:0055123", "interaction": "Direct", "SUID": 579837, "TYPE": "Direct", "Gene_A": "GO:0048565", "BEND_MAP_ID": 579837, "selected": false}, "selected": false}, {"data": {"id": "579836", "source": "579320", "target": "579322", "SCORE": 0.4078894188034386, "shared_interaction": "Direct", "source_original": "22", "Name_B": "GO:0061008", "Name_A": "GO:0055123", "target_original": "20", "Gene_B": "GO:0061008", "interaction": "Direct", "SUID": 579836, "TYPE": "Direct", "Gene_A": "GO:0055123", "BEND_MAP_ID": 579836, "selected": false}, "selected": false}, {"data": {"id": "579835", "source": "579319", "target": "579210", "SCORE": 0.5597127066197923, "shared_interaction": "Direct", "source_original": "23", "Name_B": "GO:0030855", "Name_A": "GO:0002064", "target_original": "132", "Gene_B": "GO:0030855", "interaction": "Direct", "SUID": 579835, "TYPE": "Direct", "Gene_A": "GO:0002064", "BEND_MAP_ID": 579835, "selected": false}, "selected": false}, {"data": {"id": "579834", "source": "579318", "target": "579214", "SCORE": 0.3338092216587224, "shared_interaction": "Direct", "source_original": "24", "Name_B": "GO:0034329", "Name_A": "GO:0030155", "target_original": "128", "Gene_B": "GO:0034329", "interaction": "Direct", "SUID": 579834, "TYPE": "Direct", "Gene_A": "GO:0030155", "BEND_MAP_ID": 579834, "selected": false}, "selected": false}, {"data": {"id": "579833", "source": "579318", "target": "579294", "SCORE": 0.5244337902967497, "shared_interaction": "Direct", "source_original": "24", "Name_B": "GO:0031589", "Name_A": "GO:0030155", "target_original": "48", "Gene_B": "GO:0031589", "interaction": "Direct", "SUID": 579833, "TYPE": "Direct", "Gene_A": "GO:0030155", "BEND_MAP_ID": 579833, "selected": false}, "selected": false}, {"data": {"id": "579832", "source": "579318", "target": "579293", "SCORE": 0.3238169324522186, "shared_interaction": "Direct", "source_original": "24", "Name_B": "GO:0034330", "Name_A": "GO:0030155", "target_original": "49", "Gene_B": "GO:0034330", "interaction": "Direct", "SUID": 579832, "TYPE": "Direct", "Gene_A": "GO:0030155", "BEND_MAP_ID": 579832, "selected": false}, "selected": false}, {"data": {"id": "579831", "source": "579318", "target": "579316", "SCORE": 0.3159409273542201, "shared_interaction": "Direct", "source_original": "24", "Name_B": "GO:0030217", "Name_A": "GO:0030155", "target_original": "26", "Gene_B": "GO:0030217", "interaction": "Direct", "SUID": 579831, "TYPE": "Direct", "Gene_A": "GO:0030155", "BEND_MAP_ID": 579831, "selected": false}, "selected": false}, {"data": {"id": "579830", "source": "579318", "target": "579330", "SCORE": 0.3133925808024961, "shared_interaction": "Direct", "source_original": "24", "Name_B": "GO:0051272", "Name_A": "GO:0030155", "target_original": "12", "Gene_B": "GO:0051272", "interaction": "Direct", "SUID": 579830, "TYPE": "Direct", "Gene_A": "GO:0030155", "BEND_MAP_ID": 579830, "selected": false}, "selected": false}, {"data": {"id": "579829", "source": "579318", "target": "579329", "SCORE": 0.3058113544930786, "shared_interaction": "Direct", "source_original": "24", "Name_B": "GO:2000147", "Name_A": "GO:0030155", "target_original": "13", "Gene_B": "GO:2000147", "interaction": "Direct", "SUID": 579829, "TYPE": "Direct", "Gene_A": "GO:0030155", "BEND_MAP_ID": 579829, "selected": false}, "selected": false}, {"data": {"id": "579828", "source": "579318", "target": "579328", "SCORE": 0.30077956317235577, "shared_interaction": "Direct", "source_original": "24", "Name_B": "GO:0030335", "Name_A": "GO:0030155", "target_original": "14", "Gene_B": "GO:0030335", "interaction": "Direct", "SUID": 579828, "TYPE": "Direct", "Gene_A": "GO:0030155", "BEND_MAP_ID": 579828, "selected": false}, "selected": false}, {"data": {"id": "579827", "source": "579318", "target": "579314", "SCORE": 0.7325916972942909, "shared_interaction": "Direct", "source_original": "24", "Name_B": "GO:0045785", "Name_A": "GO:0030155", "target_original": "28", "Gene_B": "GO:0045785", "interaction": "Direct", "SUID": 579827, "TYPE": "Direct", "Gene_A": "GO:0030155", "BEND_MAP_ID": 579827, "selected": false}, "selected": false}, {"data": {"id": "579826", "source": "579318", "target": "579326", "SCORE": 0.3058113544930786, "shared_interaction": "Direct", "source_original": "24", "Name_B": "GO:0040017", "Name_A": "GO:0030155", "target_original": "16", "Gene_B": "GO:0040017", "interaction": "Direct", "SUID": 579826, "TYPE": "Direct", "Gene_A": "GO:0030155", "BEND_MAP_ID": 579826, "selected": false}, "selected": false}, {"data": {"id": "579825", "source": "579318", "target": "579312", "SCORE": 0.3600032359846492, "shared_interaction": "Direct", "source_original": "24", "Name_B": "GO:1902105", "Name_A": "GO:0030155", "target_original": "30", "Gene_B": "GO:1902105", "interaction": "Direct", "SUID": 579825, "TYPE": "Direct", "Gene_A": "GO:0030155", "BEND_MAP_ID": 579825, "selected": false}, "selected": false}, {"data": {"id": "579824", "source": "579318", "target": "579311", "SCORE": 0.3941126383657026, "shared_interaction": "Direct", "source_original": "24", "Name_B": "GO:1903038", "Name_A": "GO:0030155", "target_original": "31", "Gene_B": "GO:1903038", "interaction": "Direct", "SUID": 579824, "TYPE": "Direct", "Gene_A": "GO:0030155", "BEND_MAP_ID": 579824, "selected": false}, "selected": false}, {"data": {"id": "579823", "source": "579317", "target": "579298", "SCORE": 0.6468719184839978, "shared_interaction": "Direct", "source_original": "25", "Name_B": "GO:1903707", "Name_A": "GO:1903706", "target_original": "44", "Gene_B": "GO:1903707", "interaction": "Direct", "SUID": 579823, "TYPE": "Direct", "Gene_A": "GO:1903706", "BEND_MAP_ID": 579823, "selected": false}, "selected": false}, {"data": {"id": "579822", "source": "579316", "target": "579298", "SCORE": 0.45570981619162865, "shared_interaction": "Direct", "source_original": "26", "Name_B": "GO:1903707", "Name_A": "GO:0030217", "target_original": "44", "Gene_B": "GO:1903707", "interaction": "Direct", "SUID": 579822, "TYPE": "Direct", "Gene_A": "GO:0030217", "BEND_MAP_ID": 579822, "selected": false}, "selected": false}, {"data": {"id": "579821", "source": "579316", "target": "579317", "SCORE": 0.5127439334808849, "shared_interaction": "Direct", "source_original": "26", "Name_B": "GO:1903706", "Name_A": "GO:0030217", "target_original": "25", "Gene_B": "GO:1903706", "interaction": "Direct", "SUID": 579821, "TYPE": "Direct", "Gene_A": "GO:0030217", "BEND_MAP_ID": 579821, "selected": false}, "selected": false}, {"data": {"id": "579820", "source": "579316", "target": "579314", "SCORE": 0.37531325977251934, "shared_interaction": "Direct", "source_original": "26", "Name_B": "GO:0045785", "Name_A": "GO:0030217", "target_original": "28", "Gene_B": "GO:0045785", "interaction": "Direct", "SUID": 579820, "TYPE": "Direct", "Gene_A": "GO:0030217", "BEND_MAP_ID": 579820, "selected": false}, "selected": false}, {"data": {"id": "579819", "source": "579316", "target": "579312", "SCORE": 0.6547374852708118, "shared_interaction": "Direct", "source_original": "26", "Name_B": "GO:1902105", "Name_A": "GO:0030217", "target_original": "30", "Gene_B": "GO:1902105", "interaction": "Direct", "SUID": 579819, "TYPE": "Direct", "Gene_A": "GO:0030217", "BEND_MAP_ID": 579819, "selected": false}, "selected": false}, {"data": {"id": "579818", "source": "579316", "target": "579311", "SCORE": 0.422723292470278, "shared_interaction": "Direct", "source_original": "26", "Name_B": "GO:1903038", "Name_A": "GO:0030217", "target_original": "31", "Gene_B": "GO:1903038", "interaction": "Direct", "SUID": 579818, "TYPE": "Direct", "Gene_A": "GO:0030217", "BEND_MAP_ID": 579818, "selected": false}, "selected": false}, {"data": {"id": "579817", "source": "579316", "target": "579310", "SCORE": 0.48189141397702395, "shared_interaction": "Direct", "source_original": "26", "Name_B": "GO:0045581", "Name_A": "GO:0030217", "target_original": "32", "Gene_B": "GO:0045581", "interaction": "Direct", "SUID": 579817, "TYPE": "Direct", "Gene_A": "GO:0030217", "BEND_MAP_ID": 579817, "selected": false}, "selected": false}, {"data": {"id": "579816", "source": "579315", "target": "579318", "SCORE": 0.3258918413273263, "shared_interaction": "Direct", "source_original": "27", "Name_B": "GO:0030155", "Name_A": "GO:0030098", "target_original": "24", "Gene_B": "GO:0030155", "interaction": "Direct", "SUID": 579816, "TYPE": "Direct", "Gene_A": "GO:0030098", "BEND_MAP_ID": 579816, "selected": false}, "selected": false}, {"data": {"id": "579815", "source": "579315", "target": "579298", "SCORE": 0.41407479896256577, "shared_interaction": "Direct", "source_original": "27", "Name_B": "GO:1903707", "Name_A": "GO:0030098", "target_original": "44", "Gene_B": "GO:1903707", "interaction": "Direct", "SUID": 579815, "TYPE": "Direct", "Gene_A": "GO:0030098", "BEND_MAP_ID": 579815, "selected": false}, "selected": false}, {"data": {"id": "579814", "source": "579315", "target": "579205", "SCORE": 0.30987642370923213, "shared_interaction": "Direct", "source_original": "27", "Name_B": "GO:0030099", "Name_A": "GO:0030098", "target_original": "137", "Gene_B": "GO:0030099", "interaction": "Direct", "SUID": 579814, "TYPE": "Direct", "Gene_A": "GO:0030098", "BEND_MAP_ID": 579814, "selected": false}, "selected": false}, {"data": {"id": "579813", "source": "579315", "target": "579317", "SCORE": 0.528742209369289, "shared_interaction": "Direct", "source_original": "27", "Name_B": "GO:1903706", "Name_A": "GO:0030098", "target_original": "25", "Gene_B": "GO:1903706", "interaction": "Direct", "SUID": 579813, "TYPE": "Direct", "Gene_A": "GO:0030098", "BEND_MAP_ID": 579813, "selected": false}, "selected": false}, {"data": {"id": "579812", "source": "579315", "target": "579316", "SCORE": 0.8743126779239436, "shared_interaction": "Direct", "source_original": "27", "Name_B": "GO:0030217", "Name_A": "GO:0030098", "target_original": "26", "Gene_B": "GO:0030217", "interaction": "Direct", "SUID": 579812, "TYPE": "Direct", "Gene_A": "GO:0030098", "BEND_MAP_ID": 579812, "selected": false}, "selected": false}, {"data": {"id": "579811", "source": "579315", "target": "579314", "SCORE": 0.3676721012971799, "shared_interaction": "Direct", "source_original": "27", "Name_B": "GO:0045785", "Name_A": "GO:0030098", "target_original": "28", "Gene_B": "GO:0045785", "interaction": "Direct", "SUID": 579811, "TYPE": "Direct", "Gene_A": "GO:0030098", "BEND_MAP_ID": 579811, "selected": false}, "selected": false}, {"data": {"id": "579810", "source": "579315", "target": "579312", "SCORE": 0.6607552922564192, "shared_interaction": "Direct", "source_original": "27", "Name_B": "GO:1902105", "Name_A": "GO:0030098", "target_original": "30", "Gene_B": "GO:1902105", "interaction": "Direct", "SUID": 579810, "TYPE": "Direct", "Gene_A": "GO:0030098", "BEND_MAP_ID": 579810, "selected": false}, "selected": false}, {"data": {"id": "579809", "source": "579315", "target": "579311", "SCORE": 0.38281140289369914, "shared_interaction": "Direct", "source_original": "27", "Name_B": "GO:1903038", "Name_A": "GO:0030098", "target_original": "31", "Gene_B": "GO:1903038", "interaction": "Direct", "SUID": 579809, "TYPE": "Direct", "Gene_A": "GO:0030098", "BEND_MAP_ID": 579809, "selected": false}, "selected": false}, {"data": {"id": "579808", "source": "579315", "target": "579310", "SCORE": 0.39556472408554005, "shared_interaction": "Direct", "source_original": "27", "Name_B": "GO:0045581", "Name_A": "GO:0030098", "target_original": "32", "Gene_B": "GO:0045581", "interaction": "Direct", "SUID": 579808, "TYPE": "Direct", "Gene_A": "GO:0030098", "BEND_MAP_ID": 579808, "selected": false}, "selected": false}, {"data": {"id": "579807", "source": "579314", "target": "579312", "SCORE": 0.39230754986084654, "shared_interaction": "Direct", "source_original": "28", "Name_B": "GO:1902105", "Name_A": "GO:0045785", "target_original": "30", "Gene_B": "GO:1902105", "interaction": "Direct", "SUID": 579807, "TYPE": "Direct", "Gene_A": "GO:0045785", "BEND_MAP_ID": 579807, "selected": false}, "selected": false}, {"data": {"id": "579806", "source": "579314", "target": "579311", "SCORE": 0.3354827233521347, "shared_interaction": "Direct", "source_original": "28", "Name_B": "GO:1903038", "Name_A": "GO:0045785", "target_original": "31", "Gene_B": "GO:1903038", "interaction": "Direct", "SUID": 579806, "TYPE": "Direct", "Gene_A": "GO:0045785", "BEND_MAP_ID": 579806, "selected": false}, "selected": false}, {"data": {"id": "579805", "source": "579313", "target": "579318", "SCORE": 0.617931181308013, "shared_interaction": "Direct", "source_original": "29", "Name_B": "GO:0030155", "Name_A": "GO:0007162", "target_original": "24", "Gene_B": "GO:0030155", "interaction": "Direct", "SUID": 579805, "TYPE": "Direct", "Gene_A": "GO:0007162", "BEND_MAP_ID": 579805, "selected": false}, "selected": false}, {"data": {"id": "579804", "source": "579313", "target": "579295", "SCORE": 0.36819407008166055, "shared_interaction": "Direct", "source_original": "29", "Name_B": "GO:0010810", "Name_A": "GO:0007162", "target_original": "47", "Gene_B": "GO:0010810", "interaction": "Direct", "SUID": 579804, "TYPE": "Direct", "Gene_A": "GO:0007162", "BEND_MAP_ID": 579804, "selected": false}, "selected": false}, {"data": {"id": "579803", "source": "579313", "target": "579298", "SCORE": 0.3806078827461892, "shared_interaction": "Direct", "source_original": "29", "Name_B": "GO:1903707", "Name_A": "GO:0007162", "target_original": "44", "Gene_B": "GO:1903707", "interaction": "Direct", "SUID": 579803, "TYPE": "Direct", "Gene_A": "GO:0007162", "BEND_MAP_ID": 579803, "selected": false}, "selected": false}, {"data": {"id": "579802", "source": "579313", "target": "579294", "SCORE": 0.30518554199086184, "shared_interaction": "Direct", "source_original": "29", "Name_B": "GO:0031589", "Name_A": "GO:0007162", "target_original": "48", "Gene_B": "GO:0031589", "interaction": "Direct", "SUID": 579802, "TYPE": "Direct", "Gene_A": "GO:0007162", "BEND_MAP_ID": 579802, "selected": false}, "selected": false}, {"data": {"id": "579801", "source": "579313", "target": "579317", "SCORE": 0.34127915626665195, "shared_interaction": "Direct", "source_original": "29", "Name_B": "GO:1903706", "Name_A": "GO:0007162", "target_original": "25", "Gene_B": "GO:1903706", "interaction": "Direct", "SUID": 579801, "TYPE": "Direct", "Gene_A": "GO:0007162", "BEND_MAP_ID": 579801, "selected": false}, "selected": false}, {"data": {"id": "579800", "source": "579313", "target": "579316", "SCORE": 0.33803657444986634, "shared_interaction": "Direct", "source_original": "29", "Name_B": "GO:0030217", "Name_A": "GO:0007162", "target_original": "26", "Gene_B": "GO:0030217", "interaction": "Direct", "SUID": 579800, "TYPE": "Direct", "Gene_A": "GO:0007162", "BEND_MAP_ID": 579800, "selected": false}, "selected": false}, {"data": {"id": "579799", "source": "579313", "target": "579315", "SCORE": 0.33523661870713295, "shared_interaction": "Direct", "source_original": "29", "Name_B": "GO:0030098", "Name_A": "GO:0007162", "target_original": "27", "Gene_B": "GO:0030098", "interaction": "Direct", "SUID": 579799, "TYPE": "Direct", "Gene_A": "GO:0007162", "BEND_MAP_ID": 579799, "selected": false}, "selected": false}, {"data": {"id": "579798", "source": "579313", "target": "579312", "SCORE": 0.4320389372333766, "shared_interaction": "Direct", "source_original": "29", "Name_B": "GO:1902105", "Name_A": "GO:0007162", "target_original": "30", "Gene_B": "GO:1902105", "interaction": "Direct", "SUID": 579798, "TYPE": "Direct", "Gene_A": "GO:0007162", "BEND_MAP_ID": 579798, "selected": false}, "selected": false}, {"data": {"id": "579797", "source": "579313", "target": "579311", "SCORE": 0.7087627517660752, "shared_interaction": "Direct", "source_original": "29", "Name_B": "GO:1903038", "Name_A": "GO:0007162", "target_original": "31", "Gene_B": "GO:1903038", "interaction": "Direct", "SUID": 579797, "TYPE": "Direct", "Gene_A": "GO:0007162", "BEND_MAP_ID": 579797, "selected": false}, "selected": false}, {"data": {"id": "579796", "source": "579313", "target": "579310", "SCORE": 0.4550020150674871, "shared_interaction": "Direct", "source_original": "29", "Name_B": "GO:0045581", "Name_A": "GO:0007162", "target_original": "32", "Gene_B": "GO:0045581", "interaction": "Direct", "SUID": 579796, "TYPE": "Direct", "Gene_A": "GO:0007162", "BEND_MAP_ID": 579796, "selected": false}, "selected": false}, {"data": {"id": "579795", "source": "579312", "target": "579298", "SCORE": 0.5901702599726049, "shared_interaction": "Direct", "source_original": "30", "Name_B": "GO:1903707", "Name_A": "GO:1902105", "target_original": "44", "Gene_B": "GO:1903707", "interaction": "Direct", "SUID": 579795, "TYPE": "Direct", "Gene_A": "GO:1902105", "BEND_MAP_ID": 579795, "selected": false}, "selected": false}, {"data": {"id": "579794", "source": "579312", "target": "579317", "SCORE": 0.7634354342233752, "shared_interaction": "Direct", "source_original": "30", "Name_B": "GO:1903706", "Name_A": "GO:1902105", "target_original": "25", "Gene_B": "GO:1903706", "interaction": "Direct", "SUID": 579794, "TYPE": "Direct", "Gene_A": "GO:1902105", "BEND_MAP_ID": 579794, "selected": false}, "selected": false}, {"data": {"id": "579793", "source": "579312", "target": "579311", "SCORE": 0.4788443917433983, "shared_interaction": "Direct", "source_original": "30", "Name_B": "GO:1903038", "Name_A": "GO:1902105", "target_original": "31", "Gene_B": "GO:1903038", "interaction": "Direct", "SUID": 579793, "TYPE": "Direct", "Gene_A": "GO:1902105", "BEND_MAP_ID": 579793, "selected": false}, "selected": false}, {"data": {"id": "579792", "source": "579311", "target": "579298", "SCORE": 0.4514478282586155, "shared_interaction": "Direct", "source_original": "31", "Name_B": "GO:1903707", "Name_A": "GO:1903038", "target_original": "44", "Gene_B": "GO:1903707", "interaction": "Direct", "SUID": 579792, "TYPE": "Direct", "Gene_A": "GO:1903038", "BEND_MAP_ID": 579792, "selected": false}, "selected": false}, {"data": {"id": "579791", "source": "579311", "target": "579317", "SCORE": 0.3398031387128647, "shared_interaction": "Direct", "source_original": "31", "Name_B": "GO:1903706", "Name_A": "GO:1903038", "target_original": "25", "Gene_B": "GO:1903706", "interaction": "Direct", "SUID": 579791, "TYPE": "Direct", "Gene_A": "GO:1903038", "BEND_MAP_ID": 579791, "selected": false}, "selected": false}, {"data": {"id": "579790", "source": "579310", "target": "579298", "SCORE": 0.5710102489028728, "shared_interaction": "Direct", "source_original": "32", "Name_B": "GO:1903707", "Name_A": "GO:0045581", "target_original": "44", "Gene_B": "GO:1903707", "interaction": "Direct", "SUID": 579790, "TYPE": "Direct", "Gene_A": "GO:0045581", "BEND_MAP_ID": 579790, "selected": false}, "selected": false}, {"data": {"id": "579789", "source": "579310", "target": "579317", "SCORE": 0.32077658649755497, "shared_interaction": "Direct", "source_original": "32", "Name_B": "GO:1903706", "Name_A": "GO:0045581", "target_original": "25", "Gene_B": "GO:1903706", "interaction": "Direct", "SUID": 579789, "TYPE": "Direct", "Gene_A": "GO:0045581", "BEND_MAP_ID": 579789, "selected": false}, "selected": false}, {"data": {"id": "579788", "source": "579310", "target": "579312", "SCORE": 0.4725981164228384, "shared_interaction": "Direct", "source_original": "32", "Name_B": "GO:1902105", "Name_A": "GO:0045581", "target_original": "30", "Gene_B": "GO:1902105", "interaction": "Direct", "SUID": 579788, "TYPE": "Direct", "Gene_A": "GO:0045581", "BEND_MAP_ID": 579788, "selected": false}, "selected": false}, {"data": {"id": "579787", "source": "579310", "target": "579311", "SCORE": 0.6983621899867882, "shared_interaction": "Direct", "source_original": "32", "Name_B": "GO:1903038", "Name_A": "GO:0045581", "target_original": "31", "Gene_B": "GO:1903038", "interaction": "Direct", "SUID": 579787, "TYPE": "Direct", "Gene_A": "GO:0045581", "BEND_MAP_ID": 579787, "selected": false}, "selected": false}, {"data": {"id": "579786", "source": "579309", "target": "579308", "SCORE": 0.7342704908746389, "shared_interaction": "Direct", "source_original": "33", "Name_B": "GO:0045600", "Name_A": "GO:0045598", "target_original": "34", "Gene_B": "GO:0045600", "interaction": "Direct", "SUID": 579786, "TYPE": "Direct", "Gene_A": "GO:0045598", "BEND_MAP_ID": 579786, "selected": false}, "selected": false}, {"data": {"id": "579785", "source": "579307", "target": "579306", "SCORE": 0.5710162882044281, "shared_interaction": "Direct", "source_original": "35", "Name_B": "GO:0050878", "Name_A": "GO:0042060", "target_original": "36", "Gene_B": "GO:0050878", "interaction": "Direct", "SUID": 579785, "TYPE": "Direct", "Gene_A": "GO:0042060", "BEND_MAP_ID": 579785, "selected": false}, "selected": false}, {"data": {"id": "579784", "source": "579307", "target": "579301", "SCORE": 0.6334327783662379, "shared_interaction": "Direct", "source_original": "35", "Name_B": "GO:0050817", "Name_A": "GO:0042060", "target_original": "41", "Gene_B": "GO:0050817", "interaction": "Direct", "SUID": 579784, "TYPE": "Direct", "Gene_A": "GO:0042060", "BEND_MAP_ID": 579784, "selected": false}, "selected": false}, {"data": {"id": "579783", "source": "579307", "target": "579299", "SCORE": 0.4027558841120151, "shared_interaction": "Direct", "source_original": "35", "Name_B": "R-HSA-109582", "Name_A": "GO:0042060", "target_original": "43", "Gene_B": "R-HSA-109582", "interaction": "Direct", "SUID": 579783, "TYPE": "Direct", "Gene_A": "GO:0042060", "BEND_MAP_ID": 579783, "selected": false}, "selected": false}, {"data": {"id": "579782", "source": "579306", "target": "579305", "SCORE": 0.33338750304810383, "shared_interaction": "Direct", "source_original": "36", "Name_B": "GO:0070527", "Name_A": "GO:0050878", "target_original": "37", "Gene_B": "GO:0070527", "interaction": "Direct", "SUID": 579782, "TYPE": "Direct", "Gene_A": "GO:0050878", "BEND_MAP_ID": 579782, "selected": false}, "selected": false}, {"data": {"id": "579781", "source": "579306", "target": "579299", "SCORE": 0.4610146517787321, "shared_interaction": "Direct", "source_original": "36", "Name_B": "R-HSA-109582", "Name_A": "GO:0050878", "target_original": "43", "Gene_B": "R-HSA-109582", "interaction": "Direct", "SUID": 579781, "TYPE": "Direct", "Gene_A": "GO:0050878", "BEND_MAP_ID": 579781, "selected": false}, "selected": false}, {"data": {"id": "579780", "source": "579304", "target": "579198", "SCORE": 0.5566056591386517, "shared_interaction": "Direct", "source_original": "38", "Name_B": "GO:0009611", "Name_A": "GO:0007596", "target_original": "144", "Gene_B": "GO:0009611", "interaction": "Direct", "SUID": 579780, "TYPE": "Direct", "Gene_A": "GO:0007596", "BEND_MAP_ID": 579780, "selected": false}, "selected": false}, {"data": {"id": "579779", "source": "579304", "target": "579307", "SCORE": 0.6334327783662379, "shared_interaction": "Direct", "source_original": "38", "Name_B": "GO:0042060", "Name_A": "GO:0007596", "target_original": "35", "Gene_B": "GO:0042060", "interaction": "Direct", "SUID": 579779, "TYPE": "Direct", "Gene_A": "GO:0007596", "BEND_MAP_ID": 579779, "selected": false}, "selected": false}, {"data": {"id": "579778", "source": "579304", "target": "579306", "SCORE": 0.8017354615295075, "shared_interaction": "Direct", "source_original": "38", "Name_B": "GO:0050878", "Name_A": "GO:0007596", "target_original": "36", "Gene_B": "GO:0050878", "interaction": "Direct", "SUID": 579778, "TYPE": "Direct", "Gene_A": "GO:0007596", "BEND_MAP_ID": 579778, "selected": false}, "selected": false}, {"data": {"id": "579777", "source": "579304", "target": "579305", "SCORE": 0.4603256465052544, "shared_interaction": "Direct", "source_original": "38", "Name_B": "GO:0070527", "Name_A": "GO:0007596", "target_original": "37", "Gene_B": "GO:0070527", "interaction": "Direct", "SUID": 579777, "TYPE": "Direct", "Gene_A": "GO:0007596", "BEND_MAP_ID": 579777, "selected": false}, "selected": false}, {"data": {"id": "579776", "source": "579304", "target": "579303", "SCORE": 0.7078763708878713, "shared_interaction": "Direct", "source_original": "38", "Name_B": "GO:0030168", "Name_A": "GO:0007596", "target_original": "39", "Gene_B": "GO:0030168", "interaction": "Direct", "SUID": 579776, "TYPE": "Direct", "Gene_A": "GO:0007596", "BEND_MAP_ID": 579776, "selected": false}, "selected": false}, {"data": {"id": "579775", "source": "579304", "target": "579302", "SCORE": 1.0, "shared_interaction": "Direct", "source_original": "38", "Name_B": "GO:0007599", "Name_A": "GO:0007596", "target_original": "40", "Gene_B": "GO:0007599", "interaction": "Direct", "SUID": 579775, "TYPE": "Direct", "Gene_A": "GO:0007596", "BEND_MAP_ID": 579775, "selected": false}, "selected": false}, {"data": {"id": "579774", "source": "579304", "target": "579301", "SCORE": 1.0, "shared_interaction": "Direct", "source_original": "38", "Name_B": "GO:0050817", "Name_A": "GO:0007596", "target_original": "41", "Gene_B": "GO:0050817", "interaction": "Direct", "SUID": 579774, "TYPE": "Direct", "Gene_A": "GO:0007596", "BEND_MAP_ID": 579774, "selected": false}, "selected": false}, {"data": {"id": "579773", "source": "579304", "target": "579300", "SCORE": 0.4398498902511081, "shared_interaction": "Direct", "source_original": "38", "Name_B": "GO:0034109", "Name_A": "GO:0007596", "target_original": "42", "Gene_B": "GO:0034109", "interaction": "Direct", "SUID": 579773, "TYPE": "Direct", "Gene_A": "GO:0007596", "BEND_MAP_ID": 579773, "selected": false}, "selected": false}, {"data": {"id": "579772", "source": "579304", "target": "579299", "SCORE": 0.5355875915168545, "shared_interaction": "Direct", "source_original": "38", "Name_B": "R-HSA-109582", "Name_A": "GO:0007596", "target_original": "43", "Gene_B": "R-HSA-109582", "interaction": "Direct", "SUID": 579772, "TYPE": "Direct", "Gene_A": "GO:0007596", "BEND_MAP_ID": 579772, "selected": false}, "selected": false}, {"data": {"id": "579771", "source": "579303", "target": "579307", "SCORE": 0.40502122946344643, "shared_interaction": "Direct", "source_original": "39", "Name_B": "GO:0042060", "Name_A": "GO:0030168", "target_original": "35", "Gene_B": "GO:0042060", "interaction": "Direct", "SUID": 579771, "TYPE": "Direct", "Gene_A": "GO:0030168", "BEND_MAP_ID": 579771, "selected": false}, "selected": false}, {"data": {"id": "579770", "source": "579303", "target": "579306", "SCORE": 0.5364590824111216, "shared_interaction": "Direct", "source_original": "39", "Name_B": "GO:0050878", "Name_A": "GO:0030168", "target_original": "36", "Gene_B": "GO:0050878", "interaction": "Direct", "SUID": 579770, "TYPE": "Direct", "Gene_A": "GO:0030168", "BEND_MAP_ID": 579770, "selected": false}, "selected": false}, {"data": {"id": "579769", "source": "579303", "target": "579305", "SCORE": 0.7061183550661554, "shared_interaction": "Direct", "source_original": "39", "Name_B": "GO:0070527", "Name_A": "GO:0030168", "target_original": "37", "Gene_B": "GO:0070527", "interaction": "Direct", "SUID": 579769, "TYPE": "Direct", "Gene_A": "GO:0030168", "BEND_MAP_ID": 579769, "selected": false}, "selected": false}, {"data": {"id": "579768", "source": "579303", "target": "579301", "SCORE": 0.7078763708878713, "shared_interaction": "Direct", "source_original": "39", "Name_B": "GO:0050817", "Name_A": "GO:0030168", "target_original": "41", "Gene_B": "GO:0050817", "interaction": "Direct", "SUID": 579768, "TYPE": "Direct", "Gene_A": "GO:0030168", "BEND_MAP_ID": 579768, "selected": false}, "selected": false}, {"data": {"id": "579767", "source": "579303", "target": "579300", "SCORE": 0.662123912360825, "shared_interaction": "Direct", "source_original": "39", "Name_B": "GO:0034109", "Name_A": "GO:0030168", "target_original": "42", "Gene_B": "GO:0034109", "interaction": "Direct", "SUID": 579767, "TYPE": "Direct", "Gene_A": "GO:0030168", "BEND_MAP_ID": 579767, "selected": false}, "selected": false}, {"data": {"id": "579766", "source": "579303", "target": "579299", "SCORE": 0.34228248181823084, "shared_interaction": "Direct", "source_original": "39", "Name_B": "R-HSA-109582", "Name_A": "GO:0030168", "target_original": "43", "Gene_B": "R-HSA-109582", "interaction": "Direct", "SUID": 579766, "TYPE": "Direct", "Gene_A": "GO:0030168", "BEND_MAP_ID": 579766, "selected": false}, "selected": false}, {"data": {"id": "579765", "source": "579302", "target": "579198", "SCORE": 0.5566056591386517, "shared_interaction": "Direct", "source_original": "40", "Name_B": "GO:0009611", "Name_A": "GO:0007599", "target_original": "144", "Gene_B": "GO:0009611", "interaction": "Direct", "SUID": 579765, "TYPE": "Direct", "Gene_A": "GO:0007599", "BEND_MAP_ID": 579765, "selected": false}, "selected": false}, {"data": {"id": "579764", "source": "579302", "target": "579307", "SCORE": 0.6334327783662379, "shared_interaction": "Direct", "source_original": "40", "Name_B": "GO:0042060", "Name_A": "GO:0007599", "target_original": "35", "Gene_B": "GO:0042060", "interaction": "Direct", "SUID": 579764, "TYPE": "Direct", "Gene_A": "GO:0007599", "BEND_MAP_ID": 579764, "selected": false}, "selected": false}, {"data": {"id": "579763", "source": "579302", "target": "579306", "SCORE": 0.8017354615295075, "shared_interaction": "Direct", "source_original": "40", "Name_B": "GO:0050878", "Name_A": "GO:0007599", "target_original": "36", "Gene_B": "GO:0050878", "interaction": "Direct", "SUID": 579763, "TYPE": "Direct", "Gene_A": "GO:0007599", "BEND_MAP_ID": 579763, "selected": false}, "selected": false}, {"data": {"id": "579762", "source": "579302", "target": "579305", "SCORE": 0.4603256465052544, "shared_interaction": "Direct", "source_original": "40", "Name_B": "GO:0070527", "Name_A": "GO:0007599", "target_original": "37", "Gene_B": "GO:0070527", "interaction": "Direct", "SUID": 579762, "TYPE": "Direct", "Gene_A": "GO:0007599", "BEND_MAP_ID": 579762, "selected": false}, "selected": false}, {"data": {"id": "579761", "source": "579302", "target": "579303", "SCORE": 0.7078763708878713, "shared_interaction": "Direct", "source_original": "40", "Name_B": "GO:0030168", "Name_A": "GO:0007599", "target_original": "39", "Gene_B": "GO:0030168", "interaction": "Direct", "SUID": 579761, "TYPE": "Direct", "Gene_A": "GO:0007599", "BEND_MAP_ID": 579761, "selected": false}, "selected": false}, {"data": {"id": "579760", "source": "579302", "target": "579301", "SCORE": 1.0, "shared_interaction": "Direct", "source_original": "40", "Name_B": "GO:0050817", "Name_A": "GO:0007599", "target_original": "41", "Gene_B": "GO:0050817", "interaction": "Direct", "SUID": 579760, "TYPE": "Direct", "Gene_A": "GO:0007599", "BEND_MAP_ID": 579760, "selected": false}, "selected": false}, {"data": {"id": "579759", "source": "579302", "target": "579300", "SCORE": 0.4398498902511081, "shared_interaction": "Direct", "source_original": "40", "Name_B": "GO:0034109", "Name_A": "GO:0007599", "target_original": "42", "Gene_B": "GO:0034109", "interaction": "Direct", "SUID": 579759, "TYPE": "Direct", "Gene_A": "GO:0007599", "BEND_MAP_ID": 579759, "selected": false}, "selected": false}, {"data": {"id": "579758", "source": "579302", "target": "579299", "SCORE": 0.5355875915168545, "shared_interaction": "Direct", "source_original": "40", "Name_B": "R-HSA-109582", "Name_A": "GO:0007599", "target_original": "43", "Gene_B": "R-HSA-109582", "interaction": "Direct", "SUID": 579758, "TYPE": "Direct", "Gene_A": "GO:0007599", "BEND_MAP_ID": 579758, "selected": false}, "selected": false}, {"data": {"id": "579757", "source": "579301", "target": "579306", "SCORE": 0.8017354615295075, "shared_interaction": "Direct", "source_original": "41", "Name_B": "GO:0050878", "Name_A": "GO:0050817", "target_original": "36", "Gene_B": "GO:0050878", "interaction": "Direct", "SUID": 579757, "TYPE": "Direct", "Gene_A": "GO:0050817", "BEND_MAP_ID": 579757, "selected": false}, "selected": false}, {"data": {"id": "579756", "source": "579301", "target": "579305", "SCORE": 0.4603256465052544, "shared_interaction": "Direct", "source_original": "41", "Name_B": "GO:0070527", "Name_A": "GO:0050817", "target_original": "37", "Gene_B": "GO:0070527", "interaction": "Direct", "SUID": 579756, "TYPE": "Direct", "Gene_A": "GO:0050817", "BEND_MAP_ID": 579756, "selected": false}, "selected": false}, {"data": {"id": "579755", "source": "579301", "target": "579299", "SCORE": 0.5355875915168545, "shared_interaction": "Direct", "source_original": "41", "Name_B": "R-HSA-109582", "Name_A": "GO:0050817", "target_original": "43", "Gene_B": "R-HSA-109582", "interaction": "Direct", "SUID": 579755, "TYPE": "Direct", "Gene_A": "GO:0050817", "BEND_MAP_ID": 579755, "selected": false}, "selected": false}, {"data": {"id": "579754", "source": "579300", "target": "579306", "SCORE": 0.3212414773485611, "shared_interaction": "Direct", "source_original": "42", "Name_B": "GO:0050878", "Name_A": "GO:0034109", "target_original": "36", "Gene_B": "GO:0050878", "interaction": "Direct", "SUID": 579754, "TYPE": "Direct", "Gene_A": "GO:0034109", "BEND_MAP_ID": 579754, "selected": false}, "selected": false}, {"data": {"id": "579753", "source": "579300", "target": "579305", "SCORE": 0.9158106457872947, "shared_interaction": "Direct", "source_original": "42", "Name_B": "GO:0070527", "Name_A": "GO:0034109", "target_original": "37", "Gene_B": "GO:0070527", "interaction": "Direct", "SUID": 579753, "TYPE": "Direct", "Gene_A": "GO:0034109", "BEND_MAP_ID": 579753, "selected": false}, "selected": false}, {"data": {"id": "579752", "source": "579300", "target": "579301", "SCORE": 0.4398498902511081, "shared_interaction": "Direct", "source_original": "42", "Name_B": "GO:0050817", "Name_A": "GO:0034109", "target_original": "41", "Gene_B": "GO:0050817", "interaction": "Direct", "SUID": 579752, "TYPE": "Direct", "Gene_A": "GO:0034109", "BEND_MAP_ID": 579752, "selected": false}, "selected": false}, {"data": {"id": "579751", "source": "579297", "target": "579318", "SCORE": 0.37541785766237873, "shared_interaction": "Direct", "source_original": "45", "Name_B": "GO:0030155", "Name_A": "GO:0002683", "target_original": "24", "Gene_B": "GO:0030155", "interaction": "Direct", "SUID": 579751, "TYPE": "Direct", "Gene_A": "GO:0002683", "BEND_MAP_ID": 579751, "selected": false}, "selected": false}, {"data": {"id": "579750", "source": "579297", "target": "579204", "SCORE": 0.4219945809767772, "shared_interaction": "Direct", "source_original": "45", "Name_B": "GO:0045596", "Name_A": "GO:0002683", "target_original": "138", "Gene_B": "GO:0045596", "interaction": "Direct", "SUID": 579750, "TYPE": "Direct", "Gene_A": "GO:0002683", "BEND_MAP_ID": 579750, "selected": false}, "selected": false}, {"data": {"id": "579749", "source": "579297", "target": "579298", "SCORE": 0.7005738542570857, "shared_interaction": "Direct", "source_original": "45", "Name_B": "GO:1903707", "Name_A": "GO:0002683", "target_original": "44", "Gene_B": "GO:1903707", "interaction": "Direct", "SUID": 579749, "TYPE": "Direct", "Gene_A": "GO:0002683", "BEND_MAP_ID": 579749, "selected": false}, "selected": false}, {"data": {"id": "579748", "source": "579297", "target": "579205", "SCORE": 0.3574952792842431, "shared_interaction": "Direct", "source_original": "45", "Name_B": "GO:0030099", "Name_A": "GO:0002683", "target_original": "137", "Gene_B": "GO:0030099", "interaction": "Direct", "SUID": 579748, "TYPE": "Direct", "Gene_A": "GO:0002683", "BEND_MAP_ID": 579748, "selected": false}, "selected": false}, {"data": {"id": "579747", "source": "579297", "target": "579317", "SCORE": 0.5453241201620109, "shared_interaction": "Direct", "source_original": "45", "Name_B": "GO:1903706", "Name_A": "GO:0002683", "target_original": "25", "Gene_B": "GO:1903706", "interaction": "Direct", "SUID": 579747, "TYPE": "Direct", "Gene_A": "GO:0002683", "BEND_MAP_ID": 579747, "selected": false}, "selected": false}, {"data": {"id": "579746", "source": "579297", "target": "579316", "SCORE": 0.36198011178118245, "shared_interaction": "Direct", "source_original": "45", "Name_B": "GO:0030217", "Name_A": "GO:0002683", "target_original": "26", "Gene_B": "GO:0030217", "interaction": "Direct", "SUID": 579746, "TYPE": "Direct", "Gene_A": "GO:0002683", "BEND_MAP_ID": 579746, "selected": false}, "selected": false}, {"data": {"id": "579745", "source": "579297", "target": "579315", "SCORE": 0.37471271915484733, "shared_interaction": "Direct", "source_original": "45", "Name_B": "GO:0030098", "Name_A": "GO:0002683", "target_original": "27", "Gene_B": "GO:0030098", "interaction": "Direct", "SUID": 579745, "TYPE": "Direct", "Gene_A": "GO:0002683", "BEND_MAP_ID": 579745, "selected": false}, "selected": false}, {"data": {"id": "579744", "source": "579297", "target": "579314", "SCORE": 0.3398806879488279, "shared_interaction": "Direct", "source_original": "45", "Name_B": "GO:0045785", "Name_A": "GO:0002683", "target_original": "28", "Gene_B": "GO:0045785", "interaction": "Direct", "SUID": 579744, "TYPE": "Direct", "Gene_A": "GO:0002683", "BEND_MAP_ID": 579744, "selected": false}, "selected": false}, {"data": {"id": "579743", "source": "579297", "target": "579313", "SCORE": 0.4728356734641735, "shared_interaction": "Direct", "source_original": "45", "Name_B": "GO:0007162", "Name_A": "GO:0002683", "target_original": "29", "Gene_B": "GO:0007162", "interaction": "Direct", "SUID": 579743, "TYPE": "Direct", "Gene_A": "GO:0002683", "BEND_MAP_ID": 579743, "selected": false}, "selected": false}, {"data": {"id": "579742", "source": "579297", "target": "579312", "SCORE": 0.48389531957779564, "shared_interaction": "Direct", "source_original": "45", "Name_B": "GO:1902105", "Name_A": "GO:0002683", "target_original": "30", "Gene_B": "GO:1902105", "interaction": "Direct", "SUID": 579742, "TYPE": "Direct", "Gene_A": "GO:0002683", "BEND_MAP_ID": 579742, "selected": false}, "selected": false}, {"data": {"id": "579741", "source": "579297", "target": "579311", "SCORE": 0.5227603943426484, "shared_interaction": "Direct", "source_original": "45", "Name_B": "GO:1903038", "Name_A": "GO:0002683", "target_original": "31", "Gene_B": "GO:1903038", "interaction": "Direct", "SUID": 579741, "TYPE": "Direct", "Gene_A": "GO:0002683", "BEND_MAP_ID": 579741, "selected": false}, "selected": false}, {"data": {"id": "579740", "source": "579297", "target": "579310", "SCORE": 0.35449916772833173, "shared_interaction": "Direct", "source_original": "45", "Name_B": "GO:0045581", "Name_A": "GO:0002683", "target_original": "32", "Gene_B": "GO:0045581", "interaction": "Direct", "SUID": 579740, "TYPE": "Direct", "Gene_A": "GO:0002683", "BEND_MAP_ID": 579740, "selected": false}, "selected": false}, {"data": {"id": "579739", "source": "579297", "target": "579259", "SCORE": 0.3807834553558061, "shared_interaction": "Direct", "source_original": "45", "Name_B": "GO:0045637", "Name_A": "GO:0002683", "target_original": "83", "Gene_B": "GO:0045637", "interaction": "Direct", "SUID": 579739, "TYPE": "Direct", "Gene_A": "GO:0002683", "BEND_MAP_ID": 579739, "selected": false}, "selected": false}, {"data": {"id": "579738", "source": "579297", "target": "579296", "SCORE": 0.39873055263995555, "shared_interaction": "Direct", "source_original": "45", "Name_B": "GO:0045638", "Name_A": "GO:0002683", "target_original": "46", "Gene_B": "GO:0045638", "interaction": "Direct", "SUID": 579738, "TYPE": "Direct", "Gene_A": "GO:0002683", "BEND_MAP_ID": 579738, "selected": false}, "selected": false}, {"data": {"id": "579737", "source": "579296", "target": "579298", "SCORE": 0.6318823442473412, "shared_interaction": "Direct", "source_original": "46", "Name_B": "GO:1903707", "Name_A": "GO:0045638", "target_original": "44", "Gene_B": "GO:1903707", "interaction": "Direct", "SUID": 579737, "TYPE": "Direct", "Gene_A": "GO:0045638", "BEND_MAP_ID": 579737, "selected": false}, "selected": false}, {"data": {"id": "579736", "source": "579296", "target": "579317", "SCORE": 0.3617253022554717, "shared_interaction": "Direct", "source_original": "46", "Name_B": "GO:1903706", "Name_A": "GO:0045638", "target_original": "25", "Gene_B": "GO:1903706", "interaction": "Direct", "SUID": 579736, "TYPE": "Direct", "Gene_A": "GO:0045638", "BEND_MAP_ID": 579736, "selected": false}, "selected": false}, {"data": {"id": "579735", "source": "579295", "target": "579318", "SCORE": 0.57558282543968, "shared_interaction": "Direct", "source_original": "47", "Name_B": "GO:0030155", "Name_A": "GO:0010810", "target_original": "24", "Gene_B": "GO:0030155", "interaction": "Direct", "SUID": 579735, "TYPE": "Direct", "Gene_A": "GO:0010810", "BEND_MAP_ID": 579735, "selected": false}, "selected": false}, {"data": {"id": "579734", "source": "579295", "target": "579214", "SCORE": 0.33809300541406523, "shared_interaction": "Direct", "source_original": "47", "Name_B": "GO:0034329", "Name_A": "GO:0010810", "target_original": "128", "Gene_B": "GO:0034329", "interaction": "Direct", "SUID": 579734, "TYPE": "Direct", "Gene_A": "GO:0010810", "BEND_MAP_ID": 579734, "selected": false}, "selected": false}, {"data": {"id": "579733", "source": "579295", "target": "579294", "SCORE": 0.8417590895941608, "shared_interaction": "Direct", "source_original": "47", "Name_B": "GO:0031589", "Name_A": "GO:0010810", "target_original": "48", "Gene_B": "GO:0031589", "interaction": "Direct", "SUID": 579733, "TYPE": "Direct", "Gene_A": "GO:0010810", "BEND_MAP_ID": 579733, "selected": false}, "selected": false}, {"data": {"id": "579732", "source": "579295", "target": "579293", "SCORE": 0.3242592096872016, "shared_interaction": "Direct", "source_original": "47", "Name_B": "GO:0034330", "Name_A": "GO:0010810", "target_original": "49", "Gene_B": "GO:0034330", "interaction": "Direct", "SUID": 579732, "TYPE": "Direct", "Gene_A": "GO:0010810", "BEND_MAP_ID": 579732, "selected": false}, "selected": false}, {"data": {"id": "579731", "source": "579295", "target": "579314", "SCORE": 0.468671489828504, "shared_interaction": "Direct", "source_original": "47", "Name_B": "GO:0045785", "Name_A": "GO:0010810", "target_original": "28", "Gene_B": "GO:0045785", "interaction": "Direct", "SUID": 579731, "TYPE": "Direct", "Gene_A": "GO:0010810", "BEND_MAP_ID": 579731, "selected": false}, "selected": false}, {"data": {"id": "579730", "source": "579295", "target": "579289", "SCORE": 0.7144576931901154, "shared_interaction": "Direct", "source_original": "47", "Name_B": "GO:0010811", "Name_A": "GO:0010810", "target_original": "53", "Gene_B": "GO:0010811", "interaction": "Direct", "SUID": 579730, "TYPE": "Direct", "Gene_A": "GO:0010810", "BEND_MAP_ID": 579730, "selected": false}, "selected": false}, {"data": {"id": "579729", "source": "579295", "target": "579288", "SCORE": 0.36547658435459507, "shared_interaction": "Direct", "source_original": "47", "Name_B": "GO:1901888", "Name_A": "GO:0010810", "target_original": "54", "Gene_B": "GO:1901888", "interaction": "Direct", "SUID": 579729, "TYPE": "Direct", "Gene_A": "GO:0010810", "BEND_MAP_ID": 579729, "selected": false}, "selected": false}, {"data": {"id": "579728", "source": "579294", "target": "579214", "SCORE": 0.3872317006060524, "shared_interaction": "Direct", "source_original": "48", "Name_B": "GO:0034329", "Name_A": "GO:0031589", "target_original": "128", "Gene_B": "GO:0034329", "interaction": "Direct", "SUID": 579728, "TYPE": "Direct", "Gene_A": "GO:0031589", "BEND_MAP_ID": 579728, "selected": false}, "selected": false}, {"data": {"id": "579727", "source": "579294", "target": "579293", "SCORE": 0.37336256215645863, "shared_interaction": "Direct", "source_original": "48", "Name_B": "GO:0034330", "Name_A": "GO:0031589", "target_original": "49", "Gene_B": "GO:0034330", "interaction": "Direct", "SUID": 579727, "TYPE": "Direct", "Gene_A": "GO:0031589", "BEND_MAP_ID": 579727, "selected": false}, "selected": false}, {"data": {"id": "579726", "source": "579294", "target": "579314", "SCORE": 0.39820780812673306, "shared_interaction": "Direct", "source_original": "48", "Name_B": "GO:0045785", "Name_A": "GO:0031589", "target_original": "28", "Gene_B": "GO:0045785", "interaction": "Direct", "SUID": 579726, "TYPE": "Direct", "Gene_A": "GO:0031589", "BEND_MAP_ID": 579726, "selected": false}, "selected": false}, {"data": {"id": "579725", "source": "579293", "target": "579288", "SCORE": 0.5657306195914944, "shared_interaction": "Direct", "source_original": "49", "Name_B": "GO:1901888", "Name_A": "GO:0034330", "target_original": "54", "Gene_B": "GO:1901888", "interaction": "Direct", "SUID": 579725, "TYPE": "Direct", "Gene_A": "GO:0034330", "BEND_MAP_ID": 579725, "selected": false}, "selected": false}, {"data": {"id": "579724", "source": "579292", "target": "579318", "SCORE": 0.3941126383657026, "shared_interaction": "Direct", "source_original": "50", "Name_B": "GO:0030155", "Name_A": "GO:0001952", "target_original": "24", "Gene_B": "GO:0030155", "interaction": "Direct", "SUID": 579724, "TYPE": "Direct", "Gene_A": "GO:0001952", "BEND_MAP_ID": 579724, "selected": false}, "selected": false}, {"data": {"id": "579723", "source": "579292", "target": "579214", "SCORE": 0.3655392864517741, "shared_interaction": "Direct", "source_original": "50", "Name_B": "GO:0034329", "Name_A": "GO:0001952", "target_original": "128", "Gene_B": "GO:0034329", "interaction": "Direct", "SUID": 579723, "TYPE": "Direct", "Gene_A": "GO:0001952", "BEND_MAP_ID": 579723, "selected": false}, "selected": false}, {"data": {"id": "579722", "source": "579292", "target": "579295", "SCORE": 0.7557111889281954, "shared_interaction": "Direct", "source_original": "50", "Name_B": "GO:0010810", "Name_A": "GO:0001952", "target_original": "47", "Gene_B": "GO:0010810", "interaction": "Direct", "SUID": 579722, "TYPE": "Direct", "Gene_A": "GO:0001952", "BEND_MAP_ID": 579722, "selected": false}, "selected": false}, {"data": {"id": "579721", "source": "579292", "target": "579294", "SCORE": 0.6124515393391724, "shared_interaction": "Direct", "source_original": "50", "Name_B": "GO:0031589", "Name_A": "GO:0001952", "target_original": "48", "Gene_B": "GO:0031589", "interaction": "Direct", "SUID": 579721, "TYPE": "Direct", "Gene_A": "GO:0001952", "BEND_MAP_ID": 579721, "selected": false}, "selected": false}, {"data": {"id": "579720", "source": "579292", "target": "579293", "SCORE": 0.3485350066712331, "shared_interaction": "Direct", "source_original": "50", "Name_B": "GO:0034330", "Name_A": "GO:0001952", "target_original": "49", "Gene_B": "GO:0034330", "interaction": "Direct", "SUID": 579720, "TYPE": "Direct", "Gene_A": "GO:0001952", "BEND_MAP_ID": 579720, "selected": false}, "selected": false}, {"data": {"id": "579719", "source": "579292", "target": "579314", "SCORE": 0.361542224397149, "shared_interaction": "Direct", "source_original": "50", "Name_B": "GO:0045785", "Name_A": "GO:0001952", "target_original": "28", "Gene_B": "GO:0045785", "interaction": "Direct", "SUID": 579719, "TYPE": "Direct", "Gene_A": "GO:0001952", "BEND_MAP_ID": 579719, "selected": false}, "selected": false}, {"data": {"id": "579718", "source": "579292", "target": "579291", "SCORE": 0.8089168198260271, "shared_interaction": "Direct", "source_original": "50", "Name_B": "GO:0007160", "Name_A": "GO:0001952", "target_original": "51", "Gene_B": "GO:0007160", "interaction": "Direct", "SUID": 579718, "TYPE": "Direct", "Gene_A": "GO:0001952", "BEND_MAP_ID": 579718, "selected": false}, "selected": false}, {"data": {"id": "579717", "source": "579292", "target": "579290", "SCORE": 0.7328105051985976, "shared_interaction": "Direct", "source_original": "50", "Name_B": "GO:0001954", "Name_A": "GO:0001952", "target_original": "52", "Gene_B": "GO:0001954", "interaction": "Direct", "SUID": 579717, "TYPE": "Direct", "Gene_A": "GO:0001952", "BEND_MAP_ID": 579717, "selected": false}, "selected": false}, {"data": {"id": "579716", "source": "579292", "target": "579289", "SCORE": 0.6008778568194112, "shared_interaction": "Direct", "source_original": "50", "Name_B": "GO:0010811", "Name_A": "GO:0001952", "target_original": "53", "Gene_B": "GO:0010811", "interaction": "Direct", "SUID": 579716, "TYPE": "Direct", "Gene_A": "GO:0001952", "BEND_MAP_ID": 579716, "selected": false}, "selected": false}, {"data": {"id": "579715", "source": "579292", "target": "579288", "SCORE": 0.5026271117948884, "shared_interaction": "Direct", "source_original": "50", "Name_B": "GO:1901888", "Name_A": "GO:0001952", "target_original": "54", "Gene_B": "GO:1901888", "interaction": "Direct", "SUID": 579715, "TYPE": "Direct", "Gene_A": "GO:0001952", "BEND_MAP_ID": 579715, "selected": false}, "selected": false}, {"data": {"id": "579714", "source": "579292", "target": "579287", "SCORE": 0.5026271117948884, "shared_interaction": "Direct", "source_original": "50", "Name_B": "GO:0007044", "Name_A": "GO:0001952", "target_original": "55", "Gene_B": "GO:0007044", "interaction": "Direct", "SUID": 579714, "TYPE": "Direct", "Gene_A": "GO:0001952", "BEND_MAP_ID": 579714, "selected": false}, "selected": false}, {"data": {"id": "579713", "source": "579291", "target": "579318", "SCORE": 0.38274205780787274, "shared_interaction": "Direct", "source_original": "51", "Name_B": "GO:0030155", "Name_A": "GO:0007160", "target_original": "24", "Gene_B": "GO:0030155", "interaction": "Direct", "SUID": 579713, "TYPE": "Direct", "Gene_A": "GO:0007160", "BEND_MAP_ID": 579713, "selected": false}, "selected": false}, {"data": {"id": "579712", "source": "579291", "target": "579214", "SCORE": 0.4107789247896167, "shared_interaction": "Direct", "source_original": "51", "Name_B": "GO:0034329", "Name_A": "GO:0007160", "target_original": "128", "Gene_B": "GO:0034329", "interaction": "Direct", "SUID": 579712, "TYPE": "Direct", "Gene_A": "GO:0007160", "BEND_MAP_ID": 579712, "selected": false}, "selected": false}, {"data": {"id": "579711", "source": "579291", "target": "579295", "SCORE": 0.6651348841643759, "shared_interaction": "Direct", "source_original": "51", "Name_B": "GO:0010810", "Name_A": "GO:0007160", "target_original": "47", "Gene_B": "GO:0010810", "interaction": "Direct", "SUID": 579711, "TYPE": "Direct", "Gene_A": "GO:0007160", "BEND_MAP_ID": 579711, "selected": false}, "selected": false}, {"data": {"id": "579710", "source": "579291", "target": "579294", "SCORE": 0.787822098847634, "shared_interaction": "Direct", "source_original": "51", "Name_B": "GO:0031589", "Name_A": "GO:0007160", "target_original": "48", "Gene_B": "GO:0031589", "interaction": "Direct", "SUID": 579710, "TYPE": "Direct", "Gene_A": "GO:0007160", "BEND_MAP_ID": 579710, "selected": false}, "selected": false}, {"data": {"id": "579709", "source": "579291", "target": "579293", "SCORE": 0.39388454328087347, "shared_interaction": "Direct", "source_original": "51", "Name_B": "GO:0034330", "Name_A": "GO:0007160", "target_original": "49", "Gene_B": "GO:0034330", "interaction": "Direct", "SUID": 579709, "TYPE": "Direct", "Gene_A": "GO:0007160", "BEND_MAP_ID": 579709, "selected": false}, "selected": false}, {"data": {"id": "579708", "source": "579291", "target": "579314", "SCORE": 0.3311423792195066, "shared_interaction": "Direct", "source_original": "51", "Name_B": "GO:0045785", "Name_A": "GO:0007160", "target_original": "28", "Gene_B": "GO:0045785", "interaction": "Direct", "SUID": 579708, "TYPE": "Direct", "Gene_A": "GO:0007160", "BEND_MAP_ID": 579708, "selected": false}, "selected": false}, {"data": {"id": "579707", "source": "579291", "target": "579289", "SCORE": 0.5151385143426218, "shared_interaction": "Direct", "source_original": "51", "Name_B": "GO:0010811", "Name_A": "GO:0007160", "target_original": "53", "Gene_B": "GO:0010811", "interaction": "Direct", "SUID": 579707, "TYPE": "Direct", "Gene_A": "GO:0007160", "BEND_MAP_ID": 579707, "selected": false}, "selected": false}, {"data": {"id": "579706", "source": "579291", "target": "579288", "SCORE": 0.3946867232652926, "shared_interaction": "Direct", "source_original": "51", "Name_B": "GO:1901888", "Name_A": "GO:0007160", "target_original": "54", "Gene_B": "GO:1901888", "interaction": "Direct", "SUID": 579706, "TYPE": "Direct", "Gene_A": "GO:0007160", "BEND_MAP_ID": 579706, "selected": false}, "selected": false}, {"data": {"id": "579705", "source": "579290", "target": "579295", "SCORE": 0.5198610382809422, "shared_interaction": "Direct", "source_original": "52", "Name_B": "GO:0010810", "Name_A": "GO:0001954", "target_original": "47", "Gene_B": "GO:0010810", "interaction": "Direct", "SUID": 579705, "TYPE": "Direct", "Gene_A": "GO:0001954", "BEND_MAP_ID": 579705, "selected": false}, "selected": false}, {"data": {"id": "579704", "source": "579290", "target": "579294", "SCORE": 0.40669781931554405, "shared_interaction": "Direct", "source_original": "52", "Name_B": "GO:0031589", "Name_A": "GO:0001954", "target_original": "48", "Gene_B": "GO:0031589", "interaction": "Direct", "SUID": 579704, "TYPE": "Direct", "Gene_A": "GO:0001954", "BEND_MAP_ID": 579704, "selected": false}, "selected": false}, {"data": {"id": "579703", "source": "579290", "target": "579314", "SCORE": 0.394261799113441, "shared_interaction": "Direct", "source_original": "52", "Name_B": "GO:0045785", "Name_A": "GO:0001954", "target_original": "28", "Gene_B": "GO:0045785", "interaction": "Direct", "SUID": 579703, "TYPE": "Direct", "Gene_A": "GO:0001954", "BEND_MAP_ID": 579703, "selected": false}, "selected": false}, {"data": {"id": "579702", "source": "579290", "target": "579291", "SCORE": 0.5639880952389998, "shared_interaction": "Direct", "source_original": "52", "Name_B": "GO:0007160", "Name_A": "GO:0001954", "target_original": "51", "Gene_B": "GO:0007160", "interaction": "Direct", "SUID": 579702, "TYPE": "Direct", "Gene_A": "GO:0001954", "BEND_MAP_ID": 579702, "selected": false}, "selected": false}, {"data": {"id": "579701", "source": "579290", "target": "579289", "SCORE": 0.7744852799698488, "shared_interaction": "Direct", "source_original": "52", "Name_B": "GO:0010811", "Name_A": "GO:0001954", "target_original": "53", "Gene_B": "GO:0010811", "interaction": "Direct", "SUID": 579701, "TYPE": "Direct", "Gene_A": "GO:0001954", "BEND_MAP_ID": 579701, "selected": false}, "selected": false}, {"data": {"id": "579700", "source": "579289", "target": "579318", "SCORE": 0.36677987033092163, "shared_interaction": "Direct", "source_original": "53", "Name_B": "GO:0030155", "Name_A": "GO:0010811", "target_original": "24", "Gene_B": "GO:0030155", "interaction": "Direct", "SUID": 579700, "TYPE": "Direct", "Gene_A": "GO:0010811", "BEND_MAP_ID": 579700, "selected": false}, "selected": false}, {"data": {"id": "579699", "source": "579289", "target": "579294", "SCORE": 0.575402019447103, "shared_interaction": "Direct", "source_original": "53", "Name_B": "GO:0031589", "Name_A": "GO:0010811", "target_original": "48", "Gene_B": "GO:0031589", "interaction": "Direct", "SUID": 579699, "TYPE": "Direct", "Gene_A": "GO:0010811", "BEND_MAP_ID": 579699, "selected": false}, "selected": false}, {"data": {"id": "579698", "source": "579289", "target": "579314", "SCORE": 0.559619238477541, "shared_interaction": "Direct", "source_original": "53", "Name_B": "GO:0045785", "Name_A": "GO:0010811", "target_original": "28", "Gene_B": "GO:0045785", "interaction": "Direct", "SUID": 579698, "TYPE": "Direct", "Gene_A": "GO:0010811", "BEND_MAP_ID": 579698, "selected": false}, "selected": false}, {"data": {"id": "579697", "source": "579287", "target": "579214", "SCORE": 0.5939803780284855, "shared_interaction": "Direct", "source_original": "55", "Name_B": "GO:0034329", "Name_A": "GO:0007044", "target_original": "128", "Gene_B": "GO:0034329", "interaction": "Direct", "SUID": 579697, "TYPE": "Direct", "Gene_A": "GO:0007044", "BEND_MAP_ID": 579697, "selected": false}, "selected": false}, {"data": {"id": "579696", "source": "579287", "target": "579295", "SCORE": 0.36547658435459507, "shared_interaction": "Direct", "source_original": "55", "Name_B": "GO:0010810", "Name_A": "GO:0007044", "target_original": "47", "Gene_B": "GO:0010810", "interaction": "Direct", "SUID": 579696, "TYPE": "Direct", "Gene_A": "GO:0007044", "BEND_MAP_ID": 579696, "selected": false}, "selected": false}, {"data": {"id": "579695", "source": "579287", "target": "579294", "SCORE": 0.3739468253013651, "shared_interaction": "Direct", "source_original": "55", "Name_B": "GO:0031589", "Name_A": "GO:0007044", "target_original": "48", "Gene_B": "GO:0031589", "interaction": "Direct", "SUID": 579695, "TYPE": "Direct", "Gene_A": "GO:0007044", "BEND_MAP_ID": 579695, "selected": false}, "selected": false}, {"data": {"id": "579694", "source": "579287", "target": "579293", "SCORE": 0.5657306195914944, "shared_interaction": "Direct", "source_original": "55", "Name_B": "GO:0034330", "Name_A": "GO:0007044", "target_original": "49", "Gene_B": "GO:0034330", "interaction": "Direct", "SUID": 579694, "TYPE": "Direct", "Gene_A": "GO:0007044", "BEND_MAP_ID": 579694, "selected": false}, "selected": false}, {"data": {"id": "579693", "source": "579287", "target": "579291", "SCORE": 0.5081829626530502, "shared_interaction": "Direct", "source_original": "55", "Name_B": "GO:0007160", "Name_A": "GO:0007044", "target_original": "51", "Gene_B": "GO:0007160", "interaction": "Direct", "SUID": 579693, "TYPE": "Direct", "Gene_A": "GO:0007044", "BEND_MAP_ID": 579693, "selected": false}, "selected": false}, {"data": {"id": "579692", "source": "579287", "target": "579288", "SCORE": 0.5720089469130903, "shared_interaction": "Direct", "source_original": "55", "Name_B": "GO:1901888", "Name_A": "GO:0007044", "target_original": "54", "Gene_B": "GO:1901888", "interaction": "Direct", "SUID": 579692, "TYPE": "Direct", "Gene_A": "GO:0007044", "BEND_MAP_ID": 579692, "selected": false}, "selected": false}, {"data": {"id": "579691", "source": "579286", "target": "579206", "SCORE": 0.39653988740943935, "shared_interaction": "Direct", "source_original": "56", "Name_B": "GO:0061061", "Name_A": "GO:0007507", "target_original": "136", "Gene_B": "GO:0061061", "interaction": "Direct", "SUID": 579691, "TYPE": "Direct", "Gene_A": "GO:0007507", "BEND_MAP_ID": 579691, "selected": false}, "selected": false}, {"data": {"id": "579690", "source": "579286", "target": "579285", "SCORE": 0.3433475177310426, "shared_interaction": "Direct", "source_original": "56", "Name_B": "GO:0042692", "Name_A": "GO:0007507", "target_original": "57", "Gene_B": "GO:0042692", "interaction": "Direct", "SUID": 579690, "TYPE": "Direct", "Gene_A": "GO:0007507", "BEND_MAP_ID": 579690, "selected": false}, "selected": false}, {"data": {"id": "579689", "source": "579286", "target": "579284", "SCORE": 0.32078190589624445, "shared_interaction": "Direct", "source_original": "56", "Name_B": "GO:0051146", "Name_A": "GO:0007507", "target_original": "58", "Gene_B": "GO:0051146", "interaction": "Direct", "SUID": 579689, "TYPE": "Direct", "Gene_A": "GO:0007507", "BEND_MAP_ID": 579689, "selected": false}, "selected": false}, {"data": {"id": "579688", "source": "579286", "target": "579283", "SCORE": 0.4606731724424649, "shared_interaction": "Direct", "source_original": "56", "Name_B": "GO:0060537", "Name_A": "GO:0007507", "target_original": "59", "Gene_B": "GO:0060537", "interaction": "Direct", "SUID": 579688, "TYPE": "Direct", "Gene_A": "GO:0007507", "BEND_MAP_ID": 579688, "selected": false}, "selected": false}, {"data": {"id": "579687", "source": "579286", "target": "579281", "SCORE": 0.46168905487668715, "shared_interaction": "Direct", "source_original": "56", "Name_B": "GO:0014706", "Name_A": "GO:0007507", "target_original": "61", "Gene_B": "GO:0014706", "interaction": "Direct", "SUID": 579687, "TYPE": "Direct", "Gene_A": "GO:0007507", "BEND_MAP_ID": 579687, "selected": false}, "selected": false}, {"data": {"id": "579686", "source": "579286", "target": "579278", "SCORE": 0.4260847069264851, "shared_interaction": "Direct", "source_original": "56", "Name_B": "GO:0035051", "Name_A": "GO:0007507", "target_original": "64", "Gene_B": "GO:0035051", "interaction": "Direct", "SUID": 579686, "TYPE": "Direct", "Gene_A": "GO:0007507", "BEND_MAP_ID": 579686, "selected": false}, "selected": false}, {"data": {"id": "579685", "source": "579285", "target": "579206", "SCORE": 0.7485174781525012, "shared_interaction": "Direct", "source_original": "57", "Name_B": "GO:0061061", "Name_A": "GO:0042692", "target_original": "136", "Gene_B": "GO:0061061", "interaction": "Direct", "SUID": 579685, "TYPE": "Direct", "Gene_A": "GO:0042692", "BEND_MAP_ID": 579685, "selected": false}, "selected": false}, {"data": {"id": "579684", "source": "579285", "target": "579284", "SCORE": 0.8545572568637315, "shared_interaction": "Direct", "source_original": "57", "Name_B": "GO:0051146", "Name_A": "GO:0042692", "target_original": "58", "Gene_B": "GO:0051146", "interaction": "Direct", "SUID": 579684, "TYPE": "Direct", "Gene_A": "GO:0042692", "BEND_MAP_ID": 579684, "selected": false}, "selected": false}, {"data": {"id": "579683", "source": "579285", "target": "579283", "SCORE": 0.46703979011860436, "shared_interaction": "Direct", "source_original": "57", "Name_B": "GO:0060537", "Name_A": "GO:0042692", "target_original": "59", "Gene_B": "GO:0060537", "interaction": "Direct", "SUID": 579683, "TYPE": "Direct", "Gene_A": "GO:0042692", "BEND_MAP_ID": 579683, "selected": false}, "selected": false}, {"data": {"id": "579682", "source": "579284", "target": "579206", "SCORE": 0.6170805159387243, "shared_interaction": "Direct", "source_original": "58", "Name_B": "GO:0061061", "Name_A": "GO:0051146", "target_original": "136", "Gene_B": "GO:0061061", "interaction": "Direct", "SUID": 579682, "TYPE": "Direct", "Gene_A": "GO:0051146", "BEND_MAP_ID": 579682, "selected": false}, "selected": false}, {"data": {"id": "579681", "source": "579284", "target": "579283", "SCORE": 0.5446866159307221, "shared_interaction": "Direct", "source_original": "58", "Name_B": "GO:0060537", "Name_A": "GO:0051146", "target_original": "59", "Gene_B": "GO:0060537", "interaction": "Direct", "SUID": 579681, "TYPE": "Direct", "Gene_A": "GO:0051146", "BEND_MAP_ID": 579681, "selected": false}, "selected": false}, {"data": {"id": "579680", "source": "579283", "target": "579206", "SCORE": 0.6166150395217697, "shared_interaction": "Direct", "source_original": "59", "Name_B": "GO:0061061", "Name_A": "GO:0060537", "target_original": "136", "Gene_B": "GO:0061061", "interaction": "Direct", "SUID": 579680, "TYPE": "Direct", "Gene_A": "GO:0060537", "BEND_MAP_ID": 579680, "selected": false}, "selected": false}, {"data": {"id": "579679", "source": "579282", "target": "579286", "SCORE": 0.6142960573952252, "shared_interaction": "Direct", "source_original": "60", "Name_B": "GO:0007507", "Name_A": "GO:0003007", "target_original": "56", "Gene_B": "GO:0007507", "interaction": "Direct", "SUID": 579679, "TYPE": "Direct", "Gene_A": "GO:0003007", "BEND_MAP_ID": 579679, "selected": false}, "selected": false}, {"data": {"id": "579678", "source": "579282", "target": "579339", "SCORE": 0.31505919613981187, "shared_interaction": "Direct", "source_original": "60", "Name_B": "GO:0007389", "Name_A": "GO:0003007", "target_original": "3", "Gene_B": "GO:0007389", "interaction": "Direct", "SUID": 579678, "TYPE": "Direct", "Gene_A": "GO:0003007", "BEND_MAP_ID": 579678, "selected": false}, "selected": false}, {"data": {"id": "579677", "source": "579282", "target": "579285", "SCORE": 0.3208214721439569, "shared_interaction": "Direct", "source_original": "60", "Name_B": "GO:0042692", "Name_A": "GO:0003007", "target_original": "57", "Gene_B": "GO:0042692", "interaction": "Direct", "SUID": 579677, "TYPE": "Direct", "Gene_A": "GO:0003007", "BEND_MAP_ID": 579677, "selected": false}, "selected": false}, {"data": {"id": "579676", "source": "579282", "target": "579337", "SCORE": 0.3230650750874015, "shared_interaction": "Direct", "source_original": "60", "Name_B": "GO:0048568", "Name_A": "GO:0003007", "target_original": "5", "Gene_B": "GO:0048568", "interaction": "Direct", "SUID": 579676, "TYPE": "Direct", "Gene_A": "GO:0003007", "BEND_MAP_ID": 579676, "selected": false}, "selected": false}, {"data": {"id": "579675", "source": "579282", "target": "579284", "SCORE": 0.3024473859642147, "shared_interaction": "Direct", "source_original": "60", "Name_B": "GO:0051146", "Name_A": "GO:0003007", "target_original": "58", "Gene_B": "GO:0051146", "interaction": "Direct", "SUID": 579675, "TYPE": "Direct", "Gene_A": "GO:0003007", "BEND_MAP_ID": 579675, "selected": false}, "selected": false}, {"data": {"id": "579674", "source": "579282", "target": "579283", "SCORE": 0.38869965085659214, "shared_interaction": "Direct", "source_original": "60", "Name_B": "GO:0060537", "Name_A": "GO:0003007", "target_original": "59", "Gene_B": "GO:0060537", "interaction": "Direct", "SUID": 579674, "TYPE": "Direct", "Gene_A": "GO:0003007", "BEND_MAP_ID": 579674, "selected": false}, "selected": false}, {"data": {"id": "579673", "source": "579282", "target": "579281", "SCORE": 0.3847287999293187, "shared_interaction": "Direct", "source_original": "60", "Name_B": "GO:0014706", "Name_A": "GO:0003007", "target_original": "61", "Gene_B": "GO:0014706", "interaction": "Direct", "SUID": 579673, "TYPE": "Direct", "Gene_A": "GO:0003007", "BEND_MAP_ID": 579673, "selected": false}, "selected": false}, {"data": {"id": "579672", "source": "579282", "target": "579280", "SCORE": 0.735047151531312, "shared_interaction": "Direct", "source_original": "60", "Name_B": "GO:0003206", "Name_A": "GO:0003007", "target_original": "62", "Gene_B": "GO:0003206", "interaction": "Direct", "SUID": 579672, "TYPE": "Direct", "Gene_A": "GO:0003007", "BEND_MAP_ID": 579672, "selected": false}, "selected": false}, {"data": {"id": "579671", "source": "579282", "target": "579279", "SCORE": 0.7305334150832556, "shared_interaction": "Direct", "source_original": "60", "Name_B": "GO:0003205", "Name_A": "GO:0003007", "target_original": "63", "Gene_B": "GO:0003205", "interaction": "Direct", "SUID": 579671, "TYPE": "Direct", "Gene_A": "GO:0003007", "BEND_MAP_ID": 579671, "selected": false}, "selected": false}, {"data": {"id": "579670", "source": "579282", "target": "579278", "SCORE": 0.4979692854432493, "shared_interaction": "Direct", "source_original": "60", "Name_B": "GO:0035051", "Name_A": "GO:0003007", "target_original": "64", "Gene_B": "GO:0035051", "interaction": "Direct", "SUID": 579670, "TYPE": "Direct", "Gene_A": "GO:0003007", "BEND_MAP_ID": 579670, "selected": false}, "selected": false}, {"data": {"id": "579669", "source": "579281", "target": "579206", "SCORE": 0.6221740561899437, "shared_interaction": "Direct", "source_original": "61", "Name_B": "GO:0061061", "Name_A": "GO:0014706", "target_original": "136", "Gene_B": "GO:0061061", "interaction": "Direct", "SUID": 579669, "TYPE": "Direct", "Gene_A": "GO:0014706", "BEND_MAP_ID": 579669, "selected": false}, "selected": false}, {"data": {"id": "579668", "source": "579281", "target": "579285", "SCORE": 0.46768889777140016, "shared_interaction": "Direct", "source_original": "61", "Name_B": "GO:0042692", "Name_A": "GO:0014706", "target_original": "57", "Gene_B": "GO:0042692", "interaction": "Direct", "SUID": 579668, "TYPE": "Direct", "Gene_A": "GO:0014706", "BEND_MAP_ID": 579668, "selected": false}, "selected": false}, {"data": {"id": "579667", "source": "579281", "target": "579284", "SCORE": 0.5483857529686831, "shared_interaction": "Direct", "source_original": "61", "Name_B": "GO:0051146", "Name_A": "GO:0014706", "target_original": "58", "Gene_B": "GO:0051146", "interaction": "Direct", "SUID": 579667, "TYPE": "Direct", "Gene_A": "GO:0014706", "BEND_MAP_ID": 579667, "selected": false}, "selected": false}, {"data": {"id": "579666", "source": "579281", "target": "579283", "SCORE": 0.9556723841222905, "shared_interaction": "Direct", "source_original": "61", "Name_B": "GO:0060537", "Name_A": "GO:0014706", "target_original": "59", "Gene_B": "GO:0060537", "interaction": "Direct", "SUID": 579666, "TYPE": "Direct", "Gene_A": "GO:0014706", "BEND_MAP_ID": 579666, "selected": false}, "selected": false}, {"data": {"id": "579665", "source": "579281", "target": "579278", "SCORE": 0.5651321194780283, "shared_interaction": "Direct", "source_original": "61", "Name_B": "GO:0035051", "Name_A": "GO:0014706", "target_original": "64", "Gene_B": "GO:0035051", "interaction": "Direct", "SUID": 579665, "TYPE": "Direct", "Gene_A": "GO:0014706", "BEND_MAP_ID": 579665, "selected": false}, "selected": false}, {"data": {"id": "579664", "source": "579280", "target": "579286", "SCORE": 0.40967092008123657, "shared_interaction": "Direct", "source_original": "62", "Name_B": "GO:0007507", "Name_A": "GO:0003206", "target_original": "56", "Gene_B": "GO:0007507", "interaction": "Direct", "SUID": 579664, "TYPE": "Direct", "Gene_A": "GO:0003206", "BEND_MAP_ID": 579664, "selected": false}, "selected": false}, {"data": {"id": "579663", "source": "579280", "target": "579283", "SCORE": 0.3316267045176458, "shared_interaction": "Direct", "source_original": "62", "Name_B": "GO:0060537", "Name_A": "GO:0003206", "target_original": "59", "Gene_B": "GO:0060537", "interaction": "Direct", "SUID": 579663, "TYPE": "Direct", "Gene_A": "GO:0003206", "BEND_MAP_ID": 579663, "selected": false}, "selected": false}, {"data": {"id": "579662", "source": "579280", "target": "579281", "SCORE": 0.3539661199257279, "shared_interaction": "Direct", "source_original": "62", "Name_B": "GO:0014706", "Name_A": "GO:0003206", "target_original": "61", "Gene_B": "GO:0014706", "interaction": "Direct", "SUID": 579662, "TYPE": "Direct", "Gene_A": "GO:0003206", "BEND_MAP_ID": 579662, "selected": false}, "selected": false}, {"data": {"id": "579661", "source": "579280", "target": "579278", "SCORE": 0.4786918580034969, "shared_interaction": "Direct", "source_original": "62", "Name_B": "GO:0035051", "Name_A": "GO:0003206", "target_original": "64", "Gene_B": "GO:0035051", "interaction": "Direct", "SUID": 579661, "TYPE": "Direct", "Gene_A": "GO:0003206", "BEND_MAP_ID": 579661, "selected": false}, "selected": false}, {"data": {"id": "579660", "source": "579279", "target": "579286", "SCORE": 0.4579978237220125, "shared_interaction": "Direct", "source_original": "63", "Name_B": "GO:0007507", "Name_A": "GO:0003205", "target_original": "56", "Gene_B": "GO:0007507", "interaction": "Direct", "SUID": 579660, "TYPE": "Direct", "Gene_A": "GO:0003205", "BEND_MAP_ID": 579660, "selected": false}, "selected": false}, {"data": {"id": "579659", "source": "579279", "target": "579284", "SCORE": 0.3000166675937829, "shared_interaction": "Direct", "source_original": "63", "Name_B": "GO:0051146", "Name_A": "GO:0003205", "target_original": "58", "Gene_B": "GO:0051146", "interaction": "Direct", "SUID": 579659, "TYPE": "Direct", "Gene_A": "GO:0003205", "BEND_MAP_ID": 579659, "selected": false}, "selected": false}, {"data": {"id": "579658", "source": "579279", "target": "579283", "SCORE": 0.34357725557137286, "shared_interaction": "Direct", "source_original": "63", "Name_B": "GO:0060537", "Name_A": "GO:0003205", "target_original": "59", "Gene_B": "GO:0060537", "interaction": "Direct", "SUID": 579658, "TYPE": "Direct", "Gene_A": "GO:0003205", "BEND_MAP_ID": 579658, "selected": false}, "selected": false}, {"data": {"id": "579657", "source": "579279", "target": "579281", "SCORE": 0.3658825651208912, "shared_interaction": "Direct", "source_original": "63", "Name_B": "GO:0014706", "Name_A": "GO:0003205", "target_original": "61", "Gene_B": "GO:0014706", "interaction": "Direct", "SUID": 579657, "TYPE": "Direct", "Gene_A": "GO:0003205", "BEND_MAP_ID": 579657, "selected": false}, "selected": false}, {"data": {"id": "579656", "source": "579279", "target": "579280", "SCORE": 0.9289352413292977, "shared_interaction": "Direct", "source_original": "63", "Name_B": "GO:0003206", "Name_A": "GO:0003205", "target_original": "62", "Gene_B": "GO:0003206", "interaction": "Direct", "SUID": 579656, "TYPE": "Direct", "Gene_A": "GO:0003205", "BEND_MAP_ID": 579656, "selected": false}, "selected": false}, {"data": {"id": "579655", "source": "579279", "target": "579278", "SCORE": 0.4904549227315065, "shared_interaction": "Direct", "source_original": "63", "Name_B": "GO:0035051", "Name_A": "GO:0003205", "target_original": "64", "Gene_B": "GO:0035051", "interaction": "Direct", "SUID": 579655, "TYPE": "Direct", "Gene_A": "GO:0003205", "BEND_MAP_ID": 579655, "selected": false}, "selected": false}, {"data": {"id": "579654", "source": "579278", "target": "579206", "SCORE": 0.3552614293227429, "shared_interaction": "Direct", "source_original": "64", "Name_B": "GO:0061061", "Name_A": "GO:0035051", "target_original": "136", "Gene_B": "GO:0061061", "interaction": "Direct", "SUID": 579654, "TYPE": "Direct", "Gene_A": "GO:0035051", "BEND_MAP_ID": 579654, "selected": false}, "selected": false}, {"data": {"id": "579653", "source": "579278", "target": "579285", "SCORE": 0.5006982083271224, "shared_interaction": "Direct", "source_original": "64", "Name_B": "GO:0042692", "Name_A": "GO:0035051", "target_original": "57", "Gene_B": "GO:0042692", "interaction": "Direct", "SUID": 579653, "TYPE": "Direct", "Gene_A": "GO:0035051", "BEND_MAP_ID": 579653, "selected": false}, "selected": false}, {"data": {"id": "579652", "source": "579278", "target": "579284", "SCORE": 0.5803013808521342, "shared_interaction": "Direct", "source_original": "64", "Name_B": "GO:0051146", "Name_A": "GO:0035051", "target_original": "58", "Gene_B": "GO:0051146", "interaction": "Direct", "SUID": 579652, "TYPE": "Direct", "Gene_A": "GO:0035051", "BEND_MAP_ID": 579652, "selected": false}, "selected": false}, {"data": {"id": "579651", "source": "579278", "target": "579283", "SCORE": 0.5310944962623466, "shared_interaction": "Direct", "source_original": "64", "Name_B": "GO:0060537", "Name_A": "GO:0035051", "target_original": "59", "Gene_B": "GO:0060537", "interaction": "Direct", "SUID": 579651, "TYPE": "Direct", "Gene_A": "GO:0035051", "BEND_MAP_ID": 579651, "selected": false}, "selected": false}, {"data": {"id": "579650", "source": "579277", "target": "579199", "SCORE": 0.33932522706719, "shared_interaction": "Direct", "source_original": "65", "Name_B": "GO:0045664", "Name_A": "GO:0000904", "target_original": "143", "Gene_B": "GO:0045664", "interaction": "Direct", "SUID": 579650, "TYPE": "Direct", "Gene_A": "GO:0000904", "BEND_MAP_ID": 579650, "selected": false}, "selected": false}, {"data": {"id": "579649", "source": "579277", "target": "579276", "SCORE": 0.3006047103760227, "shared_interaction": "Direct", "source_original": "65", "Name_B": "GO:0010720", "Name_A": "GO:0000904", "target_original": "66", "Gene_B": "GO:0010720", "interaction": "Direct", "SUID": 579649, "TYPE": "Direct", "Gene_A": "GO:0000904", "BEND_MAP_ID": 579649, "selected": false}, "selected": false}, {"data": {"id": "579648", "source": "579277", "target": "579274", "SCORE": 0.4908350790329202, "shared_interaction": "Direct", "source_original": "65", "Name_B": "GO:0022604", "Name_A": "GO:0000904", "target_original": "68", "Gene_B": "GO:0022604", "interaction": "Direct", "SUID": 579648, "TYPE": "Direct", "Gene_A": "GO:0000904", "BEND_MAP_ID": 579648, "selected": false}, "selected": false}, {"data": {"id": "579647", "source": "579277", "target": "579272", "SCORE": 0.7278968723398963, "shared_interaction": "Direct", "source_original": "65", "Name_B": "GO:0048858", "Name_A": "GO:0000904", "target_original": "70", "Gene_B": "GO:0048858", "interaction": "Direct", "SUID": 579647, "TYPE": "Direct", "Gene_A": "GO:0000904", "BEND_MAP_ID": 579647, "selected": false}, "selected": false}, {"data": {"id": "579646", "source": "579277", "target": "579271", "SCORE": 0.7329138690027975, "shared_interaction": "Direct", "source_original": "65", "Name_B": "GO:0120039", "Name_A": "GO:0000904", "target_original": "71", "Gene_B": "GO:0120039", "interaction": "Direct", "SUID": 579646, "TYPE": "Direct", "Gene_A": "GO:0000904", "BEND_MAP_ID": 579646, "selected": false}, "selected": false}, {"data": {"id": "579645", "source": "579277", "target": "579270", "SCORE": 0.722934488597808, "shared_interaction": "Direct", "source_original": "65", "Name_B": "GO:0032990", "Name_A": "GO:0000904", "target_original": "72", "Gene_B": "GO:0032990", "interaction": "Direct", "SUID": 579645, "TYPE": "Direct", "Gene_A": "GO:0000904", "BEND_MAP_ID": 579645, "selected": false}, "selected": false}, {"data": {"id": "579644", "source": "579277", "target": "579269", "SCORE": 0.829643239220545, "shared_interaction": "Direct", "source_original": "65", "Name_B": "GO:0048667", "Name_A": "GO:0000904", "target_original": "73", "Gene_B": "GO:0048667", "interaction": "Direct", "SUID": 579644, "TYPE": "Direct", "Gene_A": "GO:0000904", "BEND_MAP_ID": 579644, "selected": false}, "selected": false}, {"data": {"id": "579643", "source": "579276", "target": "579199", "SCORE": 0.7221563700158397, "shared_interaction": "Direct", "source_original": "66", "Name_B": "GO:0045664", "Name_A": "GO:0010720", "target_original": "143", "Gene_B": "GO:0045664", "interaction": "Direct", "SUID": 579643, "TYPE": "Direct", "Gene_A": "GO:0010720", "BEND_MAP_ID": 579643, "selected": false}, "selected": false}, {"data": {"id": "579642", "source": "579276", "target": "579275", "SCORE": 0.8445440075459693, "shared_interaction": "Direct", "source_original": "66", "Name_B": "GO:0045666", "Name_A": "GO:0010720", "target_original": "67", "Gene_B": "GO:0045666", "interaction": "Direct", "SUID": 579642, "TYPE": "Direct", "Gene_A": "GO:0010720", "BEND_MAP_ID": 579642, "selected": false}, "selected": false}, {"data": {"id": "579641", "source": "579276", "target": "579274", "SCORE": 0.34186161179037156, "shared_interaction": "Direct", "source_original": "66", "Name_B": "GO:0022604", "Name_A": "GO:0010720", "target_original": "68", "Gene_B": "GO:0022604", "interaction": "Direct", "SUID": 579641, "TYPE": "Direct", "Gene_A": "GO:0010720", "BEND_MAP_ID": 579641, "selected": false}, "selected": false}, {"data": {"id": "579640", "source": "579276", "target": "579273", "SCORE": 0.9098007388336126, "shared_interaction": "Direct", "source_original": "66", "Name_B": "GO:0050769", "Name_A": "GO:0010720", "target_original": "69", "Gene_B": "GO:0050769", "interaction": "Direct", "SUID": 579640, "TYPE": "Direct", "Gene_A": "GO:0010720", "BEND_MAP_ID": 579640, "selected": false}, "selected": false}, {"data": {"id": "579639", "source": "579276", "target": "579272", "SCORE": 0.34578524836977365, "shared_interaction": "Direct", "source_original": "66", "Name_B": "GO:0048858", "Name_A": "GO:0010720", "target_original": "70", "Gene_B": "GO:0048858", "interaction": "Direct", "SUID": 579639, "TYPE": "Direct", "Gene_A": "GO:0010720", "BEND_MAP_ID": 579639, "selected": false}, "selected": false}, {"data": {"id": "579638", "source": "579276", "target": "579271", "SCORE": 0.3489217716281768, "shared_interaction": "Direct", "source_original": "66", "Name_B": "GO:0120039", "Name_A": "GO:0010720", "target_original": "71", "Gene_B": "GO:0120039", "interaction": "Direct", "SUID": 579638, "TYPE": "Direct", "Gene_A": "GO:0010720", "BEND_MAP_ID": 579638, "selected": false}, "selected": false}, {"data": {"id": "579637", "source": "579276", "target": "579270", "SCORE": 0.3426890672520877, "shared_interaction": "Direct", "source_original": "66", "Name_B": "GO:0032990", "Name_A": "GO:0010720", "target_original": "72", "Gene_B": "GO:0032990", "interaction": "Direct", "SUID": 579637, "TYPE": "Direct", "Gene_A": "GO:0010720", "BEND_MAP_ID": 579637, "selected": false}, "selected": false}, {"data": {"id": "579636", "source": "579275", "target": "579273", "SCORE": 0.9338152247572344, "shared_interaction": "Direct", "source_original": "67", "Name_B": "GO:0050769", "Name_A": "GO:0045666", "target_original": "69", "Gene_B": "GO:0050769", "interaction": "Direct", "SUID": 579636, "TYPE": "Direct", "Gene_A": "GO:0045666", "BEND_MAP_ID": 579636, "selected": false}, "selected": false}, {"data": {"id": "579635", "source": "579275", "target": "579272", "SCORE": 0.3269334392110188, "shared_interaction": "Direct", "source_original": "67", "Name_B": "GO:0048858", "Name_A": "GO:0045666", "target_original": "70", "Gene_B": "GO:0048858", "interaction": "Direct", "SUID": 579635, "TYPE": "Direct", "Gene_A": "GO:0045666", "BEND_MAP_ID": 579635, "selected": false}, "selected": false}, {"data": {"id": "579634", "source": "579275", "target": "579271", "SCORE": 0.33017088643823583, "shared_interaction": "Direct", "source_original": "67", "Name_B": "GO:0120039", "Name_A": "GO:0045666", "target_original": "71", "Gene_B": "GO:0120039", "interaction": "Direct", "SUID": 579634, "TYPE": "Direct", "Gene_A": "GO:0045666", "BEND_MAP_ID": 579634, "selected": false}, "selected": false}, {"data": {"id": "579633", "source": "579274", "target": "579199", "SCORE": 0.3873649624139481, "shared_interaction": "Direct", "source_original": "68", "Name_B": "GO:0045664", "Name_A": "GO:0022604", "target_original": "143", "Gene_B": "GO:0045664", "interaction": "Direct", "SUID": 579633, "TYPE": "Direct", "Gene_A": "GO:0022604", "BEND_MAP_ID": 579633, "selected": false}, "selected": false}, {"data": {"id": "579632", "source": "579274", "target": "579272", "SCORE": 0.4653850222132347, "shared_interaction": "Direct", "source_original": "68", "Name_B": "GO:0048858", "Name_A": "GO:0022604", "target_original": "70", "Gene_B": "GO:0048858", "interaction": "Direct", "SUID": 579632, "TYPE": "Direct", "Gene_A": "GO:0022604", "BEND_MAP_ID": 579632, "selected": false}, "selected": false}, {"data": {"id": "579631", "source": "579274", "target": "579271", "SCORE": 0.46945731480043534, "shared_interaction": "Direct", "source_original": "68", "Name_B": "GO:0120039", "Name_A": "GO:0022604", "target_original": "71", "Gene_B": "GO:0120039", "interaction": "Direct", "SUID": 579631, "TYPE": "Direct", "Gene_A": "GO:0022604", "BEND_MAP_ID": 579631, "selected": false}, "selected": false}, {"data": {"id": "579630", "source": "579274", "target": "579270", "SCORE": 0.4613687693685534, "shared_interaction": "Direct", "source_original": "68", "Name_B": "GO:0032990", "Name_A": "GO:0022604", "target_original": "72", "Gene_B": "GO:0032990", "interaction": "Direct", "SUID": 579630, "TYPE": "Direct", "Gene_A": "GO:0022604", "BEND_MAP_ID": 579630, "selected": false}, "selected": false}, {"data": {"id": "579629", "source": "579274", "target": "579269", "SCORE": 0.4050425855947157, "shared_interaction": "Direct", "source_original": "68", "Name_B": "GO:0048667", "Name_A": "GO:0022604", "target_original": "73", "Gene_B": "GO:0048667", "interaction": "Direct", "SUID": 579629, "TYPE": "Direct", "Gene_A": "GO:0022604", "BEND_MAP_ID": 579629, "selected": false}, "selected": false}, {"data": {"id": "579628", "source": "579273", "target": "579271", "SCORE": 0.33890610558417267, "shared_interaction": "Direct", "source_original": "69", "Name_B": "GO:0120039", "Name_A": "GO:0050769", "target_original": "71", "Gene_B": "GO:0120039", "interaction": "Direct", "SUID": 579628, "TYPE": "Direct", "Gene_A": "GO:0050769", "BEND_MAP_ID": 579628, "selected": false}, "selected": false}, {"data": {"id": "579627", "source": "579272", "target": "579273", "SCORE": 0.33569953710628964, "shared_interaction": "Direct", "source_original": "70", "Name_B": "GO:0050769", "Name_A": "GO:0048858", "target_original": "69", "Gene_B": "GO:0050769", "interaction": "Direct", "SUID": 579627, "TYPE": "Direct", "Gene_A": "GO:0048858", "BEND_MAP_ID": 579627, "selected": false}, "selected": false}, {"data": {"id": "579626", "source": "579272", "target": "579271", "SCORE": 0.9928330316520769, "shared_interaction": "Direct", "source_original": "70", "Name_B": "GO:0120039", "Name_A": "GO:0048858", "target_original": "71", "Gene_B": "GO:0120039", "interaction": "Direct", "SUID": 579626, "TYPE": "Direct", "Gene_A": "GO:0048858", "BEND_MAP_ID": 579626, "selected": false}, "selected": false}, {"data": {"id": "579625", "source": "579270", "target": "579199", "SCORE": 0.43664164072551986, "shared_interaction": "Direct", "source_original": "72", "Name_B": "GO:0045664", "Name_A": "GO:0032990", "target_original": "143", "Gene_B": "GO:0045664", "interaction": "Direct", "SUID": 579625, "TYPE": "Direct", "Gene_A": "GO:0032990", "BEND_MAP_ID": 579625, "selected": false}, "selected": false}, {"data": {"id": "579624", "source": "579270", "target": "579275", "SCORE": 0.32374465210593883, "shared_interaction": "Direct", "source_original": "72", "Name_B": "GO:0045666", "Name_A": "GO:0032990", "target_original": "67", "Gene_B": "GO:0045666", "interaction": "Direct", "SUID": 579624, "TYPE": "Direct", "Gene_A": "GO:0032990", "BEND_MAP_ID": 579624, "selected": false}, "selected": false}, {"data": {"id": "579623", "source": "579270", "target": "579273", "SCORE": 0.33253825744431104, "shared_interaction": "Direct", "source_original": "72", "Name_B": "GO:0050769", "Name_A": "GO:0032990", "target_original": "69", "Gene_B": "GO:0050769", "interaction": "Direct", "SUID": 579623, "TYPE": "Direct", "Gene_A": "GO:0032990", "BEND_MAP_ID": 579623, "selected": false}, "selected": false}, {"data": {"id": "579622", "source": "579270", "target": "579272", "SCORE": 0.9929215084676203, "shared_interaction": "Direct", "source_original": "72", "Name_B": "GO:0048858", "Name_A": "GO:0032990", "target_original": "70", "Gene_B": "GO:0048858", "interaction": "Direct", "SUID": 579622, "TYPE": "Direct", "Gene_A": "GO:0032990", "BEND_MAP_ID": 579622, "selected": false}, "selected": false}, {"data": {"id": "579621", "source": "579270", "target": "579271", "SCORE": 0.9857552627740876, "shared_interaction": "Direct", "source_original": "72", "Name_B": "GO:0120039", "Name_A": "GO:0032990", "target_original": "71", "Gene_B": "GO:0120039", "interaction": "Direct", "SUID": 579621, "TYPE": "Direct", "Gene_A": "GO:0032990", "BEND_MAP_ID": 579621, "selected": false}, "selected": false}, {"data": {"id": "579620", "source": "579270", "target": "579269", "SCORE": 0.8201442529030968, "shared_interaction": "Direct", "source_original": "72", "Name_B": "GO:0048667", "Name_A": "GO:0032990", "target_original": "73", "Gene_B": "GO:0048667", "interaction": "Direct", "SUID": 579620, "TYPE": "Direct", "Gene_A": "GO:0032990", "BEND_MAP_ID": 579620, "selected": false}, "selected": false}, {"data": {"id": "579619", "source": "579269", "target": "579272", "SCORE": 0.8264843709770094, "shared_interaction": "Direct", "source_original": "73", "Name_B": "GO:0048858", "Name_A": "GO:0048667", "target_original": "70", "Gene_B": "GO:0048858", "interaction": "Direct", "SUID": 579619, "TYPE": "Direct", "Gene_A": "GO:0048667", "BEND_MAP_ID": 579619, "selected": false}, "selected": false}, {"data": {"id": "579618", "source": "579269", "target": "579271", "SCORE": 0.8329099486755379, "shared_interaction": "Direct", "source_original": "73", "Name_B": "GO:0120039", "Name_A": "GO:0048667", "target_original": "71", "Gene_B": "GO:0120039", "interaction": "Direct", "SUID": 579618, "TYPE": "Direct", "Gene_A": "GO:0048667", "BEND_MAP_ID": 579618, "selected": false}, "selected": false}, {"data": {"id": "579617", "source": "579268", "target": "579201", "SCORE": 0.9518389151428371, "shared_interaction": "Direct", "source_original": "74", "Name_B": "GO:0034248", "Name_A": "GO:0006417", "target_original": "141", "Gene_B": "GO:0034248", "interaction": "Direct", "SUID": 579617, "TYPE": "Direct", "Gene_A": "GO:0006417", "BEND_MAP_ID": 579617, "selected": false}, "selected": false}, {"data": {"id": "579616", "source": "579268", "target": "579267", "SCORE": 0.6893235831813203, "shared_interaction": "Direct", "source_original": "74", "Name_B": "GO:0017148", "Name_A": "GO:0006417", "target_original": "75", "Gene_B": "GO:0017148", "interaction": "Direct", "SUID": 579616, "TYPE": "Direct", "Gene_A": "GO:0006417", "BEND_MAP_ID": 579616, "selected": false}, "selected": false}, {"data": {"id": "579615", "source": "579268", "target": "579266", "SCORE": 0.32145633798223033, "shared_interaction": "Direct", "source_original": "74", "Name_B": "GO:1903311", "Name_A": "GO:0006417", "target_original": "76", "Gene_B": "GO:1903311", "interaction": "Direct", "SUID": 579615, "TYPE": "Direct", "Gene_A": "GO:0006417", "BEND_MAP_ID": 579615, "selected": false}, "selected": false}, {"data": {"id": "579614", "source": "579268", "target": "579264", "SCORE": 0.8669542513339615, "shared_interaction": "Direct", "source_original": "74", "Name_B": "GO:0043043", "Name_A": "GO:0006417", "target_original": "78", "Gene_B": "GO:0043043", "interaction": "Direct", "SUID": 579614, "TYPE": "Direct", "Gene_A": "GO:0006417", "BEND_MAP_ID": 579614, "selected": false}, "selected": false}, {"data": {"id": "579613", "source": "579268", "target": "579263", "SCORE": 0.6816543831377987, "shared_interaction": "Direct", "source_original": "74", "Name_B": "GO:0034249", "Name_A": "GO:0006417", "target_original": "79", "Gene_B": "GO:0034249", "interaction": "Direct", "SUID": 579613, "TYPE": "Direct", "Gene_A": "GO:0006417", "BEND_MAP_ID": 579613, "selected": false}, "selected": false}, {"data": {"id": "579612", "source": "579268", "target": "579262", "SCORE": 0.3873555352804879, "shared_interaction": "Direct", "source_original": "74", "Name_B": "GO:0061013", "Name_A": "GO:0006417", "target_original": "80", "Gene_B": "GO:0061013", "interaction": "Direct", "SUID": 579612, "TYPE": "Direct", "Gene_A": "GO:0006417", "BEND_MAP_ID": 579612, "selected": false}, "selected": false}, {"data": {"id": "579611", "source": "579268", "target": "579261", "SCORE": 0.3340778620278413, "shared_interaction": "Direct", "source_original": "74", "Name_B": "GO:0043488", "Name_A": "GO:0006417", "target_original": "81", "Gene_B": "GO:0043488", "interaction": "Direct", "SUID": 579611, "TYPE": "Direct", "Gene_A": "GO:0006417", "BEND_MAP_ID": 579611, "selected": false}, "selected": false}, {"data": {"id": "579610", "source": "579267", "target": "579201", "SCORE": 0.6464524581124718, "shared_interaction": "Direct", "source_original": "75", "Name_B": "GO:0034248", "Name_A": "GO:0017148", "target_original": "141", "Gene_B": "GO:0034248", "interaction": "Direct", "SUID": 579610, "TYPE": "Direct", "Gene_A": "GO:0017148", "BEND_MAP_ID": 579610, "selected": false}, "selected": false}, {"data": {"id": "579609", "source": "579267", "target": "579266", "SCORE": 0.35854704673482163, "shared_interaction": "Direct", "source_original": "75", "Name_B": "GO:1903311", "Name_A": "GO:0017148", "target_original": "76", "Gene_B": "GO:1903311", "interaction": "Direct", "SUID": 579609, "TYPE": "Direct", "Gene_A": "GO:0017148", "BEND_MAP_ID": 579609, "selected": false}, "selected": false}, {"data": {"id": "579608", "source": "579267", "target": "579264", "SCORE": 0.573890711059239, "shared_interaction": "Direct", "source_original": "75", "Name_B": "GO:0043043", "Name_A": "GO:0017148", "target_original": "78", "Gene_B": "GO:0043043", "interaction": "Direct", "SUID": 579608, "TYPE": "Direct", "Gene_A": "GO:0017148", "BEND_MAP_ID": 579608, "selected": false}, "selected": false}, {"data": {"id": "579607", "source": "579267", "target": "579263", "SCORE": 0.9850677810620955, "shared_interaction": "Direct", "source_original": "75", "Name_B": "GO:0034249", "Name_A": "GO:0017148", "target_original": "79", "Gene_B": "GO:0034249", "interaction": "Direct", "SUID": 579607, "TYPE": "Direct", "Gene_A": "GO:0017148", "BEND_MAP_ID": 579607, "selected": false}, "selected": false}, {"data": {"id": "579606", "source": "579267", "target": "579262", "SCORE": 0.4538975989483604, "shared_interaction": "Direct", "source_original": "75", "Name_B": "GO:0061013", "Name_A": "GO:0017148", "target_original": "80", "Gene_B": "GO:0061013", "interaction": "Direct", "SUID": 579606, "TYPE": "Direct", "Gene_A": "GO:0017148", "BEND_MAP_ID": 579606, "selected": false}, "selected": false}, {"data": {"id": "579605", "source": "579267", "target": "579261", "SCORE": 0.3774549213287148, "shared_interaction": "Direct", "source_original": "75", "Name_B": "GO:0043488", "Name_A": "GO:0017148", "target_original": "81", "Gene_B": "GO:0043488", "interaction": "Direct", "SUID": 579605, "TYPE": "Direct", "Gene_A": "GO:0017148", "BEND_MAP_ID": 579605, "selected": false}, "selected": false}, {"data": {"id": "579604", "source": "579265", "target": "579201", "SCORE": 0.8558017643368784, "shared_interaction": "Direct", "source_original": "77", "Name_B": "GO:0034248", "Name_A": "GO:0006412", "target_original": "141", "Gene_B": "GO:0034248", "interaction": "Direct", "SUID": 579604, "TYPE": "Direct", "Gene_A": "GO:0006412", "BEND_MAP_ID": 579604, "selected": false}, "selected": false}, {"data": {"id": "579603", "source": "579265", "target": "579268", "SCORE": 0.8801559404359562, "shared_interaction": "Direct", "source_original": "77", "Name_B": "GO:0006417", "Name_A": "GO:0006412", "target_original": "74", "Gene_B": "GO:0006417", "interaction": "Direct", "SUID": 579603, "TYPE": "Direct", "Gene_A": "GO:0006412", "BEND_MAP_ID": 579603, "selected": false}, "selected": false}, {"data": {"id": "579602", "source": "579265", "target": "579267", "SCORE": 0.5849335767805882, "shared_interaction": "Direct", "source_original": "77", "Name_B": "GO:0017148", "Name_A": "GO:0006412", "target_original": "75", "Gene_B": "GO:0017148", "interaction": "Direct", "SUID": 579602, "TYPE": "Direct", "Gene_A": "GO:0006412", "BEND_MAP_ID": 579602, "selected": false}, "selected": false}, {"data": {"id": "579601", "source": "579265", "target": "579264", "SCORE": 0.9865844026510775, "shared_interaction": "Direct", "source_original": "77", "Name_B": "GO:0043043", "Name_A": "GO:0006412", "target_original": "78", "Gene_B": "GO:0043043", "interaction": "Direct", "SUID": 579601, "TYPE": "Direct", "Gene_A": "GO:0006412", "BEND_MAP_ID": 579601, "selected": false}, "selected": false}, {"data": {"id": "579600", "source": "579265", "target": "579263", "SCORE": 0.5790690048255702, "shared_interaction": "Direct", "source_original": "77", "Name_B": "GO:0034249", "Name_A": "GO:0006412", "target_original": "79", "Gene_B": "GO:0034249", "interaction": "Direct", "SUID": 579600, "TYPE": "Direct", "Gene_A": "GO:0006412", "BEND_MAP_ID": 579600, "selected": false}, "selected": false}, {"data": {"id": "579599", "source": "579265", "target": "579262", "SCORE": 0.32352367325766795, "shared_interaction": "Direct", "source_original": "77", "Name_B": "GO:0061013", "Name_A": "GO:0006412", "target_original": "80", "Gene_B": "GO:0061013", "interaction": "Direct", "SUID": 579599, "TYPE": "Direct", "Gene_A": "GO:0006412", "BEND_MAP_ID": 579599, "selected": false}, "selected": false}, {"data": {"id": "579598", "source": "579264", "target": "579262", "SCORE": 0.3168199821138376, "shared_interaction": "Direct", "source_original": "78", "Name_B": "GO:0061013", "Name_A": "GO:0043043", "target_original": "80", "Gene_B": "GO:0061013", "interaction": "Direct", "SUID": 579598, "TYPE": "Direct", "Gene_A": "GO:0043043", "BEND_MAP_ID": 579598, "selected": false}, "selected": false}, {"data": {"id": "579597", "source": "579263", "target": "579266", "SCORE": 0.35327226575516835, "shared_interaction": "Direct", "source_original": "79", "Name_B": "GO:1903311", "Name_A": "GO:0034249", "target_original": "76", "Gene_B": "GO:1903311", "interaction": "Direct", "SUID": 579597, "TYPE": "Direct", "Gene_A": "GO:0034249", "BEND_MAP_ID": 579597, "selected": false}, "selected": false}, {"data": {"id": "579596", "source": "579263", "target": "579264", "SCORE": 0.5681982161776041, "shared_interaction": "Direct", "source_original": "79", "Name_B": "GO:0043043", "Name_A": "GO:0034249", "target_original": "78", "Gene_B": "GO:0043043", "interaction": "Direct", "SUID": 579596, "TYPE": "Direct", "Gene_A": "GO:0034249", "BEND_MAP_ID": 579596, "selected": false}, "selected": false}, {"data": {"id": "579595", "source": "579263", "target": "579262", "SCORE": 0.44627429805703156, "shared_interaction": "Direct", "source_original": "79", "Name_B": "GO:0061013", "Name_A": "GO:0034249", "target_original": "80", "Gene_B": "GO:0061013", "interaction": "Direct", "SUID": 579595, "TYPE": "Direct", "Gene_A": "GO:0034249", "BEND_MAP_ID": 579595, "selected": false}, "selected": false}, {"data": {"id": "579594", "source": "579263", "target": "579261", "SCORE": 0.3707293638917364, "shared_interaction": "Direct", "source_original": "79", "Name_B": "GO:0043488", "Name_A": "GO:0034249", "target_original": "81", "Gene_B": "GO:0043488", "interaction": "Direct", "SUID": 579594, "TYPE": "Direct", "Gene_A": "GO:0034249", "BEND_MAP_ID": 579594, "selected": false}, "selected": false}, {"data": {"id": "579593", "source": "579262", "target": "579266", "SCORE": 0.7936619718312697, "shared_interaction": "Direct", "source_original": "80", "Name_B": "GO:1903311", "Name_A": "GO:0061013", "target_original": "76", "Gene_B": "GO:1903311", "interaction": "Direct", "SUID": 579593, "TYPE": "Direct", "Gene_A": "GO:0061013", "BEND_MAP_ID": 579593, "selected": false}, "selected": false}, {"data": {"id": "579592", "source": "579261", "target": "579266", "SCORE": 0.7425873050739457, "shared_interaction": "Direct", "source_original": "81", "Name_B": "GO:1903311", "Name_A": "GO:0043488", "target_original": "76", "Gene_B": "GO:1903311", "interaction": "Direct", "SUID": 579592, "TYPE": "Direct", "Gene_A": "GO:0043488", "BEND_MAP_ID": 579592, "selected": false}, "selected": false}, {"data": {"id": "579591", "source": "579261", "target": "579262", "SCORE": 0.9460603829161499, "shared_interaction": "Direct", "source_original": "81", "Name_B": "GO:0061013", "Name_A": "GO:0043488", "target_original": "80", "Gene_B": "GO:0061013", "interaction": "Direct", "SUID": 579591, "TYPE": "Direct", "Gene_A": "GO:0043488", "BEND_MAP_ID": 579591, "selected": false}, "selected": false}, {"data": {"id": "579590", "source": "579260", "target": "579201", "SCORE": 0.3785888385938471, "shared_interaction": "Direct", "source_original": "82", "Name_B": "GO:0034248", "Name_A": "GO:0006402", "target_original": "141", "Gene_B": "GO:0034248", "interaction": "Direct", "SUID": 579590, "TYPE": "Direct", "Gene_A": "GO:0006402", "BEND_MAP_ID": 579590, "selected": false}, "selected": false}, {"data": {"id": "579589", "source": "579260", "target": "579268", "SCORE": 0.4035374603710158, "shared_interaction": "Direct", "source_original": "82", "Name_B": "GO:0006417", "Name_A": "GO:0006402", "target_original": "74", "Gene_B": "GO:0006417", "interaction": "Direct", "SUID": 579589, "TYPE": "Direct", "Gene_A": "GO:0006402", "BEND_MAP_ID": 579589, "selected": false}, "selected": false}, {"data": {"id": "579588", "source": "579260", "target": "579267", "SCORE": 0.39030127567257084, "shared_interaction": "Direct", "source_original": "82", "Name_B": "GO:0017148", "Name_A": "GO:0006402", "target_original": "75", "Gene_B": "GO:0017148", "interaction": "Direct", "SUID": 579588, "TYPE": "Direct", "Gene_A": "GO:0006402", "BEND_MAP_ID": 579588, "selected": false}, "selected": false}, {"data": {"id": "579587", "source": "579260", "target": "579266", "SCORE": 0.6846573922752517, "shared_interaction": "Direct", "source_original": "82", "Name_B": "GO:1903311", "Name_A": "GO:0006402", "target_original": "76", "Gene_B": "GO:1903311", "interaction": "Direct", "SUID": 579587, "TYPE": "Direct", "Gene_A": "GO:0006402", "BEND_MAP_ID": 579587, "selected": false}, "selected": false}, {"data": {"id": "579586", "source": "579260", "target": "579265", "SCORE": 0.3935162240173101, "shared_interaction": "Direct", "source_original": "82", "Name_B": "GO:0006412", "Name_A": "GO:0006402", "target_original": "77", "Gene_B": "GO:0006412", "interaction": "Direct", "SUID": 579586, "TYPE": "Direct", "Gene_A": "GO:0006402", "BEND_MAP_ID": 579586, "selected": false}, "selected": false}, {"data": {"id": "579585", "source": "579260", "target": "579264", "SCORE": 0.38613207817977707, "shared_interaction": "Direct", "source_original": "82", "Name_B": "GO:0043043", "Name_A": "GO:0006402", "target_original": "78", "Gene_B": "GO:0043043", "interaction": "Direct", "SUID": 579585, "TYPE": "Direct", "Gene_A": "GO:0006402", "BEND_MAP_ID": 579585, "selected": false}, "selected": false}, {"data": {"id": "579584", "source": "579260", "target": "579263", "SCORE": 0.384591321446933, "shared_interaction": "Direct", "source_original": "82", "Name_B": "GO:0034249", "Name_A": "GO:0006402", "target_original": "79", "Gene_B": "GO:0034249", "interaction": "Direct", "SUID": 579584, "TYPE": "Direct", "Gene_A": "GO:0006402", "BEND_MAP_ID": 579584, "selected": false}, "selected": false}, {"data": {"id": "579583", "source": "579260", "target": "579262", "SCORE": 0.8048710874324856, "shared_interaction": "Direct", "source_original": "82", "Name_B": "GO:0061013", "Name_A": "GO:0006402", "target_original": "80", "Gene_B": "GO:0061013", "interaction": "Direct", "SUID": 579583, "TYPE": "Direct", "Gene_A": "GO:0006402", "BEND_MAP_ID": 579583, "selected": false}, "selected": false}, {"data": {"id": "579582", "source": "579260", "target": "579261", "SCORE": 0.753525656885903, "shared_interaction": "Direct", "source_original": "82", "Name_B": "GO:0043488", "Name_A": "GO:0006402", "target_original": "81", "Gene_B": "GO:0043488", "interaction": "Direct", "SUID": 579582, "TYPE": "Direct", "Gene_A": "GO:0006402", "BEND_MAP_ID": 579582, "selected": false}, "selected": false}, {"data": {"id": "579581", "source": "579259", "target": "579298", "SCORE": 0.5154021087458235, "shared_interaction": "Direct", "source_original": "83", "Name_B": "GO:1903707", "Name_A": "GO:0045637", "target_original": "44", "Gene_B": "GO:1903707", "interaction": "Direct", "SUID": 579581, "TYPE": "Direct", "Gene_A": "GO:0045637", "BEND_MAP_ID": 579581, "selected": false}, "selected": false}, {"data": {"id": "579580", "source": "579259", "target": "579317", "SCORE": 0.701051964958265, "shared_interaction": "Direct", "source_original": "83", "Name_B": "GO:1903706", "Name_A": "GO:0045637", "target_original": "25", "Gene_B": "GO:1903706", "interaction": "Direct", "SUID": 579580, "TYPE": "Direct", "Gene_A": "GO:0045637", "BEND_MAP_ID": 579580, "selected": false}, "selected": false}, {"data": {"id": "579579", "source": "579259", "target": "579312", "SCORE": 0.5032305365905027, "shared_interaction": "Direct", "source_original": "83", "Name_B": "GO:1902105", "Name_A": "GO:0045637", "target_original": "30", "Gene_B": "GO:1902105", "interaction": "Direct", "SUID": 579579, "TYPE": "Direct", "Gene_A": "GO:0045637", "BEND_MAP_ID": 579579, "selected": false}, "selected": false}, {"data": {"id": "579578", "source": "579259", "target": "579296", "SCORE": 0.5806583520760779, "shared_interaction": "Direct", "source_original": "83", "Name_B": "GO:0045638", "Name_A": "GO:0045637", "target_original": "46", "Gene_B": "GO:0045638", "interaction": "Direct", "SUID": 579578, "TYPE": "Direct", "Gene_A": "GO:0045637", "BEND_MAP_ID": 579578, "selected": false}, "selected": false}, {"data": {"id": "579577", "source": "579259", "target": "579256", "SCORE": 0.5522989304079129, "shared_interaction": "Direct", "source_original": "83", "Name_B": "GO:0045639", "Name_A": "GO:0045637", "target_original": "86", "Gene_B": "GO:0045639", "interaction": "Direct", "SUID": 579577, "TYPE": "Direct", "Gene_A": "GO:0045637", "BEND_MAP_ID": 579577, "selected": false}, "selected": false}, {"data": {"id": "579576", "source": "579258", "target": "579205", "SCORE": 0.6698014222985921, "shared_interaction": "Direct", "source_original": "84", "Name_B": "GO:0030099", "Name_A": "GO:0002573", "target_original": "137", "Gene_B": "GO:0030099", "interaction": "Direct", "SUID": 579576, "TYPE": "Direct", "Gene_A": "GO:0002573", "BEND_MAP_ID": 579576, "selected": false}, "selected": false}, {"data": {"id": "579575", "source": "579258", "target": "579317", "SCORE": 0.3085673199247901, "shared_interaction": "Direct", "source_original": "84", "Name_B": "GO:1903706", "Name_A": "GO:0002573", "target_original": "25", "Gene_B": "GO:1903706", "interaction": "Direct", "SUID": 579575, "TYPE": "Direct", "Gene_A": "GO:0002573", "BEND_MAP_ID": 579575, "selected": false}, "selected": false}, {"data": {"id": "579574", "source": "579258", "target": "579312", "SCORE": 0.42718897387731314, "shared_interaction": "Direct", "source_original": "84", "Name_B": "GO:1902105", "Name_A": "GO:0002573", "target_original": "30", "Gene_B": "GO:1902105", "interaction": "Direct", "SUID": 579574, "TYPE": "Direct", "Gene_A": "GO:0002573", "BEND_MAP_ID": 579574, "selected": false}, "selected": false}, {"data": {"id": "579573", "source": "579258", "target": "579259", "SCORE": 0.4623289466793769, "shared_interaction": "Direct", "source_original": "84", "Name_B": "GO:0045637", "Name_A": "GO:0002573", "target_original": "83", "Gene_B": "GO:0045637", "interaction": "Direct", "SUID": 579573, "TYPE": "Direct", "Gene_A": "GO:0002573", "BEND_MAP_ID": 579573, "selected": false}, "selected": false}, {"data": {"id": "579572", "source": "579258", "target": "579257", "SCORE": 0.443374638402169, "shared_interaction": "Direct", "source_original": "84", "Name_B": "GO:0030225", "Name_A": "GO:0002573", "target_original": "85", "Gene_B": "GO:0030225", "interaction": "Direct", "SUID": 579572, "TYPE": "Direct", "Gene_A": "GO:0002573", "BEND_MAP_ID": 579572, "selected": false}, "selected": false}, {"data": {"id": "579571", "source": "579258", "target": "579256", "SCORE": 0.43514805675636486, "shared_interaction": "Direct", "source_original": "84", "Name_B": "GO:0045639", "Name_A": "GO:0002573", "target_original": "86", "Gene_B": "GO:0045639", "interaction": "Direct", "SUID": 579571, "TYPE": "Direct", "Gene_A": "GO:0002573", "BEND_MAP_ID": 579571, "selected": false}, "selected": false}, {"data": {"id": "579570", "source": "579258", "target": "579255", "SCORE": 0.6749981512984933, "shared_interaction": "Direct", "source_original": "84", "Name_B": "GO:0002761", "Name_A": "GO:0002573", "target_original": "87", "Gene_B": "GO:0002761", "interaction": "Direct", "SUID": 579570, "TYPE": "Direct", "Gene_A": "GO:0002573", "BEND_MAP_ID": 579570, "selected": false}, "selected": false}, {"data": {"id": "579569", "source": "579258", "target": "579254", "SCORE": 0.443374638402169, "shared_interaction": "Direct", "source_original": "84", "Name_B": "GO:0002763", "Name_A": "GO:0002573", "target_original": "88", "Gene_B": "GO:0002763", "interaction": "Direct", "SUID": 579569, "TYPE": "Direct", "Gene_A": "GO:0002573", "BEND_MAP_ID": 579569, "selected": false}, "selected": false}, {"data": {"id": "579568", "source": "579257", "target": "579256", "SCORE": 0.5172258837648491, "shared_interaction": "Direct", "source_original": "85", "Name_B": "GO:0045639", "Name_A": "GO:0030225", "target_original": "86", "Gene_B": "GO:0045639", "interaction": "Direct", "SUID": 579568, "TYPE": "Direct", "Gene_A": "GO:0030225", "BEND_MAP_ID": 579568, "selected": false}, "selected": false}, {"data": {"id": "579567", "source": "579256", "target": "579317", "SCORE": 0.34148598625151066, "shared_interaction": "Direct", "source_original": "86", "Name_B": "GO:1903706", "Name_A": "GO:0045639", "target_original": "25", "Gene_B": "GO:1903706", "interaction": "Direct", "SUID": 579567, "TYPE": "Direct", "Gene_A": "GO:0045639", "BEND_MAP_ID": 579567, "selected": false}, "selected": false}, {"data": {"id": "579566", "source": "579256", "target": "579312", "SCORE": 0.3520975486491652, "shared_interaction": "Direct", "source_original": "86", "Name_B": "GO:1902105", "Name_A": "GO:0045639", "target_original": "30", "Gene_B": "GO:1902105", "interaction": "Direct", "SUID": 579566, "TYPE": "Direct", "Gene_A": "GO:0045639", "BEND_MAP_ID": 579566, "selected": false}, "selected": false}, {"data": {"id": "579565", "source": "579255", "target": "579298", "SCORE": 0.32097334878475386, "shared_interaction": "Direct", "source_original": "87", "Name_B": "GO:1903707", "Name_A": "GO:0002761", "target_original": "44", "Gene_B": "GO:1903707", "interaction": "Direct", "SUID": 579565, "TYPE": "Direct", "Gene_A": "GO:0002761", "BEND_MAP_ID": 579565, "selected": false}, "selected": false}, {"data": {"id": "579564", "source": "579255", "target": "579205", "SCORE": 0.40829814576910883, "shared_interaction": "Direct", "source_original": "87", "Name_B": "GO:0030099", "Name_A": "GO:0002761", "target_original": "137", "Gene_B": "GO:0030099", "interaction": "Direct", "SUID": 579564, "TYPE": "Direct", "Gene_A": "GO:0002761", "BEND_MAP_ID": 579564, "selected": false}, "selected": false}, {"data": {"id": "579563", "source": "579255", "target": "579317", "SCORE": 0.3815103605745969, "shared_interaction": "Direct", "source_original": "87", "Name_B": "GO:1903706", "Name_A": "GO:0002761", "target_original": "25", "Gene_B": "GO:1903706", "interaction": "Direct", "SUID": 579563, "TYPE": "Direct", "Gene_A": "GO:0002761", "BEND_MAP_ID": 579563, "selected": false}, "selected": false}, {"data": {"id": "579562", "source": "579255", "target": "579312", "SCORE": 0.5526171135413851, "shared_interaction": "Direct", "source_original": "87", "Name_B": "GO:1902105", "Name_A": "GO:0002761", "target_original": "30", "Gene_B": "GO:1902105", "interaction": "Direct", "SUID": 579562, "TYPE": "Direct", "Gene_A": "GO:0002761", "BEND_MAP_ID": 579562, "selected": false}, "selected": false}, {"data": {"id": "579561", "source": "579255", "target": "579259", "SCORE": 0.6079743109453761, "shared_interaction": "Direct", "source_original": "87", "Name_B": "GO:0045637", "Name_A": "GO:0002761", "target_original": "83", "Gene_B": "GO:0045637", "interaction": "Direct", "SUID": 579561, "TYPE": "Direct", "Gene_A": "GO:0002761", "BEND_MAP_ID": 579561, "selected": false}, "selected": false}, {"data": {"id": "579560", "source": "579255", "target": "579296", "SCORE": 0.3788909963202288, "shared_interaction": "Direct", "source_original": "87", "Name_B": "GO:0045638", "Name_A": "GO:0002761", "target_original": "46", "Gene_B": "GO:0045638", "interaction": "Direct", "SUID": 579560, "TYPE": "Direct", "Gene_A": "GO:0002761", "BEND_MAP_ID": 579560, "selected": false}, "selected": false}, {"data": {"id": "579559", "source": "579255", "target": "579257", "SCORE": 0.47483799807188565, "shared_interaction": "Direct", "source_original": "87", "Name_B": "GO:0030225", "Name_A": "GO:0002761", "target_original": "85", "Gene_B": "GO:0030225", "interaction": "Direct", "SUID": 579559, "TYPE": "Direct", "Gene_A": "GO:0002761", "BEND_MAP_ID": 579559, "selected": false}, "selected": false}, {"data": {"id": "579558", "source": "579255", "target": "579256", "SCORE": 0.66236460013942, "shared_interaction": "Direct", "source_original": "87", "Name_B": "GO:0045639", "Name_A": "GO:0002761", "target_original": "86", "Gene_B": "GO:0045639", "interaction": "Direct", "SUID": 579558, "TYPE": "Direct", "Gene_A": "GO:0002761", "BEND_MAP_ID": 579558, "selected": false}, "selected": false}, {"data": {"id": "579557", "source": "579255", "target": "579254", "SCORE": 0.7172204605002461, "shared_interaction": "Direct", "source_original": "87", "Name_B": "GO:0002763", "Name_A": "GO:0002761", "target_original": "88", "Gene_B": "GO:0002763", "interaction": "Direct", "SUID": 579557, "TYPE": "Direct", "Gene_A": "GO:0002761", "BEND_MAP_ID": 579557, "selected": false}, "selected": false}, {"data": {"id": "579556", "source": "579254", "target": "579312", "SCORE": 0.35183712972375625, "shared_interaction": "Direct", "source_original": "88", "Name_B": "GO:1902105", "Name_A": "GO:0002763", "target_original": "30", "Gene_B": "GO:1902105", "interaction": "Direct", "SUID": 579556, "TYPE": "Direct", "Gene_A": "GO:0002763", "BEND_MAP_ID": 579556, "selected": false}, "selected": false}, {"data": {"id": "579555", "source": "579254", "target": "579259", "SCORE": 0.3925362819640471, "shared_interaction": "Direct", "source_original": "88", "Name_B": "GO:0045637", "Name_A": "GO:0002763", "target_original": "83", "Gene_B": "GO:0045637", "interaction": "Direct", "SUID": 579555, "TYPE": "Direct", "Gene_A": "GO:0002763", "BEND_MAP_ID": 579555, "selected": false}, "selected": false}, {"data": {"id": "579554", "source": "579254", "target": "579257", "SCORE": 0.5521161746466282, "shared_interaction": "Direct", "source_original": "88", "Name_B": "GO:0030225", "Name_A": "GO:0002763", "target_original": "85", "Gene_B": "GO:0030225", "interaction": "Direct", "SUID": 579554, "TYPE": "Direct", "Gene_A": "GO:0002763", "BEND_MAP_ID": 579554, "selected": false}, "selected": false}, {"data": {"id": "579553", "source": "579254", "target": "579256", "SCORE": 0.7805572198931132, "shared_interaction": "Direct", "source_original": "88", "Name_B": "GO:0045639", "Name_A": "GO:0002763", "target_original": "86", "Gene_B": "GO:0045639", "interaction": "Direct", "SUID": 579553, "TYPE": "Direct", "Gene_A": "GO:0002763", "BEND_MAP_ID": 579553, "selected": false}, "selected": false}, {"data": {"id": "579552", "source": "579253", "target": "579213", "SCORE": 0.6499402628437804, "shared_interaction": "Direct", "source_original": "89", "Name_B": "GO:0071383", "Name_A": "GO:0032870", "target_original": "129", "Gene_B": "GO:0071383", "interaction": "Direct", "SUID": 579552, "TYPE": "Direct", "Gene_A": "GO:0032870", "BEND_MAP_ID": 579552, "selected": false}, "selected": false}, {"data": {"id": "579551", "source": "579253", "target": "579252", "SCORE": 0.5641554702498268, "shared_interaction": "Direct", "source_original": "89", "Name_B": "GO:0071407", "Name_A": "GO:0032870", "target_original": "90", "Gene_B": "GO:0071407", "interaction": "Direct", "SUID": 579551, "TYPE": "Direct", "Gene_A": "GO:0032870", "BEND_MAP_ID": 579551, "selected": false}, "selected": false}, {"data": {"id": "579550", "source": "579253", "target": "579251", "SCORE": 0.642151656754739, "shared_interaction": "Direct", "source_original": "89", "Name_B": "GO:0048545", "Name_A": "GO:0032870", "target_original": "91", "Gene_B": "GO:0048545", "interaction": "Direct", "SUID": 579550, "TYPE": "Direct", "Gene_A": "GO:0032870", "BEND_MAP_ID": 579550, "selected": false}, "selected": false}, {"data": {"id": "579549", "source": "579253", "target": "579250", "SCORE": 0.5881360697219583, "shared_interaction": "Direct", "source_original": "89", "Name_B": "GO:0071396", "Name_A": "GO:0032870", "target_original": "92", "Gene_B": "GO:0071396", "interaction": "Direct", "SUID": 579549, "TYPE": "Direct", "Gene_A": "GO:0032870", "BEND_MAP_ID": 579549, "selected": false}, "selected": false}, {"data": {"id": "579548", "source": "579253", "target": "579249", "SCORE": 0.5077468732504021, "shared_interaction": "Direct", "source_original": "89", "Name_B": "GO:0043401", "Name_A": "GO:0032870", "target_original": "93", "Gene_B": "GO:0043401", "interaction": "Direct", "SUID": 579548, "TYPE": "Direct", "Gene_A": "GO:0032870", "BEND_MAP_ID": 579548, "selected": false}, "selected": false}, {"data": {"id": "579547", "source": "579251", "target": "579213", "SCORE": 0.8892249527411437, "shared_interaction": "Direct", "source_original": "91", "Name_B": "GO:0071383", "Name_A": "GO:0048545", "target_original": "129", "Gene_B": "GO:0071383", "interaction": "Direct", "SUID": 579547, "TYPE": "Direct", "Gene_A": "GO:0048545", "BEND_MAP_ID": 579547, "selected": false}, "selected": false}, {"data": {"id": "579546", "source": "579251", "target": "579252", "SCORE": 0.7016650694779811, "shared_interaction": "Direct", "source_original": "91", "Name_B": "GO:0071407", "Name_A": "GO:0048545", "target_original": "90", "Gene_B": "GO:0071407", "interaction": "Direct", "SUID": 579546, "TYPE": "Direct", "Gene_A": "GO:0048545", "BEND_MAP_ID": 579546, "selected": false}, "selected": false}, {"data": {"id": "579545", "source": "579251", "target": "579250", "SCORE": 0.6939059779457045, "shared_interaction": "Direct", "source_original": "91", "Name_B": "GO:0071396", "Name_A": "GO:0048545", "target_original": "92", "Gene_B": "GO:0071396", "interaction": "Direct", "SUID": 579545, "TYPE": "Direct", "Gene_A": "GO:0048545", "BEND_MAP_ID": 579545, "selected": false}, "selected": false}, {"data": {"id": "579544", "source": "579250", "target": "579252", "SCORE": 0.7417297521520163, "shared_interaction": "Direct", "source_original": "92", "Name_B": "GO:0071407", "Name_A": "GO:0071396", "target_original": "90", "Gene_B": "GO:0071407", "interaction": "Direct", "SUID": 579544, "TYPE": "Direct", "Gene_A": "GO:0071396", "BEND_MAP_ID": 579544, "selected": false}, "selected": false}, {"data": {"id": "579543", "source": "579249", "target": "579213", "SCORE": 0.8282028730580003, "shared_interaction": "Direct", "source_original": "93", "Name_B": "GO:0071383", "Name_A": "GO:0043401", "target_original": "129", "Gene_B": "GO:0071383", "interaction": "Direct", "SUID": 579543, "TYPE": "Direct", "Gene_A": "GO:0043401", "BEND_MAP_ID": 579543, "selected": false}, "selected": false}, {"data": {"id": "579542", "source": "579249", "target": "579252", "SCORE": 0.5979634577891165, "shared_interaction": "Direct", "source_original": "93", "Name_B": "GO:0071407", "Name_A": "GO:0043401", "target_original": "90", "Gene_B": "GO:0071407", "interaction": "Direct", "SUID": 579542, "TYPE": "Direct", "Gene_A": "GO:0043401", "BEND_MAP_ID": 579542, "selected": false}, "selected": false}, {"data": {"id": "579541", "source": "579249", "target": "579251", "SCORE": 0.7227049815025434, "shared_interaction": "Direct", "source_original": "93", "Name_B": "GO:0048545", "Name_A": "GO:0043401", "target_original": "91", "Gene_B": "GO:0048545", "interaction": "Direct", "SUID": 579541, "TYPE": "Direct", "Gene_A": "GO:0043401", "BEND_MAP_ID": 579541, "selected": false}, "selected": false}, {"data": {"id": "579540", "source": "579249", "target": "579250", "SCORE": 0.5727973338892074, "shared_interaction": "Direct", "source_original": "93", "Name_B": "GO:0071396", "Name_A": "GO:0043401", "target_original": "92", "Gene_B": "GO:0071396", "interaction": "Direct", "SUID": 579540, "TYPE": "Direct", "Gene_A": "GO:0043401", "BEND_MAP_ID": 579540, "selected": false}, "selected": false}, {"data": {"id": "579539", "source": "579248", "target": "579213", "SCORE": 0.7082993936106685, "shared_interaction": "Direct", "source_original": "94", "Name_B": "GO:0071383", "Name_A": "GO:0030522", "target_original": "129", "Gene_B": "GO:0071383", "interaction": "Direct", "SUID": 579539, "TYPE": "Direct", "Gene_A": "GO:0030522", "BEND_MAP_ID": 579539, "selected": false}, "selected": false}, {"data": {"id": "579538", "source": "579248", "target": "579253", "SCORE": 0.46011650023822626, "shared_interaction": "Direct", "source_original": "94", "Name_B": "GO:0032870", "Name_A": "GO:0030522", "target_original": "89", "Gene_B": "GO:0032870", "interaction": "Direct", "SUID": 579538, "TYPE": "Direct", "Gene_A": "GO:0030522", "BEND_MAP_ID": 579538, "selected": false}, "selected": false}, {"data": {"id": "579537", "source": "579248", "target": "579252", "SCORE": 0.5191466083155798, "shared_interaction": "Direct", "source_original": "94", "Name_B": "GO:0071407", "Name_A": "GO:0030522", "target_original": "90", "Gene_B": "GO:0071407", "interaction": "Direct", "SUID": 579537, "TYPE": "Direct", "Gene_A": "GO:0030522", "BEND_MAP_ID": 579537, "selected": false}, "selected": false}, {"data": {"id": "579536", "source": "579248", "target": "579251", "SCORE": 0.6462491951066628, "shared_interaction": "Direct", "source_original": "94", "Name_B": "GO:0048545", "Name_A": "GO:0030522", "target_original": "91", "Gene_B": "GO:0048545", "interaction": "Direct", "SUID": 579536, "TYPE": "Direct", "Gene_A": "GO:0030522", "BEND_MAP_ID": 579536, "selected": false}, "selected": false}, {"data": {"id": "579535", "source": "579248", "target": "579250", "SCORE": 0.5172663766974473, "shared_interaction": "Direct", "source_original": "94", "Name_B": "GO:0071396", "Name_A": "GO:0030522", "target_original": "92", "Gene_B": "GO:0071396", "interaction": "Direct", "SUID": 579535, "TYPE": "Direct", "Gene_A": "GO:0030522", "BEND_MAP_ID": 579535, "selected": false}, "selected": false}, {"data": {"id": "579534", "source": "579248", "target": "579249", "SCORE": 0.8441323545060416, "shared_interaction": "Direct", "source_original": "94", "Name_B": "GO:0043401", "Name_A": "GO:0030522", "target_original": "93", "Gene_B": "GO:0043401", "interaction": "Direct", "SUID": 579534, "TYPE": "Direct", "Gene_A": "GO:0030522", "BEND_MAP_ID": 579534, "selected": false}, "selected": false}, {"data": {"id": "579533", "source": "579247", "target": "579213", "SCORE": 0.8163275147075603, "shared_interaction": "Direct", "source_original": "95", "Name_B": "GO:0071383", "Name_A": "GO:0009755", "target_original": "129", "Gene_B": "GO:0071383", "interaction": "Direct", "SUID": 579533, "TYPE": "Direct", "Gene_A": "GO:0009755", "BEND_MAP_ID": 579533, "selected": false}, "selected": false}, {"data": {"id": "579532", "source": "579247", "target": "579253", "SCORE": 0.5206063038315087, "shared_interaction": "Direct", "source_original": "95", "Name_B": "GO:0032870", "Name_A": "GO:0009755", "target_original": "89", "Gene_B": "GO:0032870", "interaction": "Direct", "SUID": 579532, "TYPE": "Direct", "Gene_A": "GO:0009755", "BEND_MAP_ID": 579532, "selected": false}, "selected": false}, {"data": {"id": "579531", "source": "579247", "target": "579252", "SCORE": 0.5912066683365645, "shared_interaction": "Direct", "source_original": "95", "Name_B": "GO:0071407", "Name_A": "GO:0009755", "target_original": "90", "Gene_B": "GO:0071407", "interaction": "Direct", "SUID": 579531, "TYPE": "Direct", "Gene_A": "GO:0009755", "BEND_MAP_ID": 579531, "selected": false}, "selected": false}, {"data": {"id": "579530", "source": "579247", "target": "579251", "SCORE": 0.7133925913328117, "shared_interaction": "Direct", "source_original": "95", "Name_B": "GO:0048545", "Name_A": "GO:0009755", "target_original": "91", "Gene_B": "GO:0048545", "interaction": "Direct", "SUID": 579530, "TYPE": "Direct", "Gene_A": "GO:0009755", "BEND_MAP_ID": 579530, "selected": false}, "selected": false}, {"data": {"id": "579529", "source": "579247", "target": "579250", "SCORE": 0.5664934796825072, "shared_interaction": "Direct", "source_original": "95", "Name_B": "GO:0071396", "Name_A": "GO:0009755", "target_original": "92", "Gene_B": "GO:0071396", "interaction": "Direct", "SUID": 579529, "TYPE": "Direct", "Gene_A": "GO:0009755", "BEND_MAP_ID": 579529, "selected": false}, "selected": false}, {"data": {"id": "579528", "source": "579247", "target": "579249", "SCORE": 0.9831686580882636, "shared_interaction": "Direct", "source_original": "95", "Name_B": "GO:0043401", "Name_A": "GO:0009755", "target_original": "93", "Gene_B": "GO:0043401", "interaction": "Direct", "SUID": 579528, "TYPE": "Direct", "Gene_A": "GO:0009755", "BEND_MAP_ID": 579528, "selected": false}, "selected": false}, {"data": {"id": "579527", "source": "579247", "target": "579248", "SCORE": 0.8310176689563756, "shared_interaction": "Direct", "source_original": "95", "Name_B": "GO:0030522", "Name_A": "GO:0009755", "target_original": "94", "Gene_B": "GO:0030522", "interaction": "Direct", "SUID": 579527, "TYPE": "Direct", "Gene_A": "GO:0009755", "BEND_MAP_ID": 579527, "selected": false}, "selected": false}, {"data": {"id": "579526", "source": "579247", "target": "579246", "SCORE": 0.7294359994323, "shared_interaction": "Direct", "source_original": "95", "Name_B": "GO:0030518", "Name_A": "GO:0009755", "target_original": "96", "Gene_B": "GO:0030518", "interaction": "Direct", "SUID": 579526, "TYPE": "Direct", "Gene_A": "GO:0009755", "BEND_MAP_ID": 579526, "selected": false}, "selected": false}, {"data": {"id": "579525", "source": "579247", "target": "579245", "SCORE": 0.48110979929289943, "shared_interaction": "Direct", "source_original": "95", "Name_B": "GO:0030521", "Name_A": "GO:0009755", "target_original": "97", "Gene_B": "GO:0030521", "interaction": "Direct", "SUID": 579525, "TYPE": "Direct", "Gene_A": "GO:0009755", "BEND_MAP_ID": 579525, "selected": false}, "selected": false}, {"data": {"id": "579524", "source": "579246", "target": "579213", "SCORE": 0.5912103243815816, "shared_interaction": "Direct", "source_original": "96", "Name_B": "GO:0071383", "Name_A": "GO:0030518", "target_original": "129", "Gene_B": "GO:0071383", "interaction": "Direct", "SUID": 579524, "TYPE": "Direct", "Gene_A": "GO:0030518", "BEND_MAP_ID": 579524, "selected": false}, "selected": false}, {"data": {"id": "579523", "source": "579246", "target": "579253", "SCORE": 0.3361483007215744, "shared_interaction": "Direct", "source_original": "96", "Name_B": "GO:0032870", "Name_A": "GO:0030518", "target_original": "89", "Gene_B": "GO:0032870", "interaction": "Direct", "SUID": 579523, "TYPE": "Direct", "Gene_A": "GO:0030518", "BEND_MAP_ID": 579523, "selected": false}, "selected": false}, {"data": {"id": "579522", "source": "579246", "target": "579252", "SCORE": 0.4041324330927142, "shared_interaction": "Direct", "source_original": "96", "Name_B": "GO:0071407", "Name_A": "GO:0030518", "target_original": "90", "Gene_B": "GO:0071407", "interaction": "Direct", "SUID": 579522, "TYPE": "Direct", "Gene_A": "GO:0030518", "BEND_MAP_ID": 579522, "selected": false}, "selected": false}, {"data": {"id": "579521", "source": "579246", "target": "579251", "SCORE": 0.5029438179829645, "shared_interaction": "Direct", "source_original": "96", "Name_B": "GO:0048545", "Name_A": "GO:0030518", "target_original": "91", "Gene_B": "GO:0048545", "interaction": "Direct", "SUID": 579521, "TYPE": "Direct", "Gene_A": "GO:0030518", "BEND_MAP_ID": 579521, "selected": false}, "selected": false}, {"data": {"id": "579520", "source": "579246", "target": "579250", "SCORE": 0.3848845346403085, "shared_interaction": "Direct", "source_original": "96", "Name_B": "GO:0071396", "Name_A": "GO:0030518", "target_original": "92", "Gene_B": "GO:0071396", "interaction": "Direct", "SUID": 579520, "TYPE": "Direct", "Gene_A": "GO:0030518", "BEND_MAP_ID": 579520, "selected": false}, "selected": false}, {"data": {"id": "579519", "source": "579246", "target": "579249", "SCORE": 0.7451065680736169, "shared_interaction": "Direct", "source_original": "96", "Name_B": "GO:0043401", "Name_A": "GO:0030518", "target_original": "93", "Gene_B": "GO:0043401", "interaction": "Direct", "SUID": 579519, "TYPE": "Direct", "Gene_A": "GO:0030518", "BEND_MAP_ID": 579519, "selected": false}, "selected": false}, {"data": {"id": "579518", "source": "579246", "target": "579248", "SCORE": 0.659698025552328, "shared_interaction": "Direct", "source_original": "96", "Name_B": "GO:0030522", "Name_A": "GO:0030518", "target_original": "94", "Gene_B": "GO:0030522", "interaction": "Direct", "SUID": 579518, "TYPE": "Direct", "Gene_A": "GO:0030518", "BEND_MAP_ID": 579518, "selected": false}, "selected": false}, {"data": {"id": "579517", "source": "579246", "target": "579245", "SCORE": 0.7111165886133176, "shared_interaction": "Direct", "source_original": "96", "Name_B": "GO:0030521", "Name_A": "GO:0030518", "target_original": "97", "Gene_B": "GO:0030521", "interaction": "Direct", "SUID": 579517, "TYPE": "Direct", "Gene_A": "GO:0030518", "BEND_MAP_ID": 579517, "selected": false}, "selected": false}, {"data": {"id": "579516", "source": "579245", "target": "579213", "SCORE": 0.37601490749489436, "shared_interaction": "Direct", "source_original": "97", "Name_B": "GO:0071383", "Name_A": "GO:0030521", "target_original": "129", "Gene_B": "GO:0071383", "interaction": "Direct", "SUID": 579516, "TYPE": "Direct", "Gene_A": "GO:0030521", "BEND_MAP_ID": 579516, "selected": false}, "selected": false}, {"data": {"id": "579515", "source": "579245", "target": "579251", "SCORE": 0.3127443315101159, "shared_interaction": "Direct", "source_original": "97", "Name_B": "GO:0048545", "Name_A": "GO:0030521", "target_original": "91", "Gene_B": "GO:0048545", "interaction": "Direct", "SUID": 579515, "TYPE": "Direct", "Gene_A": "GO:0030521", "BEND_MAP_ID": 579515, "selected": false}, "selected": false}, {"data": {"id": "579514", "source": "579245", "target": "579249", "SCORE": 0.49351771823810153, "shared_interaction": "Direct", "source_original": "97", "Name_B": "GO:0043401", "Name_A": "GO:0030521", "target_original": "93", "Gene_B": "GO:0043401", "interaction": "Direct", "SUID": 579514, "TYPE": "Direct", "Gene_A": "GO:0030521", "BEND_MAP_ID": 579514, "selected": false}, "selected": false}, {"data": {"id": "579513", "source": "579245", "target": "579248", "SCORE": 0.4271318995350279, "shared_interaction": "Direct", "source_original": "97", "Name_B": "GO:0030522", "Name_A": "GO:0030521", "target_original": "94", "Gene_B": "GO:0030522", "interaction": "Direct", "SUID": 579513, "TYPE": "Direct", "Gene_A": "GO:0030521", "BEND_MAP_ID": 579513, "selected": false}, "selected": false}, {"data": {"id": "579512", "source": "579244", "target": "579243", "SCORE": 0.614642700570544, "shared_interaction": "Direct", "source_original": "98", "Name_B": "GO:0030278", "Name_A": "GO:0001649", "target_original": "99", "Gene_B": "GO:0030278", "interaction": "Direct", "SUID": 579512, "TYPE": "Direct", "Gene_A": "GO:0001649", "BEND_MAP_ID": 579512, "selected": false}, "selected": false}, {"data": {"id": "579511", "source": "579244", "target": "579242", "SCORE": 0.7078763708878713, "shared_interaction": "Direct", "source_original": "98", "Name_B": "GO:0045667", "Name_A": "GO:0001649", "target_original": "100", "Gene_B": "GO:0045667", "interaction": "Direct", "SUID": 579511, "TYPE": "Direct", "Gene_A": "GO:0001649", "BEND_MAP_ID": 579511, "selected": false}, "selected": false}, {"data": {"id": "579510", "source": "579244", "target": "579241", "SCORE": 0.3925362819640471, "shared_interaction": "Direct", "source_original": "98", "Name_B": "GO:0045668", "Name_A": "GO:0001649", "target_original": "101", "Gene_B": "GO:0045668", "interaction": "Direct", "SUID": 579510, "TYPE": "Direct", "Gene_A": "GO:0001649", "BEND_MAP_ID": 579510, "selected": false}, "selected": false}, {"data": {"id": "579509", "source": "579244", "target": "579240", "SCORE": 0.4398498902511081, "shared_interaction": "Direct", "source_original": "98", "Name_B": "GO:0045778", "Name_A": "GO:0001649", "target_original": "102", "Gene_B": "GO:0045778", "interaction": "Direct", "SUID": 579509, "TYPE": "Direct", "Gene_A": "GO:0001649", "BEND_MAP_ID": 579509, "selected": false}, "selected": false}, {"data": {"id": "579508", "source": "579244", "target": "579239", "SCORE": 0.4271318995350279, "shared_interaction": "Direct", "source_original": "98", "Name_B": "GO:0045669", "Name_A": "GO:0001649", "target_original": "103", "Gene_B": "GO:0045669", "interaction": "Direct", "SUID": 579508, "TYPE": "Direct", "Gene_A": "GO:0001649", "BEND_MAP_ID": 579508, "selected": false}, "selected": false}, {"data": {"id": "579507", "source": "579243", "target": "579211", "SCORE": 0.3132812500009815, "shared_interaction": "Direct", "source_original": "99", "Name_B": "GO:0061448", "Name_A": "GO:0030278", "target_original": "131", "Gene_B": "GO:0061448", "interaction": "Direct", "SUID": 579507, "TYPE": "Direct", "Gene_A": "GO:0030278", "BEND_MAP_ID": 579507, "selected": false}, "selected": false}, {"data": {"id": "579506", "source": "579243", "target": "579242", "SCORE": 0.8299477655256084, "shared_interaction": "Direct", "source_original": "99", "Name_B": "GO:0045667", "Name_A": "GO:0030278", "target_original": "100", "Gene_B": "GO:0045667", "interaction": "Direct", "SUID": 579506, "TYPE": "Direct", "Gene_A": "GO:0030278", "BEND_MAP_ID": 579506, "selected": false}, "selected": false}, {"data": {"id": "579505", "source": "579243", "target": "579236", "SCORE": 0.3502173334532995, "shared_interaction": "Direct", "source_original": "99", "Name_B": "GO:0032330", "Name_A": "GO:0030278", "target_original": "106", "Gene_B": "GO:0032330", "interaction": "Direct", "SUID": 579505, "TYPE": "Direct", "Gene_A": "GO:0030278", "BEND_MAP_ID": 579505, "selected": false}, "selected": false}, {"data": {"id": "579504", "source": "579243", "target": "579234", "SCORE": 0.3111139353129673, "shared_interaction": "Direct", "source_original": "99", "Name_B": "GO:0051216", "Name_A": "GO:0030278", "target_original": "108", "Gene_B": "GO:0051216", "interaction": "Direct", "SUID": 579504, "TYPE": "Direct", "Gene_A": "GO:0030278", "BEND_MAP_ID": 579504, "selected": false}, "selected": false}, {"data": {"id": "579503", "source": "579243", "target": "579233", "SCORE": 0.3808638323064951, "shared_interaction": "Direct", "source_original": "99", "Name_B": "GO:0061035", "Name_A": "GO:0030278", "target_original": "109", "Gene_B": "GO:0061035", "interaction": "Direct", "SUID": 579503, "TYPE": "Direct", "Gene_A": "GO:0030278", "BEND_MAP_ID": 579503, "selected": false}, "selected": false}, {"data": {"id": "579502", "source": "579243", "target": "579241", "SCORE": 0.4804479701367456, "shared_interaction": "Direct", "source_original": "99", "Name_B": "GO:0045668", "Name_A": "GO:0030278", "target_original": "101", "Gene_B": "GO:0045668", "interaction": "Direct", "SUID": 579502, "TYPE": "Direct", "Gene_A": "GO:0030278", "BEND_MAP_ID": 579502, "selected": false}, "selected": false}, {"data": {"id": "579501", "source": "579243", "target": "579240", "SCORE": 0.628518299383897, "shared_interaction": "Direct", "source_original": "99", "Name_B": "GO:0045778", "Name_A": "GO:0030278", "target_original": "102", "Gene_B": "GO:0045778", "interaction": "Direct", "SUID": 579501, "TYPE": "Direct", "Gene_A": "GO:0030278", "BEND_MAP_ID": 579501, "selected": false}, "selected": false}, {"data": {"id": "579500", "source": "579243", "target": "579239", "SCORE": 0.5202837911600914, "shared_interaction": "Direct", "source_original": "99", "Name_B": "GO:0045669", "Name_A": "GO:0030278", "target_original": "103", "Gene_B": "GO:0045669", "interaction": "Direct", "SUID": 579500, "TYPE": "Direct", "Gene_A": "GO:0030278", "BEND_MAP_ID": 579500, "selected": false}, "selected": false}, {"data": {"id": "579499", "source": "579242", "target": "579241", "SCORE": 0.6166289996444898, "shared_interaction": "Direct", "source_original": "100", "Name_B": "GO:0045668", "Name_A": "GO:0045667", "target_original": "101", "Gene_B": "GO:0045668", "interaction": "Direct", "SUID": 579499, "TYPE": "Direct", "Gene_A": "GO:0045667", "BEND_MAP_ID": 579499, "selected": false}, "selected": false}, {"data": {"id": "579498", "source": "579242", "target": "579240", "SCORE": 0.662123912360825, "shared_interaction": "Direct", "source_original": "100", "Name_B": "GO:0045778", "Name_A": "GO:0045667", "target_original": "102", "Gene_B": "GO:0045778", "interaction": "Direct", "SUID": 579498, "TYPE": "Direct", "Gene_A": "GO:0045667", "BEND_MAP_ID": 579498, "selected": false}, "selected": false}, {"data": {"id": "579497", "source": "579242", "target": "579239", "SCORE": 0.6628308400471665, "shared_interaction": "Direct", "source_original": "100", "Name_B": "GO:0045669", "Name_A": "GO:0045667", "target_original": "103", "Gene_B": "GO:0045669", "interaction": "Direct", "SUID": 579497, "TYPE": "Direct", "Gene_A": "GO:0045667", "BEND_MAP_ID": 579497, "selected": false}, "selected": false}, {"data": {"id": "579496", "source": "579240", "target": "579233", "SCORE": 0.4554987721531774, "shared_interaction": "Direct", "source_original": "102", "Name_B": "GO:0061035", "Name_A": "GO:0045778", "target_original": "109", "Gene_B": "GO:0061035", "interaction": "Direct", "SUID": 579496, "TYPE": "Direct", "Gene_A": "GO:0045778", "BEND_MAP_ID": 579496, "selected": false}, "selected": false}, {"data": {"id": "579495", "source": "579239", "target": "579233", "SCORE": 0.34147437818689025, "shared_interaction": "Direct", "source_original": "103", "Name_B": "GO:0061035", "Name_A": "GO:0045669", "target_original": "109", "Gene_B": "GO:0061035", "interaction": "Direct", "SUID": 579495, "TYPE": "Direct", "Gene_A": "GO:0045669", "BEND_MAP_ID": 579495, "selected": false}, "selected": false}, {"data": {"id": "579494", "source": "579239", "target": "579240", "SCORE": 0.8682948756373781, "shared_interaction": "Direct", "source_original": "103", "Name_B": "GO:0045778", "Name_A": "GO:0045669", "target_original": "102", "Gene_B": "GO:0045778", "interaction": "Direct", "SUID": 579494, "TYPE": "Direct", "Gene_A": "GO:0045669", "BEND_MAP_ID": 579494, "selected": false}, "selected": false}, {"data": {"id": "579493", "source": "579238", "target": "579212", "SCORE": 0.34957394411023257, "shared_interaction": "Direct", "source_original": "104", "Name_B": "GO:0043009", "Name_A": "GO:0001501", "target_original": "130", "Gene_B": "GO:0043009", "interaction": "Direct", "SUID": 579493, "TYPE": "Direct", "Gene_A": "GO:0001501", "BEND_MAP_ID": 579493, "selected": false}, "selected": false}, {"data": {"id": "579492", "source": "579238", "target": "579341", "SCORE": 0.34671876795814605, "shared_interaction": "Direct", "source_original": "104", "Name_B": "GO:0009792", "Name_A": "GO:0001501", "target_original": "1", "Gene_B": "GO:0009792", "interaction": "Direct", "SUID": 579492, "TYPE": "Direct", "Gene_A": "GO:0001501", "BEND_MAP_ID": 579492, "selected": false}, "selected": false}, {"data": {"id": "579491", "source": "579238", "target": "579340", "SCORE": 0.39007323435036834, "shared_interaction": "Direct", "source_original": "104", "Name_B": "GO:0048598", "Name_A": "GO:0001501", "target_original": "2", "Gene_B": "GO:0048598", "interaction": "Direct", "SUID": 579491, "TYPE": "Direct", "Gene_A": "GO:0001501", "BEND_MAP_ID": 579491, "selected": false}, "selected": false}, {"data": {"id": "579490", "source": "579238", "target": "579202", "SCORE": 0.387145035950845, "shared_interaction": "Direct", "source_original": "104", "Name_B": "GO:0001503", "Name_A": "GO:0001501", "target_original": "140", "Gene_B": "GO:0001503", "interaction": "Direct", "SUID": 579490, "TYPE": "Direct", "Gene_A": "GO:0001501", "BEND_MAP_ID": 579490, "selected": false}, "selected": false}, {"data": {"id": "579489", "source": "579238", "target": "579211", "SCORE": 0.49985506779653666, "shared_interaction": "Direct", "source_original": "104", "Name_B": "GO:0061448", "Name_A": "GO:0001501", "target_original": "131", "Gene_B": "GO:0061448", "interaction": "Direct", "SUID": 579489, "TYPE": "Direct", "Gene_A": "GO:0001501", "BEND_MAP_ID": 579489, "selected": false}, "selected": false}, {"data": {"id": "579488", "source": "579238", "target": "579339", "SCORE": 0.39237521661367064, "shared_interaction": "Direct", "source_original": "104", "Name_B": "GO:0007389", "Name_A": "GO:0001501", "target_original": "3", "Gene_B": "GO:0007389", "interaction": "Direct", "SUID": 579488, "TYPE": "Direct", "Gene_A": "GO:0001501", "BEND_MAP_ID": 579488, "selected": false}, "selected": false}, {"data": {"id": "579487", "source": "579238", "target": "579338", "SCORE": 0.4033410416600793, "shared_interaction": "Direct", "source_original": "104", "Name_B": "GO:0009952", "Name_A": "GO:0001501", "target_original": "4", "Gene_B": "GO:0009952", "interaction": "Direct", "SUID": 579487, "TYPE": "Direct", "Gene_A": "GO:0001501", "BEND_MAP_ID": 579487, "selected": false}, "selected": false}, {"data": {"id": "579486", "source": "579238", "target": "579337", "SCORE": 0.3663860215679184, "shared_interaction": "Di
```
