## Supplementary material for "Tensor-decomposition-based unsupervised feature extraction applied to prostate cancer multiomics data": output from metascape: GONetwork.style.js

```
var styles=[{"format_version": "1.0", "generated_by": "cytoscape-3.8.0", "target_cytoscapejs_version": "~2.1", "title": "ColorByCluster", "style": [{"selector": "node", "css": {"background-opacity": 1.0, "text-valign": "center", "text-halign": "right", "border-opacity": 1.0, "content": "", "border-width": 4.0, "width": 10.0, "background-color": "rgb(0,153,204)", "color": "rgb(0,0,0)", "height": 10.0, "border-color": "rgb(0,102,153)", "font-family": "Dialog.plain", "font-weight": "normal", "shape": "ellipse", "font-size": 24, "text-opacity": 1.0}}, {"selector": "node[GROUP_ID = 0.0]", "css": {"background-color": "rgb(188,189,220)"}}, {"selector": "node[GROUP_ID = 2.0]", "css": {"background-color": "rgb(55,126,184)"}}, {"selector": "node[GROUP_ID = 8.0]", "css": {"background-color": "rgb(247,129,191)"}}, {"selector": "node[GROUP_ID = 9.0]", "css": {"background-color": "rgb(153,153,153)"}}, {"selector": "node[GROUP_ID = 10.0]", "css": {"background-color": "rgb(141,211,199)"}}, {"selector": "node[GROUP_ID = 11.0]", "css": {"background-color": "rgb(255,255,179)"}}, {"selector": "node[GROUP_ID = 3.0]", "css": {"background-color": "rgb(77,175,74)"}}, {"selector": "node[GROUP_ID = 12.0]", "css": {"background-color": "rgb(190,186,218)"}}, {"selector": "node[GROUP_ID = 13.0]", "css": {"background-color": "rgb(251,128,114)"}}, {"selector": "node[GROUP_ID = 14.0]", "css": {"background-color": "rgb(128,177,211)"}}, {"selector": "node[GROUP_ID = 15.0]", "css": {"background-color": "rgb(253,180,98)"}}, {"selector": "node[GROUP_ID = 1.0]", "css": {"background-color": "rgb(228,26,28)"}}, {"selector": "node[GROUP_ID = 4.0]", "css": {"background-color": "rgb(152,78,163)"}}, {"selector": "node[GROUP_ID = 16.0]", "css": {"background-color": "rgb(179,222,105)"}}, {"selector": "node[GROUP_ID = 17.0]", "css": {"background-color": "rgb(252,205,229)"}}, {"selector": "node[GROUP_ID = 18.0]", "css": {"background-color": "rgb(217,217,217)"}}, {"selector": "node[GROUP_ID = 19.0]", "css": {"background-color": "rgb(188,128,189)"}}, {"selector": "node[GROUP_ID = 5.0]", "css": {"background-color": "rgb(255,127,0)"}}, {"selector": "node[GROUP_ID = 20.0]", "css": {"background-color": "rgb(204,235,197)"}}, {"selector": "node[GROUP_ID = 6.0]", "css": {"background-color": "rgb(255,255,51)"}}, {"selector": "node[GROUP_ID = 7.0]", "css": {"background-color": "rgb(166,86,40)"}}, {"selector": "node:selected", "css": {"background-color": "rgb(255,255,0)"}}, {"selector": "edge", "css": {"source-arrow-color": "rgb(0,0,0)", "font-family": "Dialog.plain", "font-weight": "normal", "font-size": 10, "content": "", "width": 3.0, "opacity": 0.39215686274509803, "text-opacity": 1.0, "target-arrow-shape": "none", "target-arrow-color": "rgb(0,0,0)", "source-arrow-shape": "none", "color": "rgb(0,0,0)", "line-style": "solid", "line-color": "rgb(84,39,143)"}}, {"selector": "edge[SCORE > 1]", "css": {"width": 5.0}}, {"selector": "edge[SCORE = 1]", "css": {"width": 5.0}}, {"selector": "edge[SCORE > 0.3][SCORE < 1]", "css": {"width": "mapData(SCORE,0.3,1,2.0,5.0)"}}, {"selector": "edge[SCORE = 0.3]", "css": {"width": 2.0}}, {"selector": "edge[SCORE < 0.3]", "css": {"width": 2.0}}, {"selector": "edge:selected", "css": {"line-color": "rgb(255,0,0)"}}, {"selector": "node[_GeneInGOAndHitList<=5]", "css": {"width": 20.0, "height": 20.0}}, {"selector": "node[_GeneInGOAndHitList>5][_GeneInGOAndHitList<20]", "css": {"width": "mapData(_GeneInGOAndHitList,5,20,35.0,50.0)", "height": "mapData(_GeneInGOAndHitList,5,20,35.0,50.0)"}}, {"selector": "node[_GeneInGOAndHitList>=20]", "css": {"width": 50.0, "height": 50.0}}]}, {"format_version": "1.0", "generated_by": "cytoscape-3.8.0", "target_cytoscapejs_version": "~2.1", "title": "ColorByPValue", "style": [{"selector": "node", "css": {"background-opacity": 1.0, "text-valign": "center", "text-halign": "right", "border-opacity": 1.0, "content": "", "border-width": 4.0, "width": 10.0, "background-color": "rgb(0,153,204)", "color": "rgb(0,0,0)", "height": 10.0, "border-color": "rgb(0,102,153)", "font-family": "Dialog.plain", "font-weight": "normal", "shape": "ellipse", "font-size": 24, "text-opacity": 1.0}}, {"selector": "node[LogP > -2]", "css": {"background-color": "rgb(255,255,212)"}}, {"selector": "node[LogP = -2]", "css": {"background-color": "rgb(255,255,212)"}}, {"selector": "node[LogP > -3][LogP < -2]", "css": {"background-color": "mapData(LogP,-3,-2,rgb(254,227,145),rgb(255,255,212))"}}, {"selector": "node[LogP > -4][LogP < -3]", "css": {"background-color": "mapData(LogP,-4,-3,rgb(254,196,79),rgb(254,227,145))"}}, {"selector": "node[LogP > -6][LogP < -4]", "css": {"background-color": "mapData(LogP,-6,-4,rgb(254,153,41),rgb(254,196,79))"}}, {"selector": "node[LogP > -10][LogP < -6]", "css": {"background-color": "mapData(LogP,-10,-6,rgb(217,95,14),rgb(254,153,41))"}}, {"selector": "node[LogP > -20][LogP < -10]", "css": {"background-color": "mapData(LogP,-20,-10,rgb(153,52,4),rgb(217,95,14))"}}, {"selector": "node[LogP = -20]", "css": {"background-color": "rgb(153,52,4)"}}, {"selector": "node[LogP < -20]", "css": {"background-color": "rgb(153,52,4)"}}, {"selector": "node:selected", "css": {"background-color": "rgb(255,255,0)"}}, {"selector": "edge", "css": {"source-arrow-color": "rgb(0,0,0)", "font-family": "Dialog.plain", "font-weight": "normal", "font-size": 10, "content": "", "width": 3.0, "opacity": 0.39215686274509803, "text-opacity": 1.0, "target-arrow-shape": "none", "target-arrow-color": "rgb(0,0,0)", "source-arrow-shape": "none", "color": "rgb(0,0,0)", "line-style": "solid", "line-color": "rgb(84,39,143)"}}, {"selector": "edge[SCORE > 1]", "css": {"width": 5.0}}, {"selector": "edge[SCORE = 1]", "css": {"width": 5.0}}, {"selector": "edge[SCORE > 0.3][SCORE < 1]", "css": {"width": "mapData(SCORE,0.3,1,2.0,5.0)"}}, {"selector": "edge[SCORE = 0.3]", "css": {"width": 2.0}}, {"selector": "edge[SCORE < 0.3]", "css": {"width": 2.0}}, {"selector": "edge:selected", "css": {"line-color": "rgb(255,0,0)"}}, {"selector": "node[_GeneInGOAndHitList<=5]", "css": {"width": 20.0, "height": 20.0}}, {"selector": "node[_GeneInGOAndHitList>5][_GeneInGOAndHitList<20]", "css": {"width": "mapData(_GeneInGOAndHitList,5,20,35.0,50.0)", "height": "mapData(_GeneInGOAndHitList,5,20,35.0,50.0)"}}, {"selector": "node[_GeneInGOAndHitList>=20]", "css": {"width": 50.0, "height": 50.0}}]}, {"format_version": "1.0", "generated_by": "cytoscape-3.3.0", "target_cytoscapejs_version": "~2.1", "title": "default", "style": [{"selector": "node", "css": {"text-opacity": 1.0, "text-valign": "center", "text-halign": "right", "color": "rgb(0,0,0)", "font-family": "Dialog.plain", "font-weight": "normal", "border-opacity": 1.0, "border-color": "rgb(0,102,153)", "shape": "ellipse", "font-size": 20, "content": "", "background-color": "rgb(153,204,255)", "height": 35.0, "background-opacity": 1.0, "width": 35.0, "border-width": 4.0}}, {"selector": "node[_GeneInGOAndHitList > 20]", "css": {"width": 50.0}}, {"selector": "node[_GeneInGOAndHitList = 20]", "css": {"width": 50.0}}, {"selector": "node[_GeneInGOAndHitList > 5][_GeneInGOAndHitList < 20]", "css": {"width": "mapData(_GeneInGOAndHitList,5,20,20.0,50.0)"}}, {"selector": "node[_GeneInGOAndHitList = 5]", "css": {"width": 20.0}}, {"selector": "node[_GeneInGOAndHitList < 5]", "css": {"width": 20.0}}, {"selector": "node[_GeneInGOAndHitList > 20]", "css": {"height": 50.0}}, {"selector": "node[_GeneInGOAndHitList = 20]", "css": {"height": 50.0}}, {"selector": "node[_GeneInGOAndHitList > 5][_GeneInGOAndHitList < 20]", "css": {"height": "mapData(_GeneInGOAndHitList,5,20,20.0,50.0)"}}, {"selector": "node[_GeneInGOAndHitList = 5]", "css": {"height": 20.0}}, {"selector": "node[_GeneInGOAndHitList < 5]", "css": {"height": 20.0}}, {"selector": "node:selected", "css": {"background-color": "rgb(255,255,0)"}}, {"selector": "edge", "css": {"font-size": 10, "line-style": "solid", "opacity": 0.39215686274509803, "color": "rgb(0,0,0)", "target-arrow-color": "rgb(0,0,0)", "source-arrow-color": "rgb(0,0,0)", "content": "", "text-opacity": 1.0, "target-arrow-shape": "none", "source-arrow-shape": "none", "font-family": "Dialog.plain", "font-weight": "normal", "width": 3.0, "line-color": "rgb(84,39,143)"}}, {"selector": "edge[SCORE > 1]", "css": {"width": 10.0}}, {"selector": "edge[SCORE = 1]", "css": {"width": 10.0}}, {"selector": "edge[SCORE > 0.3][SCORE < 1]", "css": {"width": "mapData(SCORE,0.3,1,2.0,10.0)"}}, {"selector": "edge[SCORE = 0.3]", "css": {"width": 2.0}}, {"selector": "edge[SCORE < 0.3]", "css": {"width": 2.0}}, {"selector": "edge:selected", "css": {"line-color": "rgb(255,0,0)"}}, {"selector": "node[_GeneInGOAndHitList<=5]", "css": {"width": 20.0, "height": 20.0}}, {"selector": "node[_GeneInGOAndHitList>5][_GeneInGOAndHitList<20]", "css": {"width": "mapData(_GeneInGOAndHitList,5,20,35.0,50.0)", "height": "mapData(_GeneInGOAndHitList,5,20,35.0,50.0)"}}, {"selector": "node[_GeneInGOAndHitList>=20]", "css": {"width": 50.0, "height": 50.0}}]}];
```
