## Supplementary material for "Tensor-decomposition-based unsupervised feature extraction applied to prostate cancer multiomics data": output from metascape: README.txt

Metascape Analysis Report Package

README.txt
--------------------
    This file

GeneListReport.pptx
--------------------
    PowerPoint presentation-ready file summarizing the key analysis results.

metascape_result.xlsx
--------------------
    Excel file containing two sheets: Annotation and Enrichment
    The "Annotation" sheet contains all gene identifiers, annotations, binary membership search and enrichment results (1/0).
    The "Enrichment" sheet contains all enriched terms and their cluster group ID.  The best term within a group is chosen as the group summary.

Evidence.csv
--------------------
    An experimental gene evidence matrix used for Gene Prioritization by Evidence Counting (GPEC) algorithm (not published yet)

Enrichment_heatmap folder
--------------------
    Available if enriched terms are found. The folder contains heatmap of select GO terms.
    To inspect the heatmap interactively, please use JTreeView program.
    Download and install JTreeView from http://jtreeview.sourceforge.net/
    Then open .cdt file to view the heatmap (.cdt, .atr, .gtr, .jtv are for this purpose).
    The Log10(p-value) behind the heatmap are in the .csv file.

Enrichment_GO folder
--------------------
    Available if enriched terms are found. The folder contains network of select GO terms.
    To inspect the network interactively, please use Cytoscape program.
    Download and install Cytoscape from http://www.cytoscape.org
    Then open .cys file to view the network (.cys, .xgmml are for this purpose).
    The GO_AllLists.csv file contains the original enrichment analysis results (before list merging),
    where the last column "GeneList" indicate the input gene list name.

    If there are multiple gene lists, _FINAL_GO.csv contains the GO enrichment results with all lists merged.
    GO_membership.csv provides the GO by Gene Lists matrix
    GeneGo_membership.csv contains the data behind clustergram, Gene by Cluster and individual GO terms.

Overlap_circos folder
--------------------
    Available if multiple gene lists are provided. The folder contains Circos plots.
    Plots are generated with Circos software (http://circos.ca), however, Circos is not required for visualization.
    .svg is the vector format, which can be open in popular browsers or manipulated in Illustrator.

Enrichment_PPI folder
--------------------
    Available if protein network is found.  The folder contains network plots.
    The networks are defined as xgmml files within the xgmml subfolder.
    The networks have been preloaded into a Cytoscape session file called MCODE_PPI.cys.
    GO_MCODE_Top3.csv file contains the GO enrichment analysis results for each network and
    MCODE components (only the top 3 GO terms are retained).

    If there are multiple gene lists, MERGE_MCODE.csv contains the MCODE components formed by all lists merged.

Contact
--------------------
If you have questions, please contact metascape.team @ gmail.com
