## Supplementary figures and images for "Tensor-decomposition-based unsupervised feature extraction applied to prostate cancer multiomics data"

### ColorByCluster.png

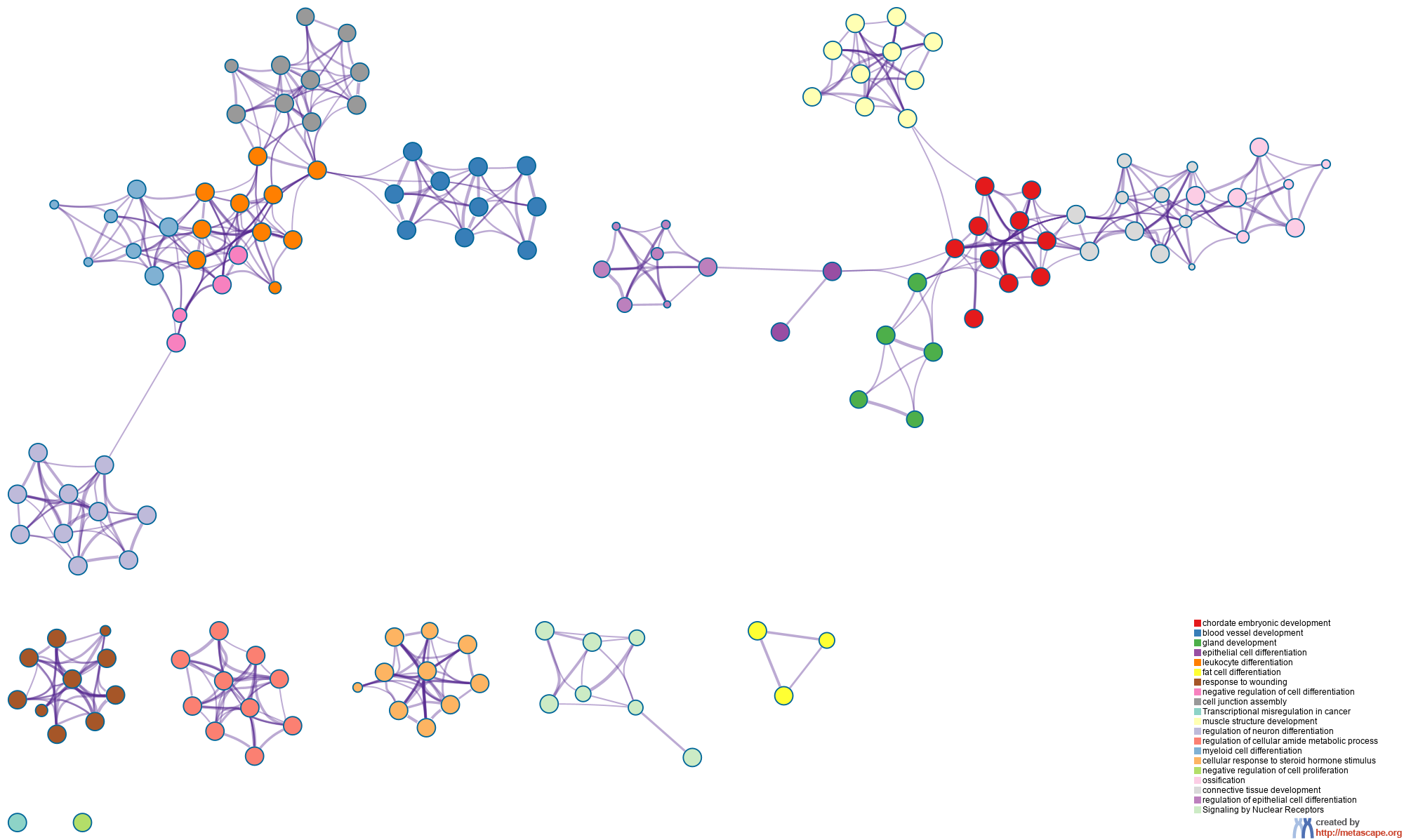

### ColorByPValue.pdf

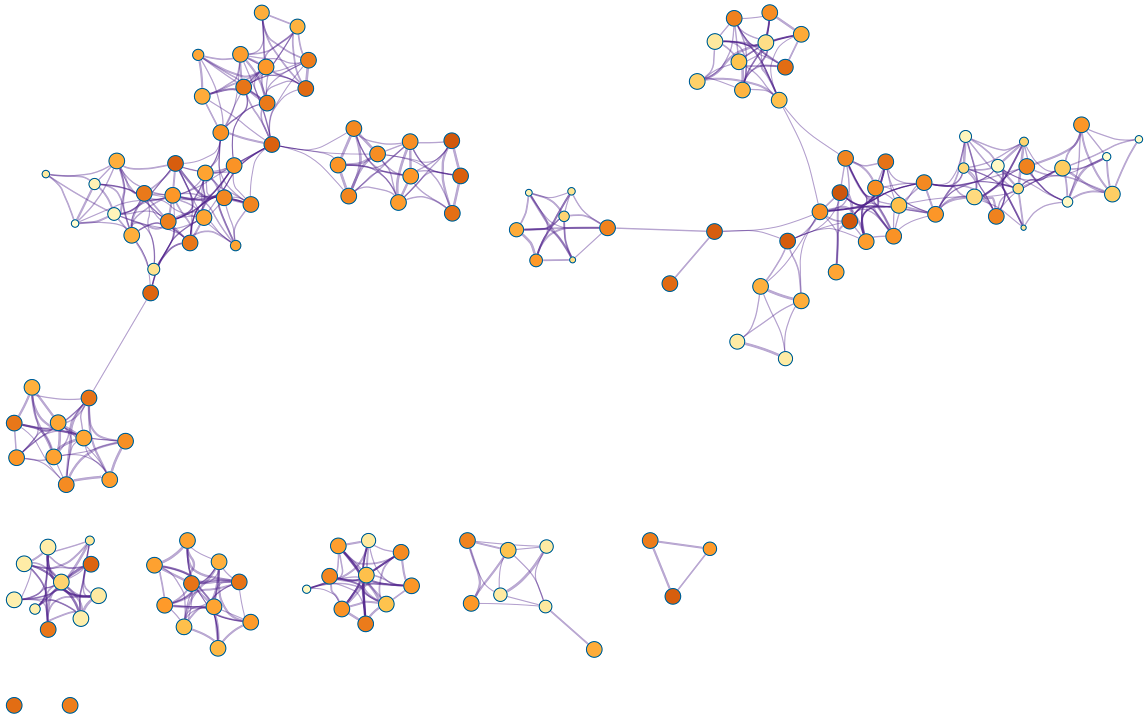

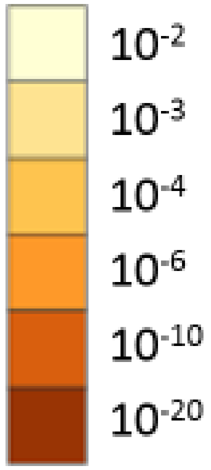


created by
http://metascape.org

### ColorByPValue.png

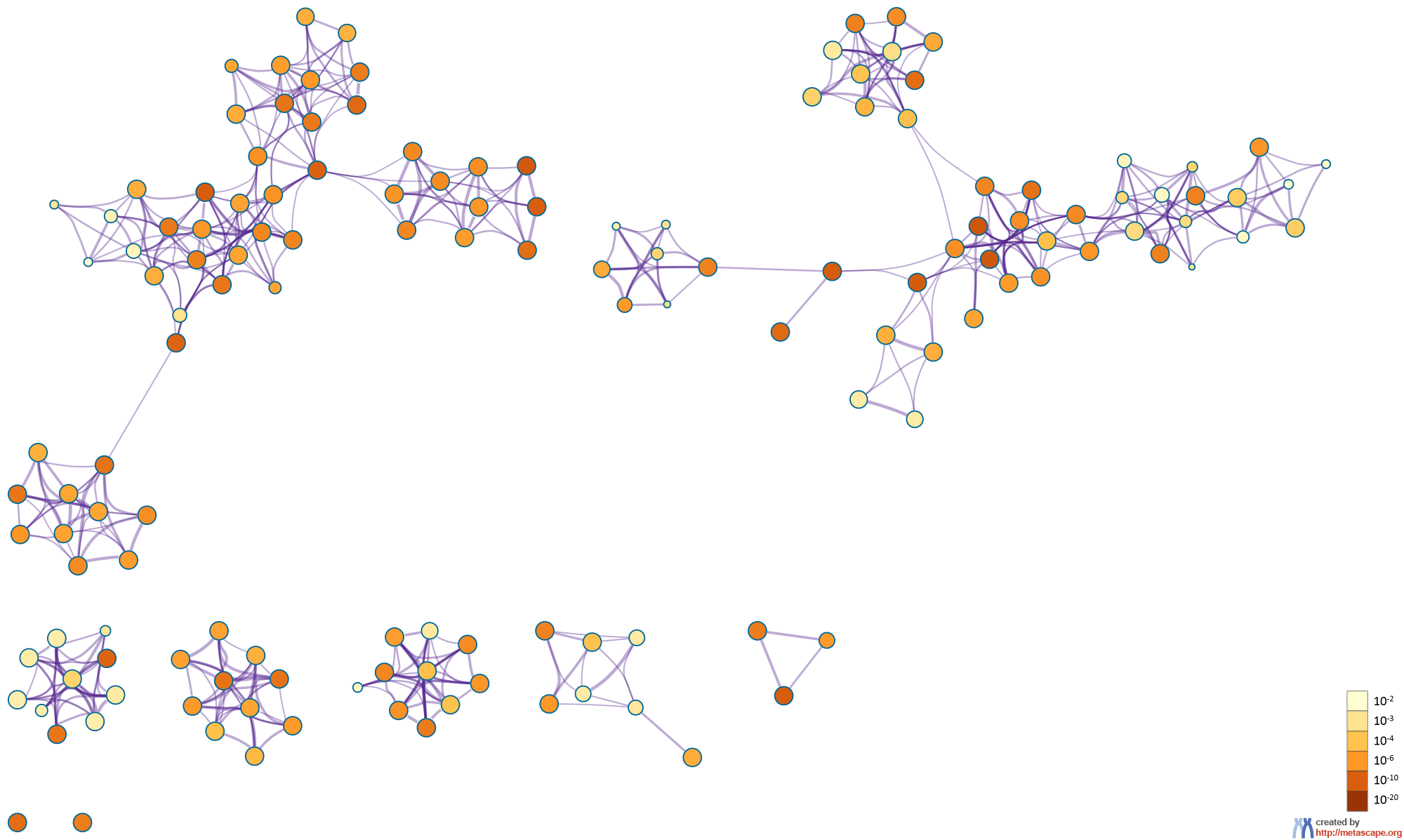

### CYS48.png

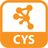

### HeatmapSelectedGO.pdf

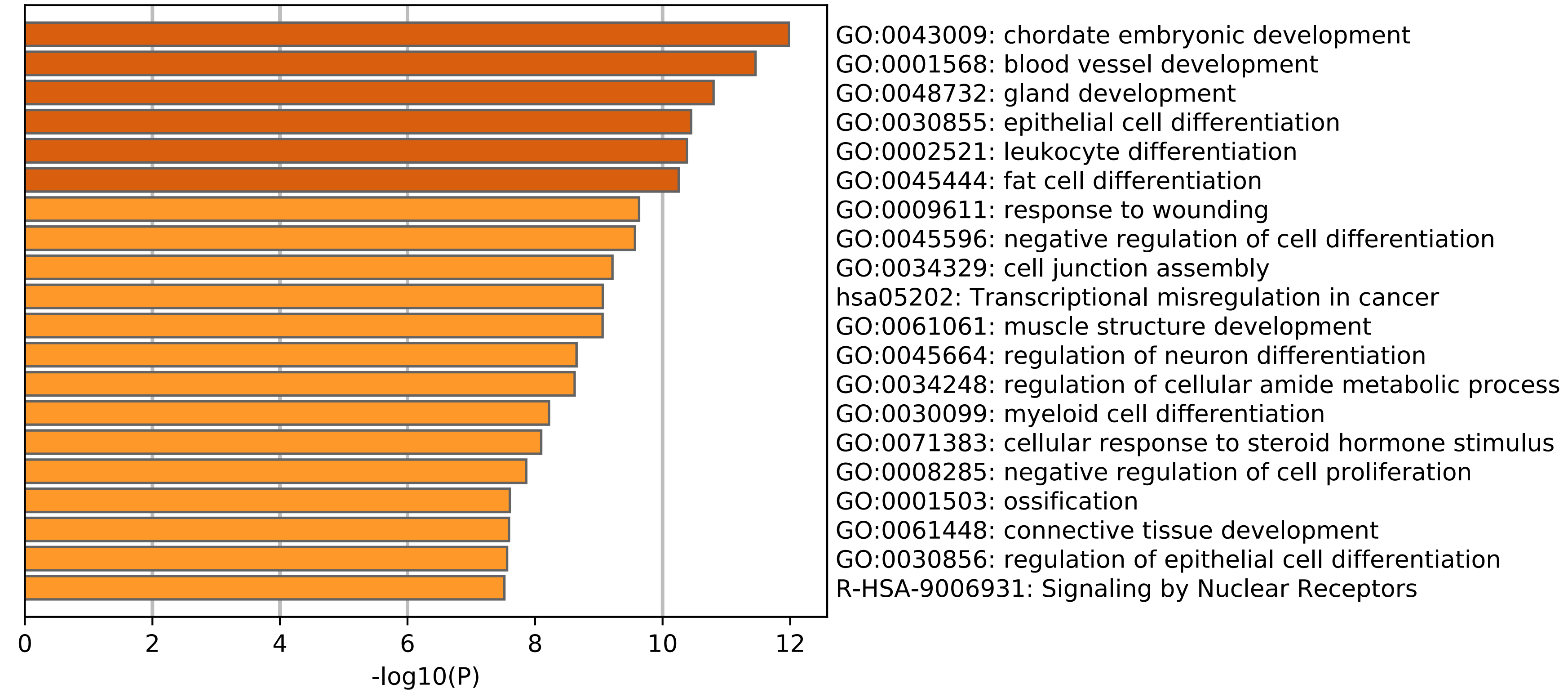

### HeatmapSelectedGO.png

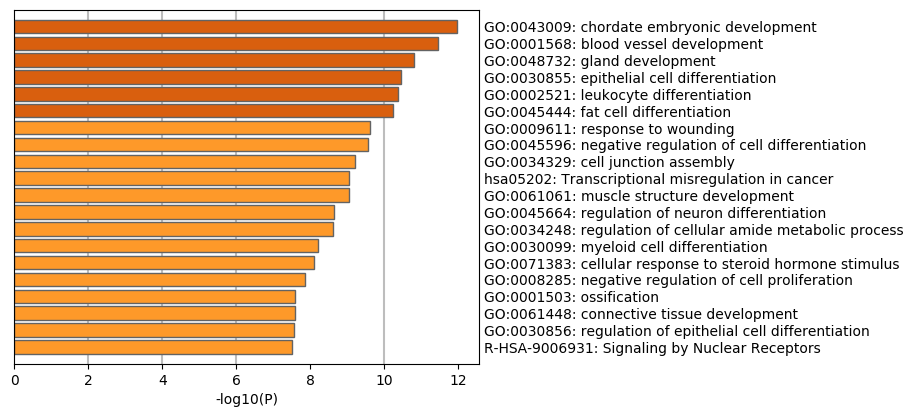

### HeatmapSelectedGO_DisGeNET.pdf

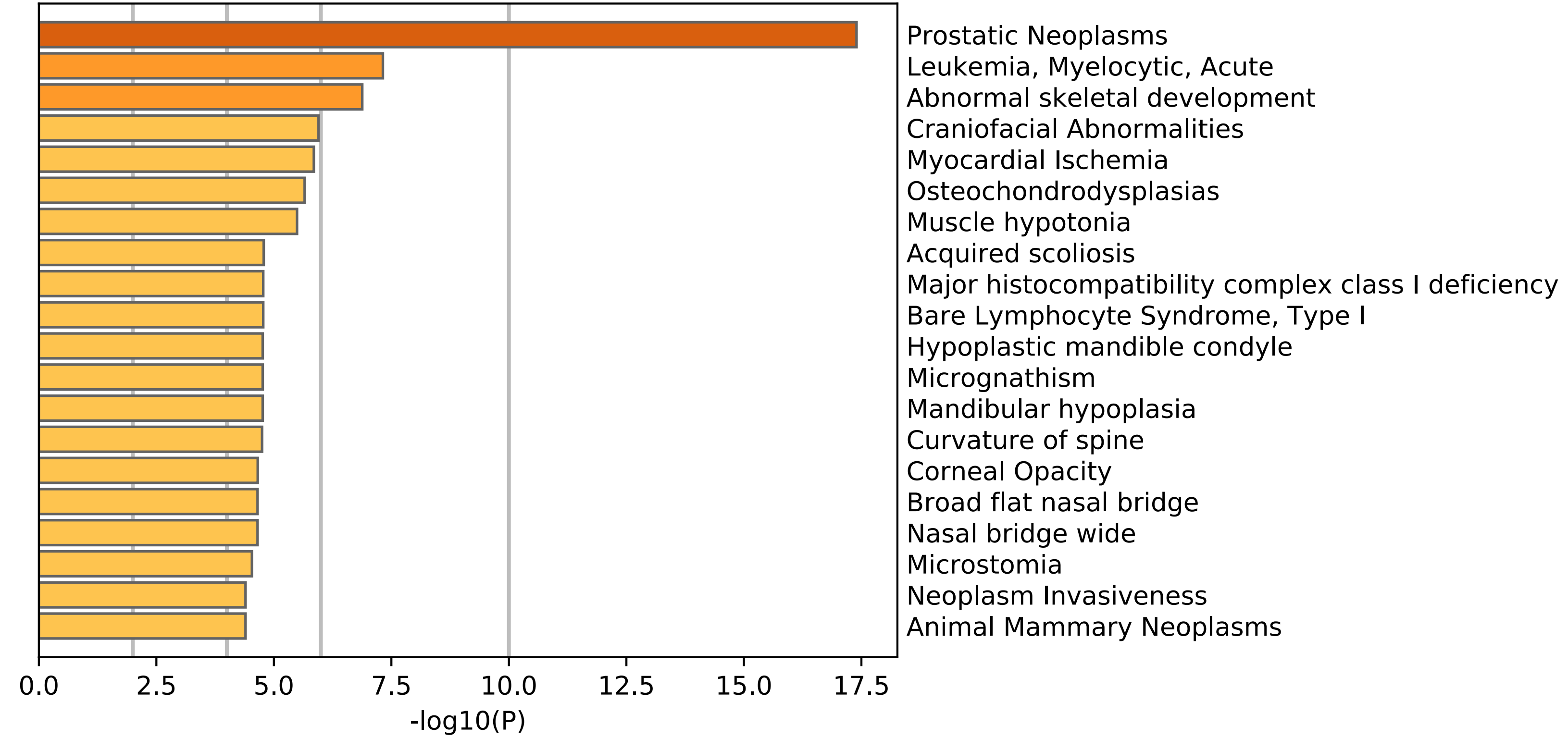

### HeatmapSelectedGO_DisGeNET.png

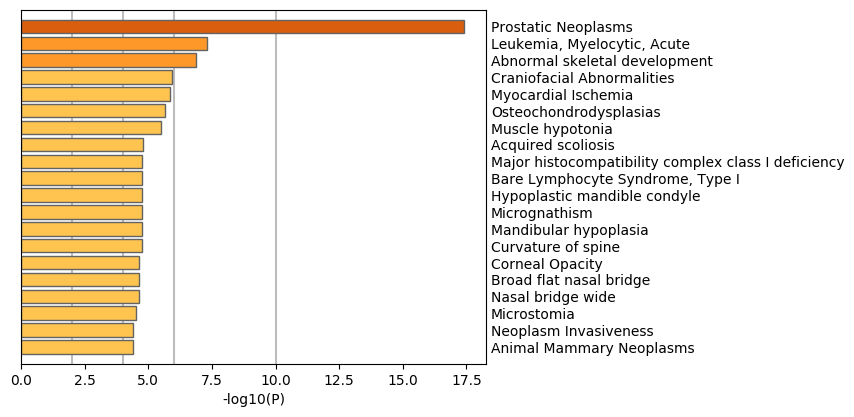

### HeatmapSelectedGO_PaGenBase.pdf

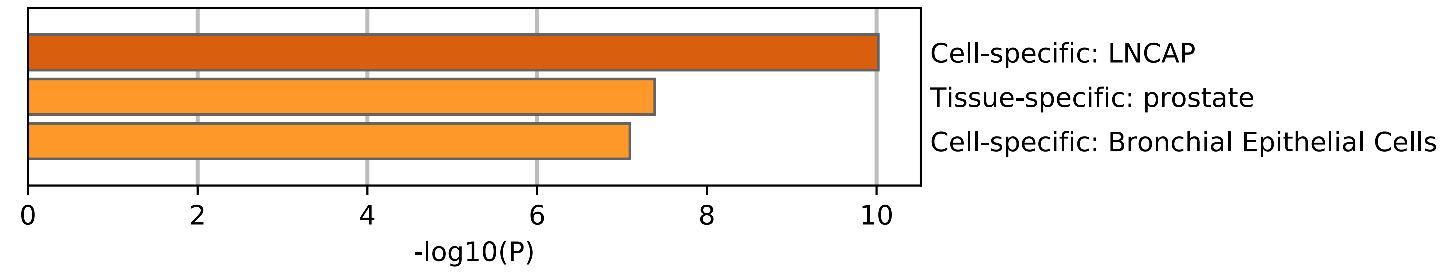

### HeatmapSelectedGO_PaGenBase.png

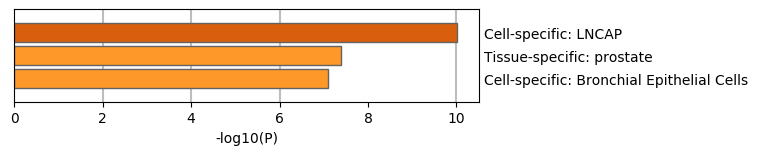

### HeatmapSelectedGO_TRRUST.pdf

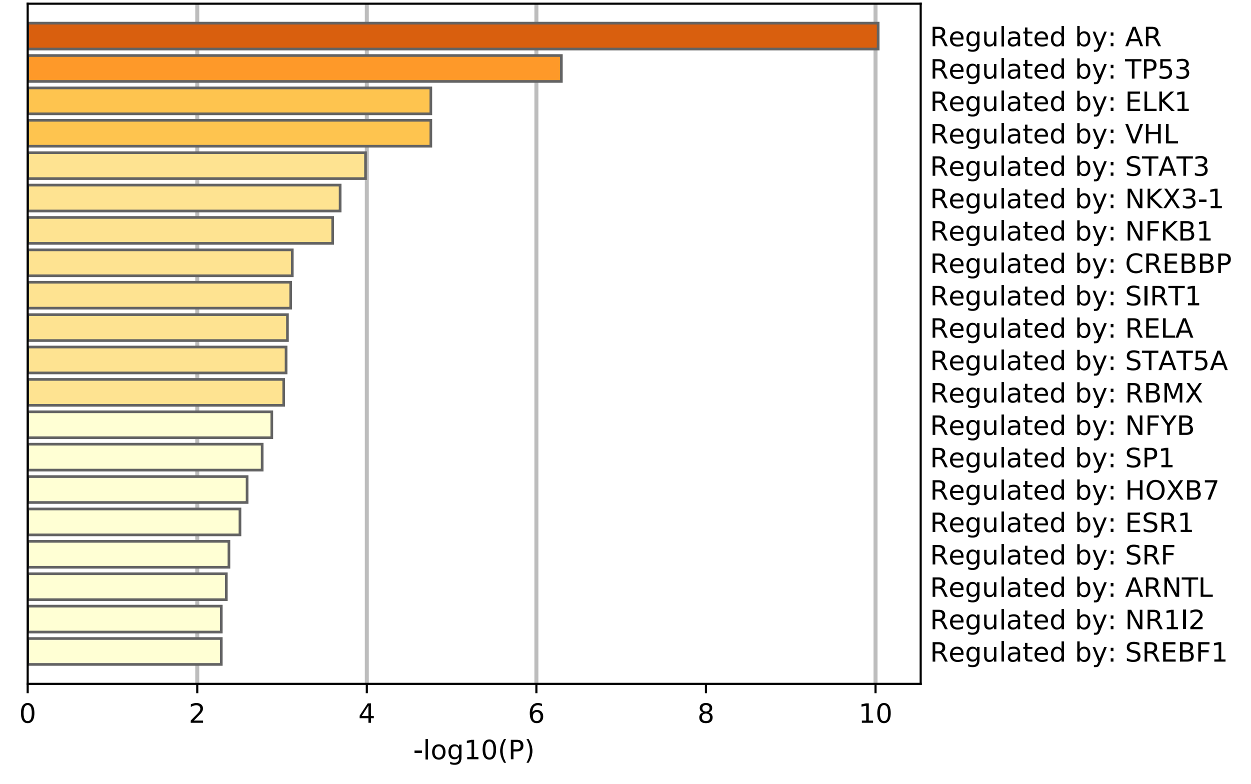

### HeatmapSelectedGO_TRRUST.png

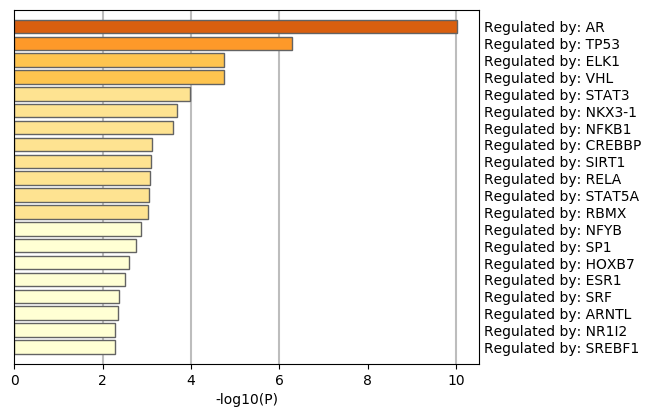

### HeatmapSelectedGOParent.pdf

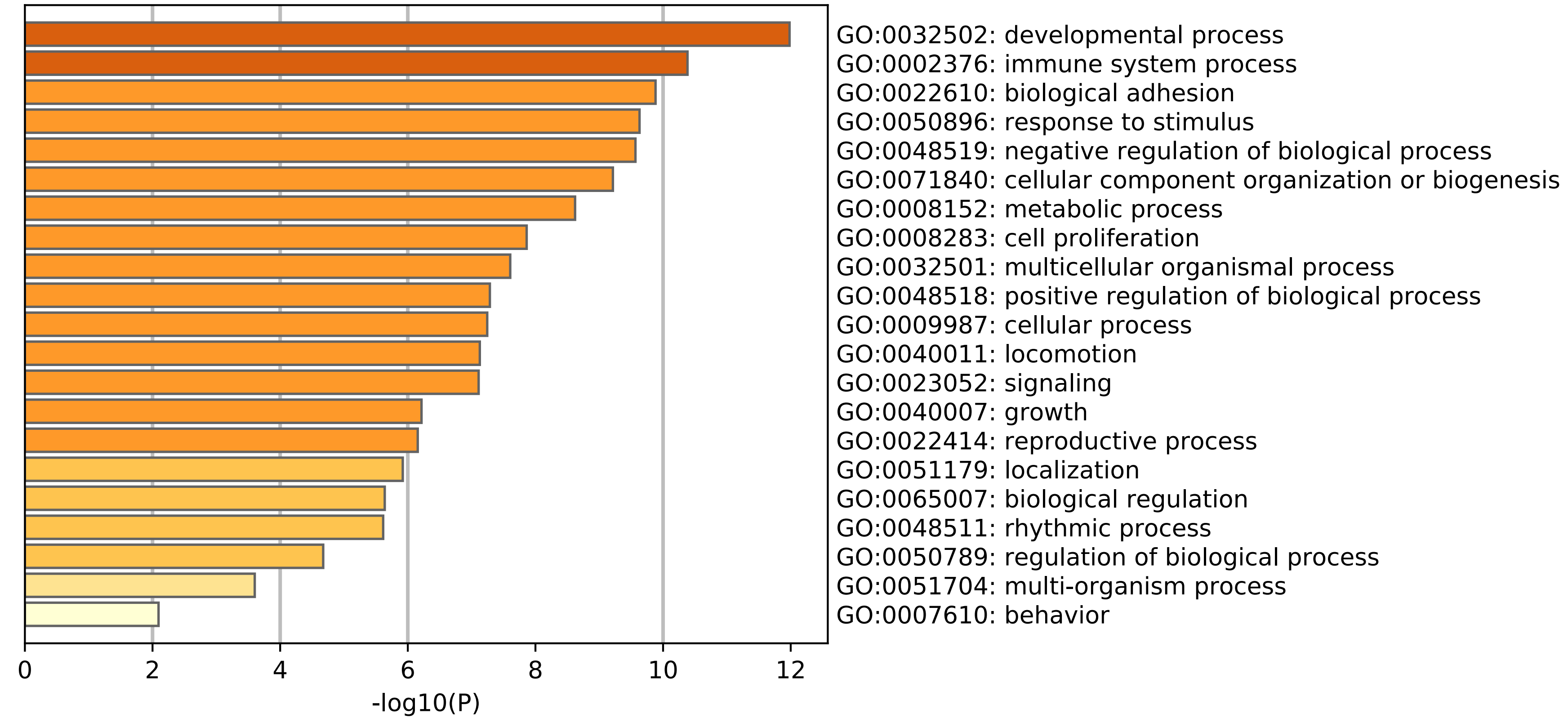

### HeatmapSelectedGOParent.png

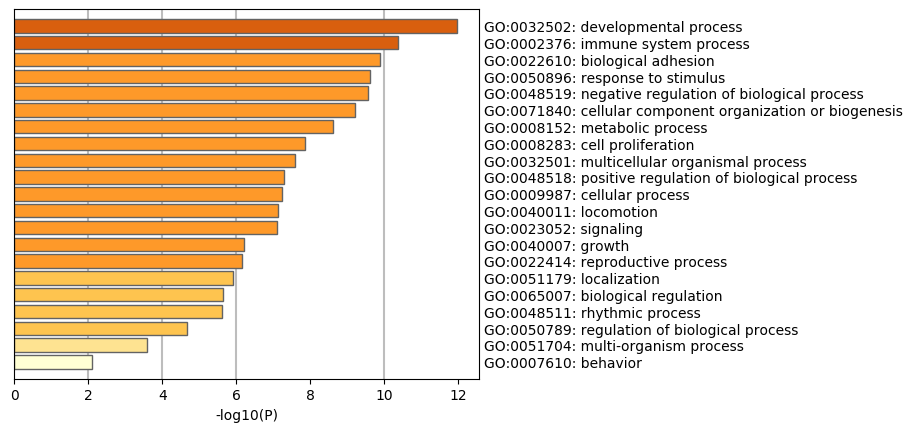

### HeatmapSelectedGOTop100.pdf

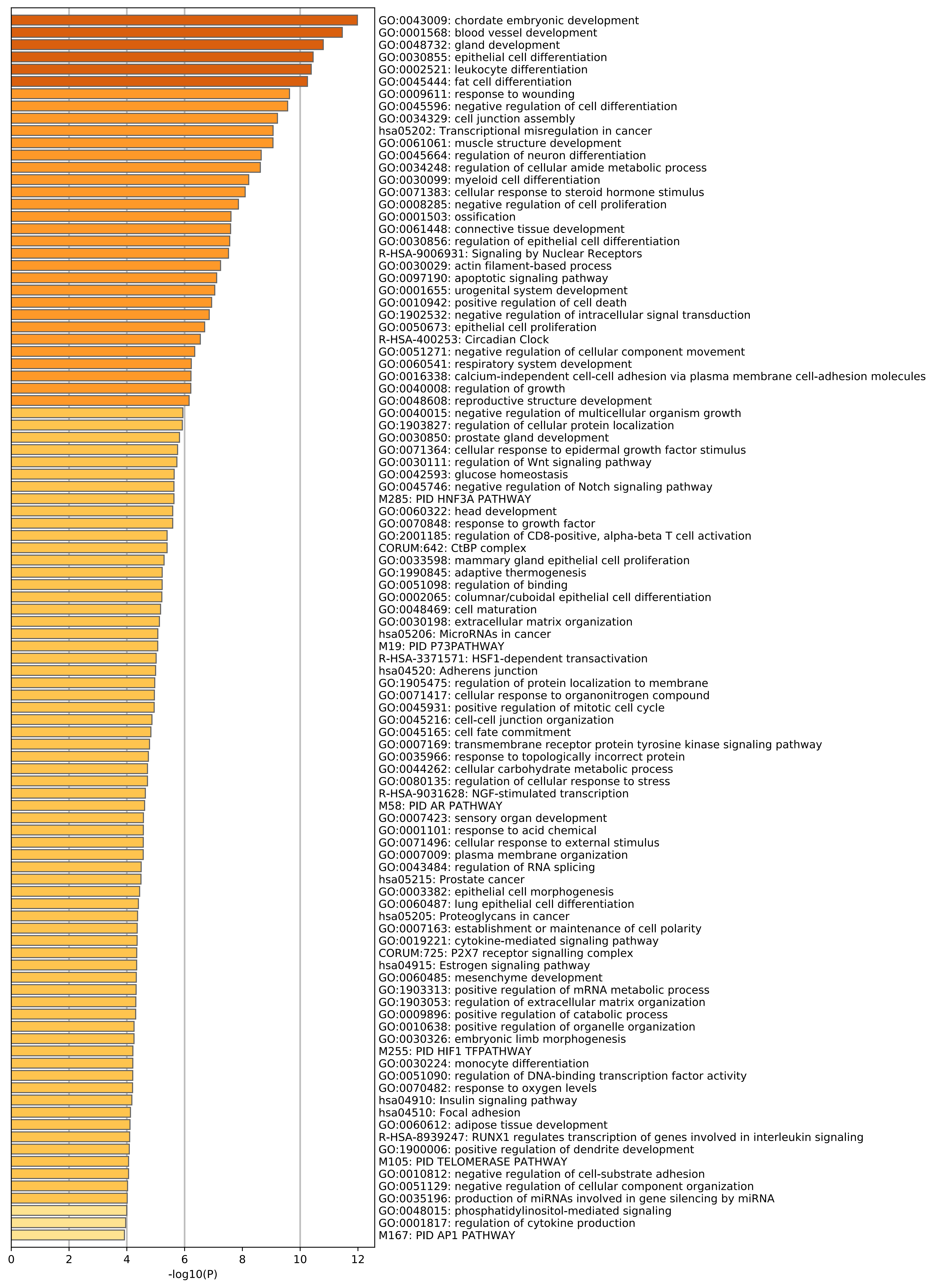

### HeatmapSelectedGOTop100.png

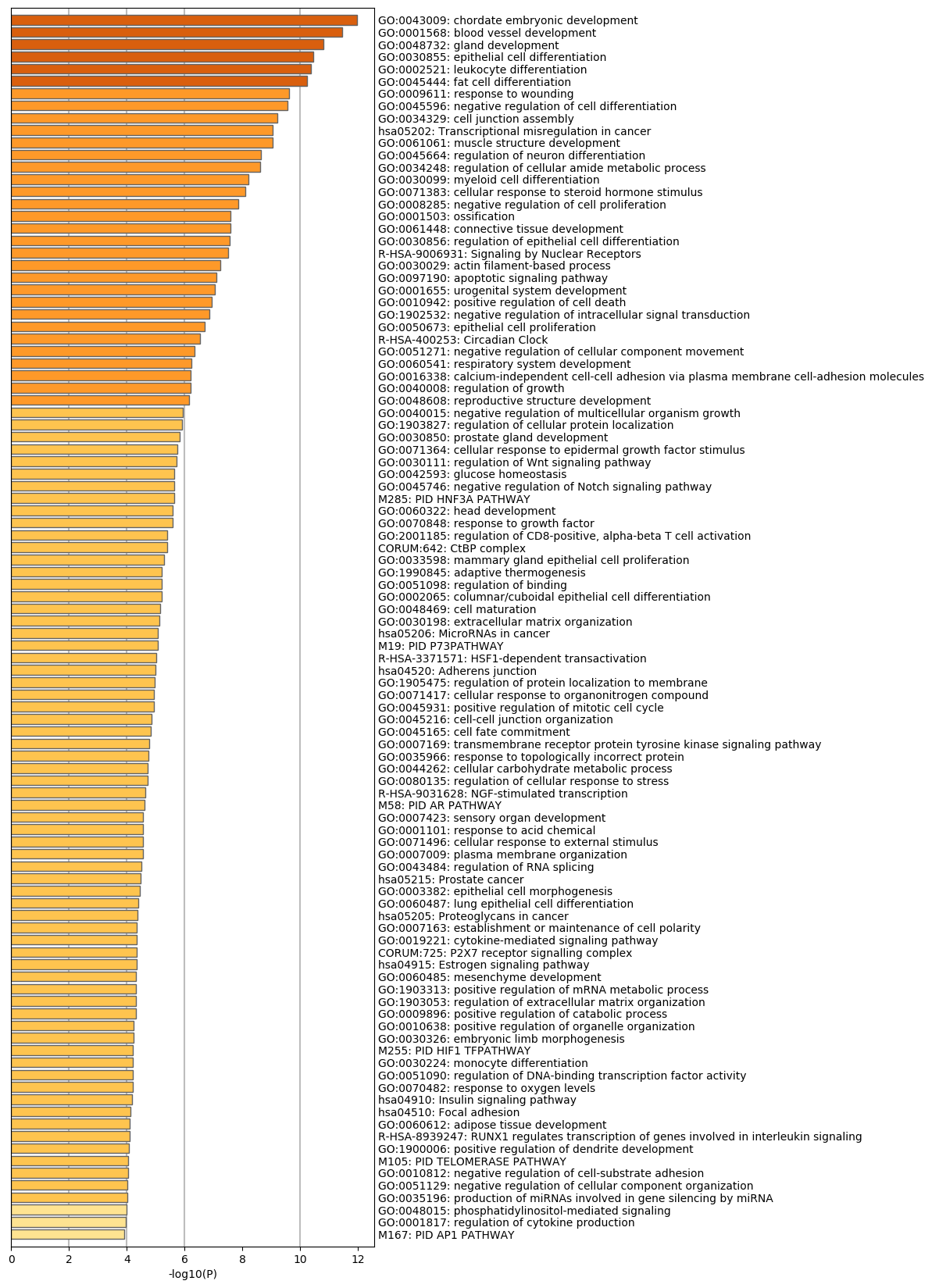

### PDF48.png

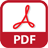

### SVG48.png

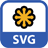

### WEB_CYS48.png

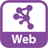
